# Interpreting the pervasive observation of U-shaped Site Frequency Spectra

**DOI:** 10.1101/2022.04.12.488084

**Authors:** Fabian Freund, Elise Kerdoncuff, Sebastian Matuszewski, Marguerite Lapierre, Marcel Hildebrandt, Jeffrey D. Jensen, Luca Ferretti, Amaury Lambert, Timothy B. Sackton, Guillaume Achaz

**Affiliations:** Institute of Plant Breeding, Seed Science and Population Genetics, University of Hohenheim, Germany; SMILE group, Center for Interdisciplinary Research in Biology (CIRB), Collège de France, CNRS UMR 7241, INSERM U 1050, Paris, France; Accenture, Vienna, Austria; Lycée Louis le Grand, Paris, France; Siemens AG, Munich, Germany; Center for Evolution & Medicine, School of Life Sciences, Arizona State University, USA; Big Data Institute, Li Ka Shing Centre for Health Information and Discovery, Nuffield Department of Medicine, University of Oxford, Oxford, UK; Institut de Biologie de l’ENS (IBENS), École Normale Supérieure, UMR CNRS 8197, INSERM U 1024, Paris, Franc; Informatics Group, Harvard University, Cambridge, MA, 02138, USA; Éco-anthropologie, Muséum National d’Histoire Naturelle, CNRS UMR 7206, Université Paris-Cité, Paris, France; Department of Genetics, University of California, Berkeley, CA 94620 USA

**Author notes:** these authors contributed equally to this work (Fabian Freund, Elise Kerdoncuff).

## Abstract

The standard neutral model of molecular evolution has traditionally been used as the null model for population genomics. We gathered a collection of 45 genome-wide site frequency spectra from a diverse set of species, most of which display an excess of low and high frequency variants compared to the expectation of the standard neutral model, resulting in U-shaped spectra. We show that multiple merger coalescent models often provide a better fit to these observations than the standard Kingman coalescent. Hence, in many circumstances these under-utilized models may serve as the more appropriate reference for genomic analyses. We further discuss the underlying evolutionary processes that may result in the widespread U-shape of frequency spectra.

## 1 Introduction

The Kingman coalescent, [Kin82], a stochastic process describing the distribution of random, bifurcating genealogical trees in a Wright-Fisher population, has been enormously impactful in the study of natural genetic variation in populations [Wak09]. Under the standard neutral theory [Kim68, Kim83], the coalescent can be used to derive expectations of neutral diversity by tracking mutations along the branches of random genealogies, and extensions can accommodate complex processes such as recombination [Hud83], population structure [WH98], and natural selection [KDH88]. The power of this approach relies on being able to compare deviations observed in real data from expectations under the coalescent model.

One common metric used to study the consistency between the assumptions of this model and the observed data is the *Site Frequency Spectrum* (SFS) - that is, the distribution of mutational frequencies, typically computed for a sample of *n* haploid genomes. Under the assumptions of the Standard Neutral Model (SNM) - including constant population size and panmixia - the expected SFS, averaged across the tree space, is given by *E*[*ξ_i_*] = *θ/i*, where *ξ_i_* is the number of sites that carry a derived variant of frequency *i/n* [Fu95]. The *θ* parameter of the SNM is defined as *θ* = 2*pNμ*, where *p* is the ploidy (typically 1 or 2), *N* the population size, and *μ* the mutation rate.

Observed SFS in natural populations are often poorly fit by this expectation, owing to violations of one or more of the underlying assumptions of the SNM, including varying population sizes, population structure, direct selection, and linkage with selected sites [JPS^+^19]. A standard procedure in population genetics is thus to first statistically test for the SNM (treated as *H*_0_, a null statistical model) and then, when rejected, fit a variety of alternative demographic and/or selection models.

In this article, we show that among a collection of genome-wide SFS from a diverse set of species, many show an unexpected excess of low and high frequency variants, resulting in a U-shaped SFS. Many possible factors may result in such a pattern of variation. These include recent migration from non-sampled populations [ME20], population structure [LBL^+^16], misorientation of ancestral and derived alleles [BD03], biased gene conversion [PATE18], recent positive selection at many targets across the genome [BWSH01], background selection [CGD18], [JCJ20], temporally-fluctuating selection [HSDB08], and various reproductive strategies [TL14].

A number of these scenarios result in an important general violation of Kingman assumptions: the presence of multiple mergers in genealogies (*i.e*., a node with more than two descendants). Under such scenarios, these distributions are better described by a more general class of models known as the Multiple Merger Coalescent (MMC) [Sag99, Pit99, DK99, MS01, Sch00]. Briefly, MMCs may arise when the number of offspring per individual has very high variance across the population. Such effects of concentrations of ancestrality (resulting in polytomies in the trees) have been reported in various species across all kingdoms of life [Mon16], and MMC-like genealogies have been observed for species ranging from bacteria (e.g. for *Mycobacterium tuberculosis* [MASSJ20, MGF20]) to viruses (e.g. for influenza [SHJ19]) to animals (e.g. for the nematode *Pristionchus pacificus* [RNW^+^14], multiple fish species, e.g. [ÁH14, NNY16, VPC^+^21]) and even to cancer cells [KVS^+^17].

Multiple neutral and selective processes can produce MMC genealogies in natural populations. Generally, the term sweepstake reproduction has been proposed for species that have rare individuals with a high reproduction rate coupled with high early-life mortality. In these species, a single or few individuals can become ancestors of a macroscopic fraction of the population by chance, thus resulting in MMC genealogies (for a review, see [Eld20]). Multiple models featuring the recurrent and rapid emergence of genotypes with high fitness also result in MMC genealogies, often modeled by the Bolthausen-Sznitman coalescent or related models, e.g. [BD13, NH13, DWF13, BBS13, Sch17]. Importantly, other biological factors can also lead to MMC-like genealogies, including large rapid demographic deviations [BBM^+^09], seed banks [CCSWB22], extinction-recolonisation in metapopulations [TV09] and range expansions [BHK21]. Yet, the frequency of MMC genealogies in nature, and more generally whether MMC models ought to be employed as a more appropriate null for certain species, remains an open question.

In this study, we collected 45 species (Table 2) from across the tree of life (bacteria, plants, invertebrates and vertebrates), for which genome-wide polymorphism data (with sample sizes of *n* > 10) were available together with an outgroup to assign ancestral and derived states. We show that MMC genealogies provide a better fit than the Kingman coalescent in many cases, even when both are combined with non-constant demography and misorientation of ancestral and derived alleles. For several species, the fit is excellent. For each species, we tested two simple MMC models: Beta-MMC [Sch03] and Psi-MMC [EW06], both tuned by a single parameter that interpolates between pure radiation to a Kingman-like tree. Demography is here tuned by a single parameter (a simple exponential growth), as is the frequency of misorientation errors. Using composite-likelihood maximization [Nie00] on genome-wide data, we explore statistical power to distinguish between these contributing factors. Finally, we discuss how MMCs may be better utilized in future population genetic analysis, and what evolutionary forces may contribute to the pervasive observation of U-shaped SFS.

**Table 1:** Model selection via two-step Bayes factor criterion. Based on 2,000 simulations for each true model assuming n = 25 individuals with 100 loci with 50 mutations each on average. For each simulation, the coalescent parameter is fixed and the growth parameter g and the allele misorientation rate e are randomly chosen (*g* ∈ [0, 11.25], *e* ∈ [0, 0.1]). The second column hows whether the parameters used for simulation were included in the inference grid. Fractions are rounded to two digits. MMC refers to cases in which neither the Psi- nor Beta-coalescent is preferred. An expanded version with enhanced sample size is provided in Table A.2. For details on simulations and inference parameters see Appendix A.5.

| True model | Within<br>the grid? | Fraction model inferred as |  |  |  |
| --- | --- | --- | --- | --- | --- |
|  |  | Kingman | Beta | Psi | MMC |
| $\alpha = 2$ | yes | 0.79 | 0.21 | | |
| $\alpha = 1.9$ | yes | 0.34 | 0.66 | | |
| $\alpha = 1.8$ | yes | 0.02 | 0.91 | 0.04 | 0.03 |
| $\alpha = 1.625$ | no | | 0.9 | 0.06 | 0.05 |
| $\alpha = 1.025$ | no | | 1 | | |
| $\Psi = 0.005$ | no | 0.55 | 0.45 | | |
| $\Psi = 0.025$ | no | 0.05 | 0.72 | 0.14 | 0.09 |
| $\Psi = 0.05$ | yes | | 0.12 | 0.82 | 0.06 |
| $\Psi = 0.075$ | no | | 0.06 | 0.91 | 0.03 |
| $\Psi = 0.1$ | yes | | 0.02 | 0.98 | |

**Table 2:** Data sets description: Taxa, Species, number of individuals (n) and number of polymorphic sites (#SNP). Best fitting model (Kingman (KM), Beta, Psi-coalescent or no preference between Multiple Merger Coalescents (MMC)), its parameters (parameters describing coalescence (Coal), growth rate (g) and misorientation (e)) and goodness-of-fit grade from Cramer’s *V* values.

| Order | Species | n | #SNP | Model | Coal | $g_{\text{Model}}$ | $e_{\text{Model}}$ | Grade |
| --- | --- | --- | --- | --- | --- | --- | --- | --- |
| Vertebrates | *Aptenodytes patagonicus* | 20 | 1,278 | Beta | 1.25 | 1.5 | 0 | B |
|  | *Athene cunicularia* | 40 | 11,268,203 | Beta | 1.8 | 1 | 0.03 | B |
|  | *Corvus cornix* | 38 | 7,167,395 | Beta | 1.95 | 1 | 0 | A |
|  | *Coturnix japonica* | 20 | 5,061,864 | Beta | 1.45 | 0.5 | 0.01 | A |
|  | *Egretta garzetta* | 10 | 9,318,499 | Beta | 1.75 | 0 | 0.02 | B |
| | *Emys orbicularis* | 20 | 515 | KM | ∅ | 0.5 | 0 | C |
| | *Ficedula albicollis* | 24 | 14,697,230 | $\Psi$ | 0.01 | 0.5 | 0.01 | A |
|  | *Gorilla gorilla gorilla* | 54 | 9,878,547 | Beta | 1.9 | 0 | 0 | B |
|  | *Homo sapiens* | 216 | 19,441,528 | Beta | 1.85 | 0 | 0 | A |
| | *Lepus granatensis* | 20 | 769 | MMC/$\Psi$ | 0.12 | 0 | 0.03 | C |
| | *Nipponia nippon* | 16 | 1,140,694 | KM | ∅ | 0 | 0.03 | D |
|  | *Pan paniscus* | 26 | 6,293,657 | Beta | 1.85 | 1 | 0 | B |
|  | *Pan troglodytes ellioti* | 20 | 10,009,190 | Beta | 1.7 | 0 | 0 | A |
|  | *Parus major* | 54 | 14,174,305 | Beta | 1.75 | 0 | 0.01 | A |
| | *Parus caeruleus* | 20 | 866 | MMC/$\Psi$ | 0.04 | 0 | 0.02 | B |
| | *Passer domesticus* | 16 | 18,501,992 | KM | ∅ | 0 | 0 | A |
| | *Phylloscopus trochilus* | 24 | 33,401,127 | KM | ∅ | 12.5 | 0 | A |
|  | *Taeniopygia guttata* | 38 | 53,263,038 | Beta | 1.75 | 4 | 0 | A |
| Invertebrates | *Armadillidium vulgare* | 20 | 23,323 | Beta | 1.7 | 0 | 0.03 | C |
|  | *Artemia franciscana* | 20 | 5,548 | Beta | 1.65 | 0 | 0.03 | B |
|  | *Caenorhabditis brenneri* | 20 | 1,339 | Beta | 1.5 | 0 | 0.06 | C |
| | *Caenorhabditis elegans* | 574 | 165 | KM | ∅ | 0 | 0.06 | D |
| | *Ciona intestinalis A* | 20 | 480 | KM | ∅ | 0 | 0.11 | B |
|  | *Ciona intestinalis B* | 20 | 1,883 | Beta | 1.65 | 0 | 0.02 | B |
|  | *Culex pipiens* | 20 | 5,442 | Beta | 1.55 | 0.5 | 0.01 | B |
|  | *Drosophila melanogaster* | 196 | 4,662,706 | Beta | 1.65 | 0.5 | 0.02 | A |
| | *Halictus scabiosae* | 22 | 712 | MMC/$\Psi$ | 0.04 | 0 | 0.01 | B |
|  | *Melitaea cinxia* | 18 | 1,695 | Beta | 1.7 | 0.5 | 0.03 | B |
| | *Messor barbarus* | 20 | 9,651 | KM | ∅ | 0.5 | 0 | C |
| | *Ostrea edulis* | 20 | 939 | MMC/$\Psi$ | 0.04 | 0 | 0.02 | B |
|  | *Physa acuta* | 18 | 4,286 | Beta | 1.5 | 0 | 0.02 | B |
| | *Sepia officinalis* | 18 | 1,740 | KM | ∅ | 0 | 0.02 | C |
| Plants | *Arabidopsis thaliana* | 345 | 10,322,757 | Beta | 1.6 | 0 | 0.07 | A |
| | *Zea mays* | 66 | 520,310 | $\Psi$ | 0.01 | 0 | 0 | A |
| Bacteria | *Acinetobacter baumannii* | 79 | 78,175 | Beta | 1.8 | 0 | 0.1 | B |
| | *Bacillus subtilis* | 38 | 105,523 | $\Psi$ | 0.14 | 0 | 0.2 | B |
| | *Chlamydia trachomatis* | 59 | 9,924 | KM | ∅ | 0 | 0.11 | D |
| | *Clostridium difficile* | 11 | 192 | KM | ∅ | 15 | 0.15 | D |
| | *Escherichia coli* | 62 | 84,222 | KM | ∅ | 0 | 0.06 | B |
| | *Helicobacter pylori* | 70 | 27,498 | $\Psi$ | 0.01 | 1 | 0.2 | B |
| | *Klebsiella pneumoniae* | 156 | 203,601 | KM | ∅ | 18.5 | 0.15 | D |
|  | *Mycobacterium tuberculosis* | 33 | 7,142 | Beta | 1.05 | 2.5 | 0 | C |
| | *Pseudomonas aeruginosa* | 86 | 90,258 | $\Psi$ | 0.06 | 3 | 0.2 | B |
| | *Staphylococcus aureus* | 152 | 30,052 | $\Psi$ | 0.01 | 1 | 0.2 | B |
|  | *Streptococcus pneumoniae* | 32 | 49,917 | Beta | 1.5 | 0 | 0.08 | C |

## 2 Materials and Methods

### 2.1 Coalescent and allele misorientation models

We compared the empirically observed SFS to the theoretical SFS expected under a variety of models. The genealogical models emerge from a discrete generation reproduction model. Each is a (random) tree with *n* leaves which approximates the genealogy for a sample of size *n* in a reproduction model in which the population size *N* is very large (*N* → ∞). One unit of time in the coalescent tree corresponds to many generations in the underlying reproduction model: for Kingman’s coalescent one time unit corresponds to *N* generations of a haploid Wright-Fisher model, or order of *N*^2^ time steps of an haploid Moran model. This correspondence affects how population size changes are reflected in the coalescent approximation (see definition below, for mathematical justification and details see [GT94, MHAJ18, Fre20]). On the genealogical tree, mutations are placed randomly via a Poisson process with rate *θ*/2.

We compared three coalescent models: Kingman’s *n*-coalescent, Psi-*n*-coalescent (also called Dirac-*n*-coalescent) with parameter Ψ ∈ [0,1] and Beta(2 – *α,α*)-*n*-coalescent with *α* ∈ [1, 2]. The parameters *α* or Ψ regulate the strength and frequency of multiple mergers: the smaller *α* or the larger the Ψ, the more frequently coalescence events are multiple mergers of increasing size. Both MMCs incorporate Kingman’s *n*-coalescent as a special case (*α* = 2 or Ψ = 0).

Both MMC coalescent models can be defined for demographic variation that stays of the same order, *i.e*. where the populations size ratio *ν_t_* = *N_t_*/*N*_0_ of the population size at time *t* in the past (in coalescent time units) is positive and finite (for large population sizes *N*). The coalescent merges any *k* of *b* (ancestral) lineages present at a time *t* with rate

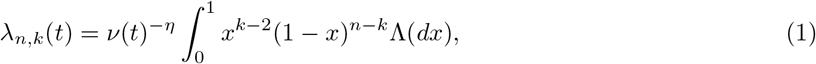

where

- Λ could be any probability distribution on [0,1] but is here either the Dirac distribution (point mass) in Ψ (Psi-coalescent) or the Beta(2 – *α, α*) distribution (Beta coalescent).
- *η* is a scaling factor reflecting how many time steps from the discrete reproduction model form one unit of coalescent time. More precisely, it is the power of *N* of the scaling factor: e.g. *η* = 2 for a Moran model and *η* = 1 for a Wright-Fisher model.

A common way of constructing the Λ-coalescent, which provides a nice interpretation of Eq (1), is the paintbox process [Pit99]: at rate *x*^-2^Λ(*dx*) per time unit, paint each lineage independently with probability *x* and merge all painted lineages simultaneously. Note that when Λ is the Dirac mass at 0, *λ_n,k_*(*t*) is nonzero only when *k* = 2, recovering Kingman’s coalescent.

We focused on exponentially growing populations, *i.e*. a population size ratio *ν*(*t*) = exp(−*gt*) for growth rate *g* ≥ 0 (see Appendix A.1 for interpretation of *g* in the initial reproduction model). As underlying reproduction models, we use modified Moran models [HM13, EW06, MHAJ18]. At each time step, in a population of size *N*, a single random individual has *U* + *G* offspring while *N* – *U* random individuals have 1 offspring (leaving *U* – 1 individuals devoid of offspring). As a consequence, the population grows from *N* to *N* + *G* individuals and *G* is chosen to fit the desired growth rate.

In a standard Moran model, *U* = 2 and *G* = 0 leaving the population size constant. However, for both MMCs, *U* is set to different values. In both cases, the mean of *U* does not grow indefinitely with *N* (for all parameters *α* and Ψ), but the resulting variance does (for *α* = 2 and Ψ = 0).

- In the Psi-*n*-coalescent (essentially [EW06, MHAJ18]), we have *U* = 2, except when a sweepstake event occurs with a small probability of order *N^γ^* (1 < *γ* ≤ 2); in this case, *U* = ⌊*N*Ψ⌋. In the coalescent time scale, one unit of time corresponds to an order of *N^γ^* time steps; this is the expected time to a sweepstake event so that *η* must equal *γ*. We chose *γ* = η = 1.5 for Ψ > 0, and *γ* = *η* = 2 for Ψ = 0 (standard Moran model) with *U* = 2 in every time step.
- In the Beta-*n*-coalescent [HM13, Fre20], *U* has distribution 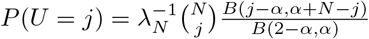, where *B* is the Beta function and *λ_N_* is the normalizing constant. Consequently, although the random variable *U* has a finite mean of at most 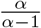, it can take large values with high probability when *α* < 2. See Appendix A.1 for more details. On the coalescent time scale, one unit of time corresponds to an order of *N^α^* time steps, so *η* = *α*. Note that *α* = 2 is the classical Moran model and thus leads to Kingman’s coalescent.

For statistical inference, we treat the observed SFS of s mutations as s independent multinomial draws from the expected SFS (see [Nie00] and [EBBF15, Eq. 11] [MHAJ18, Eq. 14]). This computes an approximate composite likelihood function of the data for any combination of growth rate (*g*) and coalescent parameter (*α* or *ψ*). However, to include the effect of misorienting the ancestral allele with the derived allele, we introduced another parameter *e*. On average, a misorientation probability of *e* lets a fraction e of the derived allele carried by *i* sequences to be falsely seen as appearing in *n* – *i* sequences. Additionally, as described in [Lap17, Section 4.2] or [BD03, p. 1620], as misorientation stems from double-mutated sites, *e* also relates to the number of sites that cannot be oriented when compared with the outgroup owing to the presence of a third allele (see Appendix A.3). We account for these two effects of *e* by swapping a fraction e of the variants at frequency *i/n* to 1 – *i/n* and we assume a Jukes-Cantor substitution model [JC^+^69] to predict for the number *s*_≠_ of non-polarizable tri-allelic variants. This leads to a slight variant of [MHAJ18, Eq. 14]. For any coalescent model with a specific set of coalescent, exponential growth and misorientation parameters, the pseudolikelihood is:

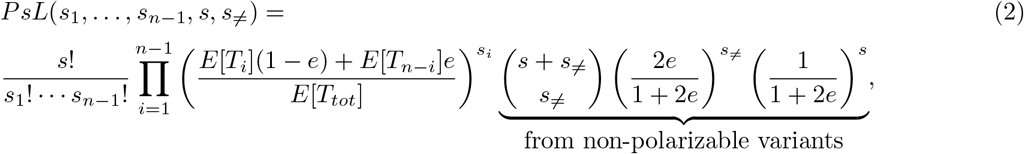

where *s*_1_,…, *s*_*n*−1_ is the observed SFS (so we observe *s_i_* sites with derived allele frequency *i/n*), 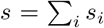 is the total number of polarizable polymorphic sites and *s*_≠_ is the number of non-polarizable sites. *E*[*T_i_*] is the expected sum of branch lengths that support *i* leaves in the genealogy and *E*[*T_tot_*] is the sum of all branch lengths. For *e* = 0, we set the term estimated from non-polarizable variants to 1. See Appendix A.3 for details on the derivation.

### 2.2 Statistical inference

To find the best-fitting parameters, we conduct a grid-search for the highest pseudolikelihood. The expected branch lengths *E*[*T_i_*] in Eq. (2) are computed as in [MHAJ18], using the approach from [SKS16]. We use the following grids with equidistant steps

**Beta:** *α* ∈ [1, 2] in steps of 0.05, *g* ∈ [0, 25] in steps of 0.05, *e* ∈ [0,0.15] in steps of 0.01.
**Psi:** Ψ ∈ [0,1] in steps of 0.05, *g, e* as for Beta above, complemented with Ψ ∈ [0,0.2] in steps of 0.01 (further expanding *g* ∈ [0, 30] by steps of 0.05 and *e* ∈ [0,0.2] by steps of 0.01) when Ψ was estimated to be close to 0.

To perform model selection between the three coalescent models, we computed the two following log Bayes factors:

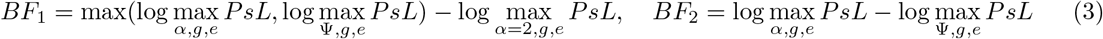

from the maximum pseudolikelihoods computed for the three models. We inferred a MMC genealogy when *BF*_1_ > log(10) and further chose a Beta coalescent or a Psi-coalescent when (additionally) *BF*_2_ > log(10) or *BF*_2_ < – log(10) respectively.

For the best fitting parameter combinations either over the full parameter space or restricted to the Kingman coalescent with growth and allele misorientation (*i.e*., fixing *α* = 2 or Ψ = 0), we assessed the goodness-of-fit of the observed data. First, we graphically compare the observed SFS with the expected SFS, approximated as 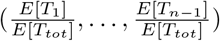. Second, we quantified the (lack of) fit of the data by Cramér’s *V*, a goodness-of-fit measure which accounts for different sample sizes and different numbers of polymorphic sites. See Appendix A.4 for details.

### 2.3 Data

We collected 45 genome-wide SFS that are described in Tables 2 and A.5. The collected SFS come from public data sets or private communications. For 20 data sets, SFS were extracted from whole genome SNP data, including both coding and non-coding regions. For 16 data sets, they were extracted only from transcriptomes (equivalent to coding regions). For 9 bacterial data sets, the SFS were extracted from the core genome. Supplementary files 1 and 2 provide the shapes of the empirically-observed SFS.

## 3 Results

We have first demonstrated the power of the methodology using extensive simulations, and then applied it to 45 real SFS computed from a very large variety of taxa.

### 3.1 Statistical performance

Using simulations, we first assess the power of the method to retrieve the correct model and then its power to estimate the parameters. Briefly, for each simulation, we simulated 100 independent loci for each parameter combination, sampling over the coalescent parameter (*α* or Ψ), the growth rate of the demographic model (*g*), and the misorientation probability (*e*). For each locus, we then simulate SNPs under an infinite sites model, with a mutation rate such that on average 50 sites are segregating for each locus. This simulation setup is described in further detail in Appendix A.5.

Applied to the simulated data, our method performs well. Even for small datasets (*n* = 25), the model selection approach based on Bayes factors computed from Eq. (2) identifies the correct multiple merger model in most cases (Table 1), as long as multiple mergers occur with reasonable frequency. As the rate of multiple mergers becomes very low (*α* ≈ 2 or Ψ ≈ 0), mis-identifications are more common (Table 1). However, even when our model prefers the beta-coalescent for data simulated with *α* = 2, in 96% of such cases (with *n* = 100; 71% with *n* = 20), we estimate *α* ≥ 1.9, suggesting that even when model mis-idenfication occurs, parameter estimation remains reliable (Table A.3). Over the range of parameter combinations, larger sample sizes lead to smaller errors, as expected. This selection approach is conservative with respect to departures from the standard Kingman coalescent, as we choose a Kingman genealogy model if the Bayes factor does not distinctively point towards an MMC model.

Parameter estimation within both the Beta- and Psi-coalescent models works well for multi-locus data for large enough samples, especially for the allele misorientation rate *e* and for the coalescent parameter *α* or *ψ* (Figure 1, Figures A.2–A.4). The growth rate, in contrast, is only estimated well for situations where the simulated growth rate was low (Figures A.9, A.12, A.15, A.18).

**Figure 1:**
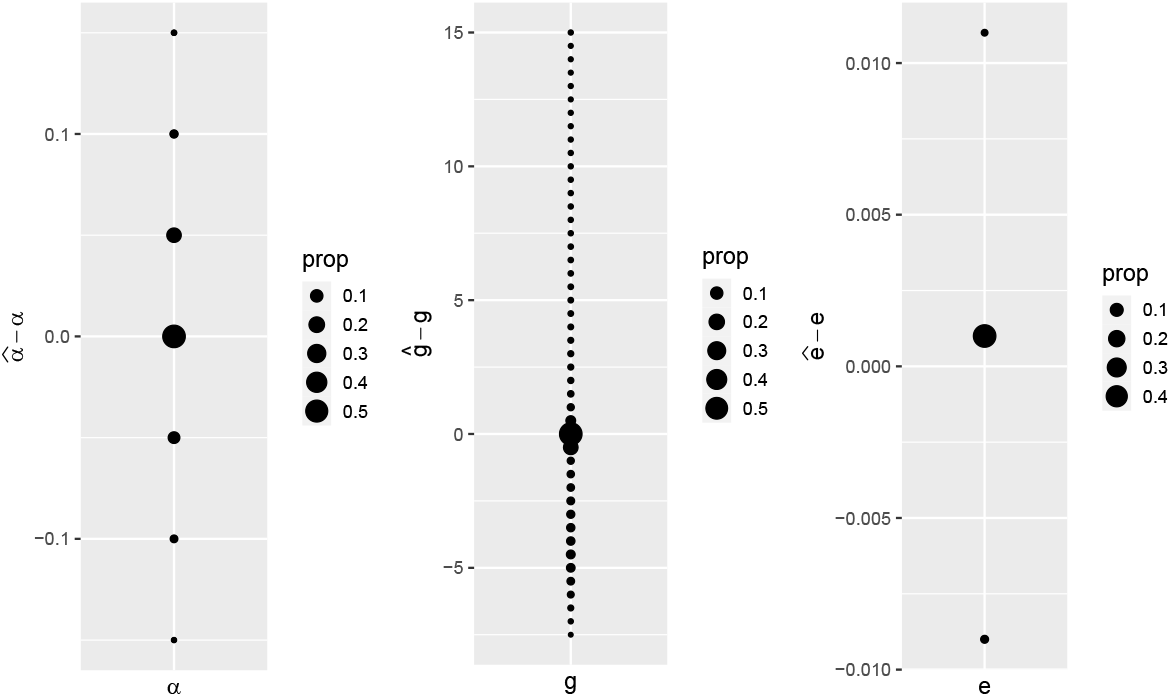
Error for estimating parameters for Beta coalescents with exponential growth and allele misorientation across the parameter grid for (*α, g, e*). Sample size *n* = 100, 50 independent loci with 100 mutations on average. 500 simulations were performed per parameter triplet.

### 3.2 Data analysis

The simulations demonstrate that the method is able to retrieve the correct model, and also correctly estimate the parameters of the MMC, provided that there is enough signal in the data. Next, we applied the method to 45 real SFS from 45 distantly related taxa. We first tested how many datasets are better fit by an MMC model than by a Kingman model, then tested the goodness of the MMC fit and estimated MMC parameters for real data.

#### MMC fits better than Kingman

First, we assessed the fit of each SFS to both MMC models and the Kingman coalescent, with exponential growth and misorientation. Using the Bayes Factor criterion, we selected the best fitting model for each empirical SFS in our dataset (Table 2). A large majority (73%) of the SFS produce a better fit to MMC models than to the standard Kingman coalescent model. The best model is most frequently the Beta-coalescent (51%), followed by the Kingman coalescent (27%) and the Psi-coalescent (13%). In a few cases, both MMC models produce a better fit than the coalescent, but we cannot distinguish the best fitting MMC (9%).

#### MMC is sometimes a good fit

While we show that MMC models produce better fits than the Kingman coalescent across many species, this could be because no model fits well. To test whether the best fit coalescent model is indeed a good model to predict the observed SFS, we calculated Cramer’s *V*, a measure of goodness-of-fit appropriate for variable contingency tables (e.g., SFS with different sample sizes across species). Combined with visual inspections, (supplementary files 1,2), we designed grade categories from ‘very accurate’ fit to ‘very poor’ fit, as following: A: *V* ∈ [0: 0.033[, B: *V* ∈ [0.033: 0.066[, C: *V* ∈ [0.066: 0.1 [and D: *V* ∈ [0.1: ∞[. Importantly, the MMC models fit well to 71% of data sets: 32/45 SFS have grades A or B on Table 3. This demonstrates that not only is MMC a better choice than Kingman on statistical grounds but also that it appears as a good model to predict patterns of diversity for a large majority of species.

**Table 3:** Distribution of goodness-of-fit grades of the best-fitting models for the 45 collected SFS. Calculated from Cramér’s *V*, A: *V* ∈ [0: 0.033[, B: *V* ∈ [0.033: 0.066[, C: *V* ∈ [0.066: 0.1 [and D: *V* ∈ [0.1: ∞[.

| Model \ Grade | A | B | C | D | Total |
| --- | --- | --- | --- | --- | --- |
| Kingman | 2 | 2 | 3 | 5 | 12 |
| Beta | 8 | 11 | 4 |  | 23 |
| Psi | 2 | 4 |  |  | 6 |
| MMC |  | 3 | 1 |  | 4 |
| Total | 12 | 20 | 8 | 5 | 45 |

#### The amount of multiple mergers greatly varies among species

The MMC models we use vary in the extent of multiple mergers, from star-like to Kingman-like, scaled by a single parameter (*α* and Ψ respectively for the Beta- and Psi-coalescent). To determine whether the model fits suggest an appreciable level of multiple mergers, we next explore the estimated parameters for MMC models. Of the 45 empirical SFS we analyzed, 68% (31/45) have 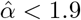 under the Beta-coalescent, which suggests a non-trivial frequency of multiple mergers, and implies something that is not captured by the SNM is occurring in these species (Table A.4, *α* estimates of all data sets, including those where the Kingman or Psi-coalescent are the best fit model). Nonetheless, estimates of *α* and Ψ are both skewed towards values that approach the Kingman coalescent (2 and 0, respectively), despite covering the full range of values across the tree of life (Fig 2, Fig A.7).

**Figure 2:**
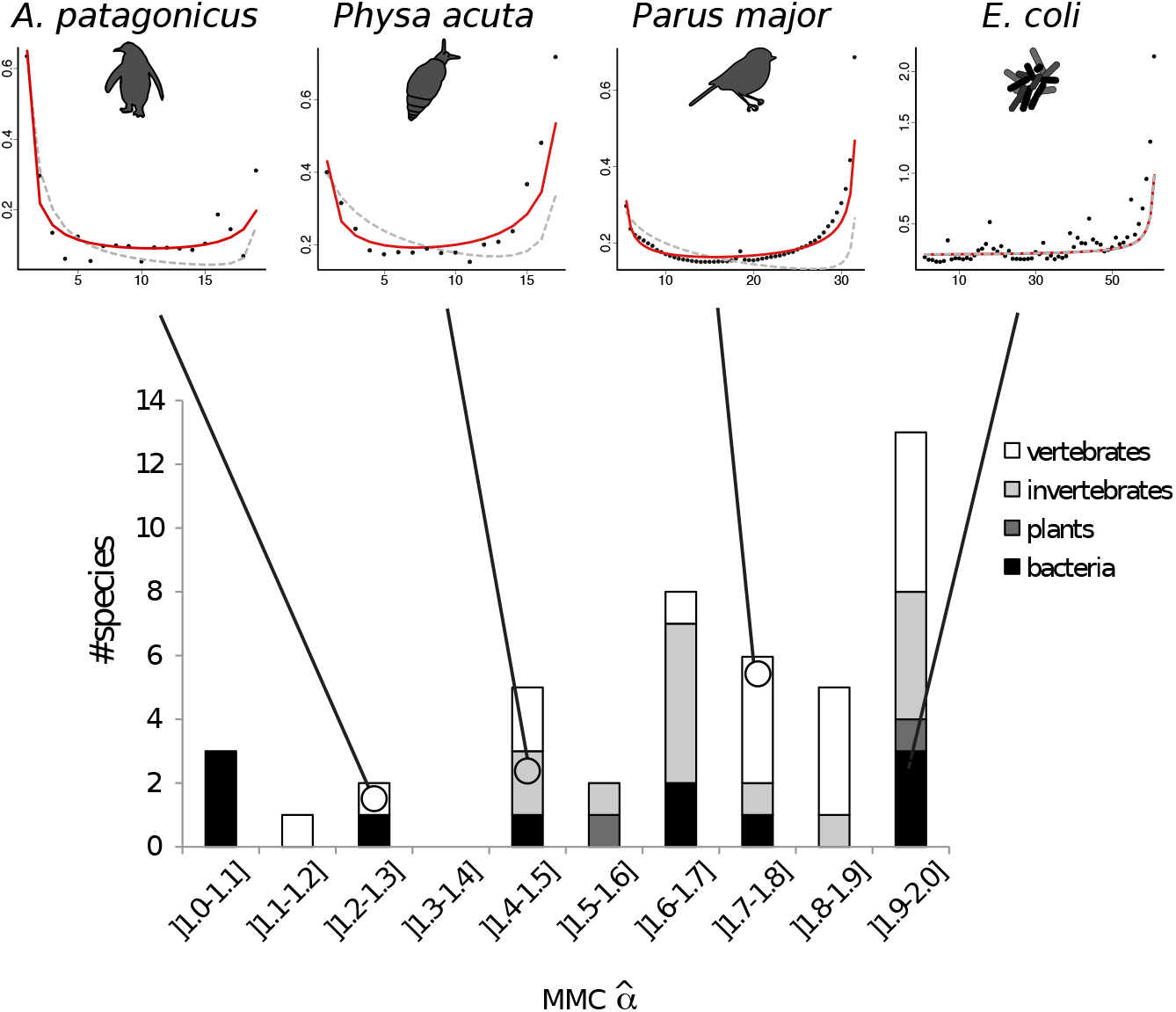
Distribution of *α* in function of the order of the species. The four top panels represent transformed *ϕ*-SFS (*ϕ_i_* = *iξ_i_* as in [Ach09, LLA17]) for four species from different taxa: two vertebrates *Aptenodytes patagonicus* (left) and *Parus major* (center right) an invertebrate *Physa acuta* (center left), and a bacteria *Escherichia coli* (right). Black dots are the observed values, grey dotted lines are the best fits under the Kingman’s coalescent model and red lines are the best fits under a Beta-coalescent model.

#### Assuming a Kingman coalescent leads to an overestimation of the growth rate

One potential impact of using the standard Kingman coalescent instead of better-fitting MMC models is the incorrect estimation of other parameters, including aspects of demography. To explore this issue, we compared the estimated growth rate and misorientation error assuming a Kingman model rather than an MMC model. We observe that the growth parameters are often higher when inferred under the the Kingman coalescent than in either of the MMC models (Table A.4), although estimates of g tend to converge in empirical datasets where the MMC parameter estimate approaches Kingman (Fig A.5a). This mirrors previous results of compensating the effect of MMC when inferring under a Kingman coalescent by estimating a higher growth rate in our scenario without allele misorientation, see e.g. [MHAJ18].

In contrast, the allele misorientation parameters *e* are almost identical between the Kingman model and the MMC (Fig A.5b), which may be a consequence of adding a second, coalescent-model-free estimation method for e to the pseudolikelihood 2. This suggests that for datasets with frequent multiple mergers, assuming a Kingman model may lead to overestimating *g*, but is not likely to impact estimates of *e*.

#### Both MMC models have similar parameter estimates

Finally, we compare the estimations of both MMC models to see whether using one or the other would result in qualitatively different conclusions. The parameters inferred under the two MMC models are highly correlated. The multiple merger parameters *α* of the Beta-coalescent and Ψ of the Psi-coalescent are negatively correlated, as expected from their definitions (Fig A.6a, Spearman correlation: *ρ* = –0.73). The estimated growth and misorientation parameters are highly positively correlated (Spearman correlations *ρ* = 0.74 and *ρ* = 0.96). The case of *Clostridium difficile* is a notable exception. The best model inferred is the Kingman, consistent with 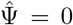 inferred for the Psi-coalescent, but for the Beta-coalescent 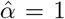, the strongest MMC component, is estimated. However, this discrepancy is likely due to statistical noise: the data set is very small (192 mutations in a sample size *n* =11) and the species has a very low recombination rate.

### 3.3 Code and data availability

All simulation and analysis codeis available upon request and additionally will be made available as a public GitHub repository upon publication.

## 4 Discussion

In this study, we show that unfolded SFS for large variety of species show a characteristic U-shape, which is inconsistent with the expectations of the standard neutral model using the Kingman coalescent. One possible explanation for this observation is the prevalence of MMC and MMC-like genealogies in real populations. To explore the role of MMCs in these data, we develop a statistical framework to detect MMC models. Using simulated data, we show this approach has good power to detect the correct MMC model and estimate its parameters, provided that the data are informative enough. Using real SFS collected from 45 species across the tree of life, we further show the MMC models are a better fit than the Kingman coalescent in most species, even when population growth and orientation errors are additionally modeling, although in some cases the MMC parameter suggests approximately Kingman behavior. In the following, we discuss some possible biological implications of these observations.

### Chosen multiple-merger models, alternatives and limitations

We chose two commonly used haploid multiple-merger models, the Beta- and the Psi-coalescent, which were previously associated with sweepstake reproduction in the literature [EW06, Sch03]. However, these MMC models may also originate either from alternative neutral processes or from selective processes. Indeed, the Beta *n*-coalescent with *α* = 1 is known as the Bolthausen-Sznitman *n*-coalescent and it (resp. a slight variant of it) emerges in a variety of models with rapid selection [BD13, NH13, DWF13, BBS13, Sch17]. The Beta-coalescent has also been associated with range expansions [BHK21]. In addition, Psi *n*-coalescents have been successfully used as proxy models for detecting regions experiencing positive selection [HJ20].

While Beta- and Psi-coalescent models are linked to several biological properties potentially present in a considerable number of species, these are not the only MMCs used to model biological populations. For instance, in the modified Moran models presented above, one can let the Ψ be random, leading to another more general class of MMC that also belongs to the family of Λ-coalescents [HM13], which is a generally good candidate for sweepstakes reproduction. Other alternative models exist that more closely mimic recurrent selective sweeps [DS05] or appear as variants of Psi- and Beta-coalescents, but for diploid reproduction [BCEH16, BLS18, KB19].

We have chosen to evaluate two simple classes of coalescent processes which interpolate between the two extreme tree shapes – a purely bifurcating Kingman tree (Ψ = 0 or *α* = 2) and a star-shaped tree (Ψ = 1 or *α* = 0). Alternative multiple merger models could potentially be (mis)identified as Beta- or Psi-coalescents, as previously shown [FSJ21]. Our method should thus still be able to detect multiple merger signals even if caused by processes that lead to another MMC. Assessing further which MMC models are best fitting for biological populations could be informative [MGF20]. In this regard, our inference approach is based on computing *E*(*T_i_*) from Eq. (2) via the method from [SKS16], so it can easily be extended to incorporate most multiple merger models (any Λ- or Ξ-coalescent) and any demographic histories, by replacing the Markov transition rate matrix of the coalescent and the population size profile *ν*.

To assess the quality of our inference method, we used a simplified approach where unlinked loci are assumed to be independent. This is not always true for MMC models (see [BBE13] and Appendix A.8), especially for Psi-coalescents caused by strong sweepstake reproduction events with Ψ well above 0. Thus, the real error rates of our techniques could be higher than anticipated by our simulation study. However, this potential increase in error rates can be offset by the presence of datasets that are larger than those assumed in our simulation study. Additionally, due to our reliance on the expected SFS entries – which are averages over the tree space – our inference method (and also our goodness-of-fit assessment) should perform worse (given identical sample sizes and mutation counts) when used on species with small genomes and low recombination rates. This tendency is clearly visible in the goodness-of-fit tests of multiple bacterial data sets.

### Non-extreme demography alone cannot generate U-shaped SFS

The Kingman coalescent for a population undergoing non-extreme demographic changes corresponds simply to a monotonic time rescaling of the standard Kingman coalescent. Non-extreme changes mean that the order of the number of generations compressed to form one unit of coalescent time (which depends on the probability that a coalescence event happens in a generation) is not affected by the demographic changes. For the MMC models employed, this is for instance satisfied if the population size stays of the same order (N) throughout generations. If this is true, changes in population size correspond to changes in waiting times, but not topology, of the tree. The expected SFS for a large population and a large sample is a linear function of the expected population-level waiting times *c_k_* (for the next coalescence of *k* lineages) with a simple analytical form:

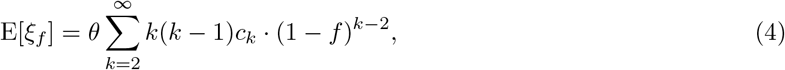

where *ξ_f_* is the number of variants at frequency *f*. Since the expected waiting times are positive *c_k_* > 0, all coefficients in this expansion are positive. This means that the spectrum has a positive value, negative derivative, positive convexity (second derivative), etc., so it is a completely monotonic function (‘no bumps’). More details are provided in Appendix A.11. As it is monotonically decreasing with *i*, U-shaped spectra cannot occur as a result of any non-extreme demographic dynamics alone. Note however that extreme changes in population size violate this and may lead to multiple merger genealogies [BBM^+^09, CPSJ22].

### Alternative processes leading to *U*-shaped SFS, further confounding factors

Our model directly incorporates MMC genealogies, exponential growth combined with allele misorientation as sources of the U-shape of the SFS. However, other potential factors can also influence the SFS and produce SFS with similar shapes. We further discuss here two particularly notable factors, population structure (e.g. gene flow or admixture) and biased gene conversion.

First, to explore population structure, we performed a PCA analysis of all datasets, followed by a *k*-means clustering (results in Table A.6). Importantly, among the 11 species that display a clear pattern of genetic structure, only 6 have an observed U-shaped SFS that is well fitted (grades A and B) by an MMC model. Furthermore, among the 14 species with no clear structure, 10 have an observed U-shaped SFS well fitted by an MMC model. This suggests that population structure is not the main cause of the U-shape of the observed SFS. Additionally, many species with clear structure have low goodness-of-fit grades (C and D), suggesting that none of the models we compare are a good fit to these datasets. We however note that 8/11 species with a clear structure pattern are Bacteria. Indeed for the small genomes with low recombination rate (in Bacteria recombination preserves long distance linkage), the apparent structure does not necessarily equate with population structure, but may instead arise from the limited number of genealogies. At the limit, a single Kingman tree would result in a clear structure pattern due its long internal branches.

To check for the effect of biased gene conversion, we built alternative SFS only based on a subset of unbiased mutations that are immune to biased gene conversion (details in A.7, the unbiased SFS are added in supplementary files 1,2). Many of these unbiased SFS were only slightly changed, and many kept their U-shape. However 6 species (A. *cunicularia, F. albicollis, E. garzetta, P. maior, O. edulis, P. troglodytes e*., all but one vertebrates) lost their U-shape. Two have a small sample size (*E. garzetta*) or a low multiple merger component estimate (*F. albicollis*). For these species, it is nonetheless possible that the U-shape is caused by biased gene conversion.

In a very conservative approach, among the 17 data sets showing robust and strong MMC signals (category A, B in Table 2, with *α* ≤ 1.8 or Ψ ≥ 0.04 and sample size ≥ 20), 6 cases may arise due to structured genetic diversity (A. *baumannii, D. melanogaster, H. pilori, O. edulis, P. aeruginosa* and *S. aureus*) and 3 more lose their characteristic U-shape when biased gene conversion is accounted for (A. *cunicularia, P. maior, P. troglodytes*; *O. edulis* being in common). Thus, 8 species have strong support for MMC models with population growth. We believe that at least for these cases (and likely for more), neutral sweepstake reproduction, frequent selection, or other factors that can produce MMC-like genealogies ought to be seriously considered as underlying drivers of their genetic diversity.

Importantly and more generally, among the 32 species that display a good statistical fit (with grades A and B), 28 point to MMC models whereas only 4 point to a Kingman coalescent. Noting that MMC models encompass the Kingman coalescent as a special case, our results support the view that MMC models may often constitute better reference models.

### MMC and biological properties

Although we only analyzed a small number of species sampled non-uniformly across the tree of life, we often observed signatures of multiple merger-like events. Reassuringly, our analysis supports multiple merger genealogies for *Mycobacterium tuberculosis*, which was recently proposed in [MASSJ20] and [MGF20] (the non-optimal goodness-of-fit likely stems from a small and essentially nonrecombining genome). The strongest multiple merger effects estimated within the class of Beta coalescents (*α* ≤ 1.1) were found in two bacterial pathogens with low or intermediate recombination rates (M. *tuberculosis* and *P. aeruginosa*). There also does not seem to be a meaningful correlation between MMC effects and overall genetic diversity (Figure A.21, Table A.7). We stress that links between MMC model parameters and biological properties are not always obvious. For example, while reproduction sweepstakes can lead to both Beta- and Psi-coalescents, it is not straightforward to translate the parameters *α* and Ψ into realistic offspring distributions. For instance the Psi-coalescent model hypothesizes that an occasional individual contributes a fraction Ψ of the next generation, though examples of such a single-individual contribution are not biologically likely. Still, the coalescent approximations do fit well to data. Importantly, different reproduction models can result in the same model on the *coalescent time scale*. The large families of the MMC models could result from the rapid accumulation of coalescences over multiple generations instead of in a single one.

### Conclusion

We analyzed genomic data for 45 species across the tree of life, and showed that many exhibit a U-shaped SFS. By developing a statistical approach to distinguish the genetic signatures of different potential sources of this U-shape: allele misorientation and MMC genealogies, together with exponential population growth, our results show that while some U-shaped SFS are well-described by only allele misorientation, the majority are better described by models that include an MMC component (28 point to MMC and only 4 to Kingman coalescent, with the rest inconclusive). However, distinguishing true MMC from MMC-like processes remains challenging. For example, both biased gene conversion (evident for 6 species) and population structure (clear for 11 species, many of which had no U-shapes) could also generate U-shaped SFS, and appear to be plausible explanations for the observed data of certain species. MMC models with simple growth nonetheless represent an excellent fit for at least 8 species.

This study thus invites both closer inspection for the species at hand, but also suggests that MMC genealogies may appear in a wider range of species than previously reported (e.g., a few marine species and multiple human pathogens). For such species, their biological properties likely render MMC rather than Kingman models as the more fruitful analysis framework, highlighting the importance of further developing both theory and statistical inference procedures under these lesser-used models [Wak13].

## Supporting information

Supplementary 3 - per chromosome Droso

Supplementary 3 - BIC plots for DAPC

Supplementary 3 - PCA plots w. DAPC colours

Supplementary 1 - SFS and fits

Supplementary 2 - tranformed SFS and fits

## Ackowledgments

We thank NSF for support through the DEB-1754397 to Timothy B. Sackton and DFG for support through FR 3633/2-1 (within Priority Program 1590: Probabilistic Structures in Evolution) to Fabian Freund. Jeffrey D. Jensen was supported by National Institutes of Health grant R35GM139383. We would like to thank Allison J Shultz and Brian J Arnold for help with producing VCFs for several bird species. The authors acknowledge the support by the state of Baden-Wurttemberg (Germany) through bwHPC.

## Appendix A Appendix

### A.1 Reproduction models linked to MMC and time scalings

The coalescent approximations from the main text are the coalescent limits for population size *N* → ∞ (with changed time-scale) of genealogical trees given some reproduction model. We focus on Cannings models [Can74] of reproduction, which are discrete-generation models, usually with fixed population size, and exchangeable offspring numbers between individuals. This is a standard model choice, see e.g. [Sag99], and the modified Moran models present in the Methods are Cannings models. Different reproduction models can lead to the same coalescent limit, e.g. the Wright-Fisher and Moran model both lead to Kingman’s coalescent. If the coalescent limit is identical for two constant population size reproduction models (and the number of generations to form one coalescent time unit is of order *N^η^*), we can describe the limit as in Eq. (1) for both models. Thus, adding population size changes can still lead to a difference in coalescent limit via changing the power *η* of the population size ratio *ν*. For instance, *η* = 2 for the standard Moran model but *η* = 1 for the Wright-Fisher model (Λ the point mass in 0 in both cases). In the case of exponential growth (on the coalescent time scale), we see that the factor influenced by *η* in Eq. (1) equals *ν*(*t*)^−η^ = exp(*ηgt*). This means that we can still interpret parameters assuming one reproduction model (model 1) leading to the coalescent (with scale parameter *η* = *x*_1_) under the assumption of an alternative reproduction model (model 2 with scale parameter *η* = *x*_2_) by simply re-scaling the exponential growth parameter *g* from model 1 as 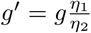. For instance, a growth rate of 2*g* in the Wright-Fisher model corresponds to a growth rate of *g* in the Moran model.

For our two MMC models, this also means that we could analyze the models based on alternative reproduction models. For instance, we set *γ* = 1.5 for the discrete reproduction model leading to the Psi-coalescent, but we could also choose any other 1 < *γ* < 2. For the Beta-coalescent, there is indeed a very appealing alternative reproduction model due to Schweinsberg [Sch03]. This alternative model assumes, for 1 < *α* < 2, that each individual at each generation independently produces a number of offspring following a power law distribution with tail parameter *α* (infinite variance), and that the next generation (the individuals surviving long enough to reproduce) is sampled from these offspring. In this model, one unit of coalescent time corresponds to an order of *N*^*α*–1^ generations. As discussed above, if *α* > 1, we can interpret any growth rate *g* when seeing the Beta-coalescent as the genealogy model based on the modified Moran model with growth rate 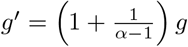 under Schweinsberg’s model.

### A.2 Properties of the reproduction model underlying the Beta-coalescent

The modified Moran model with distribution given on p.4 leading to the Beta(2 – *α, α*)-coalescent was introduced in [HM13] and the properties of *U* have been additionally analyzed in [IM02]. *U*, or more precisely *U_N_* since it depends on *N*, is distributed as the number of lineages merged at the first merger in a Beta(2 – *α,α*)-coalescent starting with sample size *N*. Since when increasing the sample size, the first merger can only include more lineages, *U_N_* ≤ *U_M_* holds for *M* ≥ *N* (we can assume that, with increasing sample sizes, coalescent events just add branches to the tree from smaller sample sizes, see [Pit99]). This then also holds for the expected values, so 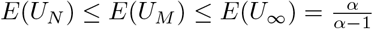, where *U*_∞_ is the limit of *U* for *N* → ∞. See [HM13, p.9] for the existence of the limit, whose properties including its mean are described on the cited page combined with [IM02, p.226], including its infinite variance. See Table A.1 for some properties of *U_N_* for different *N* and *α*, computed from the definition of *U_N_* and the listed properties of its limit.

**Table A.1:** Properties of *U_N_* for the modified Moran model underlying the Beta-coalescents. *x_min_*: Minimal integer x such that *P*(*U* ≤ *x*) ≥ 0.99.

| $N$ | $\alpha$ | $E(U_N)$ | $\sqrt{\text{Var}(U_N)}$ | $P(U \leq x_{\min}) \geq 0.99$ |
| --- | --- | --- | --- | --- |
| 5000 | 1.1 | 6.54 | 46.87 | 62 |
| 10000 | 1.1 | 6.83 | 64.24 | 62 |
| 25000 | 1.1 | 7.20 | 97.26 | 62 |
| ∞ | 1.1 | 11.0 | - | 62 |
| 5000 | 1.5 | 2.96 | 9.39 | 15 |
| 10000 | 1.5 | 2.97 | 11.27 | 15 |
| 25000 | 1.5 | 2.98 | 14.29 | 15 |
| ∞ | 1.5 | 3.00 | - | 15 |
| 5000 | 1.9 | 2.11 | 1.39 | 4 |
| 10000 | 1.9 | 2.11 | 1.50 | 4 |
| 25000 | 1.9 | 2.11 | 1.64 | 4 |
| ∞ | 1.9 | 2.11 | - | 4 |

### A.3 Mathematical derivation of the pseudolikelihood function Eq. (2)

We follow the derivation from [EBBF15, Eq. 11]. We want to compute the likelihood of seeing the observed SFS *s*_1_,…, *s*_*n*–1_ under a given coalescent (here a Beta-n-coalescent or a Psi-n-coalescent with exponential growth, but the derivation works for any coalescent model). Let 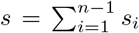 be the number of observed segregating sites. We assume the fixed-*s* approach, e.g. we assume that the distribution of the SFS is given by placing s mutations at random on the genealogical tree. Under the fixed-s assumption, the probability of observing the SFS is given by the multinomial distribution

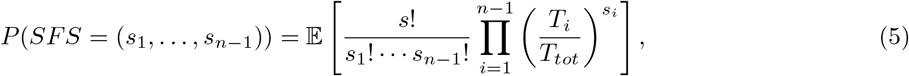

since a segregating site has mutant allele frequency *i* if it lands on a branch that supports i leaves (*T_i_* is the sum of lengths of branches supporting *i* leaves, 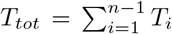 is the total length of the genealogy). Under further assumptions of independence of the different fractions 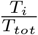 of the total branch length and approximating 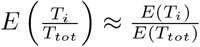, we have a further approximation

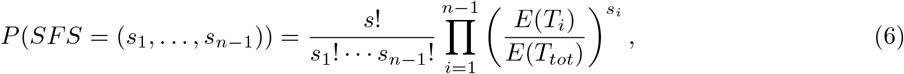

Next, we consider the addition of a misorientation probability, *e*, describing the switch of ancestral and derived states. Eq. (6) constitutes a multinomial distribution, which can be interpreted as throwing *s* balls into compartments 1,…, *n* – 1, where compartment *i* is hit with probability 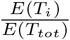. Misorienting the allele in this interpretation means that a ball that originally lands in compartment *i* is placed in compartment *n* – *i* instead. If this happens with probability *e*, a ball consequently lands in compartment *i* with probability 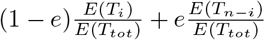. So the probability to observe a specific SFS when ancestral and derived types can be confused is

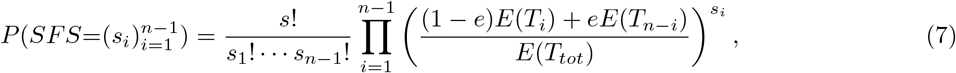

where 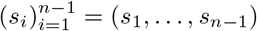 is the observed SFS. This is again a multinomial distribution.

Simulations showed that inferring parameters via a pseudolikelihood approach based on Eq. 7 tends to overestimate *e* to fit the U-shape. To counteract this, we couple this equation with an alternative estimation of *e* by using polymorphic sites discarded in the process of polarizing the SFS due to having a third allele in the outgroup. As described in [Lap17, Section 4.2] or [BD03, p. 1620], these sites carry information about *e*. Let *S*_0_ = *S* + *S*_≠_ be the total number of biallelic SNPs in the sample, where *S* is the (random) number of sites where the outgroup does not show a third allele not observed in the sample (left and central trees in figure A.1) and *S*_≠_ the number of sites where it does (right trees in Figure A.1). Observe that s is the observed outcome of S, the total sum of the observed SFS.

**Figure A.1:**
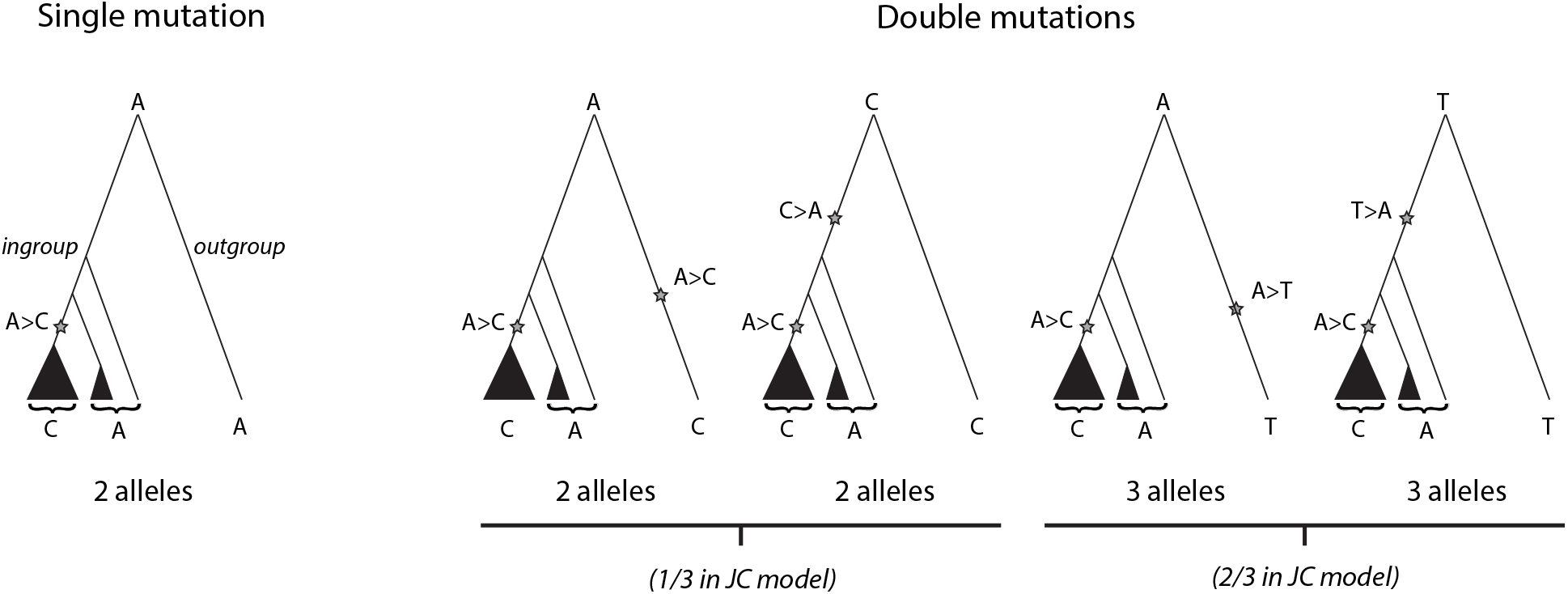
Sketch of trees with mutations to illustrate how tri-allelic sites relates to the probability of misorientation. Left and central trees have one of their variant equal to the outgroup (counted in *s*) whereas the left tree have 3 differents alleles (counted in *s*_≠_). Under a Jukes and Cantor setting, the expected number of misoriented variants (central trees) equals half the number of tri-allelic sites.

Consider a polymorphic site in the sample (ingroup). If one of the allele is the same than the outgroup, there was either a single mutation (most likely for closely related outgroup) or two mutations, the second one masking the effect of the first (central trees of figure A.1). In the Jukes-Cantor model where all mutations are equally likely, if the probability of having a specific mutation *X* > *Y* masking the effect of the first is *p*, then the probabilties of observing 2 alelles is *p* and the probability of observing 3 alleles is 2*p*. We emphasise that while we assume an infinite sites model within the sample, we allow reverse mutations here due to the considerably longer branch lengths.

Following this, we can compute the probability *P*(*S*_≠_ = *s*_≠_|*S*_0_ = *s* + *s*_≠_) that we observe exactly *s*_≠_ sites which are biallelic within the sample but have a third allele for the outgroup. This is simply binomial sampling from *S*_0_ biallelic sites with success probability 2*p*. We can also express this probability in terms of the misorientation probability *e*. Let *v* ∈ *SFS* be the event that a bialllelic site (variant) can be polarized via outgroup (which means it has one of the two alleles of the sample also for the outgroup) and *mis*(*v*) the event that the ancestral state of v is misidentified. The probability that a site of the SFS can be polarized (displaying one allele similar to the outgroup) is 1 – *p*. We have

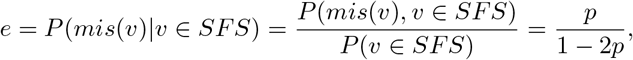

and thus equivalently 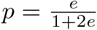. This leads to

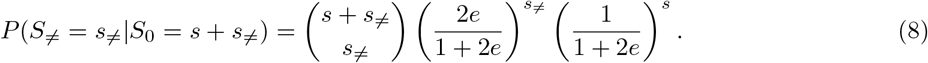

We now assume a composite likelihood, multiplying Eqs. (7) and (8). Conditional on observing *s* + *s*_≠_ segregating sites from which *s* can be polarized via outgroup and form the SFS, the pseudo-likelihood of observing a specific SFS is given by Eq. (2).

#### Remark A.1.

*Eq*. (8) *shows that S given S*_0_ *is binomially distributed with S*_0_ *draws with success probability* 1 – 2*p* = (1 + 2*e*)^-1^. *Let X*(*S*_0_) *be a r.v. with this distribution. We will use this to simulate S*_0_ *based on S and the mis-classification probability e: The maximum likelihood estimate for the number of trials S*_0_ *of the binomial r.v. X*(*S*_0_) ~ *Bin*(*S*_0_, (1 + 2*e*)^-1^), *in the sense of maximising P*(*X*(*S*_0_) = *s*), *is* 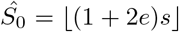, *since*

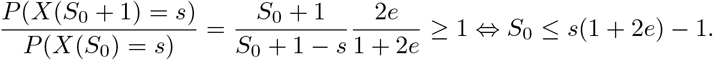

*We will use this estimate to simulate a reasonable s*_≠_. *If we simulate a SFS with s mutations, and we flip each mutation in it with frequency e from class i to n – i, we then simulate s_≠_ as a binomial draw from* 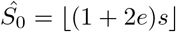 *Bernoulli r.v.’s with success probability* 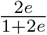. *This is denoted as the* 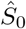 *approach*.

### A.4 Cramer’s *V* as a goodness-of-fit measure

Our assumptions leading to Equations (6), (7) can be interpreted that each variant observed for the SFS is sampled from a multinomial distribution from the ‘true’ allele frequency spectrum. In the following, we denote the multinomial approximation of the SFS entry frequencies, the ‘true’ spectrum, by (*p*_1_,…, *p*_*n*–1_). Since assuming sampling from a multinomial distribution is also the statistical model behind the *χ*^2^ goodness-of-fit test, we chose the effect size measure Cramer’s *V* [Cra16, ch. 21] of this test, defined as

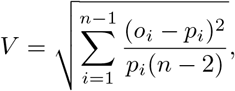

to quantify the lack of goodness of fit (o¿ is the observed frequency of mutations with frequency *i/n* among all mutations). This measure can be interpreted as a dimensionless version of the *χ*^2^ test statistic, since the mutation counts do not enter, just the mutation frequencies and the additional factor *n* – 2 corrects for unequal sample sizes.

### A.5 Assessing estimation errors

#### A.5.1 Simulation and inference setup

As a rough approximation of a genome, we simulated 100 independent loci (ignoring the fine structure of weakly physically linked loci and long range LD, see Appendix A.8). This means that the genealogical trees of the loci are independent and follow the same tree distribution, e.g. realisations of a Beta coalescent with exponential growth with rate *g* and coalescent parameter *α*. The mutations on each tree are independent of all trees (and mutations on other trees) and given by a Poisson process with rate 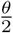. We assumed three different sample sizes *n* = 20, *n* = 25 and *n* = 100. For each locus, we set the mutation rate so that on average 50 mutations appear, *i.e*. we set *θ* = 100/*E*[*T_tot_*] (generalized Watterson estimate), where *T_tot_* is the sum of all branch lengths of the locus’ genealogy. Mutations are interpreted under the infinite-sites model, resulting in simulated SNP sequences (ancestral vs. derived type). For each SNP, we then flip ancestral and derived allele with probability *e*. We simulate 500 SNP sequences as described above for each combination of coalescent parameter *α* or Ψ, growth rate *g* and misorientation probability *e* from the following two sets (the first set has *α*, Ψ and *g* on the inference grid, the second uses off-grid values).

- Set 1: equidistant *α* ∈ {1, 1.05, 1.1,…, 2} and Ψ ∈ {0.05,0.1, 0.2,0.3,0.4,0.5,0.6,0.7,0.8, 0.9} Set 2: *α* ∈ {1.025,1.325}, Ψ ∈ {0.025, 0.075} (additionally Ψ = 0.005 for *n* = 25)
- Set1: *g* ∈ {0,0.5,1,10}, Set 2: *g* ∈ {0.25, 2.25,11.25}
- Set 1: *e* ∈ {0,0.01, 0.05, 0.1} (essentially on grid), Set 2: e ∈ {0,0.015,0.045,0.095}

To infer via Eq. (2), we also need the total number of segregating sites *s* + *s*_≠_, adding the number of segregating positions not included in the SFS due to not being able to polarize them. For this, we use the 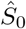 approach described in Remark A.1.

#### A.5.2 Parameter and model selection accuracy

First, for *n* = 20 and *n* = 100, we estimate parameters using Eq. 2 using the same coalescent model (Beta or Psi) on equidistant grids with *α* ∈ {1,1.05,1.1,…, 2} or Ψ ∈ {0, 0.5,0.1,…, 1}, *g* ∈ {0, 0.05,…, 25}, *e* ∈ {0.001,0.011,…, 0.201}. For this, we only used the on-grid values (Set 1). Results are shown in Figures 1, A.2 – A.4, A.8 – A.19.

Second, we assess the error of our model selection approach based on approximated Bayes factors for *n* ∈ { 20, 25, 100}. For this, we fixed different values of Ψ and *α* from Set 1 and Set 2 including *α* = 2.

We then picked 2,000 simulations at random from all parameter combinations with this fixed coalescent parameter (as described above) and performed model selection via Bayes factors as described in the method described in the main document (section 2.2). The maximum was taken on the same equidistant grids as for the parameter estimation. Expanded results are provided in Tables A.2 and A.3.

#### A.5.3 Parameter estimation accuracy - results

For inferring parameters under the Beta-coalescent or the Psi-coalescent, Figures 1, A.2–A.4 show the error distribution of all three parameters for *n* ∈ {20,100} across all simulation parameter choices. While *g* cannot be estimated precisely in some cases, *e*, Ψ and, to a lesser degree, *α*, can generally be estimated rather well, especially if sample size *n* = 100.

These errors distribute over the different parameter settings as shown in Figures A.8 – A.19. Most notably, large errors when estimating growth rates only happen if the growth rate is also large. For Psi-coalescents, we see that choosing Ψ between grid points are still mostly captured by the adjacent Ψ grid points.

**Table A.2:** Model selection via two-step Bayes factor criterion. Based on 2,000 simulations for each true model assuming 100 loci with 50 observed mutations. For each simulation, the coalescent parameter is fixed and the growth parameter g and the allele misorientation rate e are randomly chosen (*g* ∈ [0,11.25], *e* ∈ [0,0.1]). The column grid shows whether the parameters used for simulation were included in the inference grid. For details on both simulations and inference parameters see Appendix A.5. Fractions are rounded to two digits.

| sample size | true model | grid | Fraction model inferred as |  |  |  |
| --- | --- | --- | --- | --- | --- | --- |
|  |  |  | Kingman | Beta | Psi | MMC |
| $n = 20$ | $\alpha = 2$ | yes | 0.77 | 0.22 | 0.00 | 0.00 |
| | $\alpha = 1.9$ | yes | 0.37 | 0.58 | 0.02 | 0.02 |
| | $\alpha = 1.8$ | yes | 0.06 | 0.79 | 0.09 | 0.08 |
| | $\alpha = 1.625$ | no | 0 | 0.82 | 0.09 | 0.08 |
| | $\alpha = 1.025$ | no | | 0.99 | 0.01 | 0 |
| | $\alpha = 1$ | yes | | 0.99 | 0 | 0 |
| | $\Psi = 0.025$ | no | 0.14 | 0.59 | 0.15 | 0.12 |
| | $\Psi = 0.05$ | yes | 0.01 | 0.17 | 0.70 | 0.11 |
| | $\Psi = 0.075$ | no | | 0.12 | 0.82 | 0.06 |
| | $\Psi = 0.1$ | yes | | 0.04 | 0.94 | 0.02 |
| $n = 25$ | $\alpha = 2$ | yes | 0.79 | 0.21 | | |
| | $\alpha = 1.9$ | yes | 0.34 | 0.66 | 0 | 0 |
| | $\alpha = 1.8$ | yes | 0.02 | 0.91 | 0.04 | 0.03 |
| | $\alpha = 1.625$ | no | | 0.9 | 0.06 | 0.05 |
| | $\alpha = 1.025$ | no | | 1 | | 0 |
| | $\alpha = 1$ | yes | | 1 | | 0 |
| | $\Psi = 0.005$ | no | 0.55 | 0.45 | | |
| | $\Psi = 0.025$ | no | 0.05 | 0.72 | 0.14 | 0.09 |
| | $\Psi = 0.05$ | yes | 0 | 0.12 | 0.82 | 0.06 |
| | $\Psi = 0.075$ | no | | 0.06 | 0.91 | 0.03 |
| | $\Psi = 0.1$ | yes | | 0.02 | 0.98 | 0 |
| $n = 100$ | $\alpha = 2$ | yes | 0.87 | 0.13 | | |
| | $\alpha = 1.9$ | yes | 0.12 | 0.88 | | |
| | $\alpha = 1.8$ | yes | | 1 | | |
| | $\alpha = 1.625$ | no | | 1 | | |
| | $\alpha = 1.025$ | no | | 1 | | |
| | $\alpha = 1$ | yes | | 1 | | |
| | $\Psi = 0.025$ | no | | 0.92 | 0.06 | 0.02 |
| | $\Psi = 0.05$ | yes | | | 1 | |
| | $\Psi = 0.075$ | no | | | 1 | |
| | $\Psi = 0.1$ | yes | | | 1 | |

### A.6 Visual correlations between the estimated parameters

We provide a graphical view of the correlation between the parameters inferred between the Kingman coalescent and the Beta-coalescent (Figure A.5) or between both MMC models (Figure A.6).

**Figure A.2:**
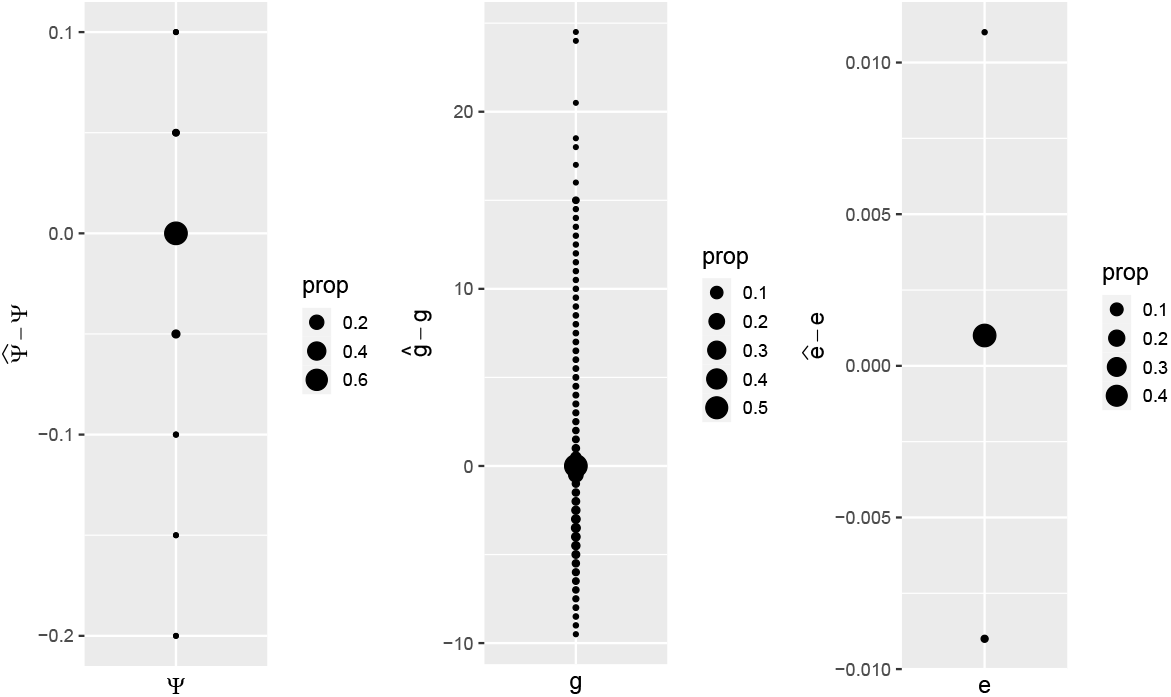
Error for estimating parameters for Psi-coalescents (*n* = 100, with growth and misclassification) across all simulation scenarios

**Figure A.3:**
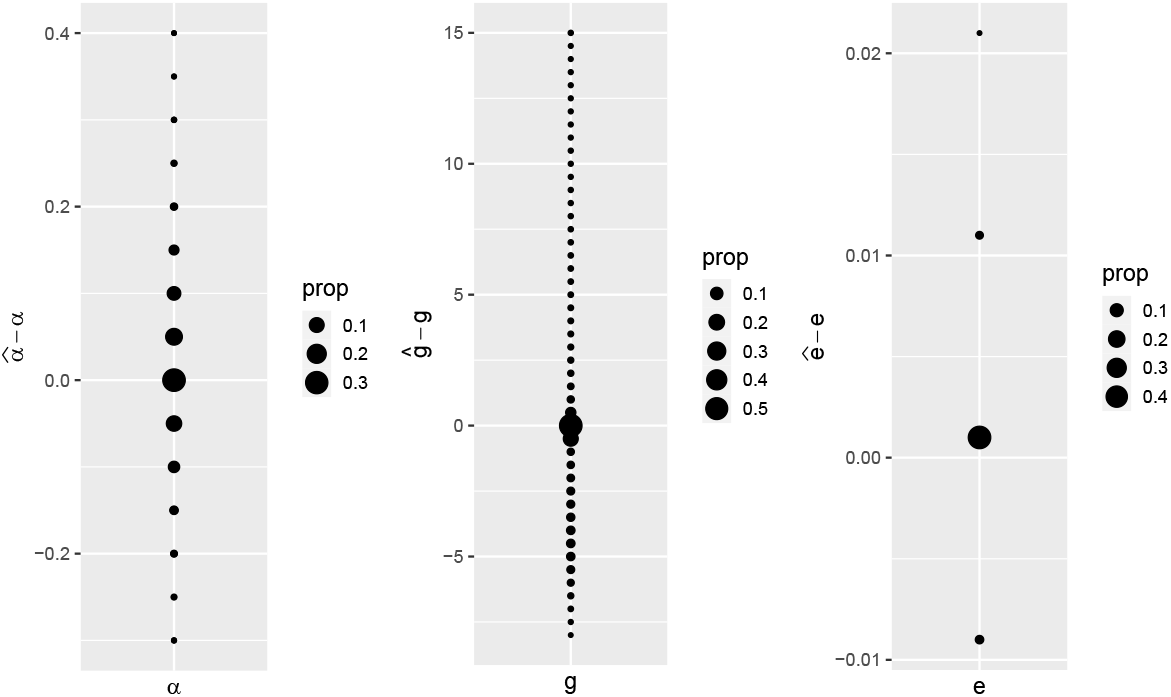
Error for estimating parameters for Beta-coalescents (n = 20, with growth and misclassification) across all simulation scenarios

**Table A.3:** Fractions of estimated parameters of model-misidentified coalescent simulations with *α* = 2. If the two-step Bayes factor model inference recorded “MMC”, the Beta parameter is reported.

| sample size | $n = 20$ | $n = 100$ |
| --- | --- | --- |
| $\alpha = 1.75$ | 0.01 | |
| $\alpha = 1.8$ | 0.04 | |
| $\alpha = 1.85$ | 0.22 | 0.04 |
| $\alpha = 1.9$ | 0.37 | 0.53 |
| $\alpha = 1.95$ | 0.34 | 0.43 |
| $\Psi = 0.05$ | 0.02 | |

**Figure A.4:**
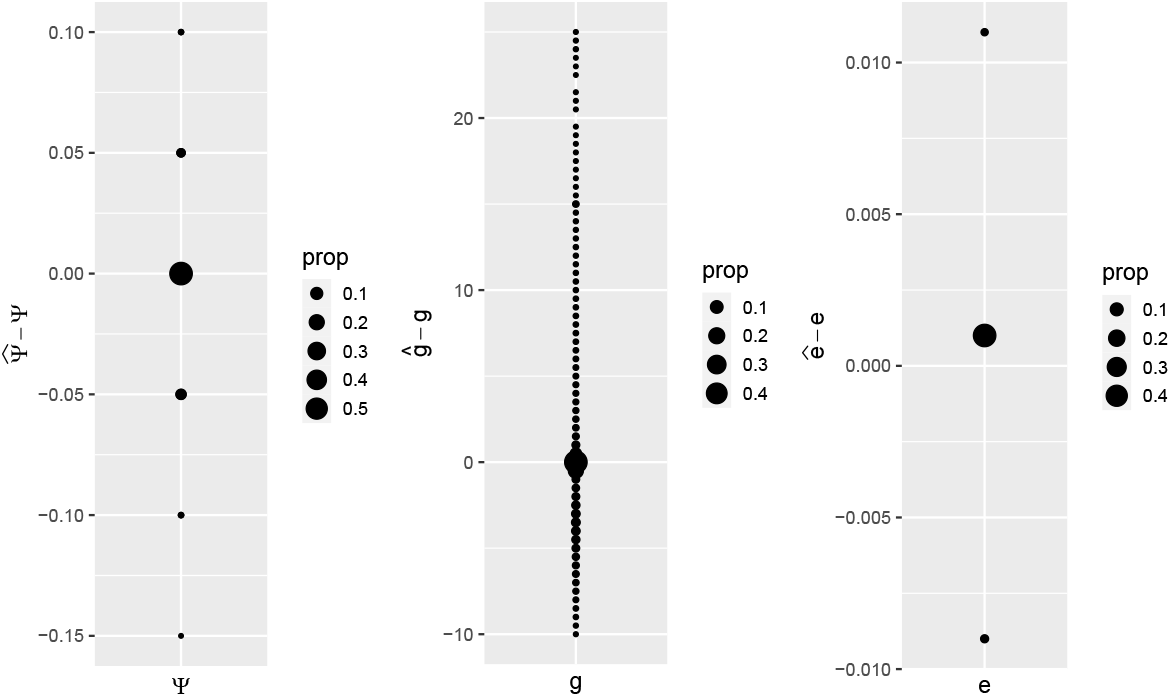
Error for estimating parameters for Psi-coalescents (*n* = 20, with growth and misclassification) across all simulation scenarios

**Figure A.5:**
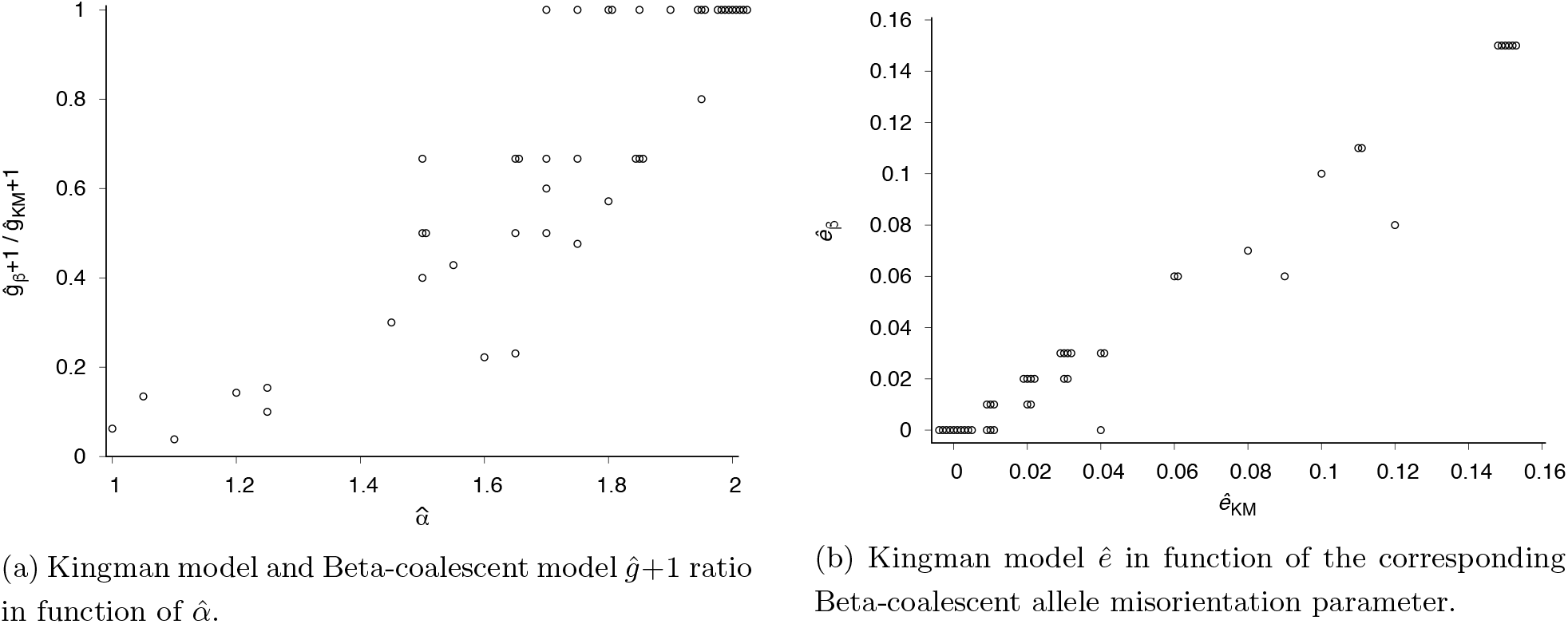
Comparison of parameters between Kingman model and Beta-coalescent model.

### A.7 Correction for GC-bias

We use the approach from [PATE18] and consider the subset of SNPs corresponding to *A* ↔ *T* and *G* ↔ *C* substitutions, which are not affected by biased gene conversion. We overlaid these neutralized SFS to the observed SFS and the predictions of the fitted models in supplementary files 1,2.

### A.8 Non-independence of unlinked loci under multiple merger genealogies

Here, we address the issue that physically unlinked loci in multiple merger genealogies still have dependent genetic diversity. For Λ-coalescents, which all our coalescent models are, the issue can be easily understood within the approximate multi-unlinked-locus model from the appendix of [Kos18]. In this model, multiple mergers result from large families appearing in a short amount of evolutionary time (see also a more thorough explanation in [MGF20]), so these families affect not only one, but all loci. Due to the model definition of MMC, each ancestral lineage can join one of such events with the same probability. Thus, if this probability is high, there will be a merger of similar size at each or nearly each locus in the genome, introducing a dependency between loci. The strength of this dependency should be correlated to the probability with which an ancestral lineage merges, following Remark 1 in [Kos18]. This probability x is generated by a Poisson process whose rate is proportional to *x*^-2^Λ(*dx*), where Λ is the associated measure of the coalescent (a Beta distribution or the point mass in Ψ for our model classes). This probability is rather small for Beta coalescents, but high for high Ψ values, see also Figure A.20.

**Figure A.6:**
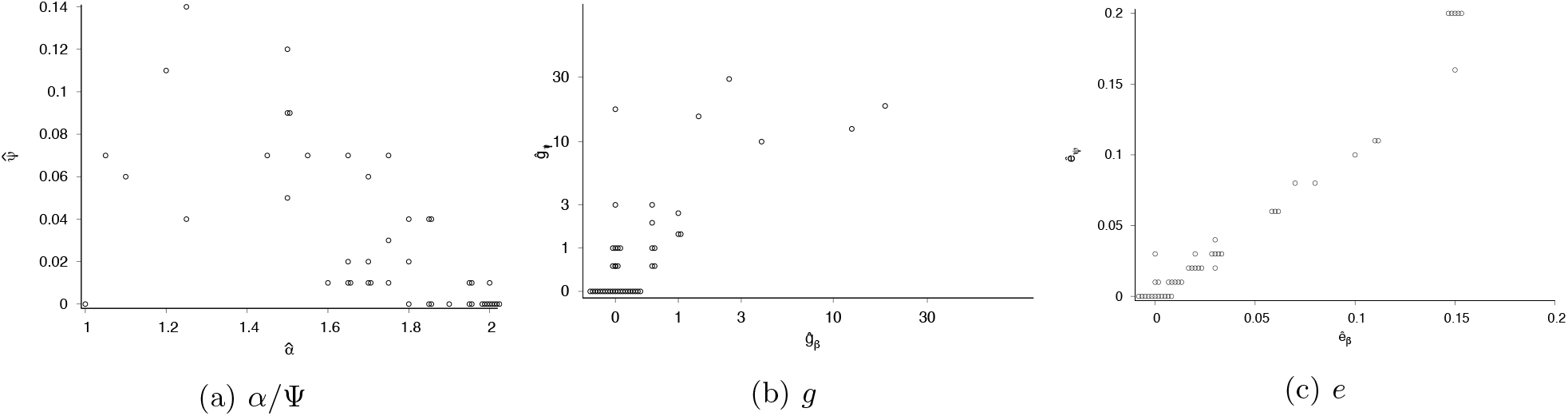
Comparison of Beta-coalescent (*x*-axis) and Psi-coalescent (*y*-axis) parameters inferred from each SFS.

### A.9 Population structure scans

We performed two simple checks for population structure: PCA and `find.clusters` from the `R` package `adegenet` [JA11]. For PCA, we coded alleles as 0 and 1, imputed missing data as the mean allele at the site, to then perform a double-centered PCA: PCA, as implemented in `adegenet`, was performed on the SNP matrix after subtracting row means and column means (and adding the overall mean), see [GJQP+19, p.20]. The approach behind `find.clusters` is to first perform a standard PCA and then group individuals by running the *k*-means clustering algorithm on the principal component coordinates for different numbers of clusters. Based on the goodness-of-fit criterion BIC, we chose the ‘optimal’ *k* as the smallest value of *k* that is visibly a local minimum (essentially the elbow criterion). For large data sets of more than 1 million SNPs, we performed the analysis with a reduced data set by filtering down the number of SNPs by only retaining each *x*th SNP where 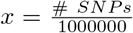, rounded to the lower integer. Results are shown in Supplementary file 3 and Table A.6. For diploids, PCA and `find.clusters` results were not qualitatively affected by performing them on either haplotypes or diploid genotypes. For *D. melanogaster*, we performed the population structure scan separately on Chromosome 2L, 2R, 3L, 3R (with filtering down as described above). For the human data, we omitted the X and Y chromosomes.

### A.10 Nucleotide diversity across the genome

We recorded the (sample) mean and standard deviation of the per-site nucleotide diversity in non-overlapping windows of 15,000 sites along the genome (resp. the sequenced part of it). For computation, we used the R package `pegas` [Par10] and `vcftools` [DAA^+^11] (for haploid data presented as vcf files, we used J. Dutheils fork of `vcftools` https://github.com/jydu/vcftools). For the human data, we omitted the X and Y chromosomes and for *D. melanogaster*, we used chromosomes 2L, 2R, 3L, 3R.

Results are shown in Table A.7 and Figure A.21.

### A.11 Effect of non-extreme demography on the SFS

The expected SFS (*E*(*S*_1_),…, *E*(*S*_*n*–1_)) for a given genealogy tree (conditional on waiting times and topology) is a linear function of the waiting times *C_k_* for the next coalescence event of the sample genealogy if *k* ancestral lineages are present, with coefficients *kP_n,k_*(*i*) dependent on the topology, where *P_n,k_*(*i*) is the probability that a random branch at level *k* has *i* descendants in the sample ([Fu95, FLW^+^17, SW08]). For a sample from a population with panmictic, neutral dynamics and finite variance in offspring number, corresponding to a Kingman coalescent where time is rescaled by a deterministic strictly monotonic function, all tree topologies are equally probable and independent of waiting times. The expected coefficients are given by 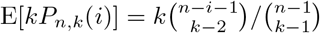. It should be noted that the assumption of monotonic time change ensures that the genealogy stays bifurcating: extreme changes in population size violate this and may lead to multiple merger genealogies.

The SFS for a large population described by the time-rescaled Kingman coalescent can be obtained as the large sample limit *n* → ∞ of the above spectrum [FKR^+^18]. For large *n*, the probability that a random lineage at level *k* takes a fraction *f* of the descendants is E[*P_n,k_*(*fn*)] → (*k* – 1)(1 – *f*)^*k*−2^*df*. Hence the continuous expected SFS is given by equation (4), which depends on the expected population-level waiting times *c_k_* = E[*C_k_*] > 0 for *k* = 2… ∞. The positivity of all coefficients in this expansion implies that for a finite expected TMRCA 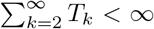, the expected SFS for populations with non-extreme demography is an absolutely monotonic function of 1 – *f* in [0, 1), and therefore a completely monotonic function of the frequency *f* in (0, 1]. This is the case for all non-extreme demographies with bounded past population size, since all of them have finite expected TMRCA.

### A.12 Psi-coalescent graphs

**Figure A.7:**
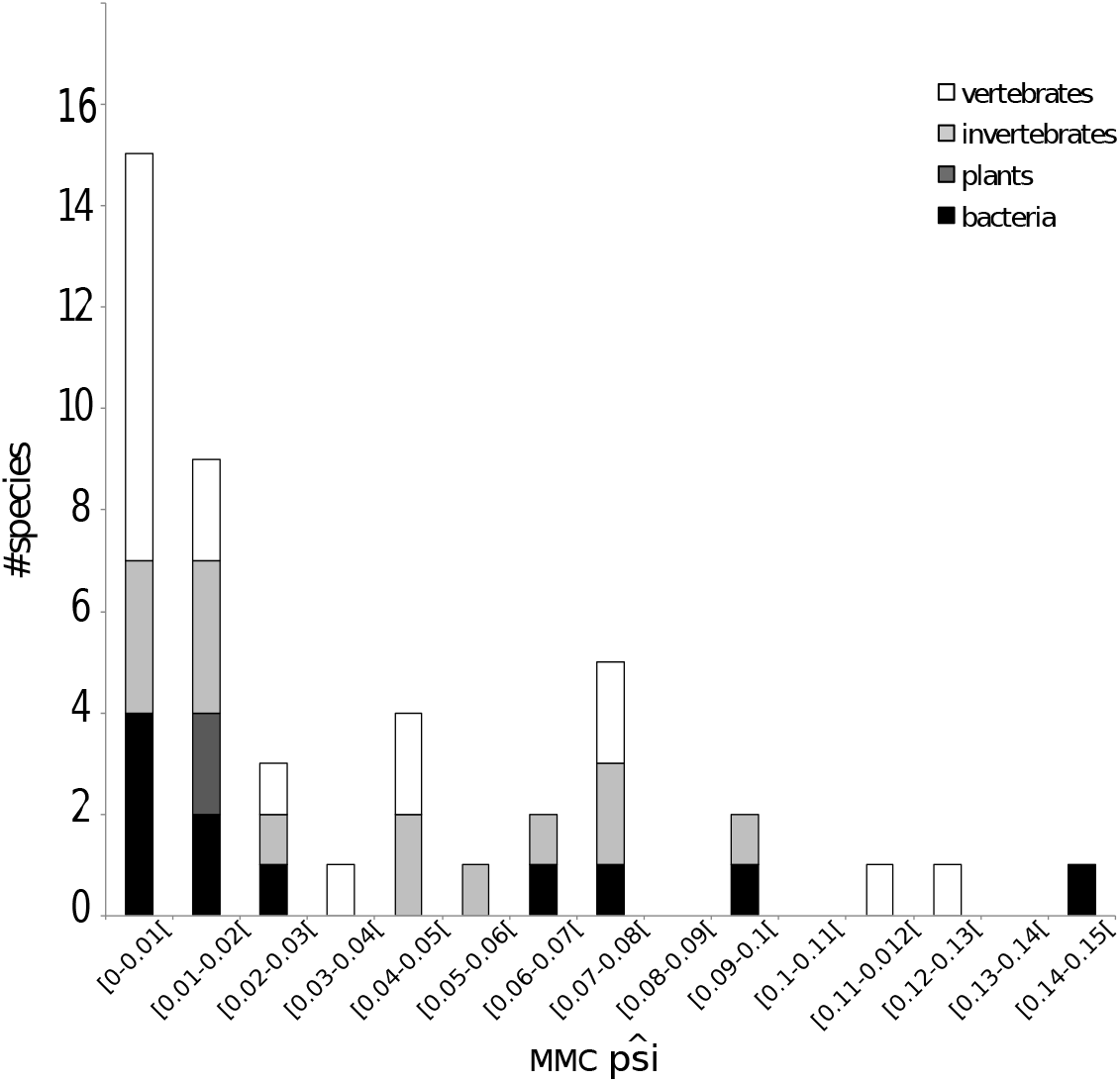
Distribution of psi parameter in function of the order of the species (white: vertebrates, light grey: invertebrates, dark grey: plants, black: bacteria).

**Figure A.8:**
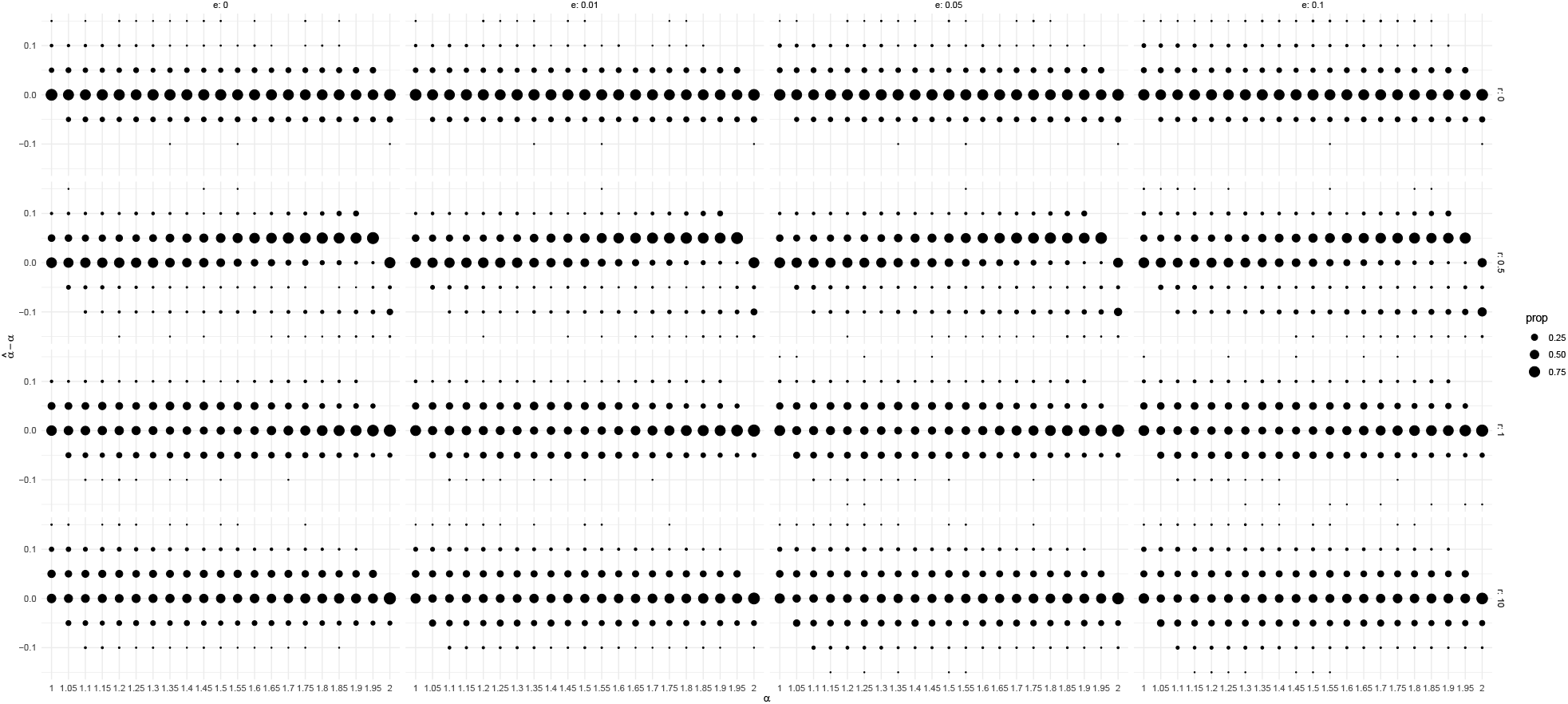
Error for estimating coalescent parameter *α* for Beta-coalescents with growth and misclassification (*n* = 100). Growth rate is denoted by *g*.

**Table A.4:**
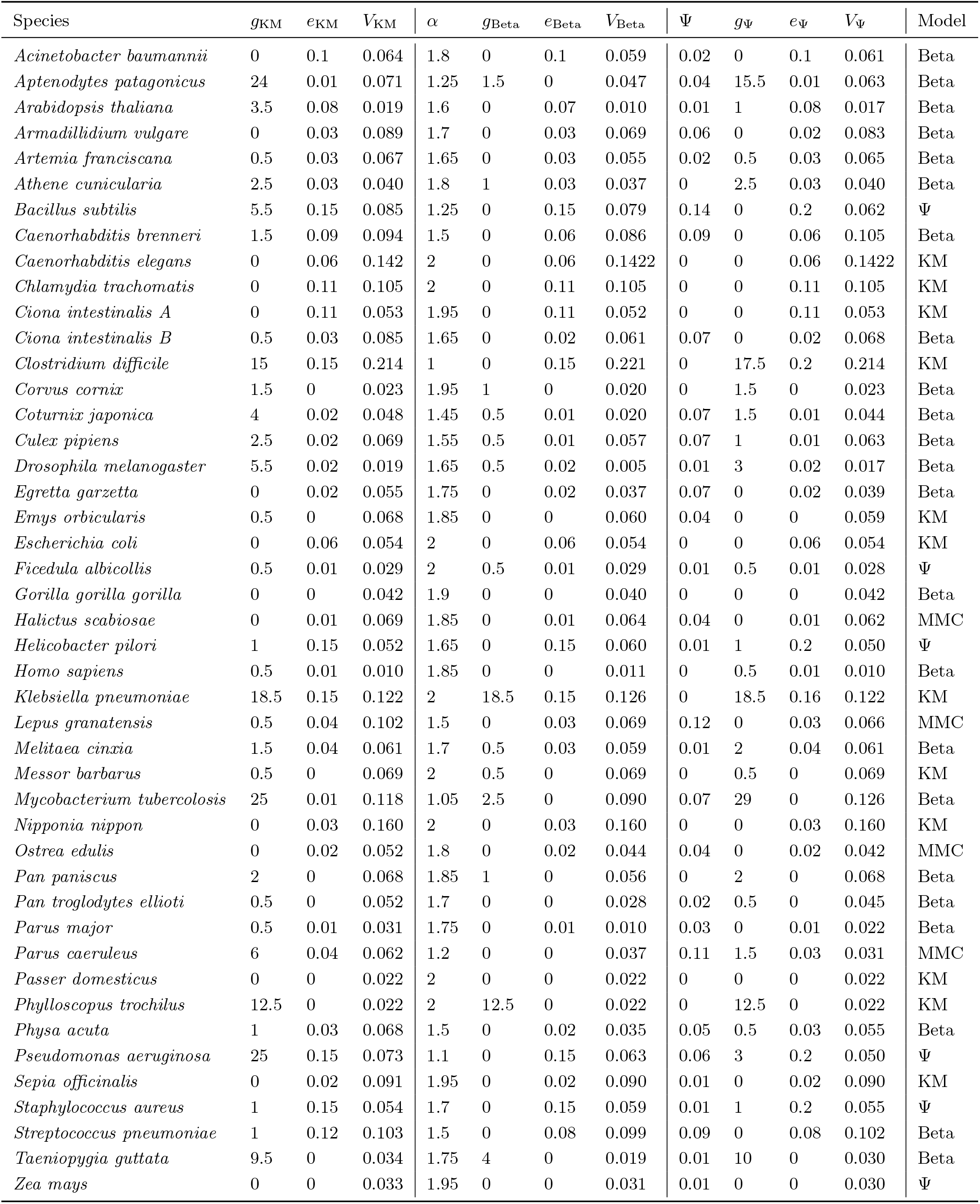
Parameters Estimations

**Figure A.9:**
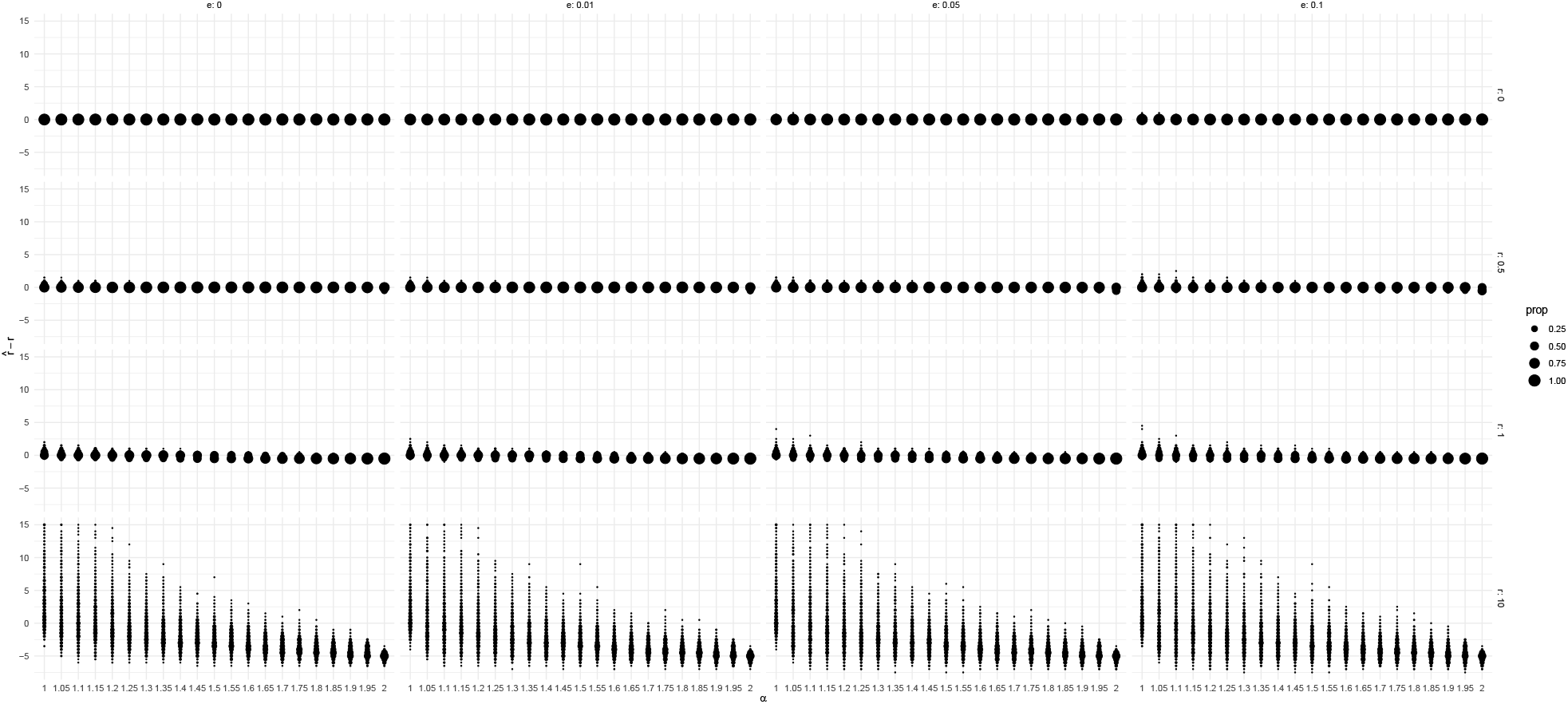
Error for estimating growth rate *g* for Beta-coalescents with growth and misclassification (*n* = 100)

**Figure A.10:**
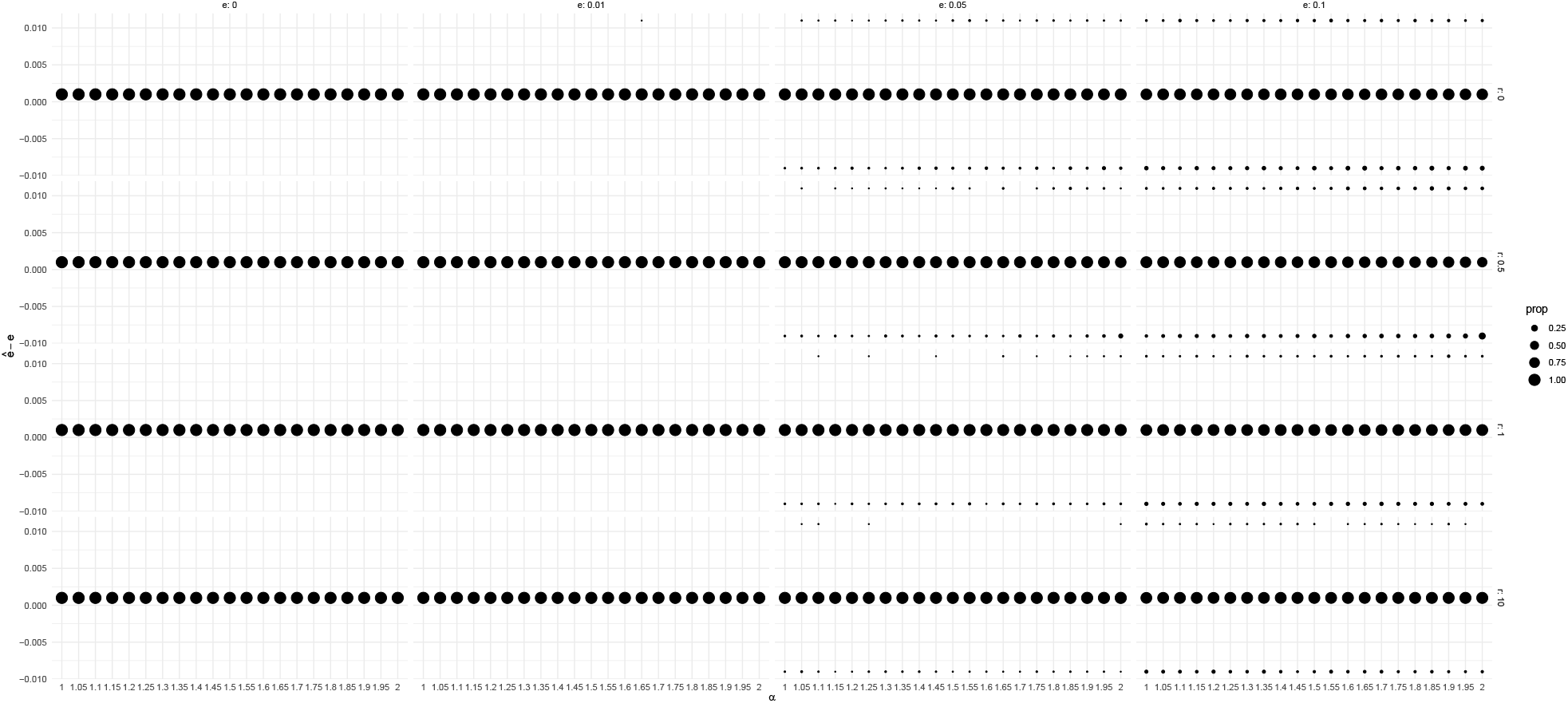
Error for estimating misorientation rate *e* for Beta-coalescents with growth and misclassification (*n* = 100). Growth rate is denoted by *g*.

**Figure A.11:**
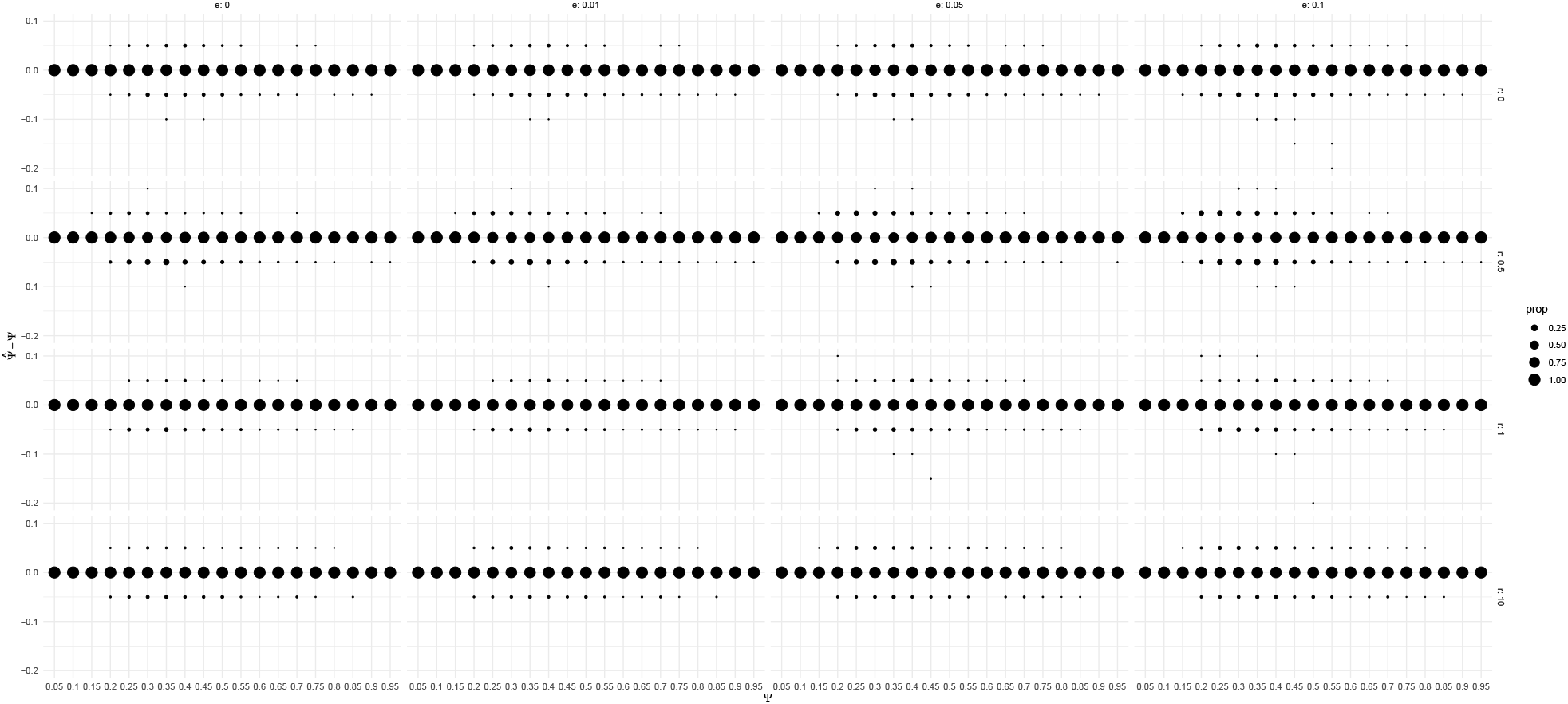
Error for estimating coalescent parameter Ψ for Psi-coalescents with growth and misclassification (*n* = 100). Growth rate is denoted by *g*.

**Figure A.12:**
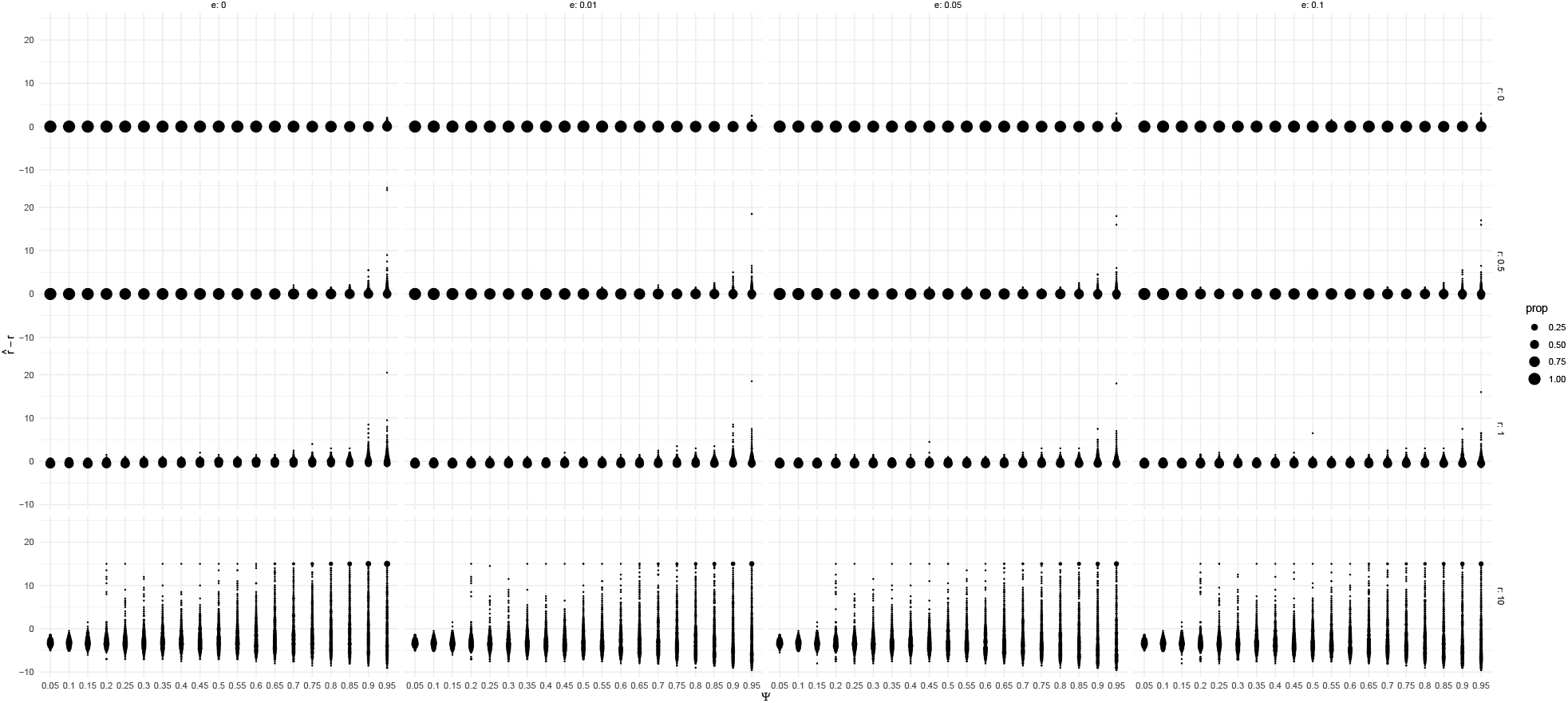
Error for estimating growth rate *g* for Psi-coalescents with growth and misclassification (*n* = 100).

**Figure A.13:**
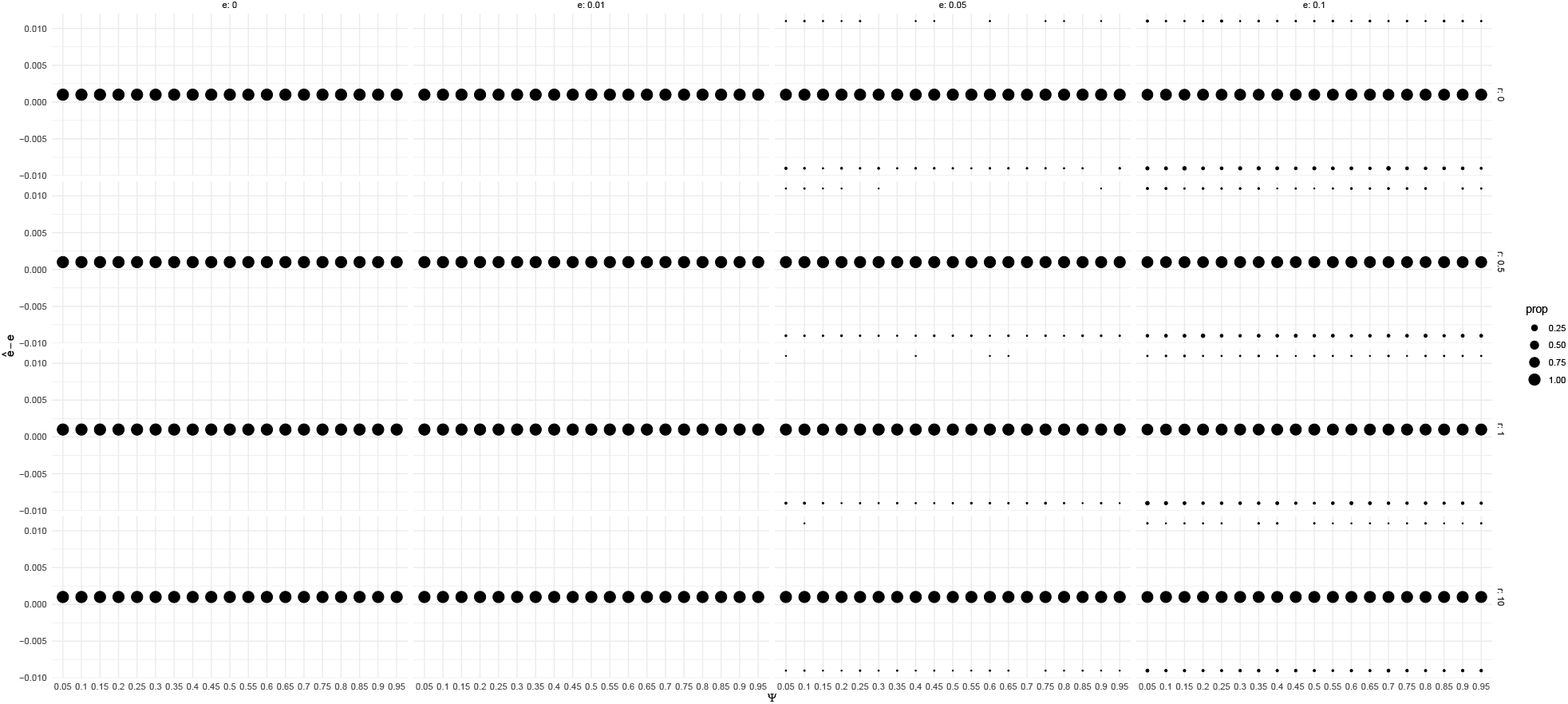
Error for estimating misorientation rate *e* for Psi-coalescents with growth and misclassification (*n* = 100). Growth rate is denoted by *g*.

**Figure A.14:**
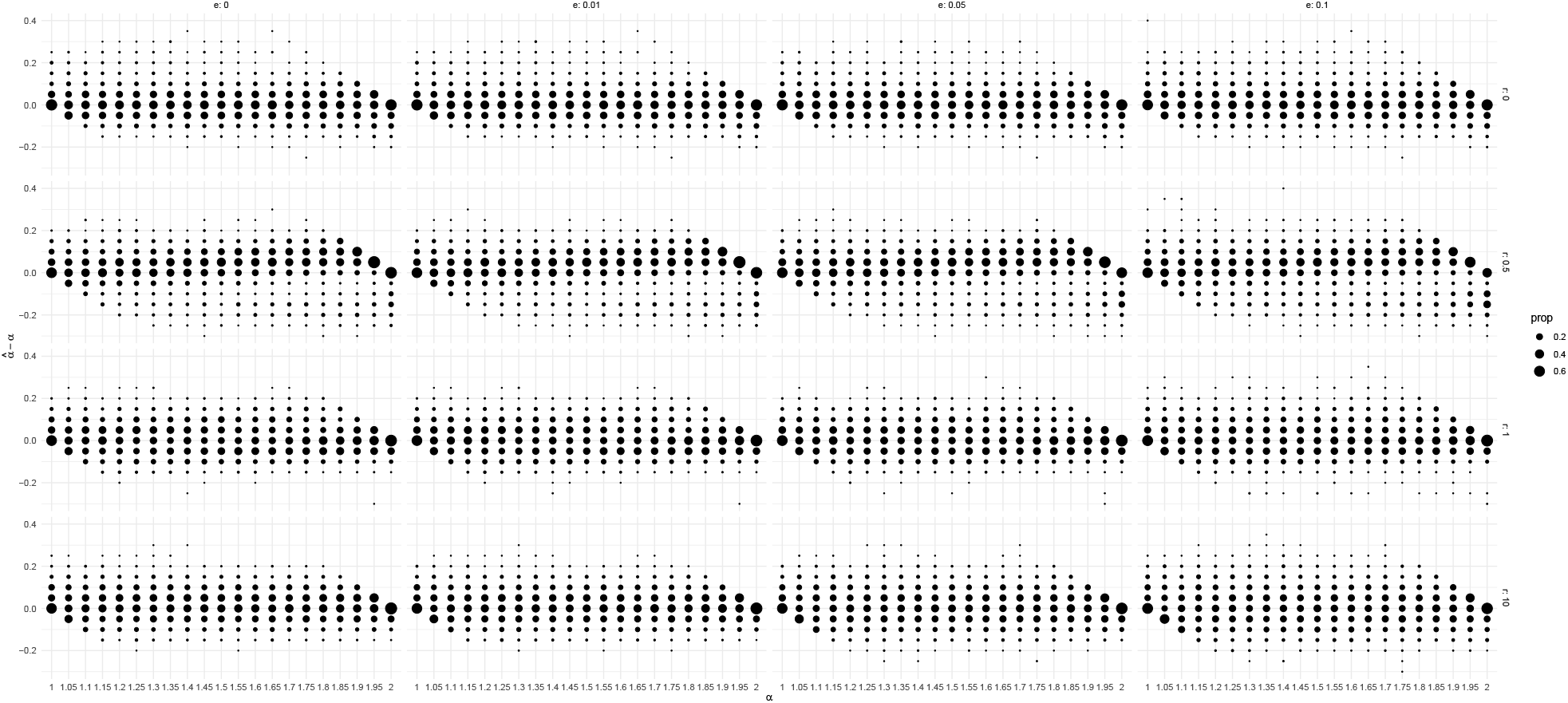
Error for estimating coalescent parameter *α* for Beta coalescents with growth and misclassification (*n* = 20). Growth rate is denoted by *g*.

**Figure A.15:**
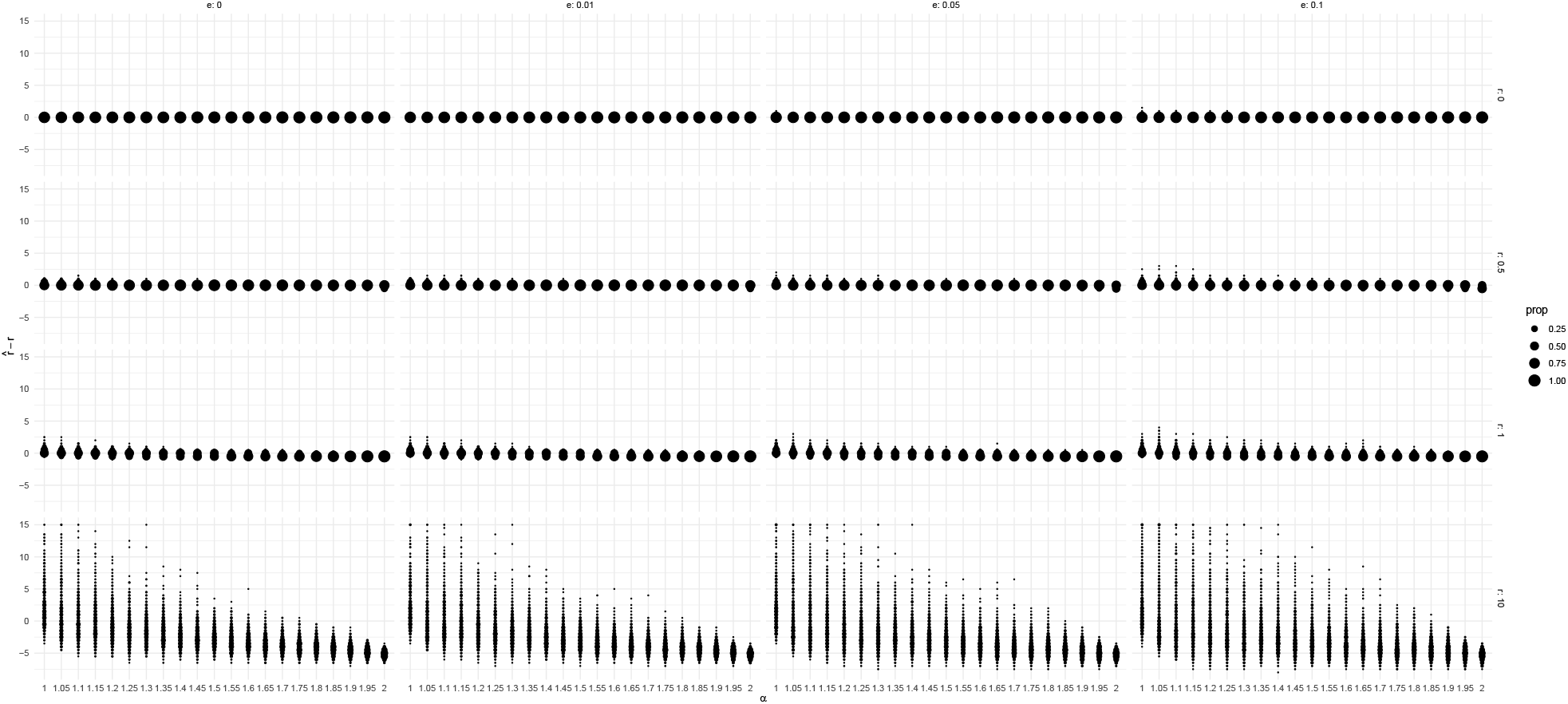
Error for estimating growth rate *g* for Beta-coalescents with growth and misclassification (*n* = 20)

**Figure A.16:**
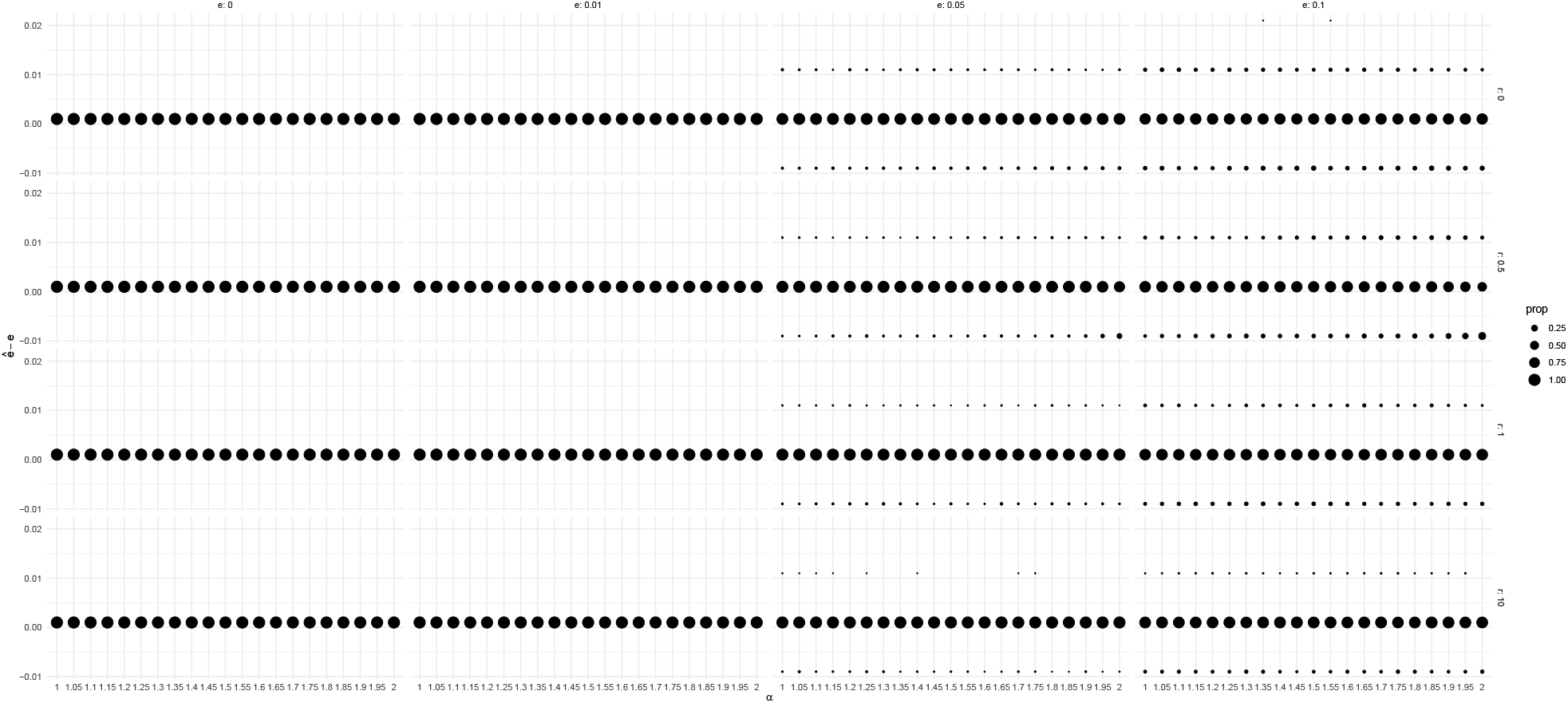
Error for estimating misorientation rate *e* for Beta-coalescents with growth and misclassification (*n* = 20). Growth rate is denoted by *g*.

**Figure A.17:**
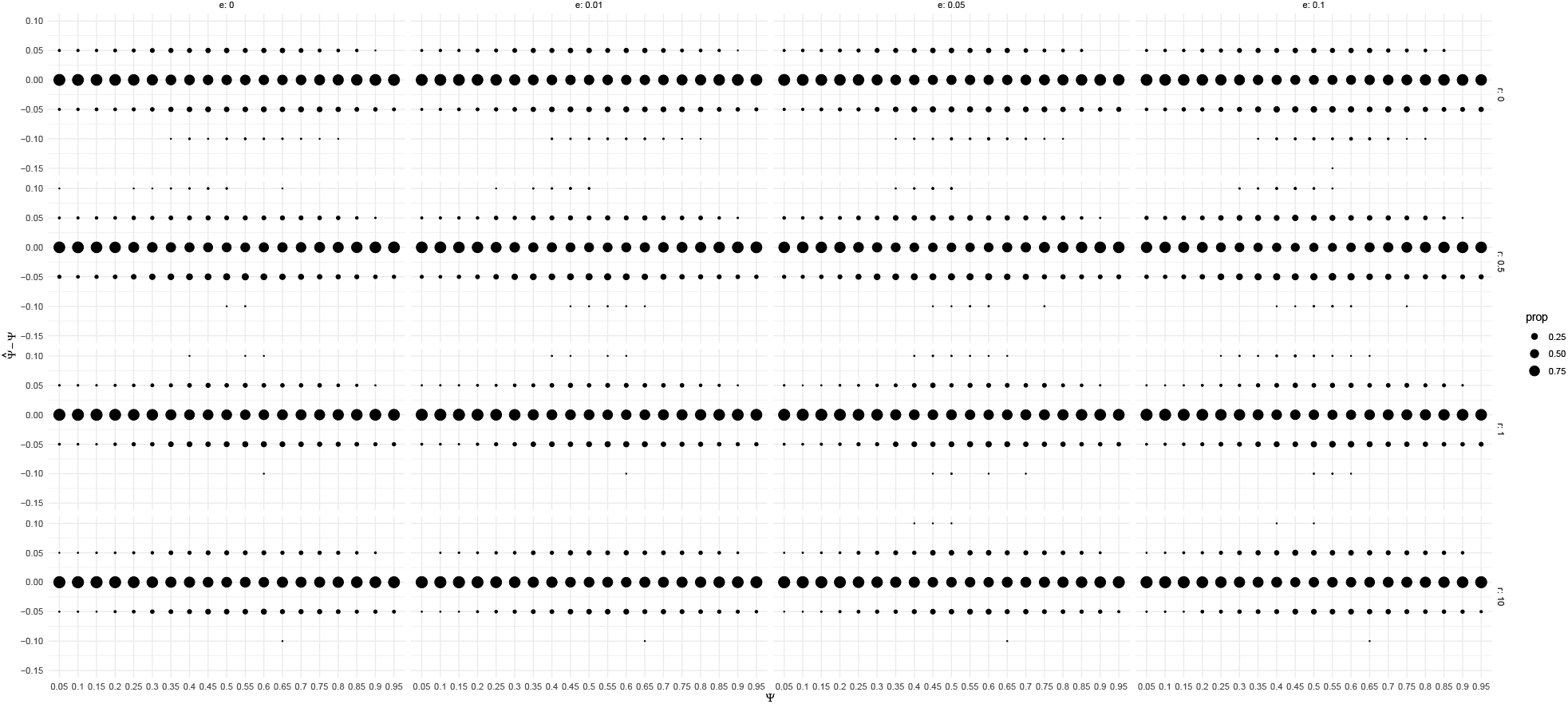
Error for estimating coalescent parameter Ψ for Psi-coalescents with growth and misclassification (*n* = 20). Growth rate is denoted by *g*.

**Figure A.18:**
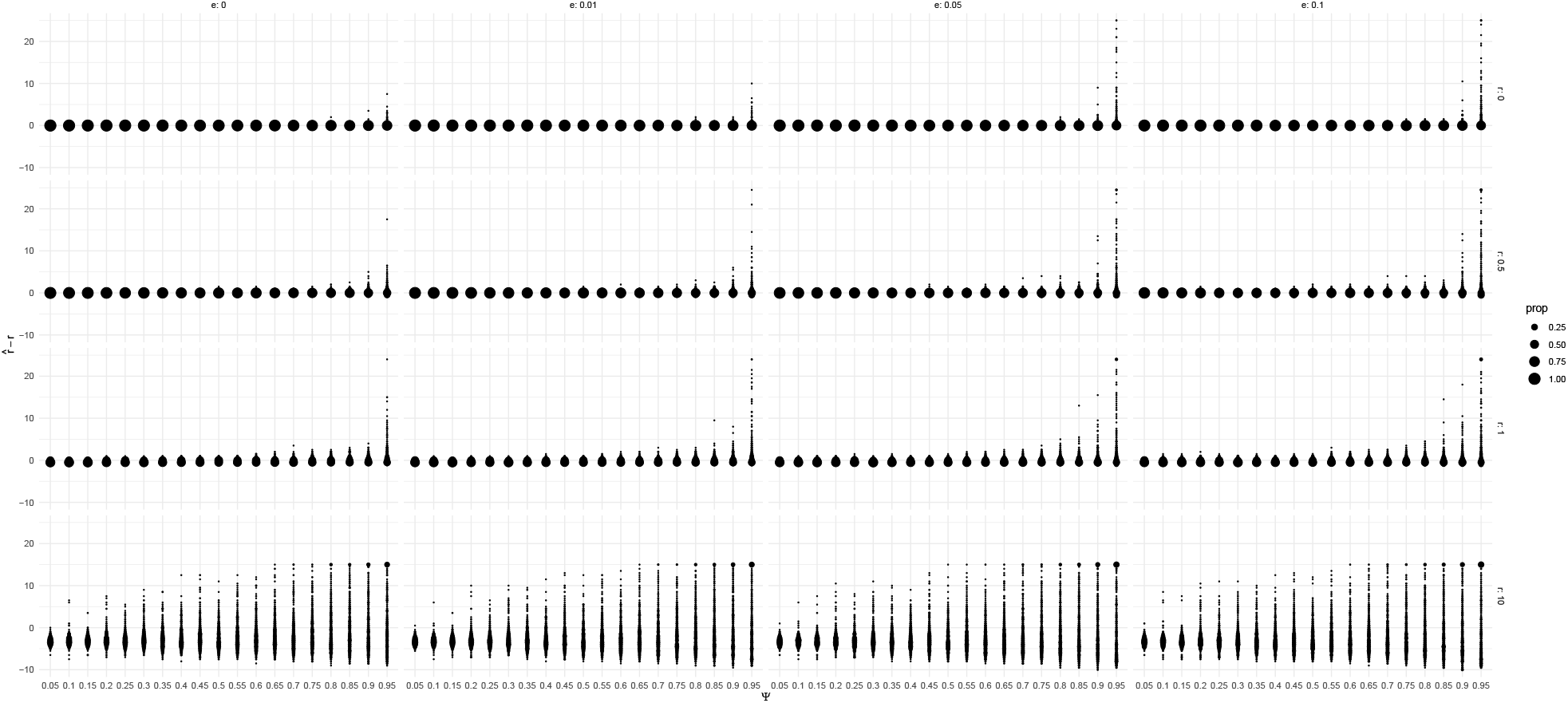
Error for estimating growth rate *g* for Psi-coalescents with growth and misclassification (*n* = 20).

**Figure A.19:**
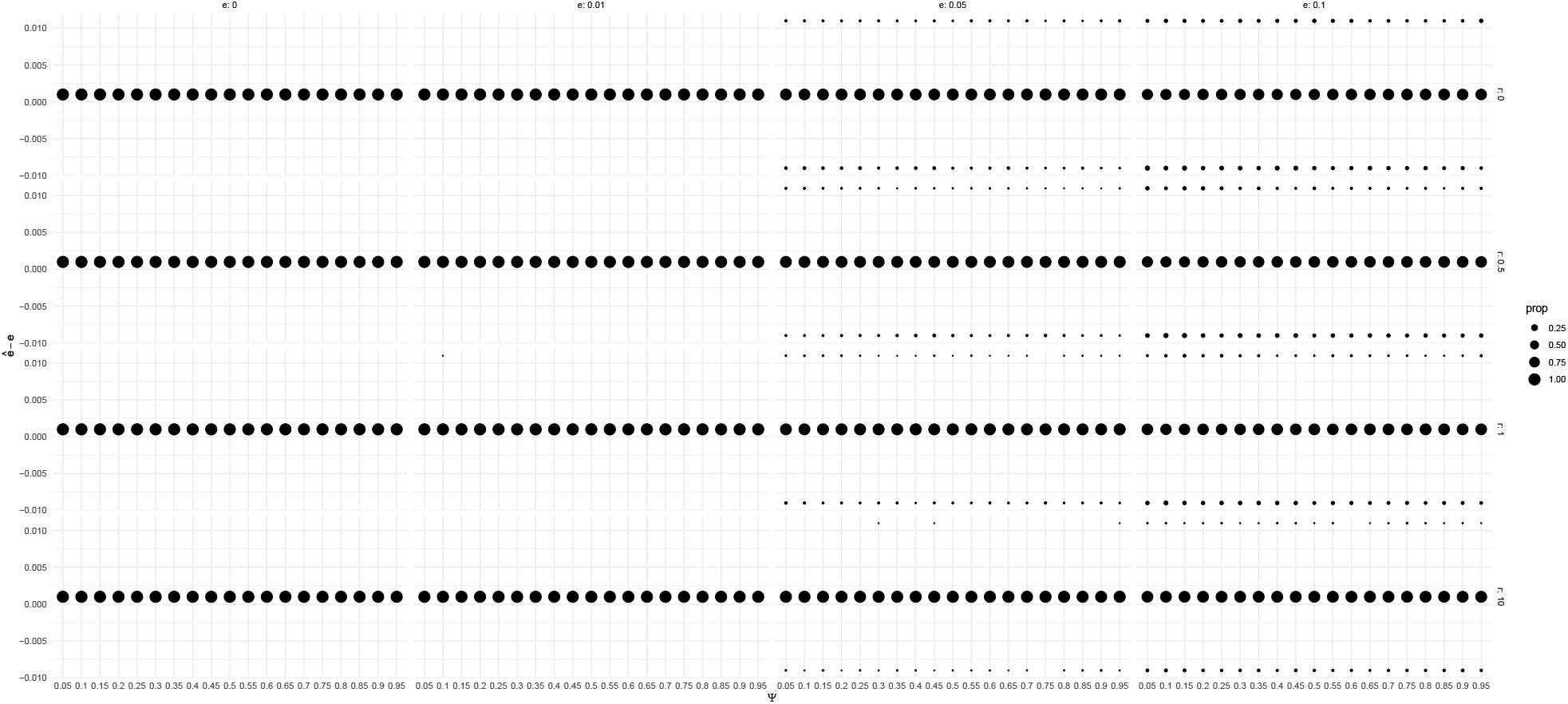
Error for estimating misorientation rate e for Psi-coalescents with growth and misclassification (*n* = 20). Growth rate is denoted by *g*.

**Figure A.20:**
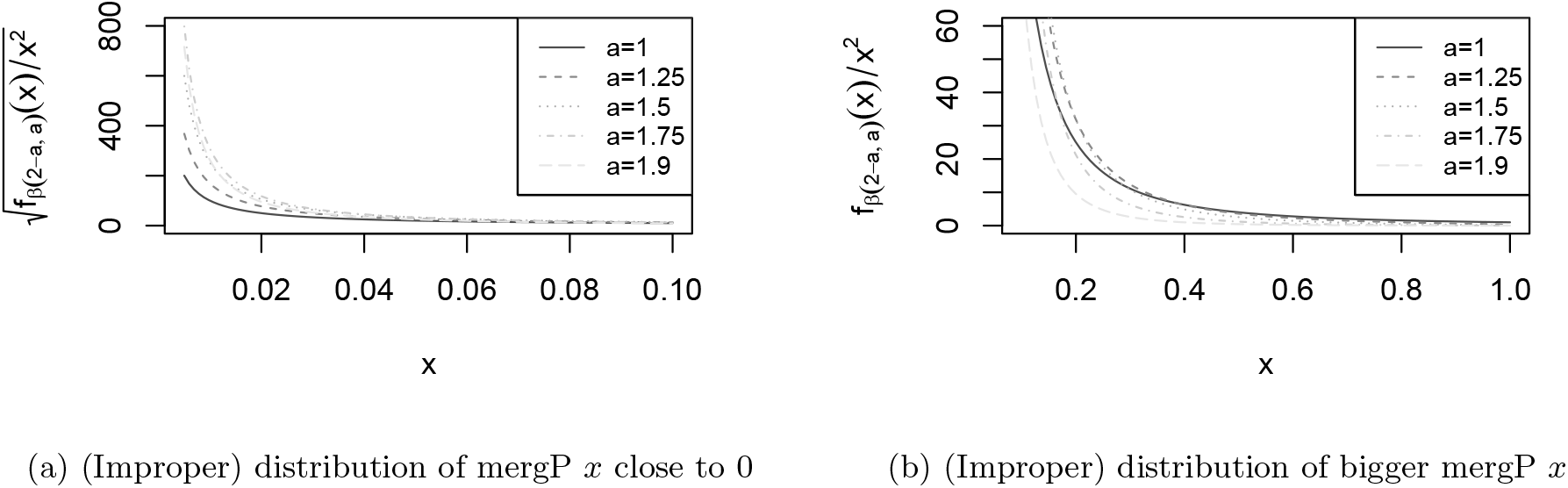
Distribution of merger rates of Beta-coalescents: Each lineage merges with merger probability *x* (abbeviated as mergP), where *x* is chosen with rate *x*^−2^ * Λ(*dx*), where Λ is a Beta distribution with parameters 2 – *a* and a. Mergers only are realized if at least two lineages merge. The figures depict the corresponding (improper) density *x*^−2^ * *f_β_*(2 – *a, a*), where *f_β_* is the density of the Beta distribution used. The detailed (Poisson) construction can be found in [Pit99].

**Table A.5:**
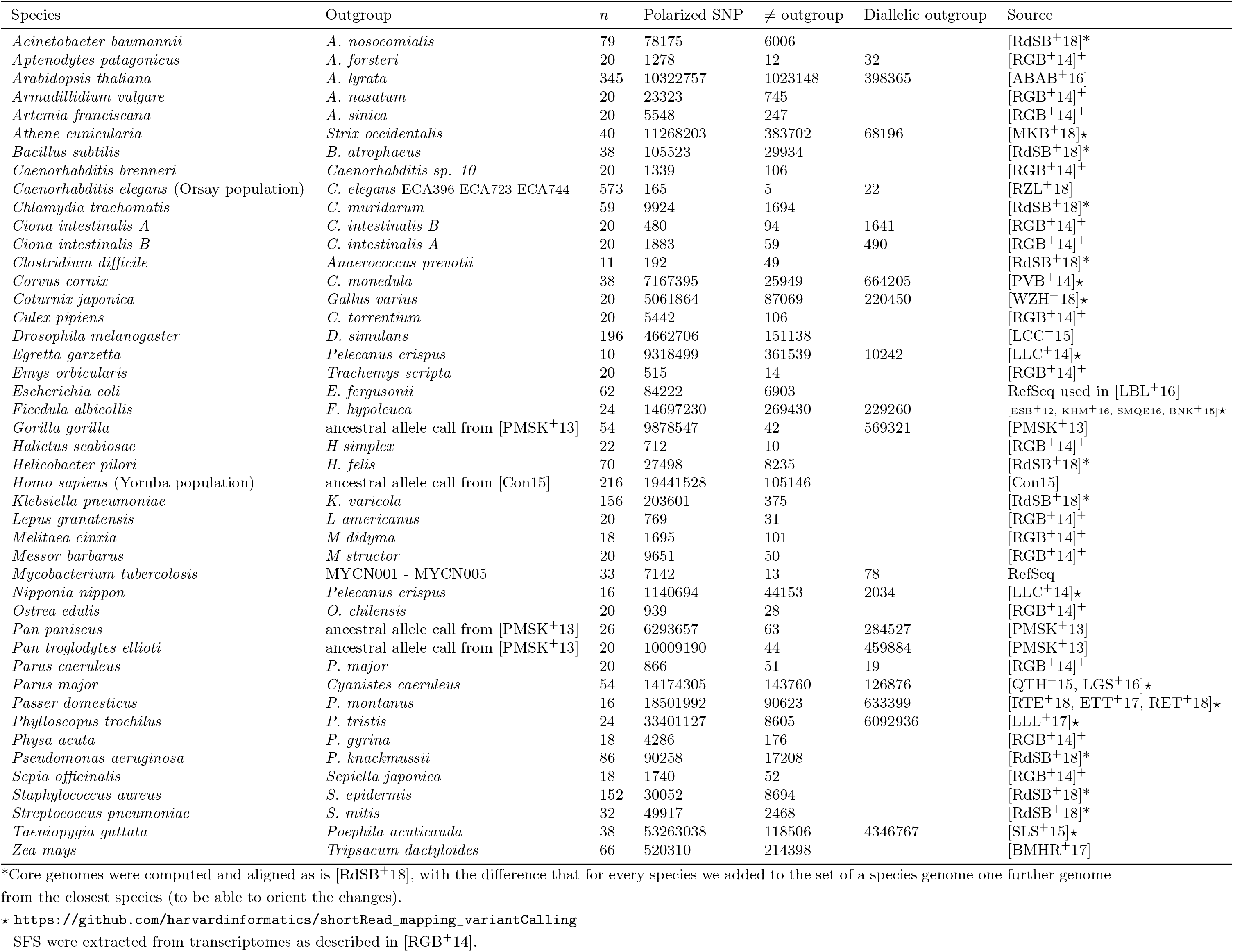
Data set information

**Figure A.21:**
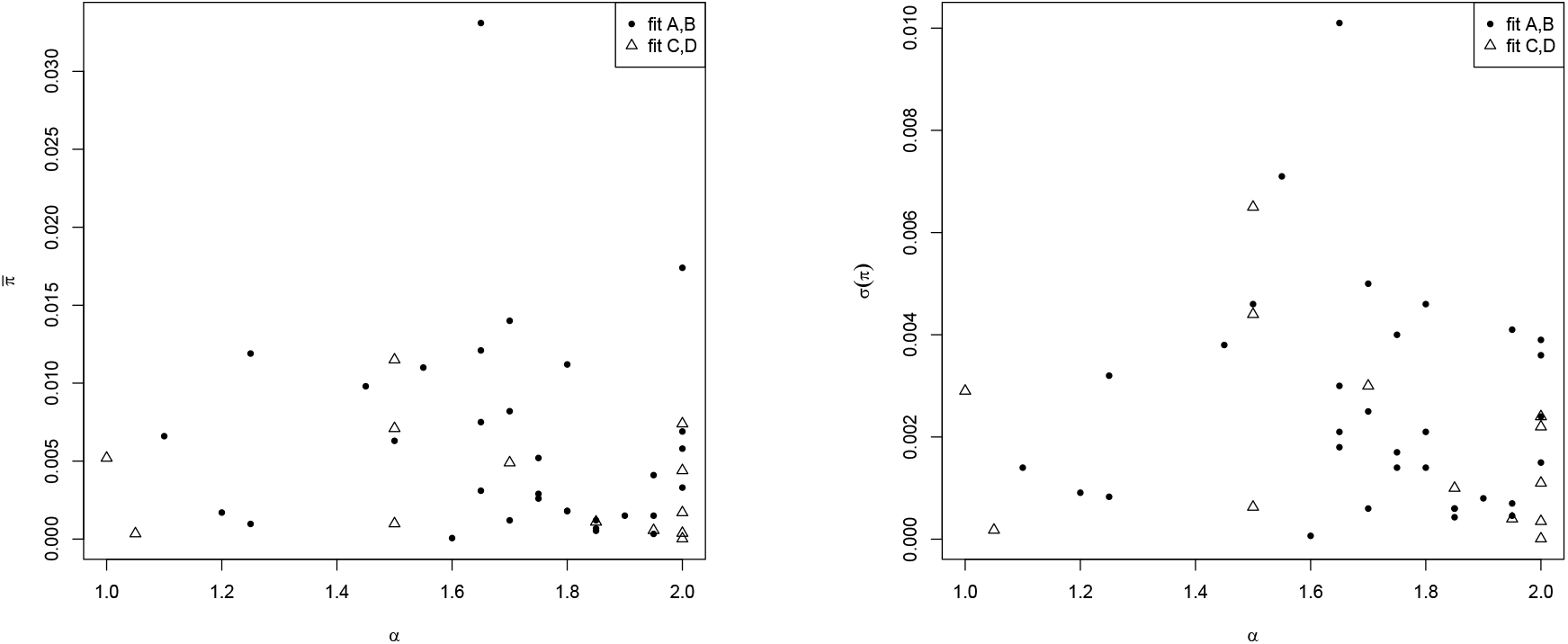
Comparison of estimated *α* parameter 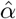 (*x*-axis) and mean 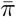 (standard deviation *σ*(*π*)) of windowed nucleotide diversity *π* (*y*-axis). See Sect. A.10) for details.

**Table A.6:**
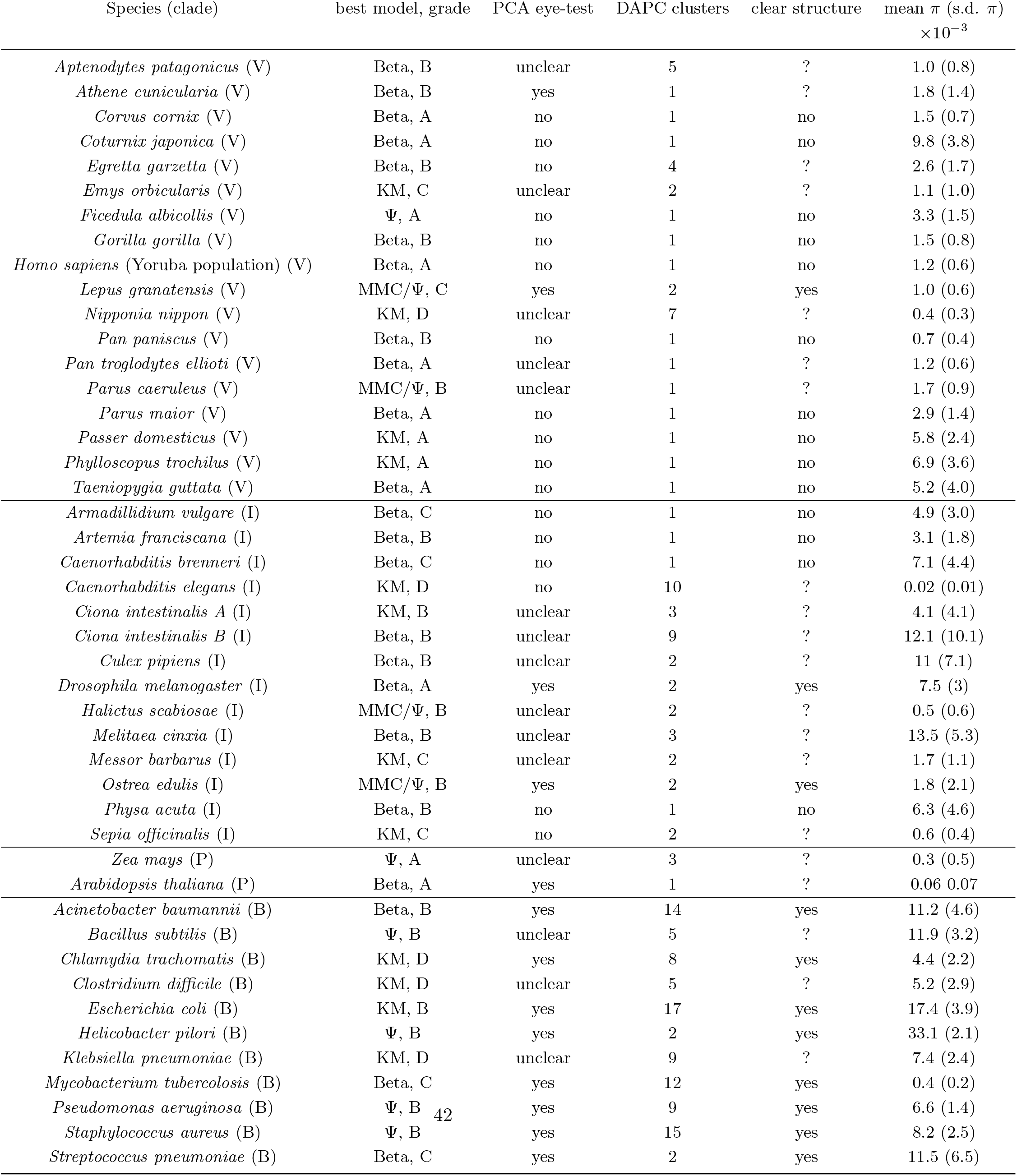
Population structure inference and genetic diversity *π* (nucleotide diversity per site in 15 kb windows) for the fitted data sets. Model fitted, grade of fit, and biological order repeated from Table 2. Number of clusters: *k* inferred by BIC criterion for subsequent *k*-means clustering. PCA eye-test: Visual inspectation of PCA plot. Clear structure: “yes” if PCA eye-test=yes and DAPC cluster=1, “no” if PCA eye-test=no and DAPC cluster>1, otherwise “?”.

**Table A.7:** Correlation coefficient *ρ* of estimated *α* parameter 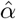 and mean 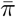 (standard deviation *σ*(*π*)) of windowed nucleotide diversity *π* (Sect. A.10)

| $x$ | $\rho(\hat{\alpha}, x)$ all data sets | $\rho(\alpha, x)$ data sets with fits A,B |
| --- | --- | --- |
| $\bar{\pi}$ | -.11 | -.11 |
| $\sigma(\pi)$ | -.12 | -.06 |

