## Supplementary figures and images for "Interpreting the pervasive observation of U-shaped Site Frequency Spectra"

### Supplementary 3 - per chromosome Droso

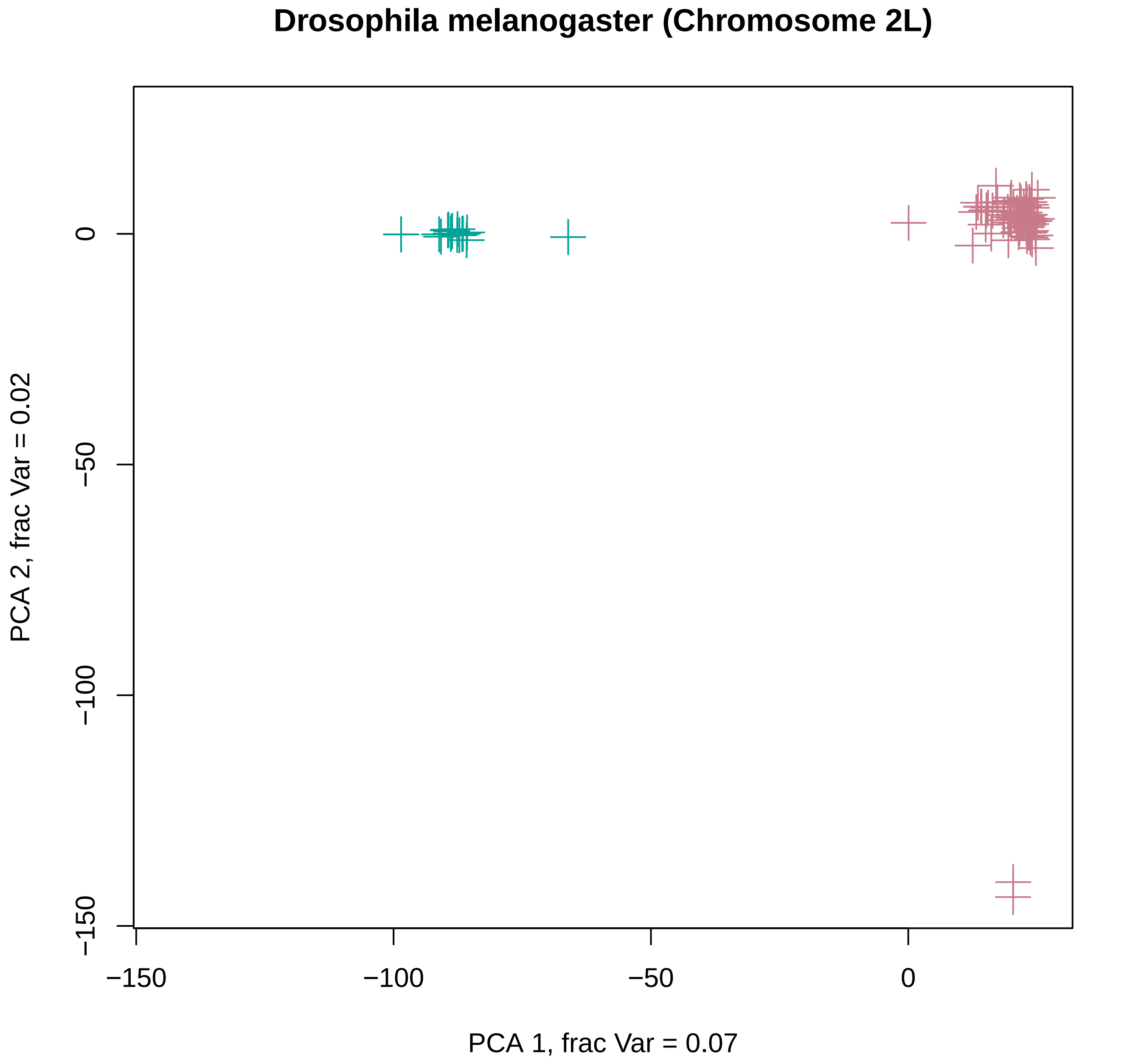

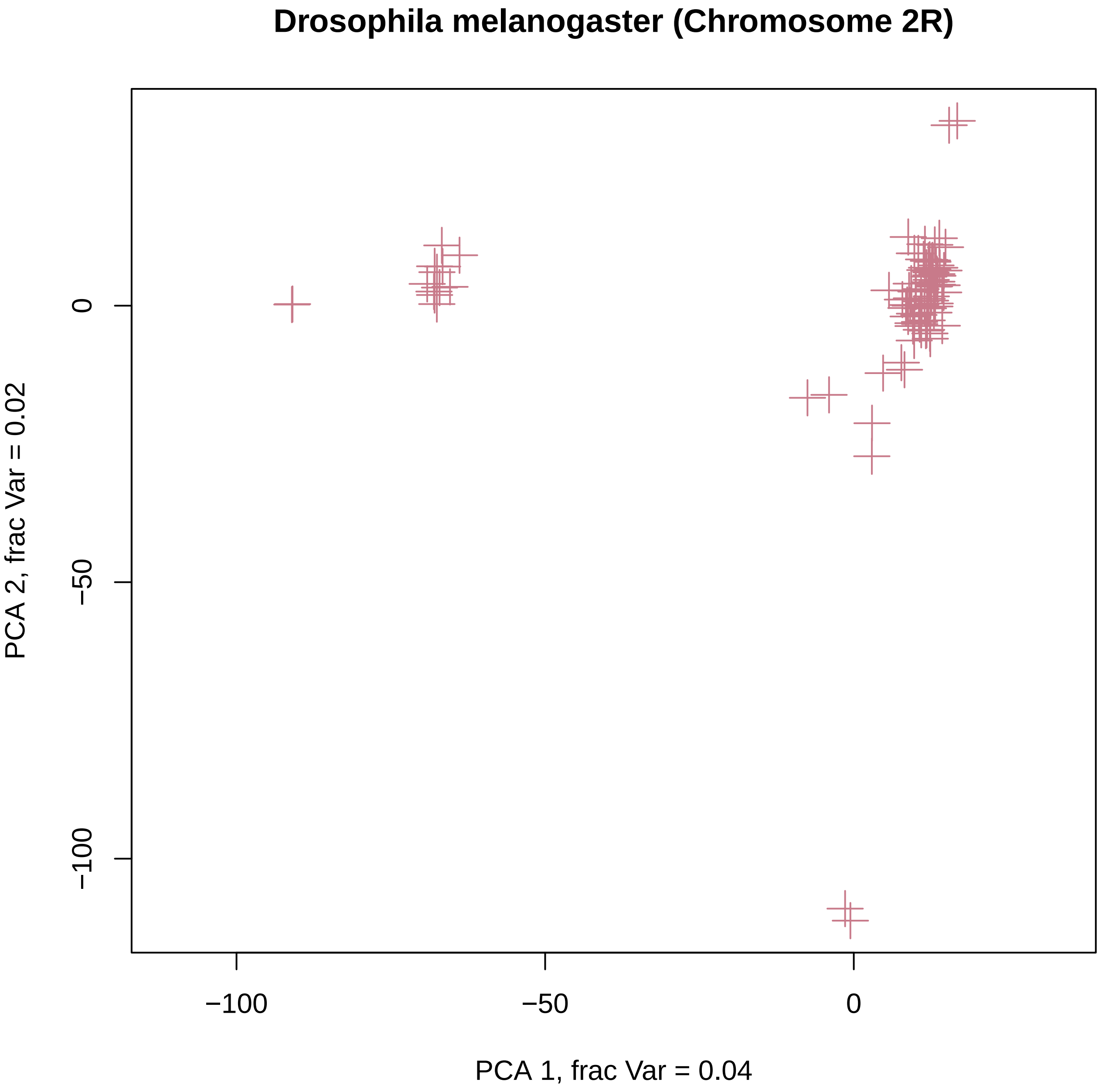

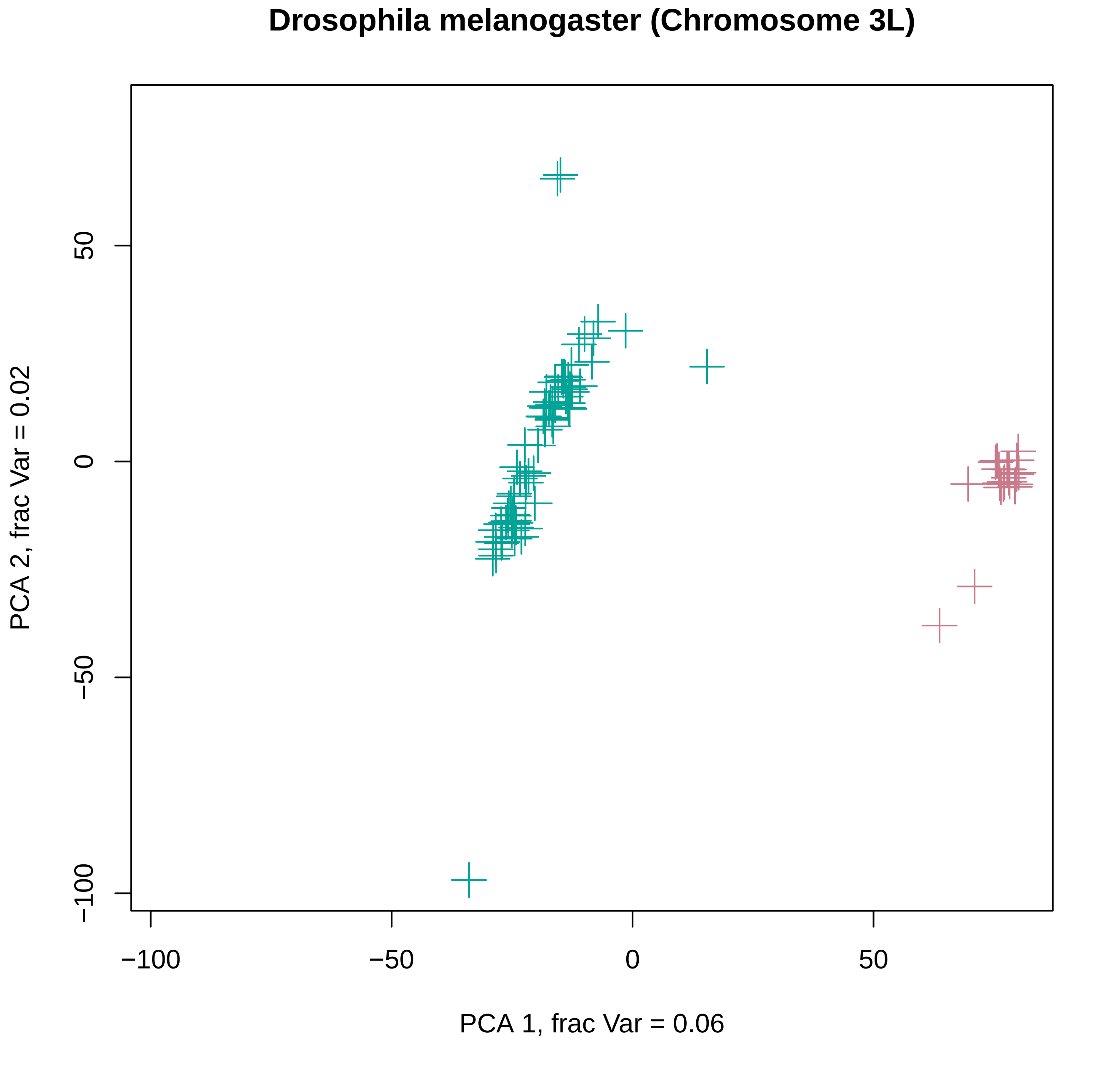

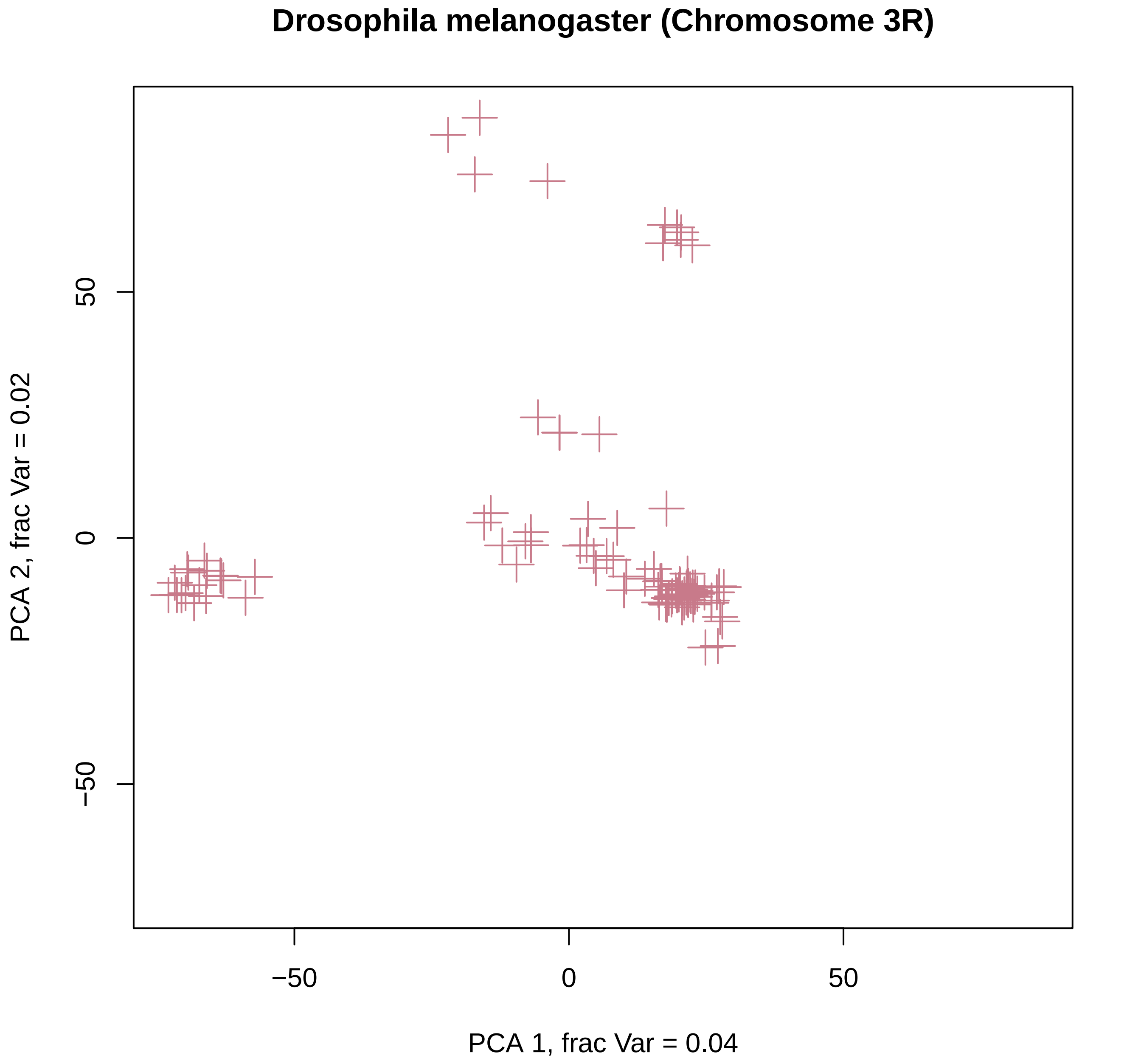

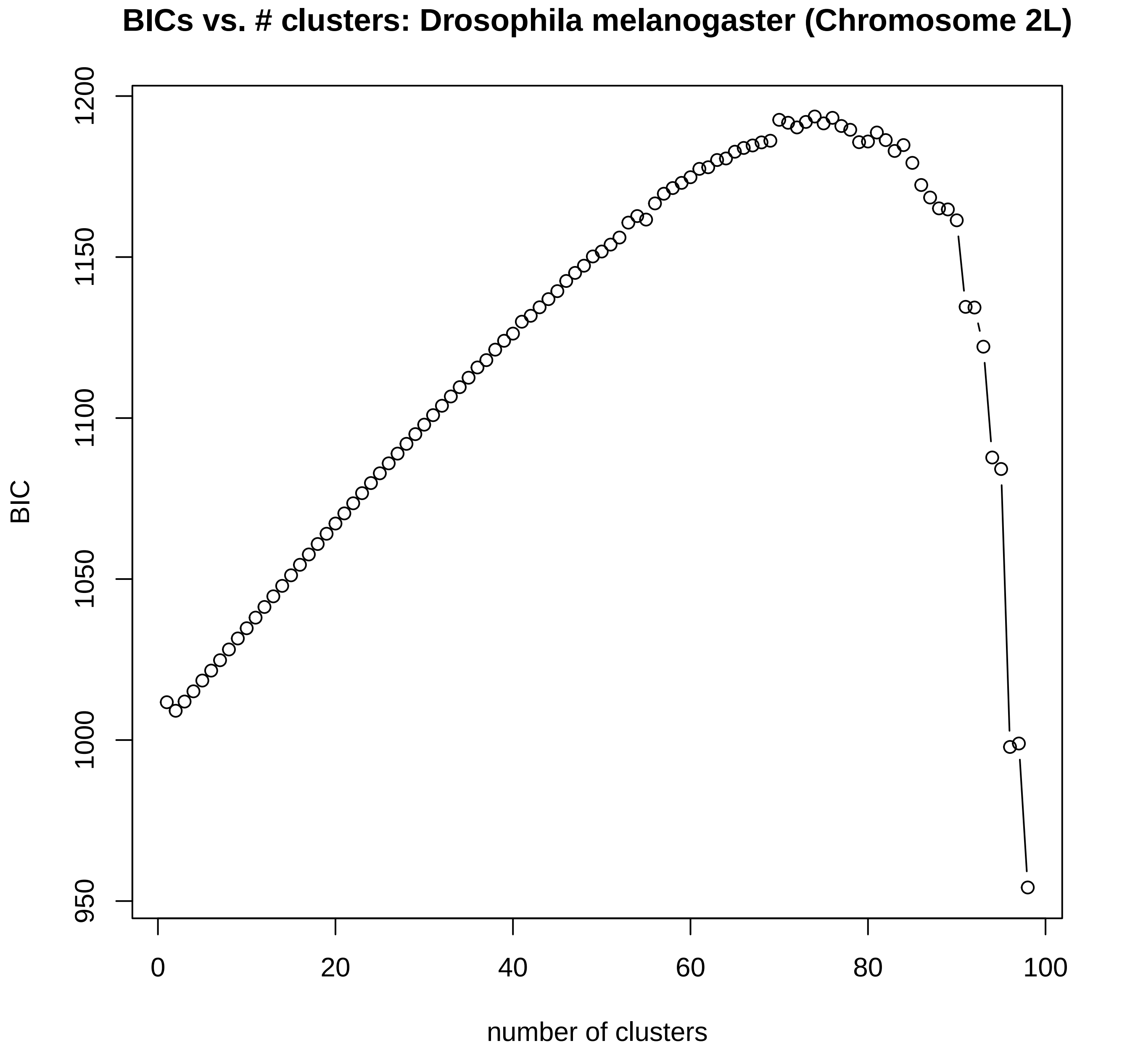

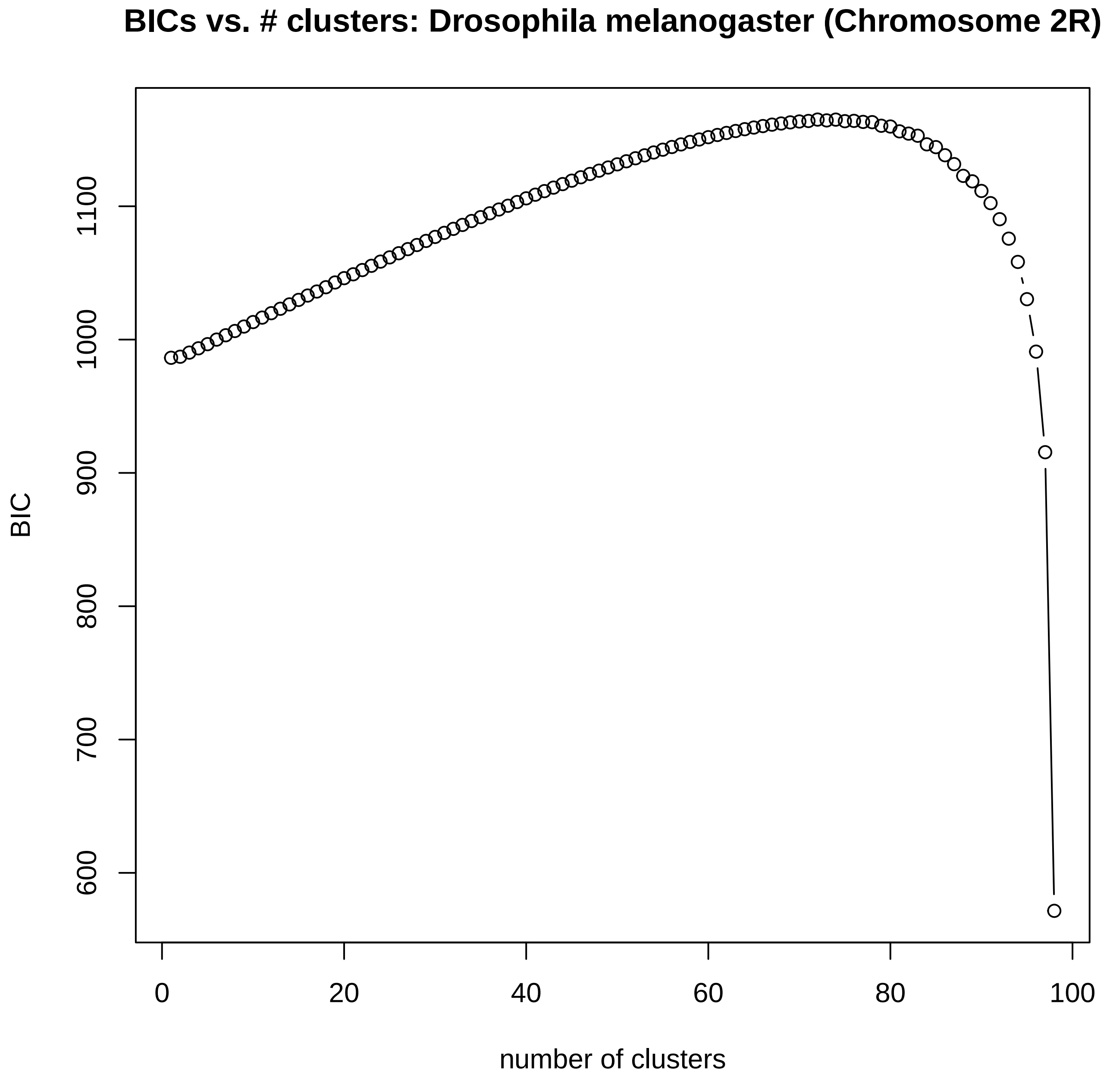

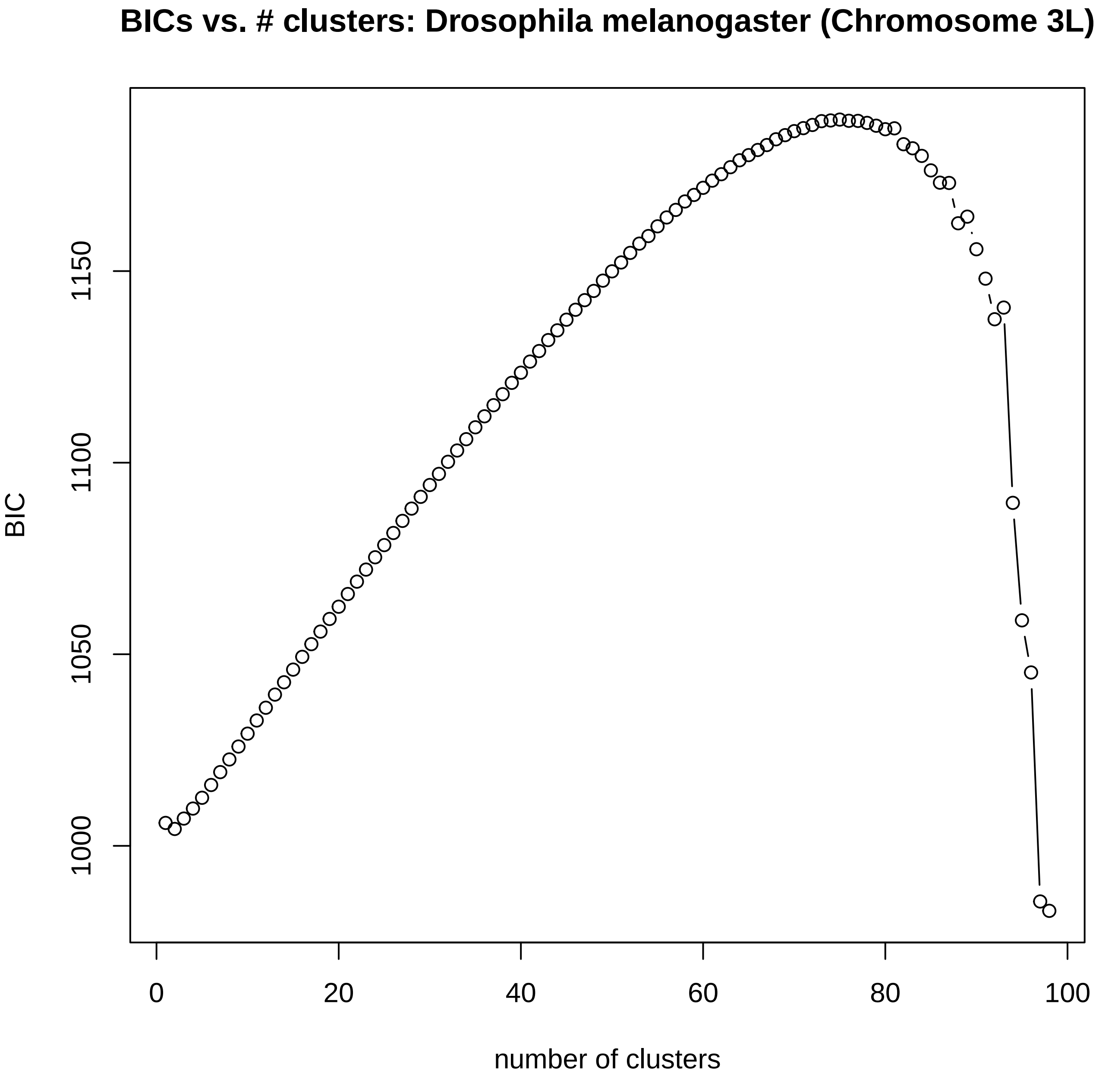

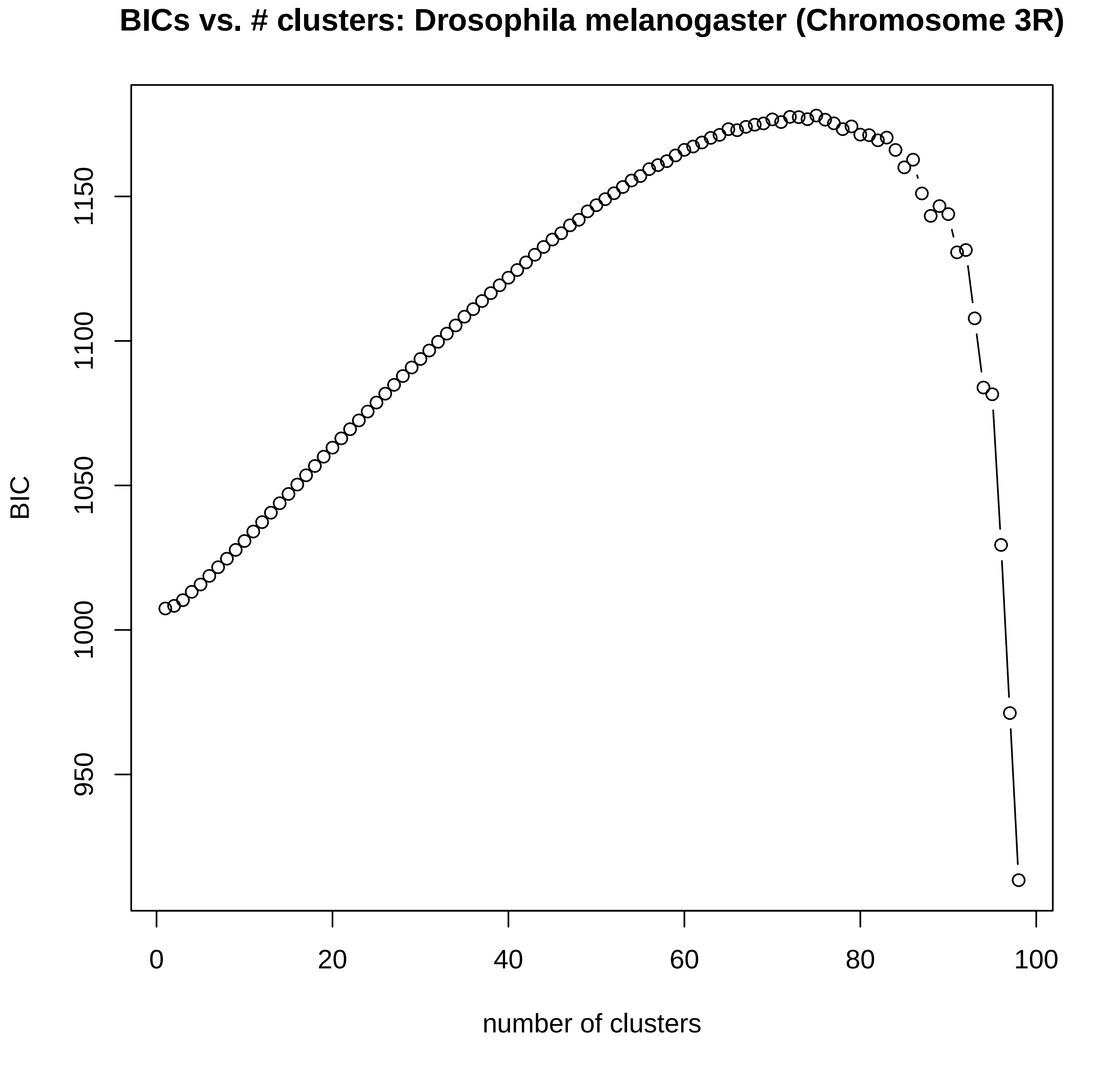
