## Supplementary 3 - BIC plots for DAPC for "Interpreting the pervasive observation of U-shaped Site Frequency Spectra"

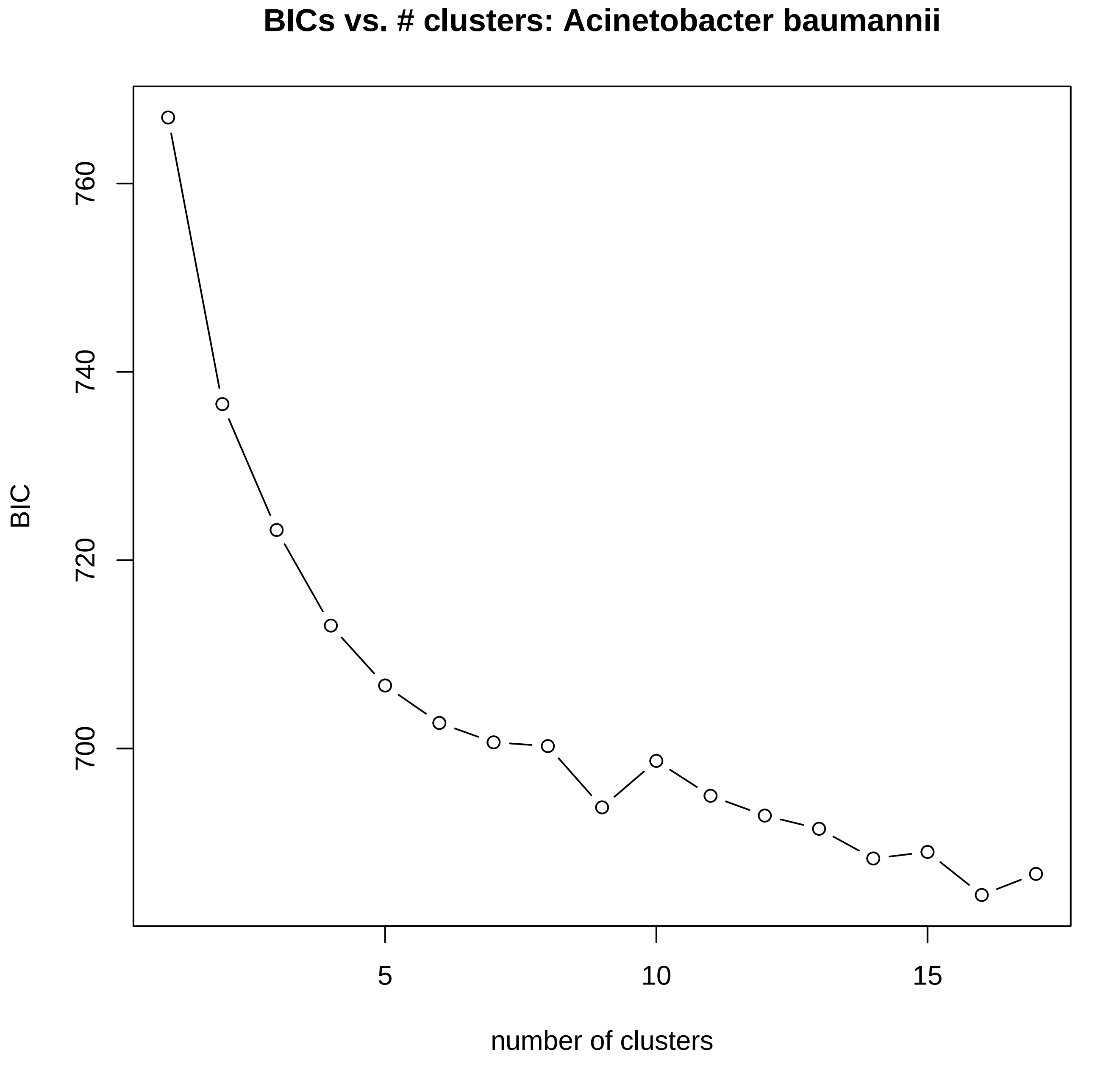
BICs vs. # clusters: Acinetobacter baumannii
BIC
760
740
720
700
5
10
15
number of clusters

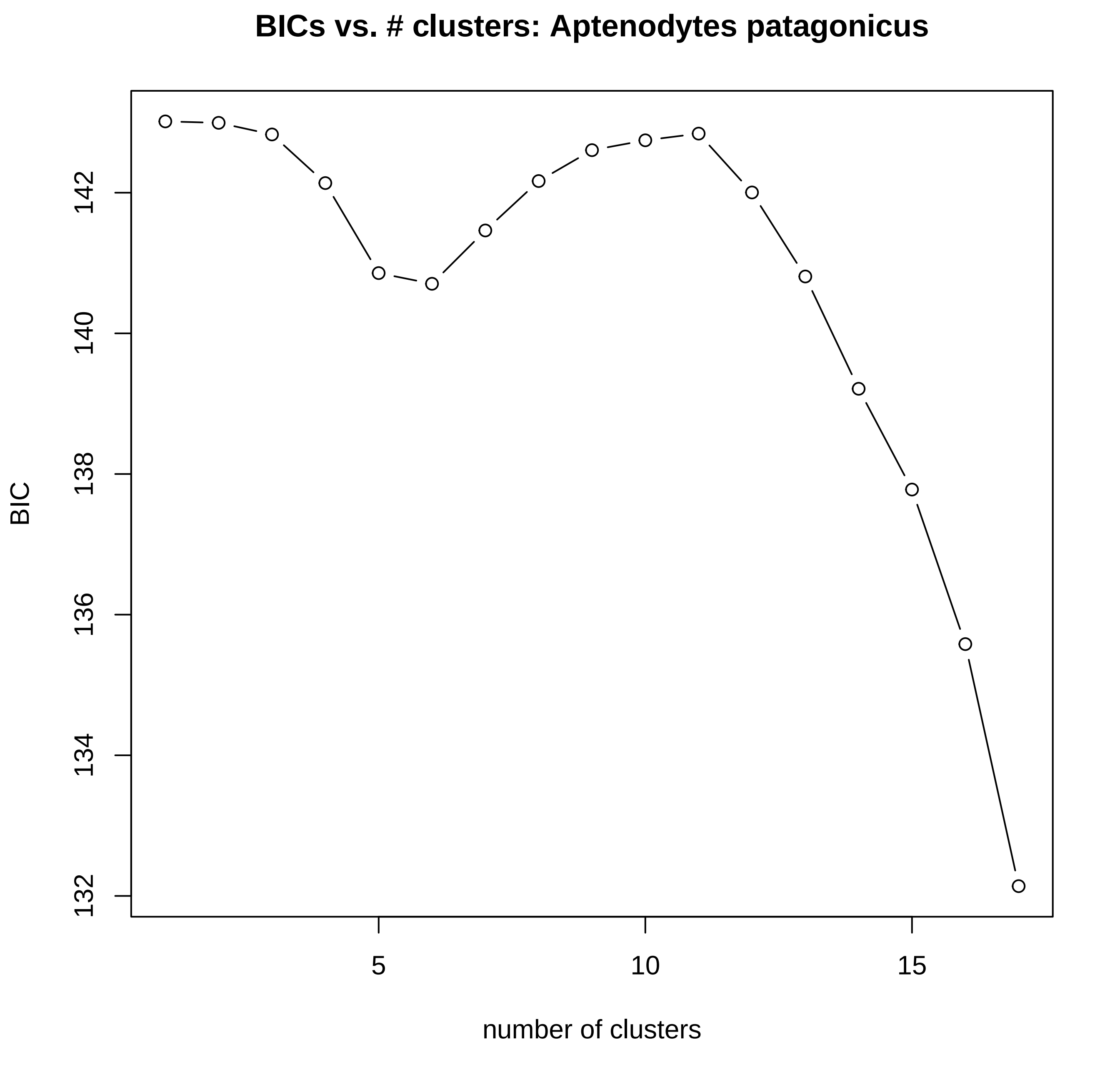

BICs vs. # clusters: Aptenodytes patagonicus
BIC
132
134
136
138
140
142
5
10
15
number of clusters

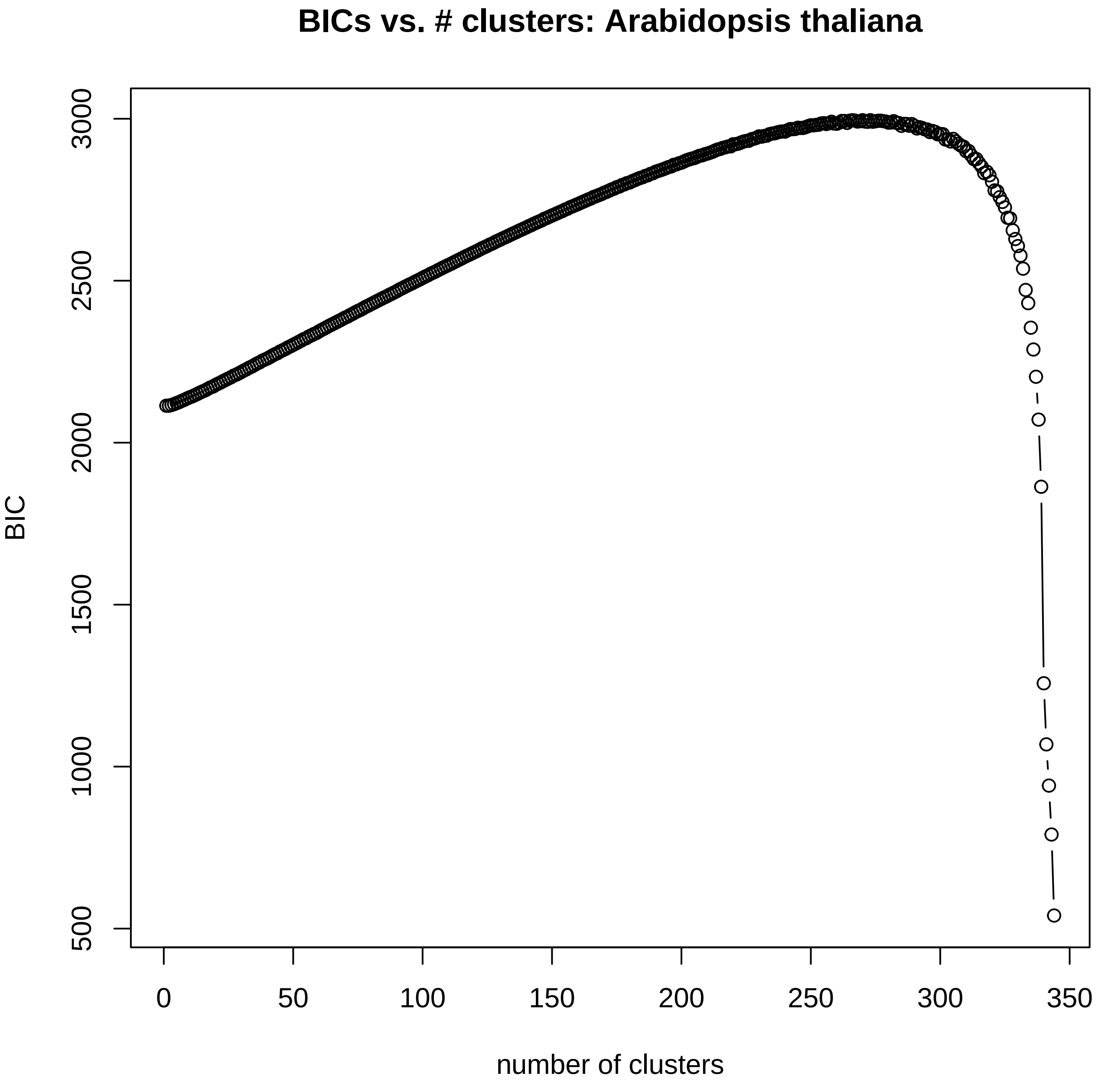
BICs vs. # clusters: Arabidopsis thaliana
BIC
number of clusters
500
1000
1500
2000
2500
3000
0
50
100
150
200
250
300
350

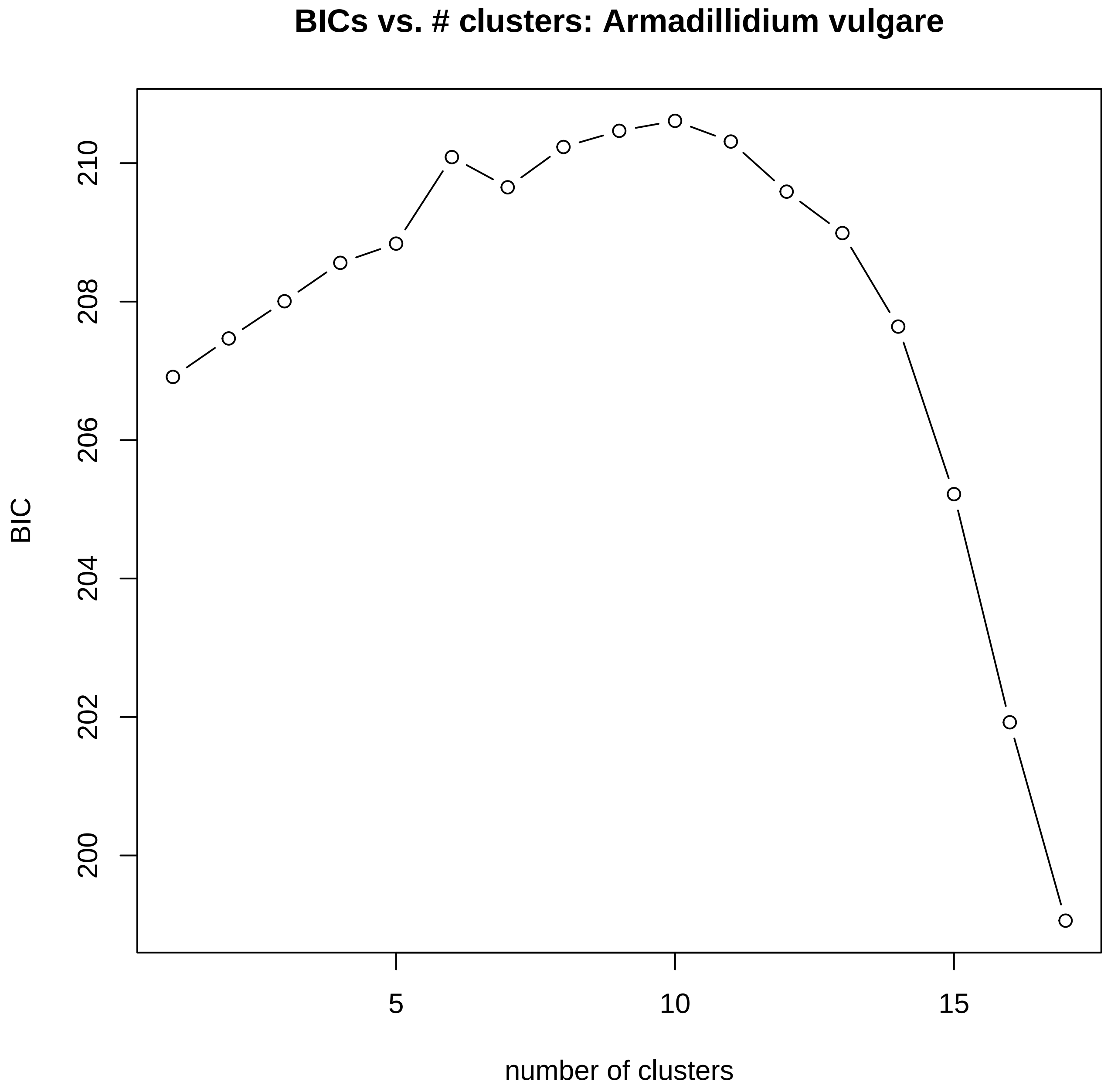
BICs vs. # clusters: Armadillidium vulgare
BIC
200
202
204
206
208
210
5
10
15
number of clusters

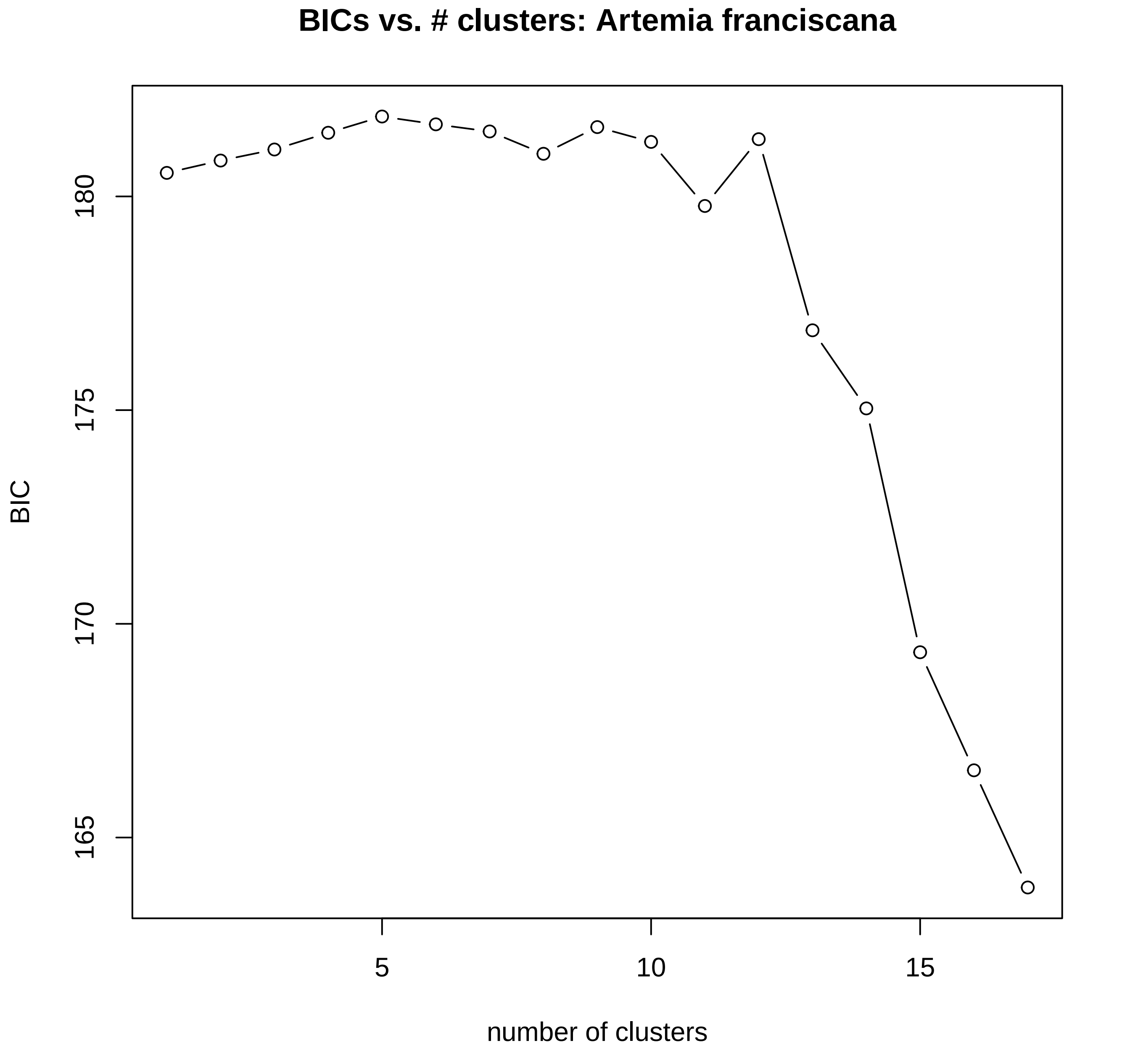
BICs vs. # clusters: Artemia franciscana
BIC
180
175
170
165
5
10
15
number of clusters

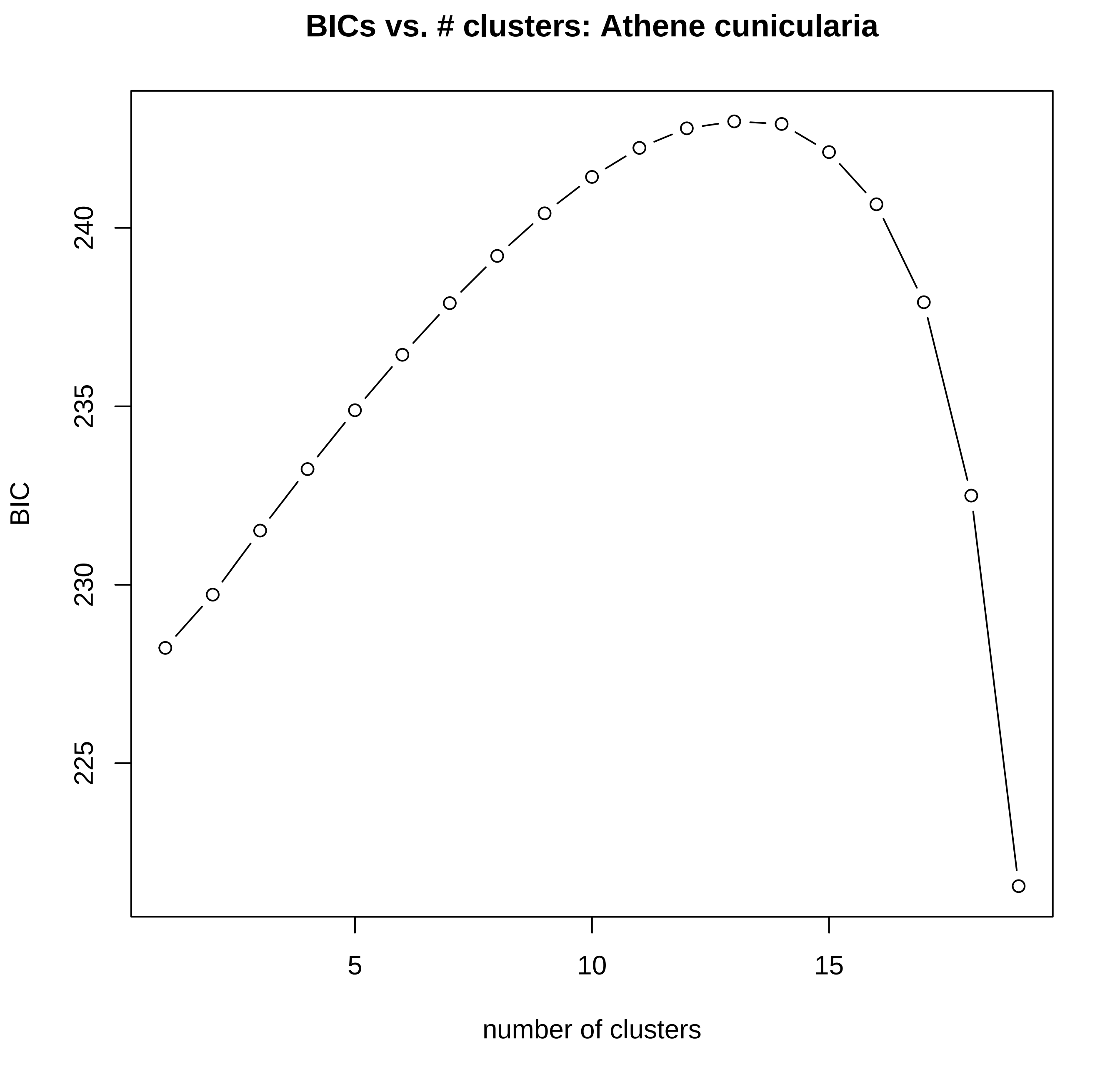

BICs vs. # clusters: Athene cunicularia
BIC
225
230
235
240
5
10
15
number of clusters

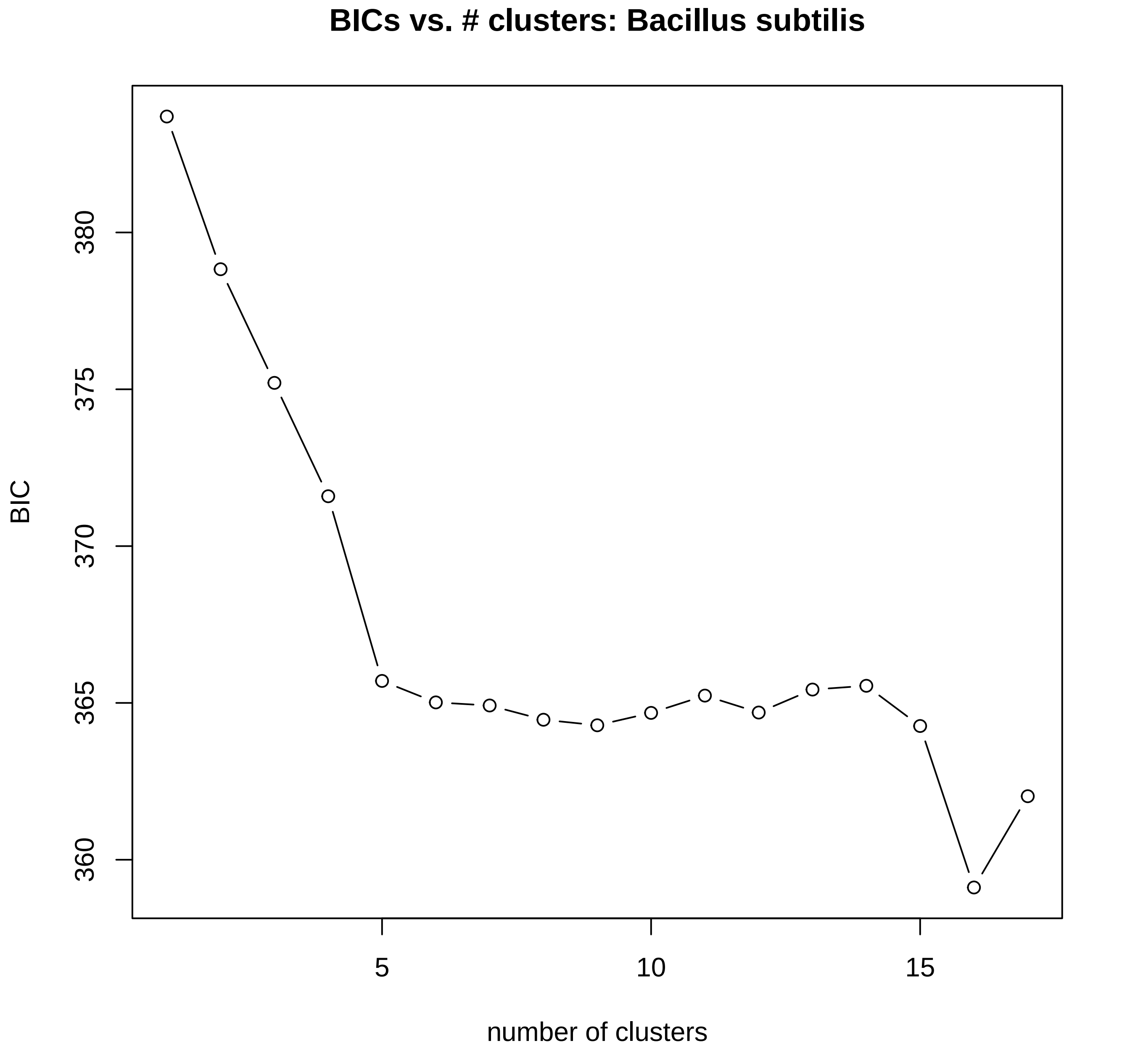
BICs vs. # clusters: Bacillus subtilis
BIC
380
375
370
365
360
5
10
15
number of clusters

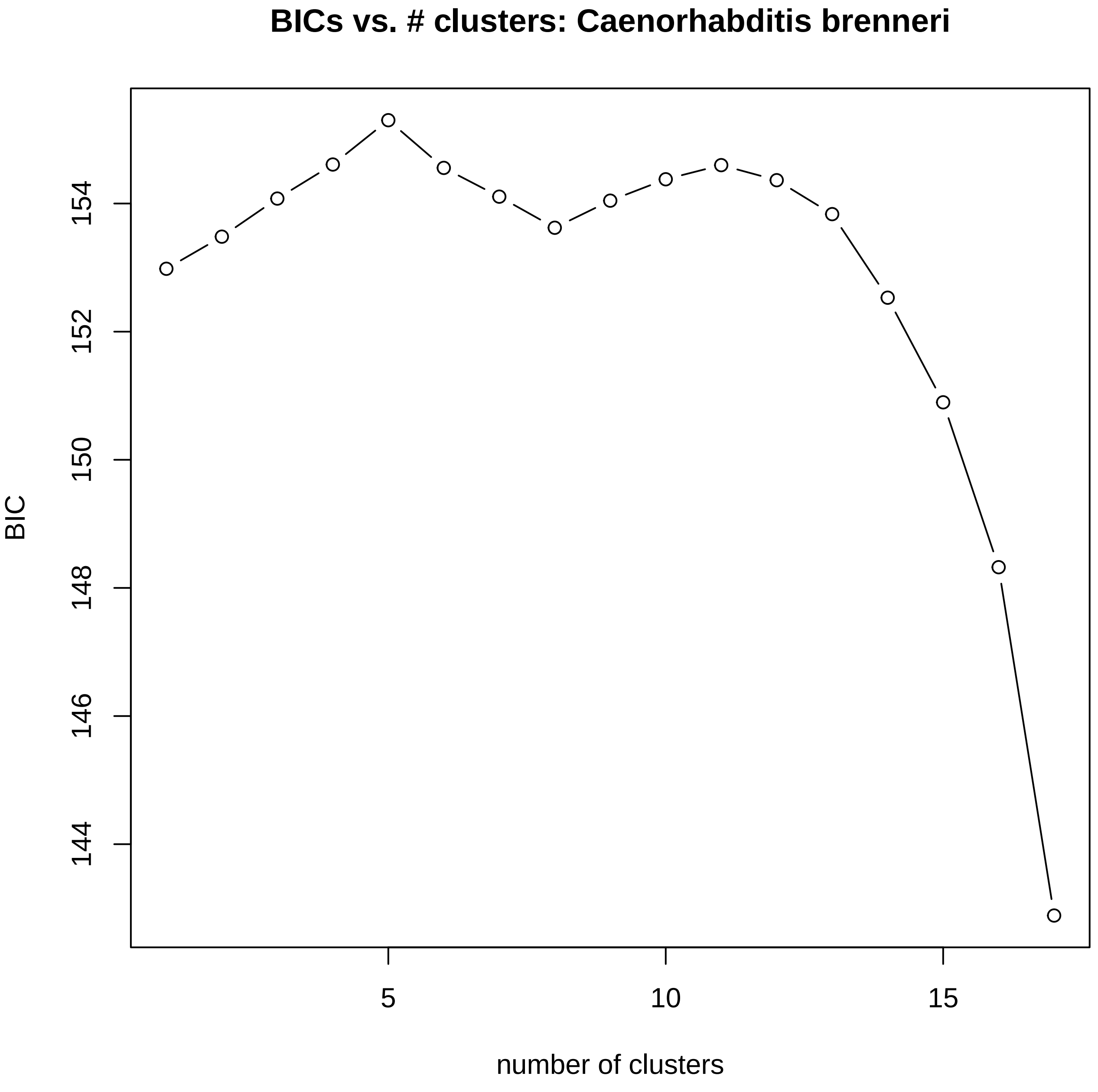
BICs vs. # clusters: Caenorhabditis brenneri
BIC
154
152
150
148
146
144
5
10
15
number of clusters

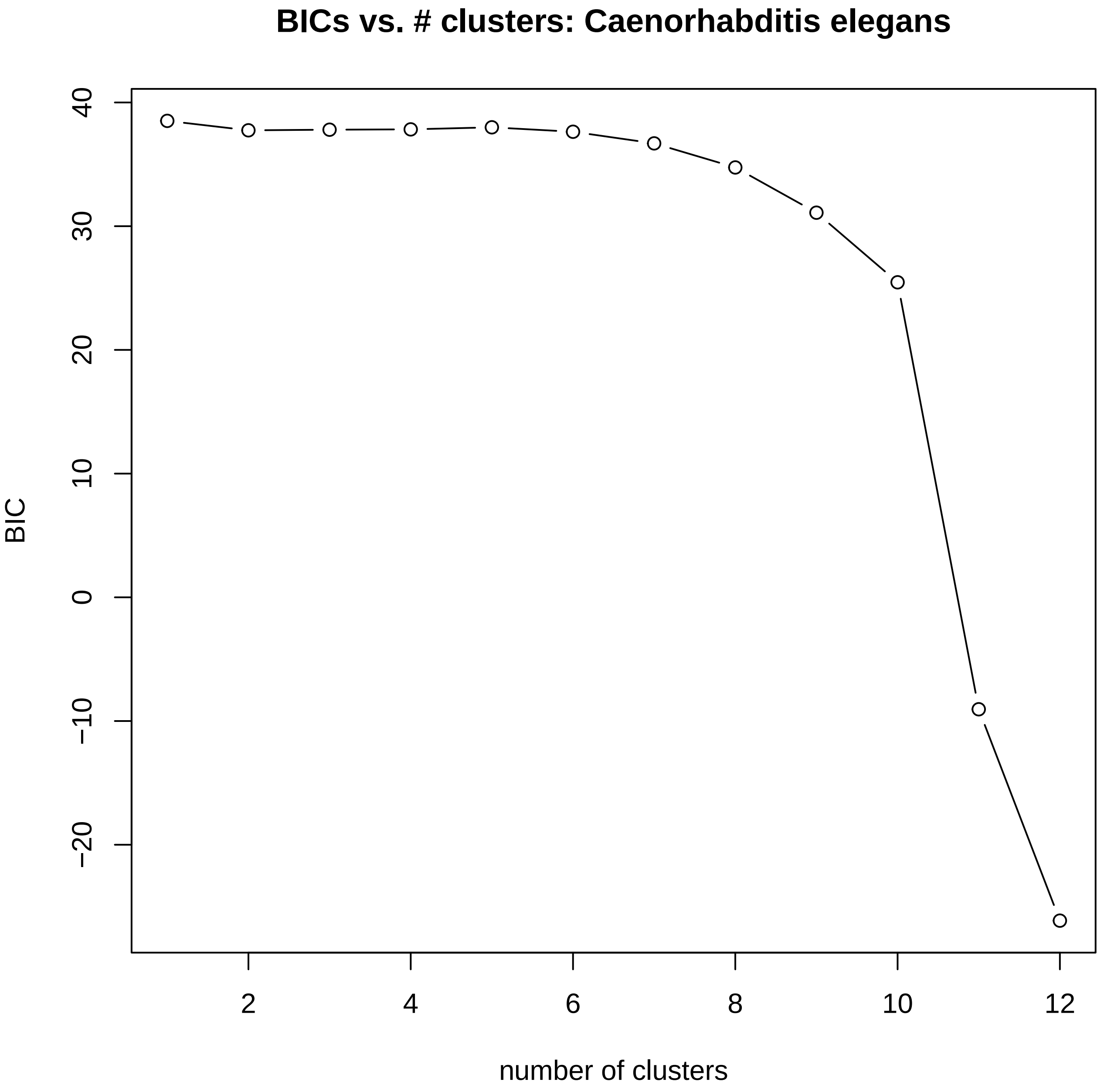
BICs vs. # clusters: Caenorhabditis elegans
BIC
40
30
20
10
0
−10
−20
2
4
6
8
10
12
number of clusters

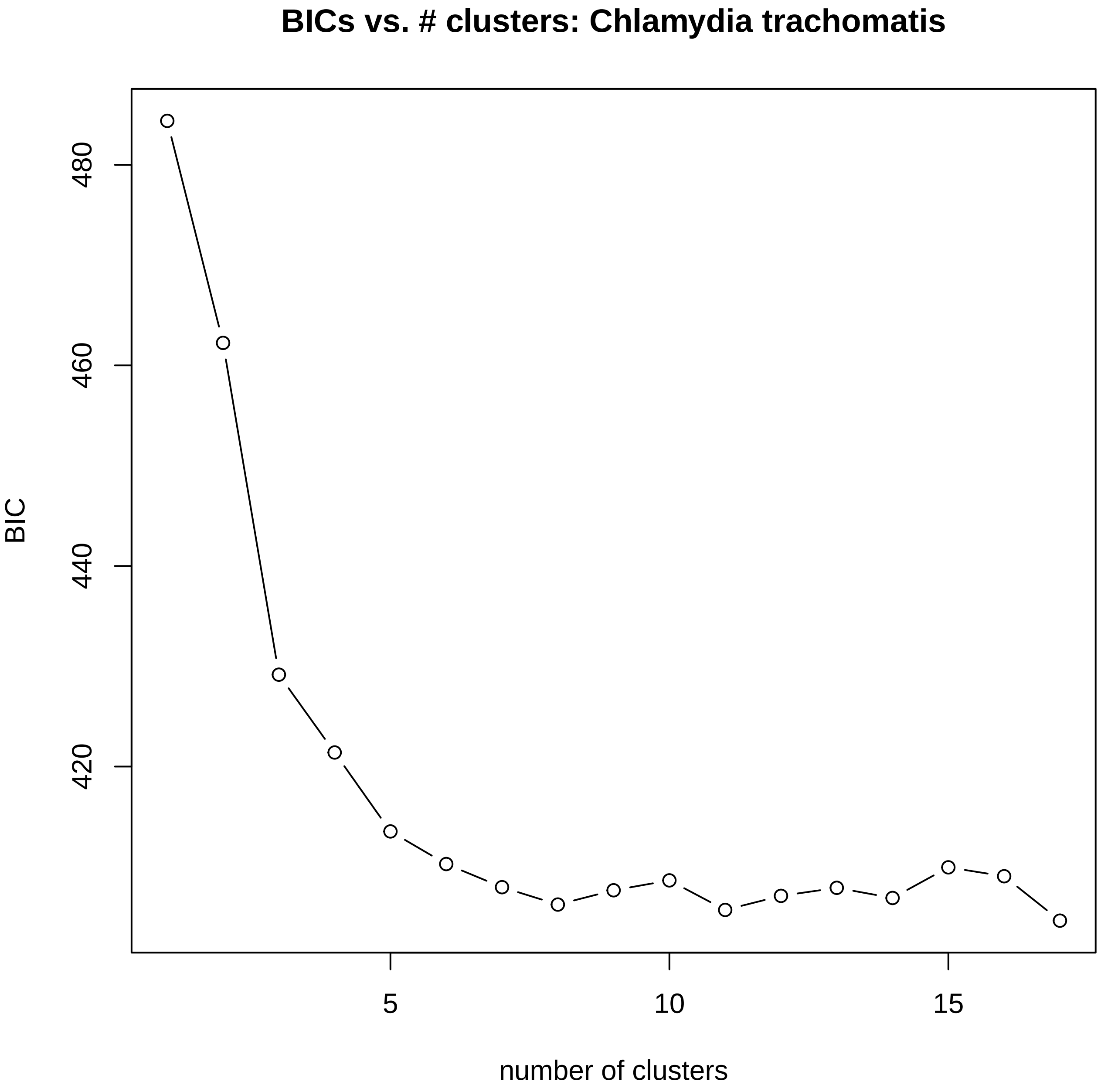

BICs vs. # clusters: Chlamydia trachomatis
BIC
480
460
440
420
5
10
15
number of clusters

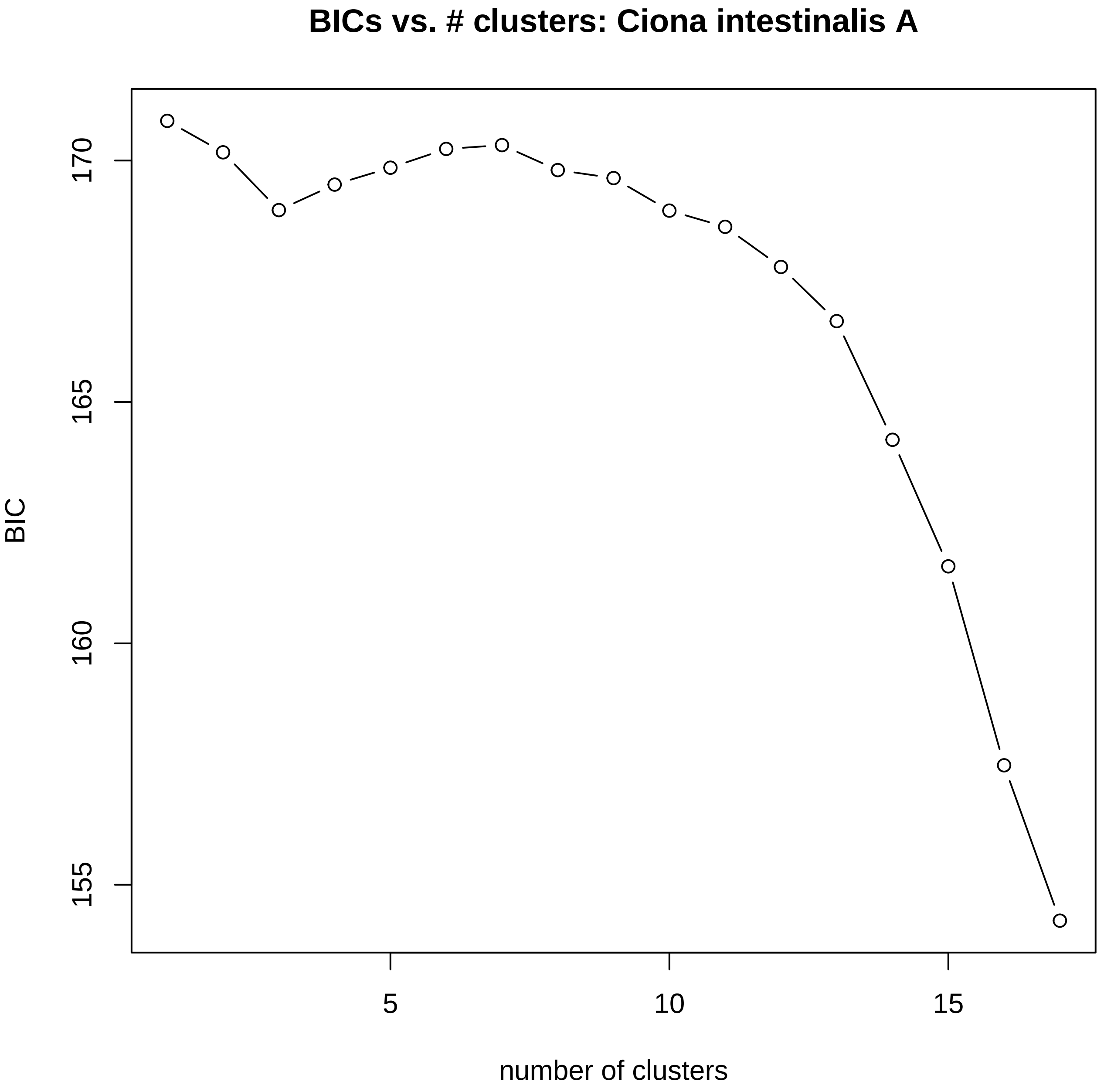

BICs vs. # clusters: Ciona intestinalis A
BIC
170
165
160
155
5
10
15
number of clusters

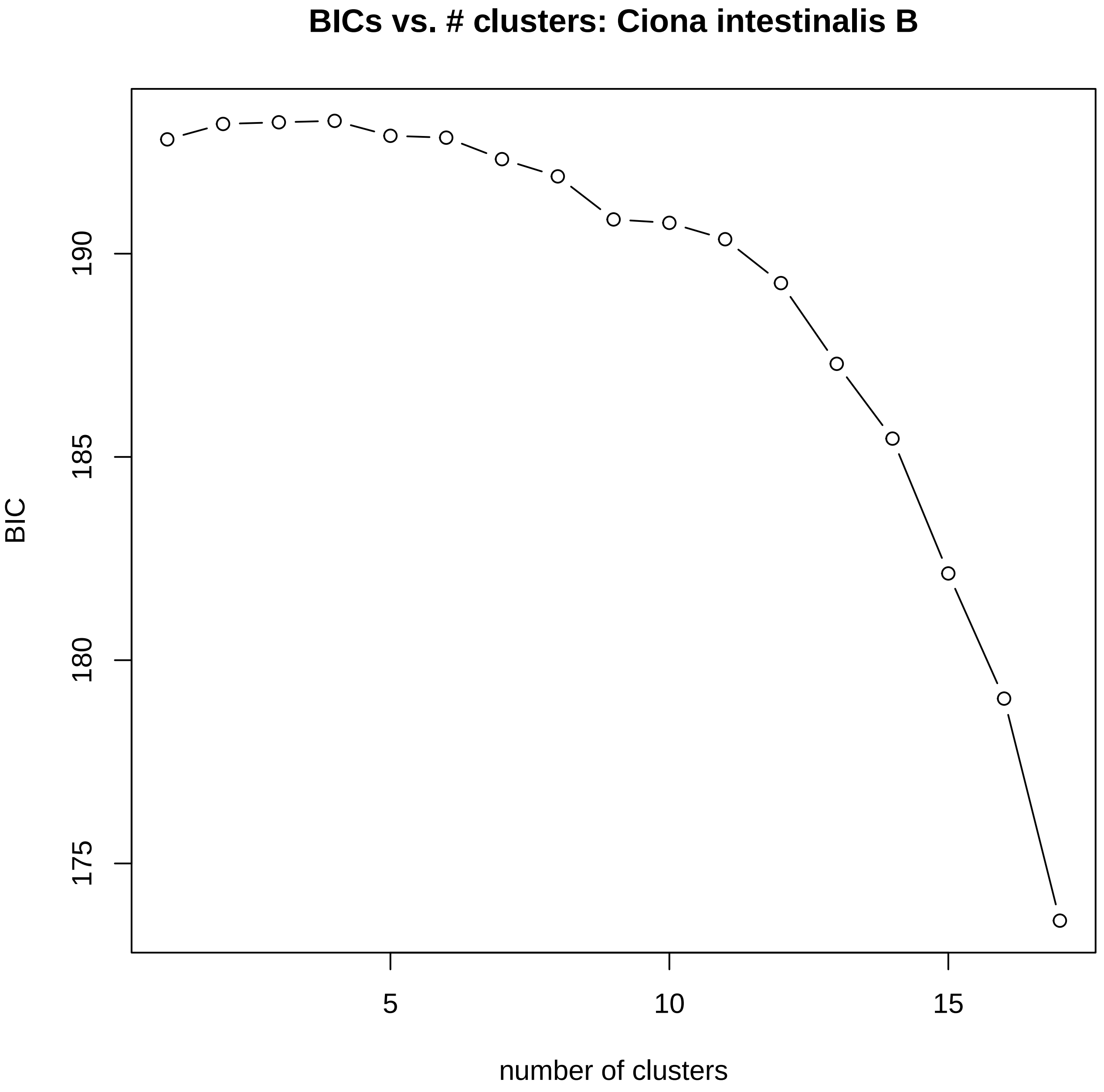
BICs vs. # clusters: Ciona intestinalis B
BIC
175
180
185
190
5
10
15
number of clusters

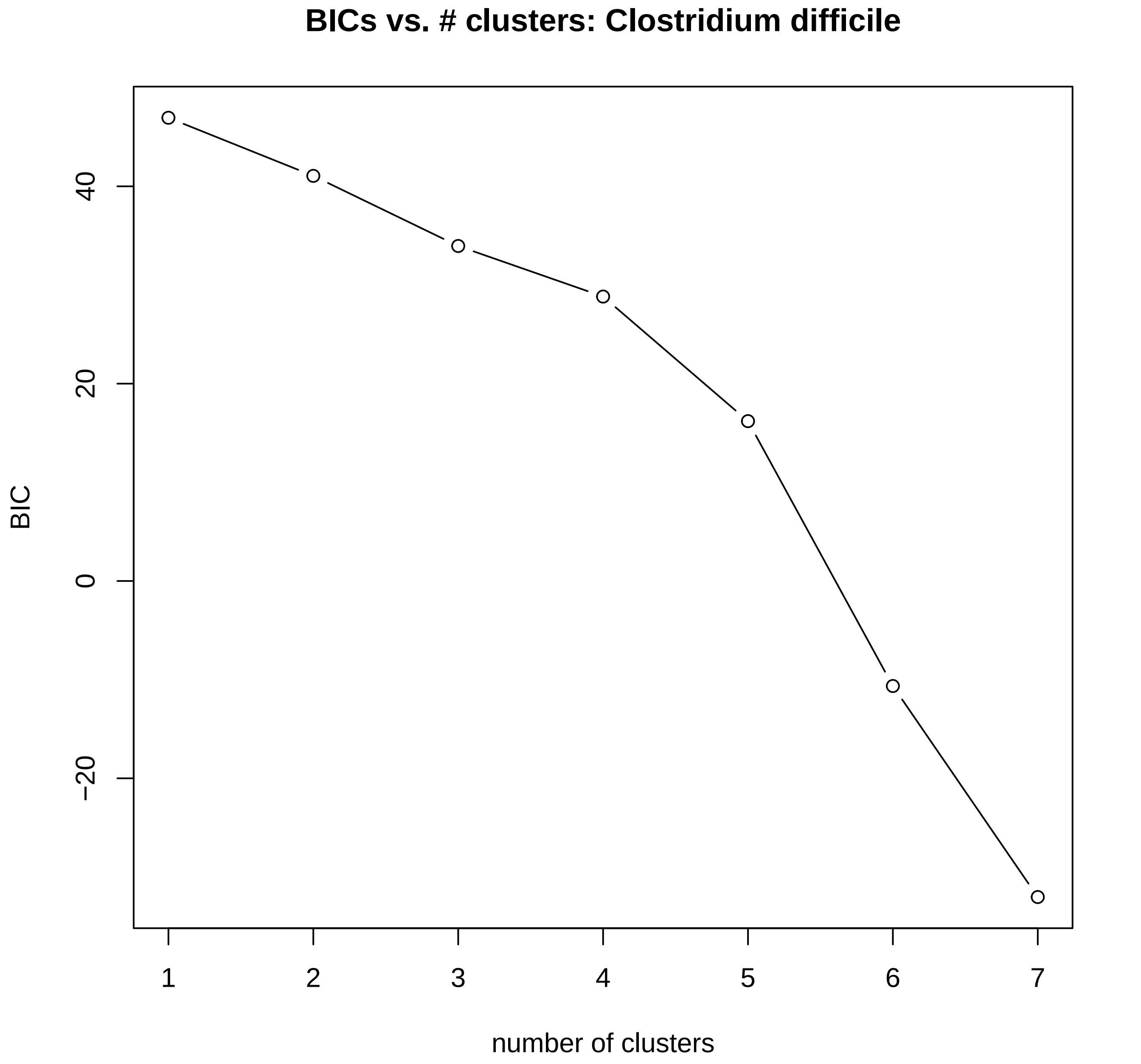

BICs vs. # clusters: Clostridium difficile
BIC
40
20
0
−20
1
2
3
4
5
6
7
number of clusters

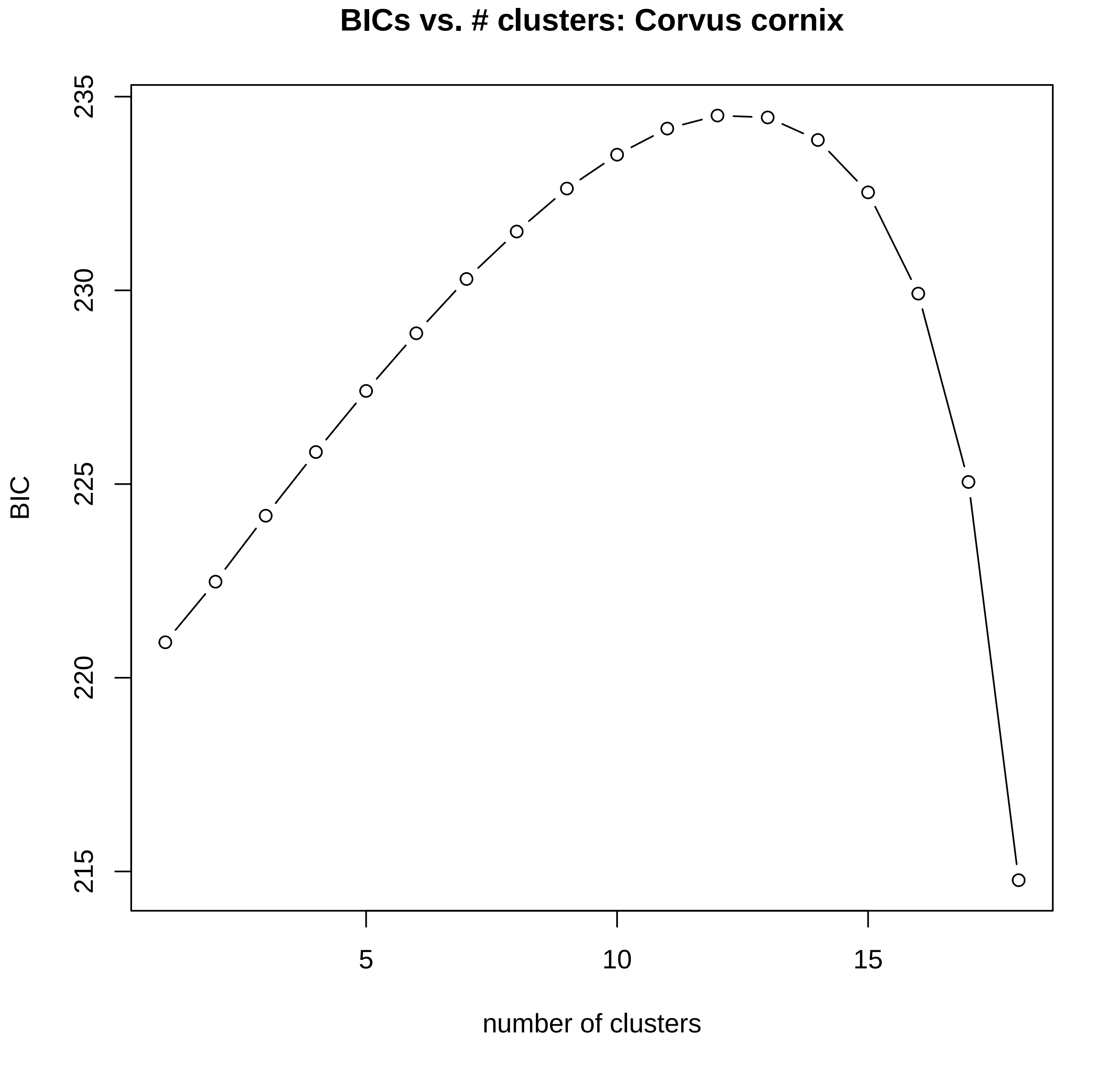

BICs vs. # clusters: Corvus cornix
BIC
215
220
225
230
235
5
10
15
number of clusters

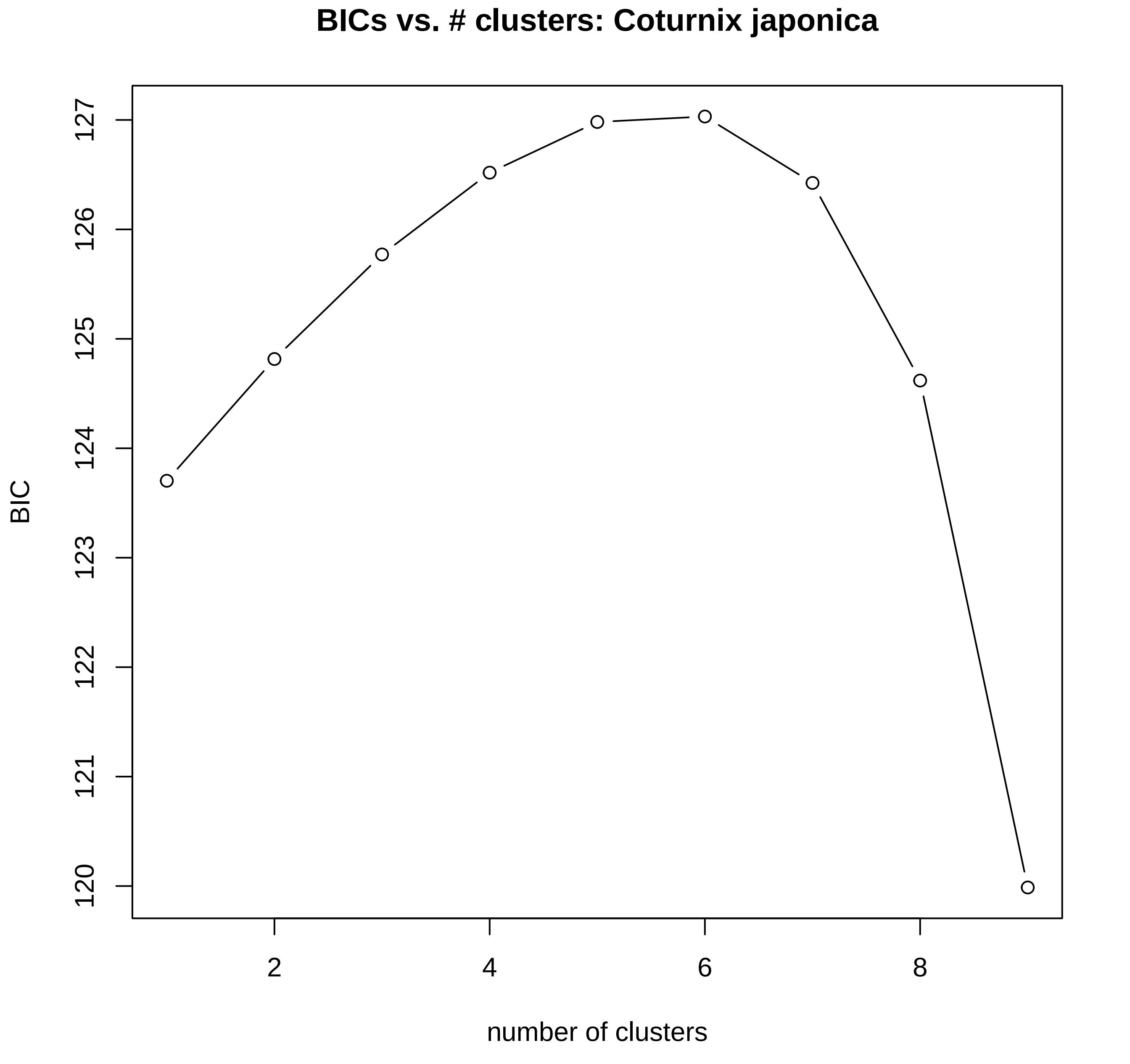
BICs vs. # clusters: Coturnix japonica
BIC
120
121
122
123
124
125
126
127
2
4
6
8
number of clusters

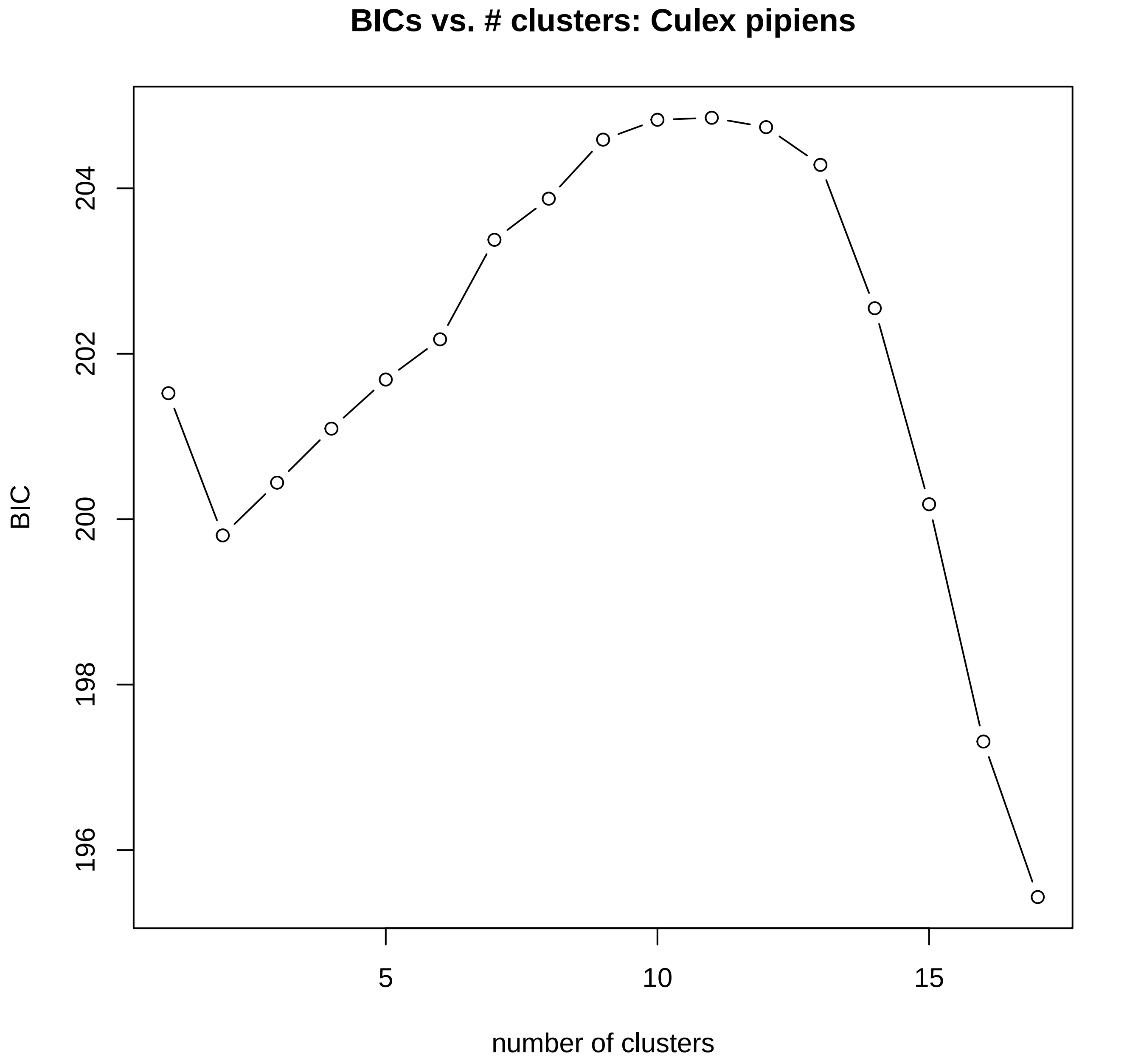

BICs vs. # clusters: Culex pipiens
BIC
196
198
200
202
204
5
10
15
number of clusters

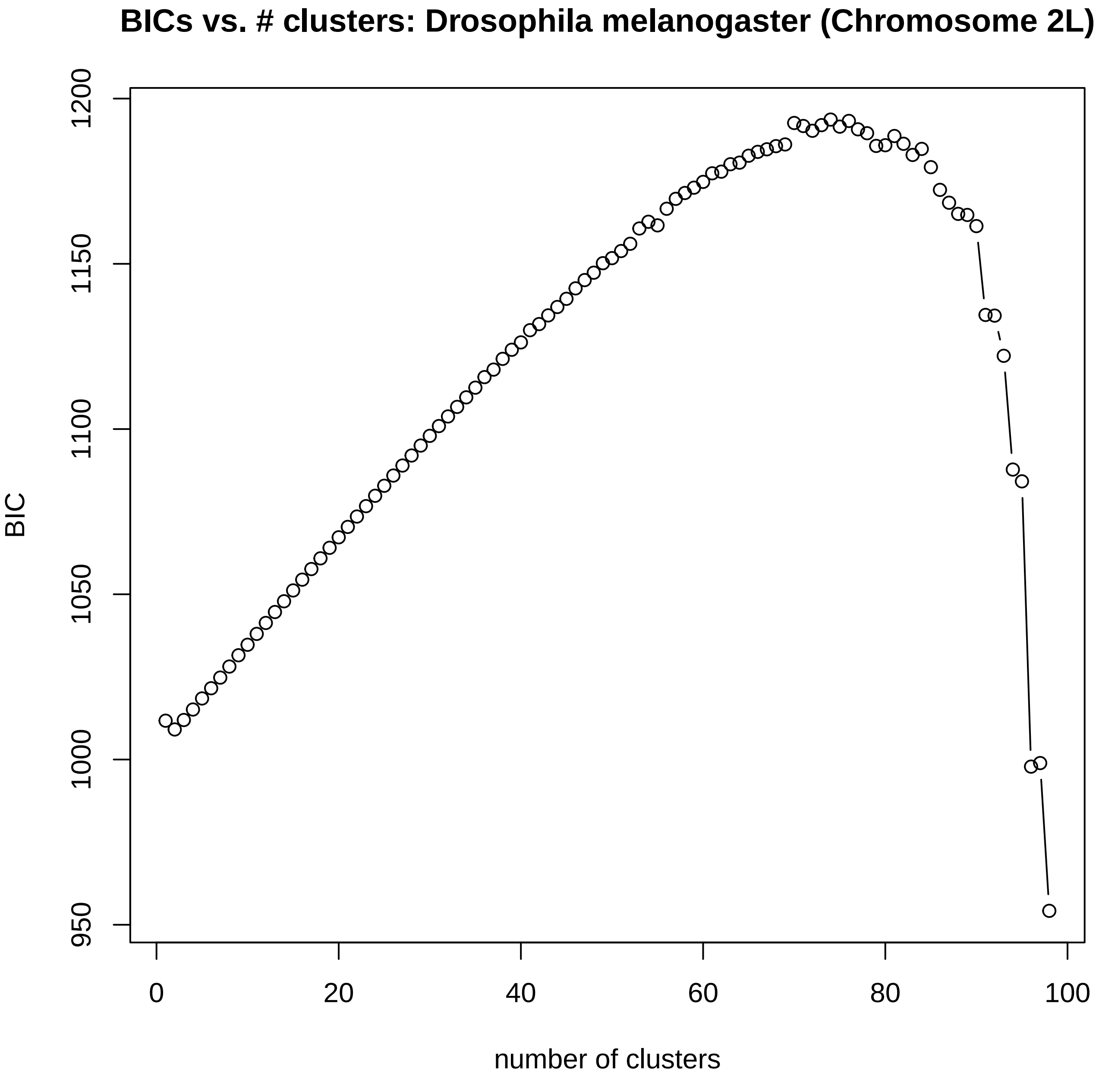

BICs vs. # clusters: Drosophila melanogaster (Chromosome 2L)
BIC
1200
1150
1100
1050
1000
950
0
20
40
60
80
100
number of clusters

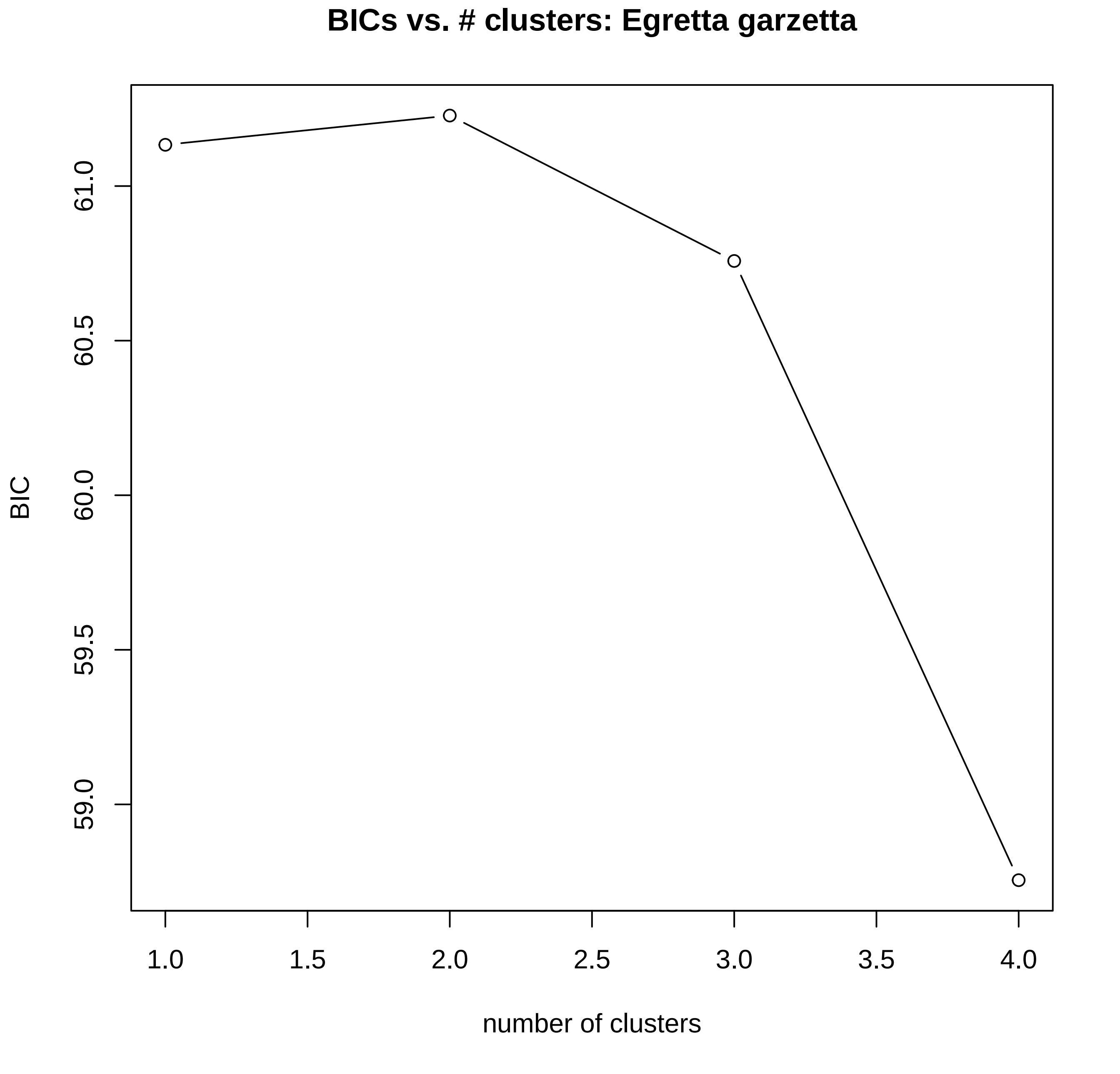

BICs vs. # clusters: Egretta garzetta
BIC
61.0
60.5
60.0
59.5
59.0
1.0
1.5
2.0
2.5
3.0
3.5
4.0
number of clusters

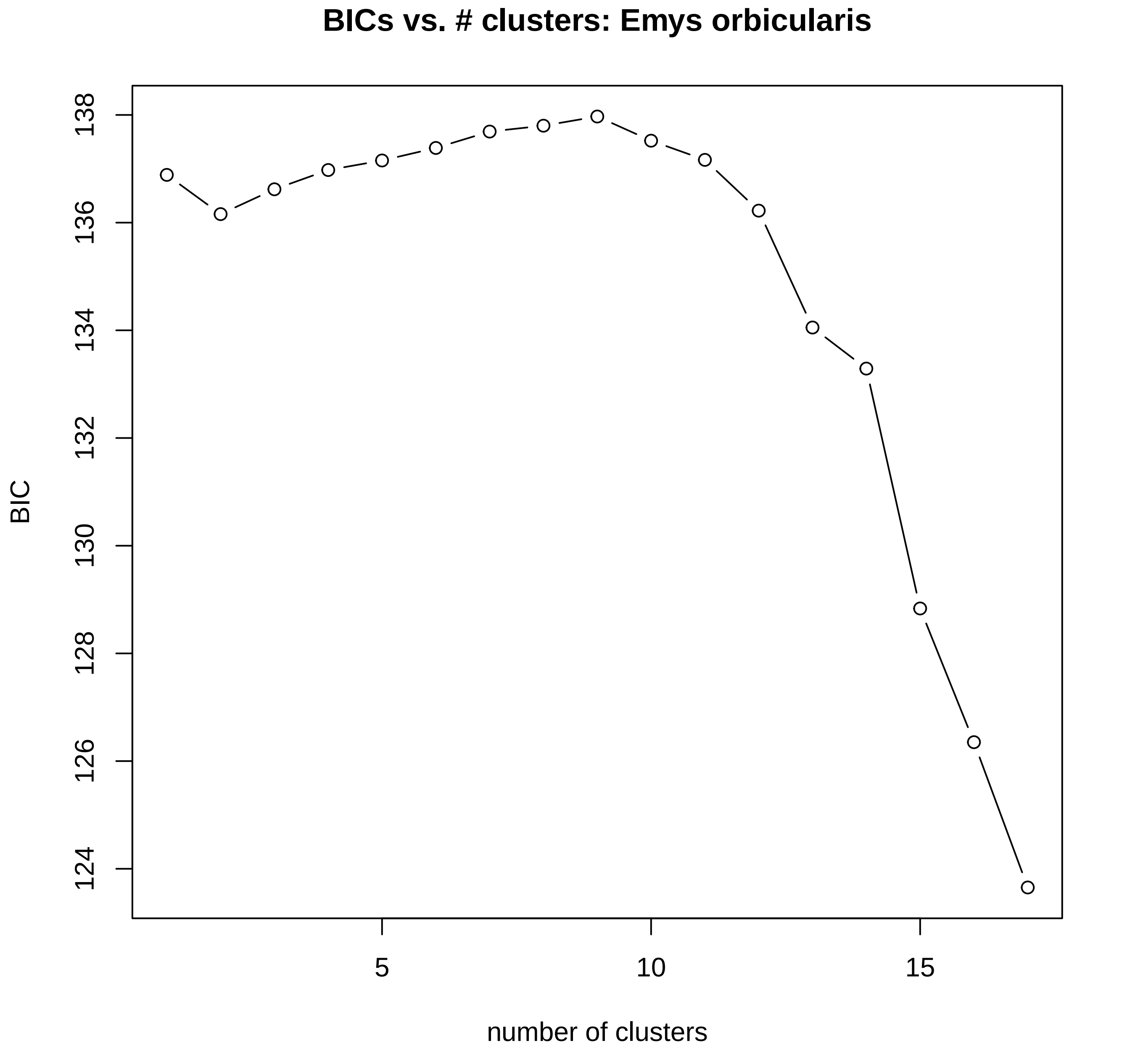
BICs vs. # clusters: Emys orbicularis
BIC
138
136
134
132
130
128
126
124
5
10
15
number of clusters

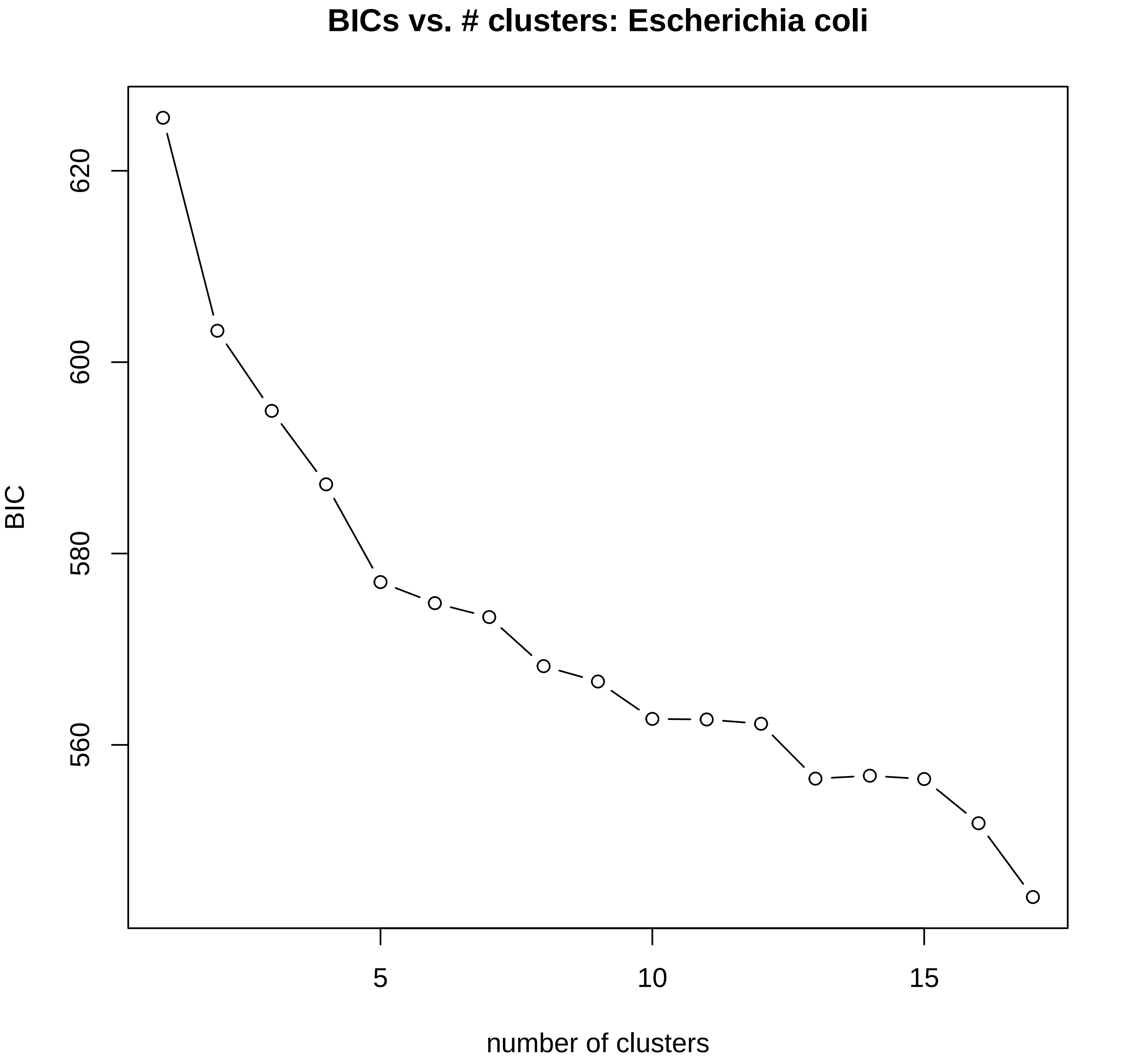

BICs vs. # clusters: Escherichia coli
BIC
620
600
580
560
5
10
15
number of clusters

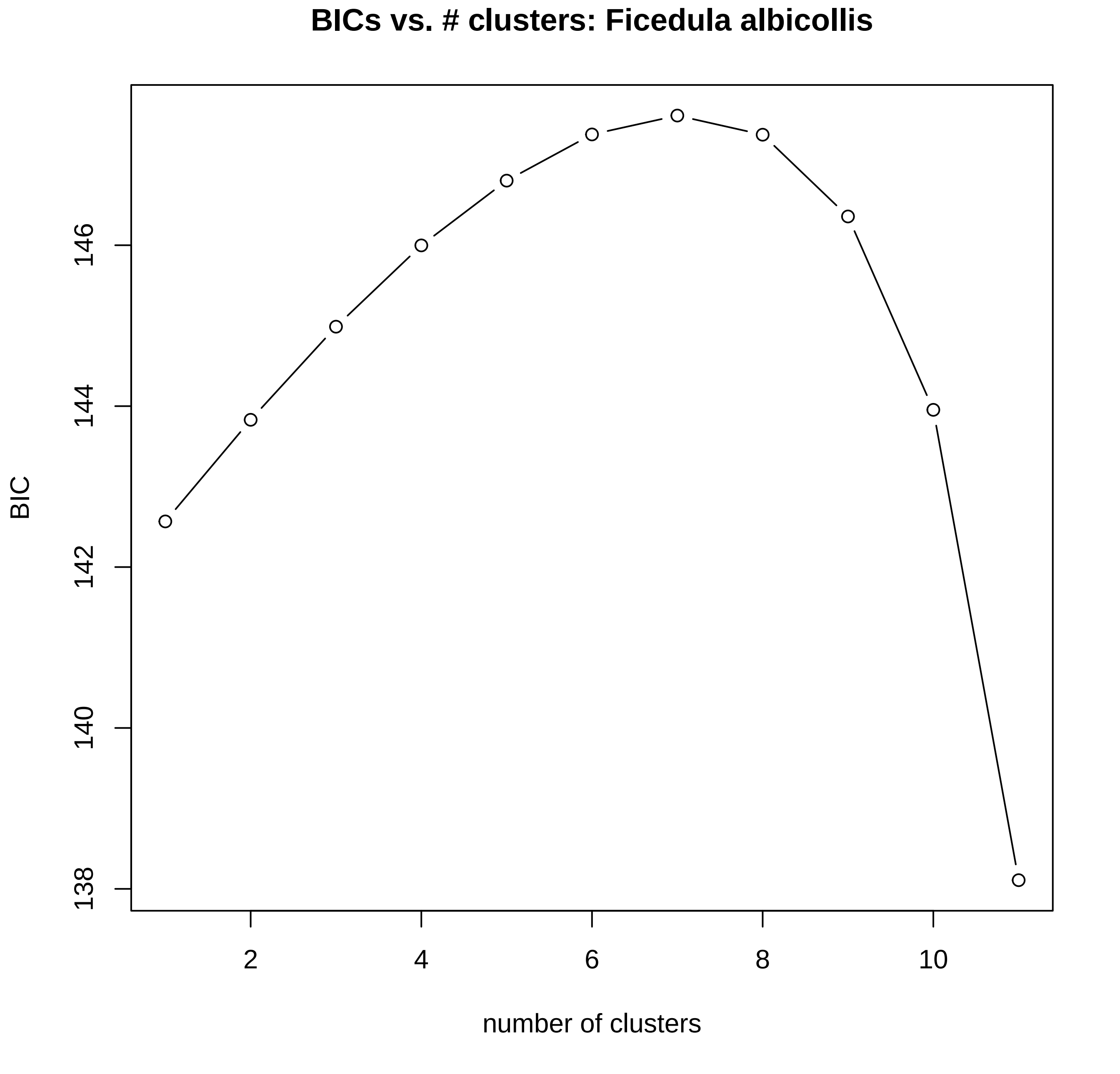

BICs vs. # clusters: Ficedula albicollis
BIC
138
140
142
144
146
2
4
6
8
10
number of clusters

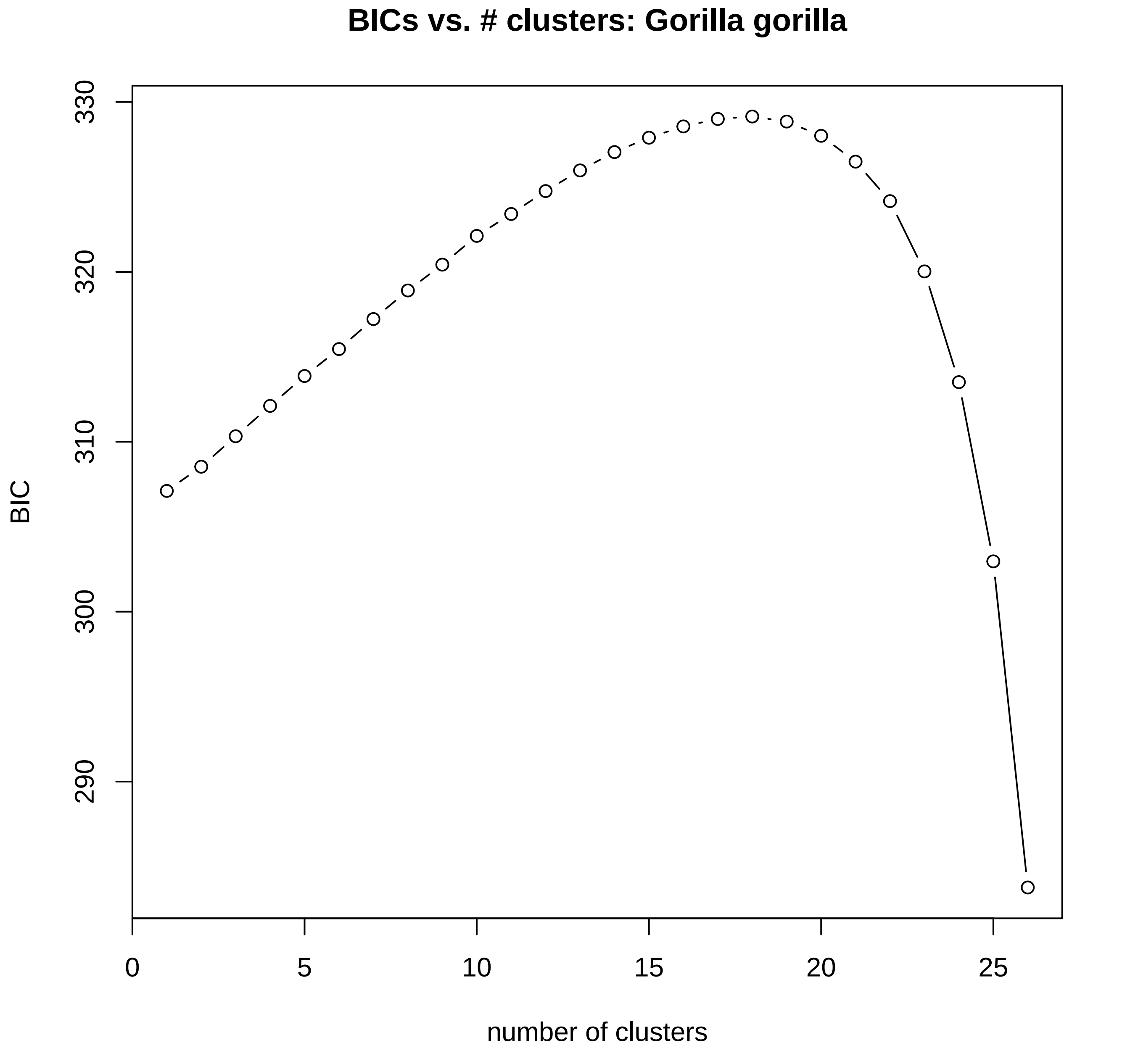

BICs vs. # clusters: Gorilla gorilla
BIC
330
320
310
300
290
0
5
10
15
20
25
number of clusters

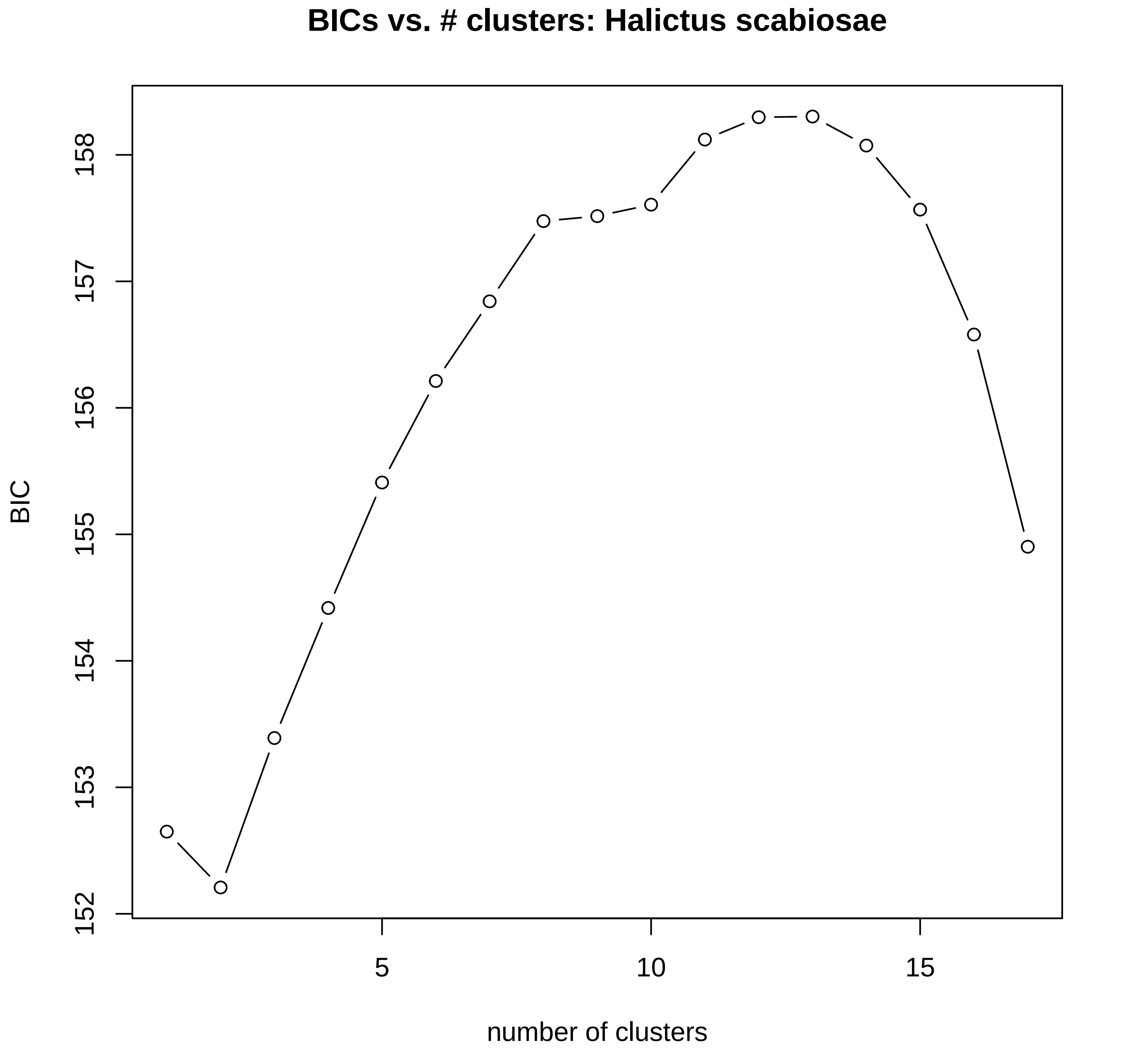

BICs vs. # clusters: Halictus scabiosae
BIC
152
153
154
155
156
157
158
5
10
15
number of clusters

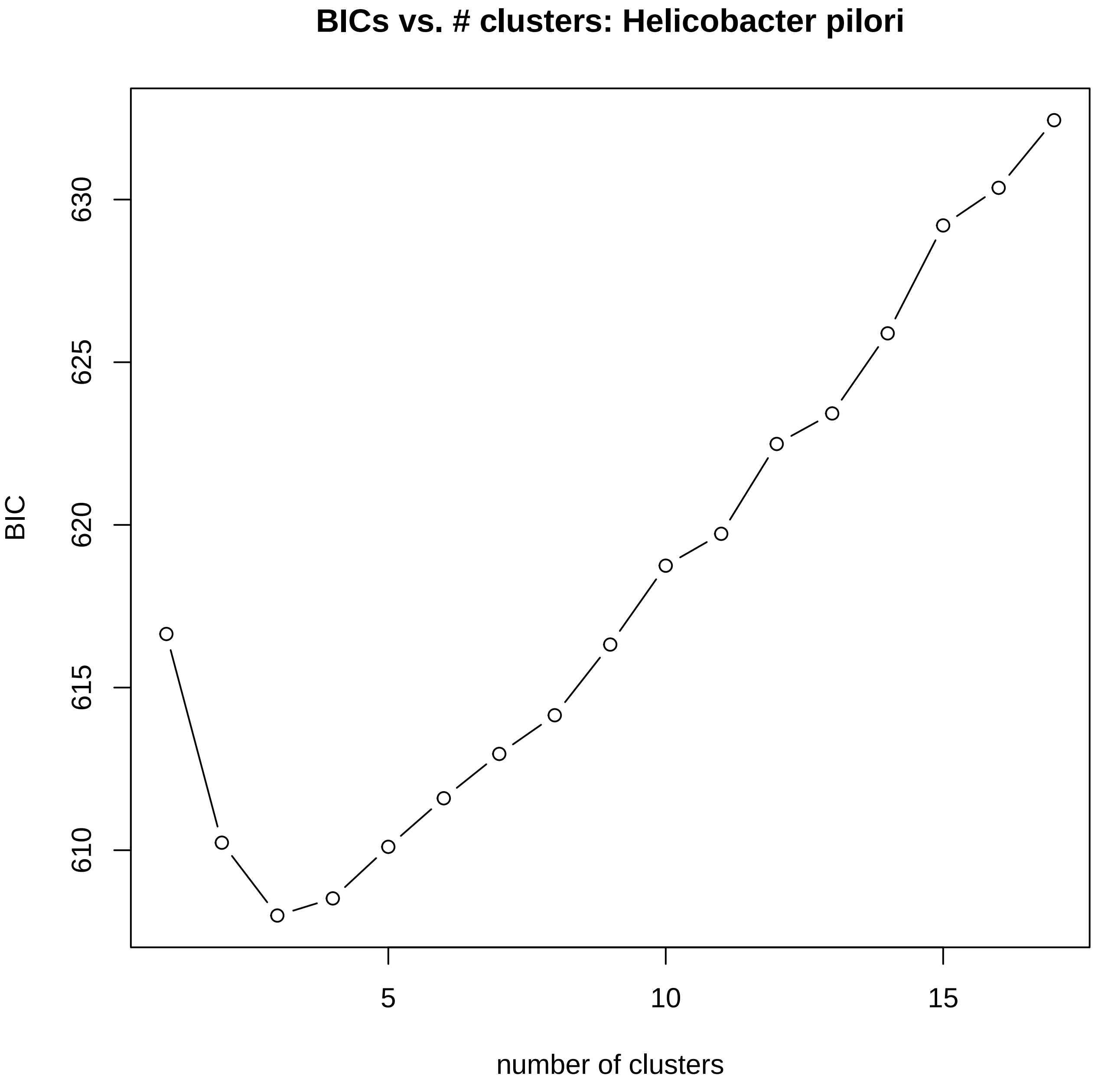
BICs vs. # clusters: Helicobacter pilori
BIC
610
615
620
625
630
5
10
15
number of clusters

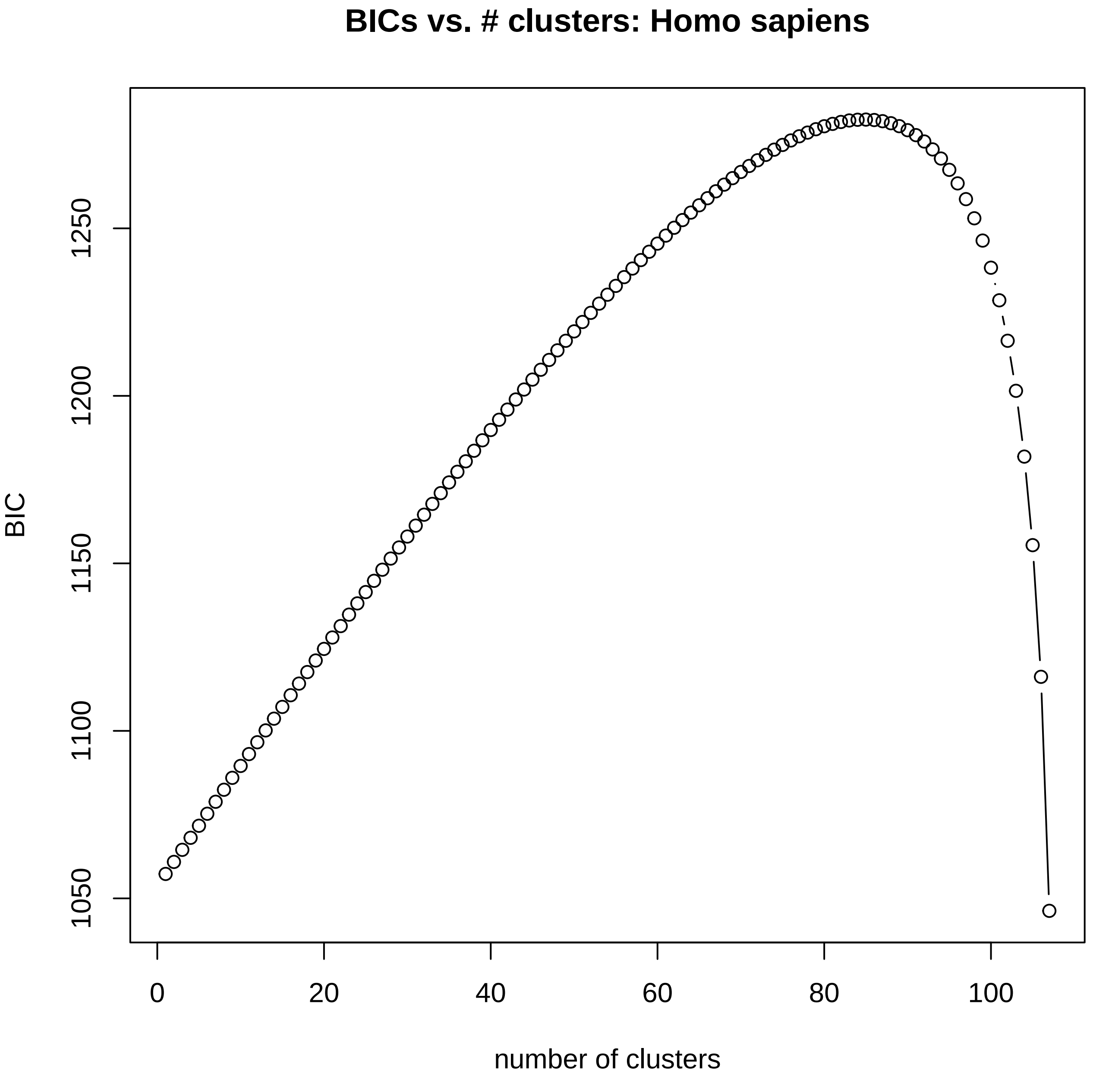
BICs vs. # clusters: Homo sapiens
BIC
1050
1100
1150
1200
1250
0
20
40
60
80
100
number of clusters

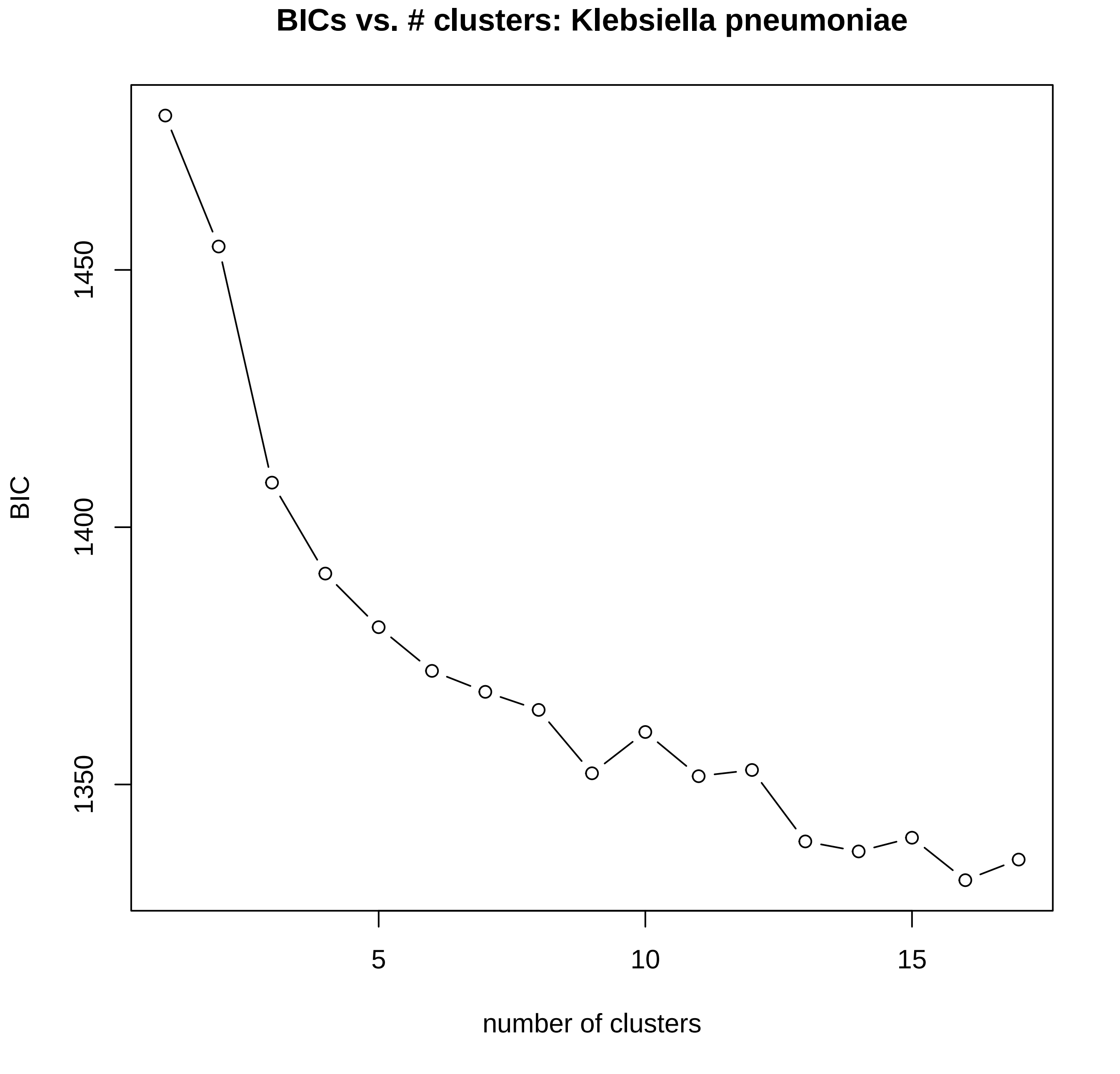
BICs vs. # clusters: Klebsiella pneumoniae
BIC
1450
1400
1350
5
10
15
number of clusters

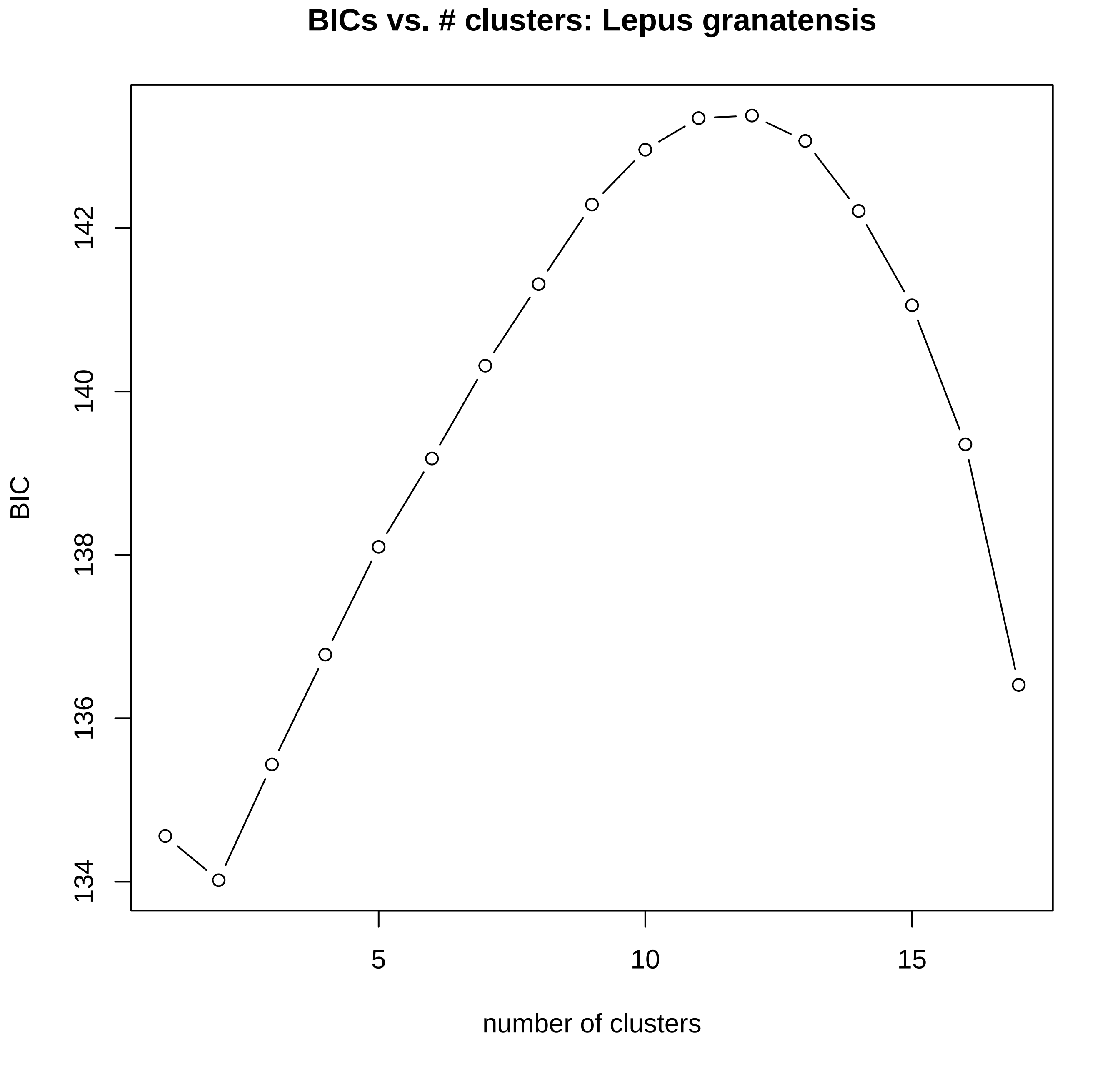
BICs vs. # clusters: Lepus granatensis
BIC
134
136
138
140
142
5
10
15
number of clusters

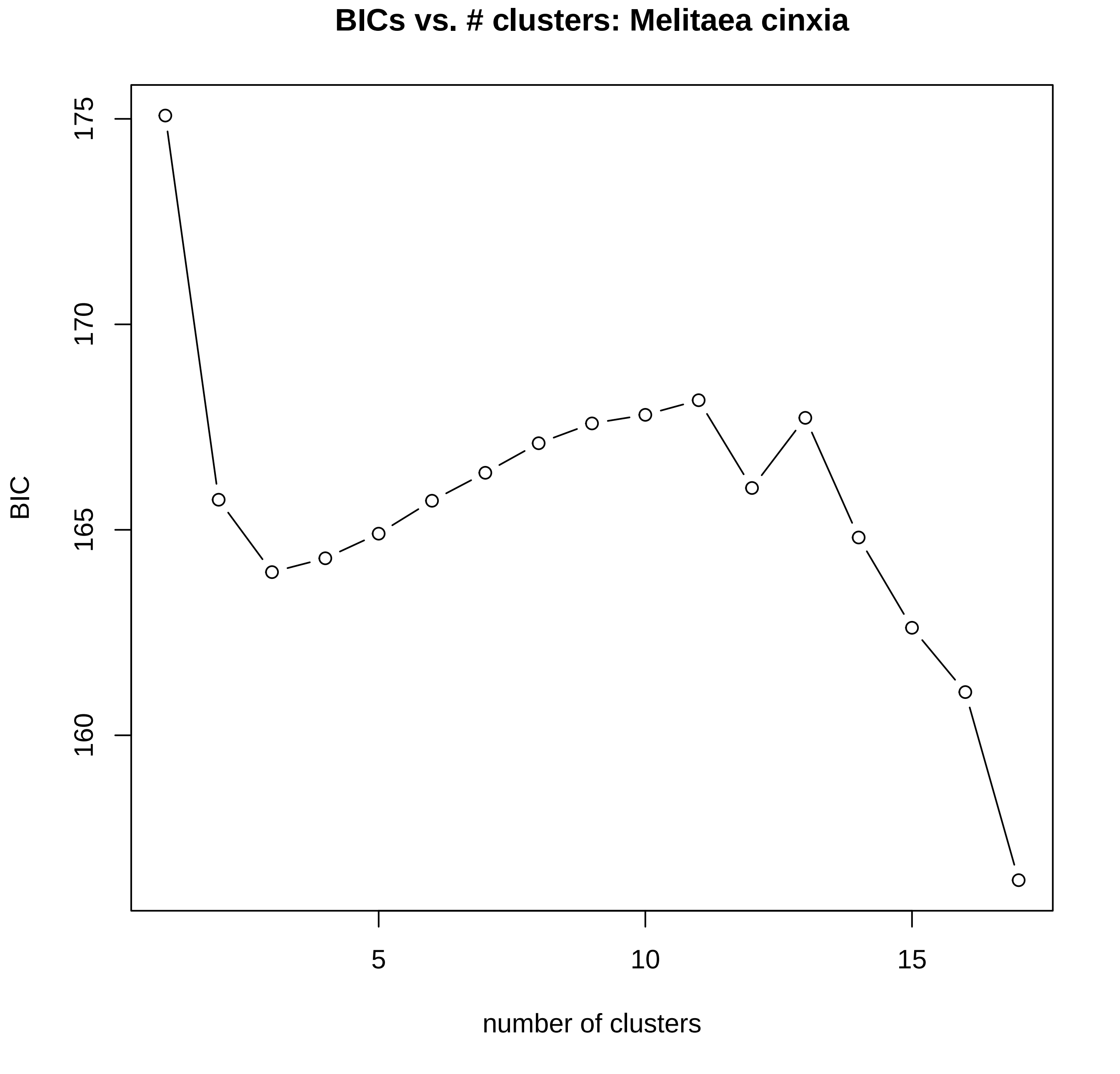
BICs vs. # clusters: Melitaea cinxia
BIC
175
170
165
160
5
10
15
number of clusters

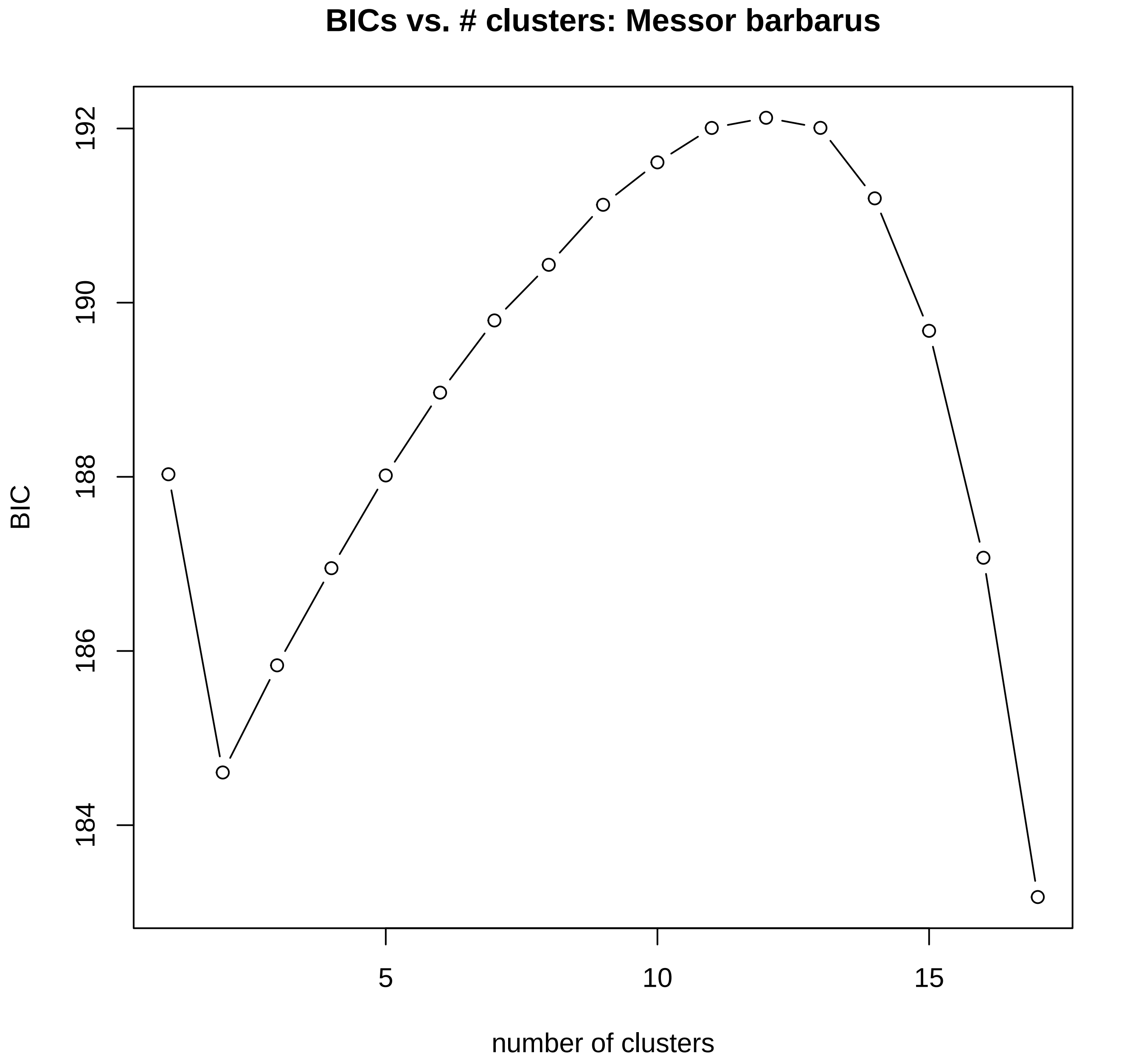

BICs vs. # clusters: Messor barbarus
192
190
188
186
184
BIC
5
10
15
number of clusters

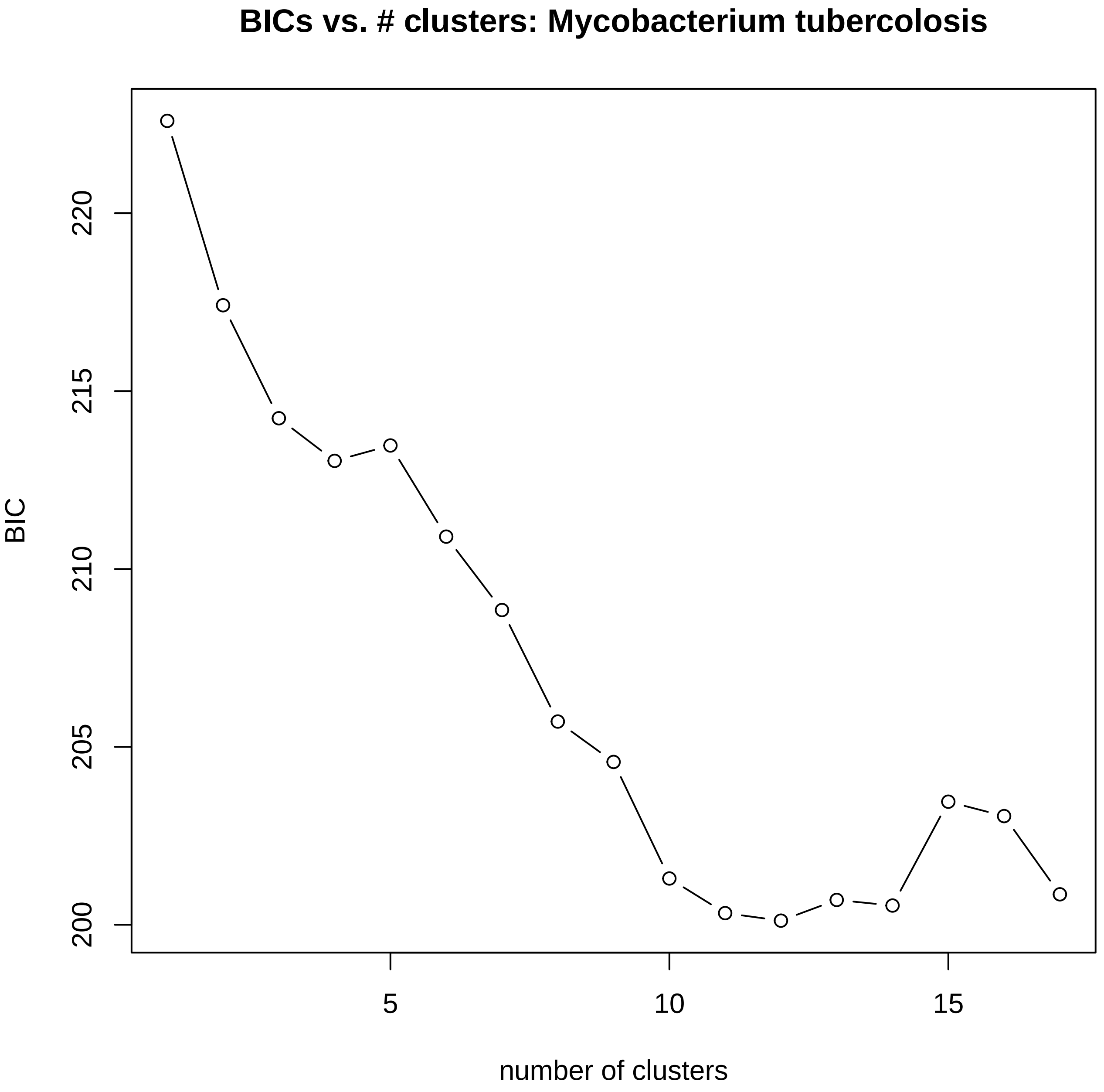

BICs vs. # clusters: Mycobacterium tubercolosis
BIC
220
215
210
205
200
5
10
15
number of clusters

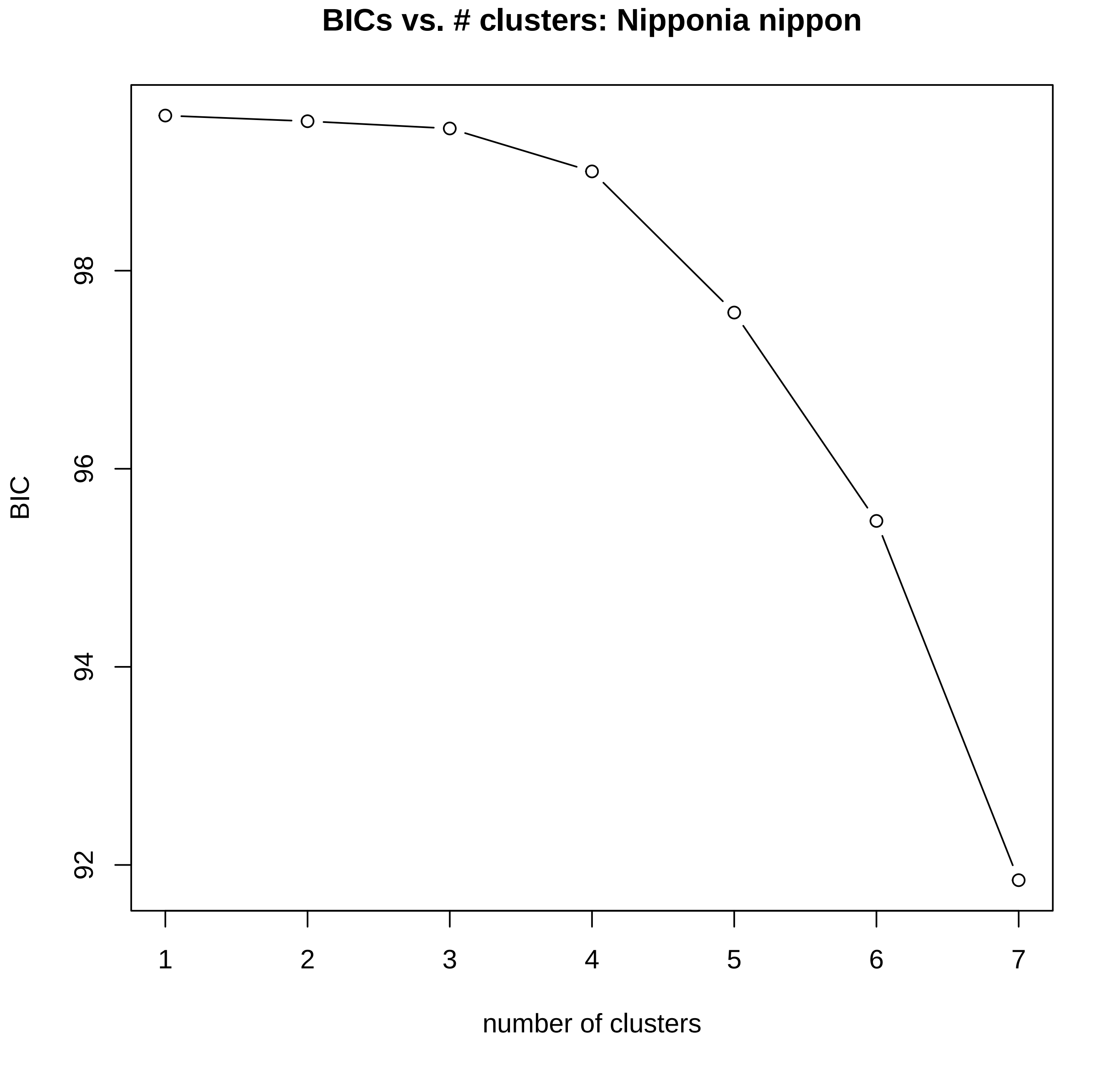

BICs vs. # clusters: Nipponia nippon
BIC
98
96
94
92
1
2
3
4
5
6
7
number of clusters

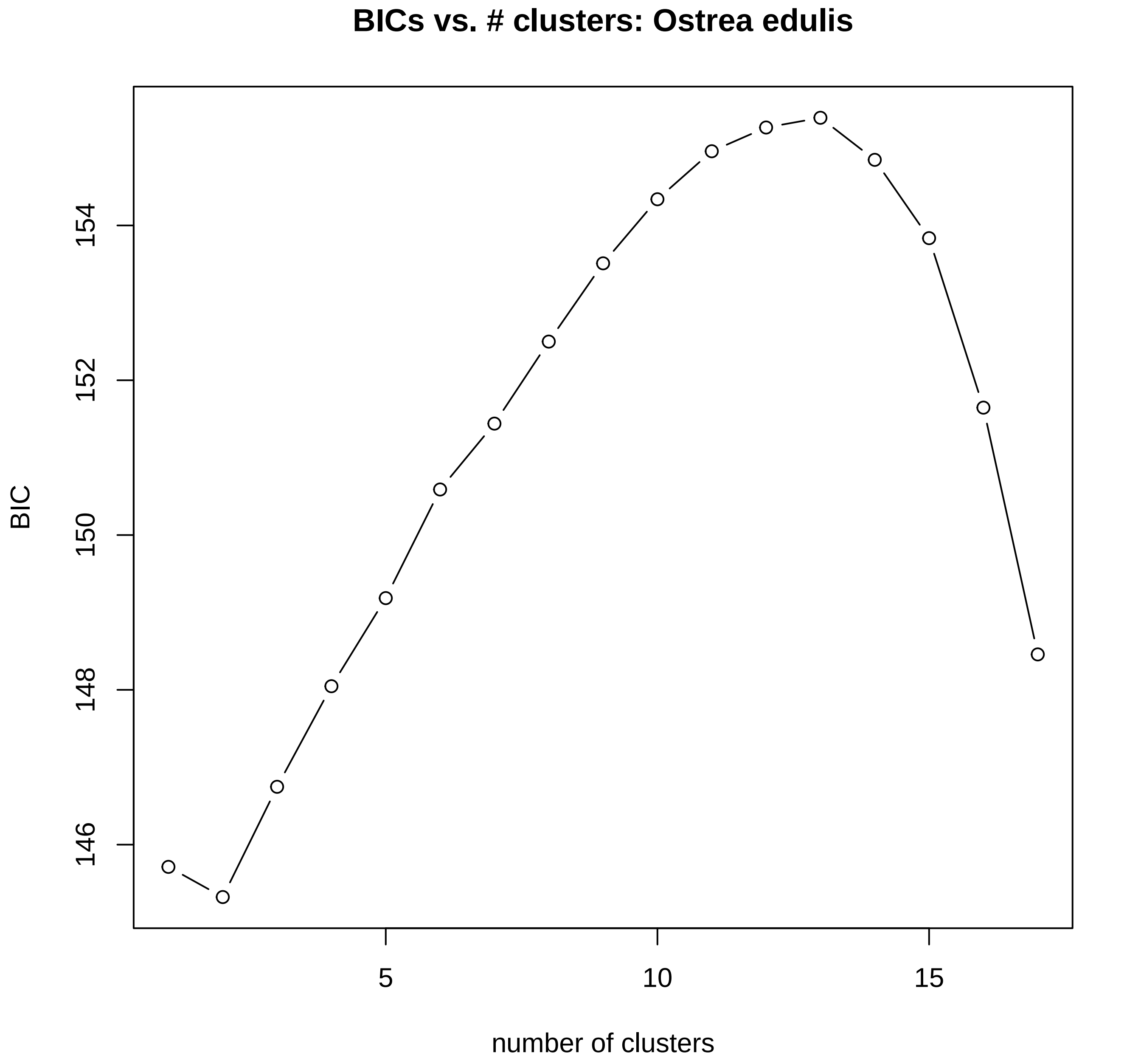
BICs vs. # clusters: Ostrea edulis
BIC
146
148
150
152
154
5
10
15
number of clusters

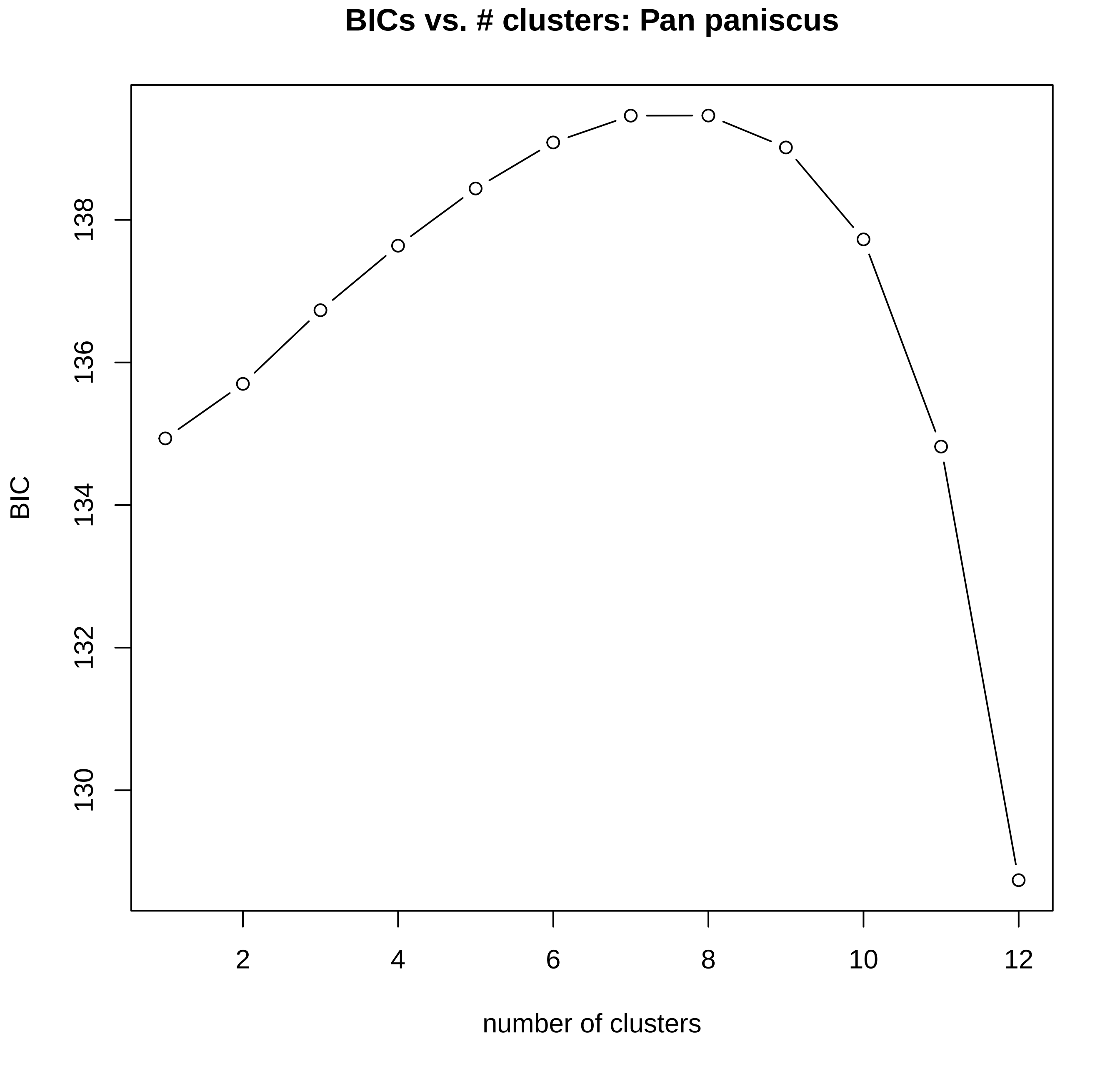

BICs vs. # clusters: Pan paniscus
BIC
138
136
134
132
130
2
4
6
8
10
12
number of clusters

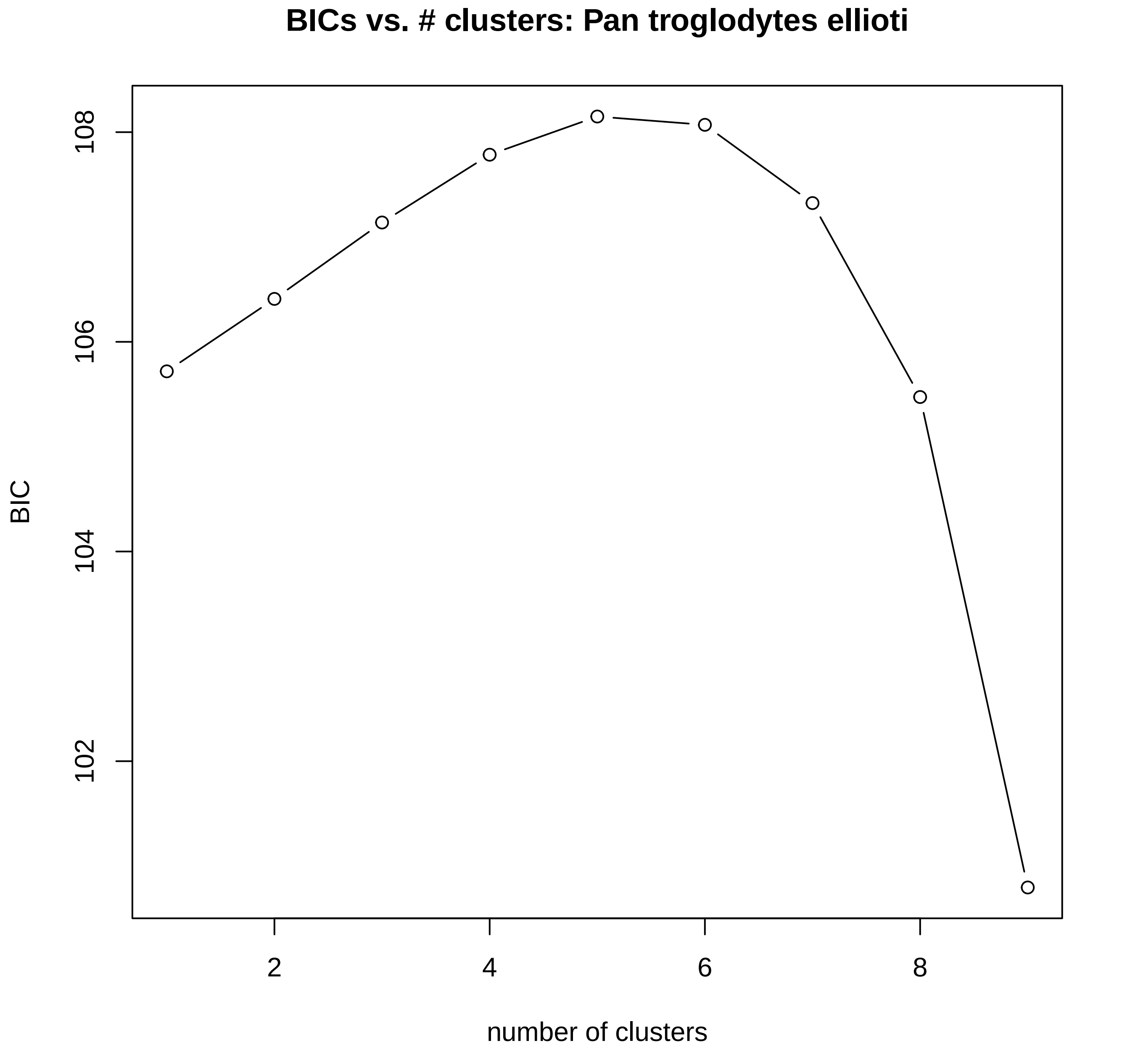
BICs vs. # clusters: Pan troglodytes ellioti
BIC
108
106
104
102
2
4
6
8
number of clusters

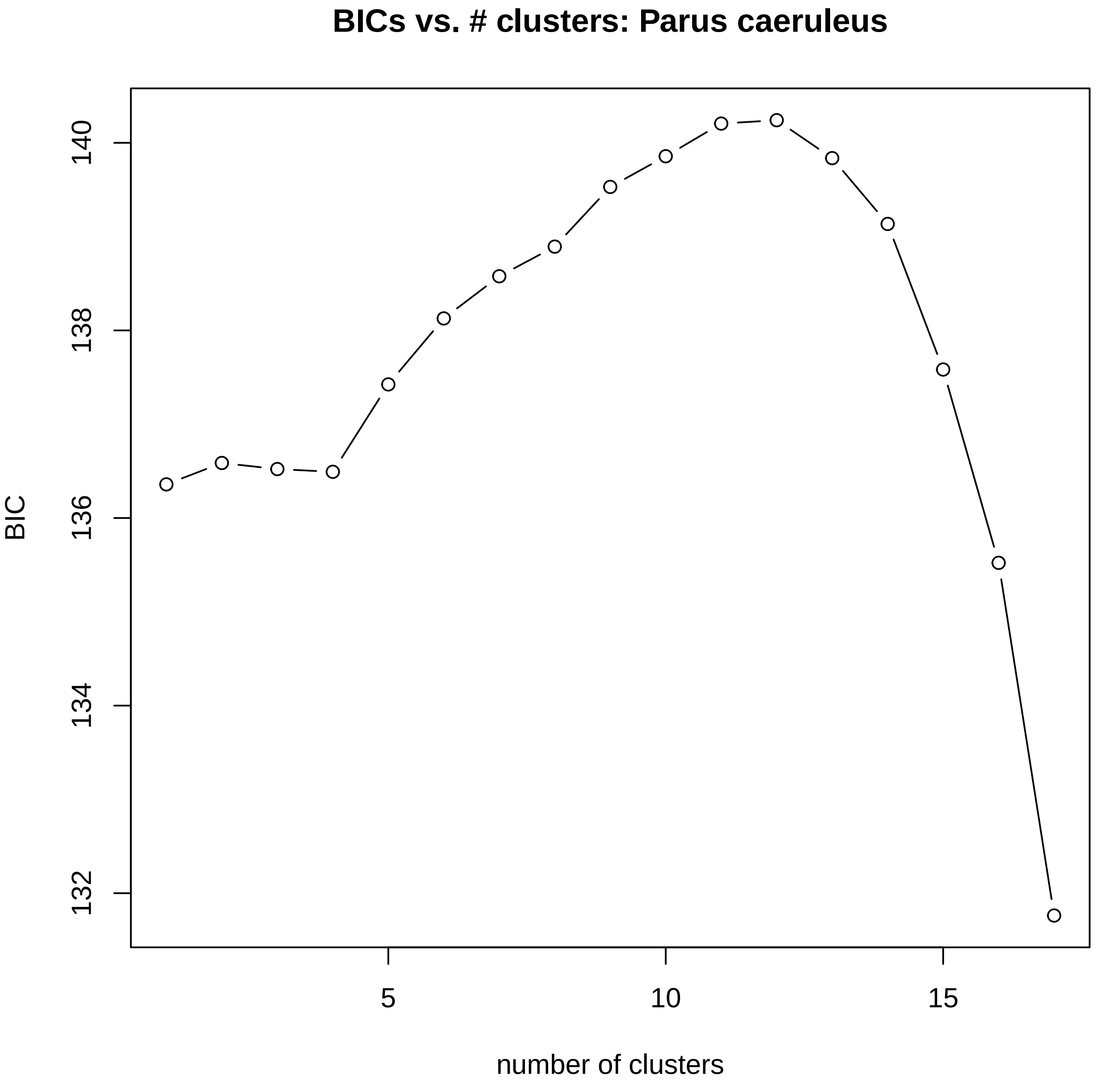

BICs vs. # clusters: Parus caeruleus
BIC
140
138
136
134
132
5
10
15
number of clusters

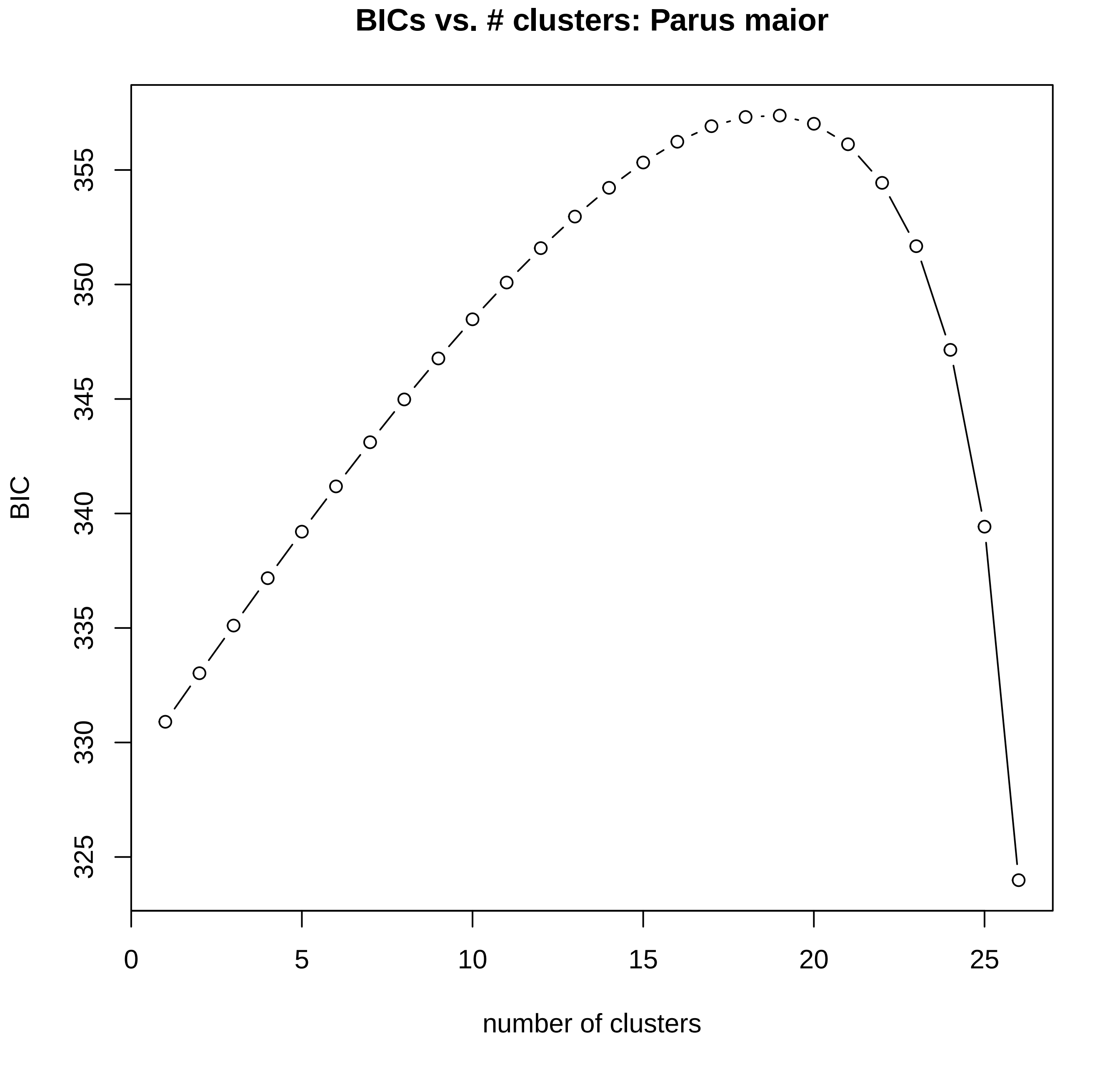

BICs vs. # clusters: Parus maior
BIC
325
330
335
340
345
350
355
0
5
10
15
20
25
number of clusters

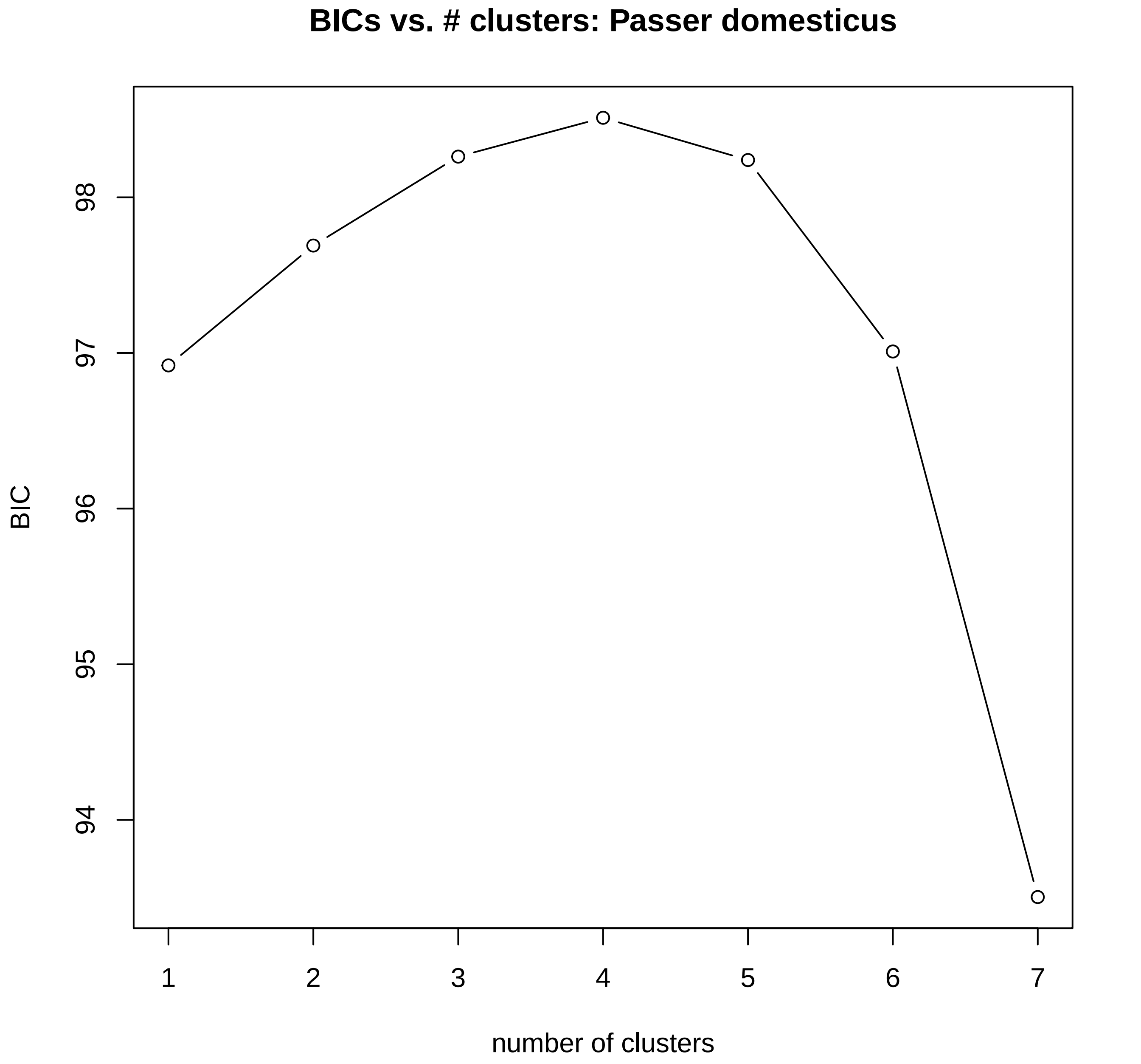
BICs vs. # clusters: Passer domesticus
BIC
98
97
96
95
94
1
2
3
4
5
6
7
number of clusters

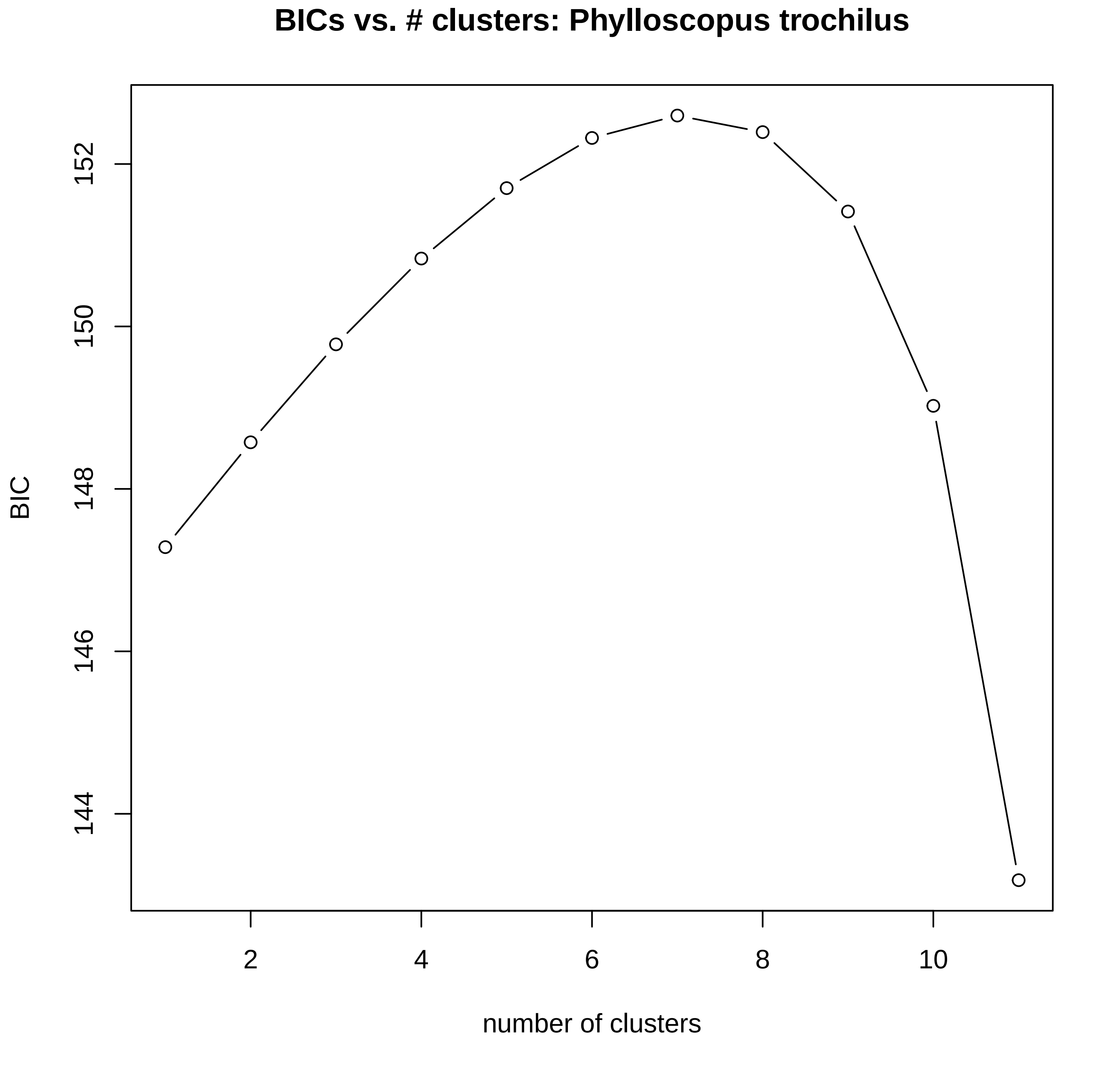

BICs vs. # clusters: Phylloscopus trochilus
BIC
144
146
148
150
152
2
4
6
8
10
number of clusters

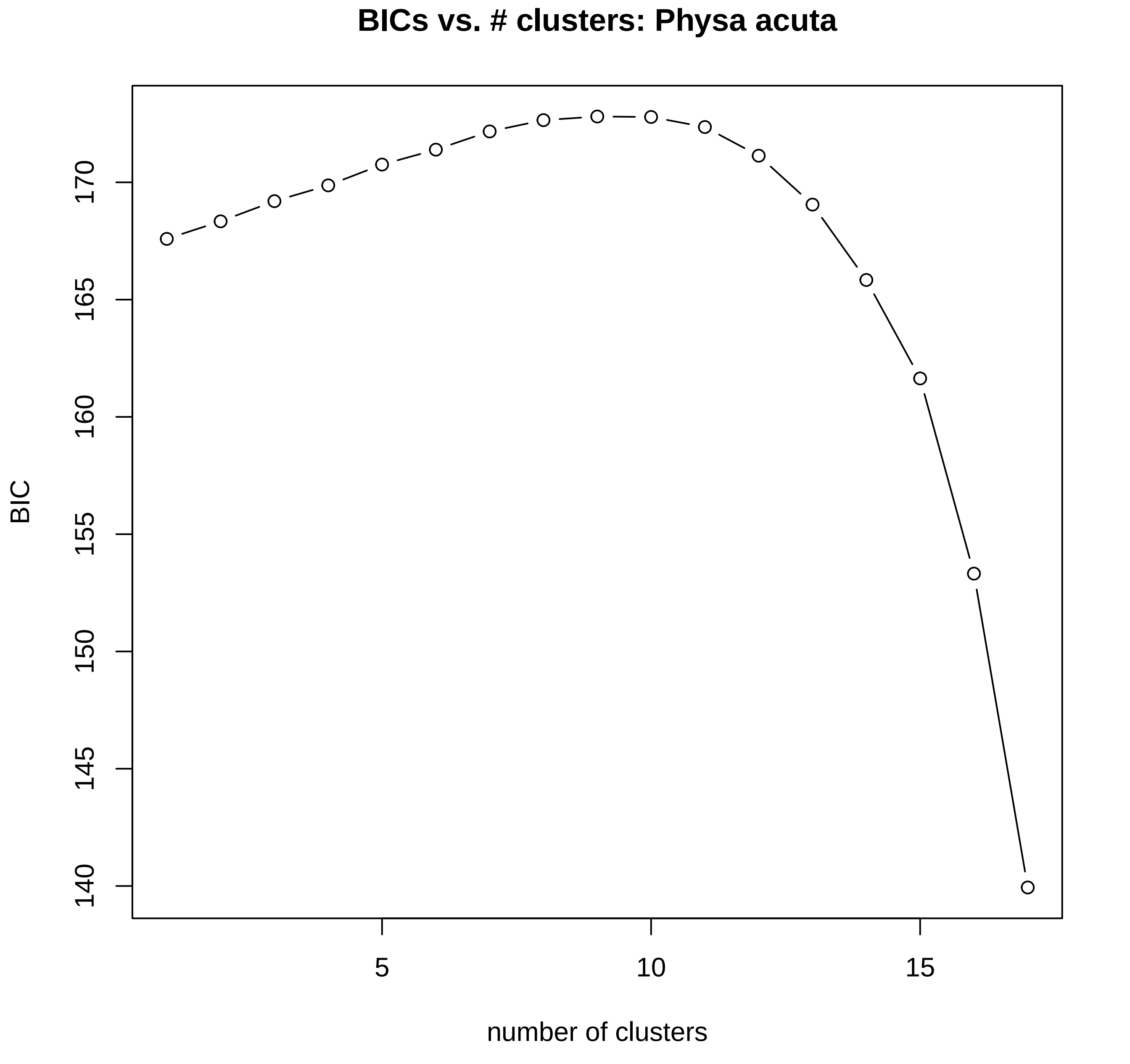
BICs vs. # clusters: Physa acuta
BIC
number of clusters
140
145
150
155
160
165
170
5
10
15

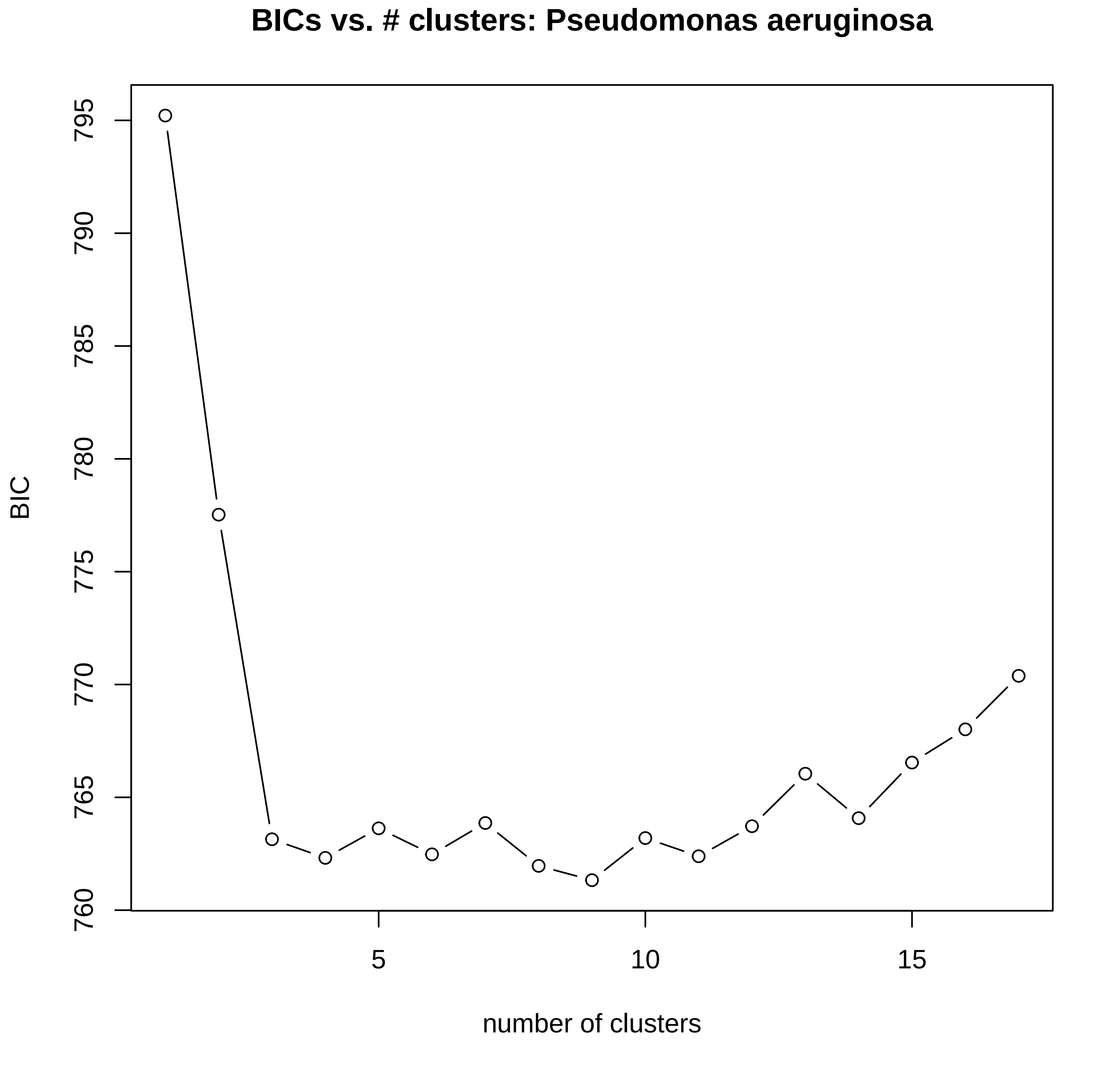
BICs vs. # clusters: Pseudomonas aeruginosa
BIC
795
790
785
780
775
770
765
760
5
10
15
number of clusters

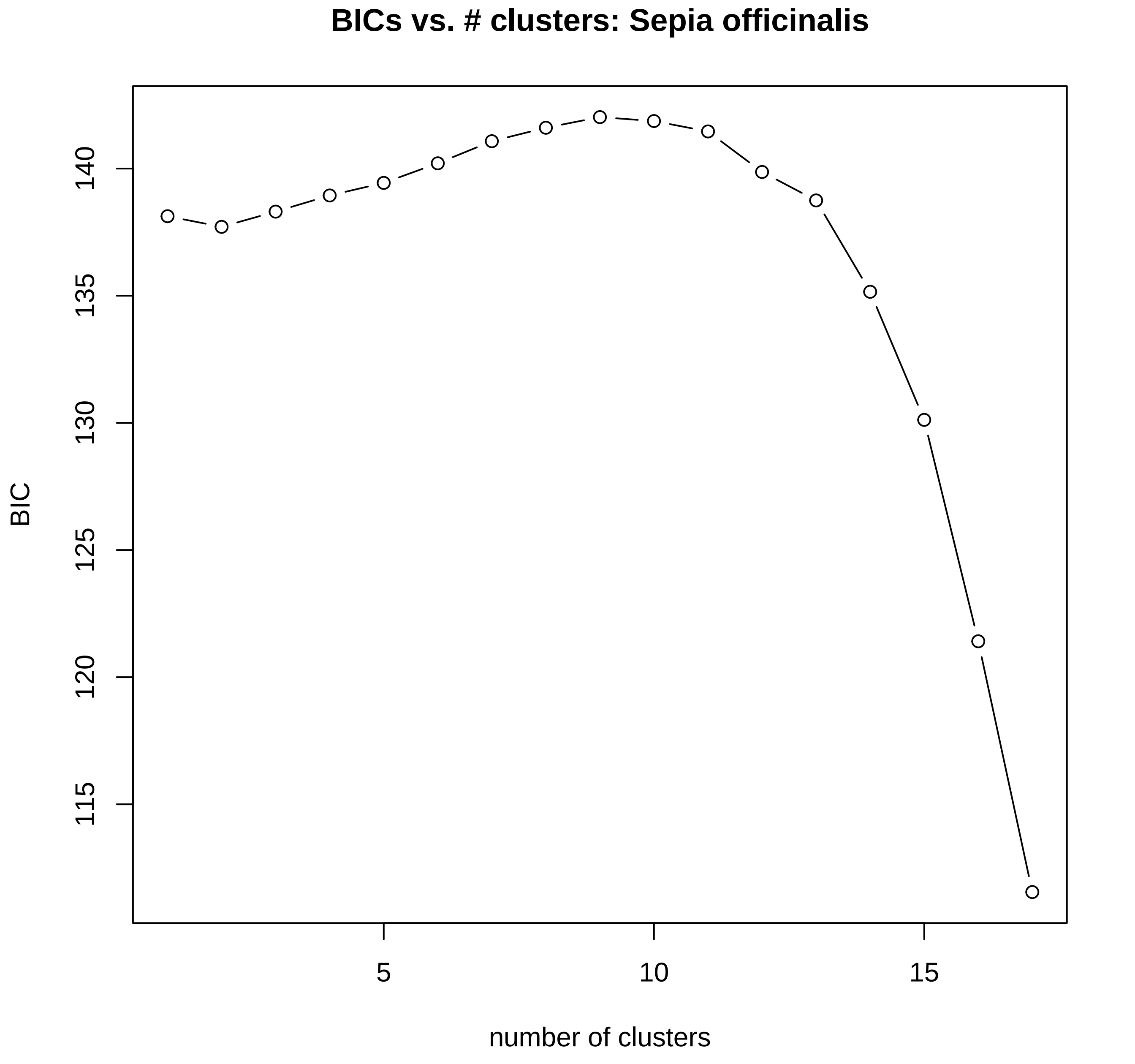

BICs vs. # clusters: Sepia officinalis
BIC
140
135
130
125
120
115
5
10
15
number of clusters

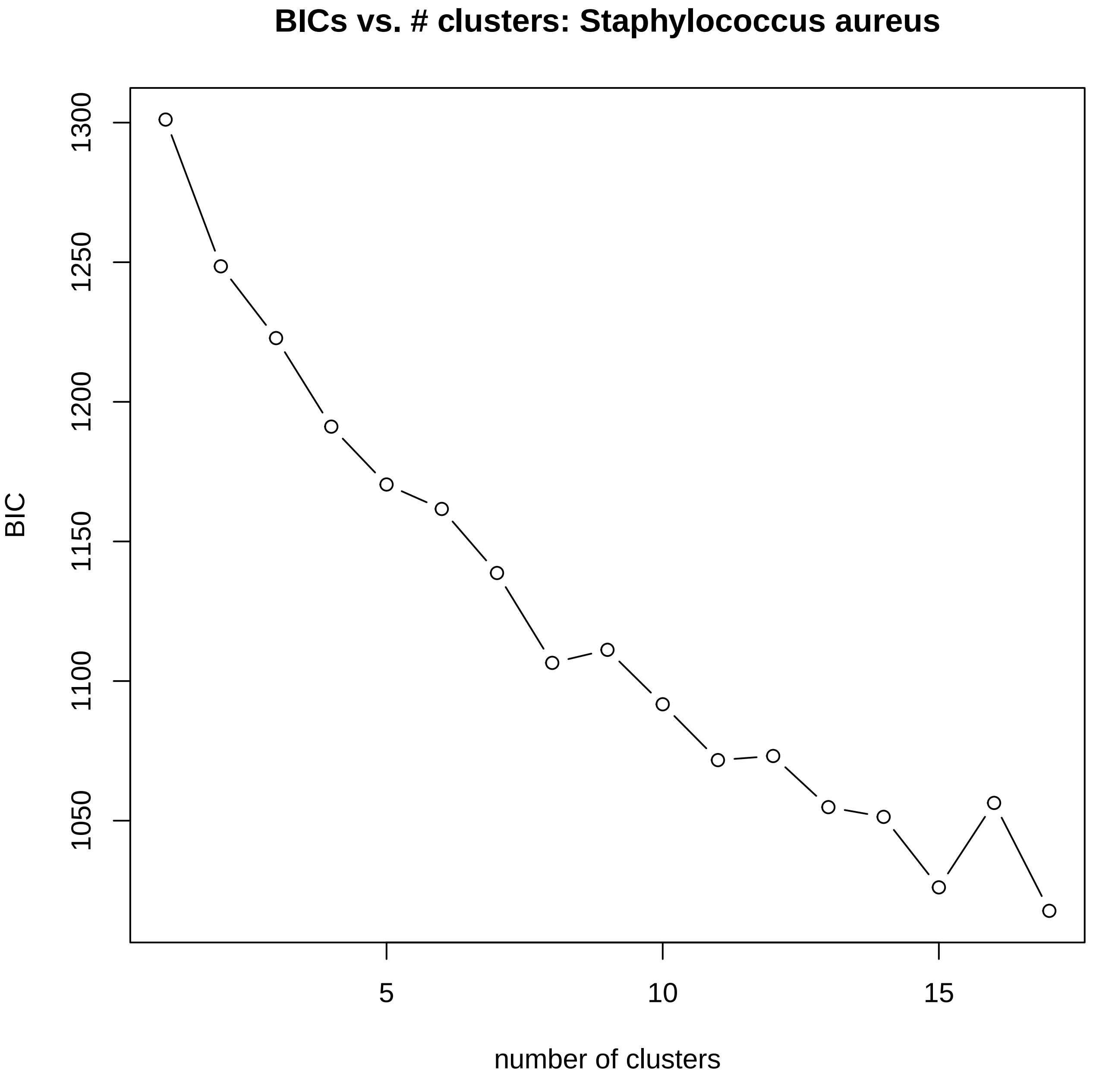
BICs vs. # clusters: Staphylococcus aureus
BIC
1300
1250
1200
1150
1100
1050
5
10
15
number of clusters

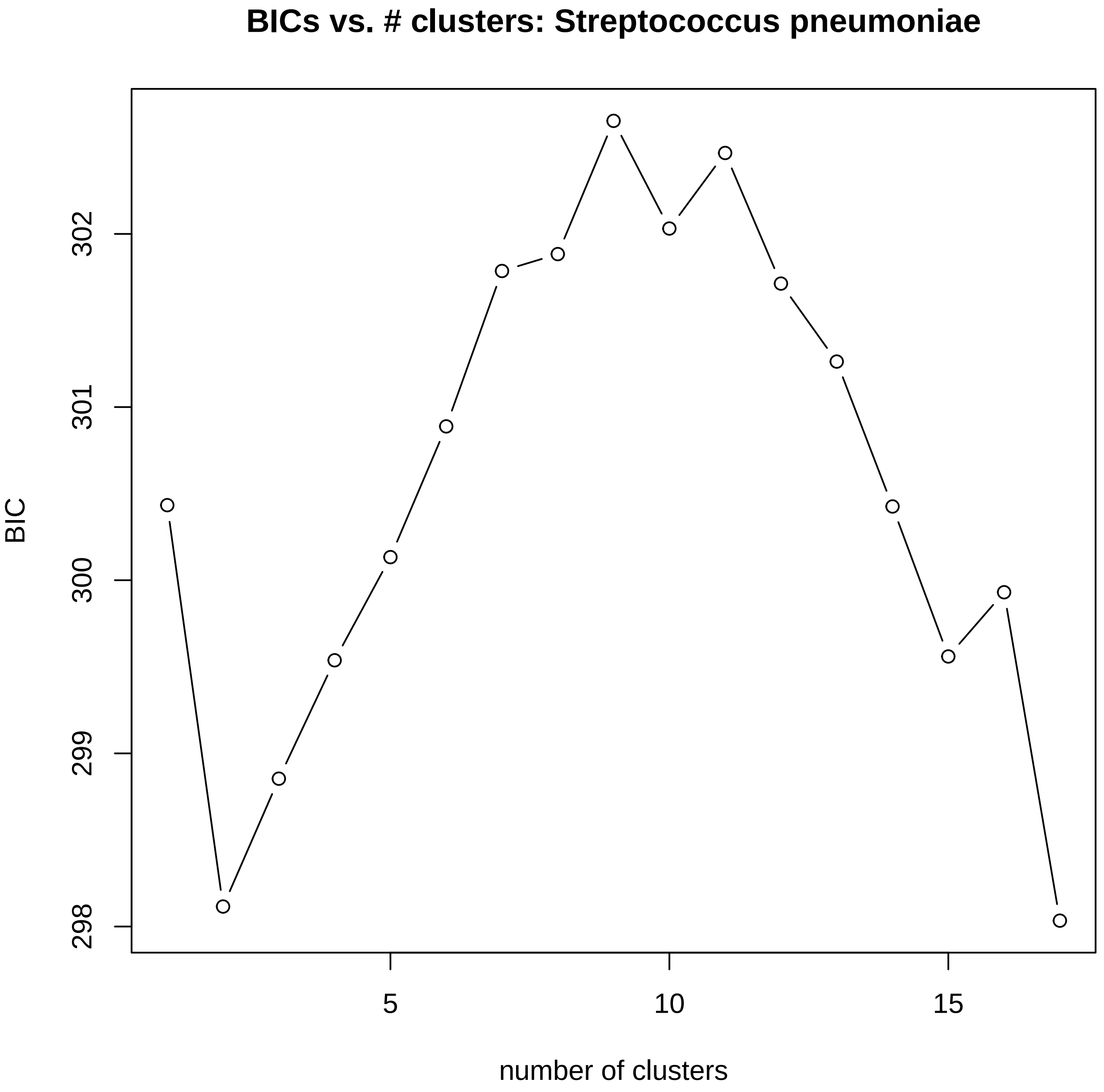
BICs vs. # clusters: Streptococcus pneumoniae
BIC
302
301
300
299
298
5
10
15
number of clusters

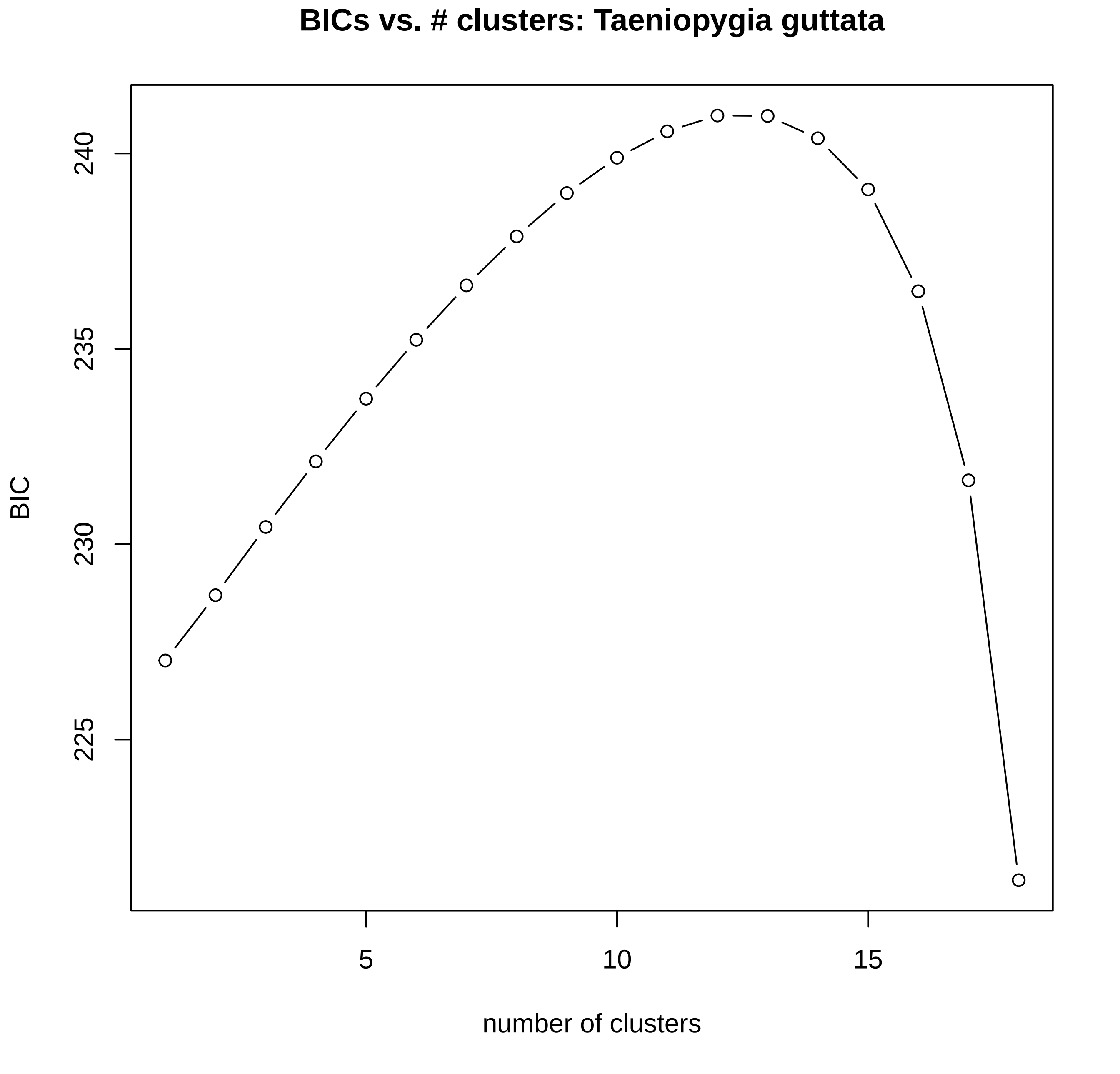
BICs vs. # clusters: Taeniopygia guttata
BIC
240
235
230
225
5
10
15
number of clusters

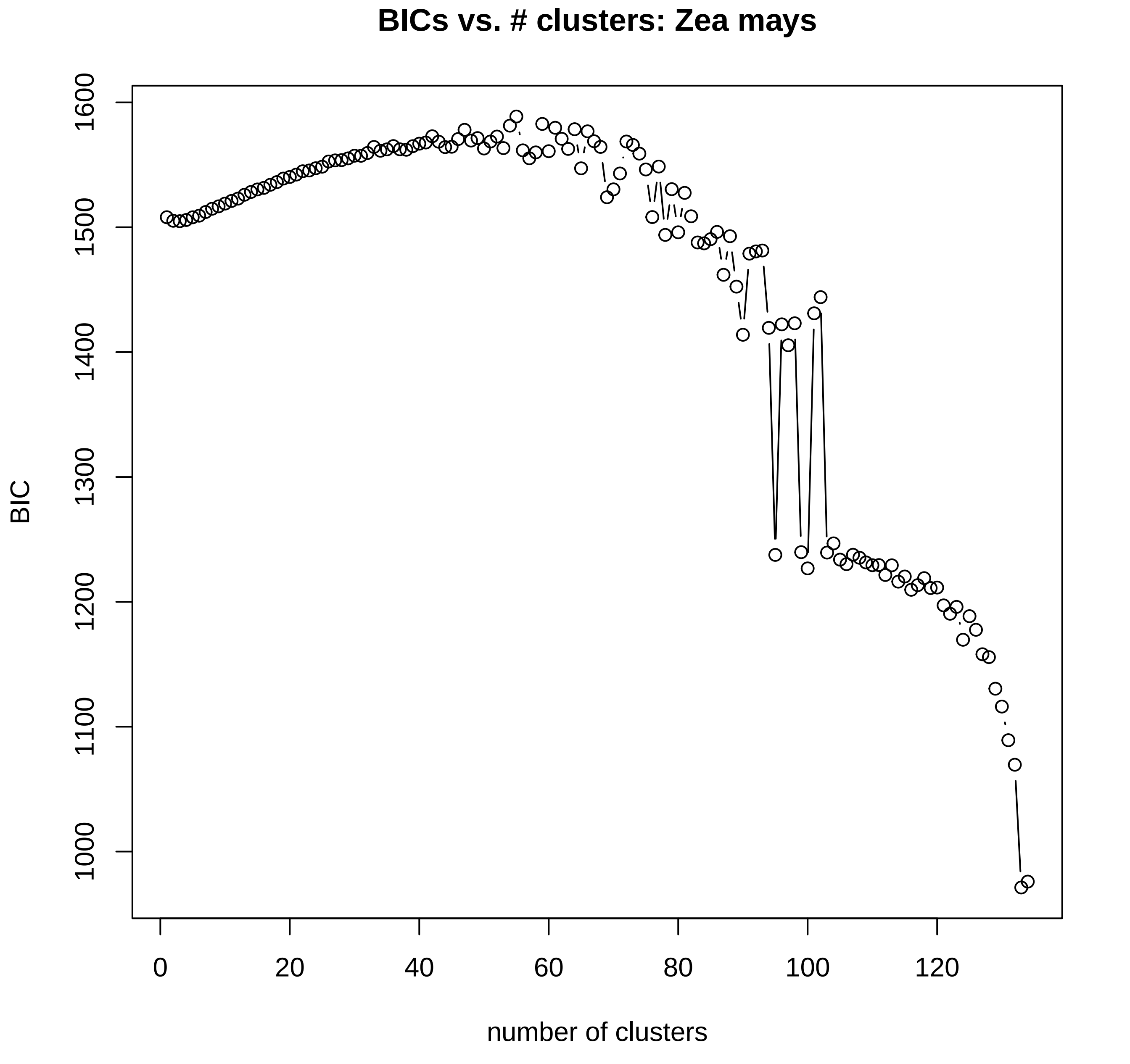

BICs vs. # clusters: Zea mays
BIC
1000
1100
1200
1300
1400
1500
1600
number of clusters
0
20
40
60
80
100
120
