## Supplementary 3 - PCA plots w. DAPC colours for "Interpreting the pervasive observation of U-shaped Site Frequency Spectra"

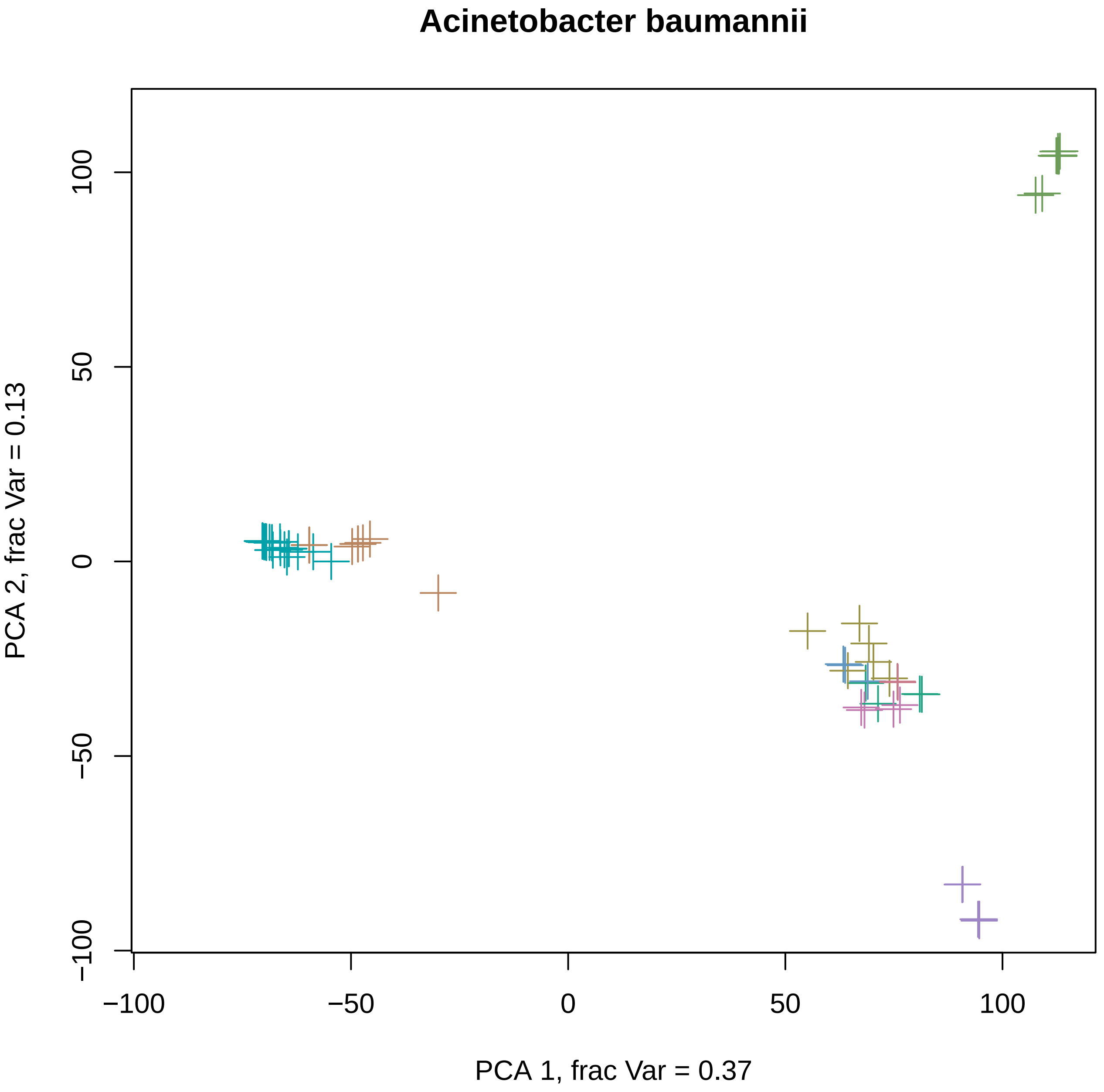
Acinetobacter baumannii
PCA 1, frac Var = 0.37
PCA 2, frac Var = 0.13
−100
−50
0
50
100

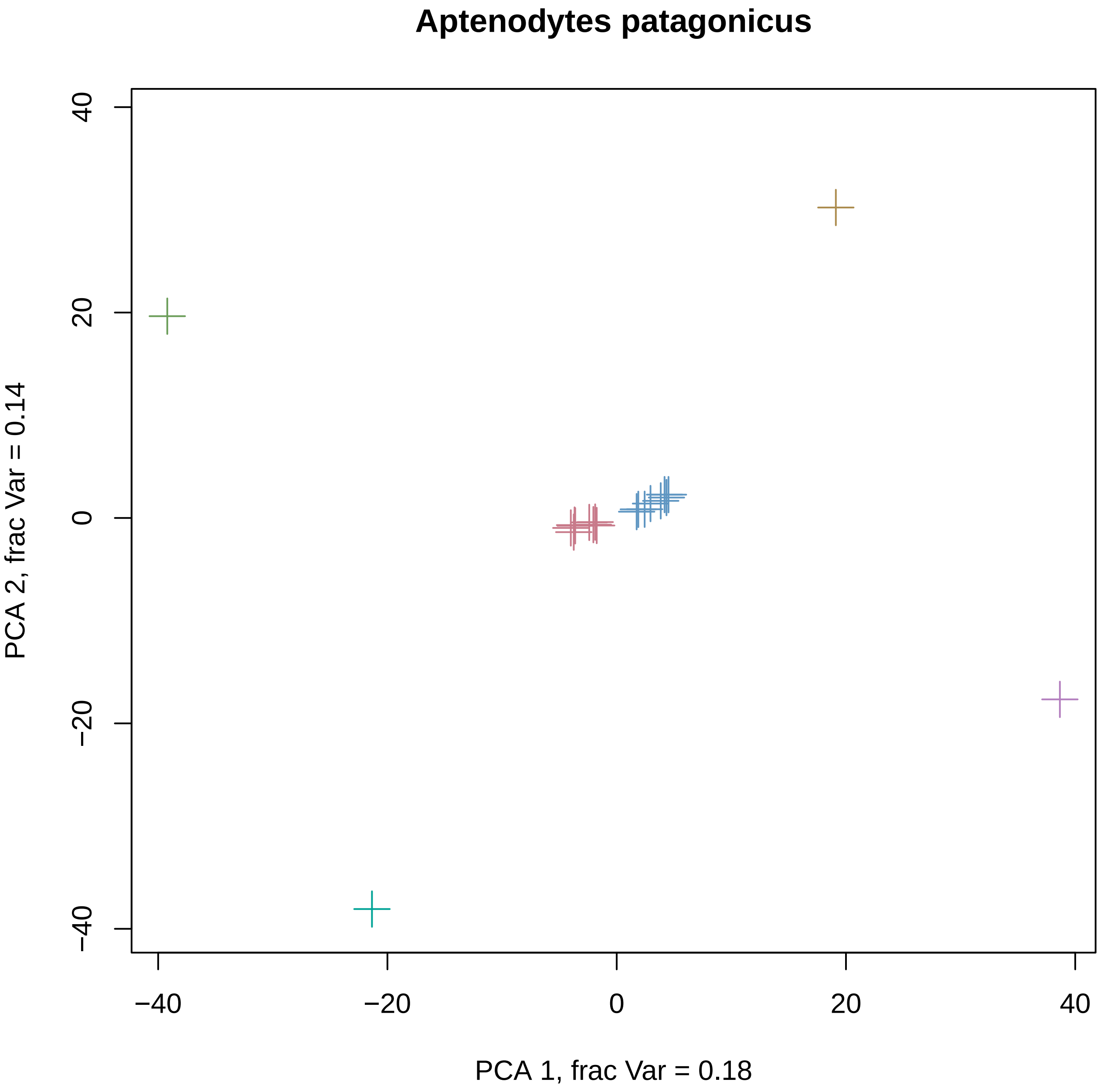

Aptenodytes patagonicus
PCA 1, frac Var = 0.18
PCA 2, frac Var = 0.14
−40
−20
0
20
40

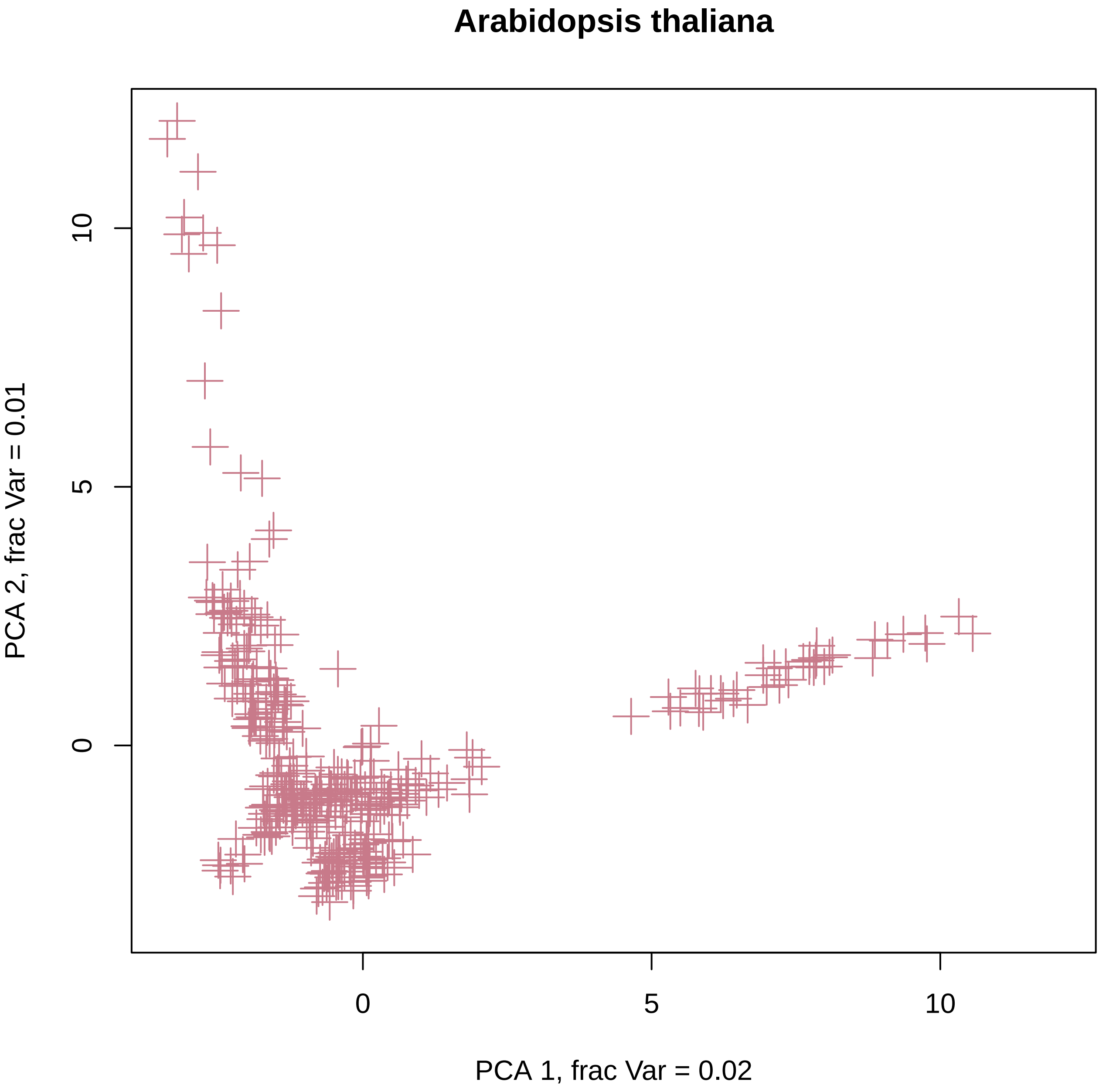
Arabidopsis thaliana
PCA 2, frac Var = 0.01
PCA 1, frac Var = 0.02
0
5
10
0
5
10

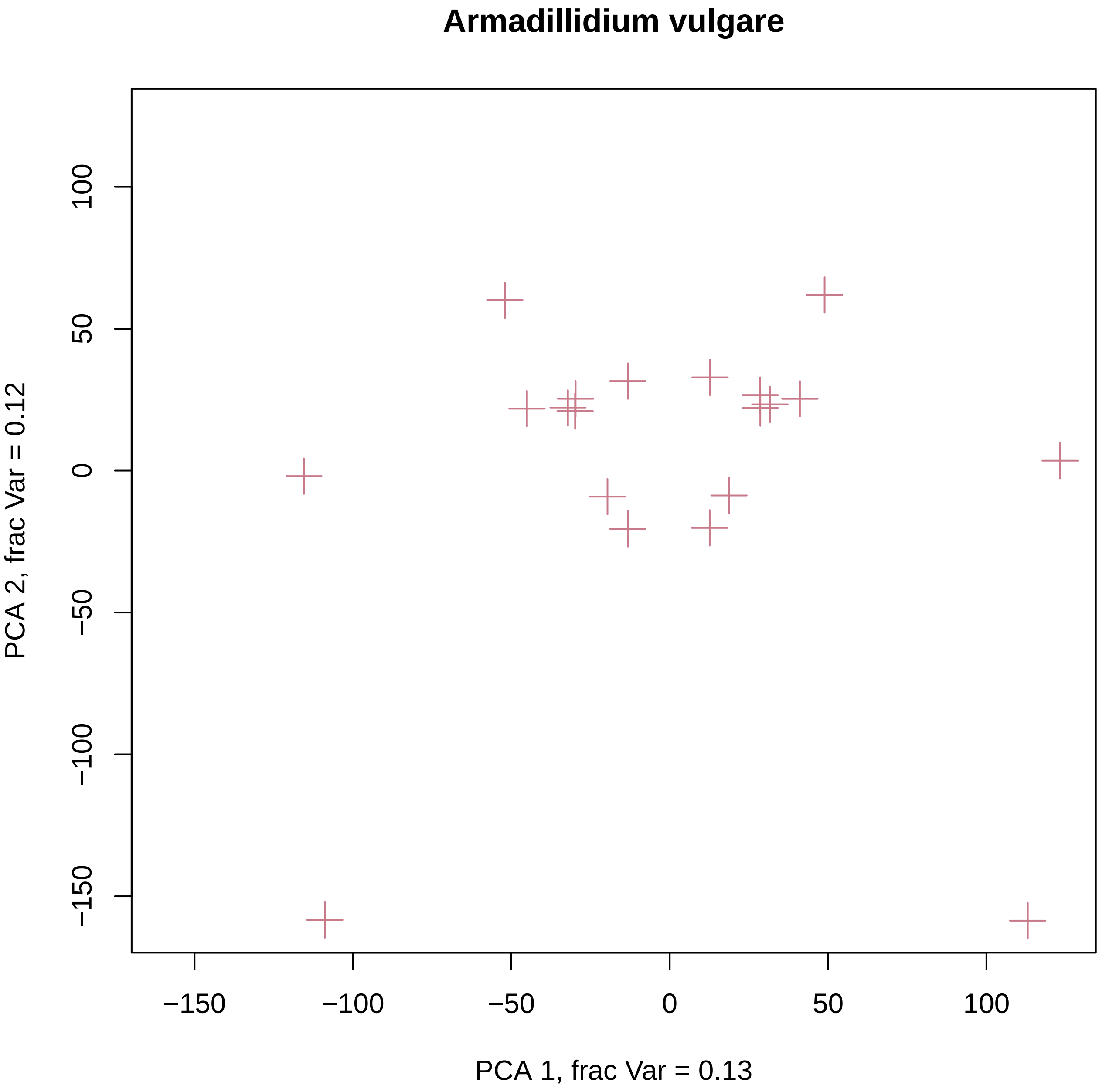
Armadillidium vulgare
PCA 1, frac Var = 0.13
PCA 2, frac Var = 0.12
−150
−100
−50
0
50
100

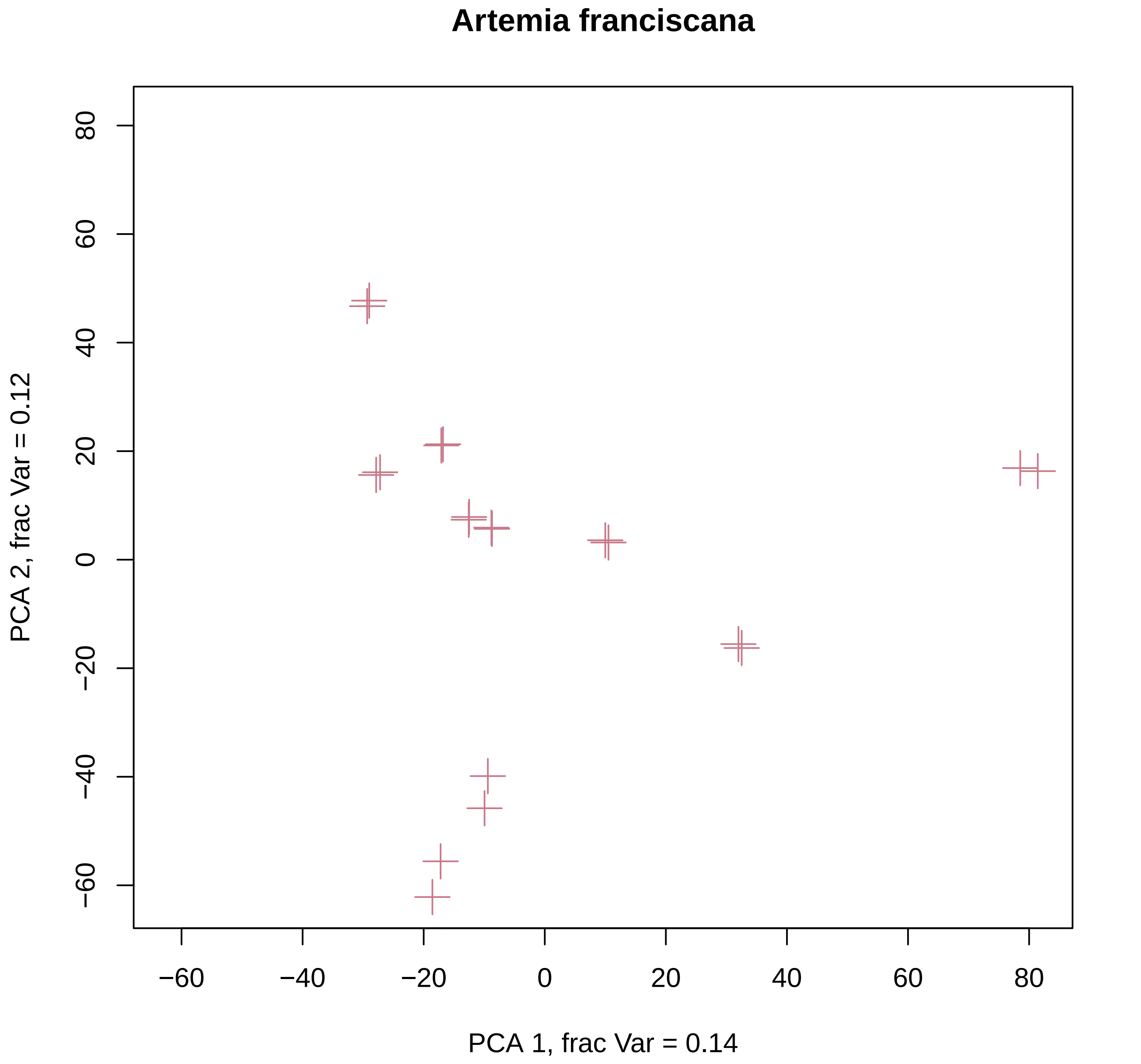

Artemia franciscana
PCA 1, frac Var = 0.14
PCA 2, frac Var = 0.12
−60
−40
−20
0
20
40
60
80

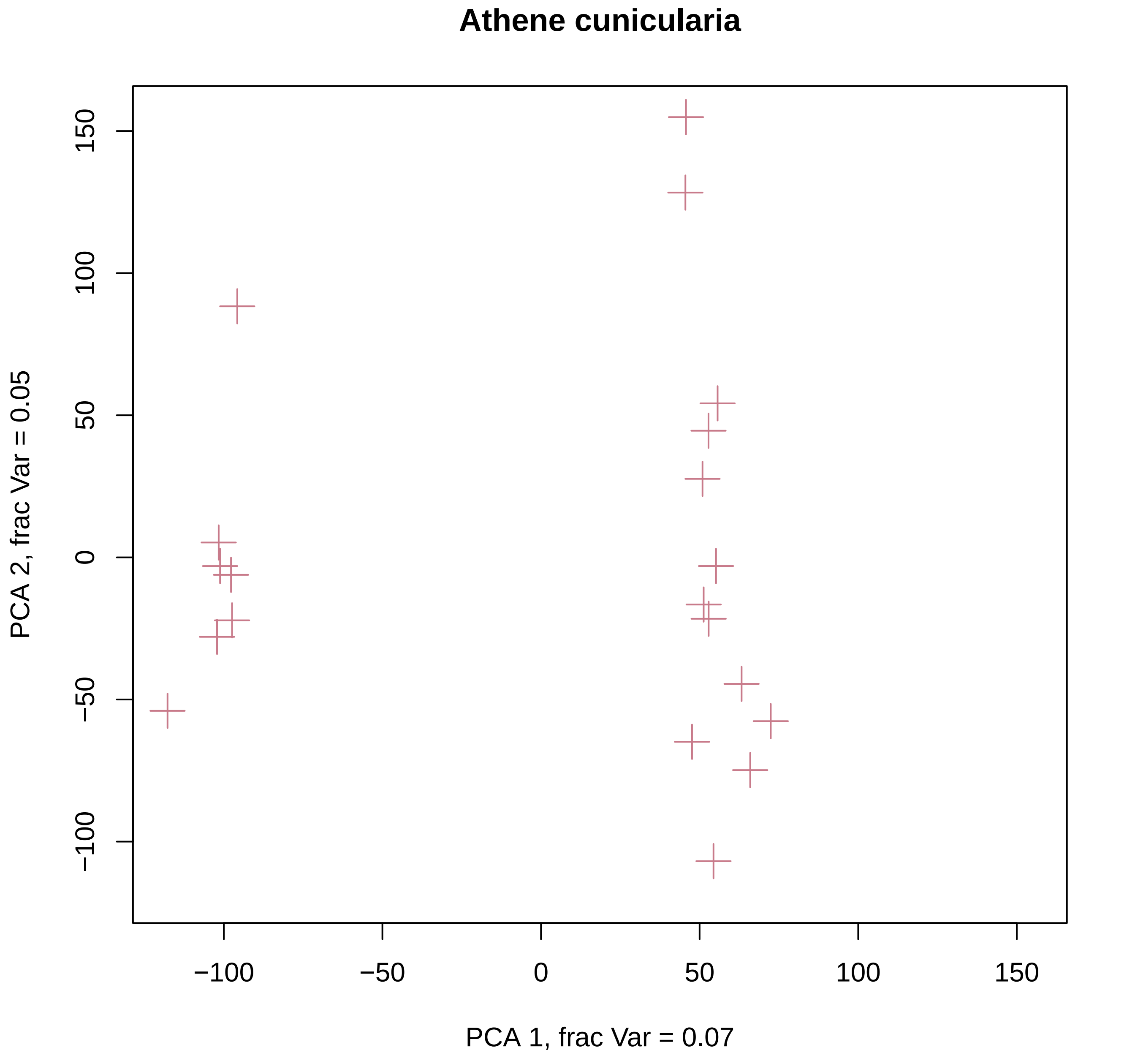
Athene cunicularia
PCA 2, frac Var = 0.05
150
100
50
0
−50
−100
−100
−50
0
50
100
150
PCA 1, frac Var = 0.07

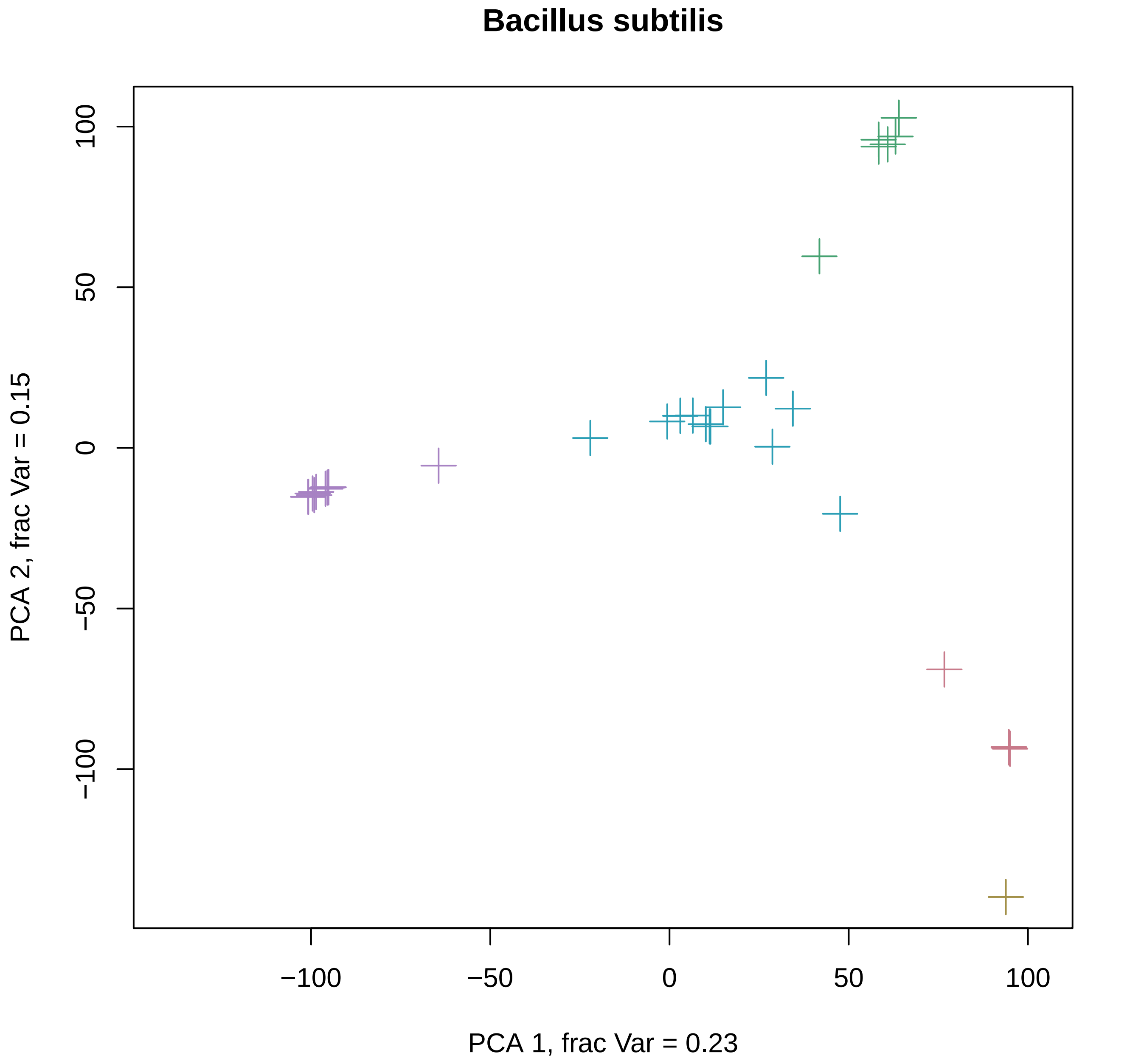

Bacillus subtilis
100
50
0
−50
−100
PCA 2, frac Var = 0.15
−100
−50
0
50
100
PCA 1, frac Var = 0.23

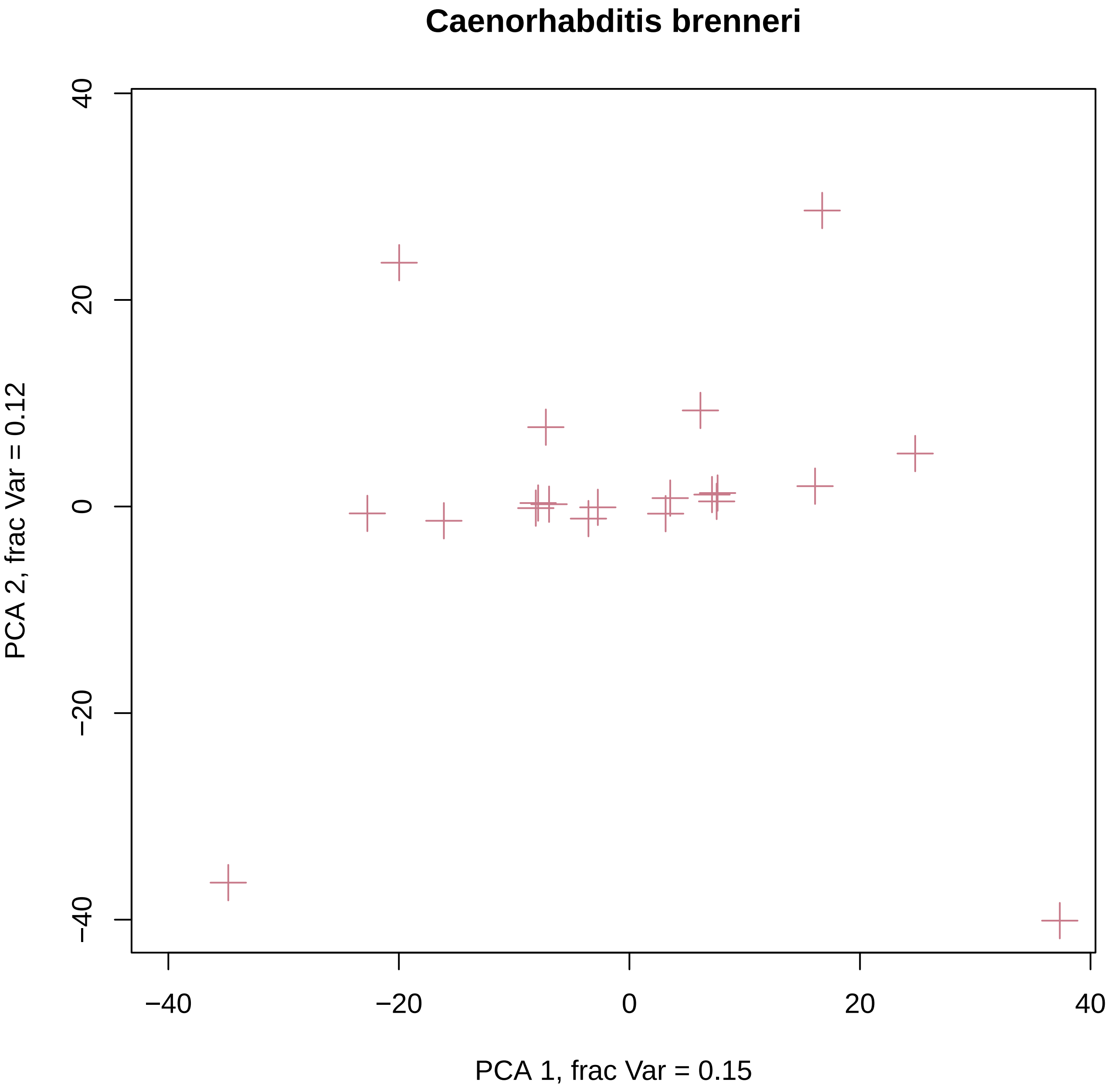
Caenorhabditis brenneri
PCA 2, frac Var = 0.12
PCA 1, frac Var = 0.15
−40
−20
0
20
40
−40
−20
0
20
40

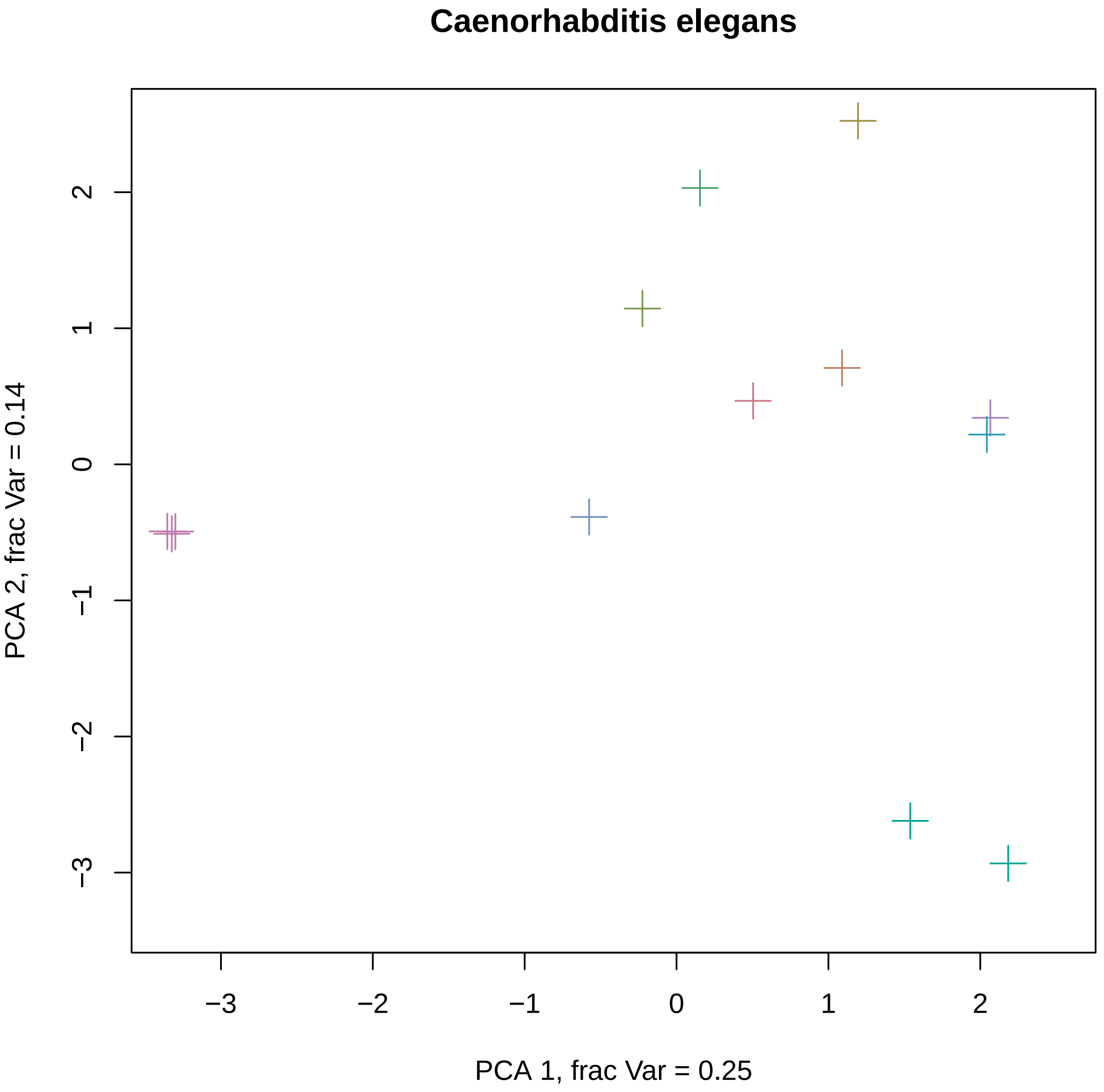

Caenorhabditis elegans
PCA 1, frac Var = 0.25
PCA 2, frac Var = 0.14
−3
−2
−1
0
1
2

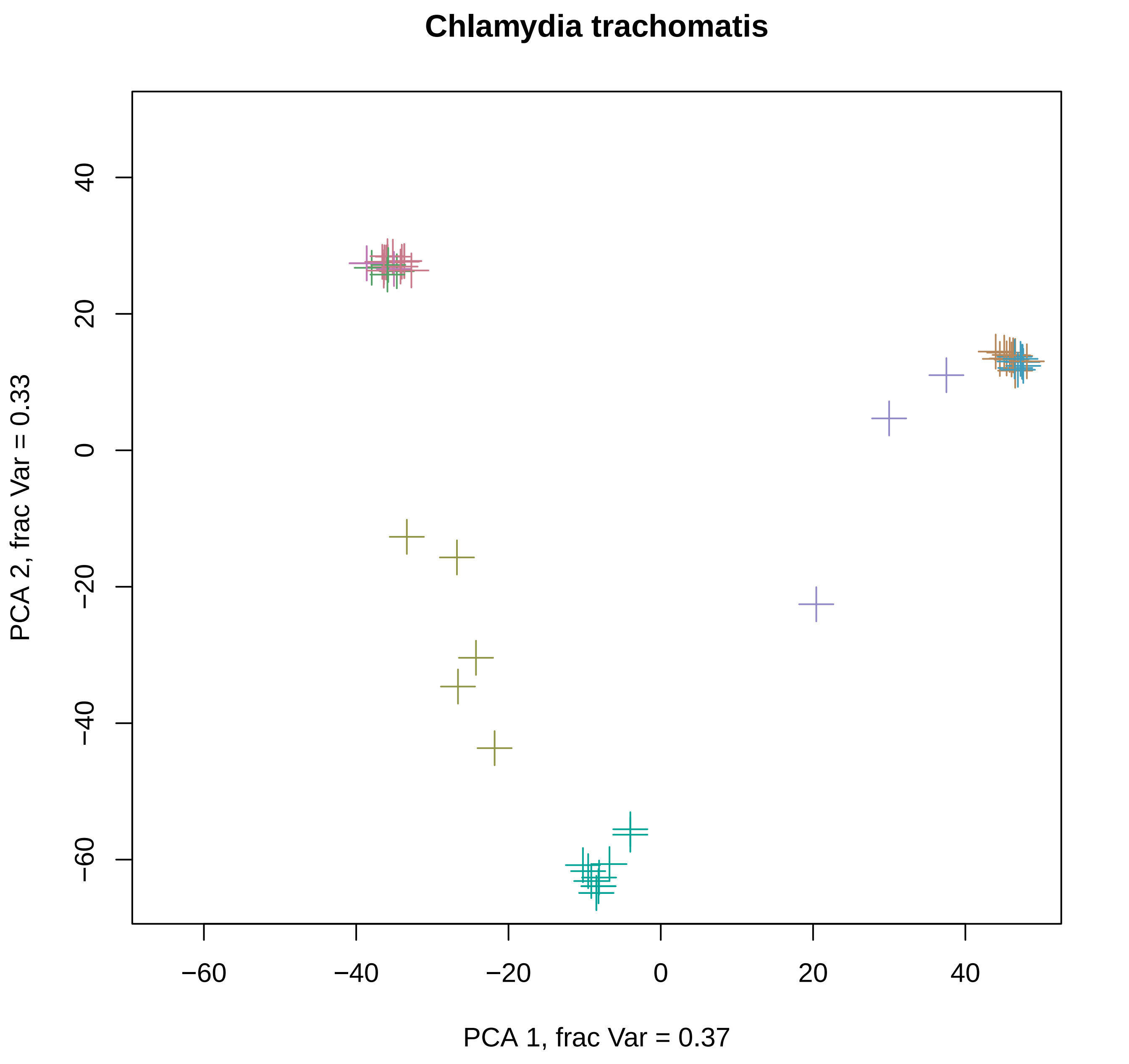

Chlamydia trachomatis
PCA 2, frac Var = 0.33
PCA 1, frac Var = 0.37
40
20
0
−20
−40
−60
−60
−40
−20
0
20
40

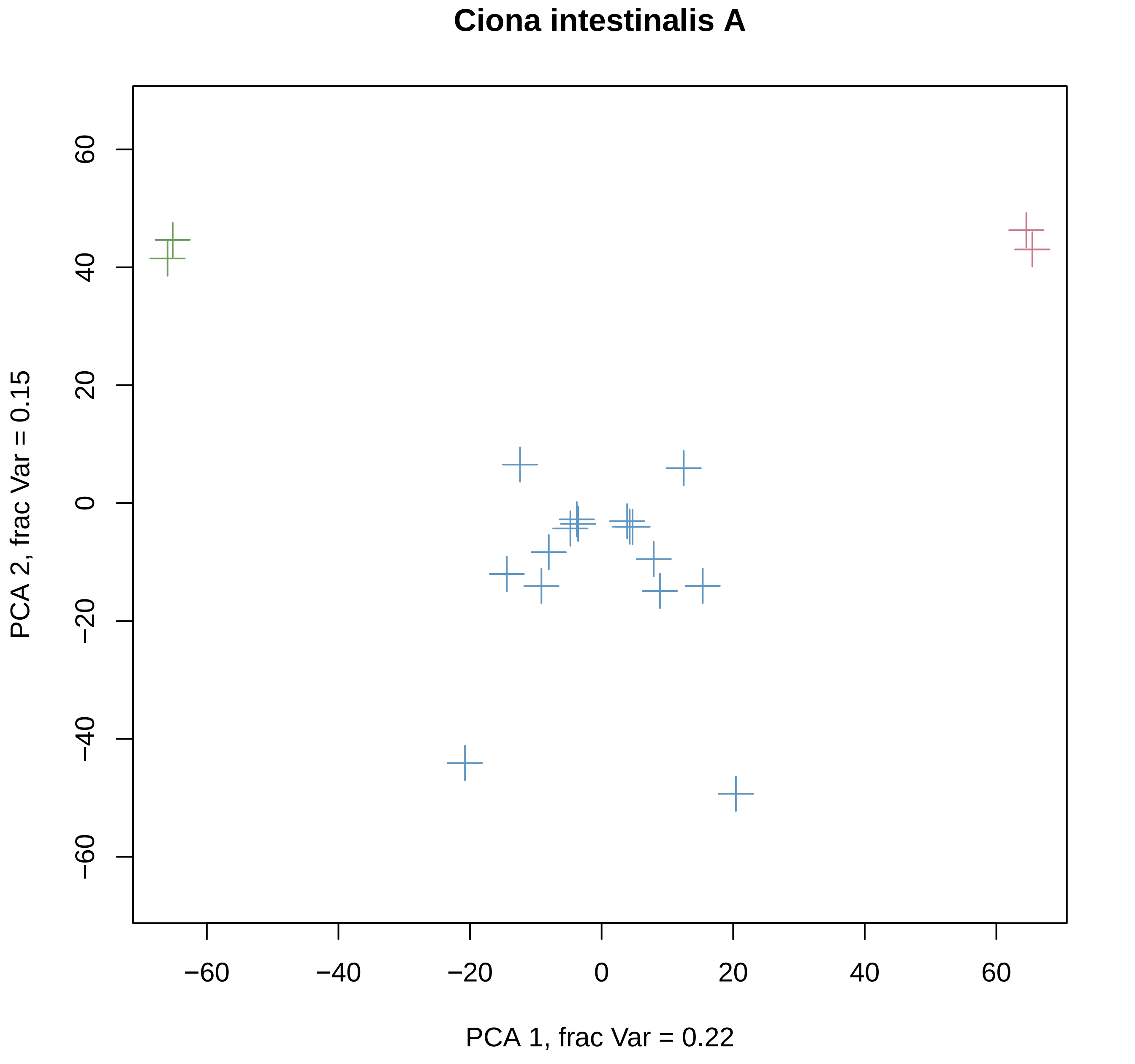
Ciona intestinalis A
PCA 1, frac Var = 0.22
PCA 2, frac Var = 0.15
−60
−40
−20
0
20
40
60

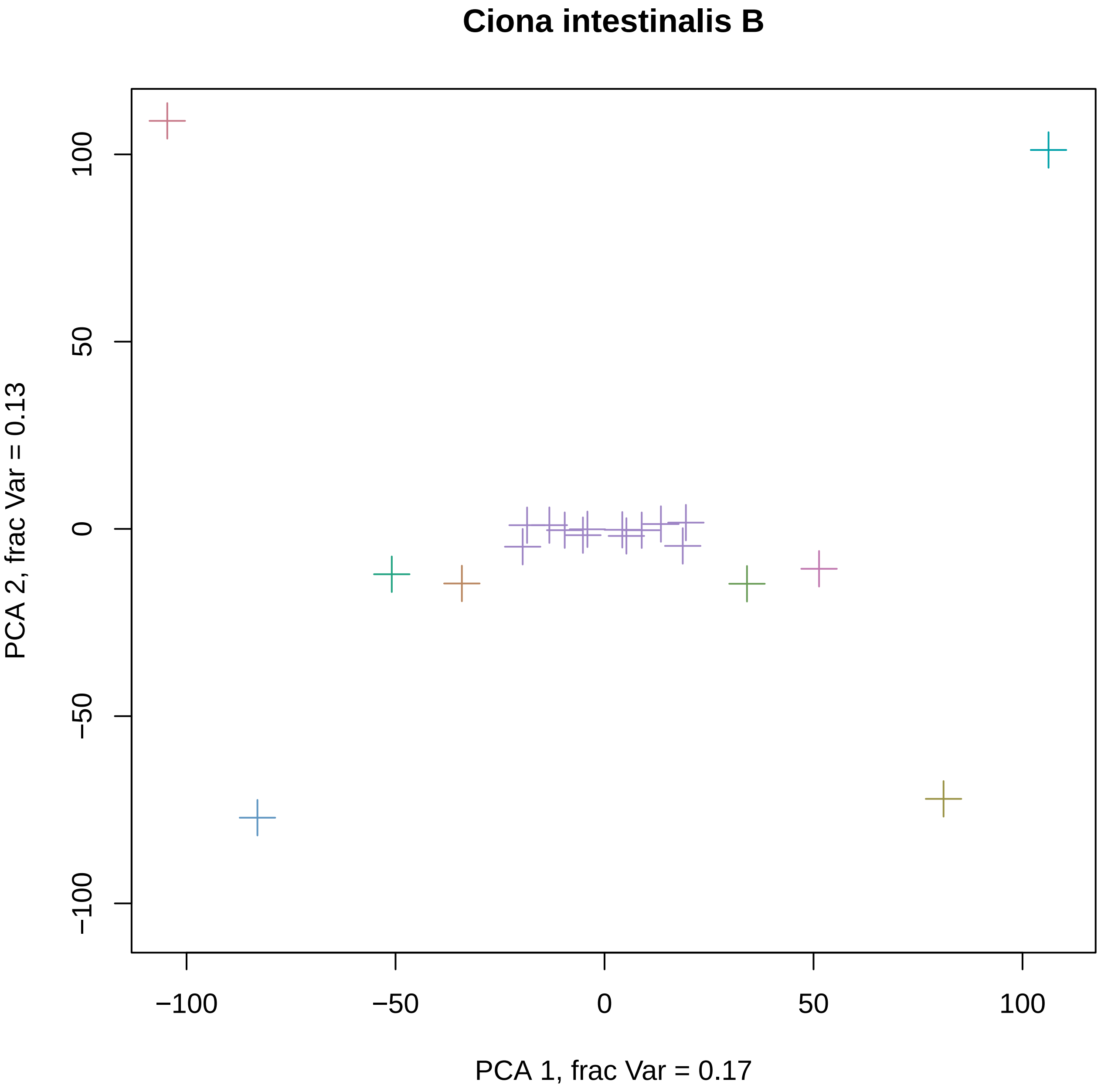
Ciona intestinalis B
PCA 1, frac Var = 0.17
PCA 2, frac Var = 0.13
−100
−50
0
50
100
−100
−50
0
50
100

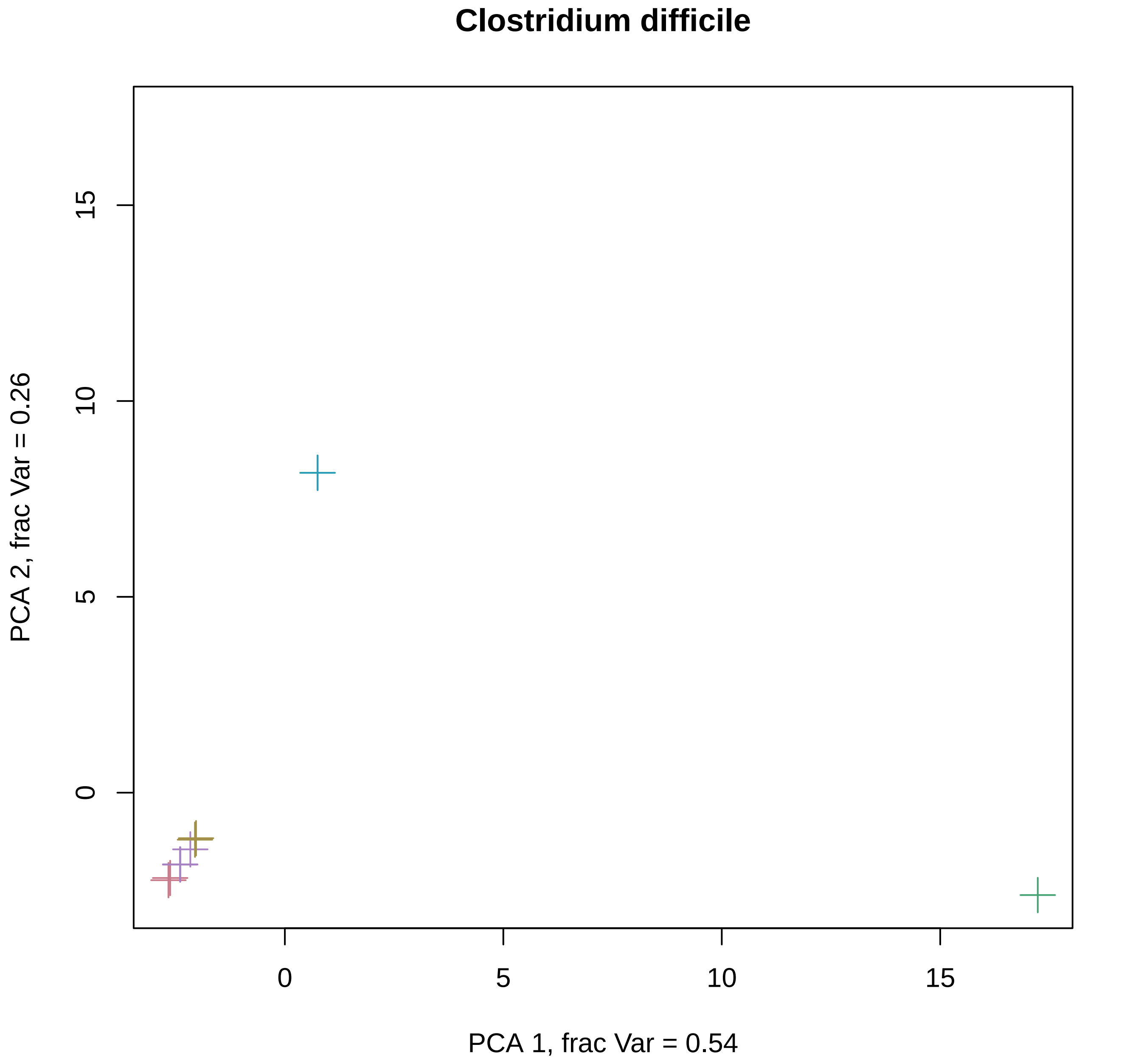

Clostridium difficile
PCA 2, frac Var = 0.26
15
10
5
0
0
5
10
15
PCA 1, frac Var = 0.54

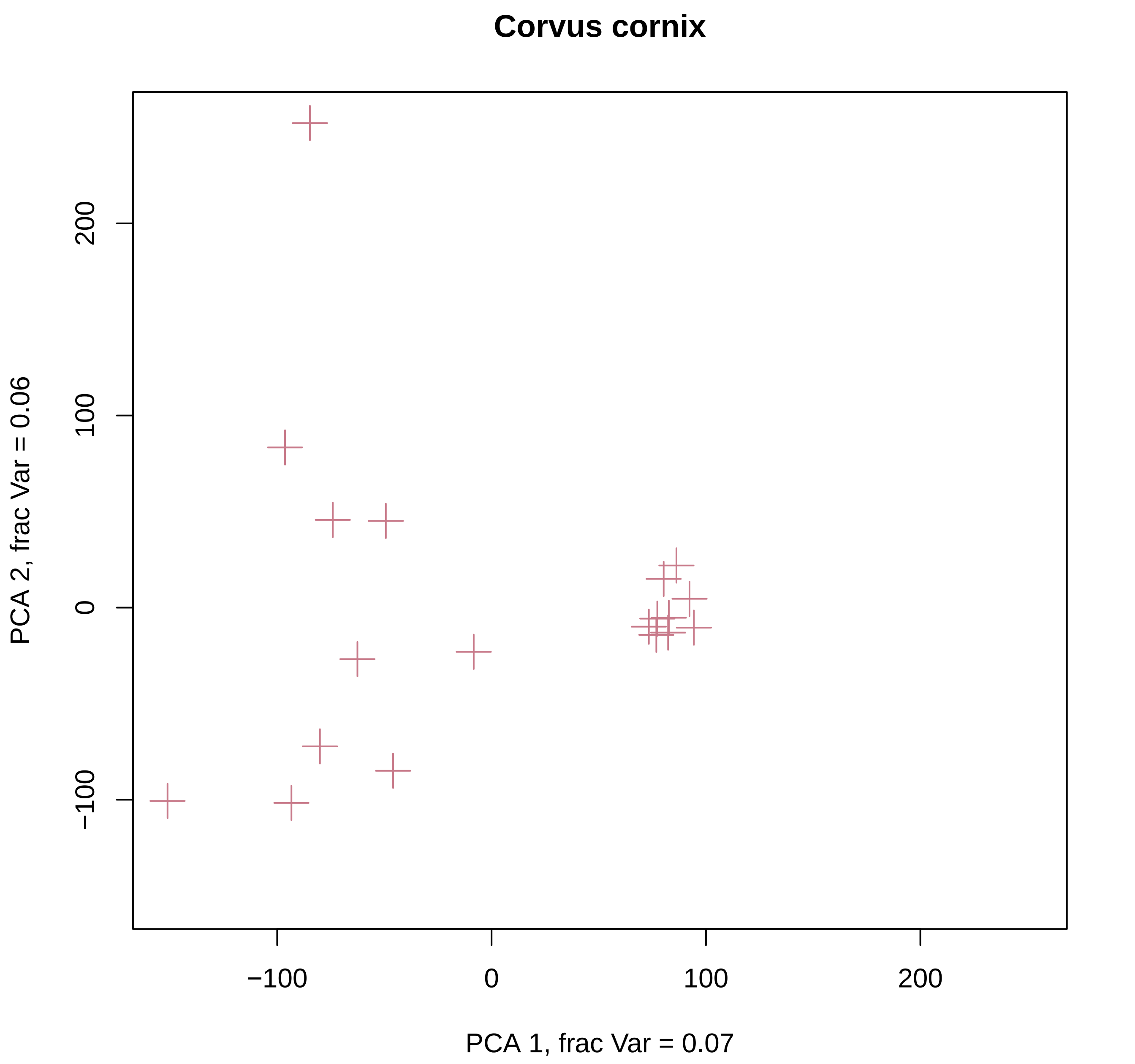

Corvus cornix
200
100
0
−100
PCA 2, frac Var = 0.06
−100
0
100
200
PCA 1, frac Var = 0.07

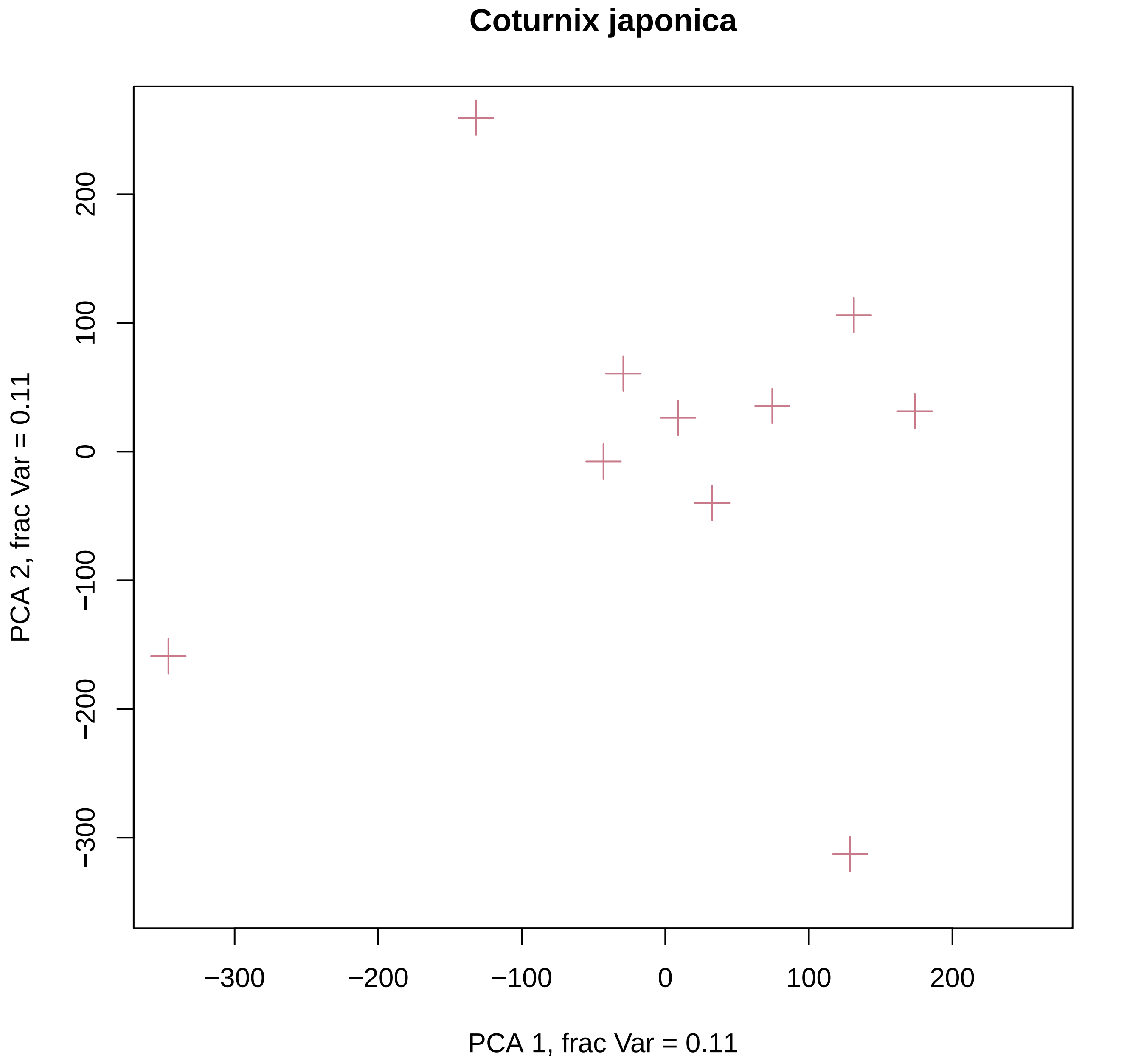

Coturnix japonica
PCA 1, frac Var = 0.11
PCA 2, frac Var = 0.11
−300
−200
−100
0
100
200

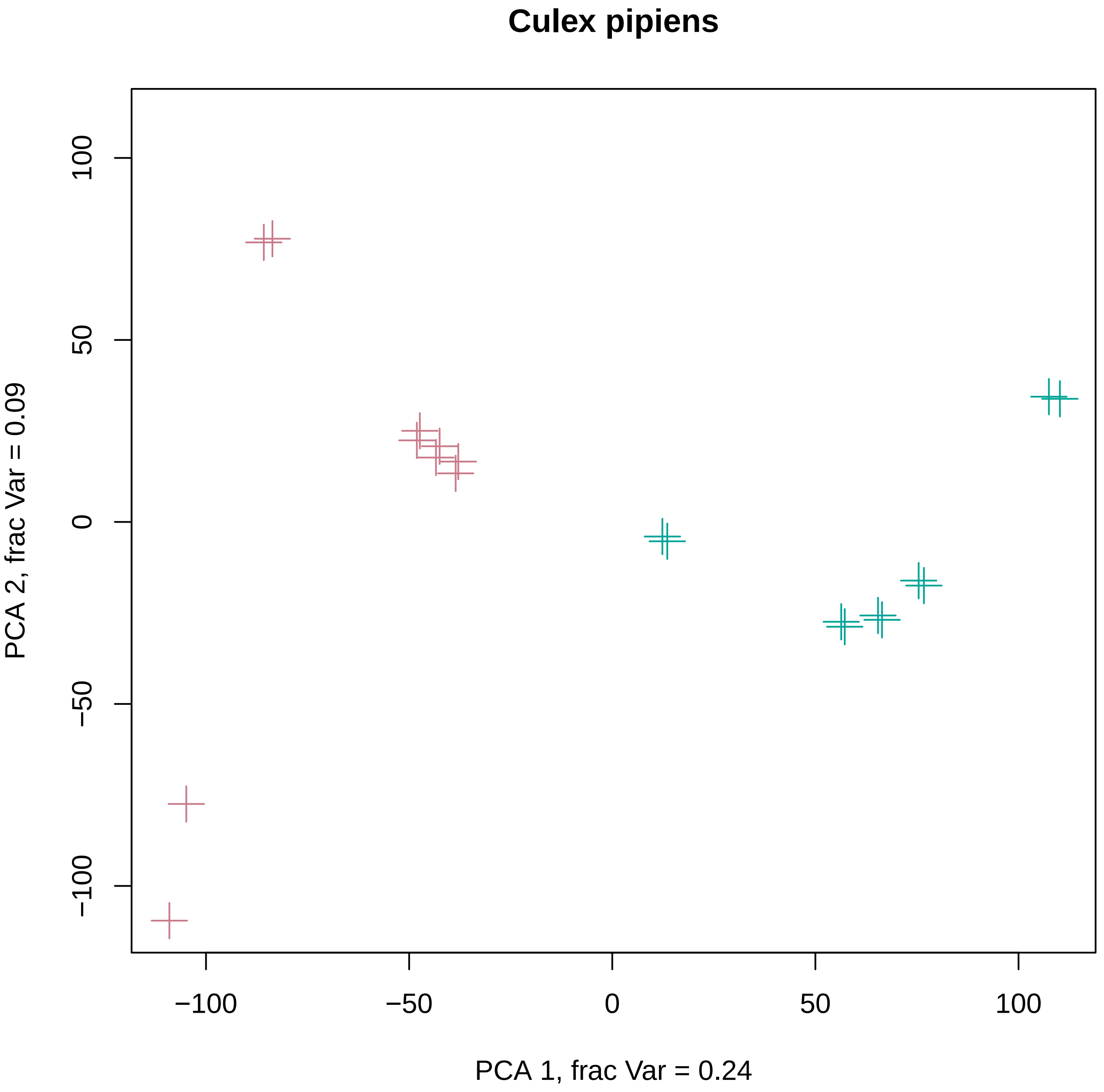

Culex pipiens
PCA 1, frac Var = 0.24
PCA 2, frac Var = 0.09
−100
−50
0
50
100

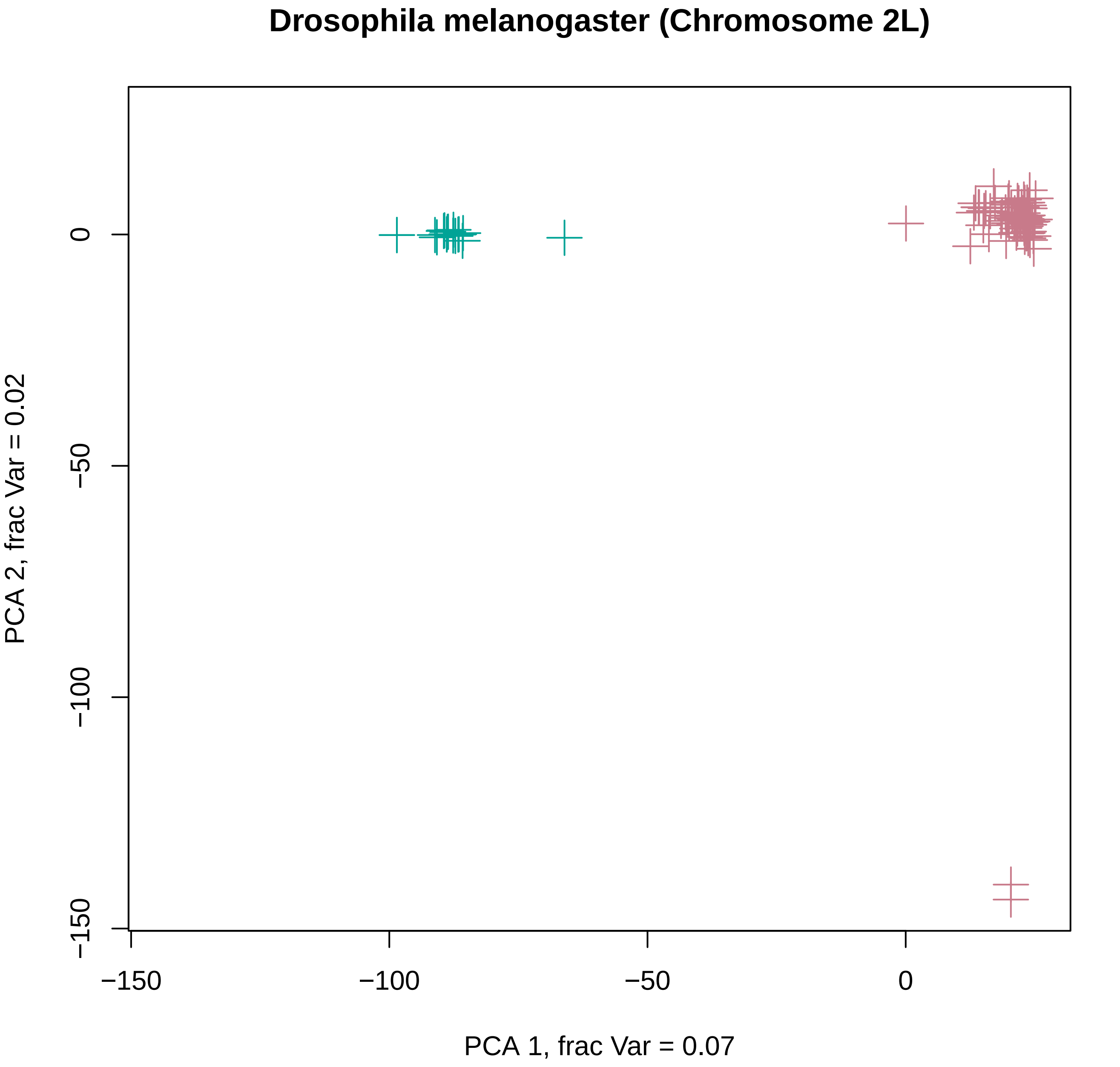

Drosophila melanogaster (Chromosome 2L)
0
−50
−100
−150
−150
−100
−50
0
PCA 1, frac Var = 0.07
PCA 2, frac Var = 0.02

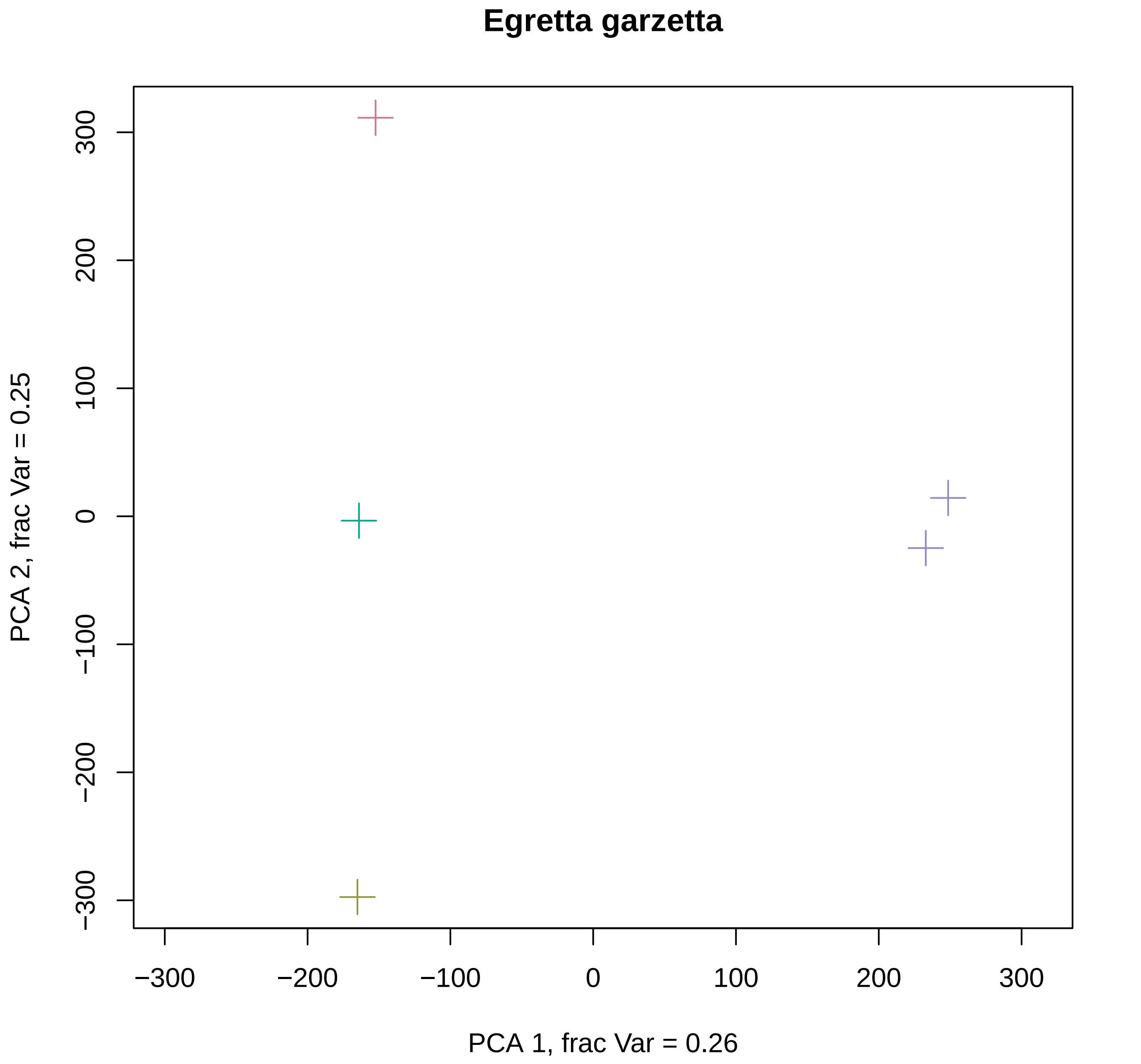

Egretta garzetta
300
200
100
0
−100
−200
−300
PCA 2, frac Var = 0.25
−300
−200
−100
0
100
200
300
PCA 1, frac Var = 0.26

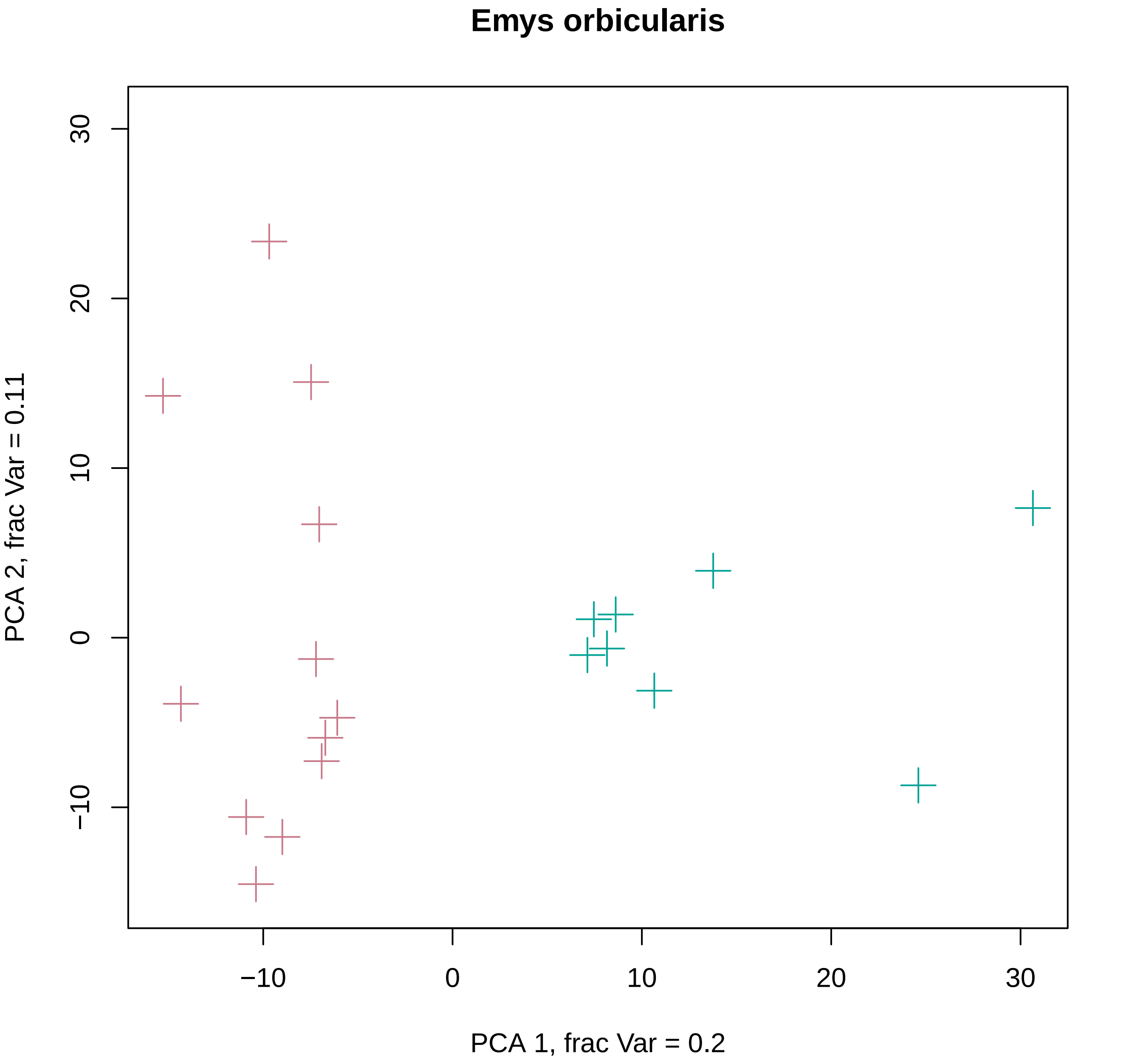
Emys orbicularis
PCA 1, frac Var = 0.2
PCA 2, frac Var = 0.11
−10
0
10
20
30
−10
0
10
20
30

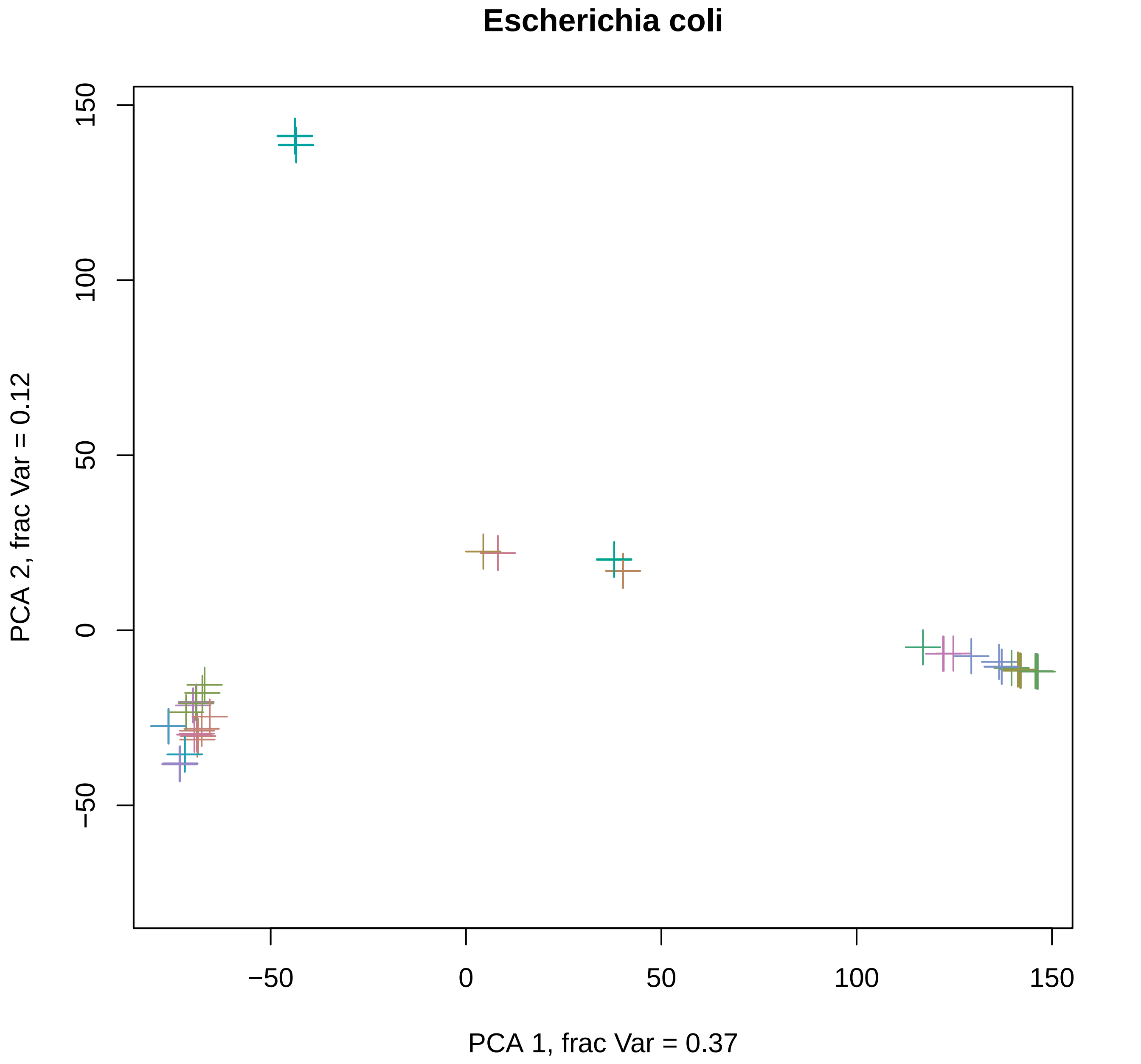

Escherichia coli
PCA 1, frac Var = 0.37
PCA 2, frac Var = 0.12
−50
0
50
100
150
−50
0
50
100
150

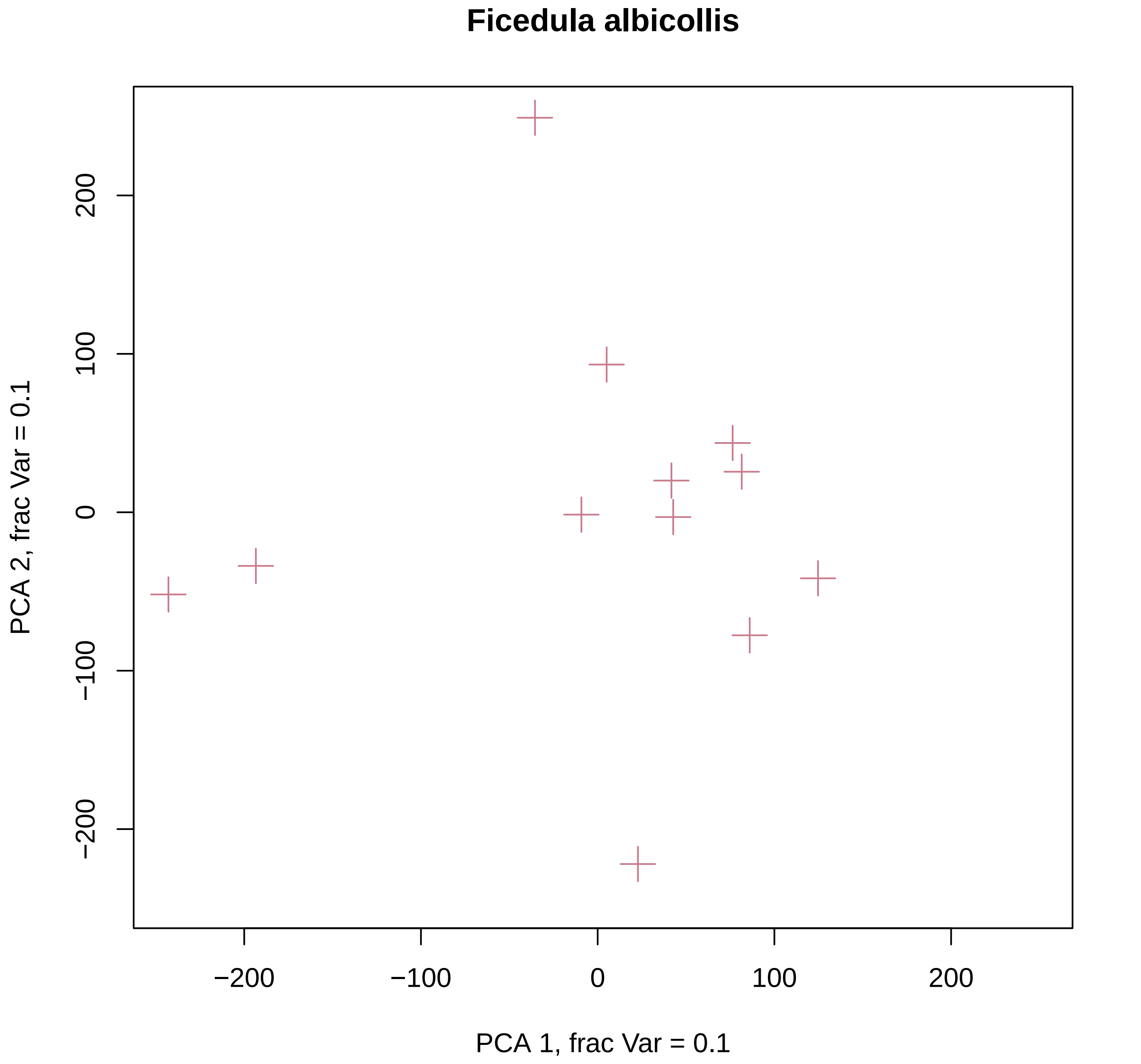
Ficedula albicollis
PCA 1, frac Var = 0.1
PCA 2, frac Var = 0.1
−200
−100
0
100
200
−200
−100
0
100
200

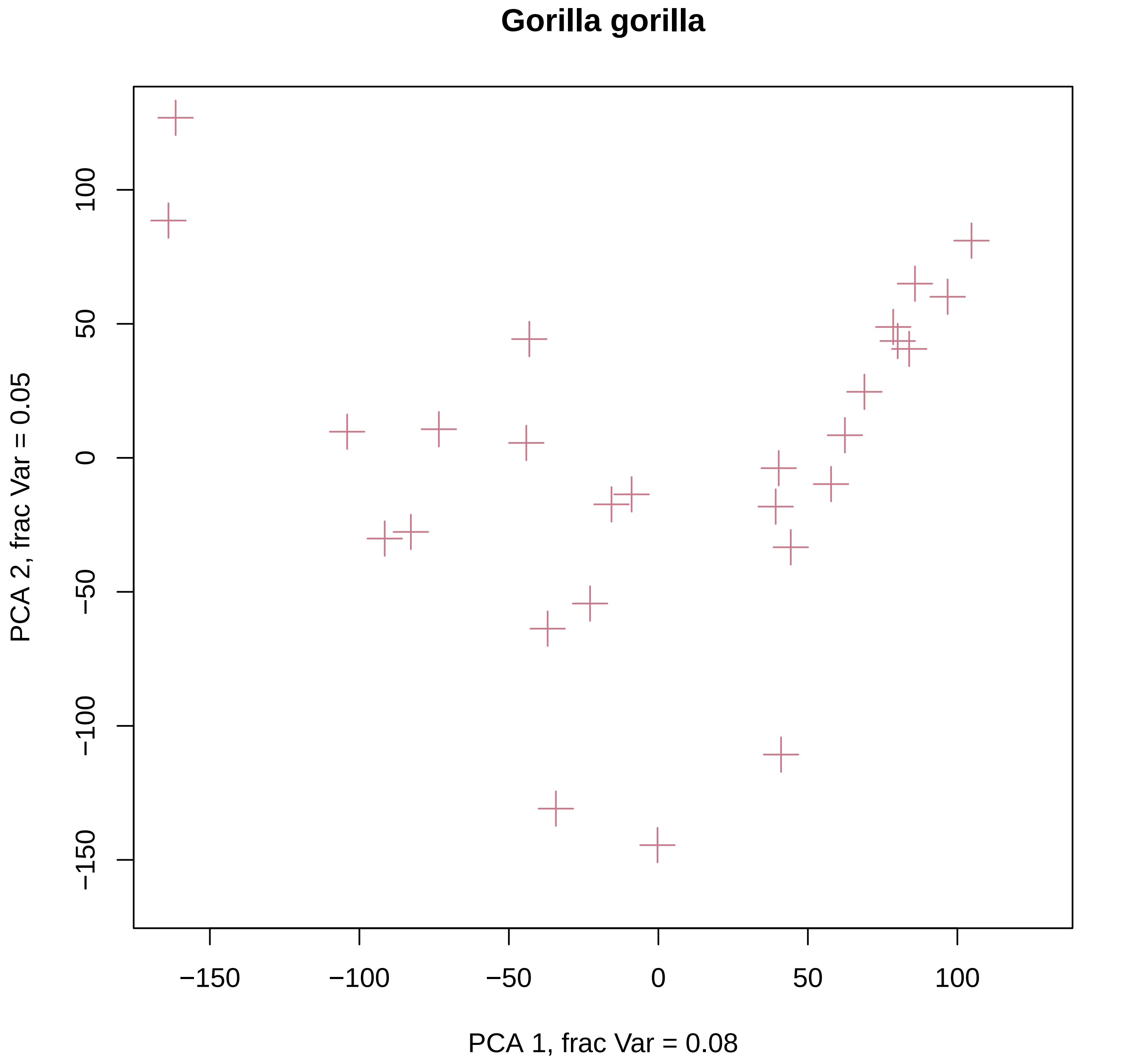

Gorilla gorilla
100
50
0
−50
−100
−150
PCA 2, frac Var = 0.05
−150
−100
−50
0
50
100
PCA 1, frac Var = 0.08

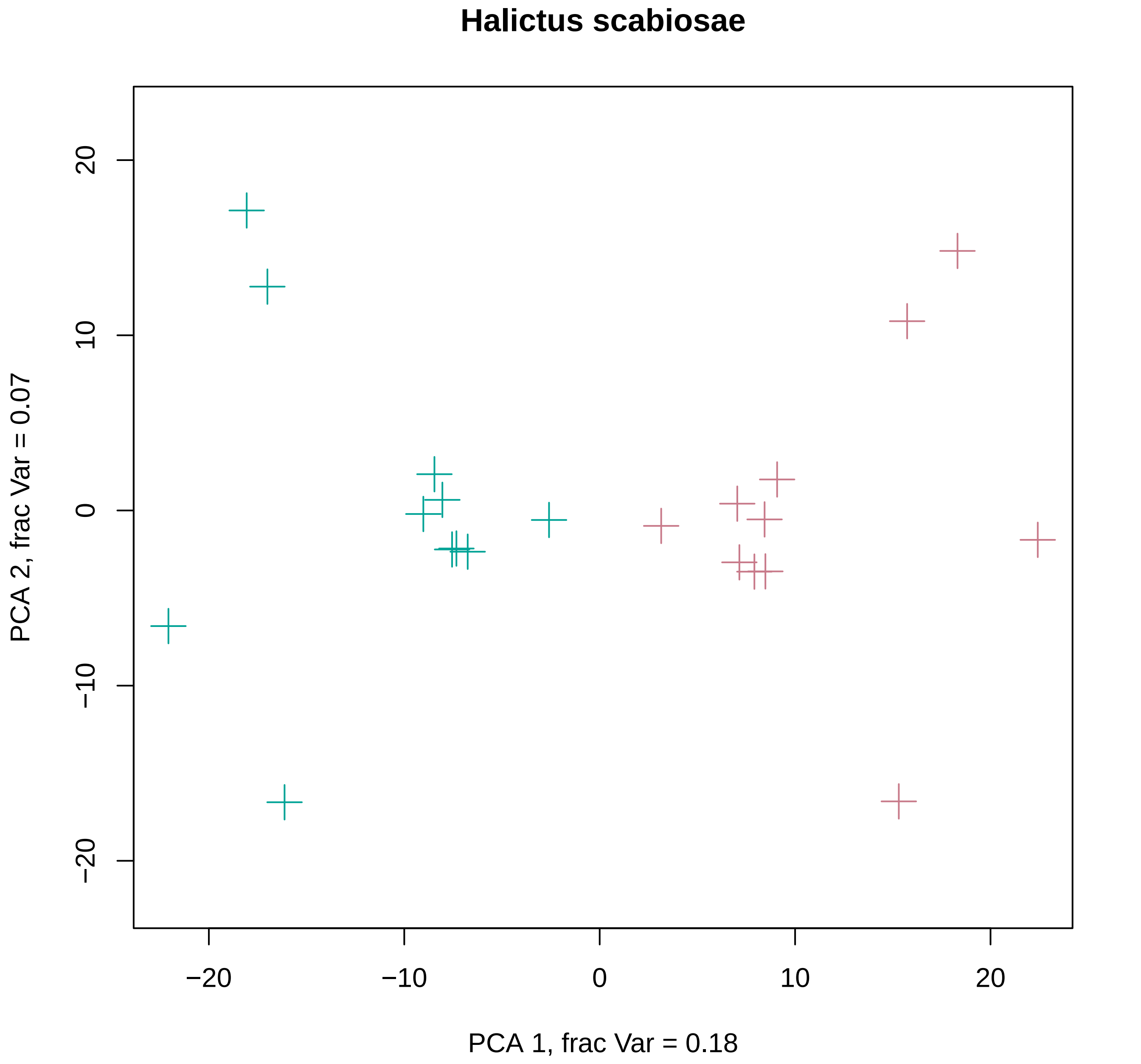

Halictus scabiosae
20
10
0
−10
−20
PCA 2, frac Var = 0.07
−20
−10
0
10
20
PCA 1, frac Var = 0.18

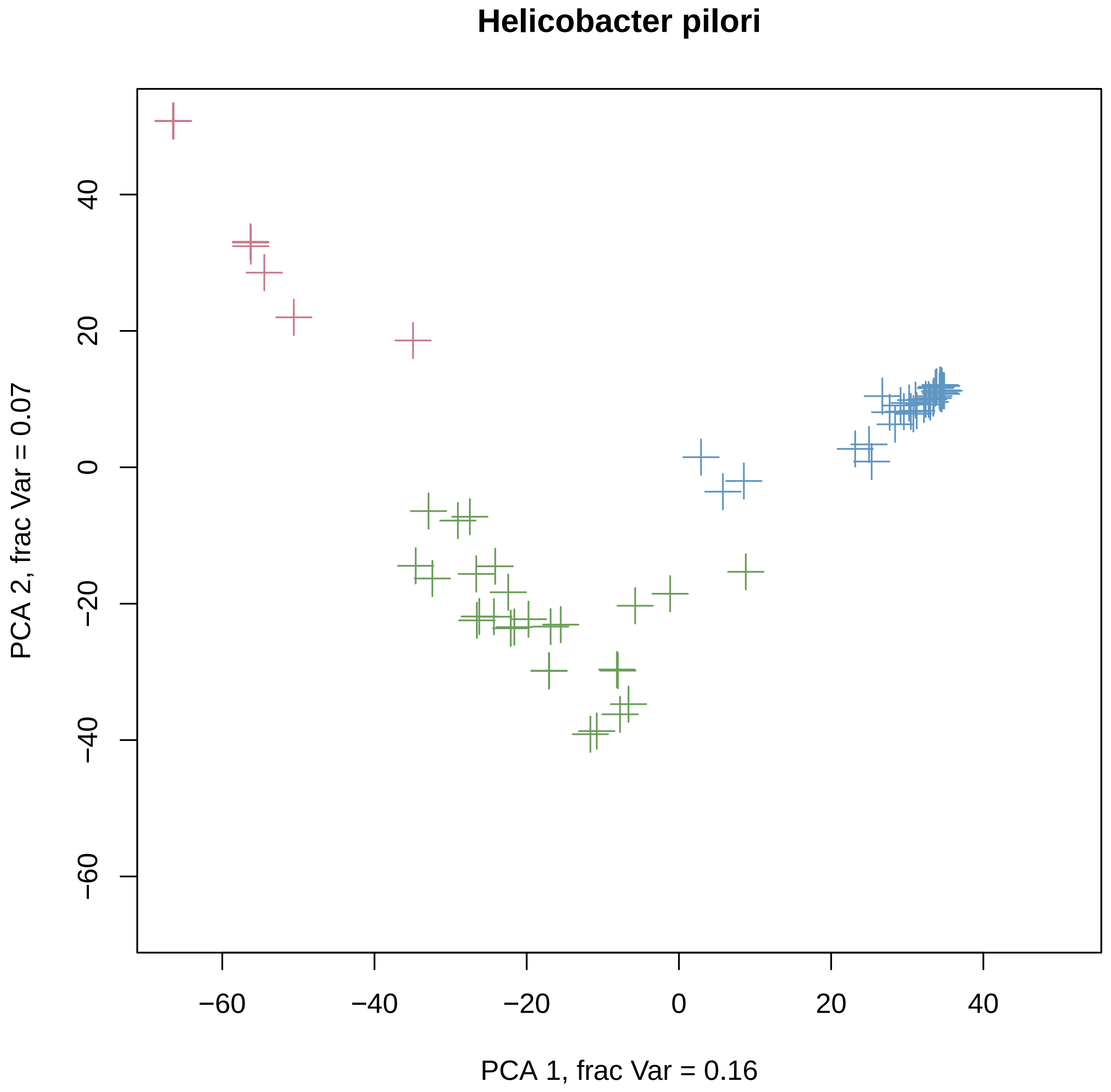
Helicobacter pilori
PCA 1, frac Var = 0.16
PCA 2, frac Var = 0.07
−60
−40
−20
0
20
40

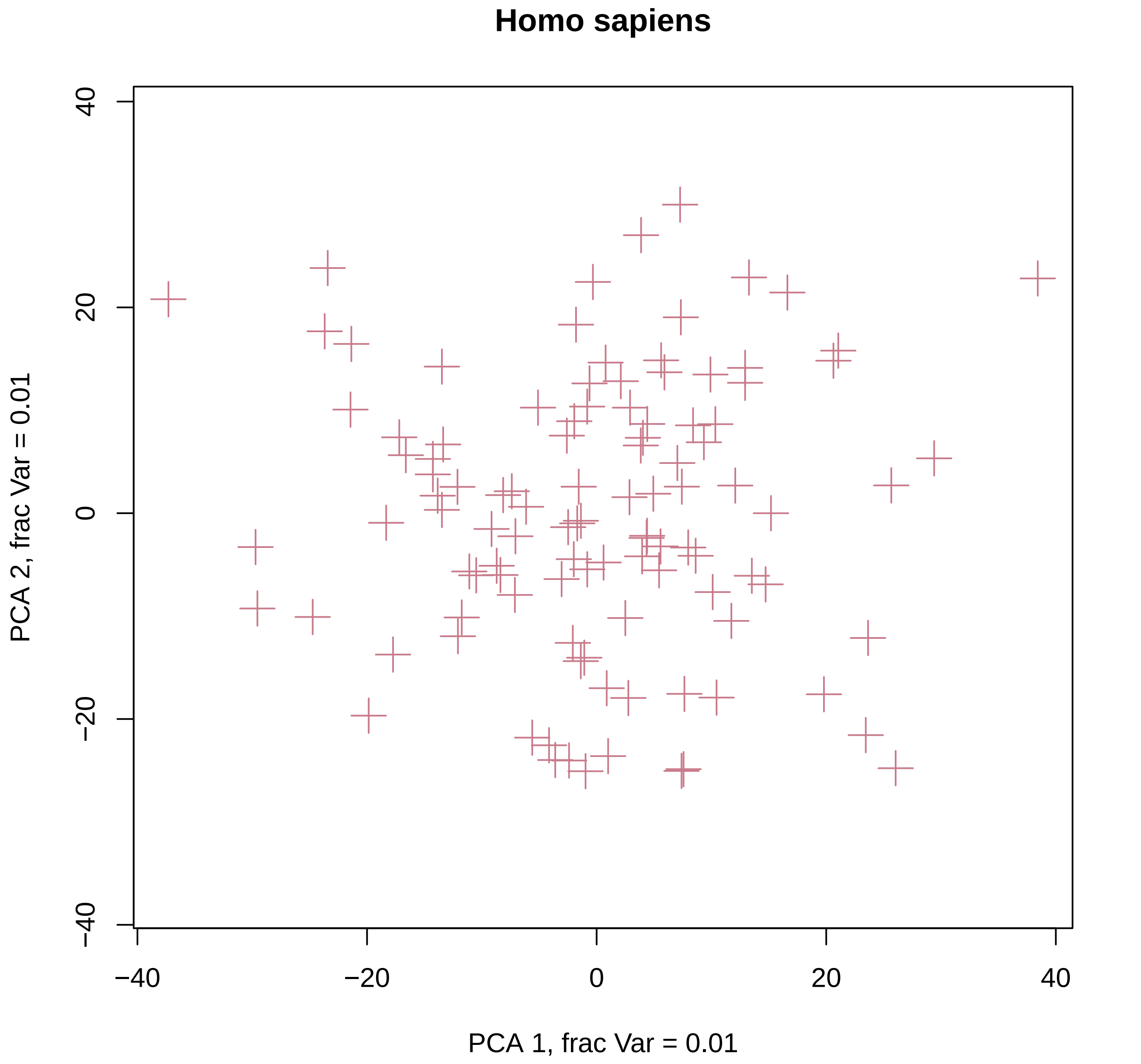
Homo sapiens
PCA 2, frac Var = 0.01
PCA 1, frac Var = 0.01
40
20
0
−20
−40
−40
−20
0
20
40

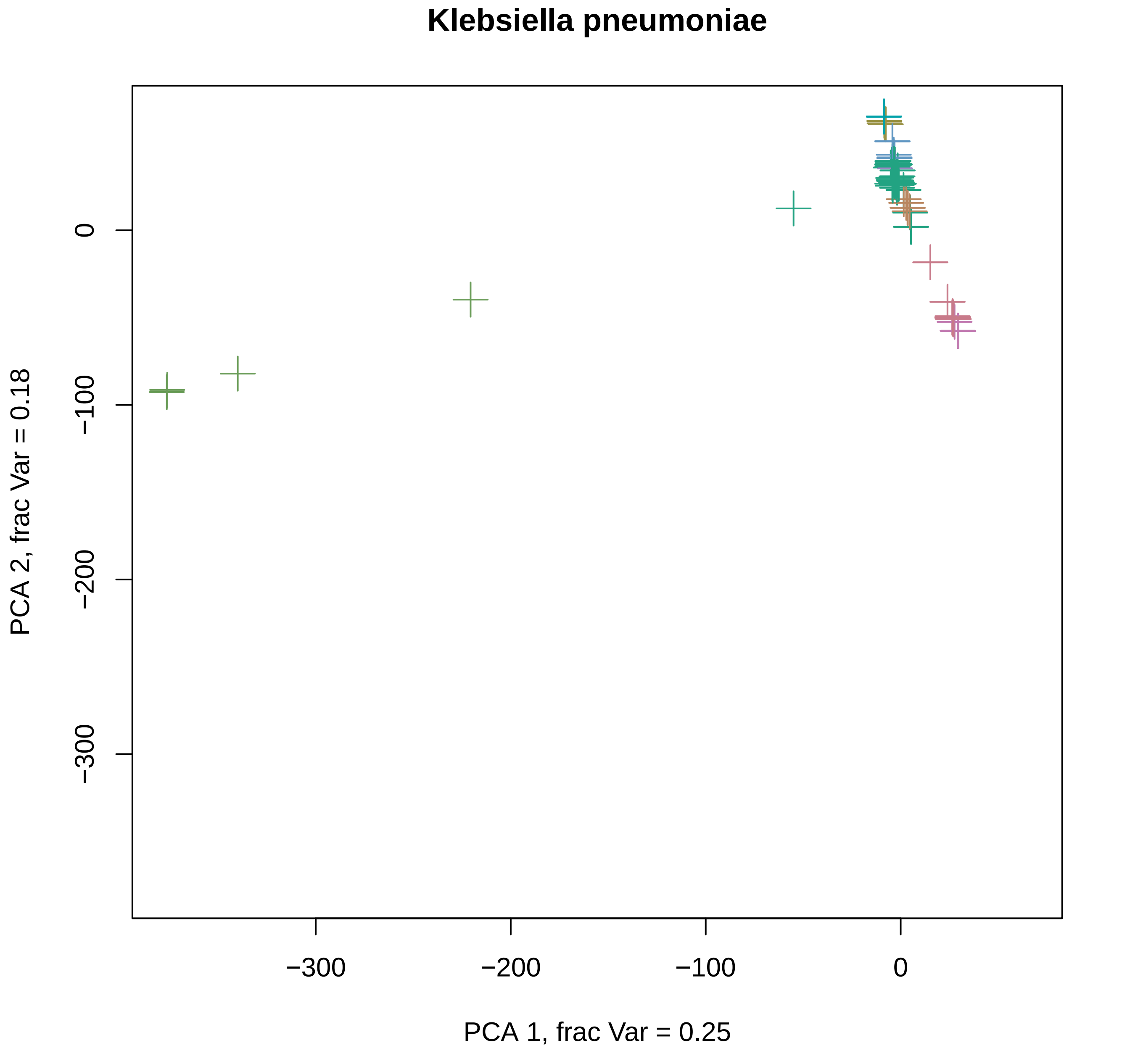
Klebsiella pneumoniae
0
−100
−200
−300
−300
−200
−100
0
PCA 1, frac Var = 0.25
PCA 2, frac Var = 0.18

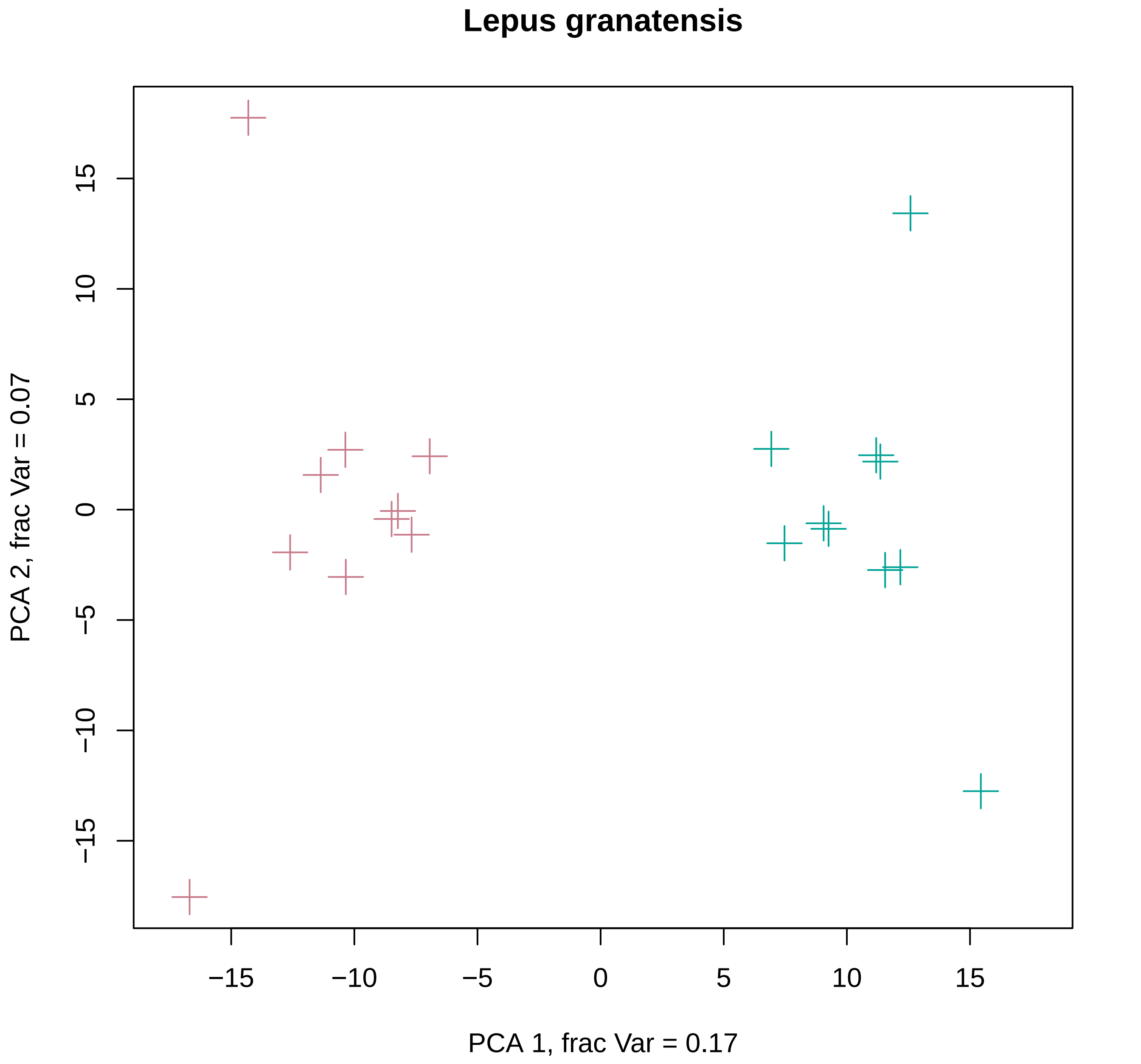

Lepus granatensis
PCA 2, frac Var = 0.07
PCA 1, frac Var = 0.17
−15
−10
−5
0
5
10
15

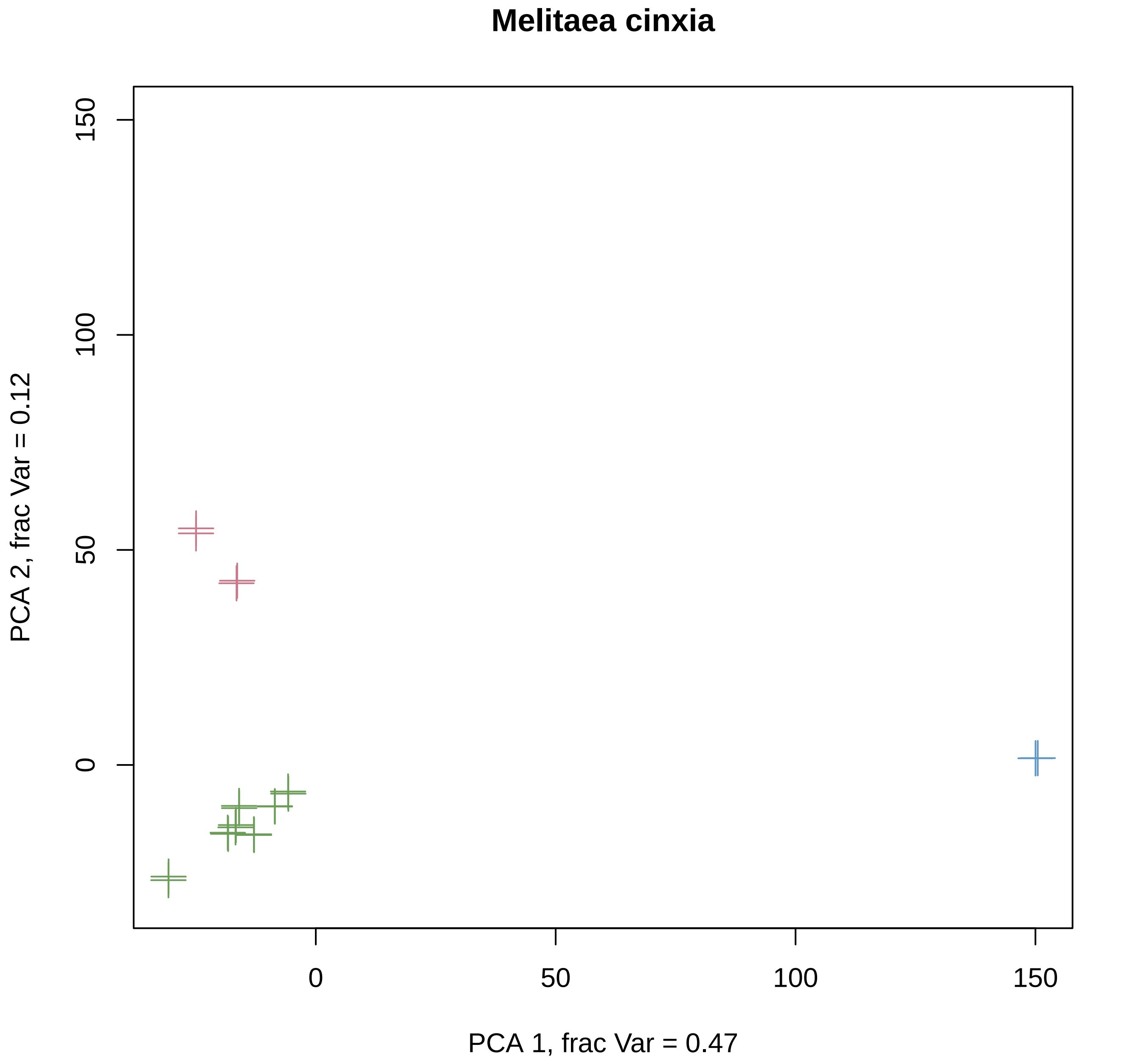

Melitaea cinxia
PCA 2, frac Var = 0.12
0
50
100
150
PCA 1, frac Var = 0.47

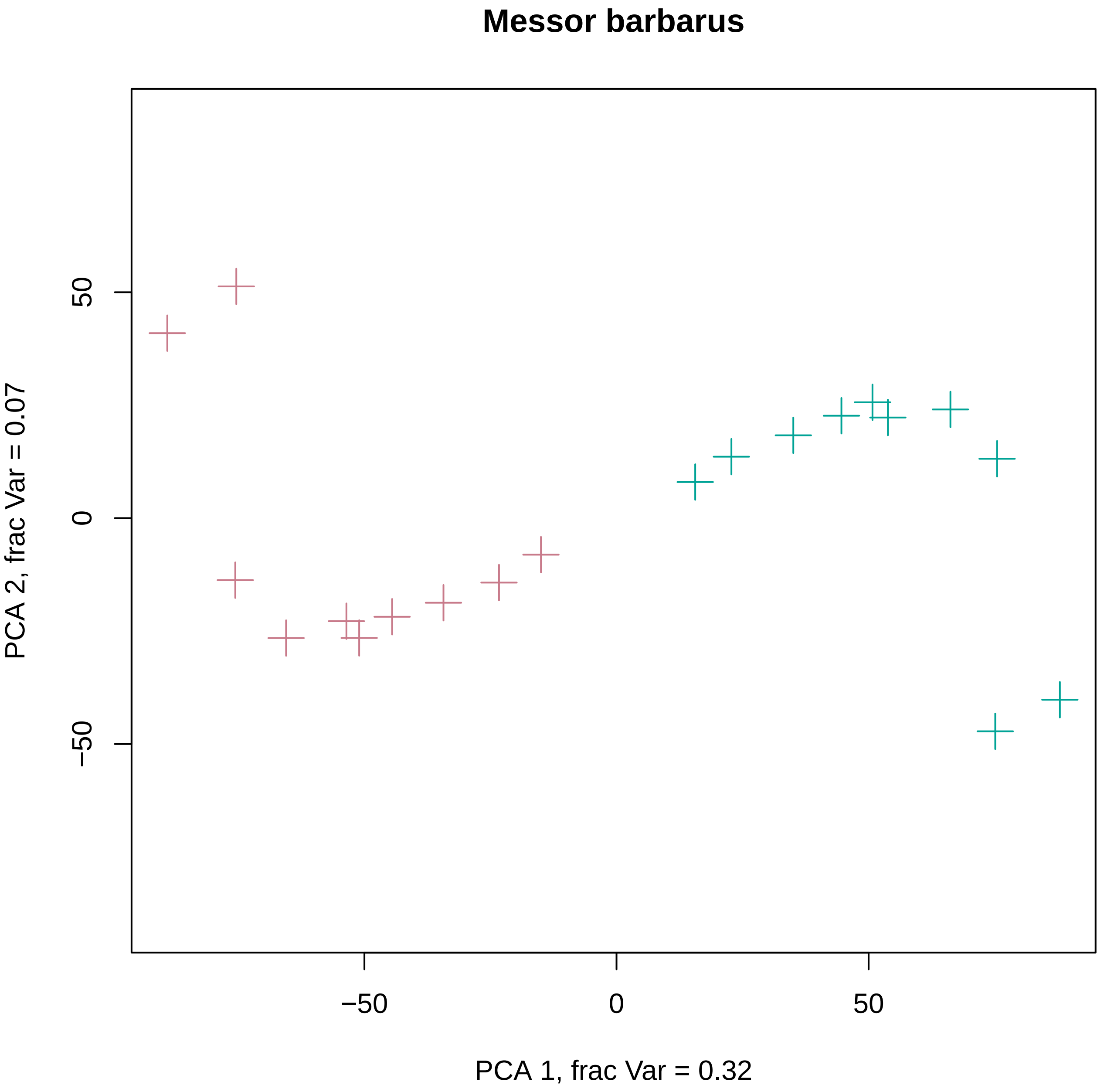

Messor barbarus
PCA 1, frac Var = 0.32
PCA 2, frac Var = 0.07
−50
0
50
−50
0
50

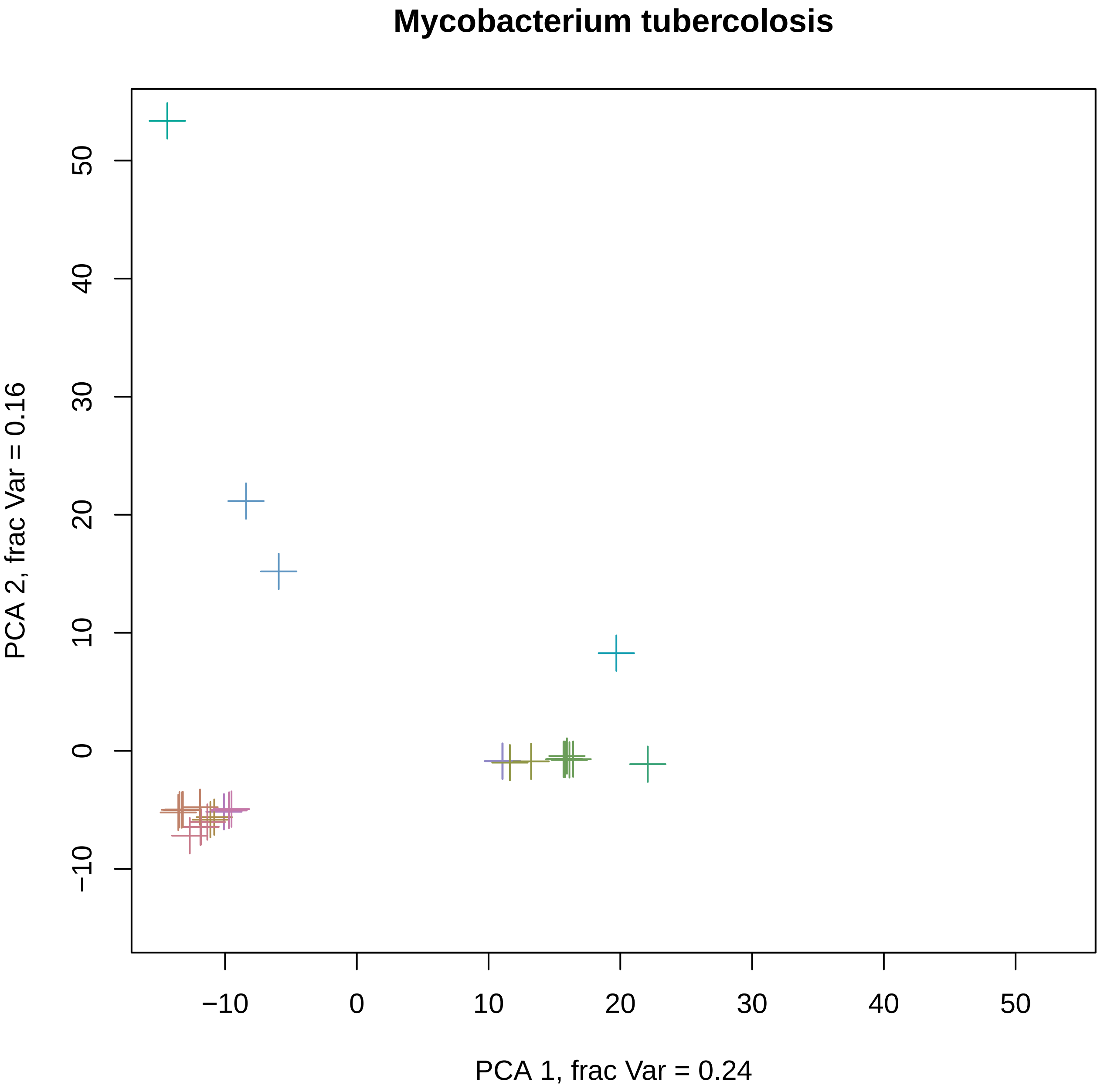
Mycobacterium tubercolosis
PCA 1, frac Var = 0.24
PCA 2, frac Var = 0.16
−10
0
10
20
30
40
50

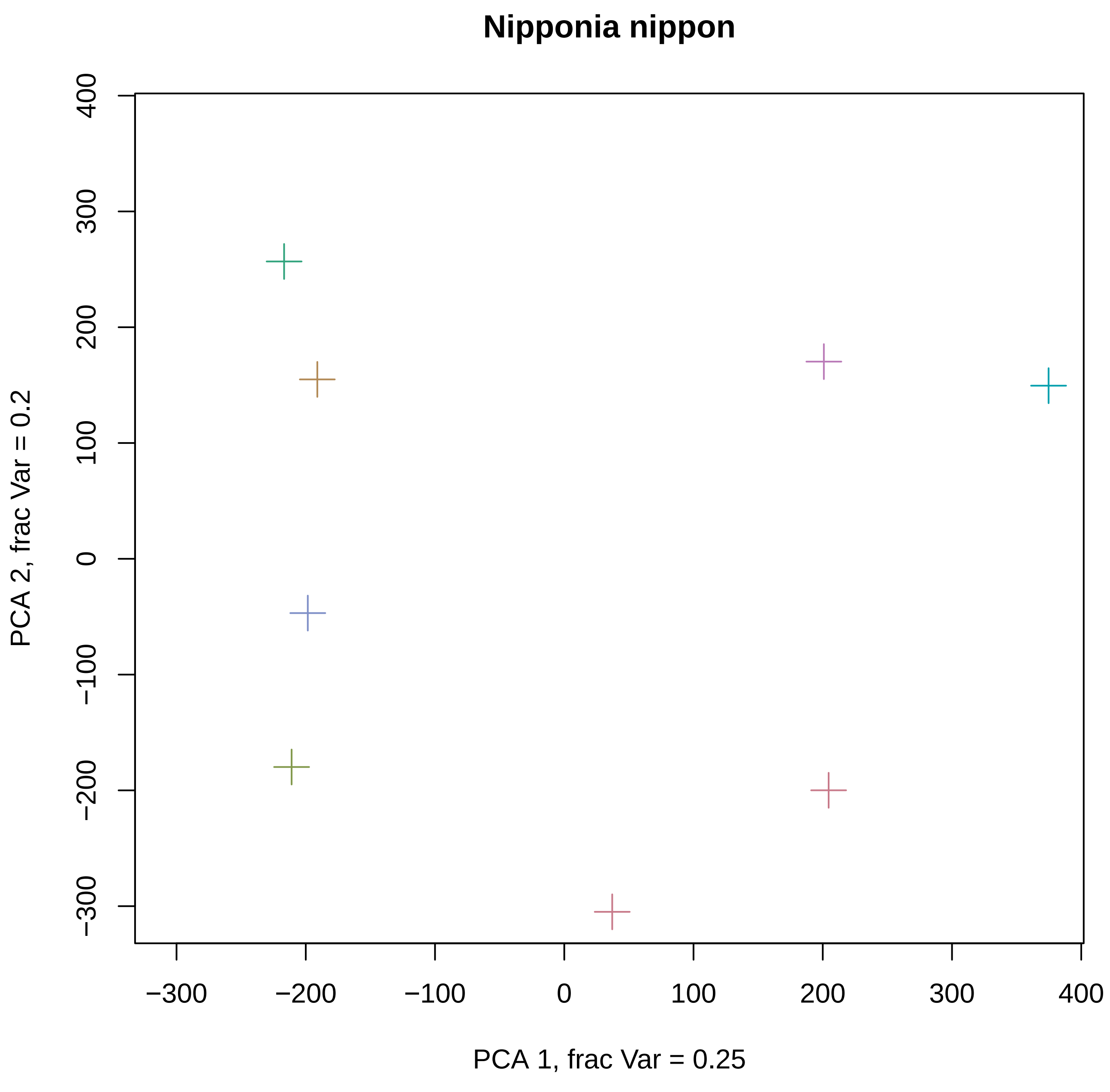
Nipponia nippon
PCA 1, frac Var = 0.25
PCA 2, frac Var = 0.2
−300
−200
−100
0
100
200
300
400

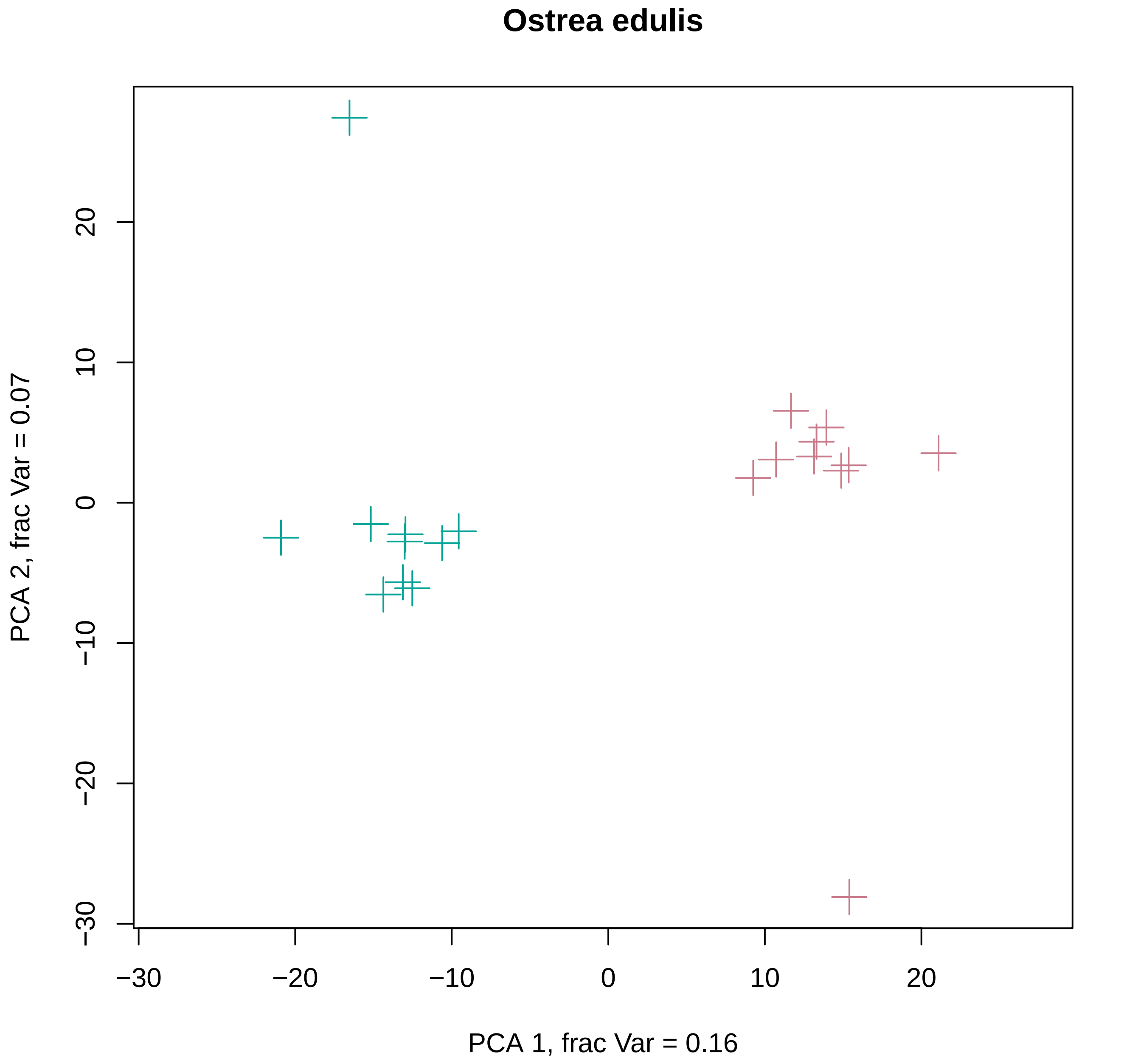
Ostrea edulis
PCA 1, frac Var = 0.16
PCA 2, frac Var = 0.07
−30
−20
−10
0
10
20

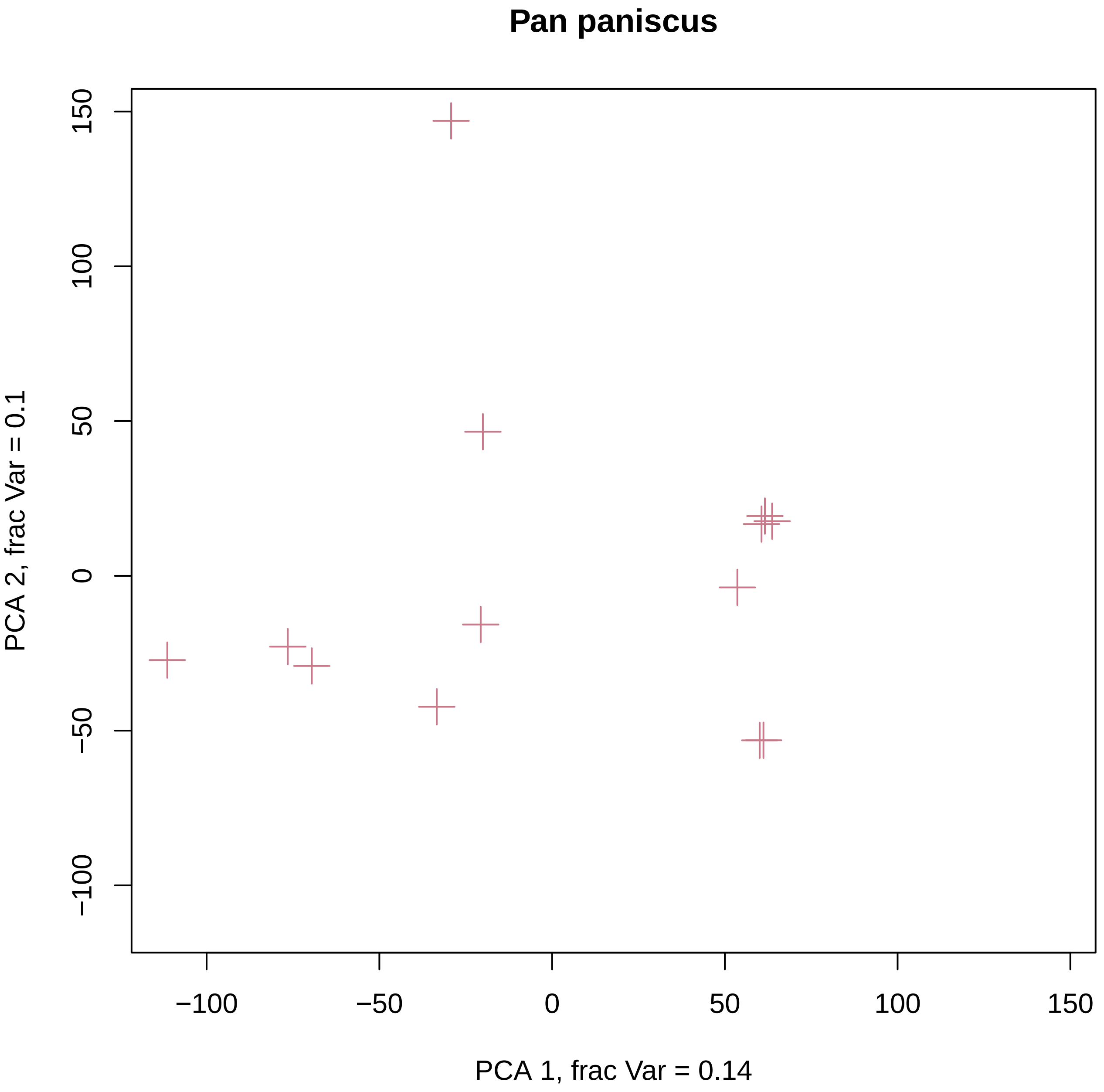

Pan paniscus
PCA 1, frac Var = 0.14
PCA 2, frac Var = 0.1
−100
−50
0
50
100
150

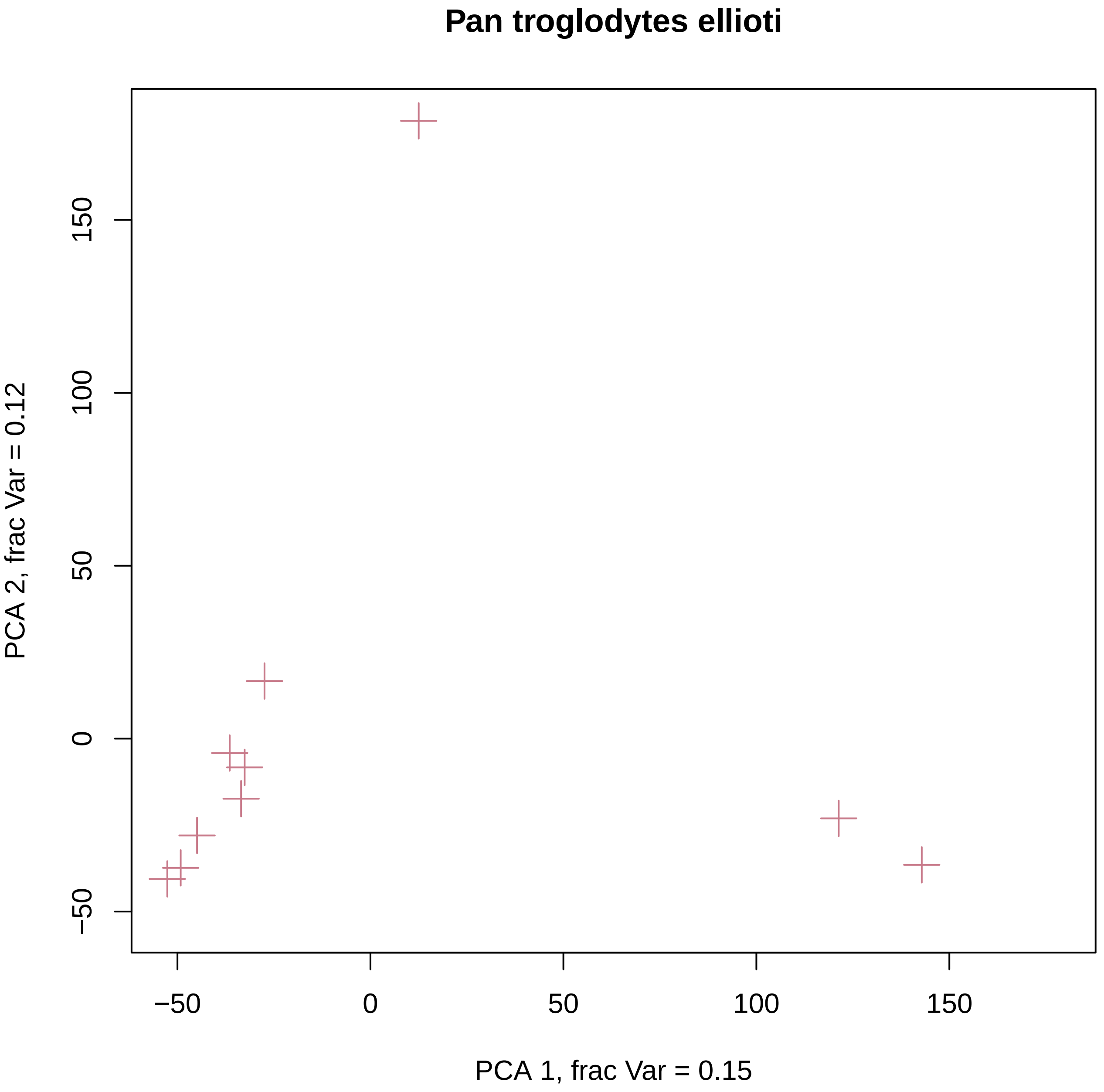
Pan troglodytes ellioti
PCA 1, frac Var = 0.15
PCA 2, frac Var = 0.12
−50
0
50
100
150
−50
0
50
100
150

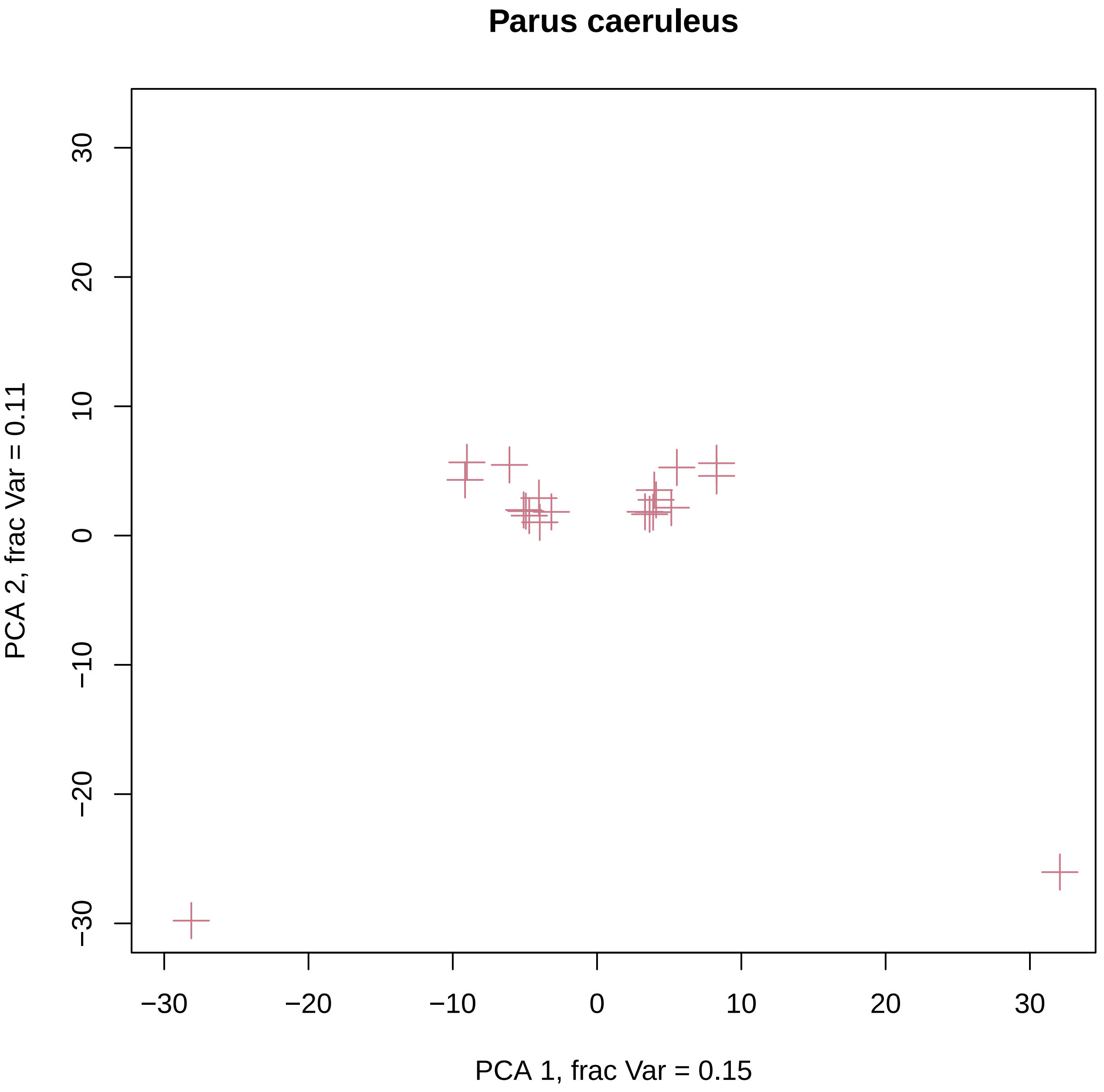
Parus caeruleus
PCA 1, frac Var = 0.15
PCA 2, frac Var = 0.11
−30
−20
−10
0
10
20
30

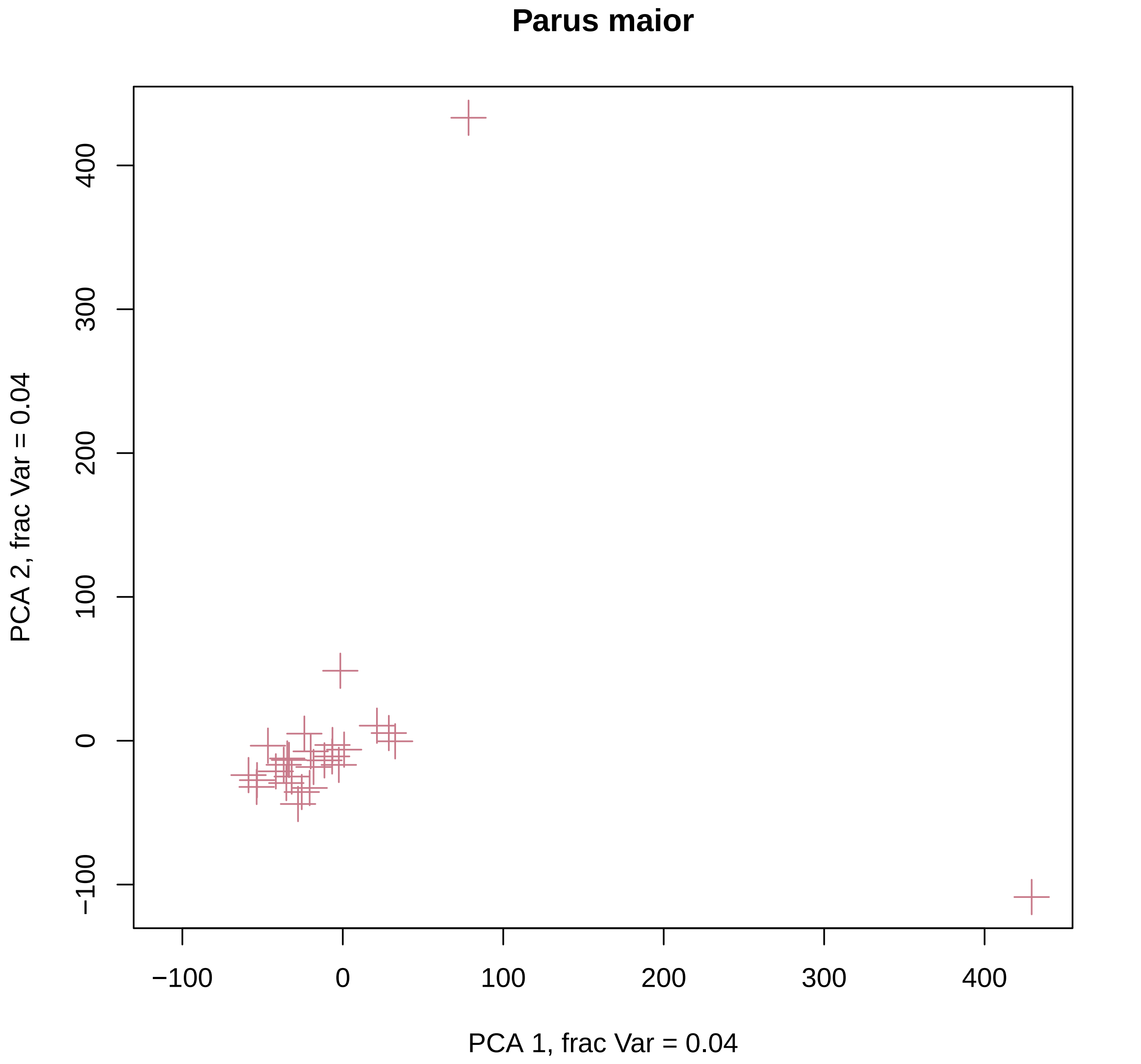
Parus maior
PCA 2, frac Var = 0.04
−100
0
100
200
300
400
−100
0
100
200
300
400
PCA 1, frac Var = 0.04

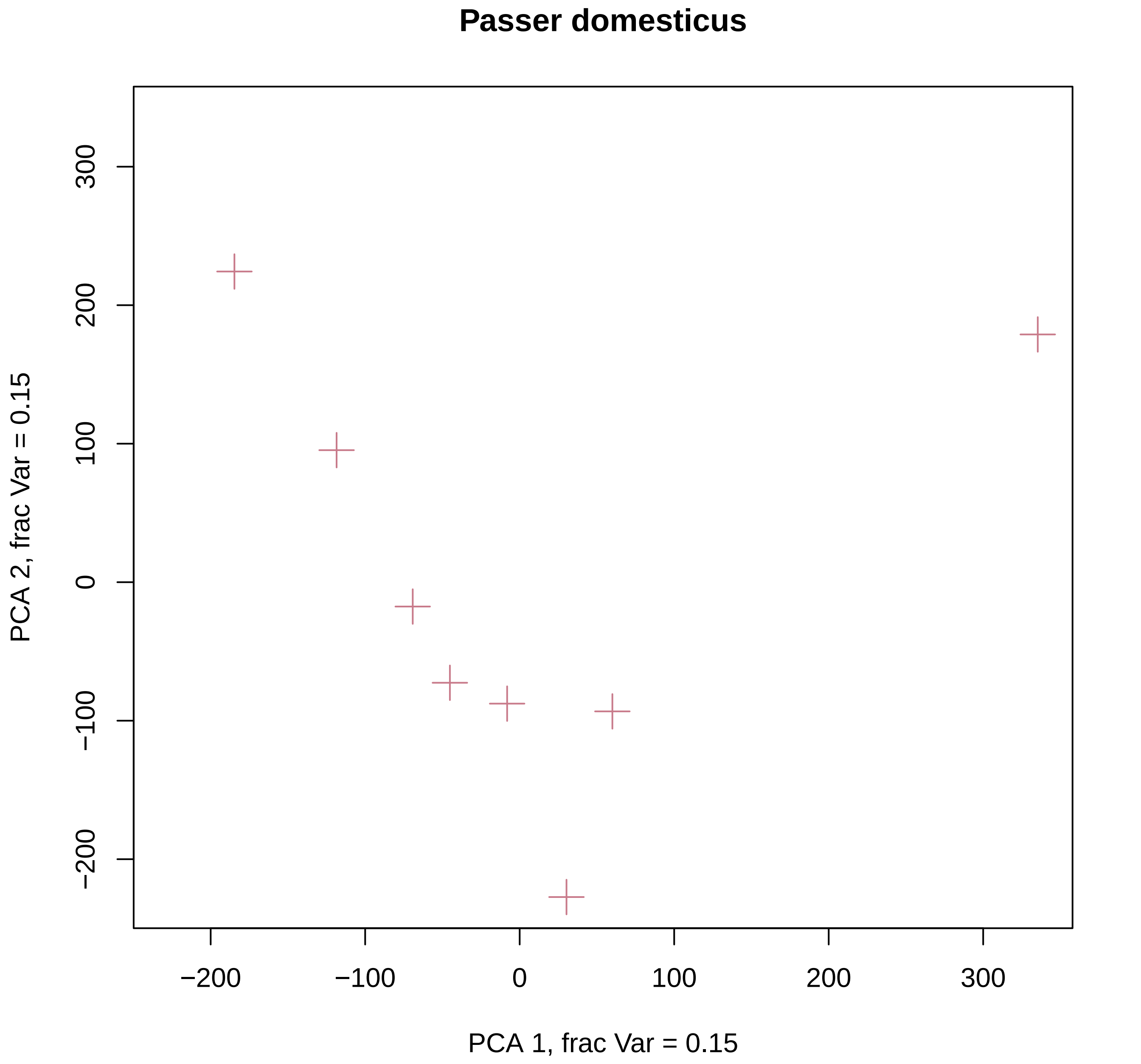
Passer domesticus
PCA 1, frac Var = 0.15
PCA 2, frac Var = 0.15
−200
−100
0
100
200
300
−200
−100
0
100
200
300

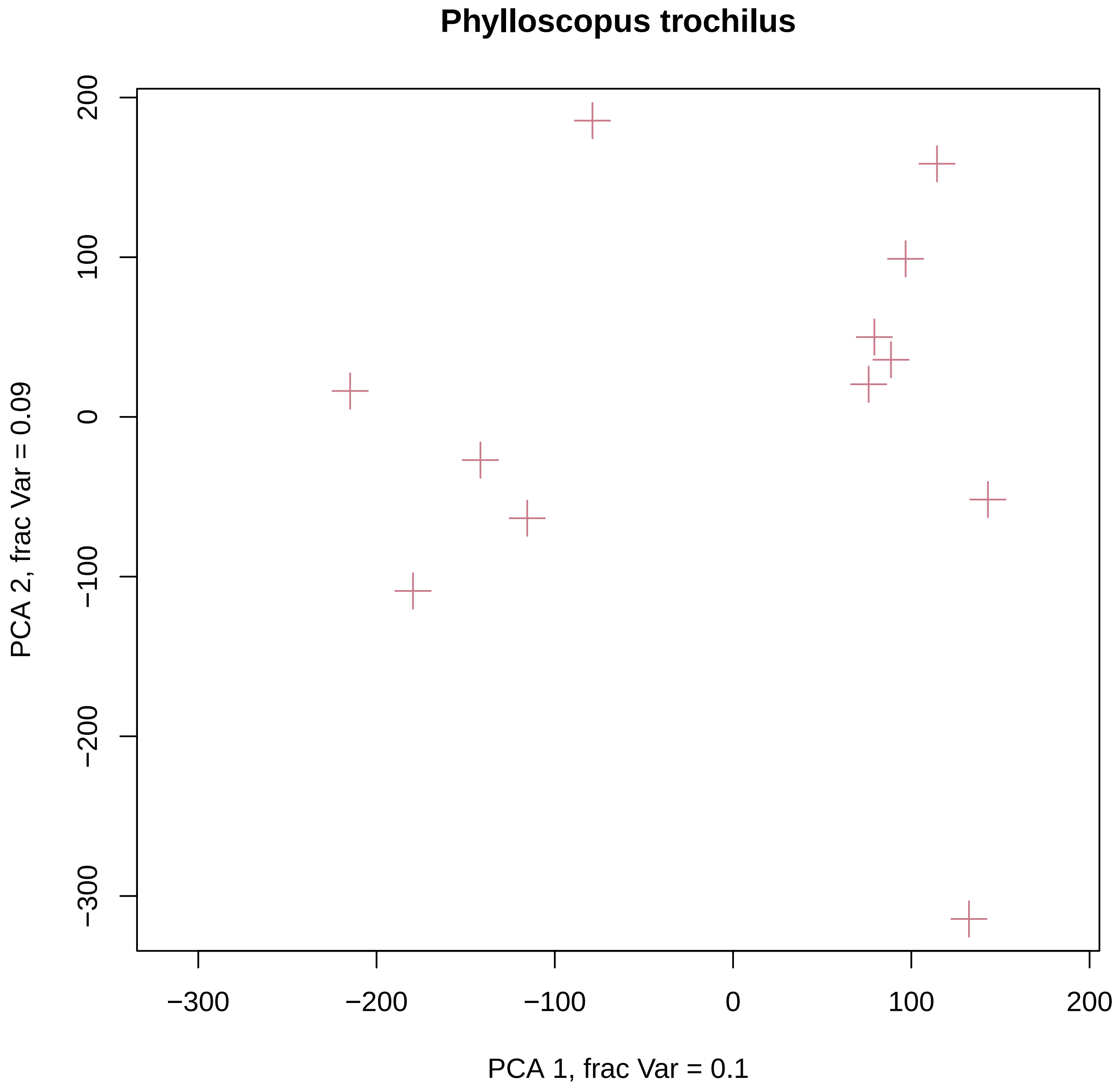
Phylloscopus trochilus
PCA 1, frac Var = 0.1
PCA 2, frac Var = 0.09
−300
−200
−100
0
100
200

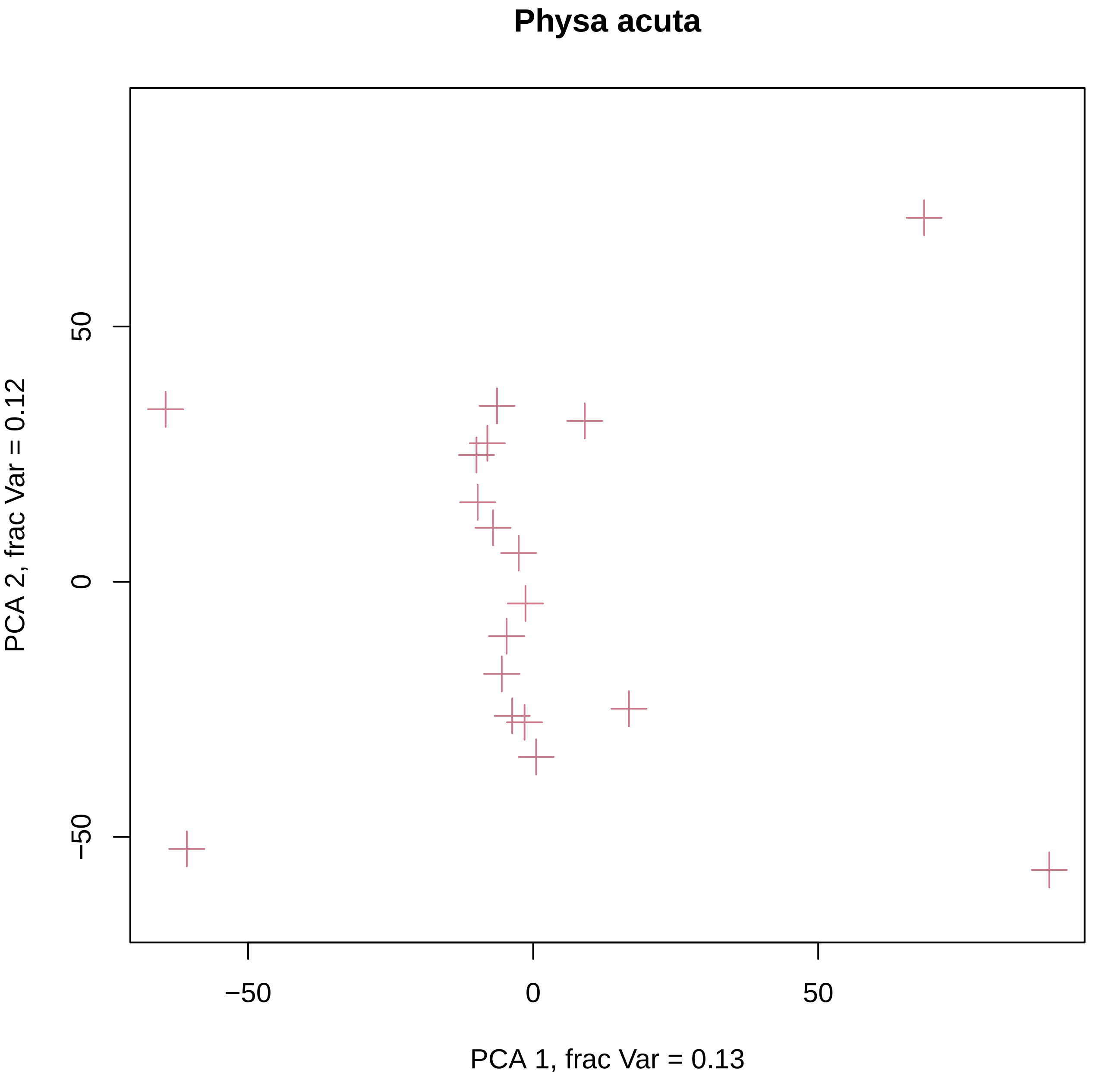
Physa acuta
PCA 1, frac Var = 0.13
PCA 2, frac Var = 0.12
−50
0
50
−50
0
50

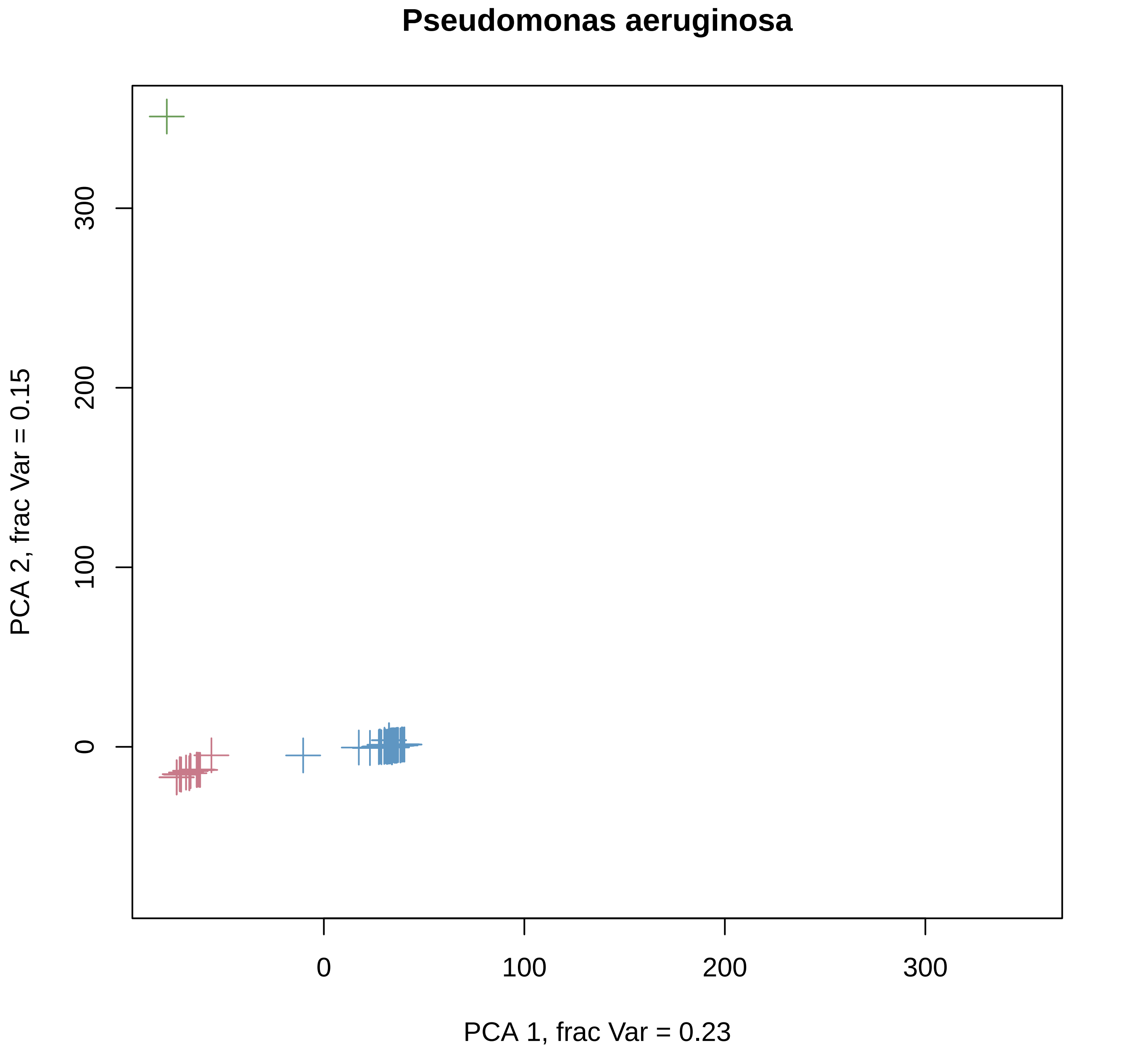
Pseudomonas aeruginosa
PCA 1, frac Var = 0.23
PCA 2, frac Var = 0.15
0
100
200
300
0
100
200
300

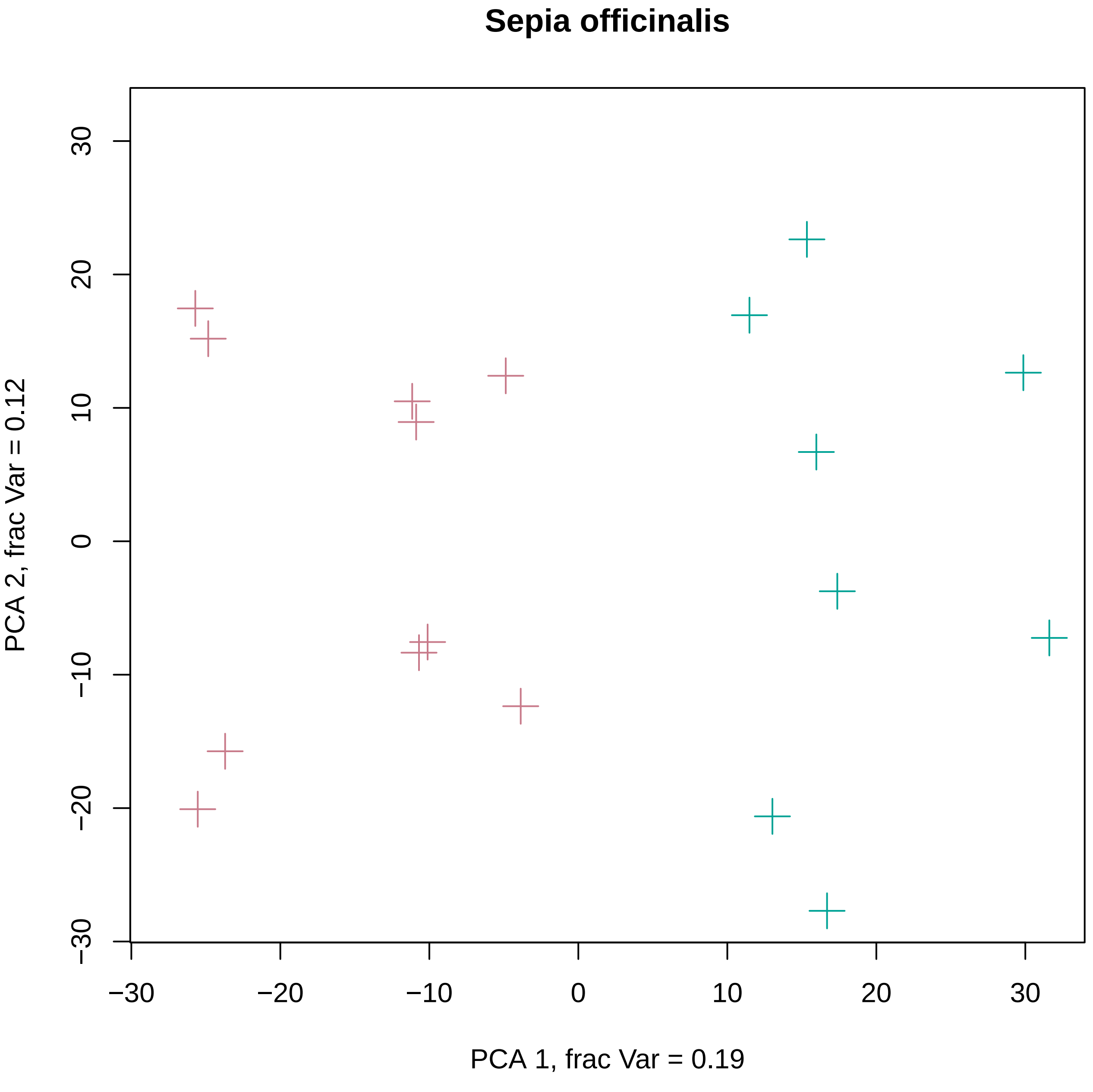

Sepia officinalis
PCA 1, frac Var = 0.19
PCA 2, frac Var = 0.12
−30
−20
−10
0
10
20
30

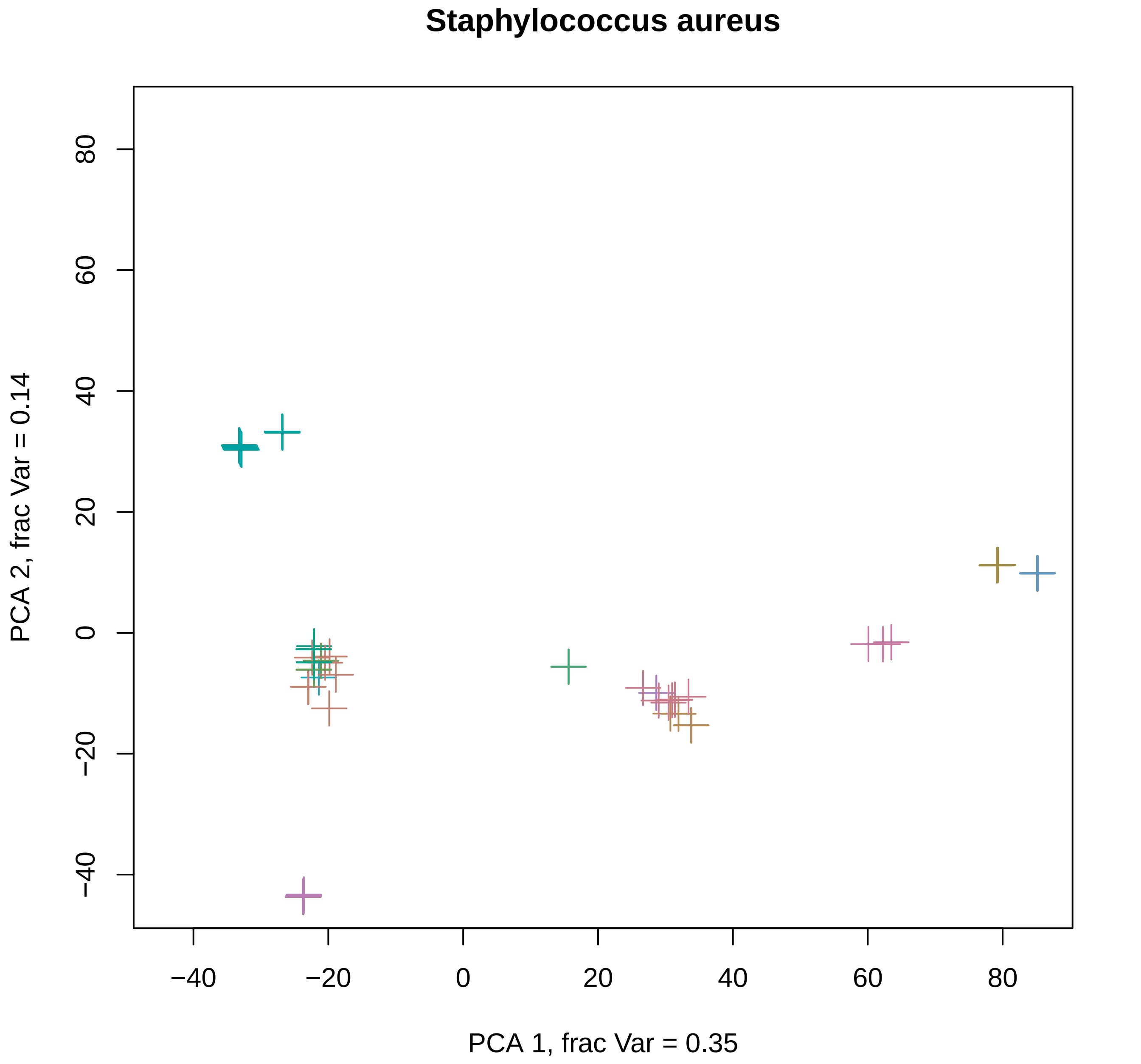
Staphylococcus aureus
80
60
40
20
0
−20
−40
PCA 2, frac Var = 0.14
−40
−20
0
20
40
60
80
PCA 1, frac Var = 0.35

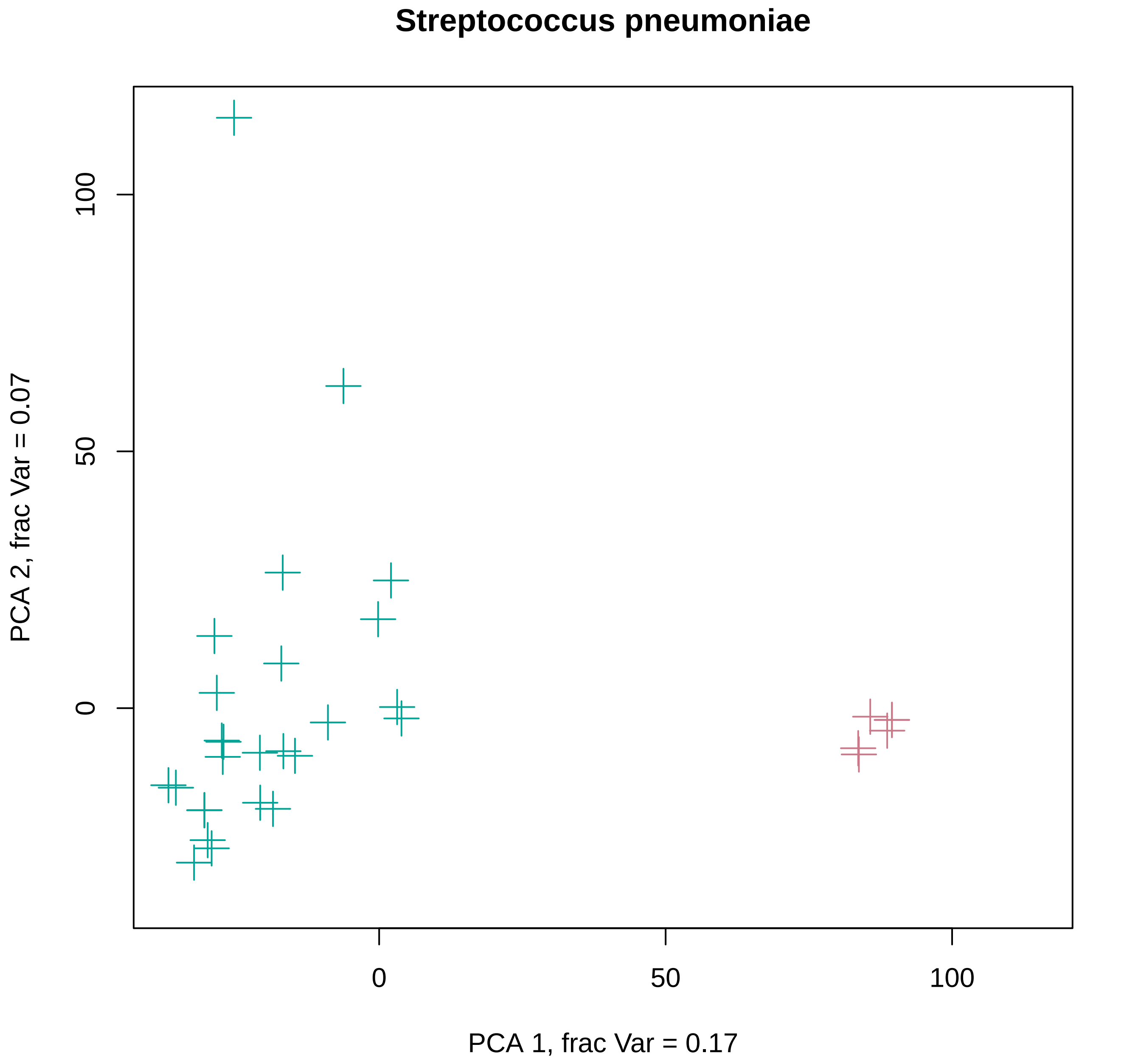
Streptococcus pneumoniae
PCA 1, frac Var = 0.17
PCA 2, frac Var = 0.07
0
50
100
0
50
100

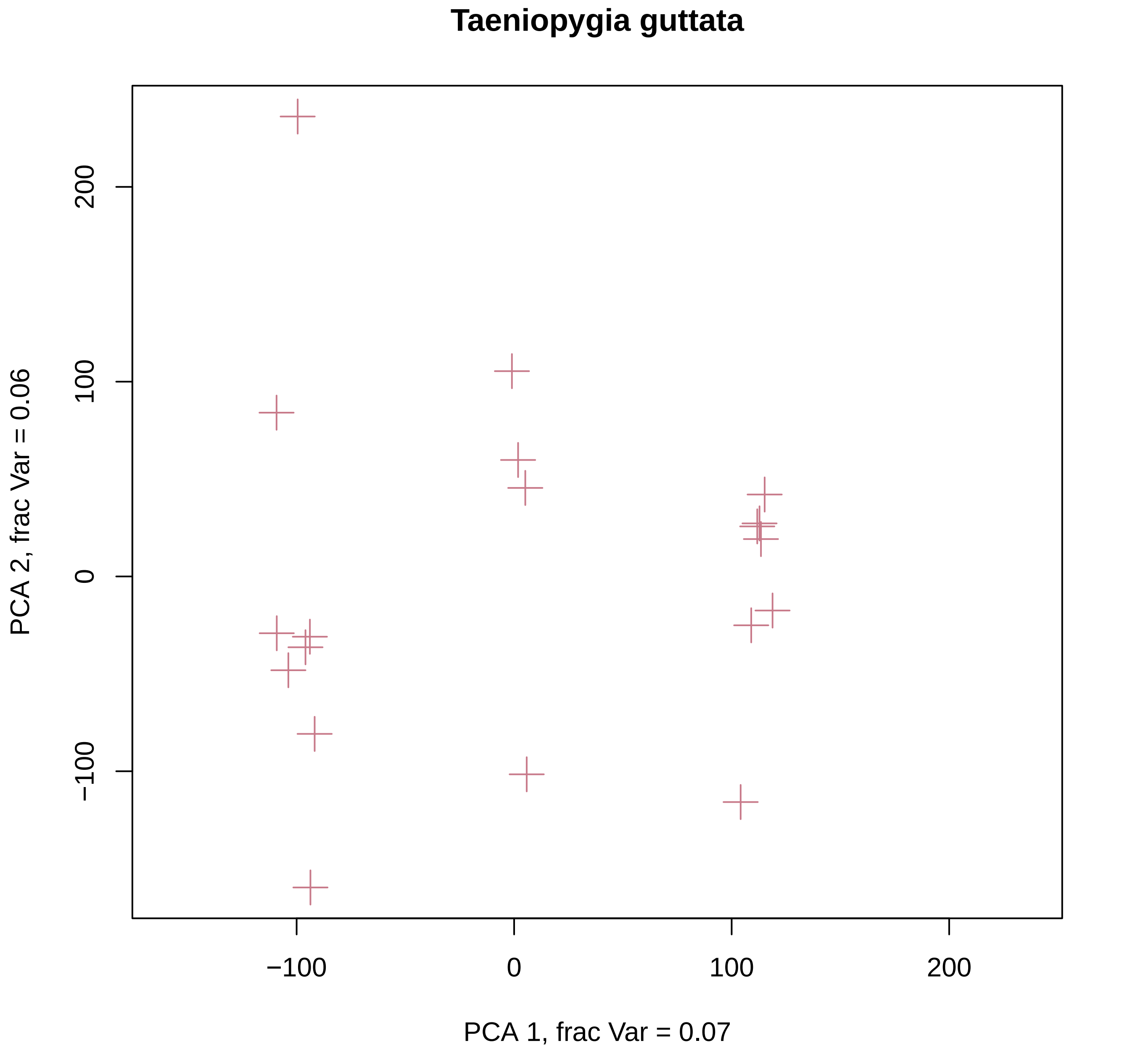
Taeniopygia guttata
PCA 2, frac Var = 0.06
PCA 1, frac Var = 0.07
200
100
0
−100
−100
0
100
200

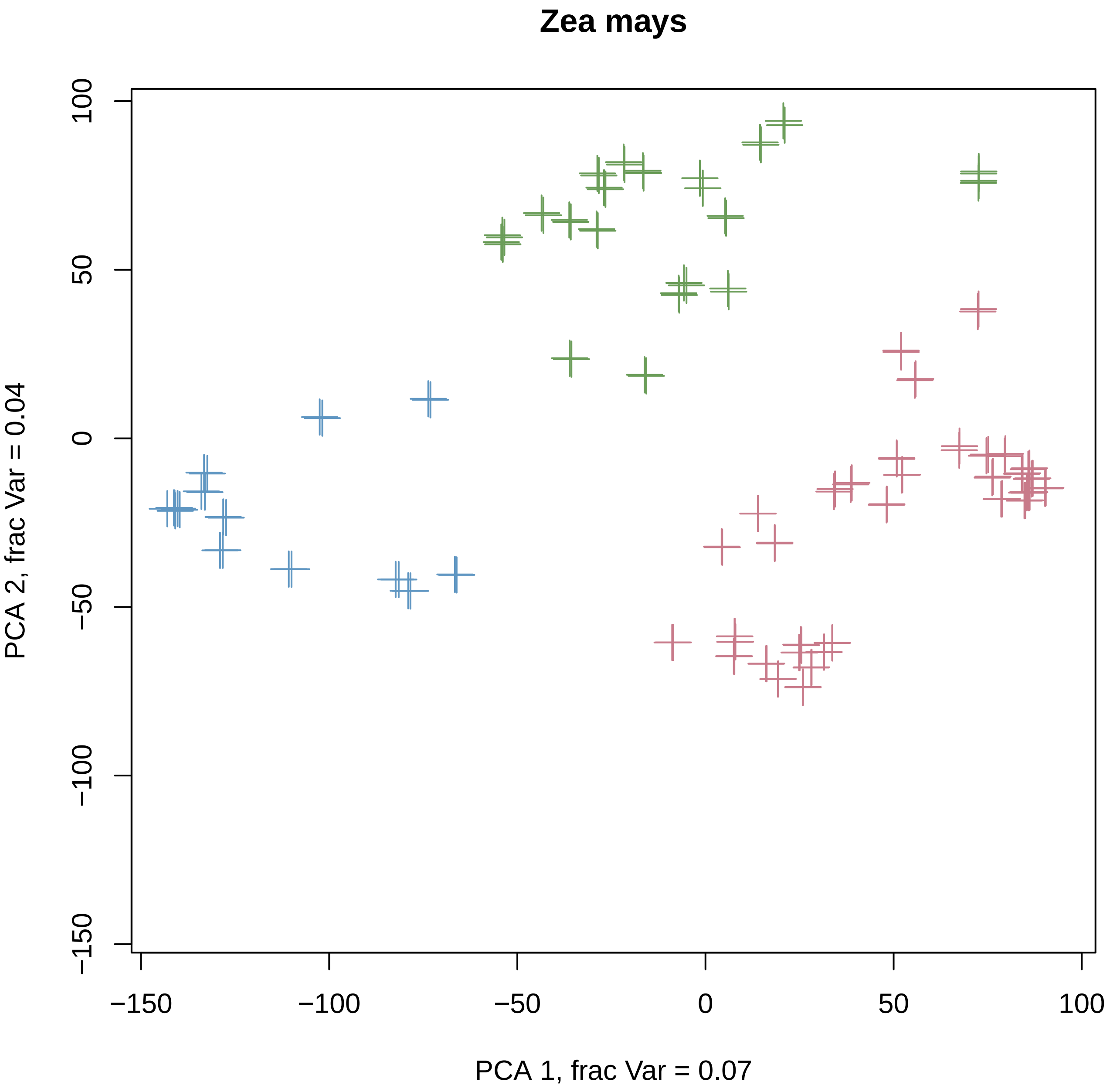
Zea mays
PCA 1, frac Var = 0.07
PCA 2, frac Var = 0.04
−150
−100
−50
0
50
100
