## Supplementary 1 - SFS and fits for "Interpreting the pervasive observation of U-shaped Site Frequency Spectra"

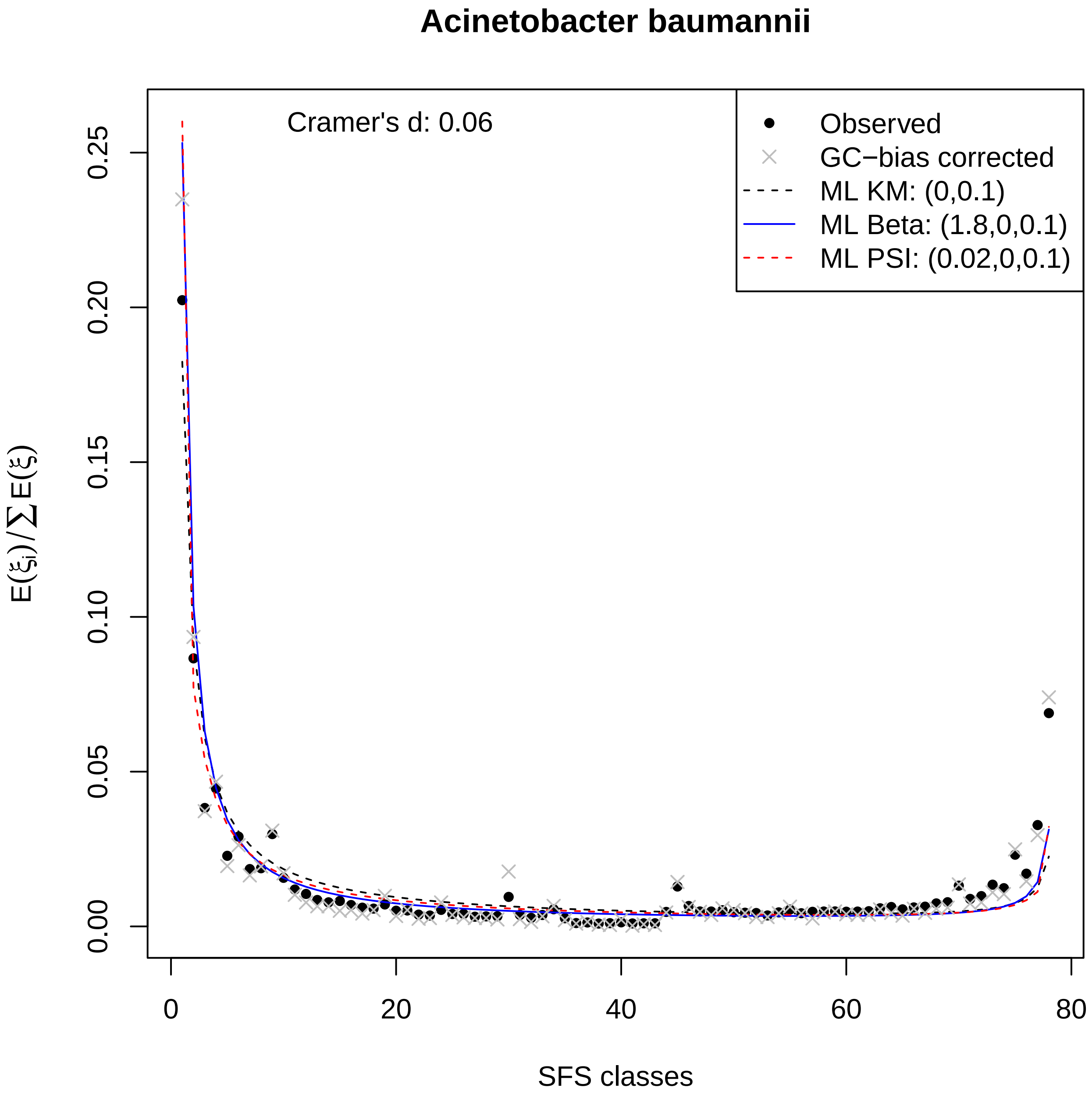

Acinetobacter baumannii
Cramer's d: 0.06
Observed
GC−bias corrected
ML KM: (0,0.1)
ML Beta: (1.8,0,0.1)
ML PSI: (0.02,0,0.1)
0.00
0.05
0.10
0.15
0.20
0.25
0
20
40
60
80
$E(\xi_i)/\sum E(\xi)$
SFS classes

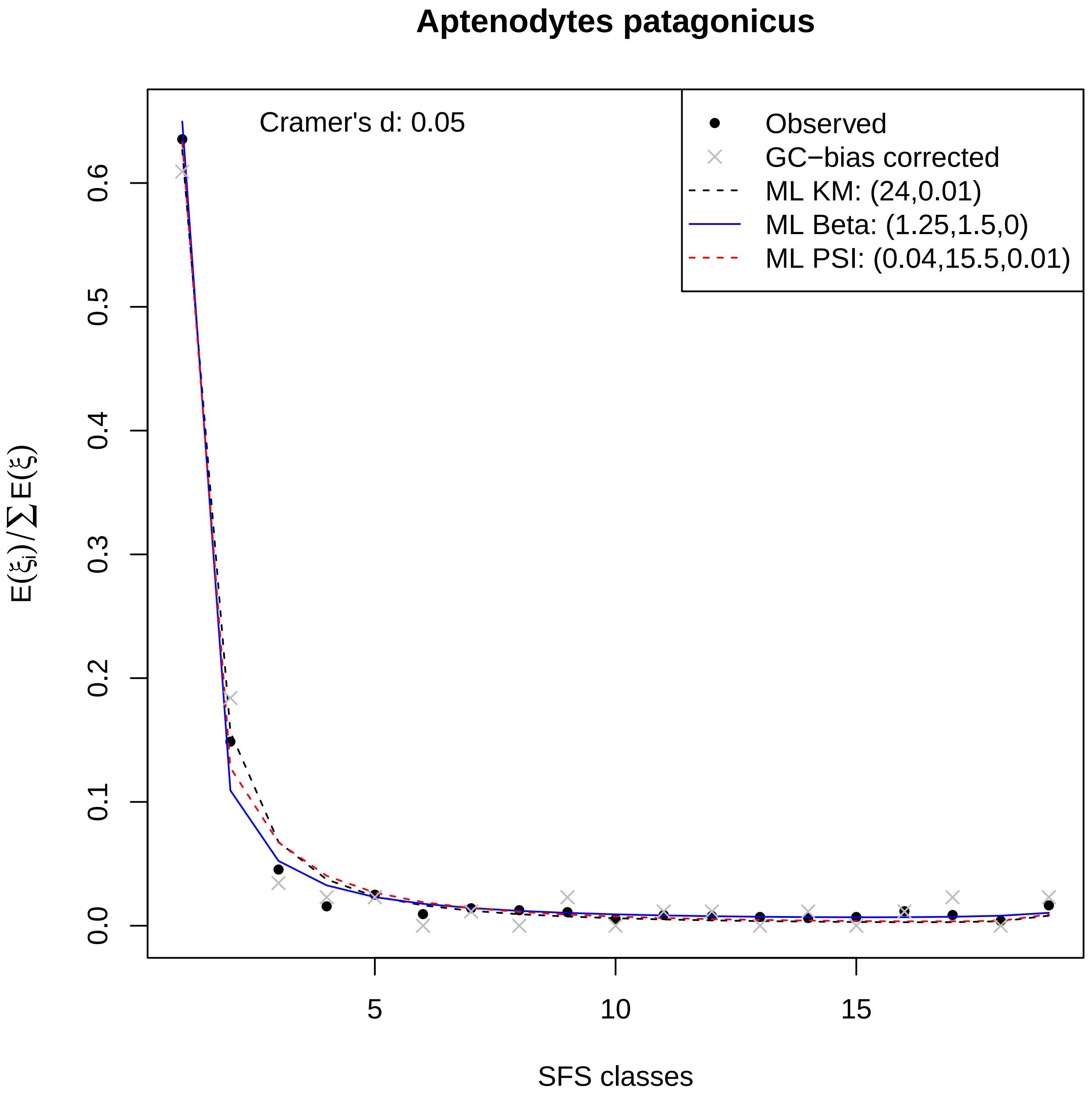

Aptenodytes patagonicus
Cramer's d: 0.05
Observed
GC−bias corrected
ML KM: (24,0.01)
ML Beta: (1.25,1.5,0)
ML PSI: (0.04,15.5,0.01)
0.0
0.1
0.2
0.3
0.4
0.5
0.6
5
10
15
$E(\xi_i)/\sum E(\xi)$
SFS classes

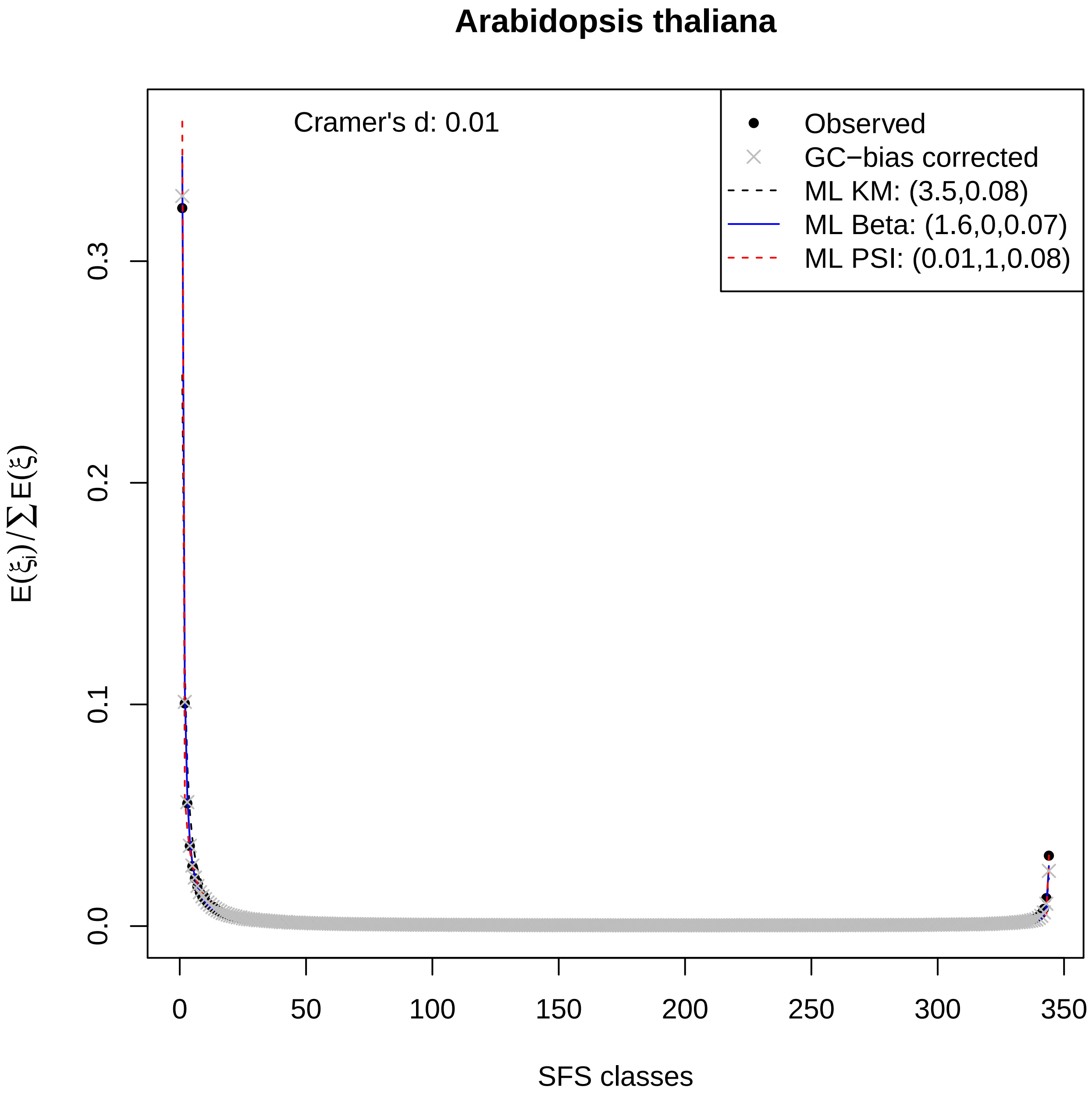
Arabidopsis thaliana
Cramer's d: 0.01
Observed
GC−bias corrected
ML KM: (3.5,0.08)
ML Beta: (1.6,0,0.07)
ML PSI: (0.01,1,0.08)
0.0
0.1
0.2
0.3
$E(\xi_i)/\sum E(\xi)$
0
50
100
150
200
250
300
350
SFS classes

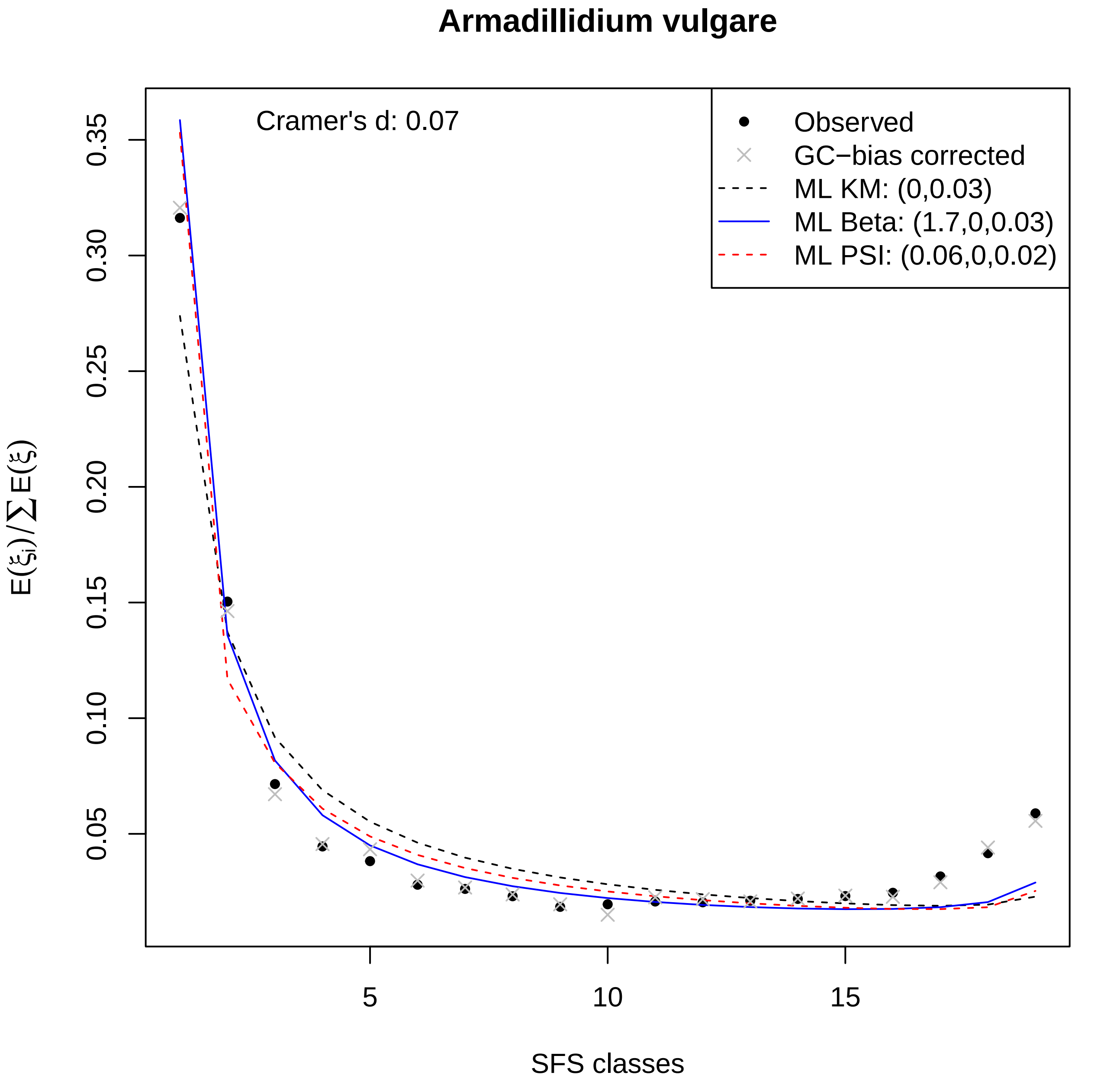
Armadillidium vulgare
Cramer's d: 0.07
Observed
GC−bias corrected
ML KM: (0,0.03)
ML Beta: (1.7,0,0.03)
ML PSI: (0.06,0,0.02)
0.35
0.30
0.25
0.20
0.15
0.10
0.05
5
10
15
SFS classes

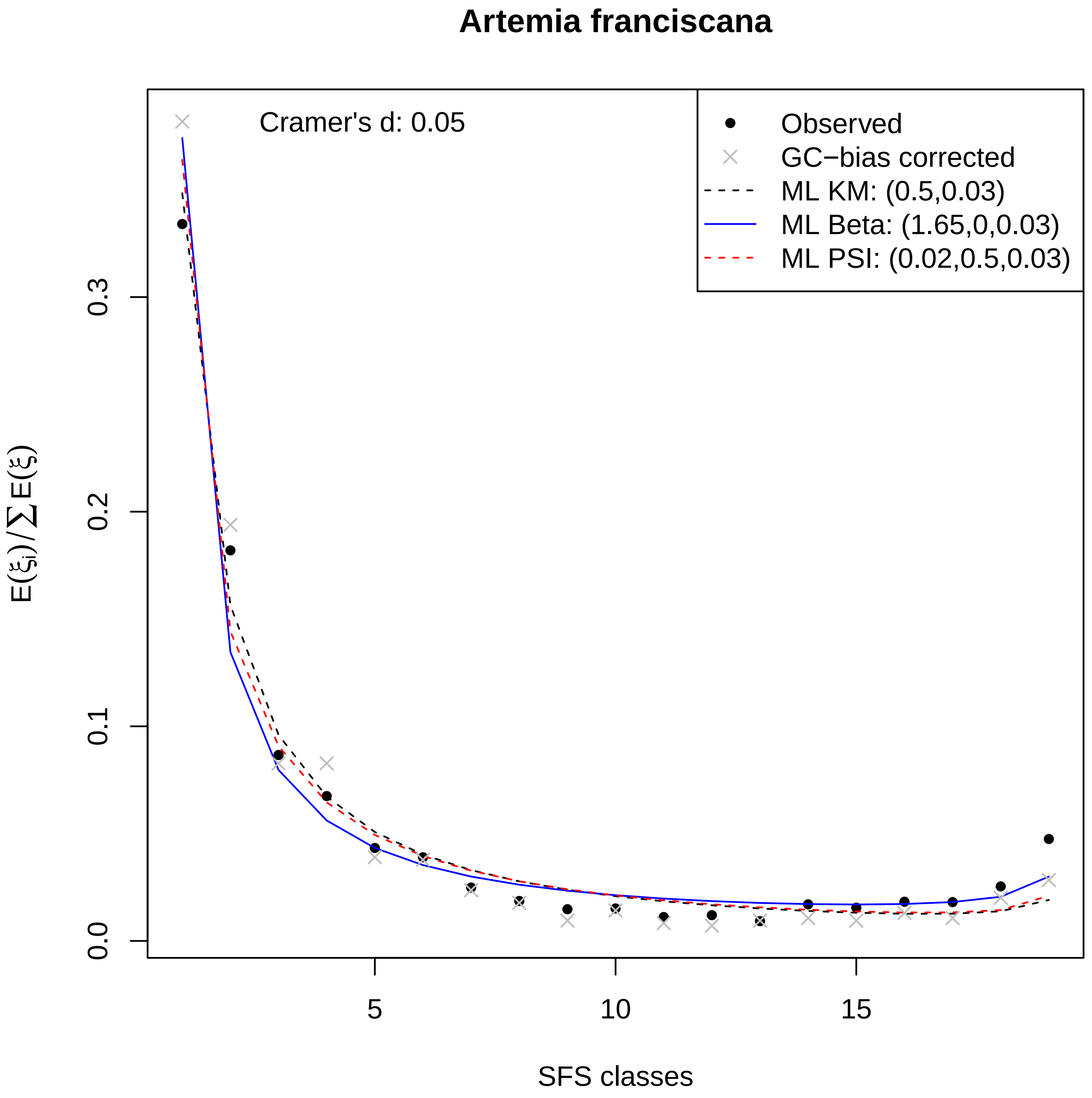

Artemia franciscana
Cramer's d: 0.05
Observed
GC−bias corrected
ML KM: (0.5,0.03)
ML Beta: (1.65,0,0.03)
ML PSI: (0.02,0.5,0.03)
0.0
0.1
0.2
0.3
5
10
15
SFS classes
$E(\xi_i)/\sum E(\xi)$

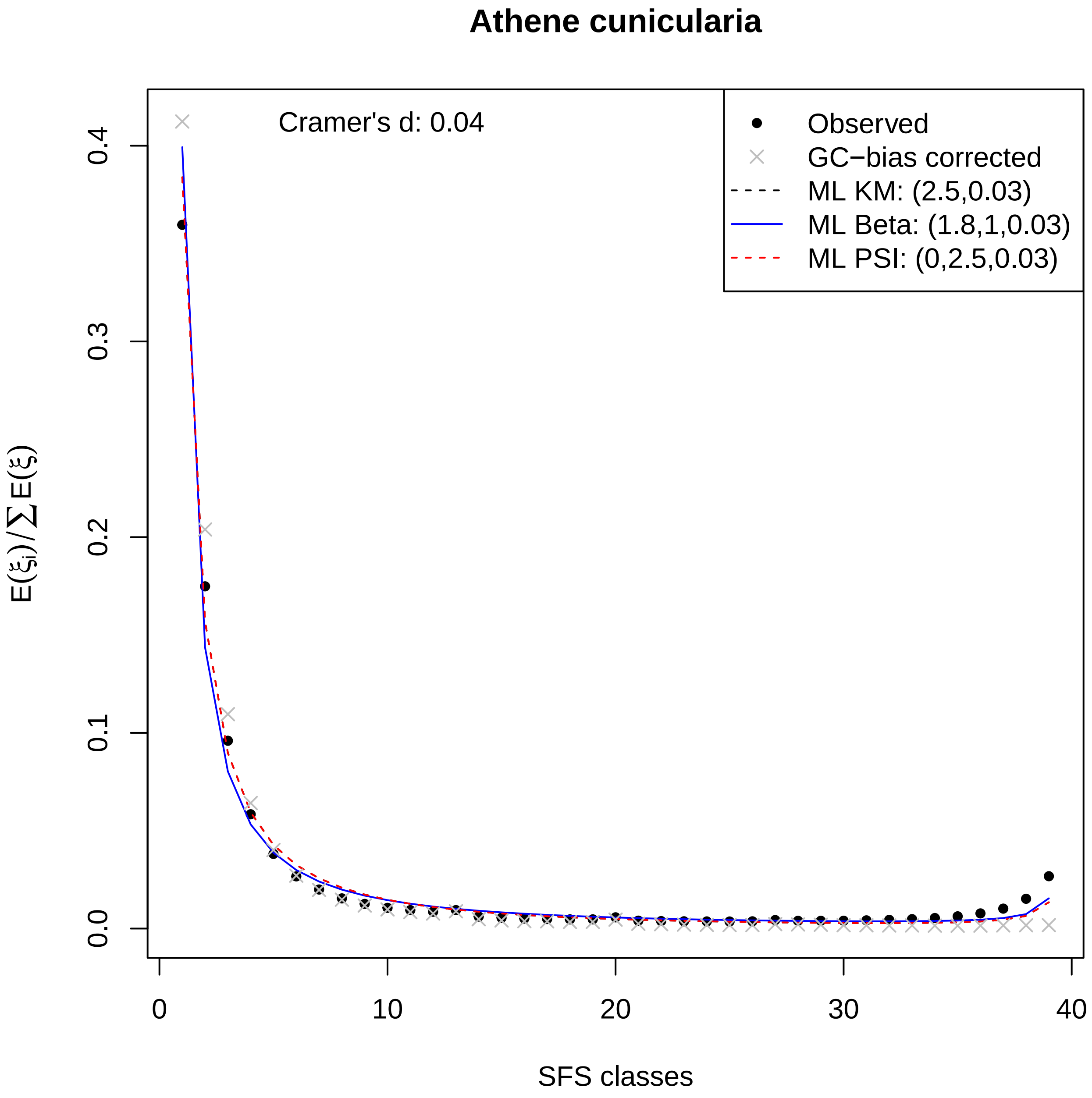
Athene cunicularia
Cramer's d: 0.04
Observed
GC−bias corrected
ML KM: (2.5,0.03)
ML Beta: (1.8,1,0.03)
ML PSI: (0,2.5,0.03)
$E(\xi_i)/\sum E(\xi)$
0.0
0.1
0.2
0.3
0.4
0
10
20
30
40
SFS classes

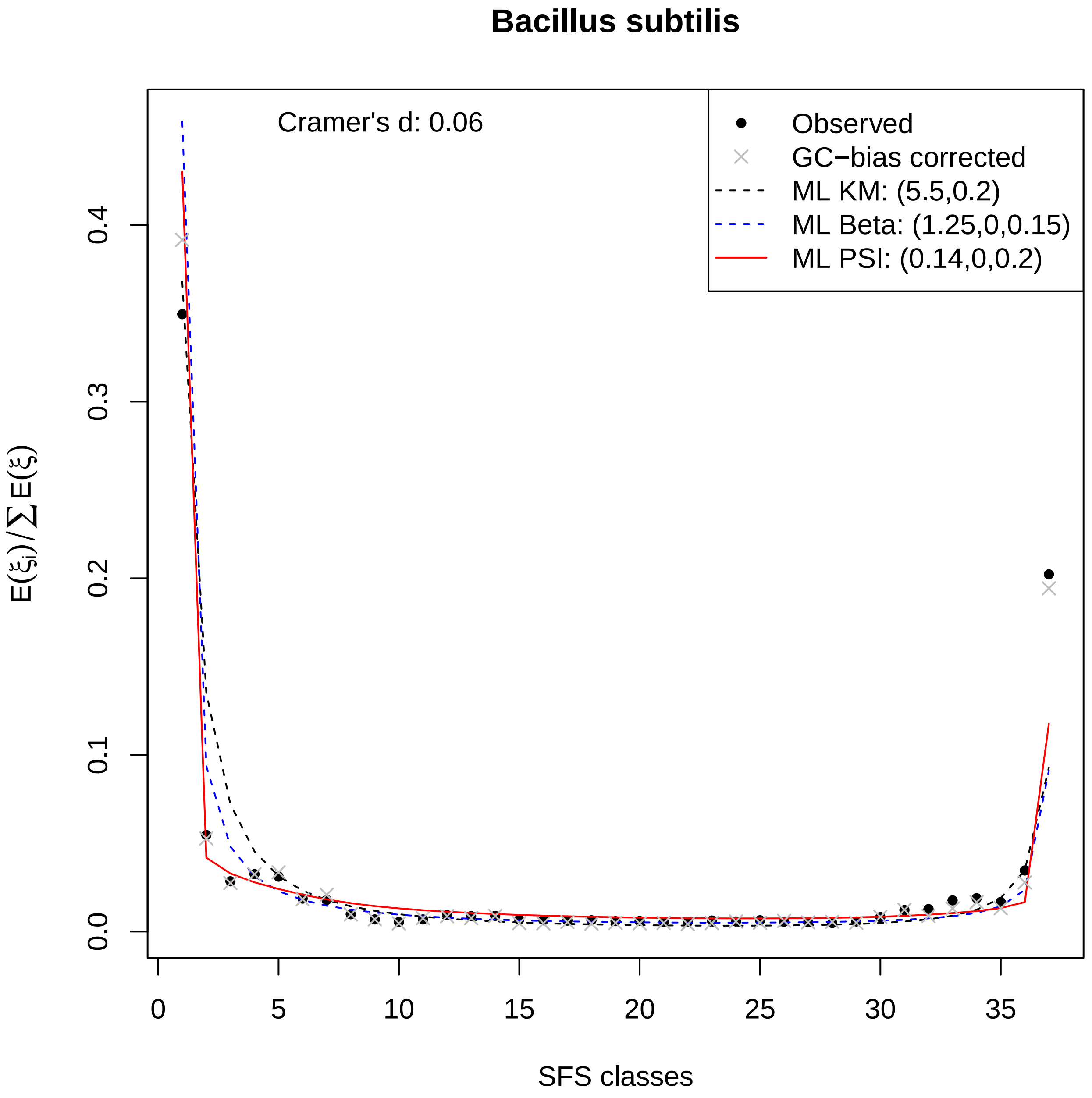

Bacillus subtilis
Cramer's d: 0.06
Observed
GC−bias corrected
ML KM: (5.5,0.2)
ML Beta: (1.25,0,0.15)
ML PSI: (0.14,0,0.2)
0.0
0.1
0.2
0.3
0.4
$E(\xi_i)/\sum E(\xi)$
0
5
10
15
20
25
30
35
SFS classes

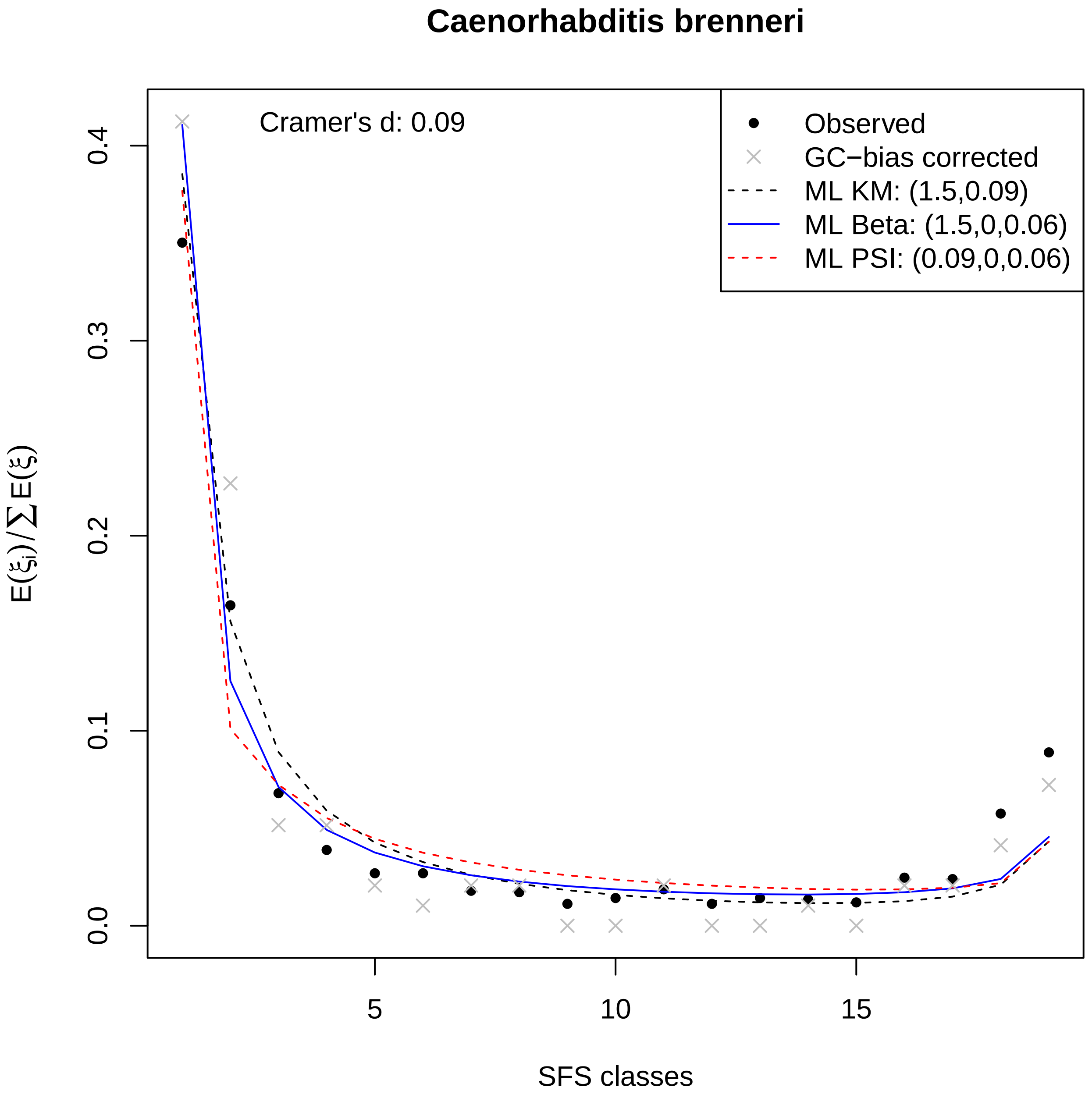
Caenorhabditis brenneri
Cramer's d: 0.09
Observed
GC−bias corrected
ML KM: (1.5,0.09)
ML Beta: (1.5,0,0.06)
ML PSI: (0.09,0,0.06)
0.0
0.1
0.2
0.3
0.4
5
10
15
SFS classes
E(ξi)/∑E(ξ)

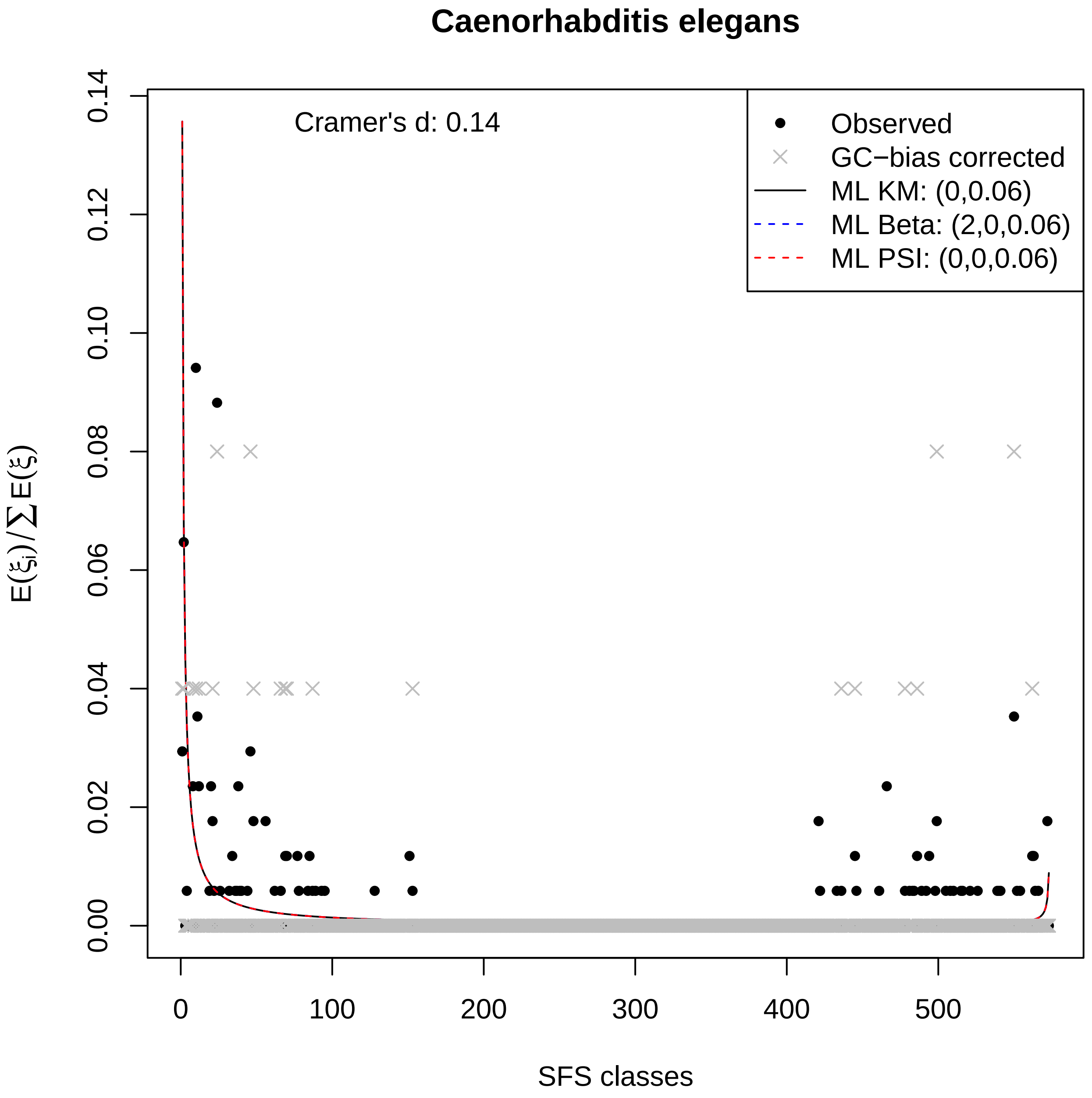
Caenorhabditis elegans
Cramer's d: 0.14
Observed
GC−bias corrected
ML KM: (0,0.06)
ML Beta: (2,0,0.06)
ML PSI: (0,0,0.06)
0.00
0.02
0.04
0.06
0.08
0.10
0.12
0.14
$E(\xi_i)/\sum E(\xi)$
0
100
200
300
400
500
SFS classes

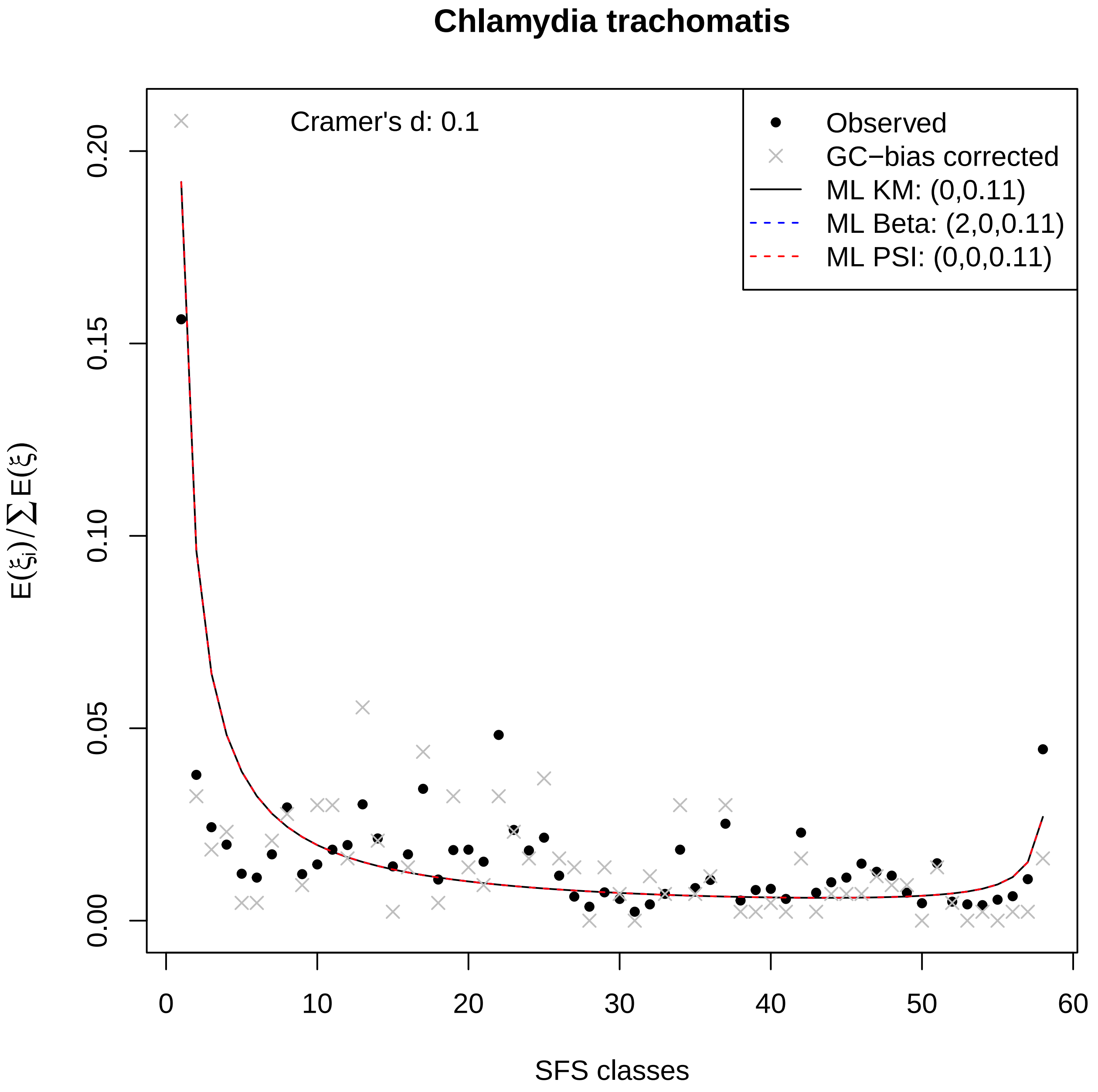

Chlamydia trachomatis
Cramer's d: 0.1
Observed
GC−bias corrected
ML KM: (0,0.11)
ML Beta: (2,0,0.11)
ML PSI: (0,0,0.11)
0.20
0.15
0.10
0.05
0.00
0
10
20
30
40
50
60
SFS classes
E(ξi)/∑E(ξ)

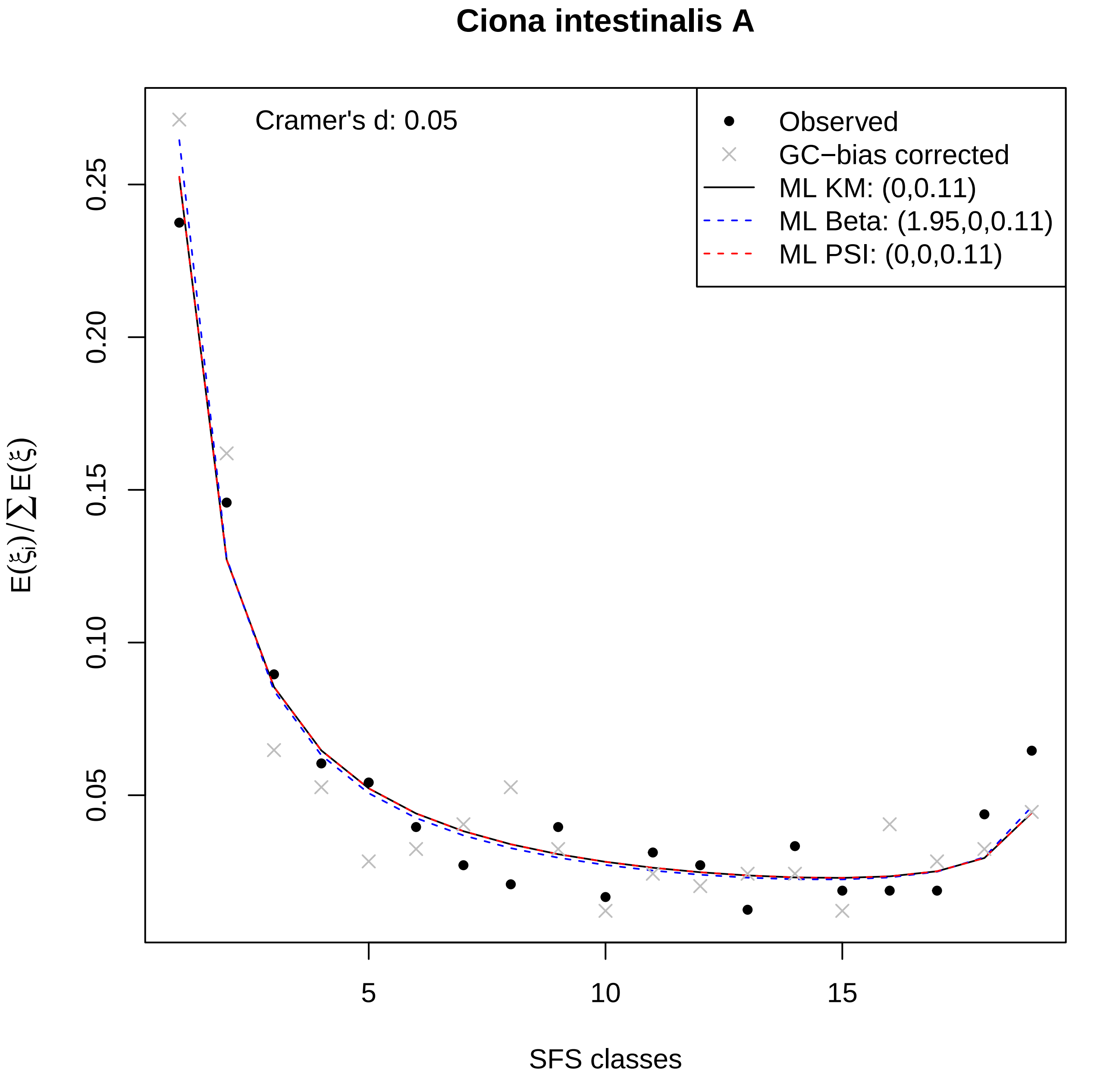

Ciona intestinalis A
Cramer's d: 0.05
Observed
GC−bias corrected
ML KM: (0,0.11)
ML Beta: (1.95,0,0.11)
ML PSI: (0,0,0.11)
0.05
0.10
0.15
0.20
0.25
5
10
15
SFS classes
$E(\xi_i)/\sum E(\xi)$

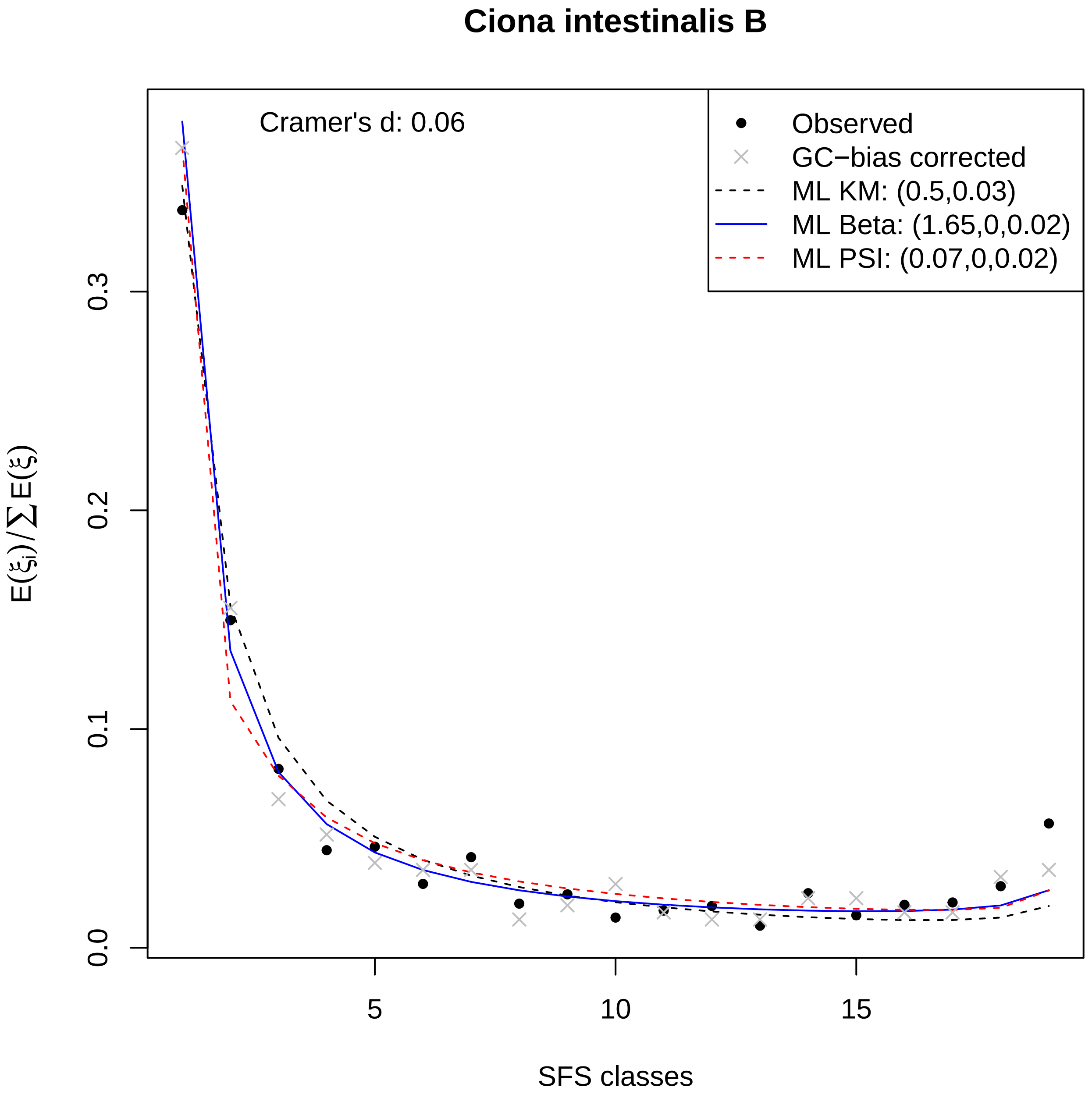

Ciona intestinalis B
Cramer's d: 0.06
Observed
GC−bias corrected
ML KM: (0.5,0.03)
ML Beta: (1.65,0,0.02)
ML PSI: (0.07,0,0.02)
0.0
0.1
0.2
0.3
5
10
15
SFS classes
$E(\xi_i)/\sum E(\xi)$

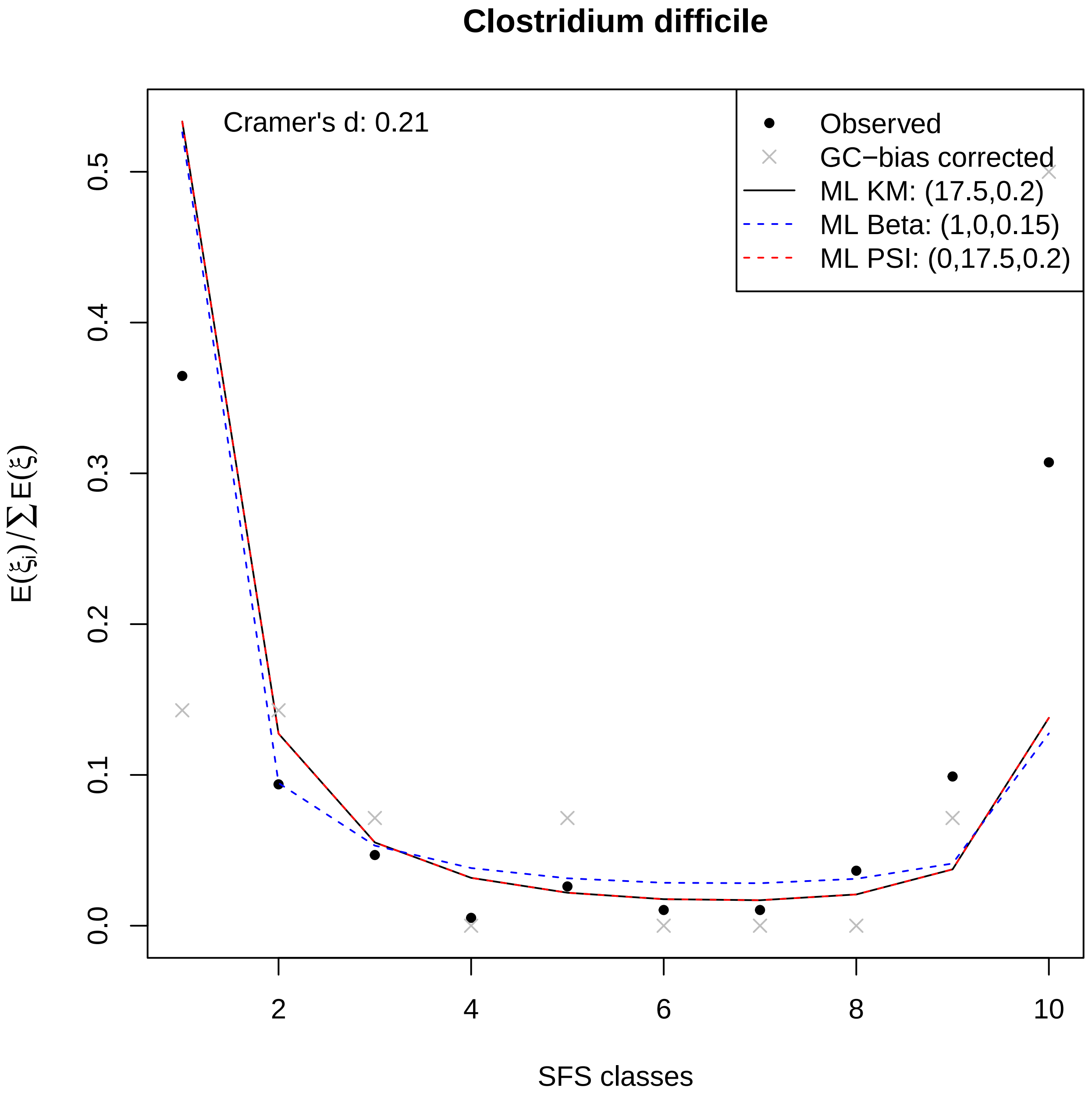

Clostridium difficile
Cramer's d: 0.21
Observed
GC−bias corrected
ML KM: (17.5,0.2)
ML Beta: (1,0,0.15)
ML PSI: (0,17.5,0.2)
0.0
0.1
0.2
0.3
0.4
0.5
2
4
6
8
10
SFS classes

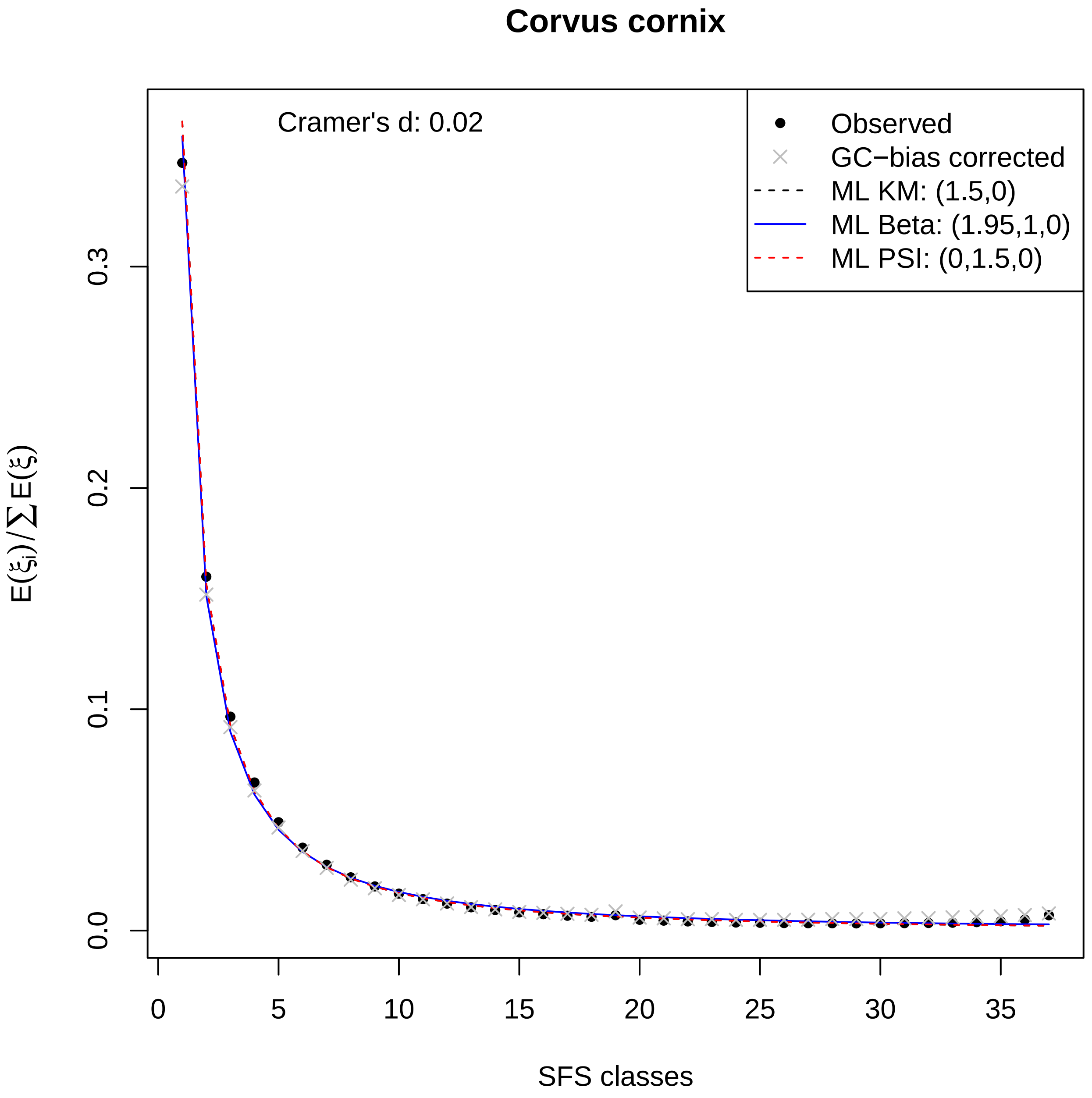

Corvus cornix
Cramer's d: 0.02
Observed
GC−bias corrected
ML KM: (1.5,0)
ML Beta: (1.95,1,0)
ML PSI: (0,1.5,0)
0.0
0.1
0.2
0.3
0
5
10
15
20
25
30
35
SFS classes
$E(\xi_i)/\sum E(\xi)$

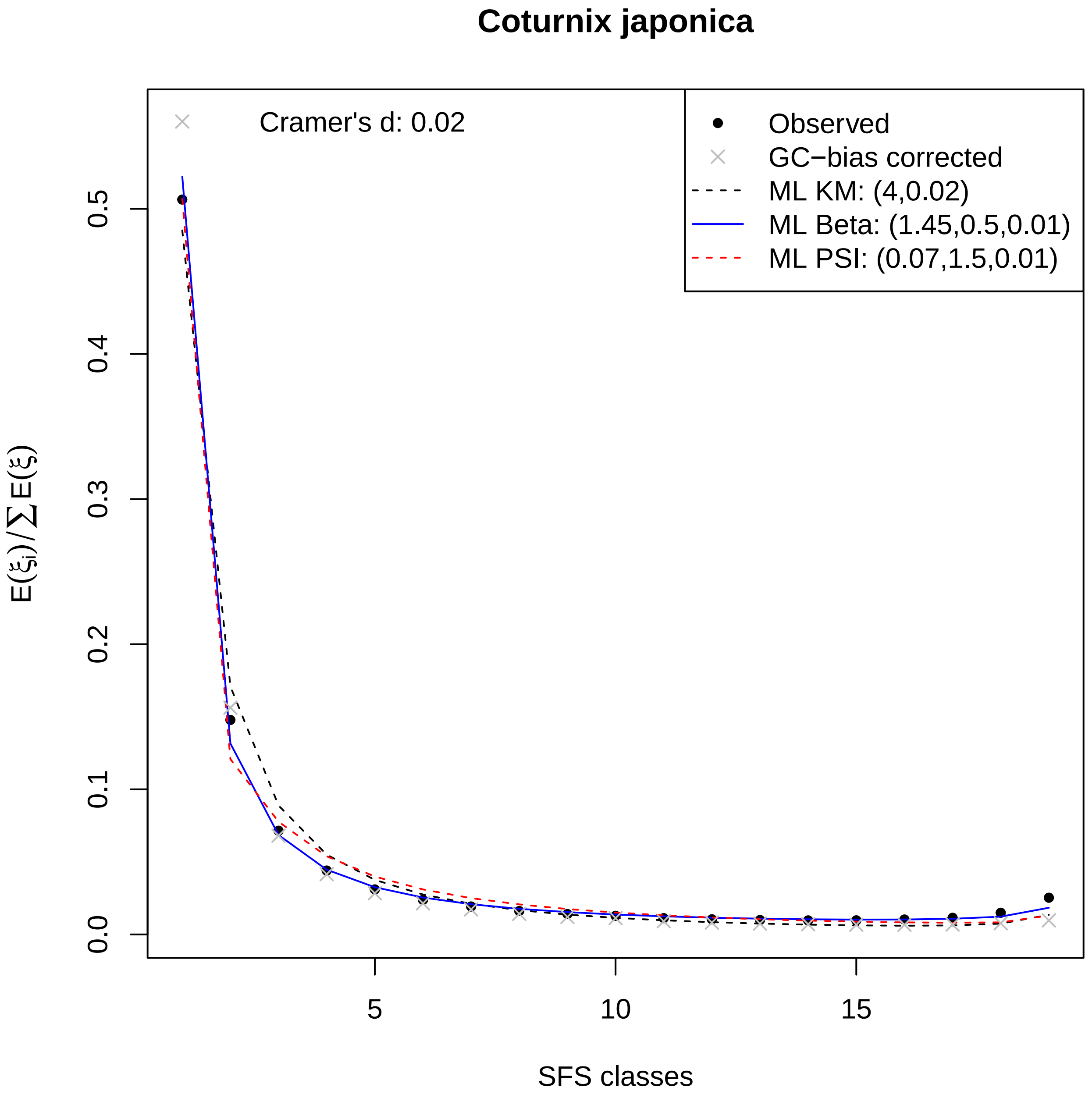

Coturnix japonica
Cramer's d: 0.02
Observed
GC−bias corrected
ML KM: (4,0.02)
ML Beta: (1.45,0.5,0.01)
ML PSI: (0.07,1.5,0.01)
$E(\xi_i)/\sum E(\xi)$
0.0
0.1
0.2
0.3
0.4
0.5
5
10
15
SFS classes

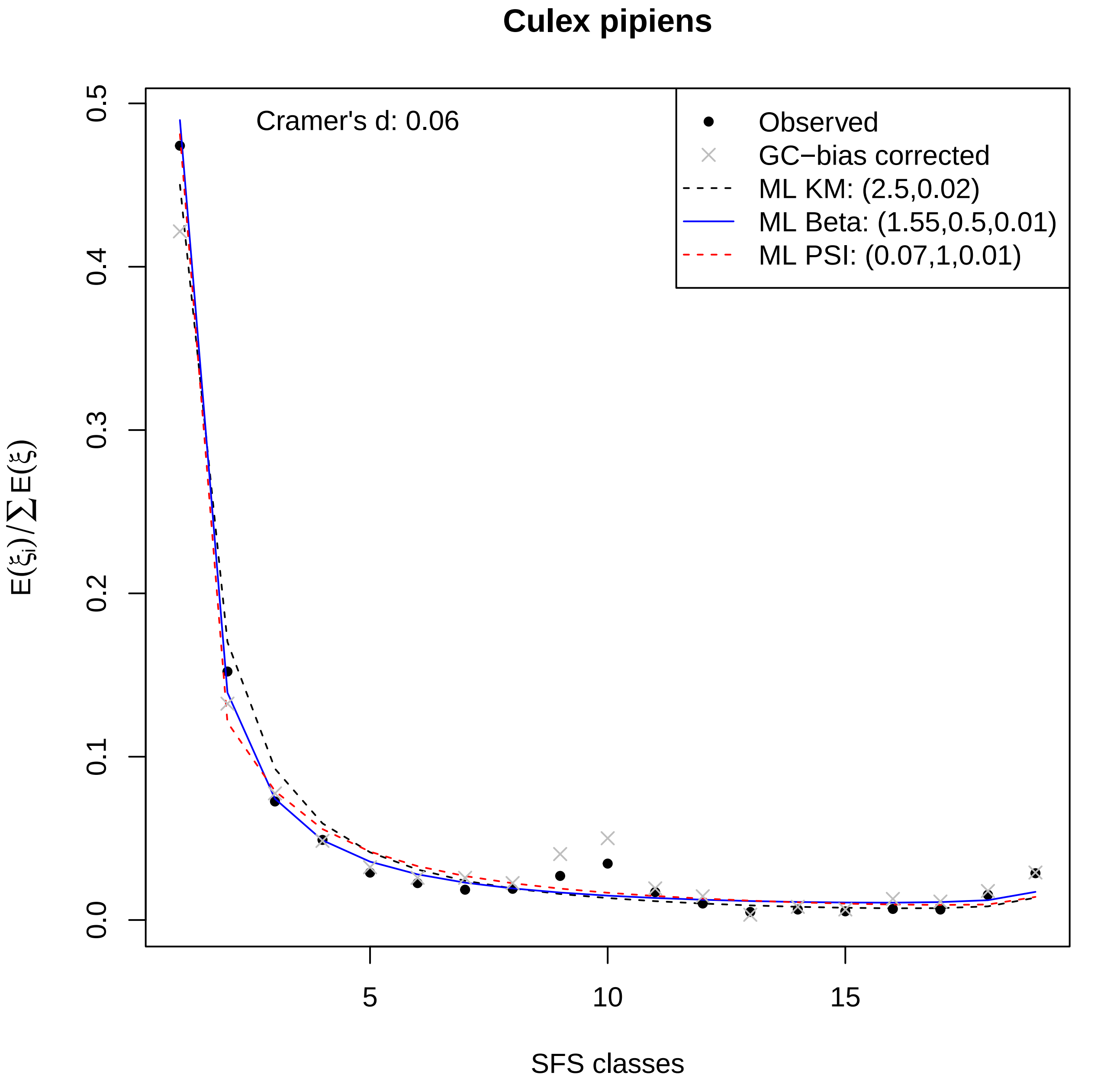

Culex pipiens
Cramer's d: 0.06
Observed
GC−bias corrected
ML KM: (2.5,0.02)
ML Beta: (1.55,0.5,0.01)
ML PSI: (0.07,1,0.01)
$E(\xi_i)/\sum E(\xi)$
SFS classes
0.0
0.1
0.2
0.3
0.4
0.5
5
10
15

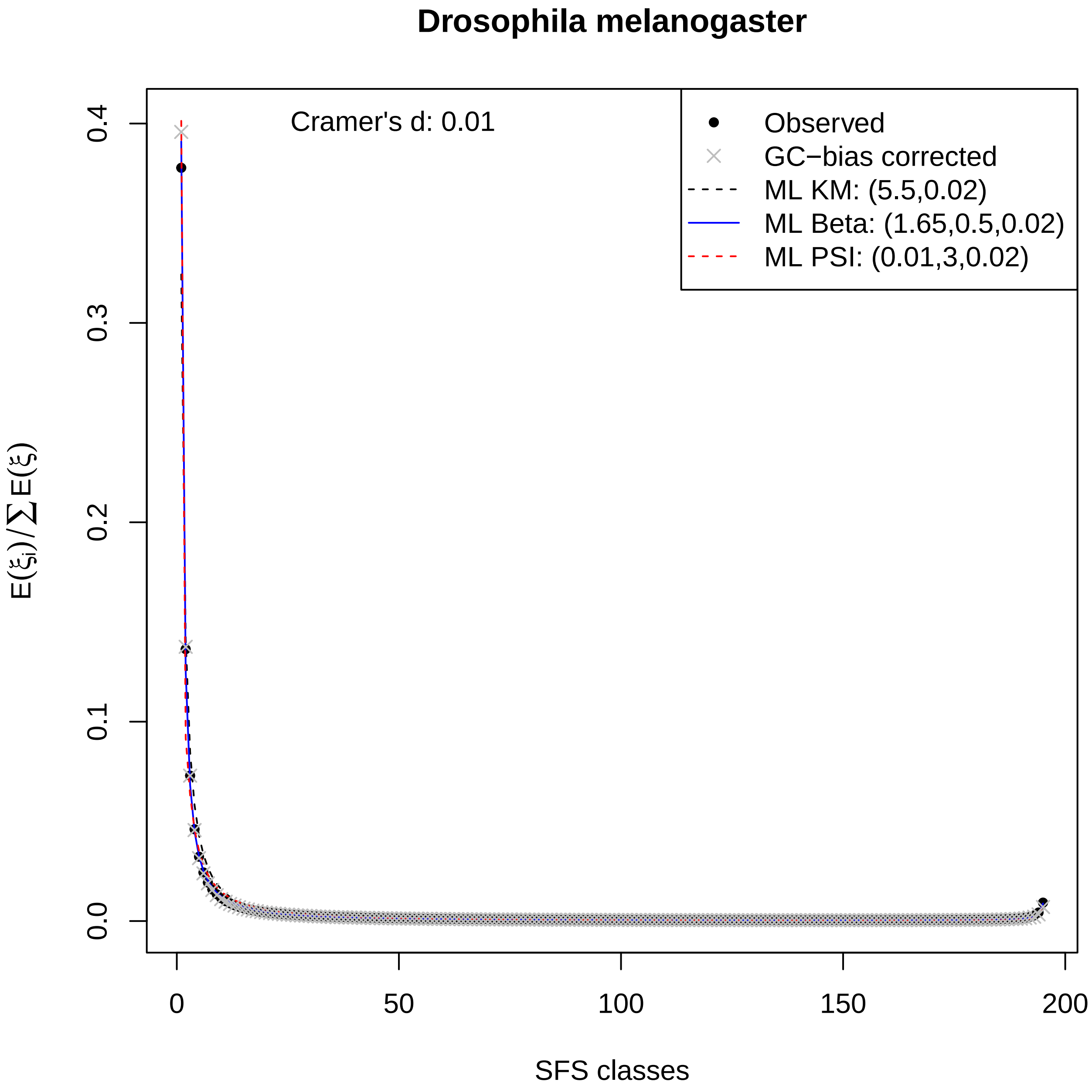

Drosophila melanogaster
Cramer's d: 0.01
Observed
GC−bias corrected
ML KM: (5.5,0.02)
ML Beta: (1.65,0.5,0.02)
ML PSI: (0.01,3,0.02)
0.0
0.1
0.2
0.3
0.4
0
50
100
150
200
SFS classes
$E(\xi_i)/\sum E(\xi)$

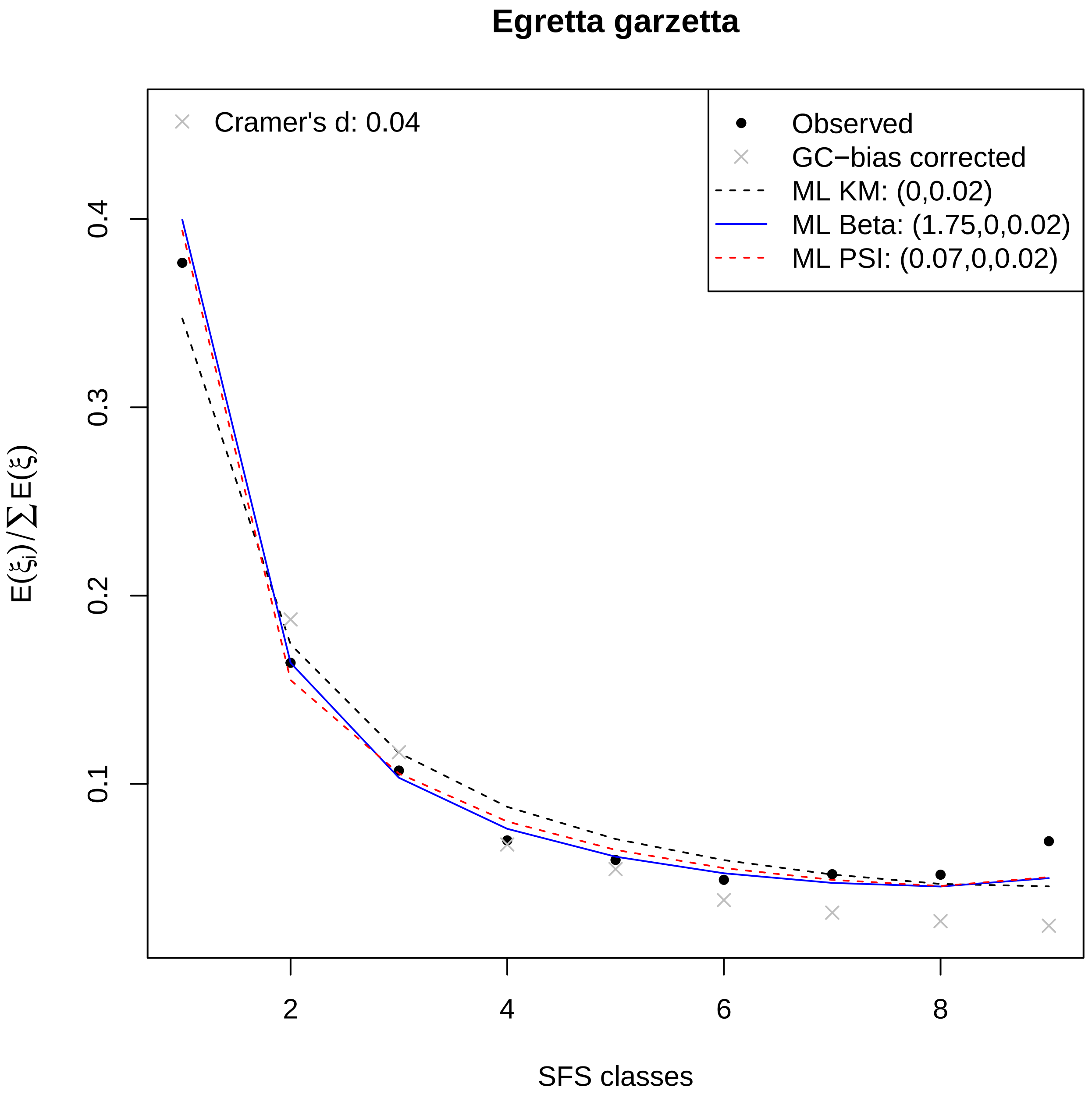
Egretta garzetta
Cramer's d: 0.04
Observed
GC−bias corrected
ML KM: (0,0.02)
ML Beta: (1.75,0,0.02)
ML PSI: (0.07,0,0.02)
$E(\xi_i)/\sum E(\xi)$
0.1
0.2
0.3
0.4
2
4
6
8
SFS classes

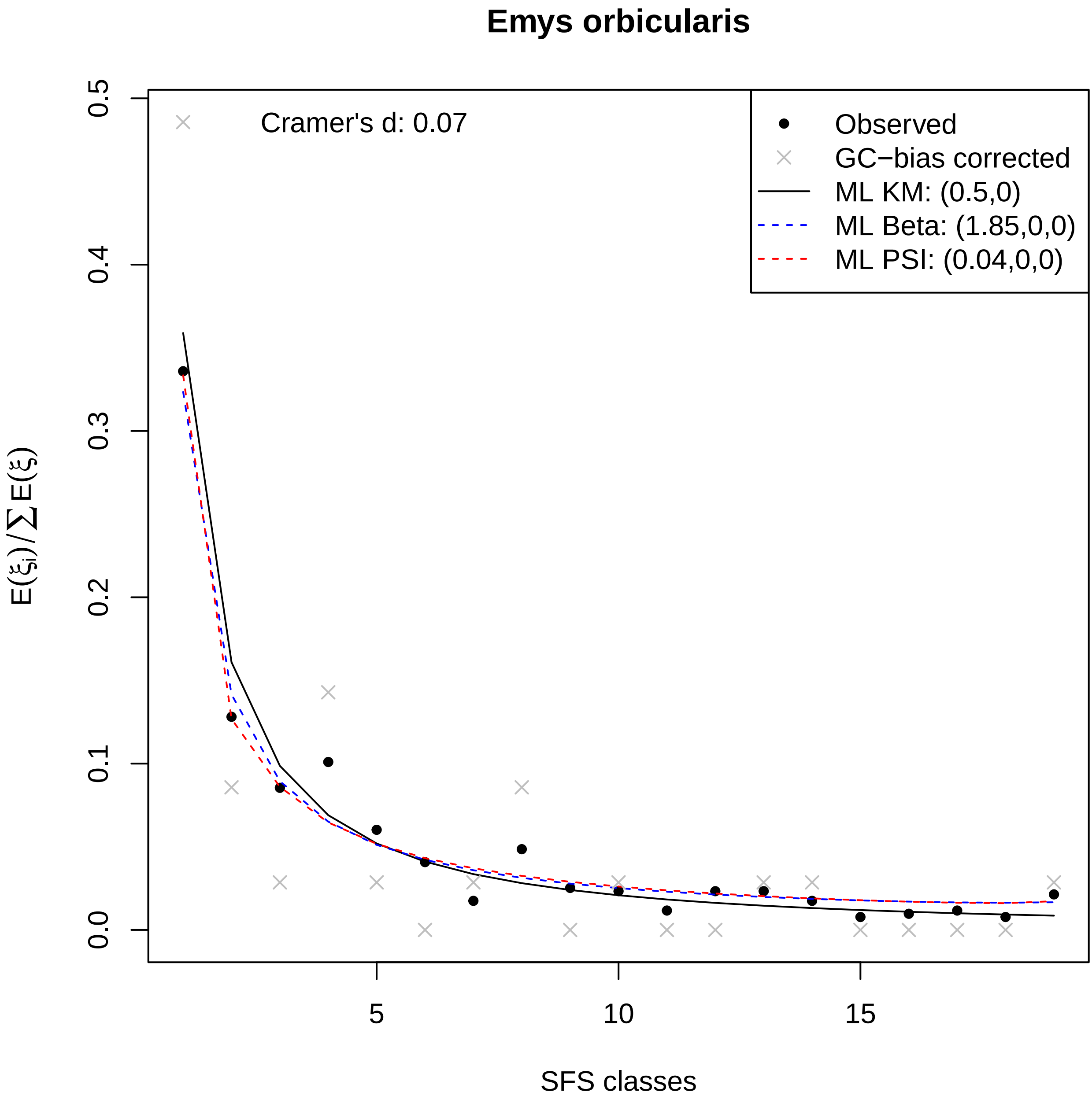
Emys orbicularis
Cramer's d: 0.07
Observed
GC−bias corrected
ML KM: (0.5,0)
ML Beta: (1.85,0,0)
ML PSI: (0.04,0,0)
$E(\xi_i)/\sum E(\xi)$
SFS classes
0.0
0.1
0.2
0.3
0.4
0.5
5
10
15

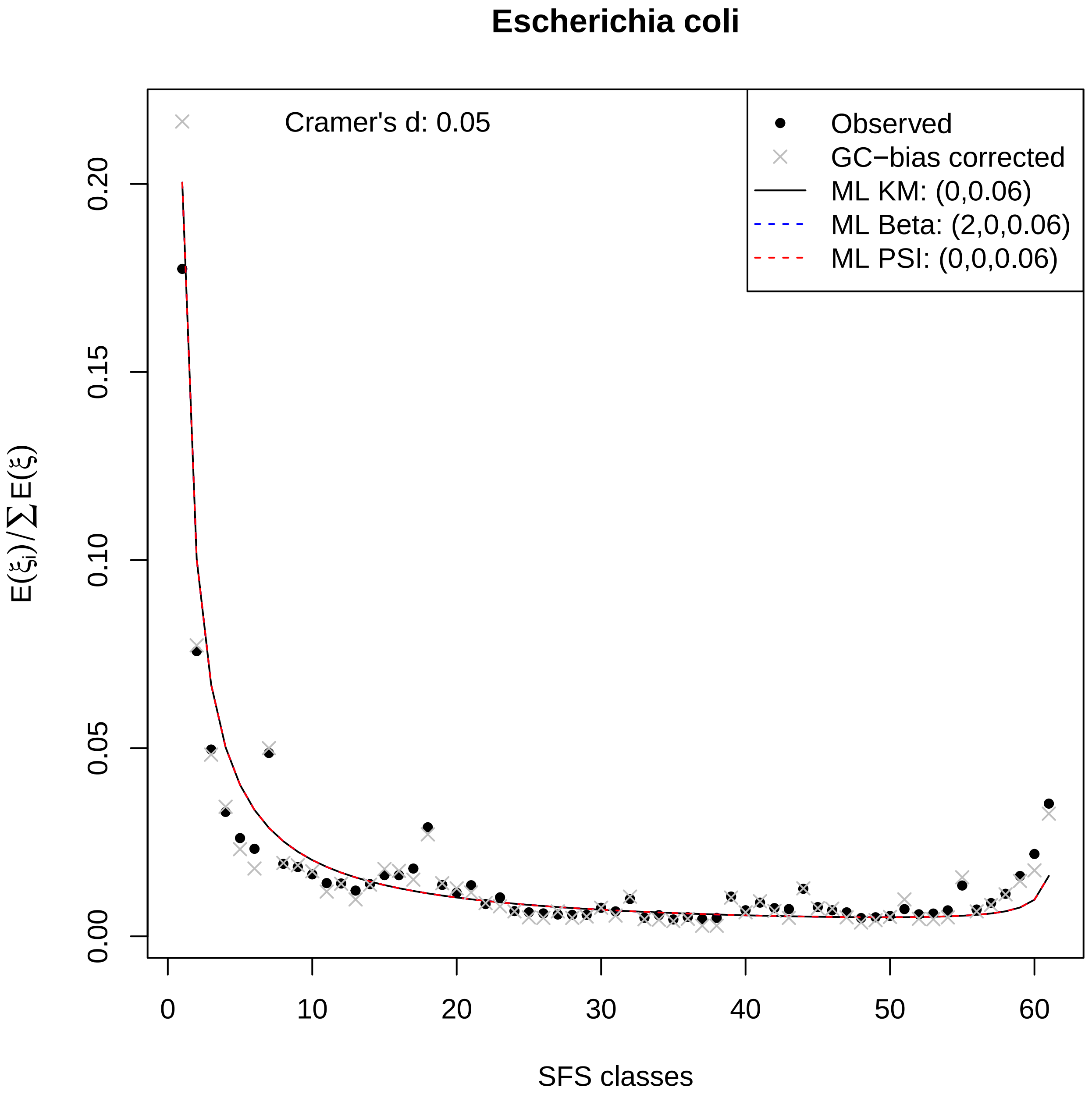

Escherichia coli
Cramer's d: 0.05
Observed
GC−bias corrected
ML KM: (0,0.06)
ML Beta: (2,0,0.06)
ML PSI: (0,0,0.06)
0.00
0.05
0.10
0.15
0.20
0
10
20
30
40
50
60
$E(\xi_i)/\sum E(\xi)$
SFS classes

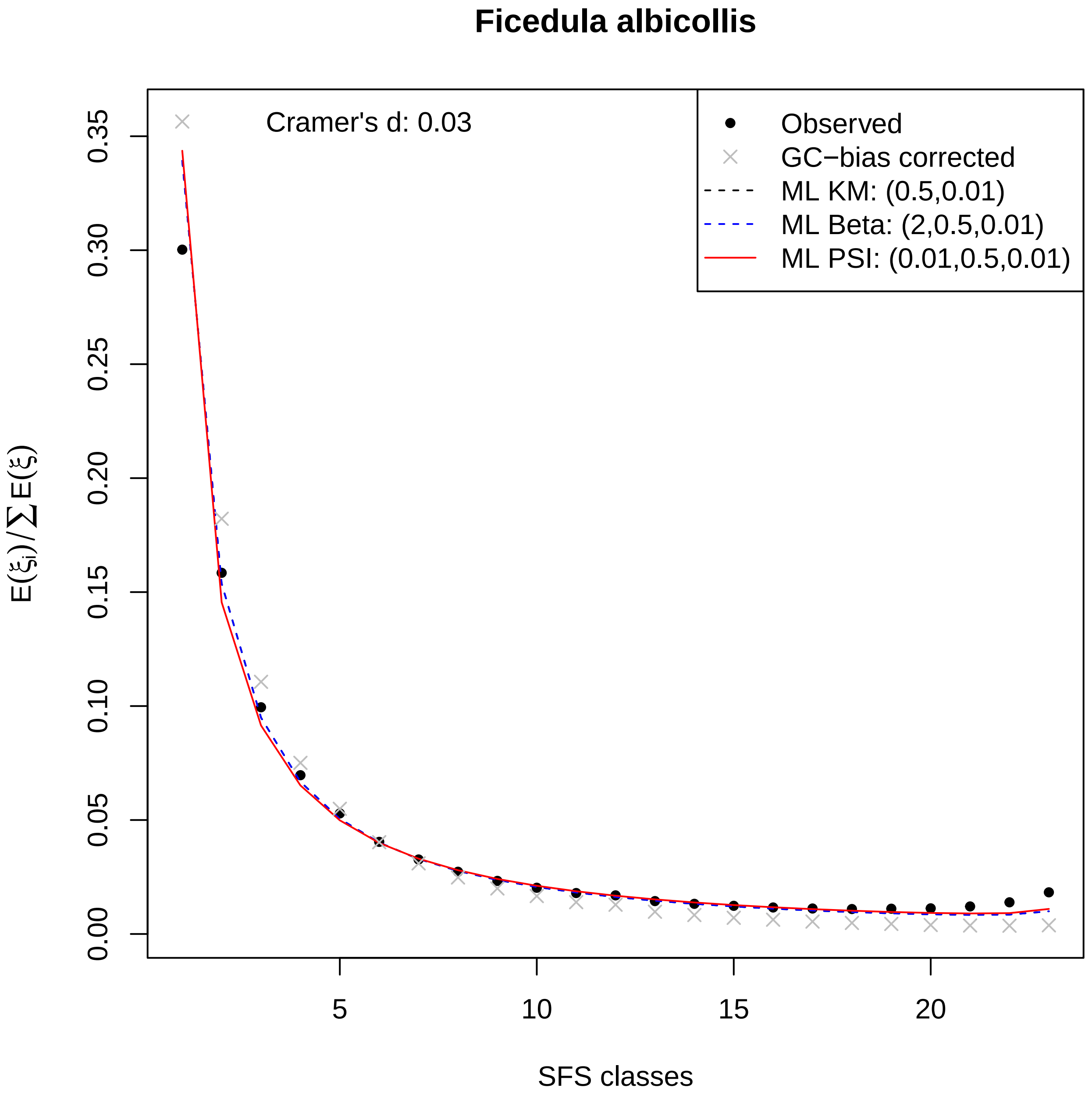

Ficedula albicollis
Cramer's d: 0.03
Observed
GC−bias corrected
ML KM: (0.5,0.01)
ML Beta: (2,0.5,0.01)
ML PSI: (0.01,0.5,0.01)
0.00
0.05
0.10
0.15
0.20
0.25
0.30
0.35
5
10
15
20
SFS classes
$E(\xi_i)/\sum E(\xi)$

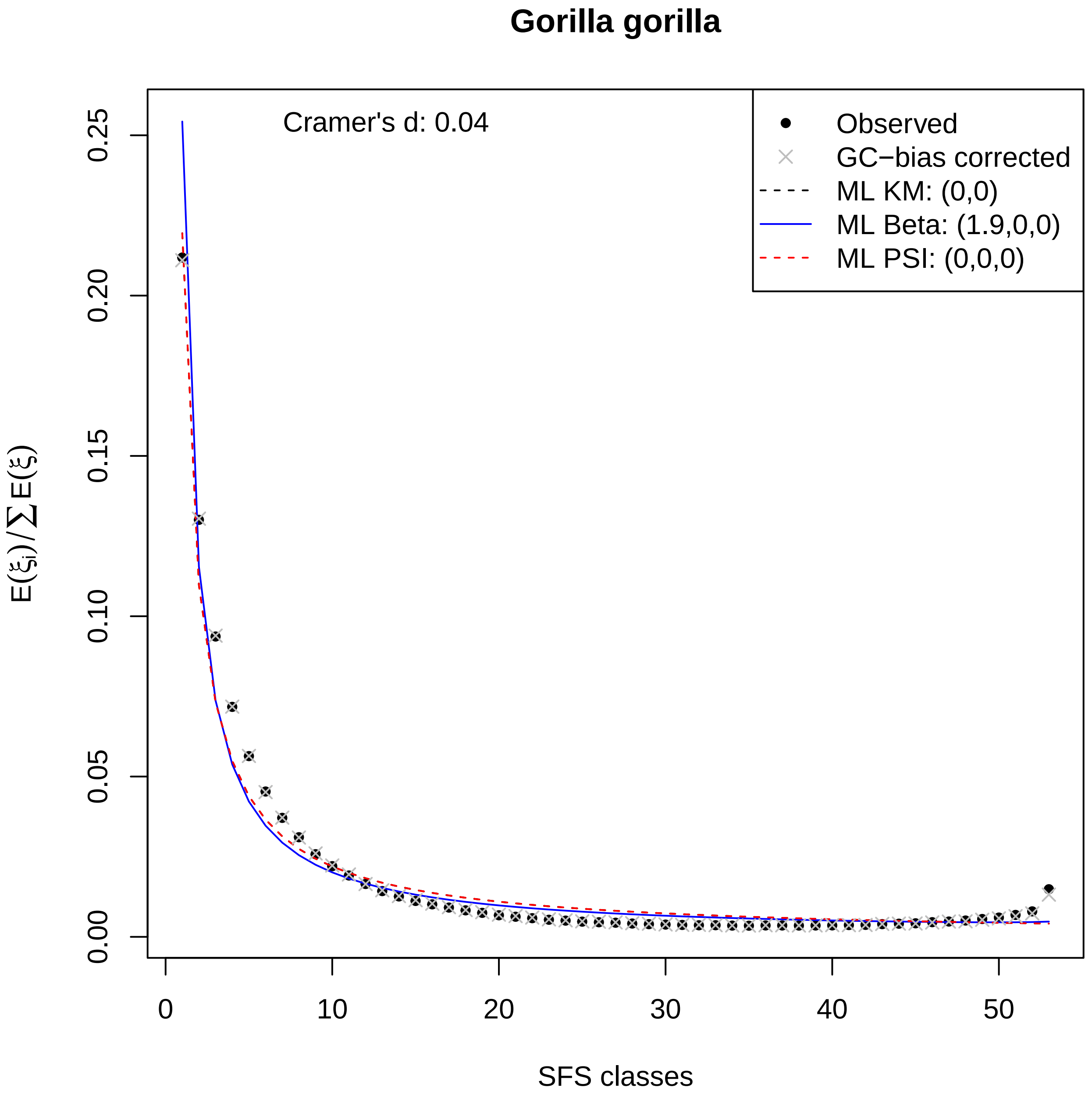
Gorilla gorilla
Cramer's d: 0.04
Observed
GC−bias corrected
ML KM: (0,0)
ML Beta: (1.9,0,0)
ML PSI: (0,0,0)
$E(\xi_i)/\sum E(\xi)$
SFS classes
0.00
0.05
0.10
0.15
0.20
0.25
0
10
20
30
40
50

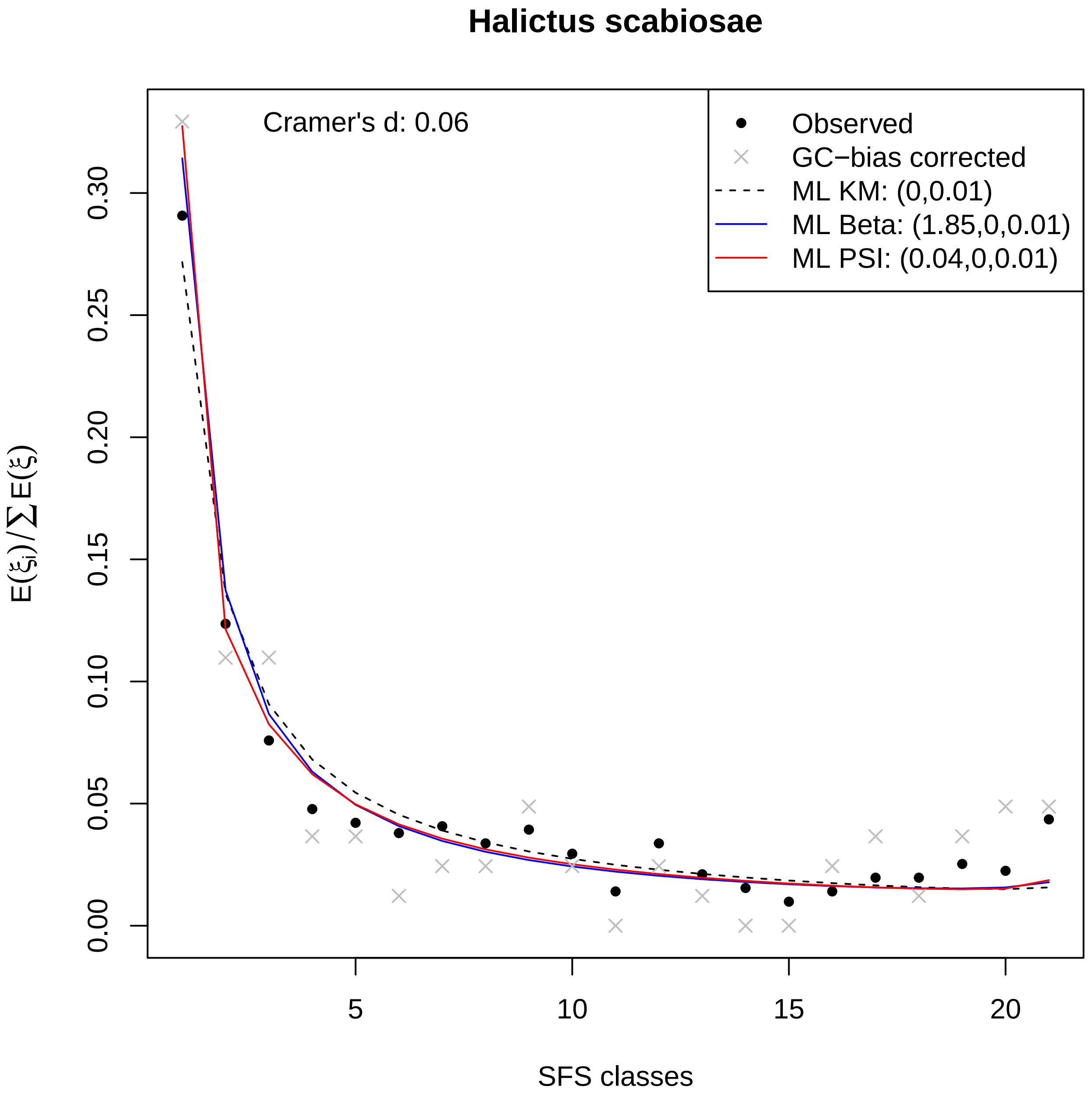

Halictus scabiosae
Cramer's d: 0.06
Observed
GC−bias corrected
ML KM: (0,0.01)
ML Beta: (1.85,0,0.01)
ML PSI: (0.04,0,0.01)
$E(\xi_i)/\sum E(\xi)$
0.00
0.05
0.10
0.15
0.20
0.25
0.30
5
10
15
20
SFS classes

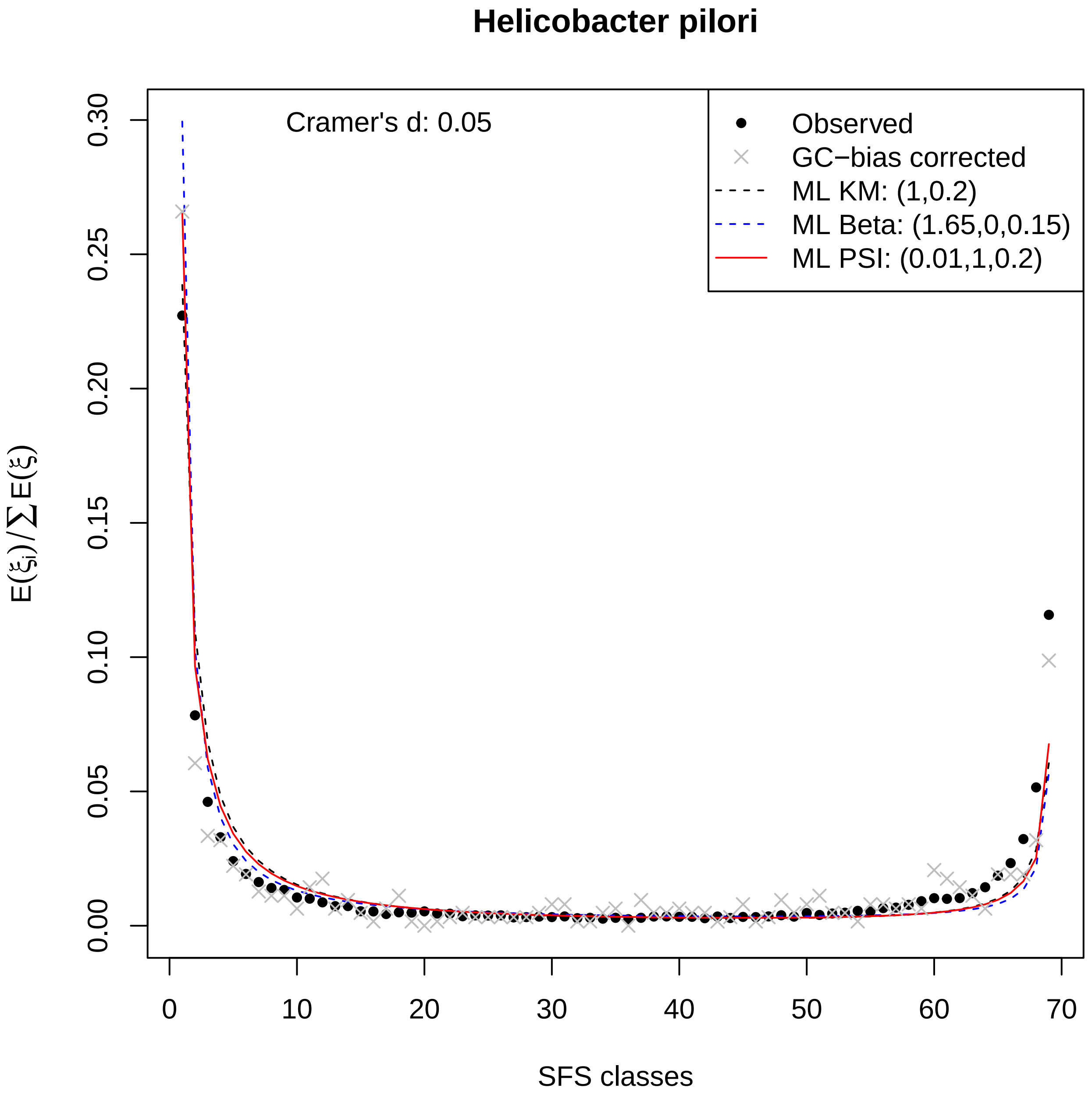

Helicobacter pilori
Cramer's d: 0.05
Observed
GC−bias corrected
ML KM: (1,0.2)
ML Beta: (1.65,0,0.15)
ML PSI: (0.01,1,0.2)
$E(\xi_i)/\sum E(\xi)$
SFS classes
0.00
0.05
0.10
0.15
0.20
0.25
0.30
0
10
20
30
40
50
60
70

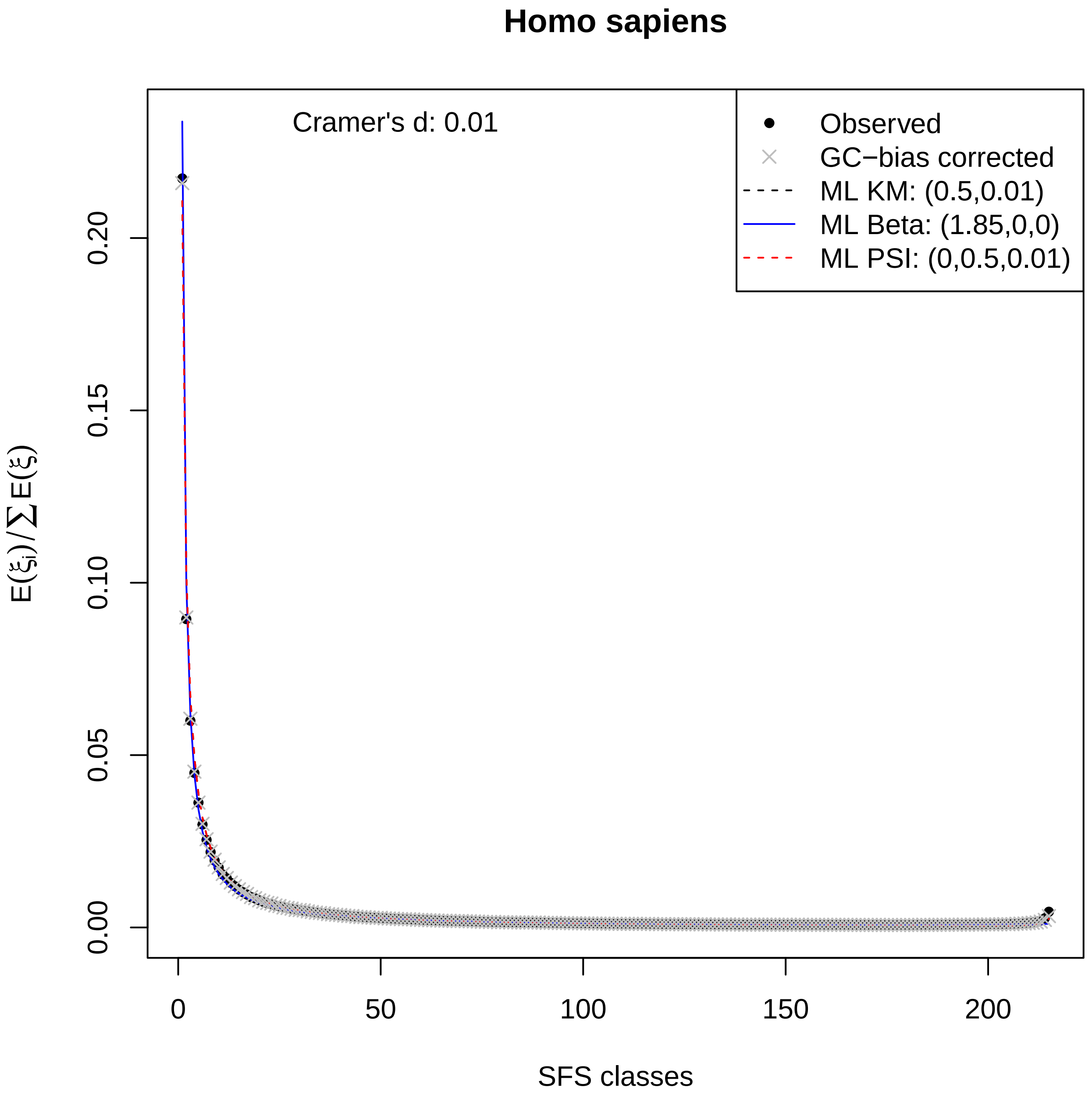
Homo sapiens
Cramer's d: 0.01
Observed
GC−bias corrected
ML KM: (0.5,0.01)
ML Beta: (1.85,0,0)
ML PSI: (0,0.5,0.01)
$E(\xi_i)/\sum E(\xi)$
0.00
0.05
0.10
0.15
0.20
0
50
100
150
200
SFS classes

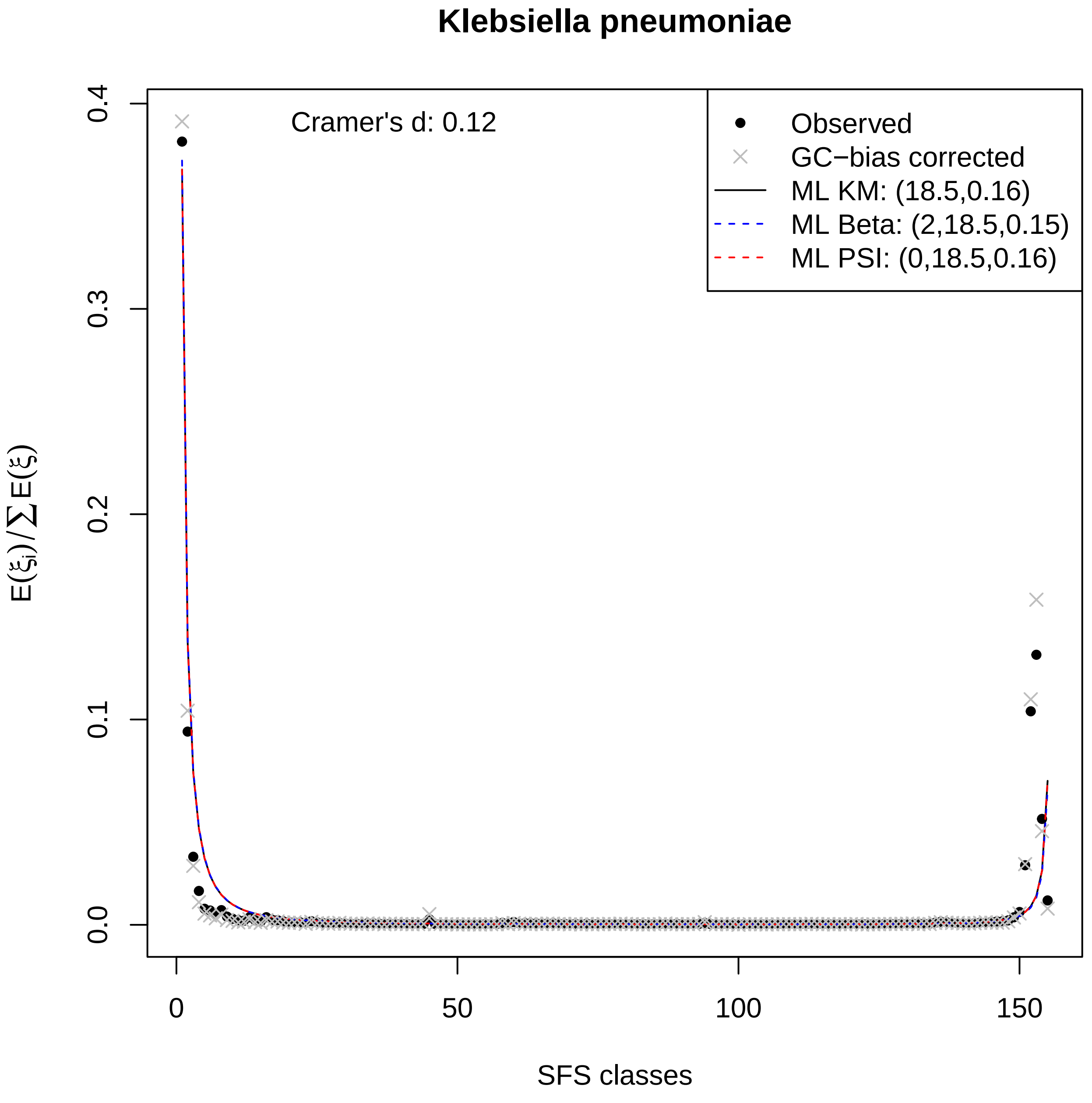

Klebsiella pneumoniae
Cramer's d: 0.12
Observed
GC−bias corrected
ML KM: (18.5,0.16)
ML Beta: (2,18.5,0.15)
ML PSI: (0,18.5,0.16)
0.0
0.1
0.2
0.3
0.4
$E(\xi_i)/\sum E(\xi)$
0
50
100
150
SFS classes

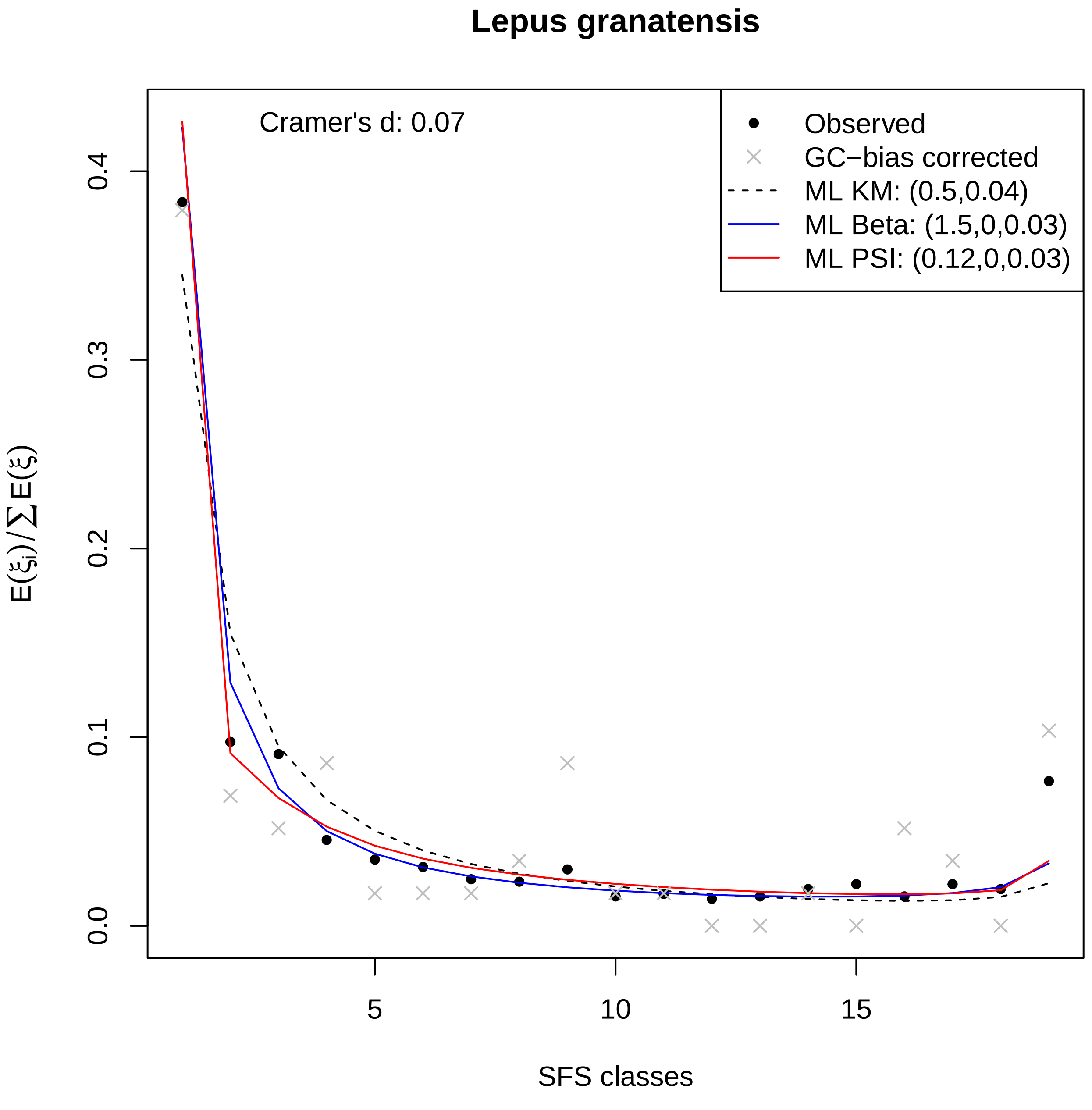

Lepus granatensis
Cramer's d: 0.07
Observed
GC−bias corrected
ML KM: (0.5,0.04)
ML Beta: (1.5,0,0.03)
ML PSI: (0.12,0,0.03)
0.0
0.1
0.2
0.3
0.4
5
10
15
SFS classes
E(ξi)/∑E(ξ)

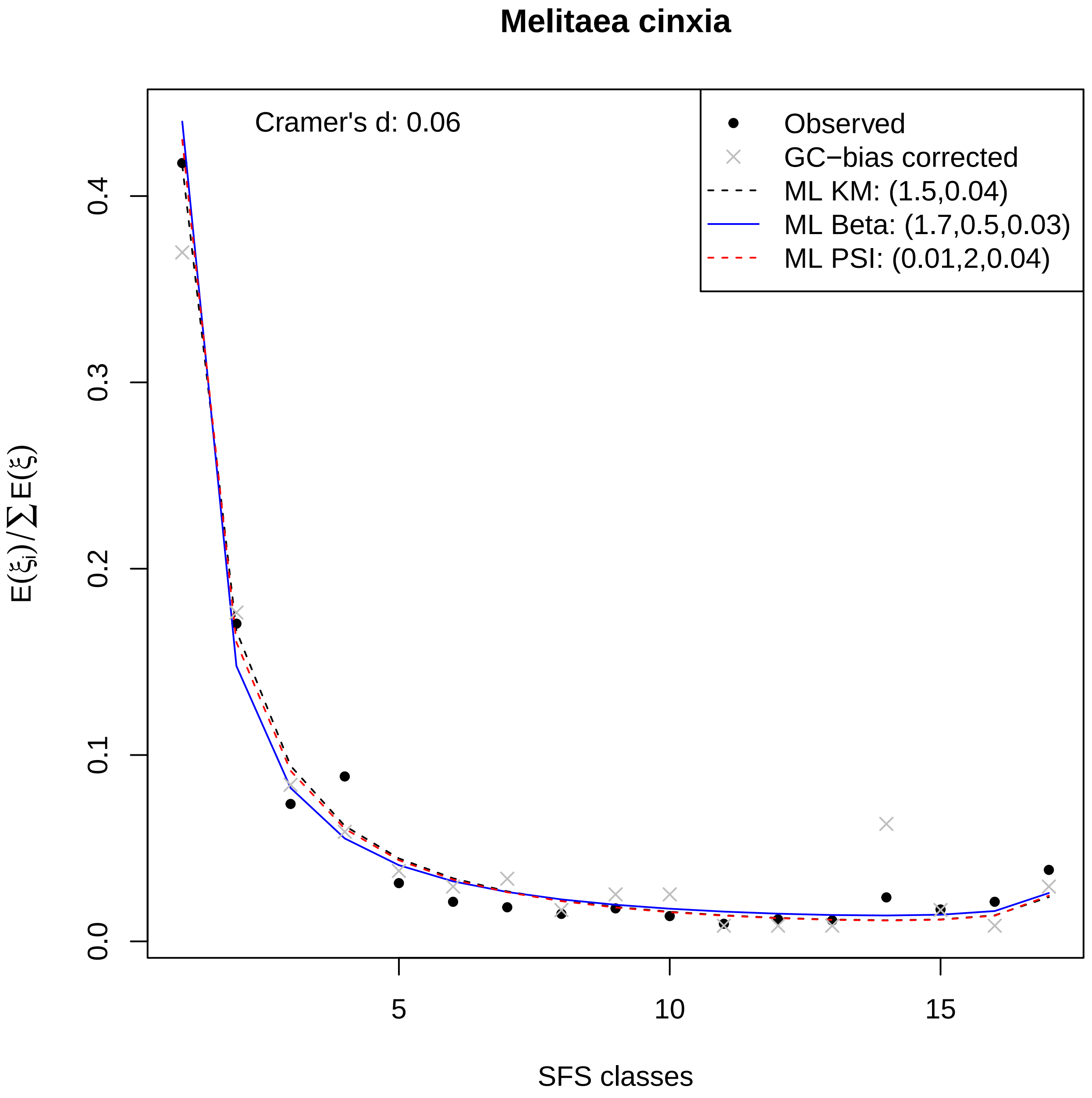
Melitaea cinxia
Cramer's d: 0.06
Observed
GC−bias corrected
ML KM: (1.5,0.04)
ML Beta: (1.7,0.5,0.03)
ML PSI: (0.01,2,0.04)
0.0
0.1
0.2
0.3
0.4
5
10
15
SFS classes
$E(\xi_i)/\sum E(\xi)$

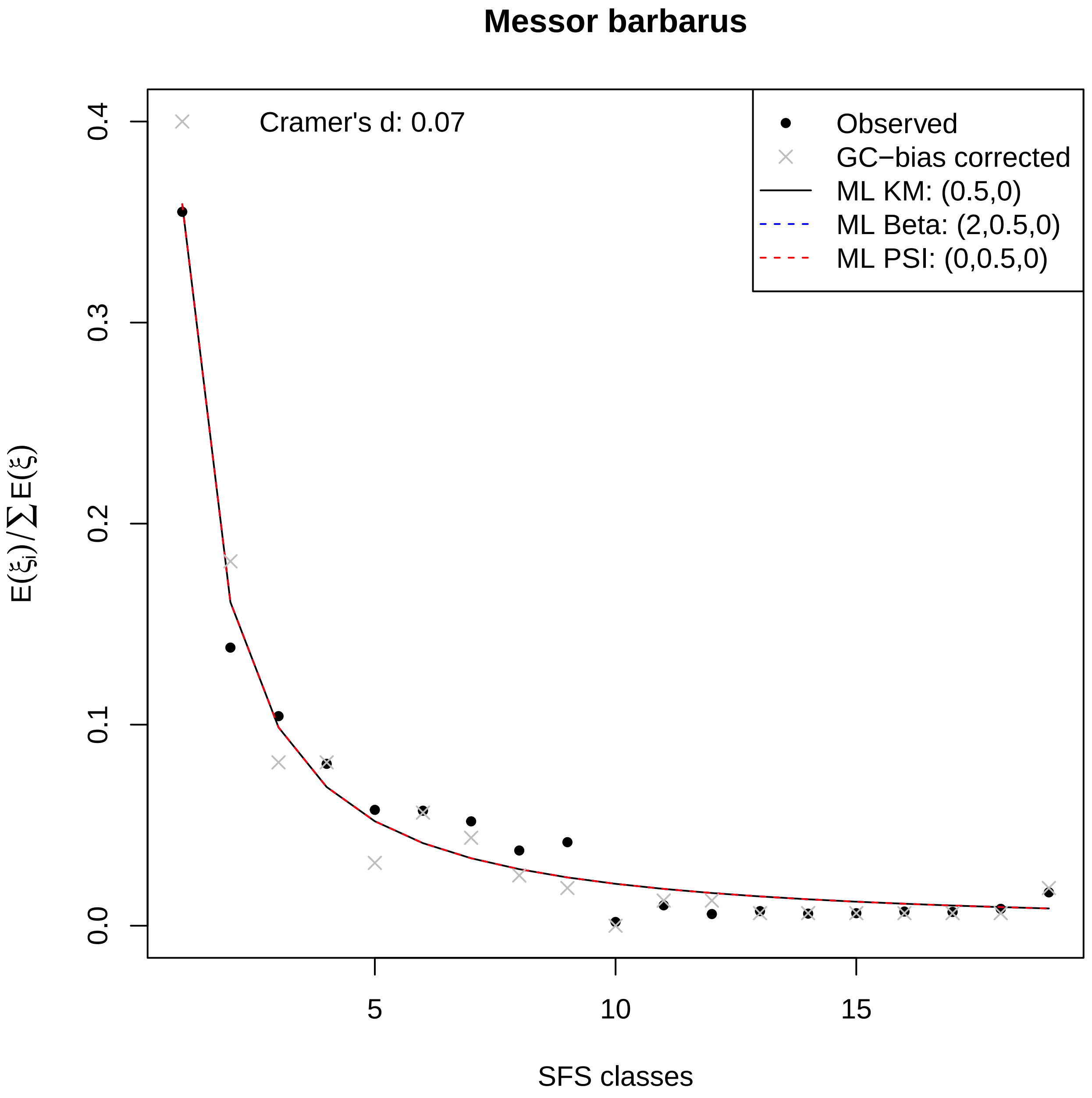

Messor barbarus
Cramer's d: 0.07
Observed
GC−bias corrected
ML KM: (0.5,0)
ML Beta: (2,0.5,0)
ML PSI: (0,0.5,0)
$E(\xi_i)/\sum E(\xi)$
0.0
0.1
0.2
0.3
0.4
5
10
15
SFS classes

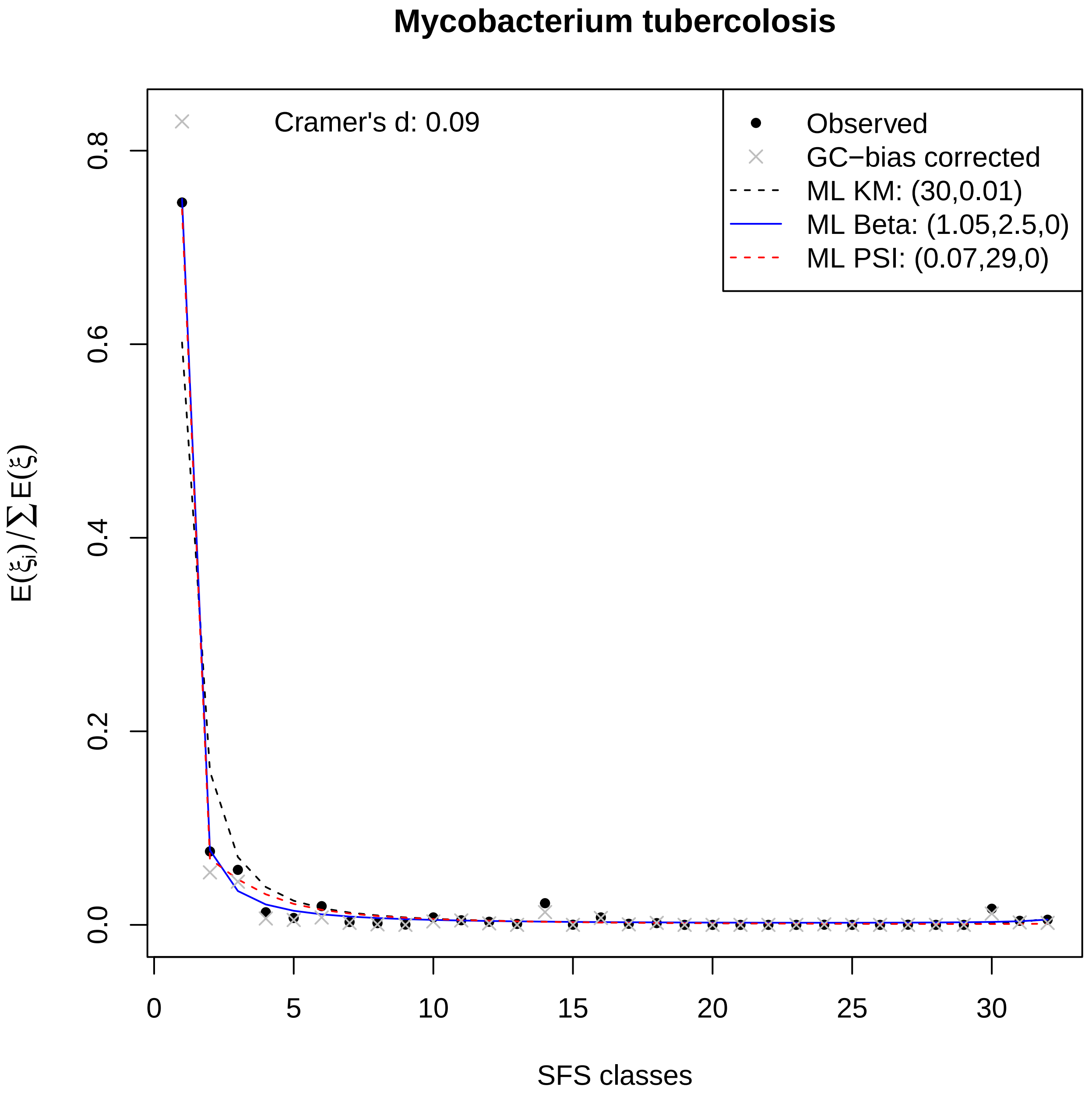
Mycobacterium tubercolosis
Cramer's d: 0.09
Observed
GC−bias corrected
ML KM: (30,0.01)
ML Beta: (1.05,2.5,0)
ML PSI: (0.07,29,0)
0.0
0.2
0.4
0.6
0.8
0
5
10
15
20
25
30
SFS classes
$E(\xi_i)/\sum E(\xi)$

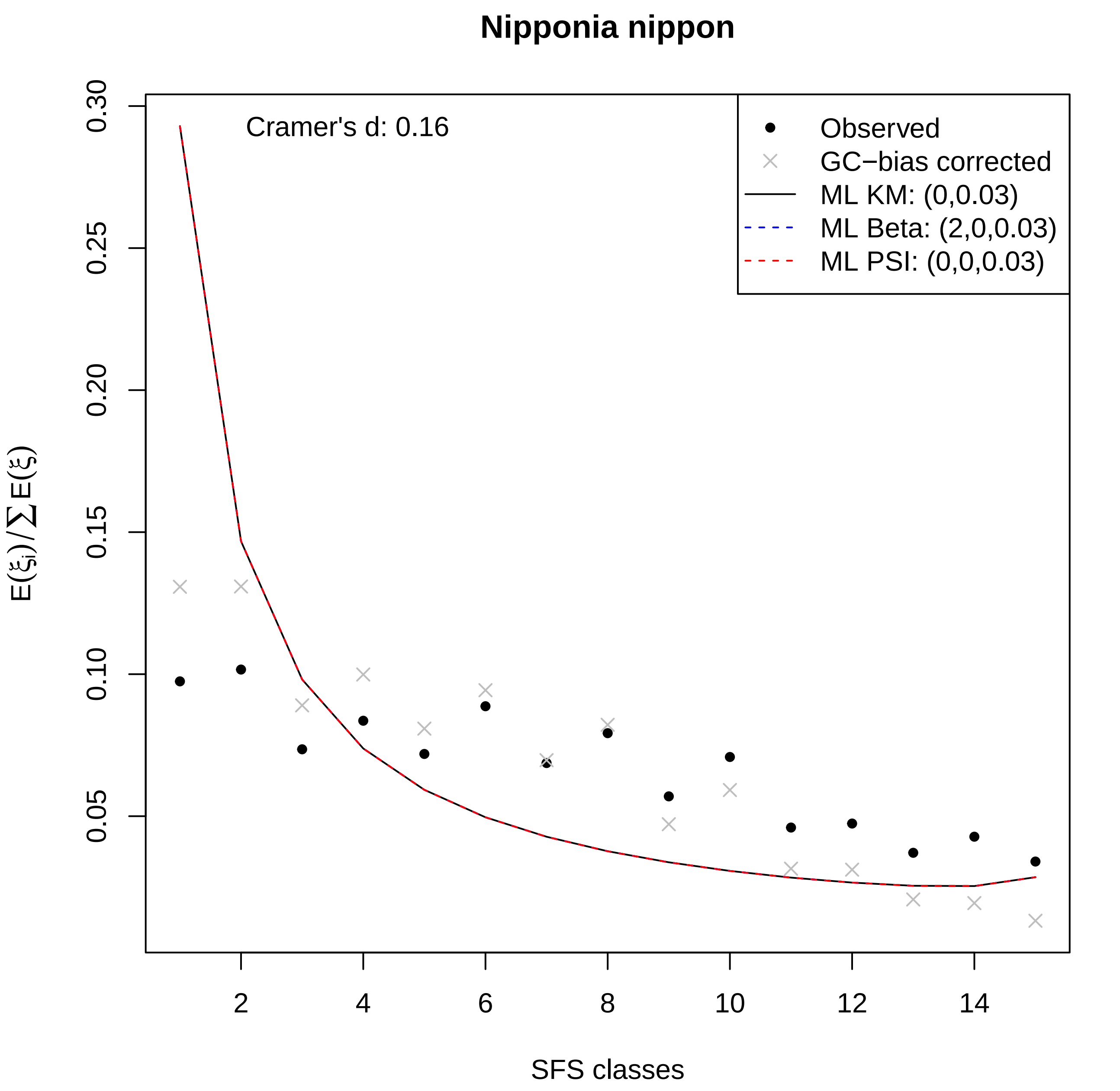

Nipponia nippon
Cramer's d: 0.16
Observed
GC−bias corrected
ML KM: (0,0.03)
ML Beta: (2,0,0.03)
ML PSI: (0,0,0.03)
0.05
0.10
0.15
0.20
0.25
0.30
2
4
6
8
10
12
14
SFS classes
$E(\xi_i)/\sum E(\xi)$

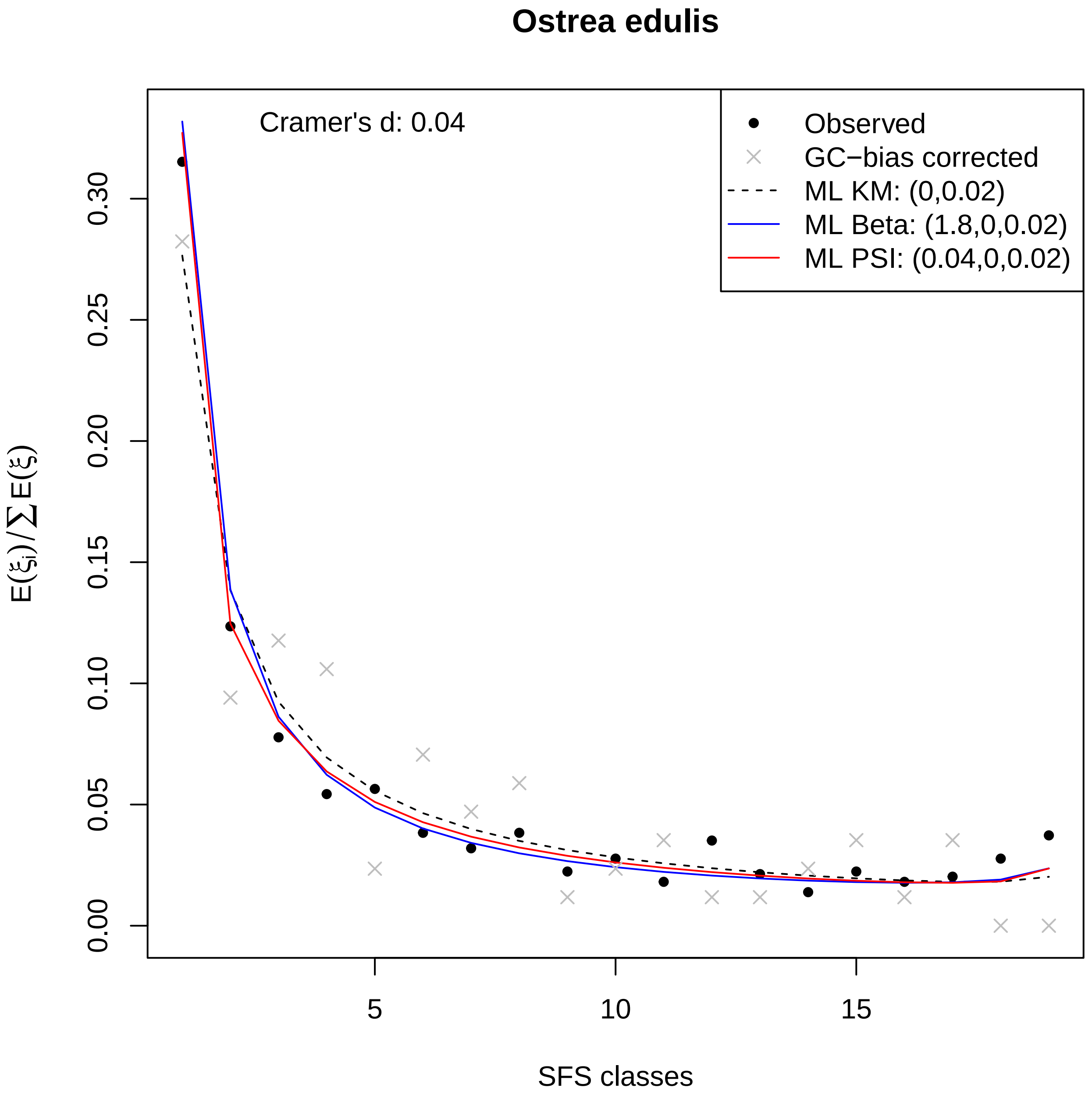

Ostrea edulis
Cramer's d: 0.04
Observed
GC−bias corrected
ML KM: (0,0.02)
ML Beta: (1.8,0,0.02)
ML PSI: (0.04,0,0.02)
$E(\xi_i)/\sum E(\xi)$
0.00
0.05
0.10
0.15
0.20
0.25
0.30
5
10
15
SFS classes

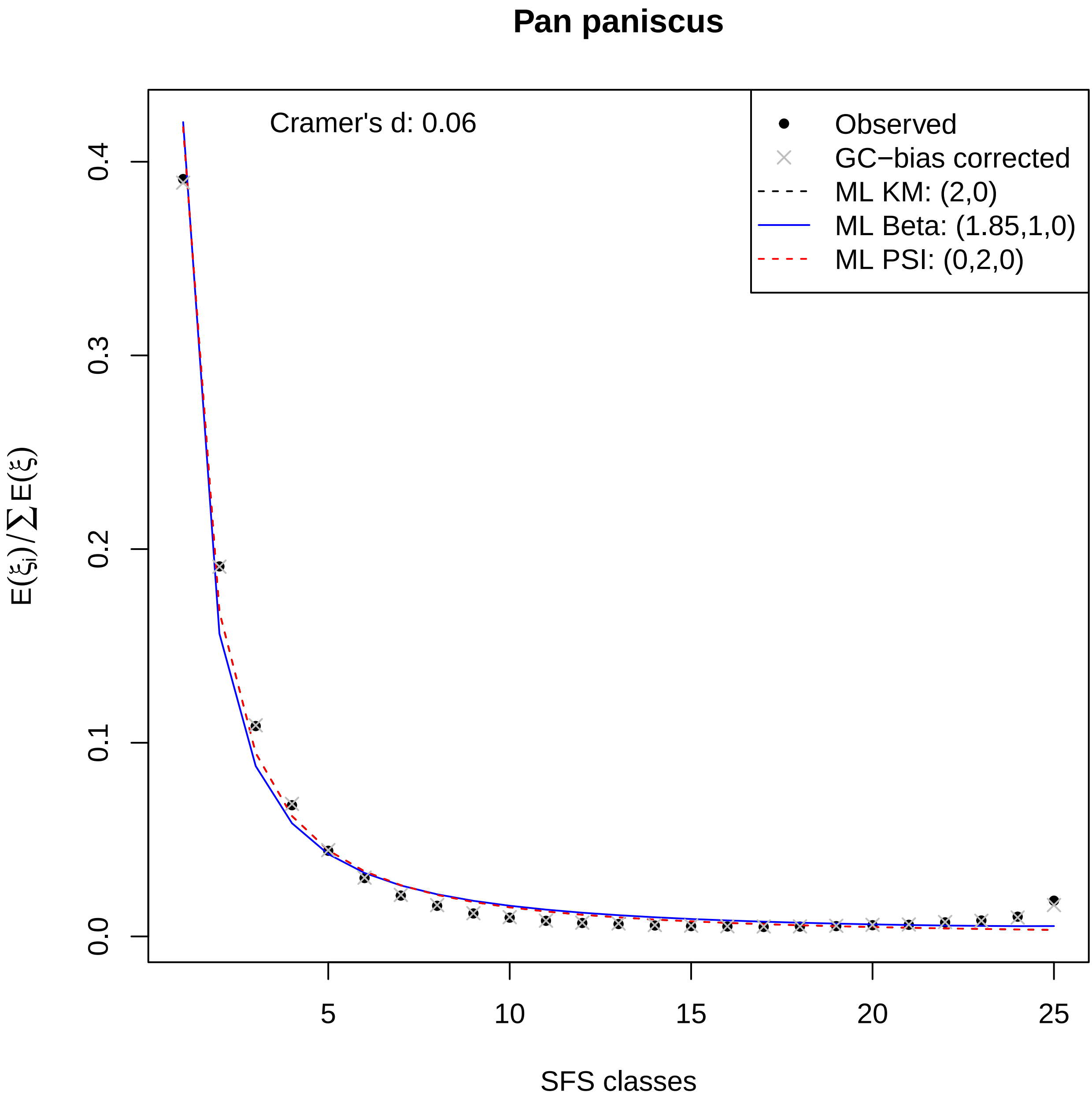

Pan paniscus
Cramer's d: 0.06
Observed
GC−bias corrected
ML KM: (2,0)
ML Beta: (1.85,1,0)
ML PSI: (0,2,0)
0.0
0.1
0.2
0.3
0.4
5
10
15
20
25
SFS classes

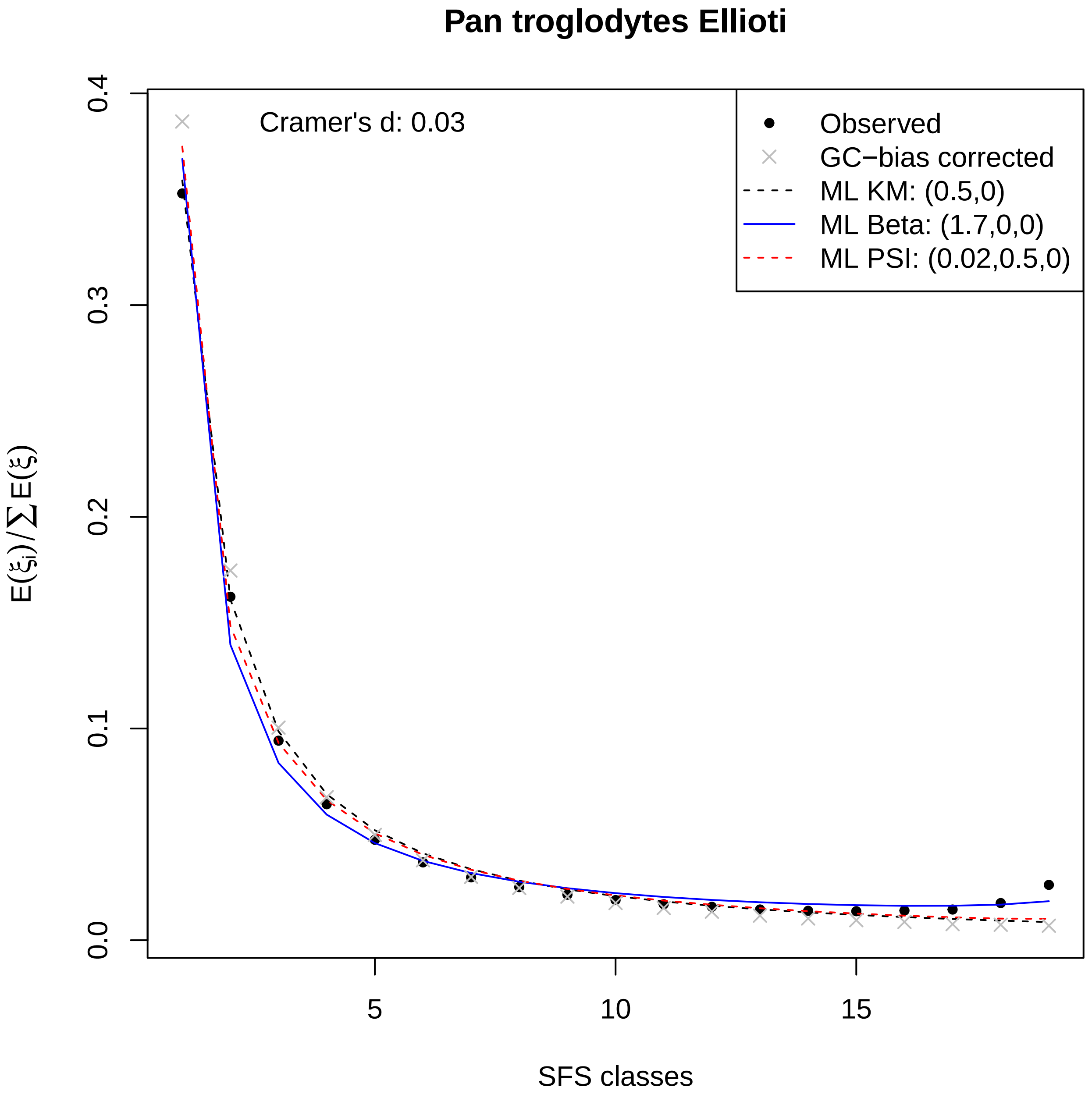

Pan troglodytes Ellioti
Cramer's d: 0.03
Observed
GC−bias corrected
ML KM: (0.5,0)
ML Beta: (1.7,0,0)
ML PSI: (0.02,0.5,0)
0.0
0.1
0.2
0.3
0.4
5
10
15
SFS classes

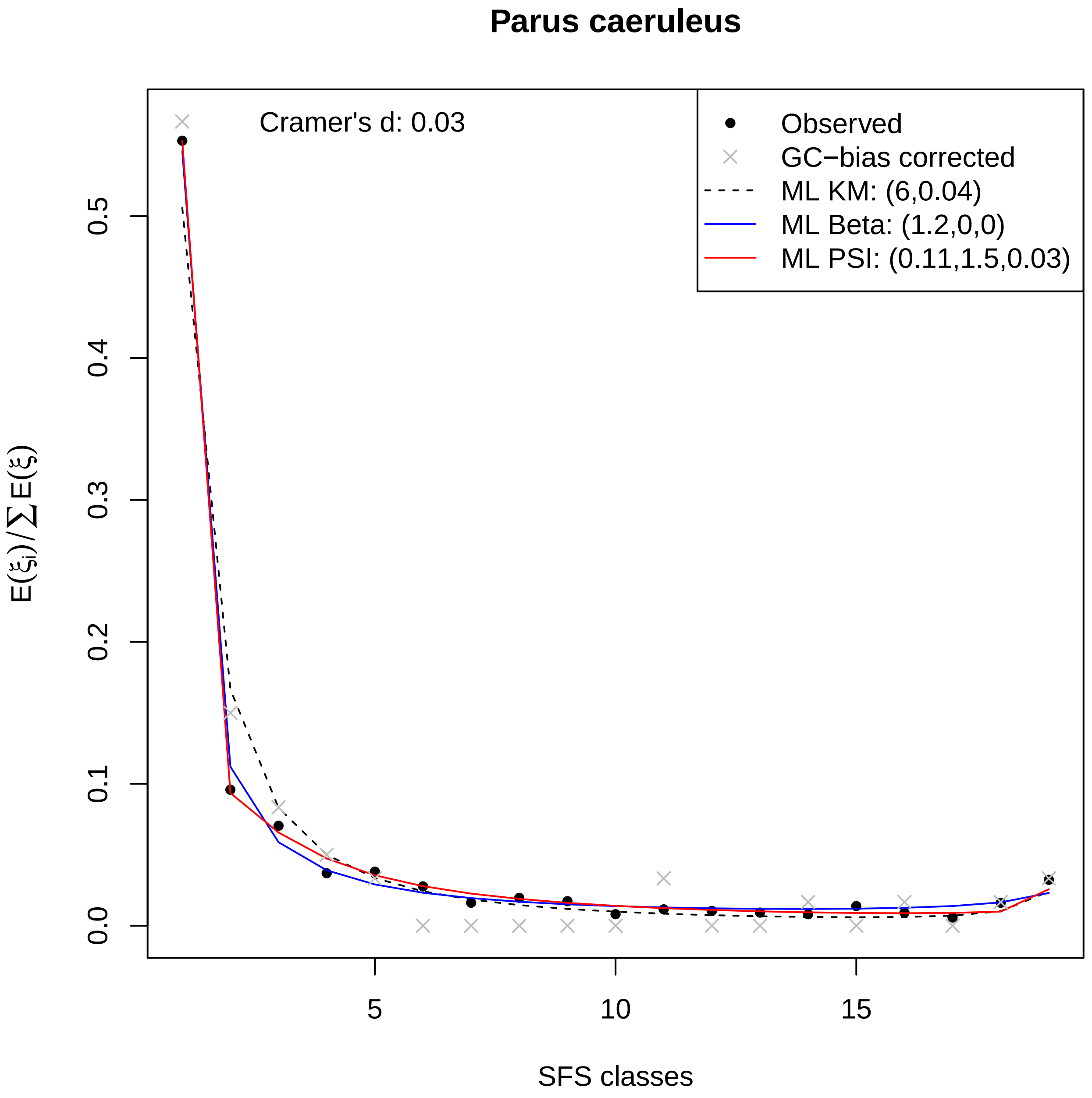

Parus caeruleus
Cramer's d: 0.03
Observed
GC−bias corrected
ML KM: (6,0.04)
ML Beta: (1.2,0,0)
ML PSI: (0.11,1.5,0.03)
0.0
0.1
0.2
0.3
0.4
0.5
5
10
15
SFS classes
$E(\xi_i)/\sum E(\xi)$

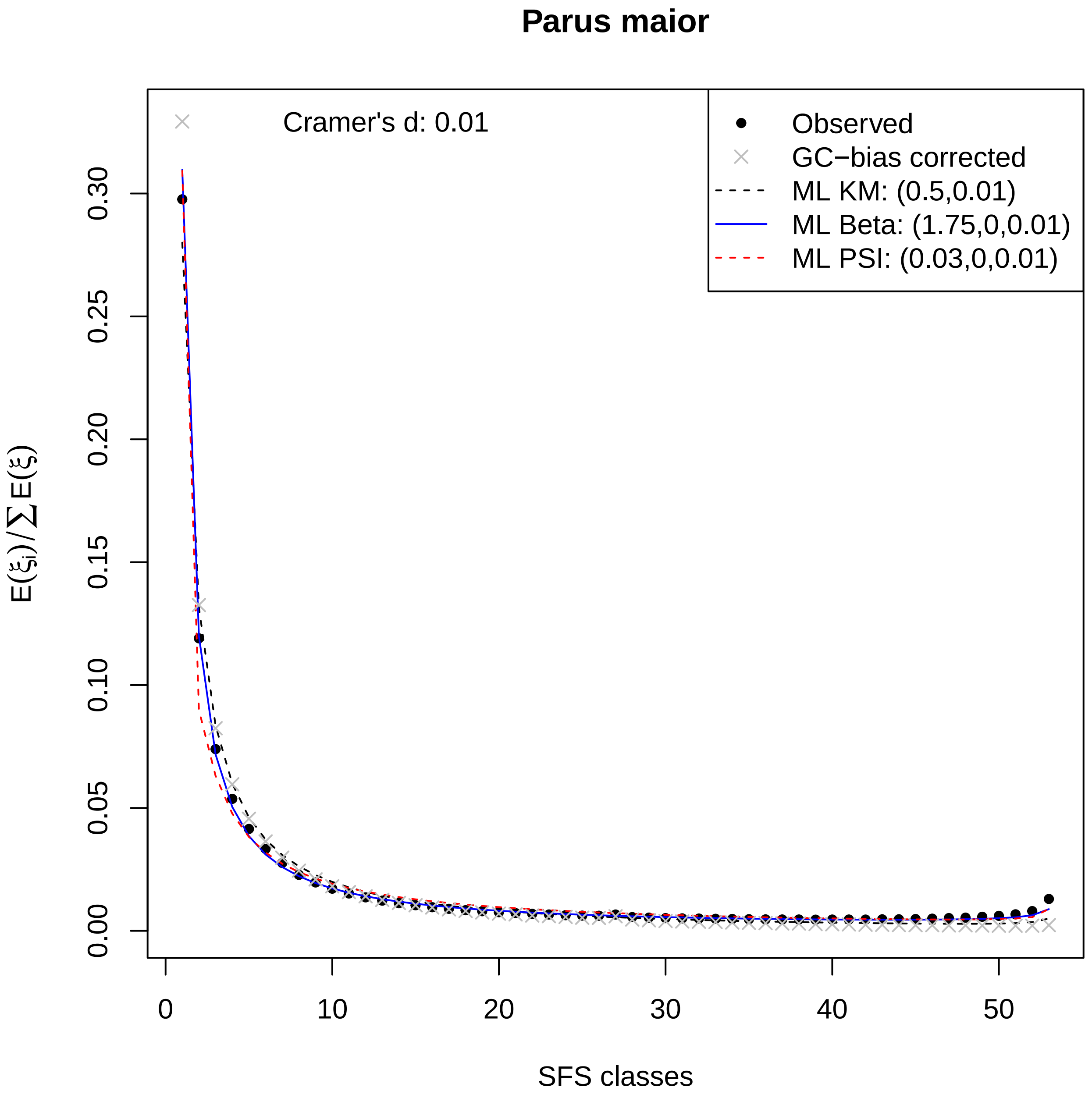

Parus maior
Cramer's d: 0.01
Observed
GC−bias corrected
ML KM: (0.5,0.01)
ML Beta: (1.75,0,0.01)
ML PSI: (0.03,0,0.01)
0.00
0.05
0.10
0.15
0.20
0.25
0.30
$E(\xi_i)/\sum E(\xi)$
0
10
20
30
40
50
SFS classes

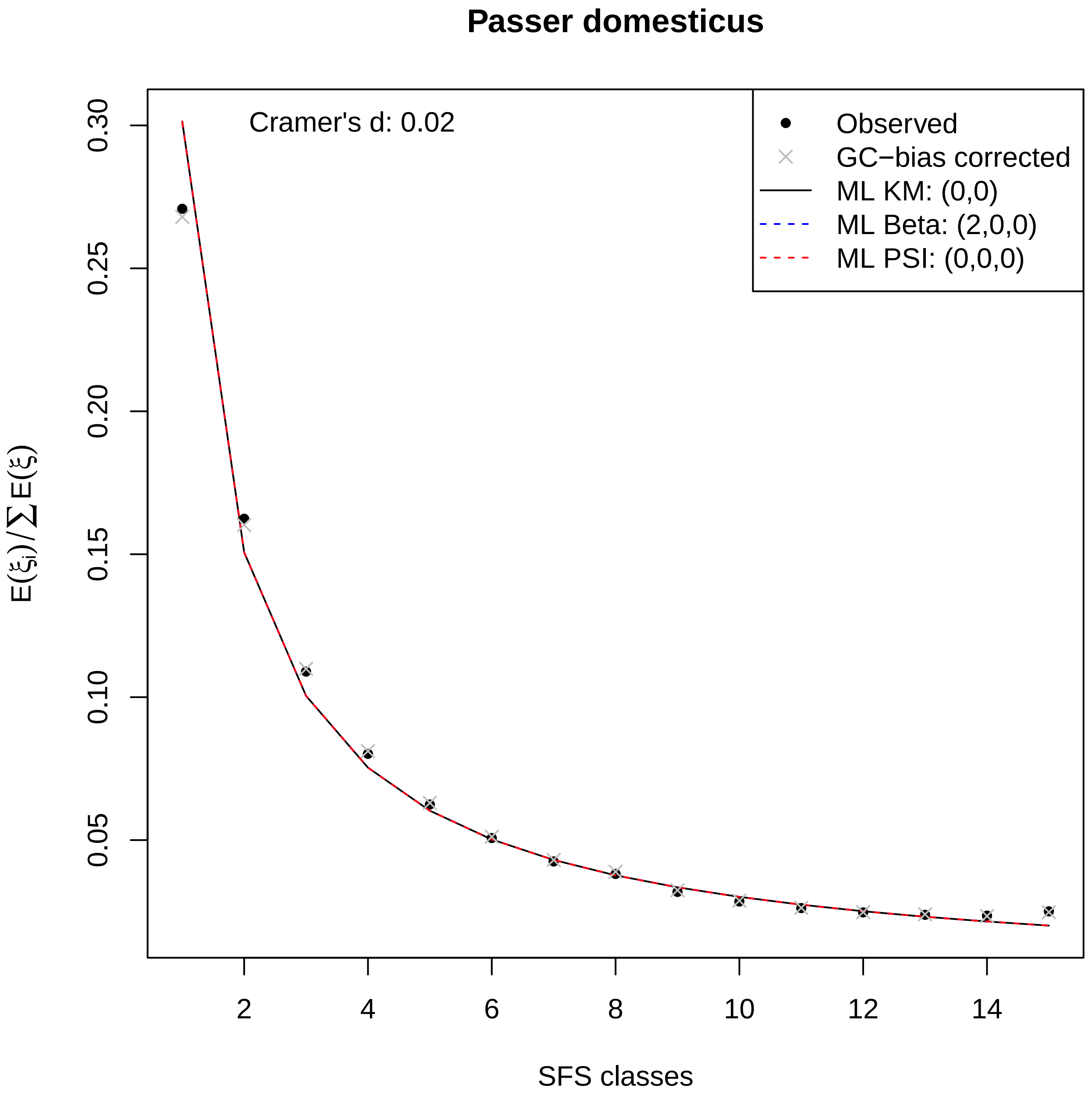
Passer domesticus
Cramer's d: 0.02
Observed
GC−bias corrected
ML KM: (0,0)
ML Beta: (2,0,0)
ML PSI: (0,0,0)
$E(\xi_i)/\sum E(\xi)$
SFS classes
0.05
0.10
0.15
0.20
0.25
0.30
2
4
6
8
10
12
14

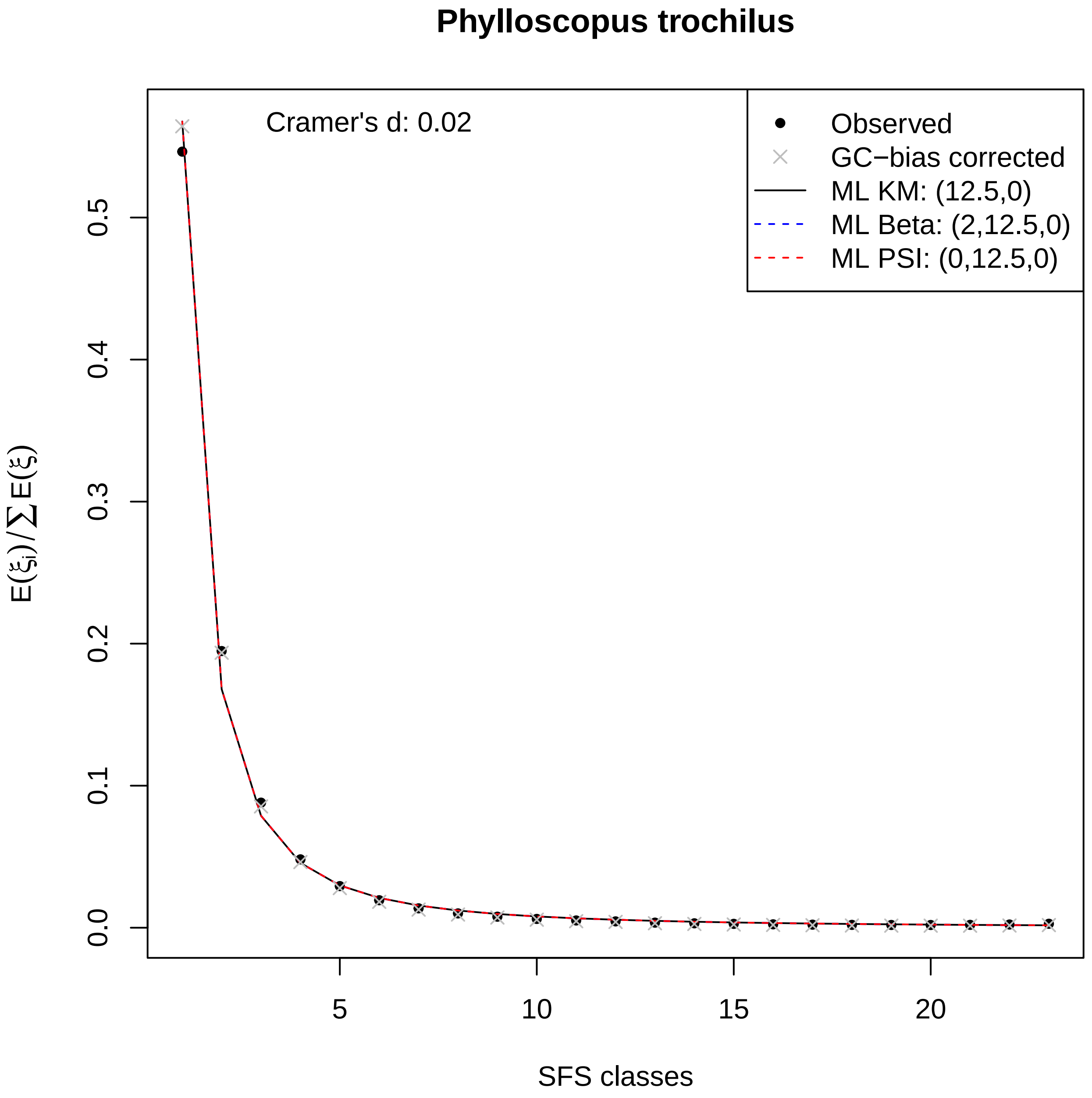
Phylloscopus trochilus
Cramer's d: 0.02
Observed
GC−bias corrected
ML KM: (12.5,0)
ML Beta: (2,12.5,0)
ML PSI: (0,12.5,0)
$E(\xi_i)/\sum E(\xi)$
SFS classes
0.0
0.1
0.2
0.3
0.4
0.5
5
10
15
20

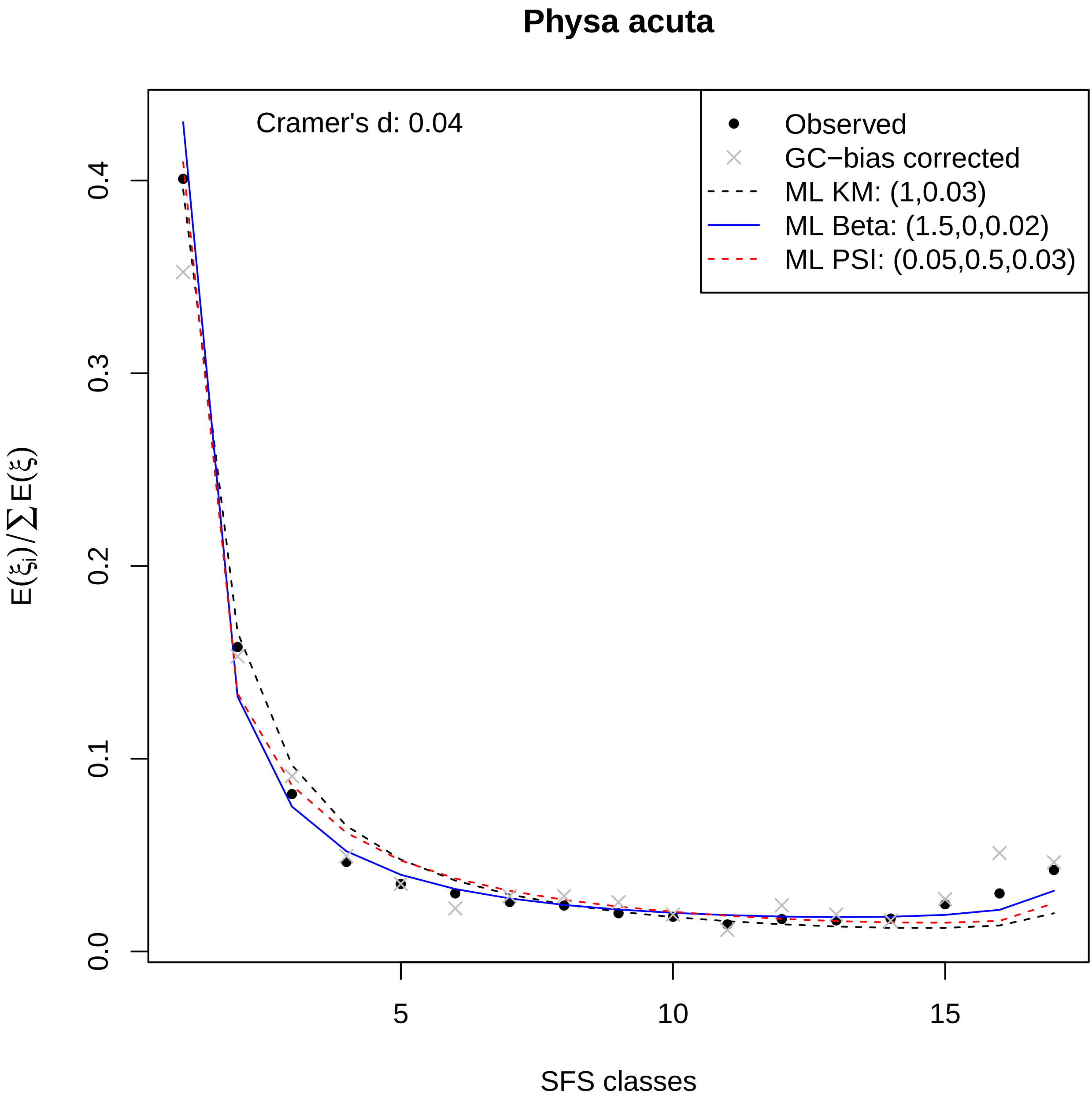
Physa acuta
Cramer's d: 0.04
Observed
GC−bias corrected
ML KM: (1,0.03)
ML Beta: (1.5,0,0.02)
ML PSI: (0.05,0.5,0.03)
0.0
0.1
0.2
0.3
0.4
5
10
15
SFS classes
$E(\xi_i)/\sum E(\xi)$

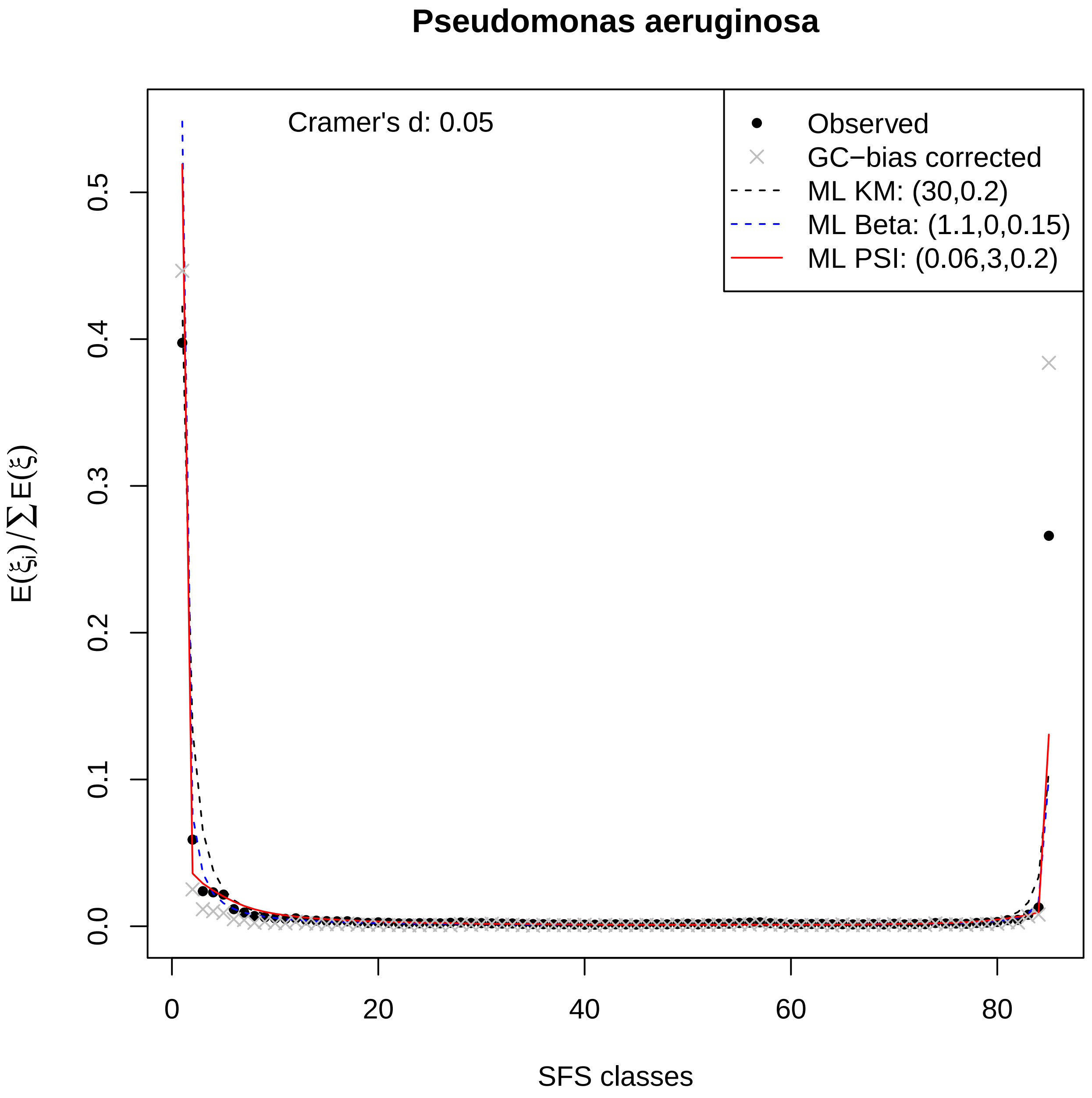

Pseudomonas aeruginosa
Cramer's d: 0.05
Observed
GC−bias corrected
ML KM: (30,0.2)
ML Beta: (1.1,0,0.15)
ML PSI: (0.06,3,0.2)
$E(\xi_i)/\sum E(\xi)$
0.0
0.1
0.2
0.3
0.4
0.5
0
20
40
60
80
SFS classes

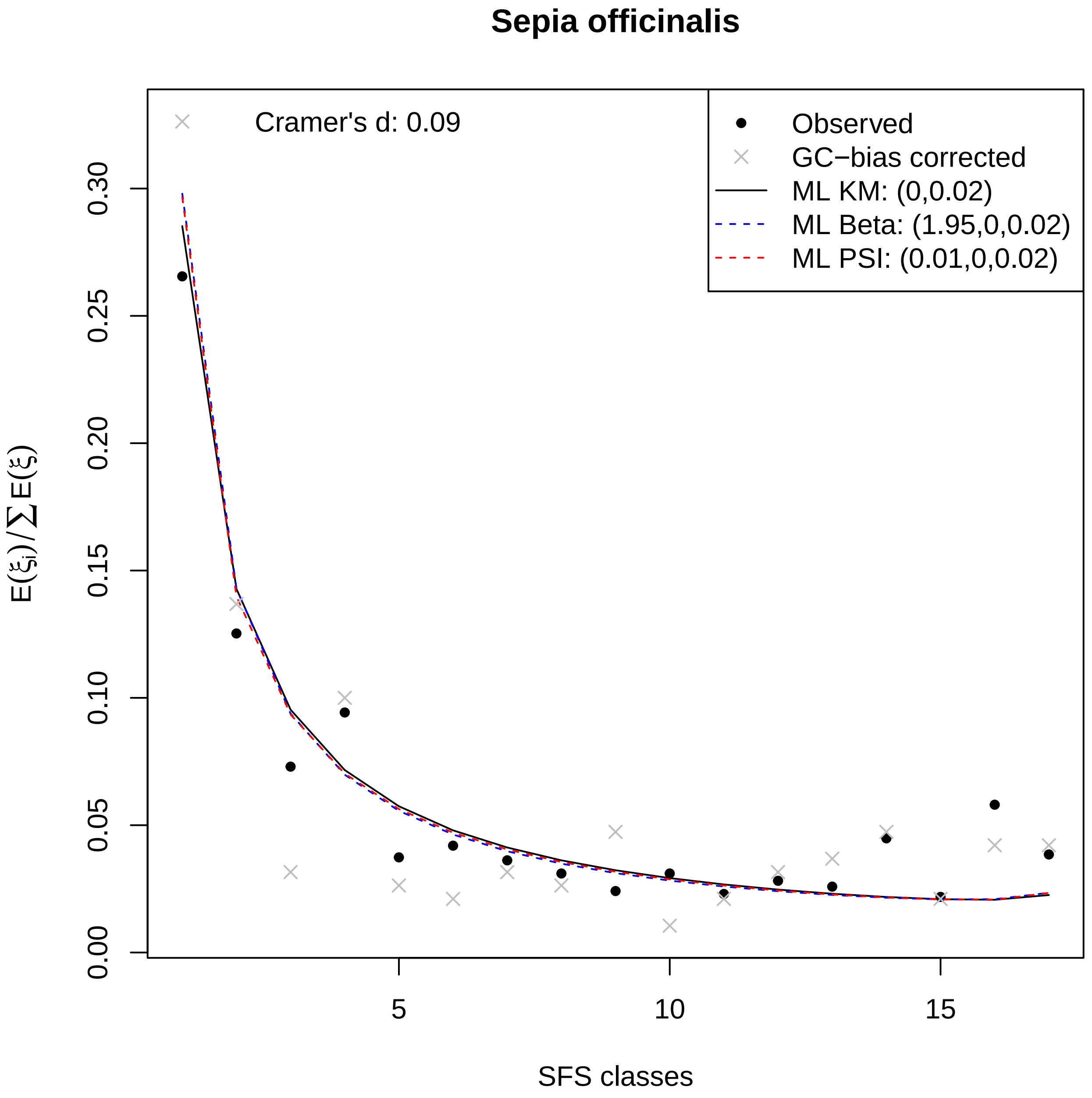

Sepia officinalis
Cramer's d: 0.09
Observed
GC−bias corrected
ML KM: (0,0.02)
ML Beta: (1.95,0,0.02)
ML PSI: (0.01,0,0.02)
0.00
0.05
0.10
0.15
0.20
0.25
0.30
5
10
15
SFS classes
$E(\xi_i)/\sum E(\xi)$

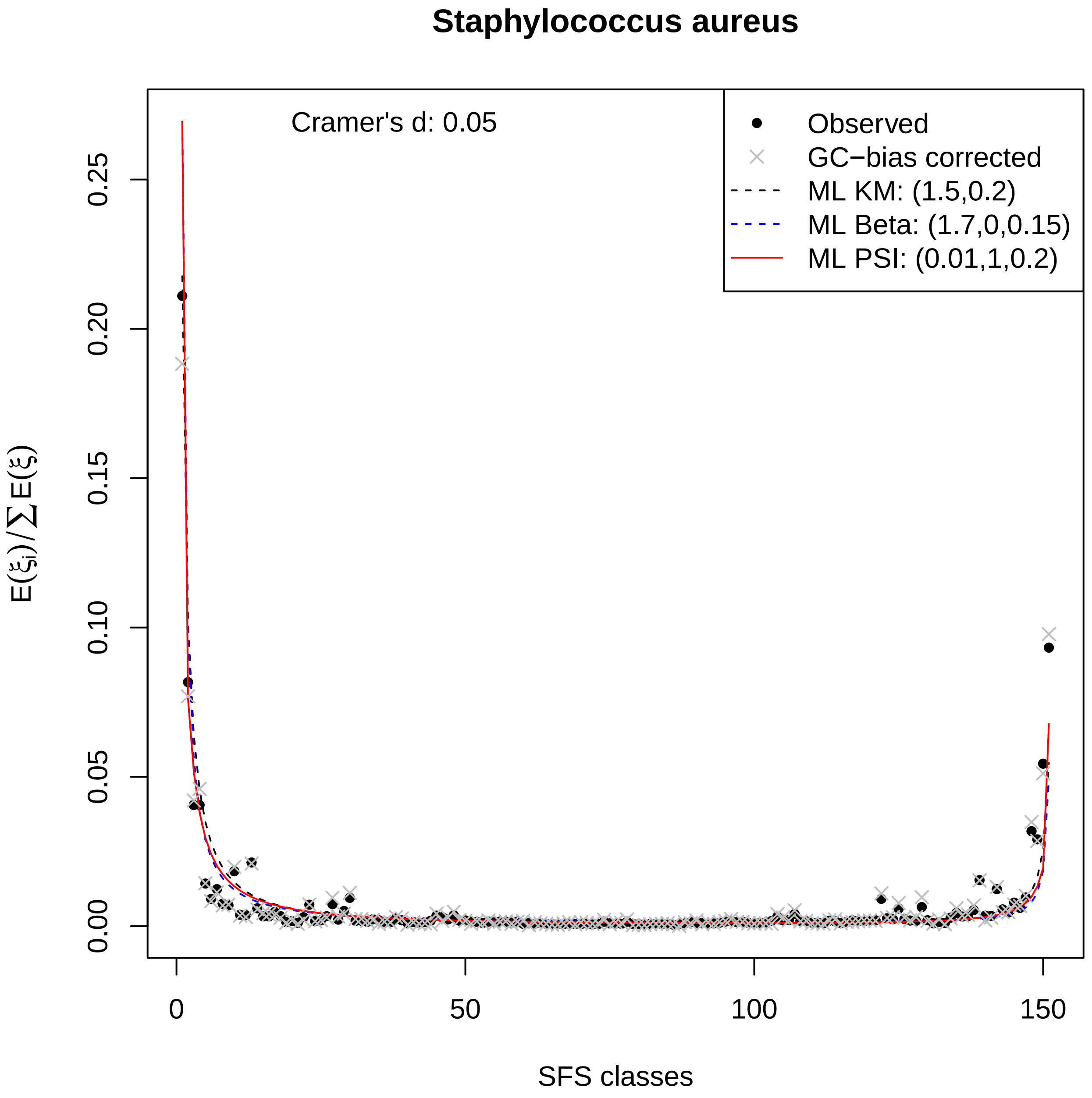

Staphylococcus aureus
Cramer's d: 0.05
Observed
GC−bias corrected
ML KM: (1.5,0.2)
ML Beta: (1.7,0,0.15)
ML PSI: (0.01,1,0.2)
0.00
0.05
0.10
0.15
0.20
0.25
0
50
100
150
SFS classes
$E(\xi_i)/\sum E(\xi)$

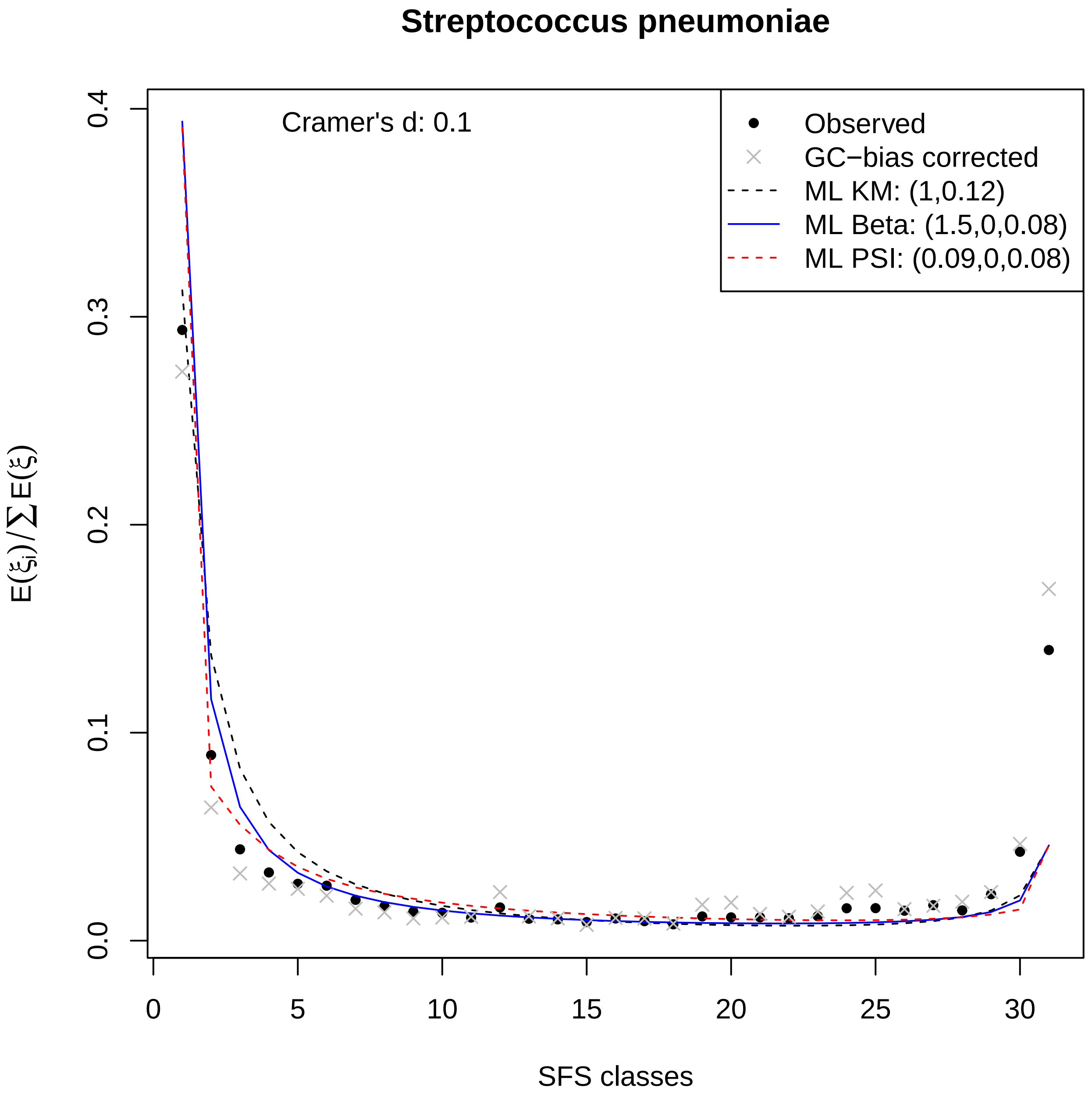
Streptococcus pneumoniae
Cramer's d: 0.1
Observed
GC−bias corrected
ML KM: (1,0.12)
ML Beta: (1.5,0,0.08)
ML PSI: (0.09,0,0.08)
$E(\xi_i)/\sum E(\xi)$
SFS classes
0.0
0.1
0.2
0.3
0.4
0
5
10
15
20
25
30

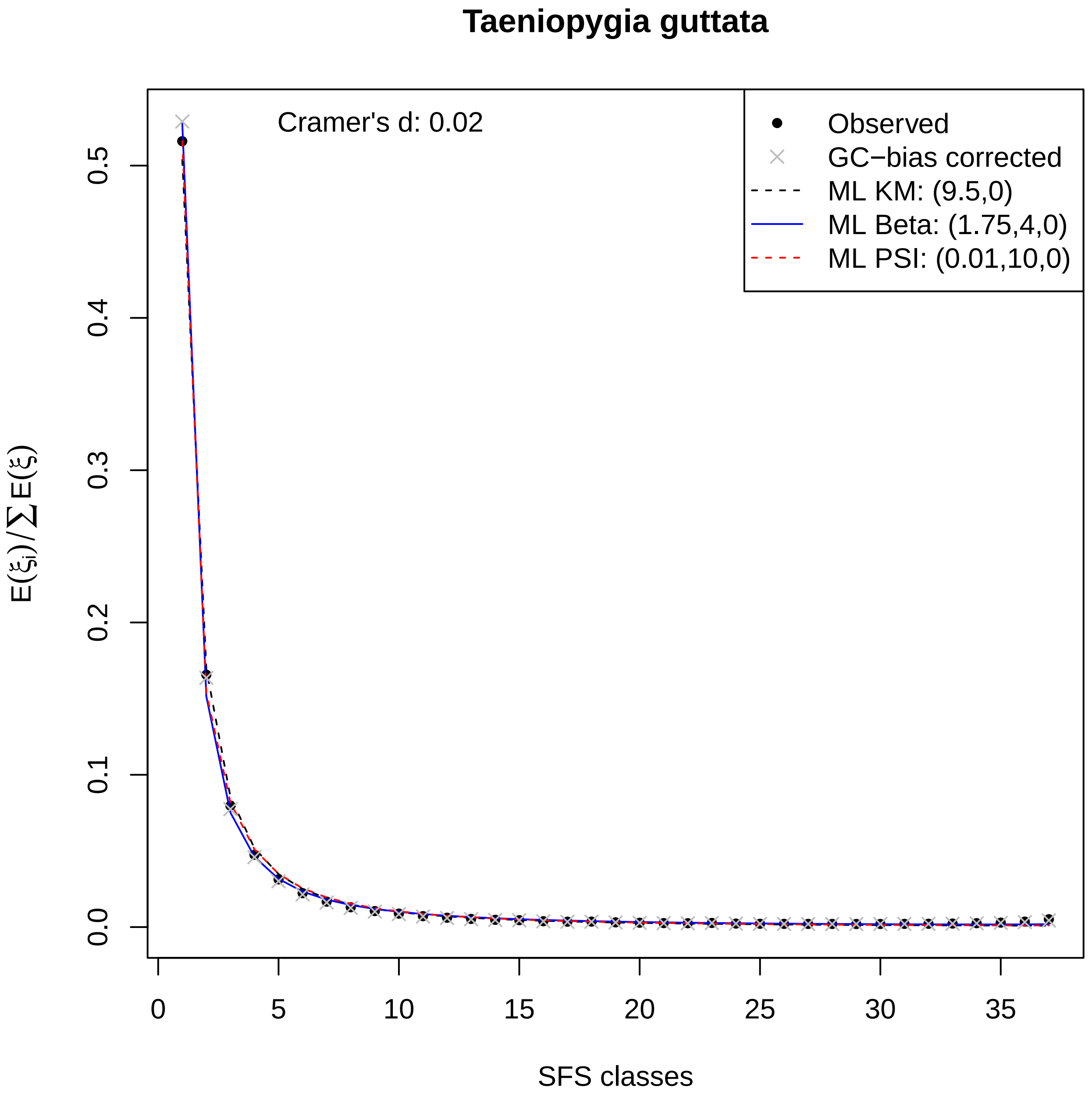

Taeniopygia guttata
Cramer's d: 0.02
Observed
GC−bias corrected
ML KM: (9.5,0)
ML Beta: (1.75,4,0)
ML PSI: (0.01,10,0)
$E(\xi_i)/\sum E(\xi)$
SFS classes
0.0
0.1
0.2
0.3
0.4
0.5
0
5
10
15
20
25
30
35

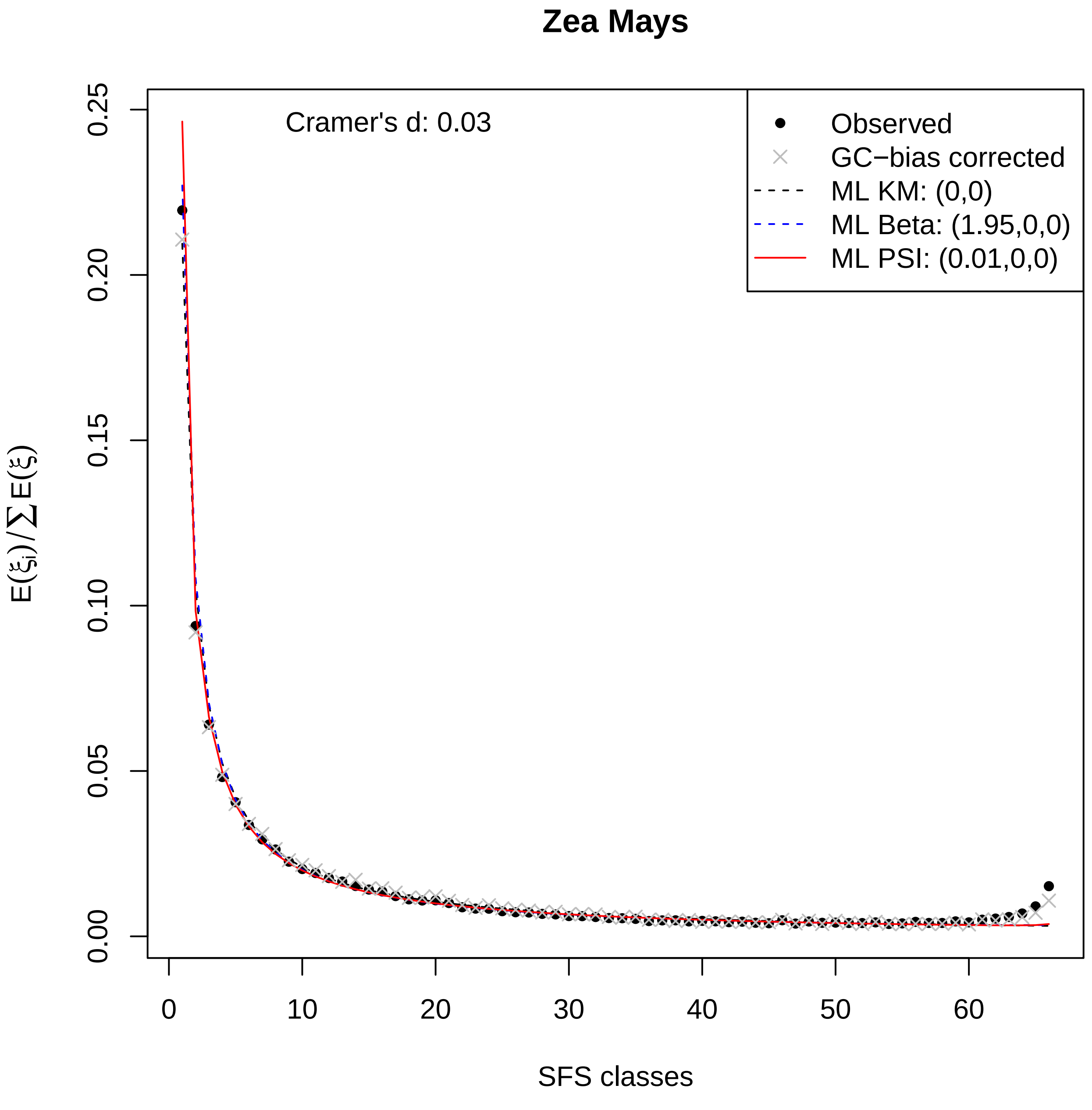

Zea Mays
Cramer's d: 0.03
Observed
GC−bias corrected
ML KM: (0,0)
ML Beta: (1.95,0,0)
ML PSI: (0.01,0,0)
0.25
0.20
0.15
0.10
0.05
0.00
0
10
20
30
40
50
60
$E(\xi_i)/\sum E(\xi)$
SFS classes
