## Supplementary 2 - tranformed SFS and fits for "Interpreting the pervasive observation of U-shaped Site Frequency Spectra"

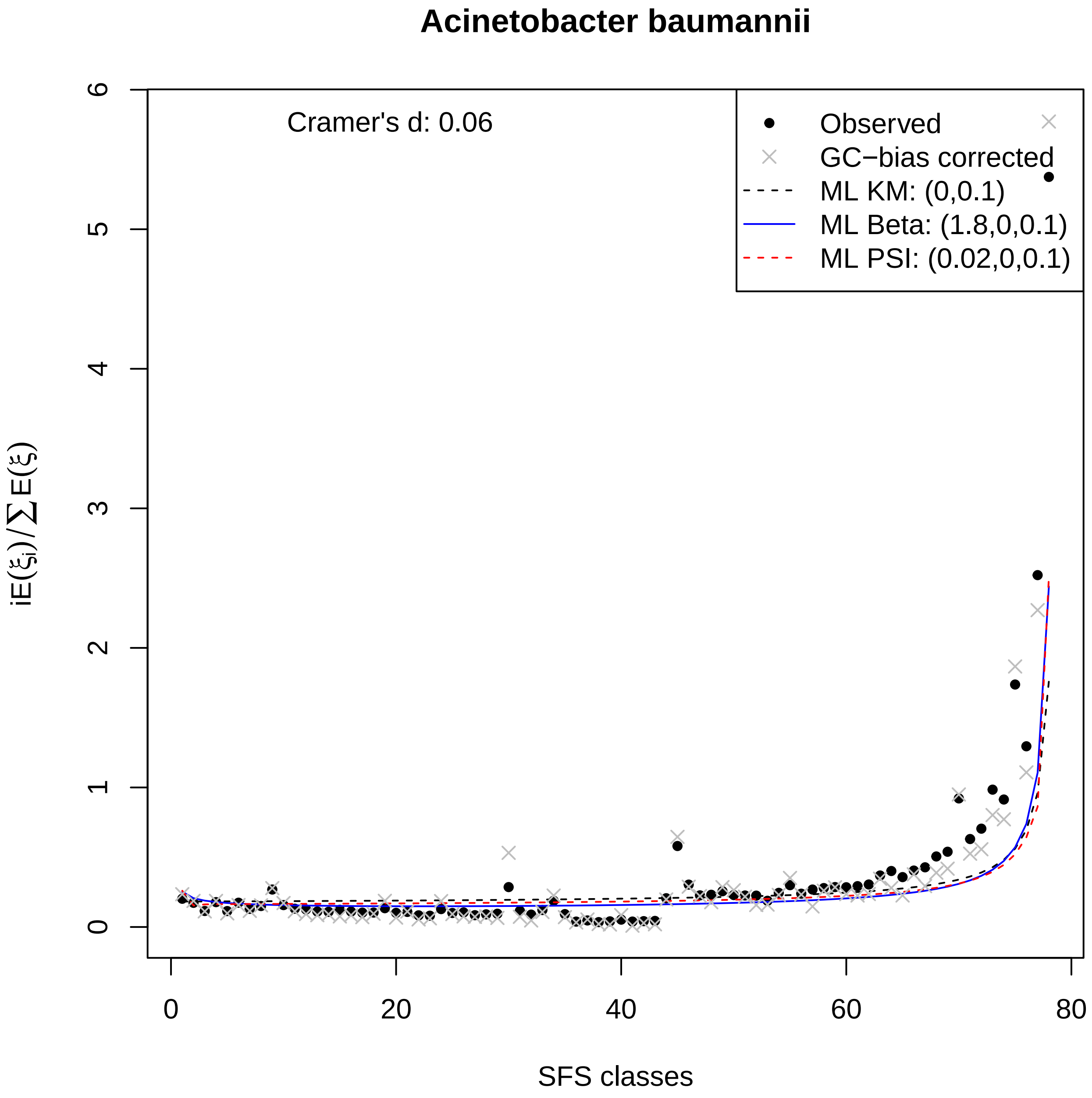

Acinetobacter baumannii
Cramer's d: 0.06
Observed
GC−bias corrected
ML KM: (0,0.1)
ML Beta: (1.8,0,0.1)
ML PSI: (0.02,0,0.1)
$iE(\xi_i)/\sum E(\xi)$
SFS classes
0
1
2
3
4
5
6
0
20
40
60
80

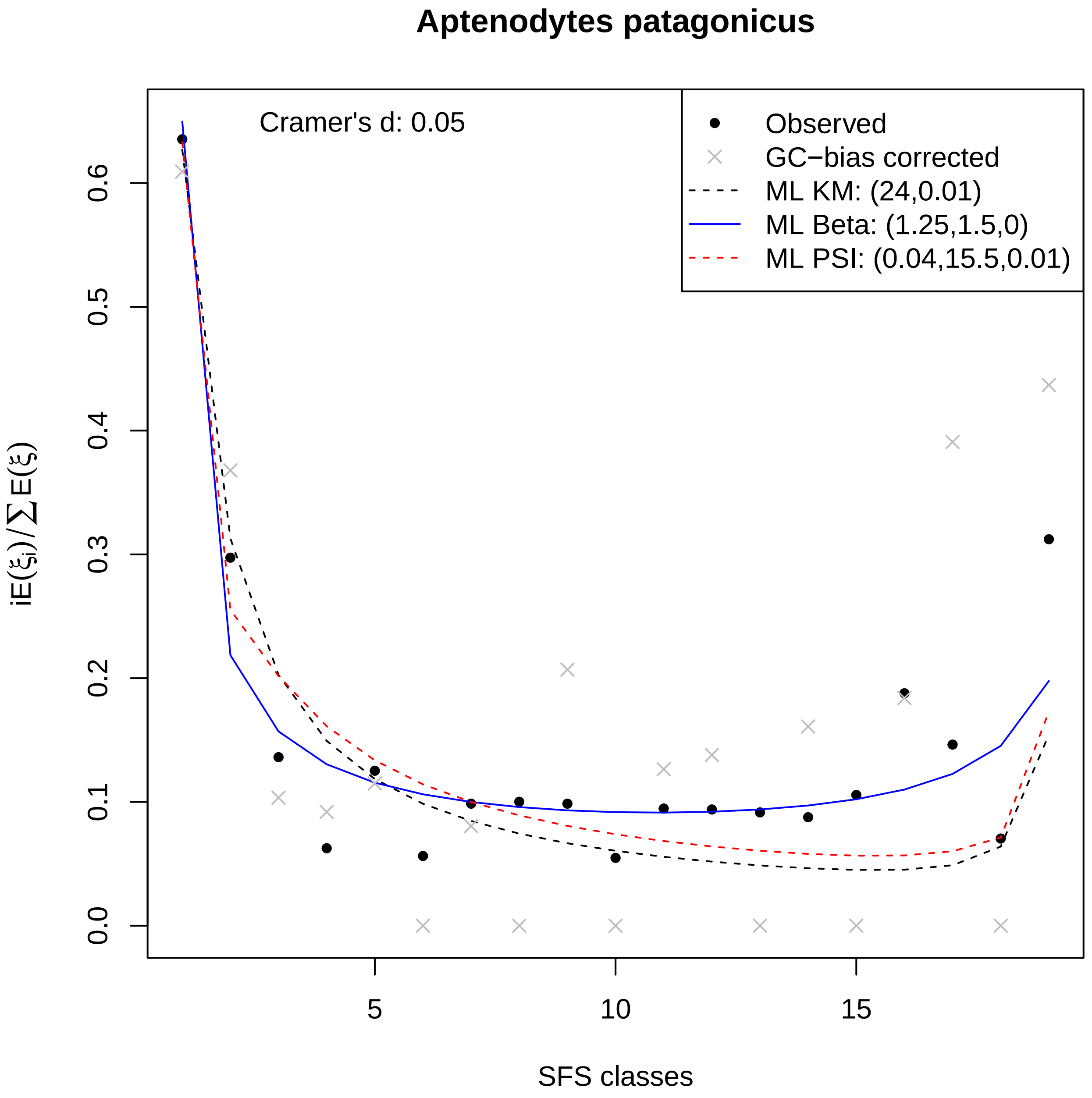
Aptenodytes patagonicus
Cramer's d: 0.05
Observed
GC−bias corrected
ML KM: (24,0.01)
ML Beta: (1.25,1.5,0)
ML PSI: (0.04,15.5,0.01)
iE(ξi)/∑E(ξ)
SFS classes
0.0
0.1
0.2
0.3
0.4
0.5
0.6
5
10
15

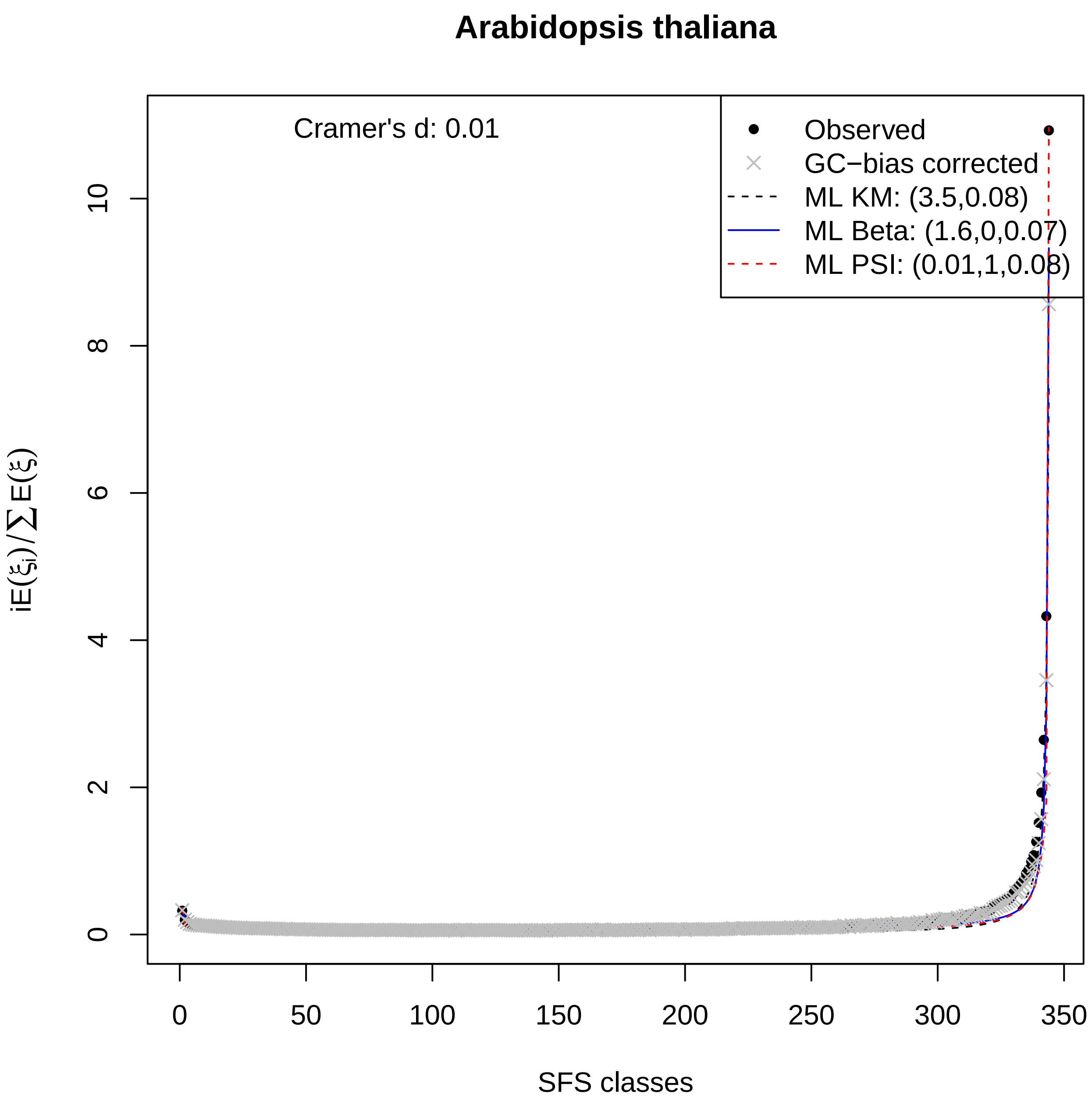
Arabidopsis thaliana
Cramer's d: 0.01
Observed
GC−bias corrected
ML KM: (3.5,0.08)
ML Beta: (1.6,0,0.07)
ML PSI: (0.01,1,0.08)
iE(ξi)/∑E(ξ)
SFS classes
0
2
4
6
8
10
0
50
100
150
200
250
300
350

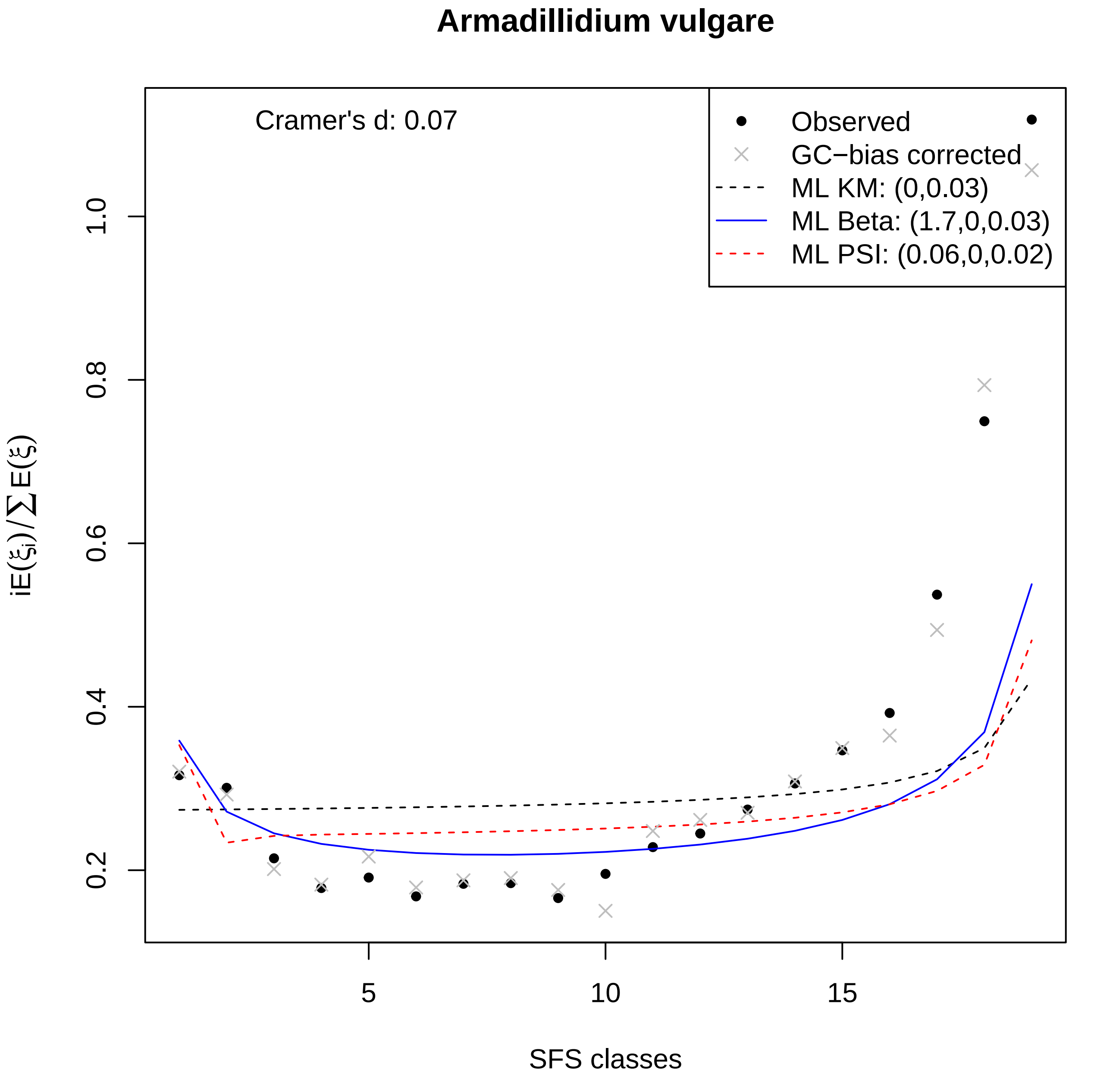
Armadillidium vulgare
Cramer's d: 0.07
Observed
GC−bias corrected
ML KM: (0,0.03)
ML Beta: (1.7,0,0.03)
ML PSI: (0.06,0,0.02)
$iE(\xi_i)/\sum E(\xi)$
0.2
0.4
0.6
0.8
1.0
5
10
15
SFS classes

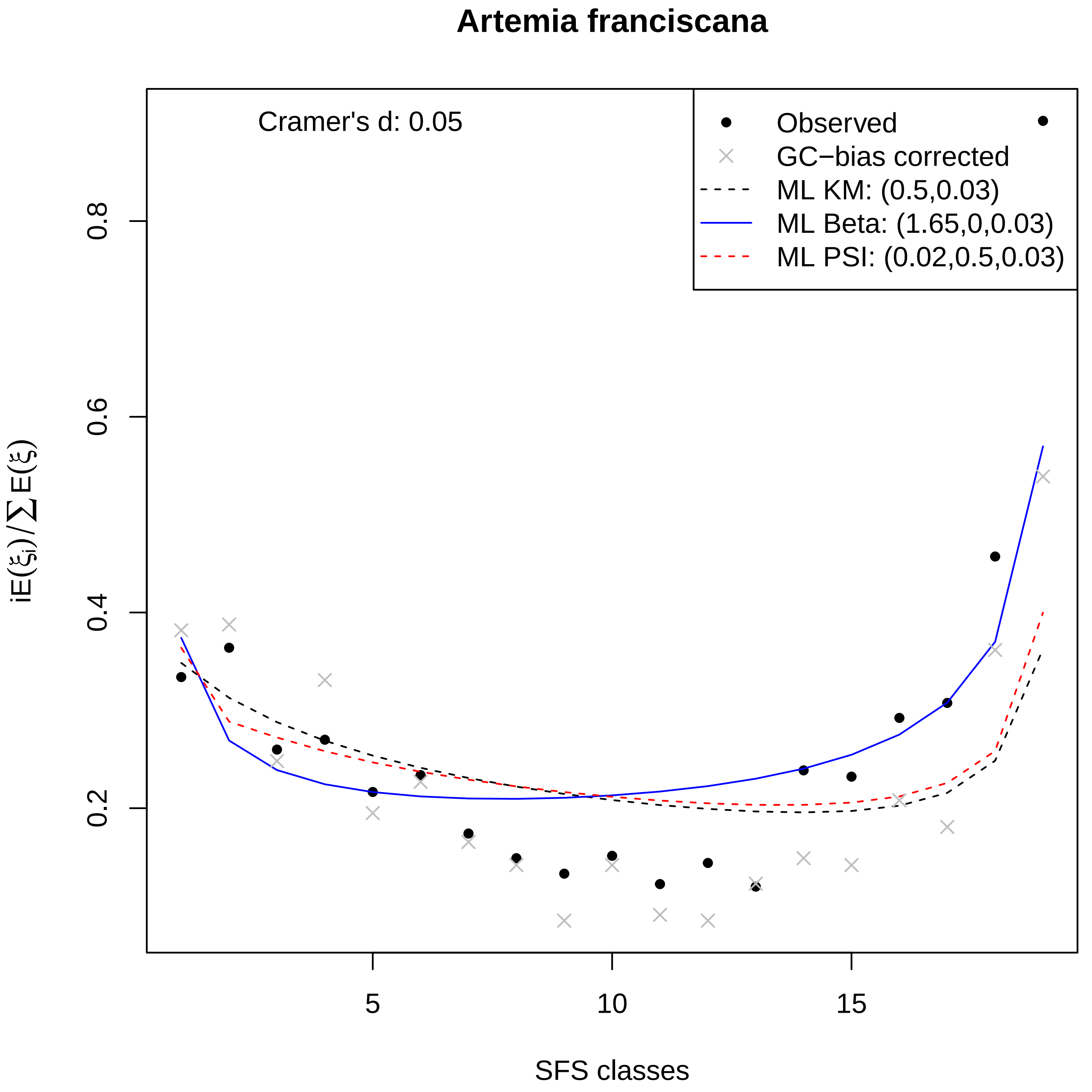

Artemia franciscana
Cramer's d: 0.05
Observed
GC−bias corrected
ML KM: (0.5,0.03)
ML Beta: (1.65,0,0.03)
ML PSI: (0.02,0.5,0.03)
0.2
0.4
0.6
0.8
5
10
15
SFS classes
$iE(\xi_i)/\sum E(\xi)$

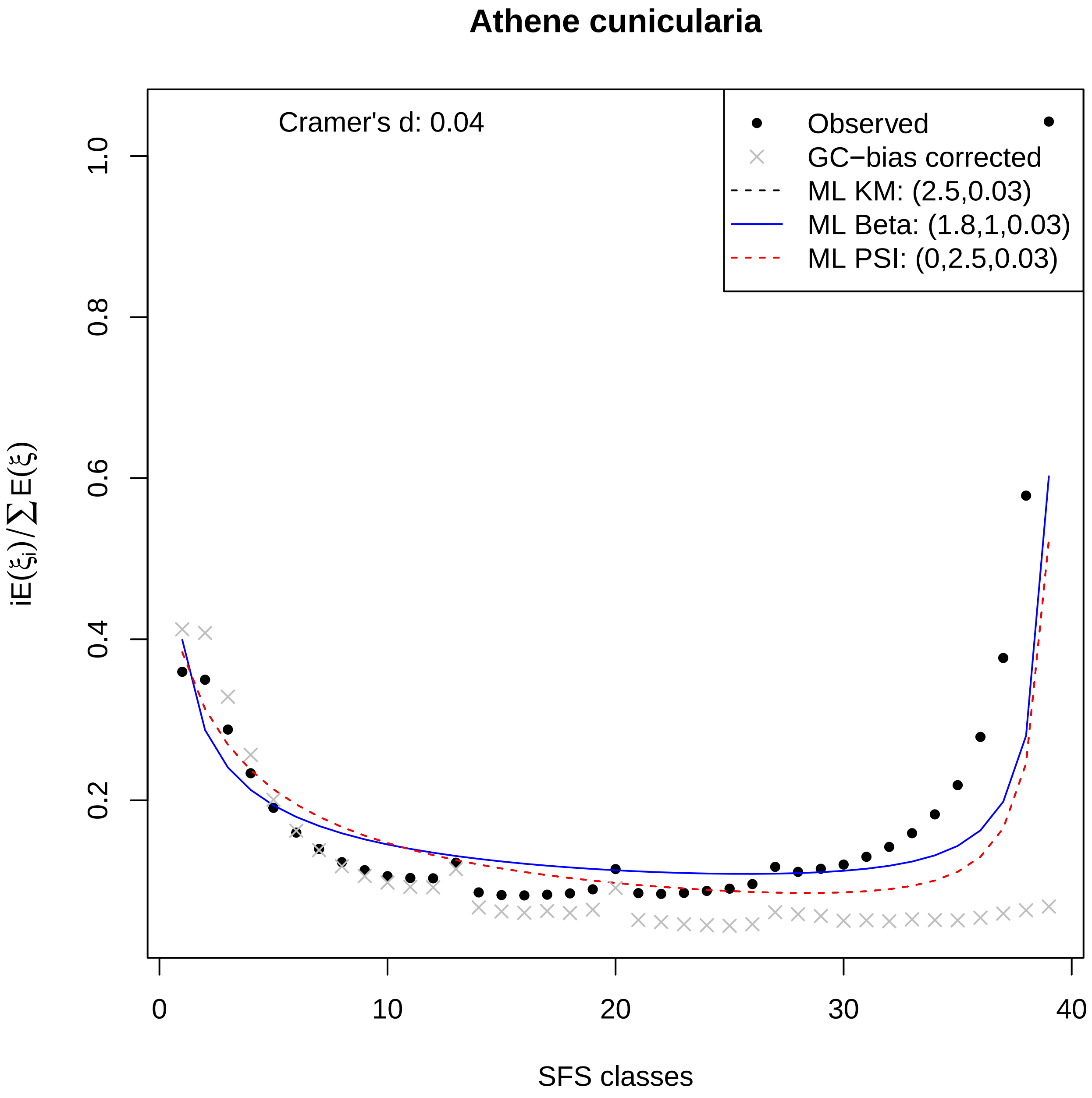

Athene cunicularia
Cramer's d: 0.04
Observed
GC−bias corrected
ML KM: (2.5,0.03)
ML Beta: (1.8,1,0.03)
ML PSI: (0,2.5,0.03)
iE(ξi)/∑E(ξ)
0.2
0.4
0.6
0.8
1.0
0
10
20
30
40
SFS classes

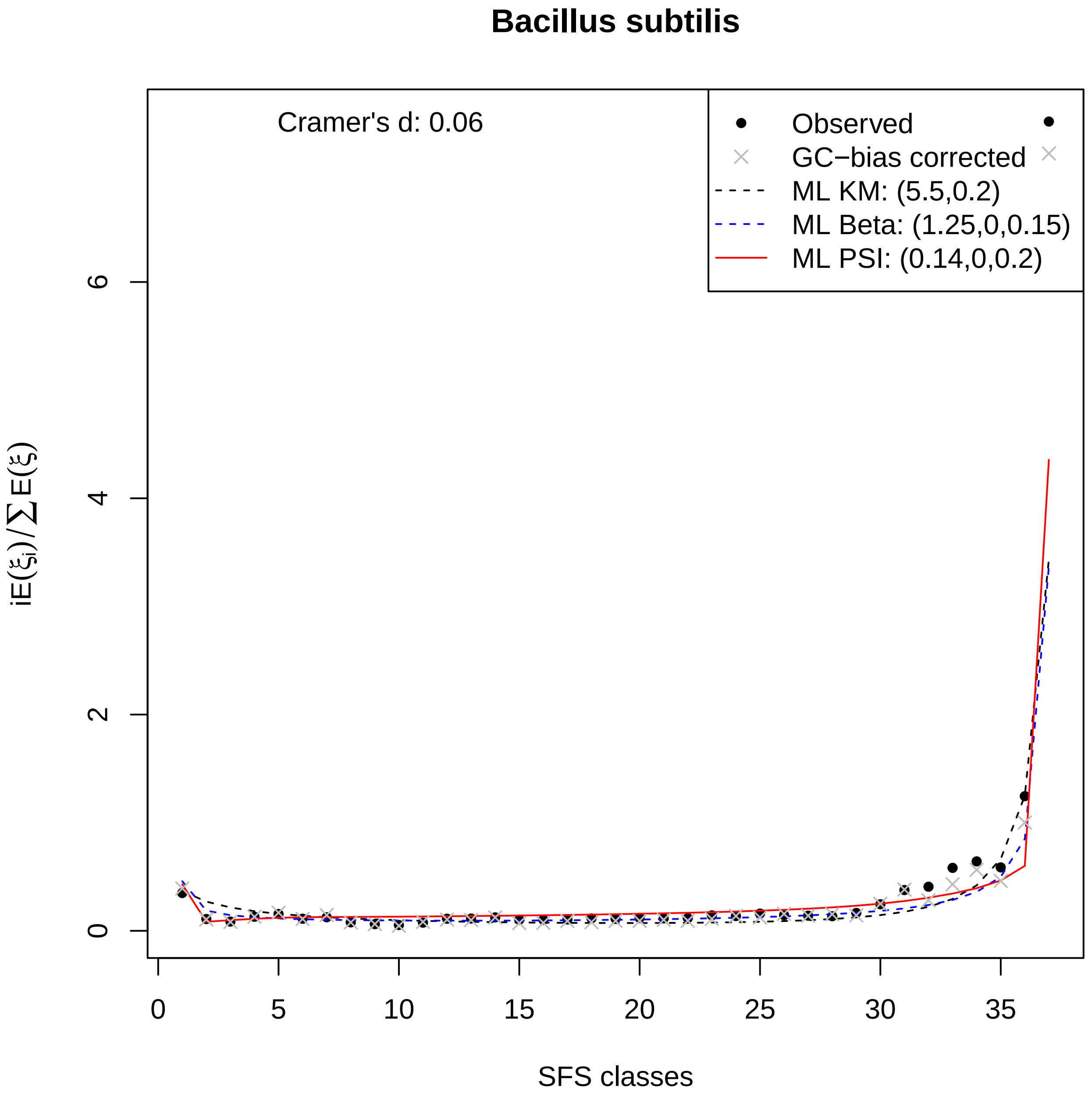

Bacillus subtilis
Cramer's d: 0.06
Observed
GC−bias corrected
ML KM: (5.5,0.2)
ML Beta: (1.25,0,0.15)
ML PSI: (0.14,0,0.2)
0
2
4
6
iE(ξi)/∑E(ξ)
0
5
10
15
20
25
30
35
SFS classes

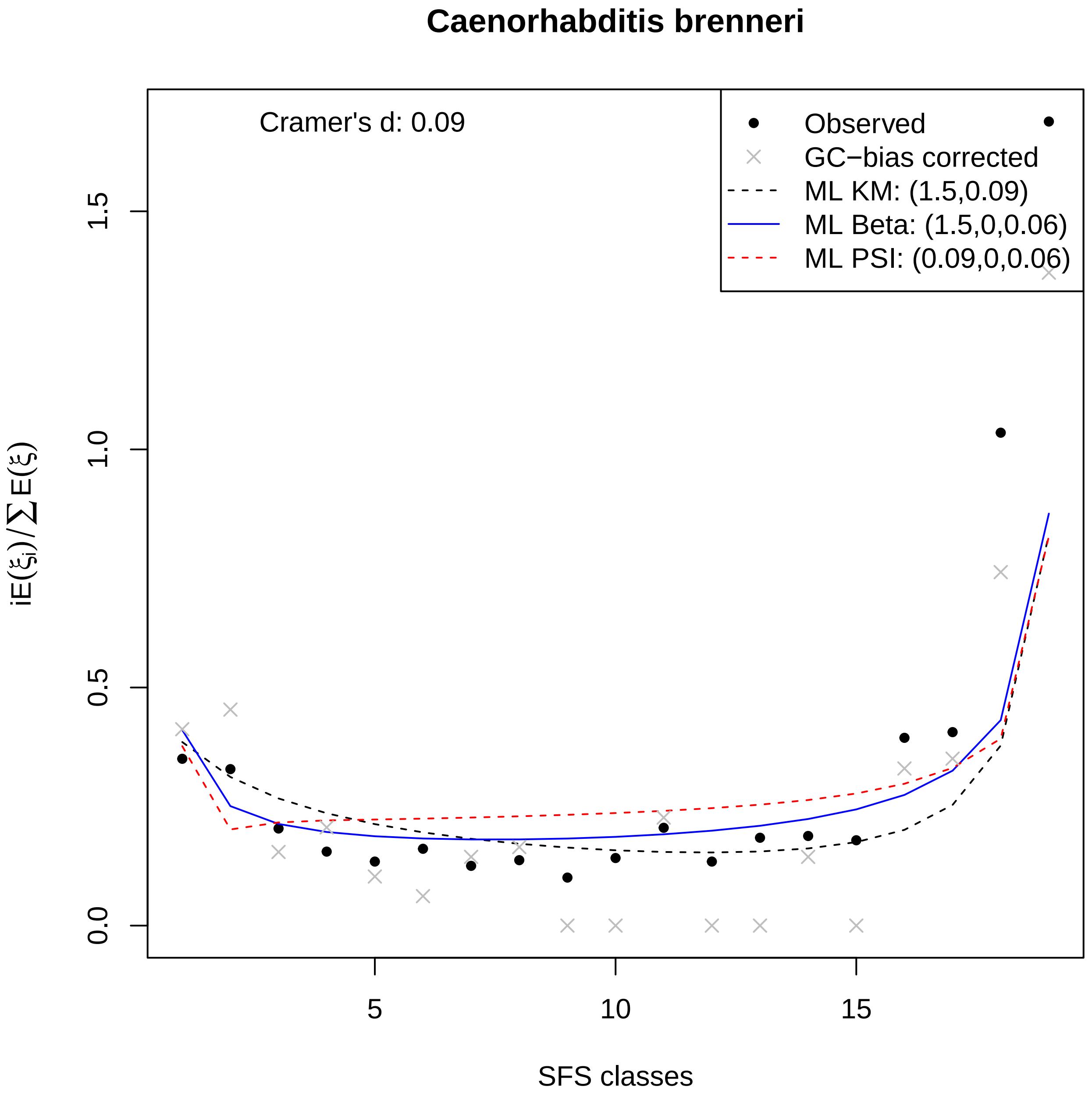
Caenorhabditis brenneri
Cramer's d: 0.09
Observed
GC−bias corrected
ML KM: (1.5,0.09)
ML Beta: (1.5,0,0.06)
ML PSI: (0.09,0,0.06)
0.0
0.5
1.0
1.5
5
10
15
SFS classes
$iE(\xi_i)/\sum E(\xi)$

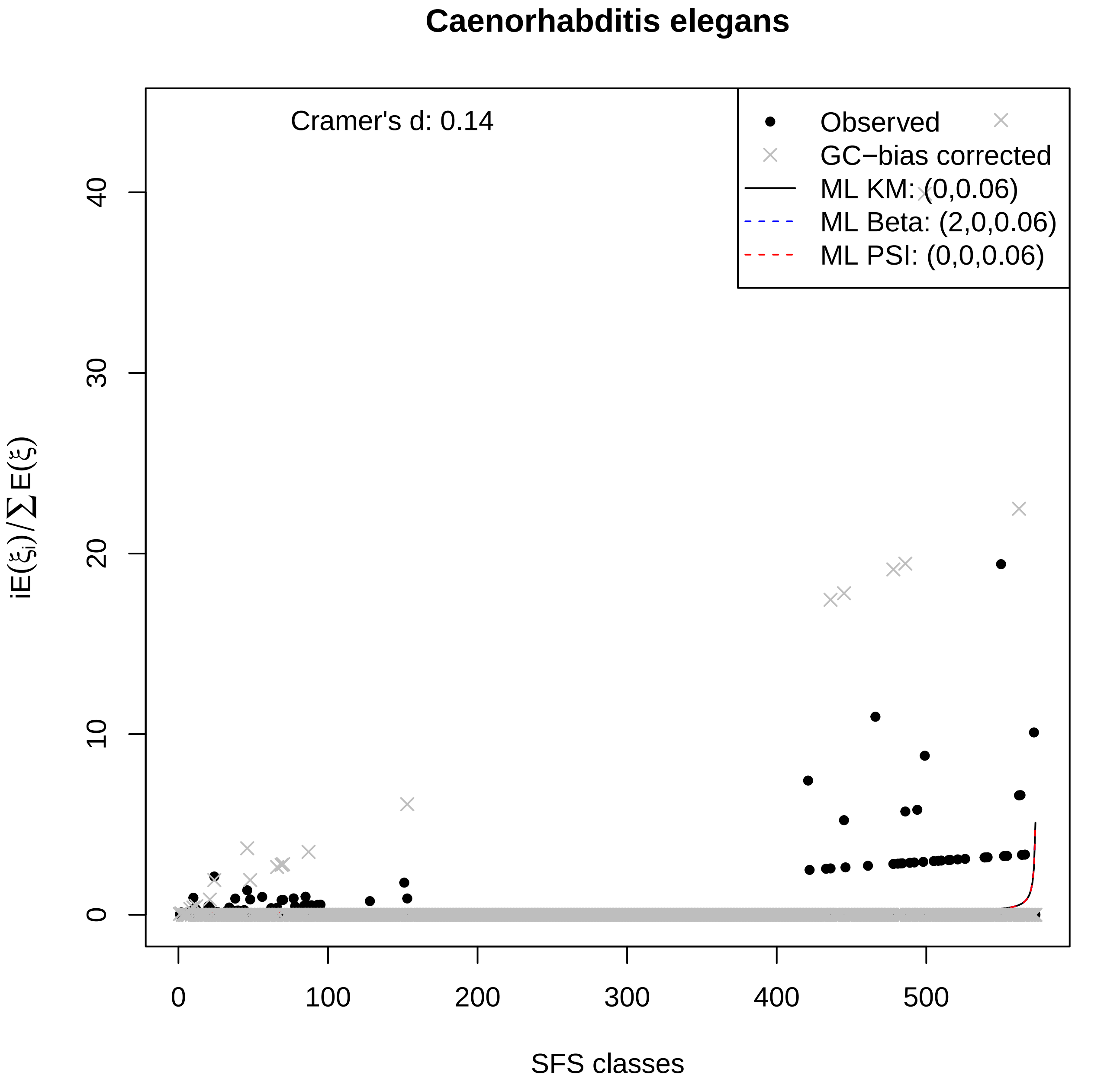

Caenorhabditis elegans
Cramer's d: 0.14
Observed
GC−bias corrected
ML KM: (0,0.06)
ML Beta: (2,0,0.06)
ML PSI: (0,0,0.06)
$iE(\xi_i)/\sum E(\xi)$
SFS classes
0
10
20
30
40
0
100
200
300
400
500

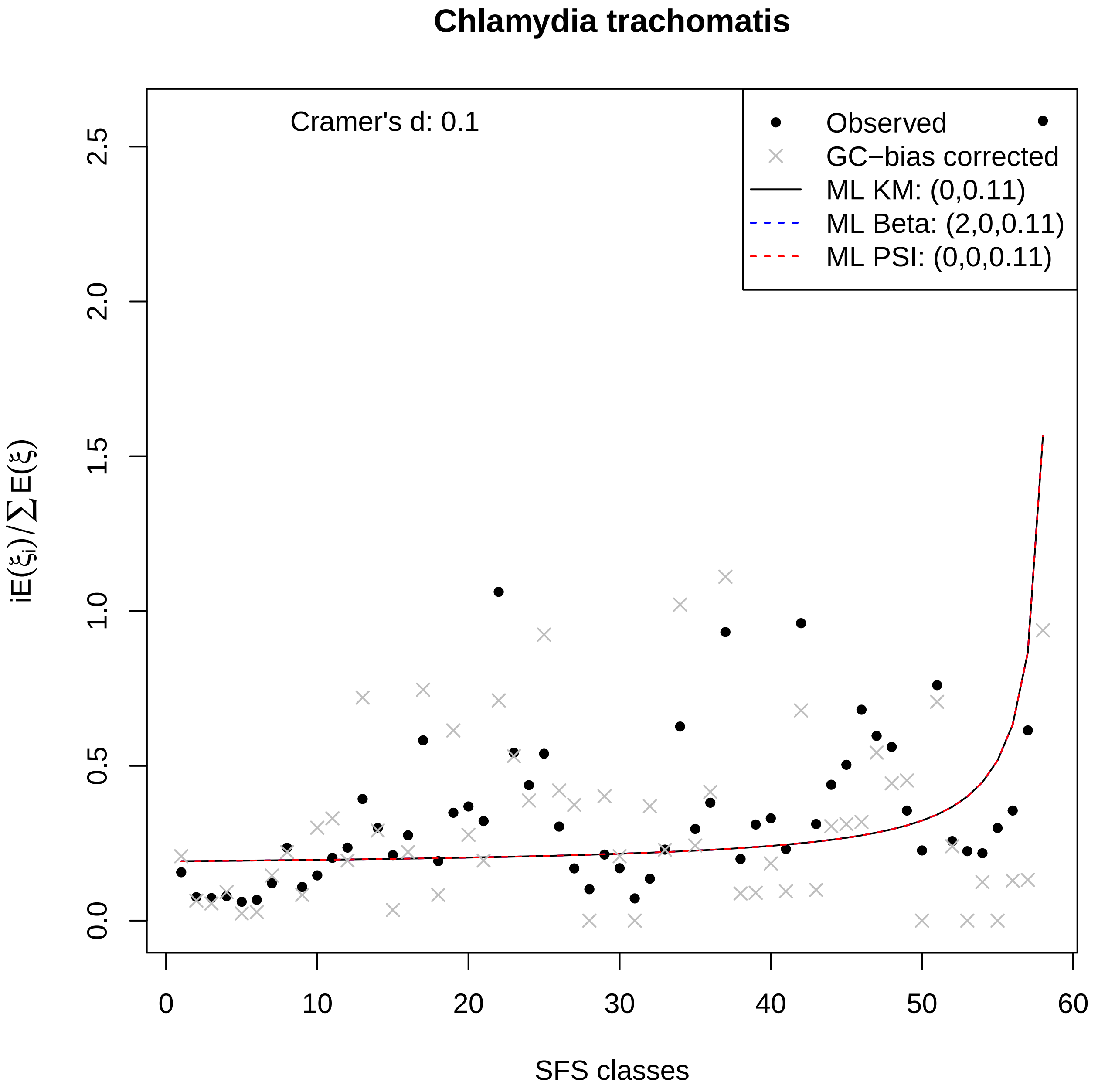

Chlamydia trachomatis
Cramer's d: 0.1
Observed
GC−bias corrected
ML KM: (0,0.11)
ML Beta: (2,0,0.11)
ML PSI: (0,0,0.11)
0.0
0.5
1.0
1.5
2.0
2.5
0
10
20
30
40
50
60
SFS classes

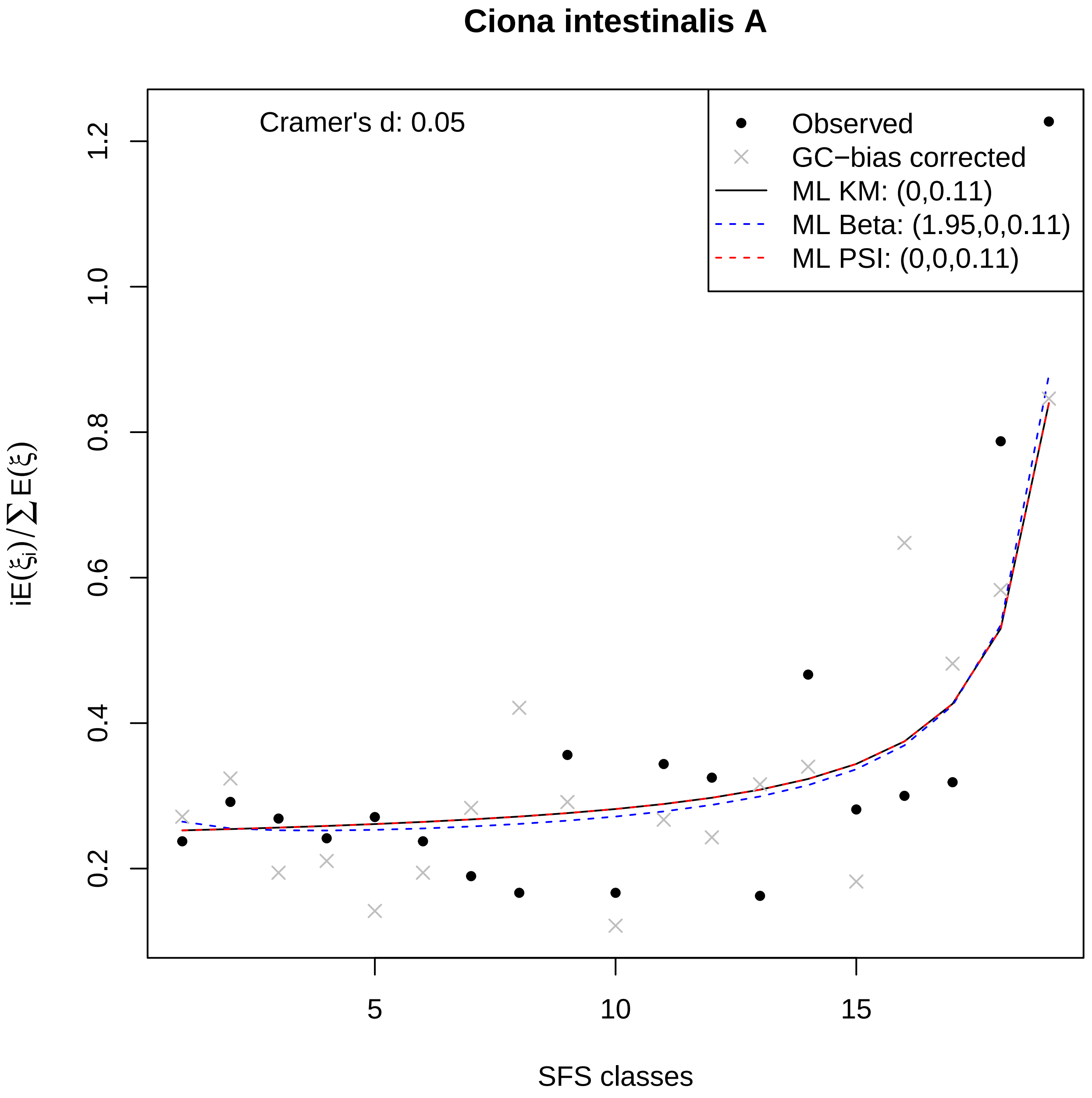

Ciona intestinalis A
Cramer's d: 0.05
Observed
GC−bias corrected
ML KM: (0,0.11)
ML Beta: (1.95,0,0.11)
ML PSI: (0,0,0.11)
$iE(\xi_i)/\sum E(\xi)$
0.2
0.4
0.6
0.8
1.0
1.2
5
10
15
SFS classes

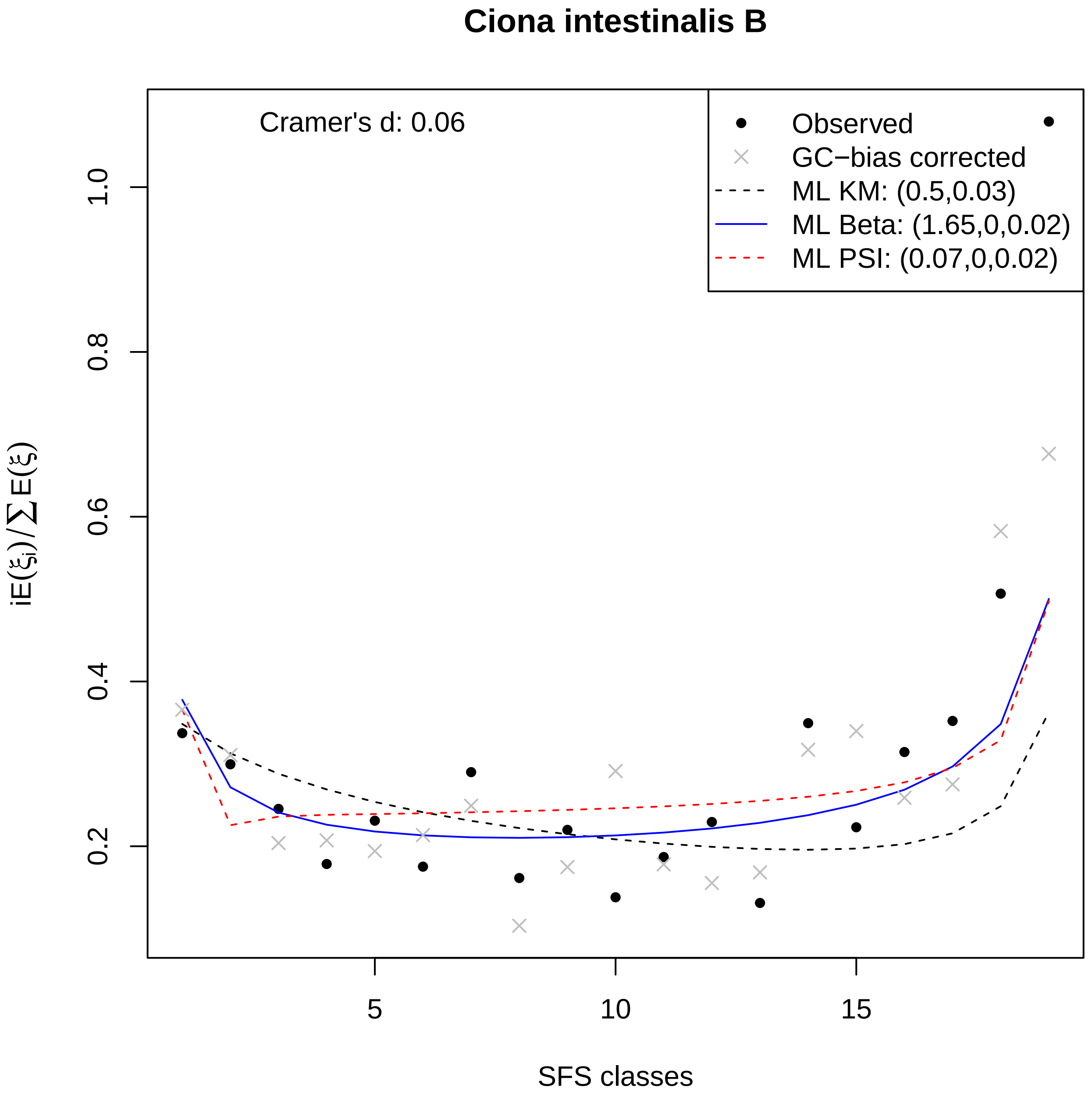
Ciona intestinalis B
Cramer's d: 0.06
Observed
GC−bias corrected
ML KM: (0.5,0.03)
ML Beta: (1.65,0,0.02)
ML PSI: (0.07,0,0.02)
iE(ξi)/∑E(ξ)
SFS classes
0.2
0.4
0.6
0.8
1.0
5
10
15

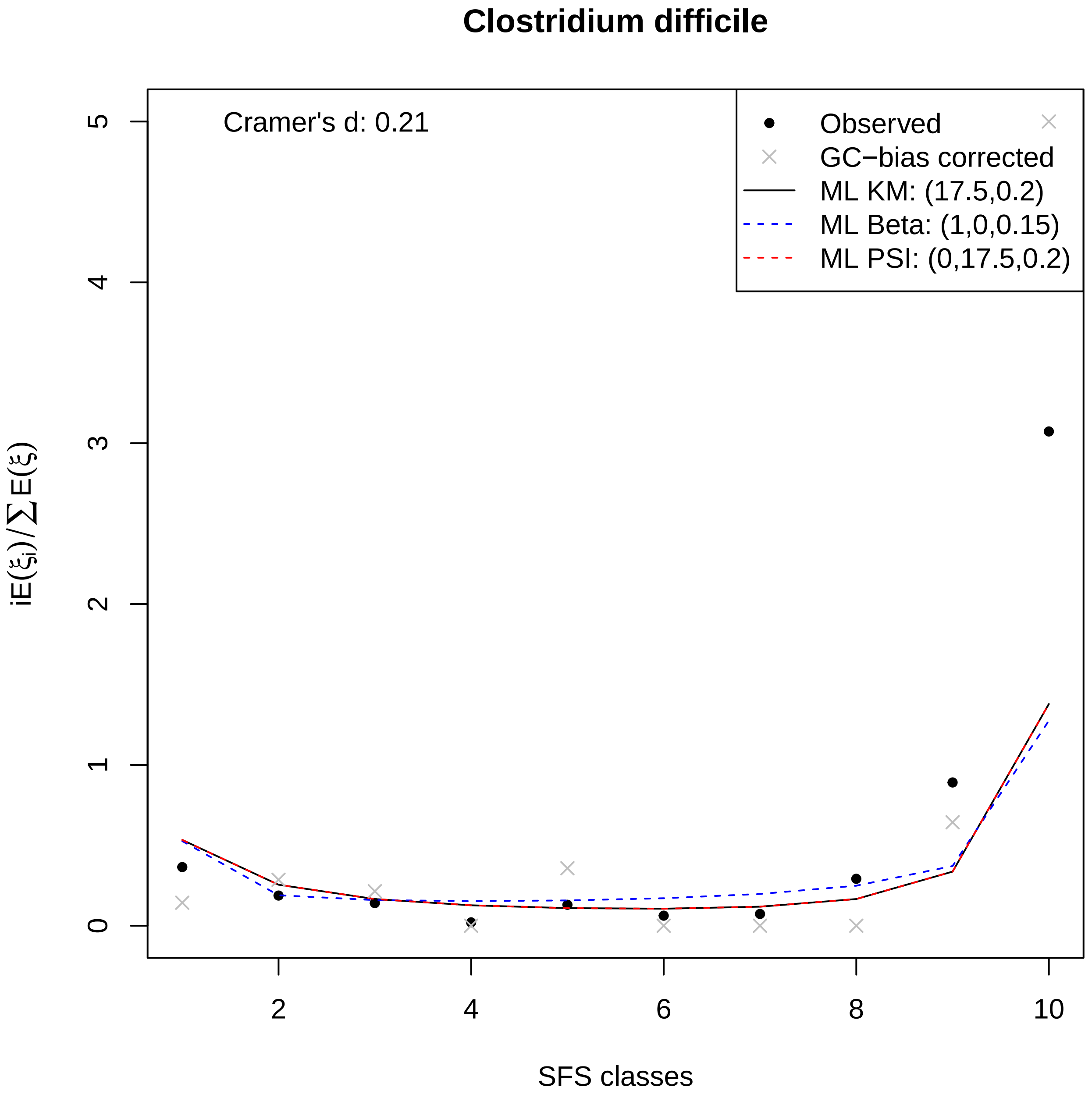
Clostridium difficile
Cramer's d: 0.21
Observed
GC−bias corrected
ML KM: (17.5,0.2)
ML Beta: (1,0,0.15)
ML PSI: (0,17.5,0.2)
iE(ξi)/∑E(ξ)
SFS classes
0
1
2
3
4
5
2
4
6
8
10

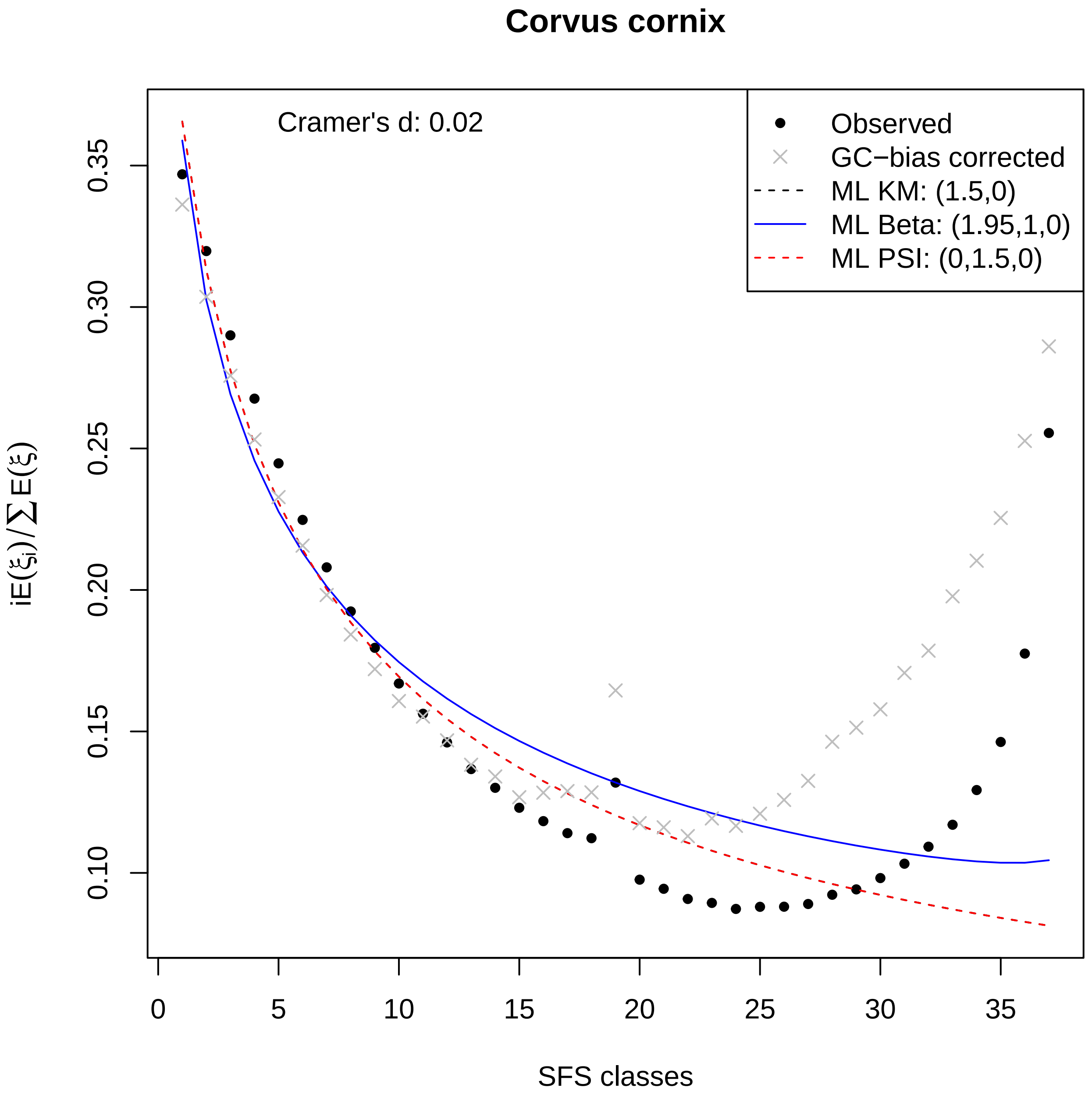

Corvus cornix
Cramer's d: 0.02
Observed
GC−bias corrected
ML KM: (1.5,0)
ML Beta: (1.95,1,0)
ML PSI: (0,1.5,0)
0.35
0.30
0.25
0.20
0.15
0.10
0
5
10
15
20
25
30
35
SFS classes
$iE(\xi_i)/\sum E(\xi)$

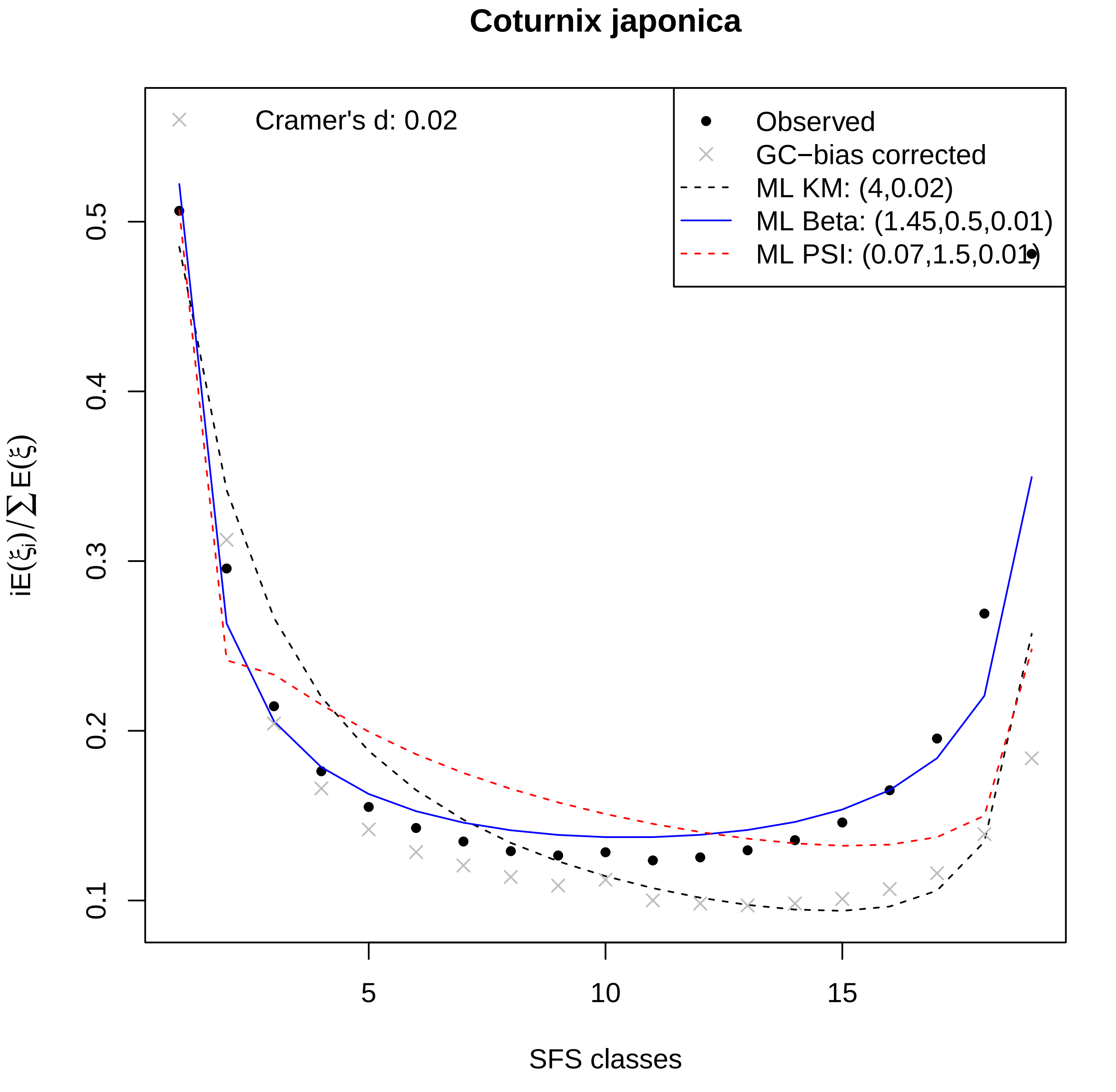

Coturnix japonica
Cramer's d: 0.02
Observed
GC−bias corrected
ML KM: (4,0.02)
ML Beta: (1.45,0.5,0.01)
ML PSI: (0.07,1.5,0.01)
0.1
0.2
0.3
0.4
0.5
5
10
15
$iE(\xi_i)/\sum E(\xi)$
SFS classes

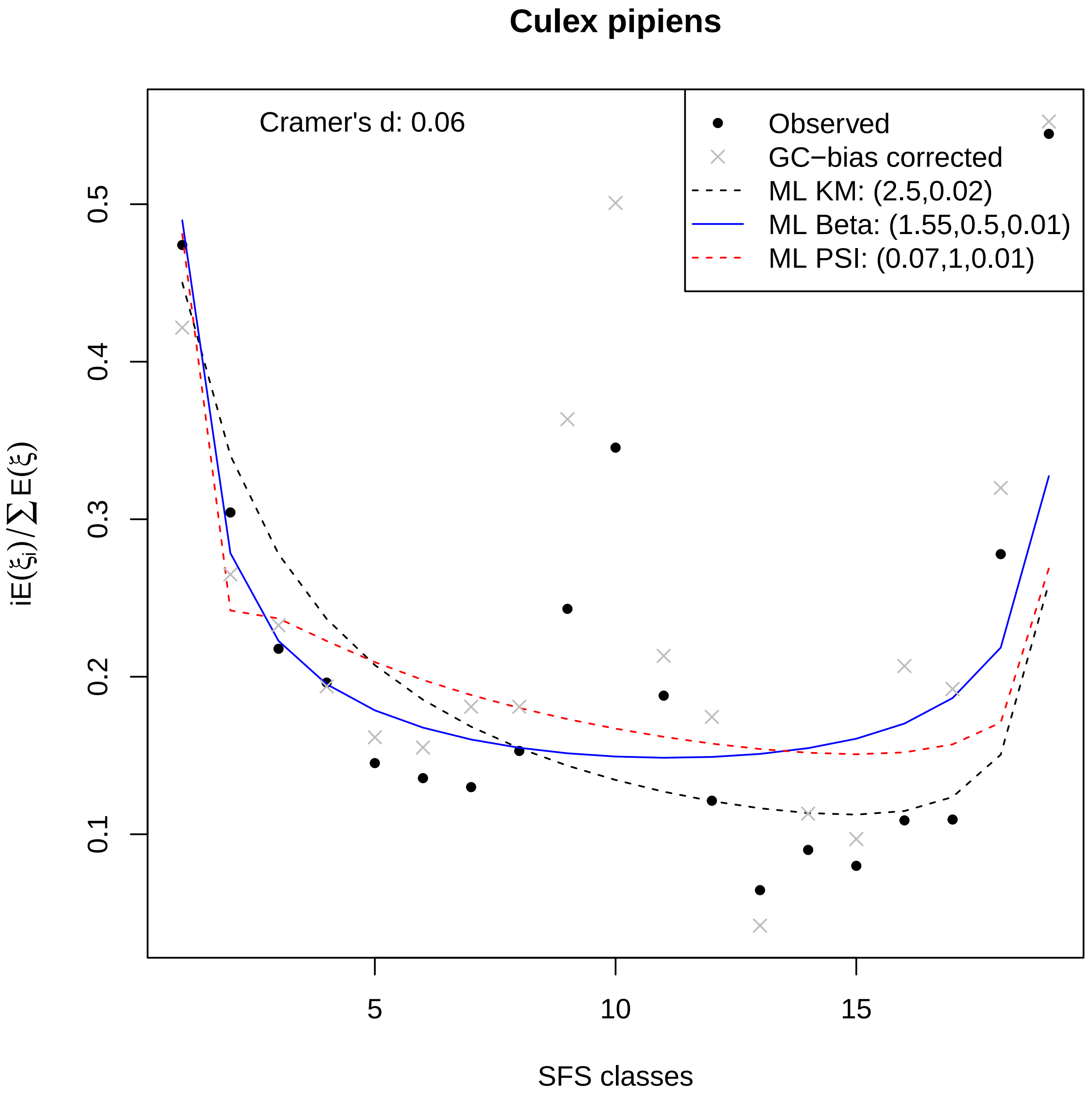
Culex pipiens
Cramer's d: 0.06
Observed
GC−bias corrected
ML KM: (2.5,0.02)
ML Beta: (1.55,0.5,0.01)
ML PSI: (0.07,1,0.01)
0.1
0.2
0.3
0.4
0.5
5
10
15
SFS classes
iE(ξi)/∑E(ξ)

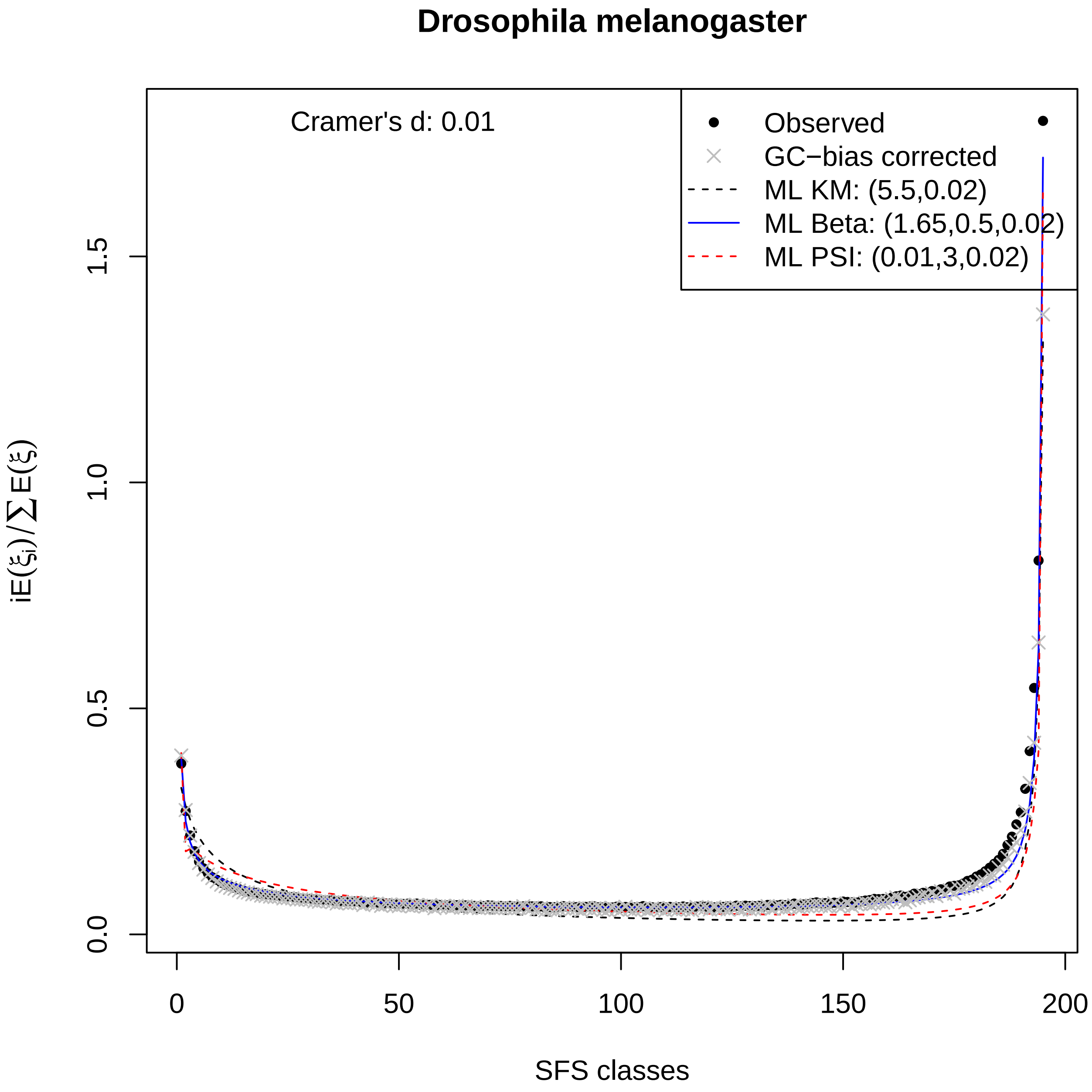
Drosophila melanogaster
Cramer's d: 0.01
Observed
GC−bias corrected
ML KM: (5.5,0.02)
ML Beta: (1.65,0.5,0.02)
ML PSI: (0.01,3,0.02)
$iE(\xi_i)/\sum E(\xi)$
0.0
0.5
1.0
1.5
0
50
100
150
200
SFS classes

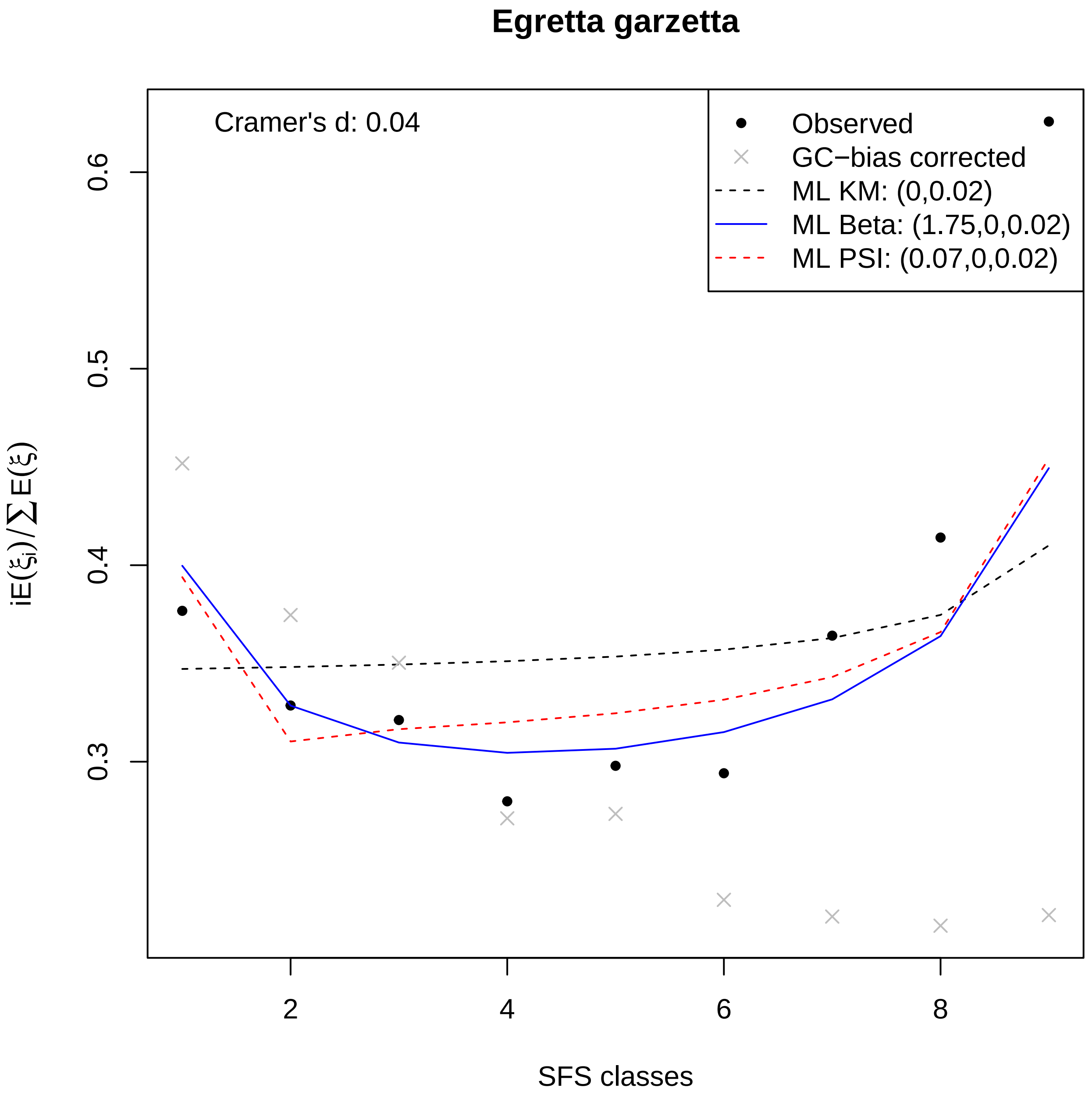

Egretta garzetta
Cramer's d: 0.04
Observed
GC−bias corrected
ML KM: (0,0.02)
ML Beta: (1.75,0,0.02)
ML PSI: (0.07,0,0.02)
0.3
0.4
0.5
0.6
2
4
6
8
SFS classes
$iE(\xi_i)/\sum E(\xi)$

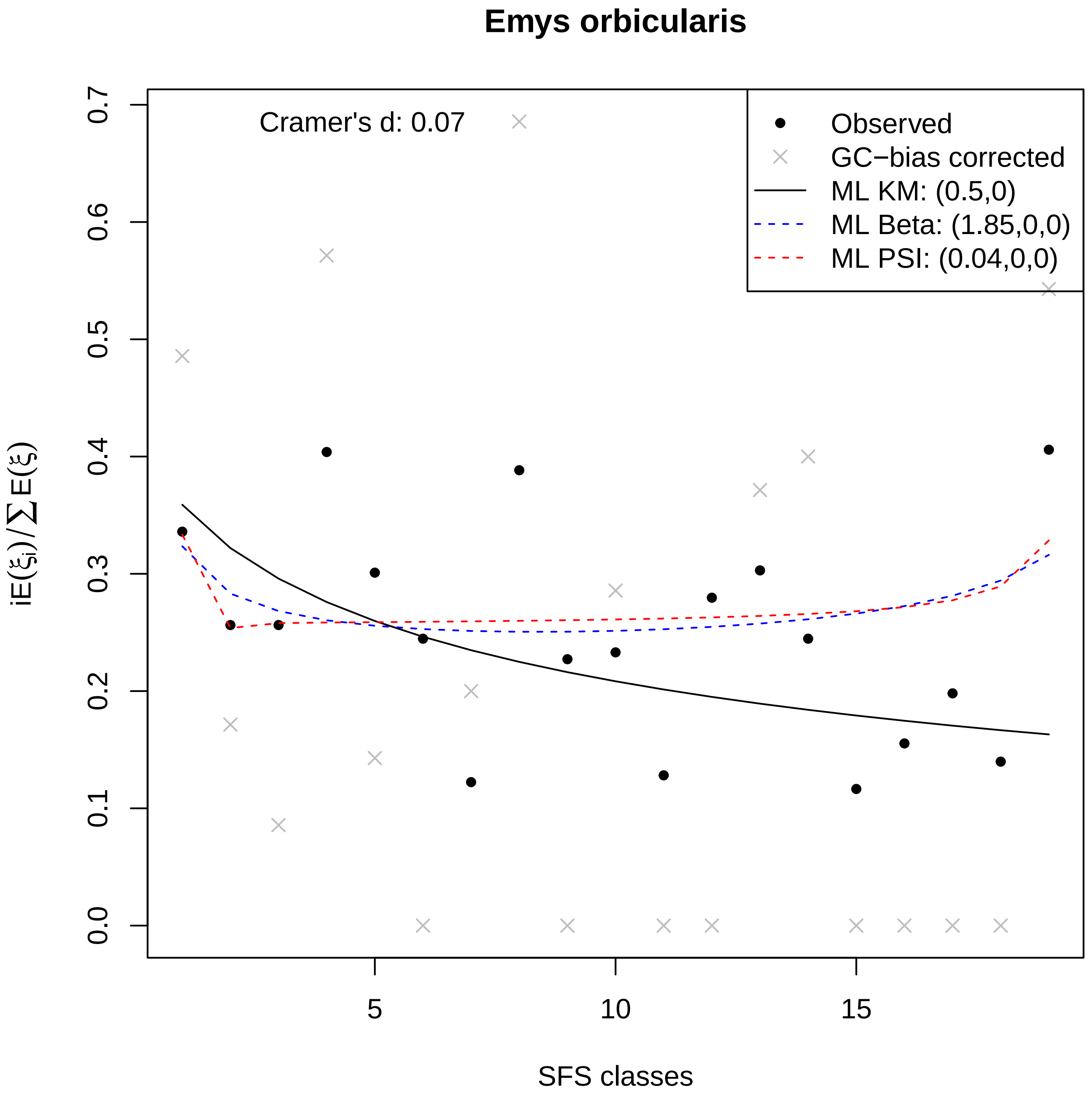

Emys orbicularis
Cramer's d: 0.07
Observed
GC−bias corrected
ML KM: (0.5,0)
ML Beta: (1.85,0,0)
ML PSI: (0.04,0,0)
0.0
0.1
0.2
0.3
0.4
0.5
0.6
0.7
5
10
15
SFS classes
iE(ξi)/∑E(ξ)

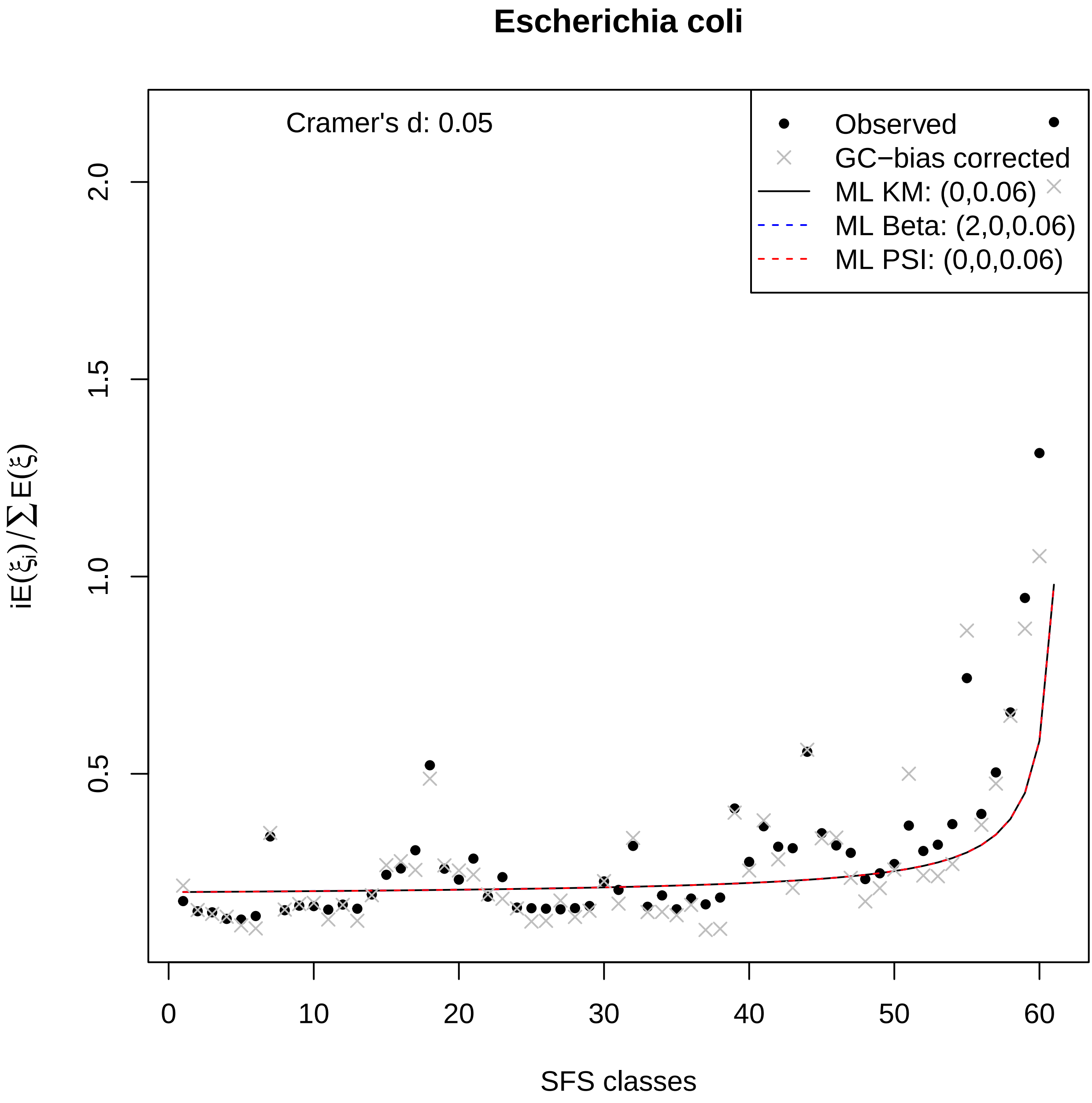

Escherichia coli
Cramer's d: 0.05
Observed
GC−bias corrected
ML KM: (0,0.06)
ML Beta: (2,0,0.06)
ML PSI: (0,0,0.06)
iE(ξi)/∑E(ξ)
SFS classes
0
10
20
30
40
50
60
0.5
1.0
1.5
2.0

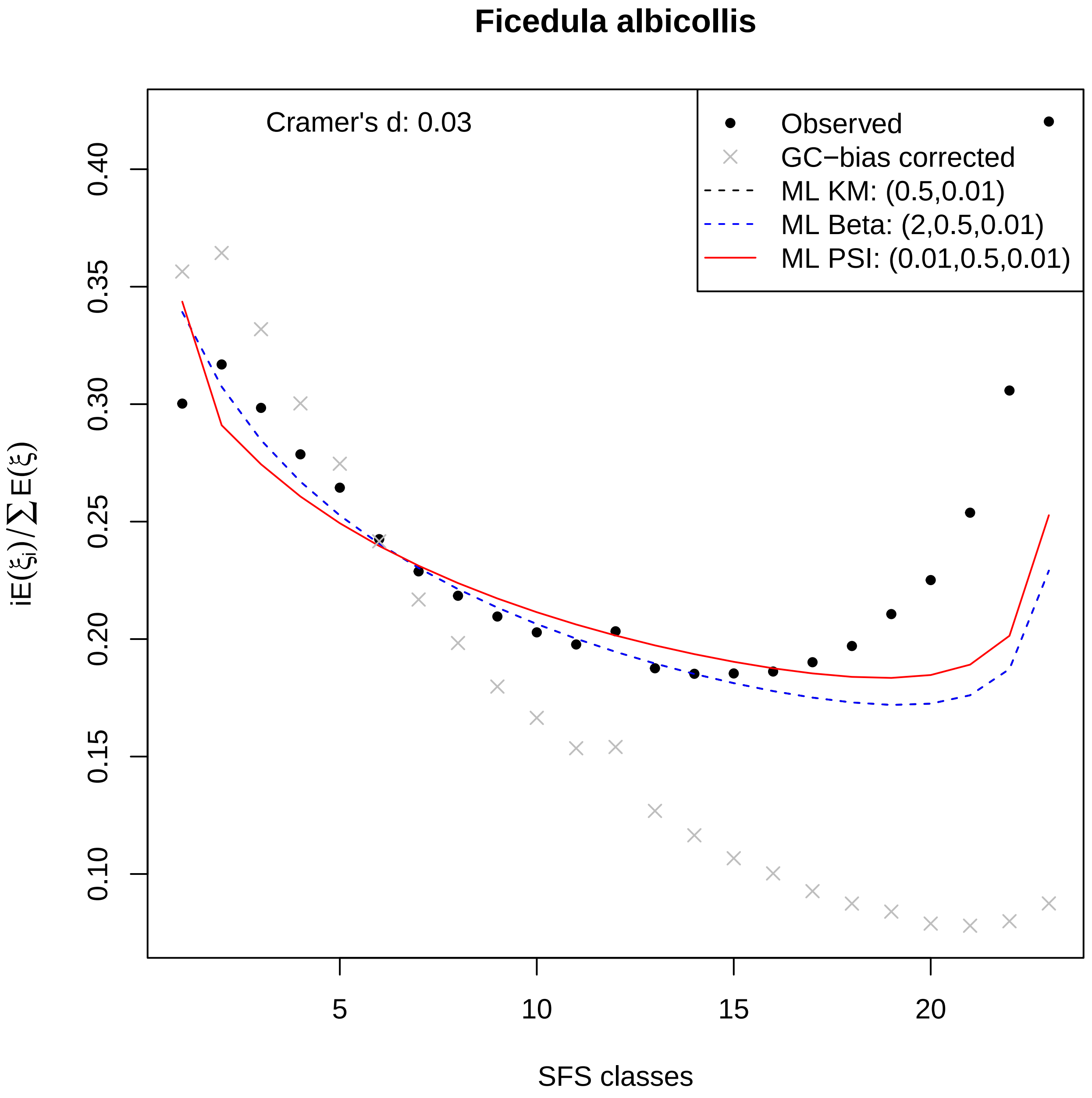

Ficedula albicollis
Cramer's d: 0.03
Observed
GC−bias corrected
ML KM: (0.5,0.01)
ML Beta: (2,0.5,0.01)
ML PSI: (0.01,0.5,0.01)
0.10
0.15
0.20
0.25
0.30
0.35
0.40
$iE(\xi_i)/\sum E(\xi)$
5
10
15
20
SFS classes

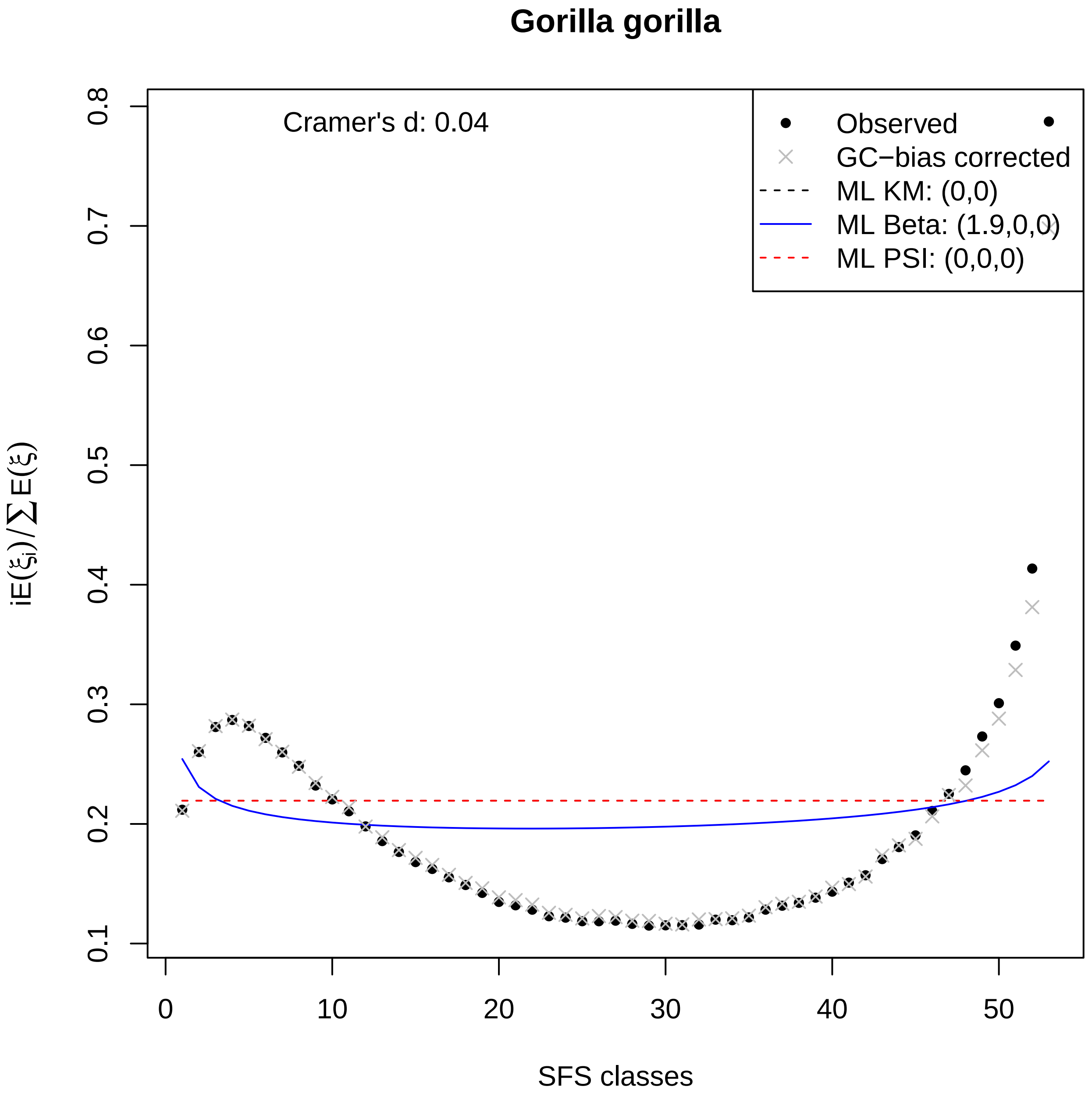
Gorilla gorilla
Cramer's d: 0.04
Observed
GC−bias corrected
ML KM: (0,0)
ML Beta: (1.9,0,0)
ML PSI: (0,0,0)
iE(ξi)/∑E(ξ)
SFS classes
0.1
0.2
0.3
0.4
0.5
0.6
0.7
0.8
0
10
20
30
40
50

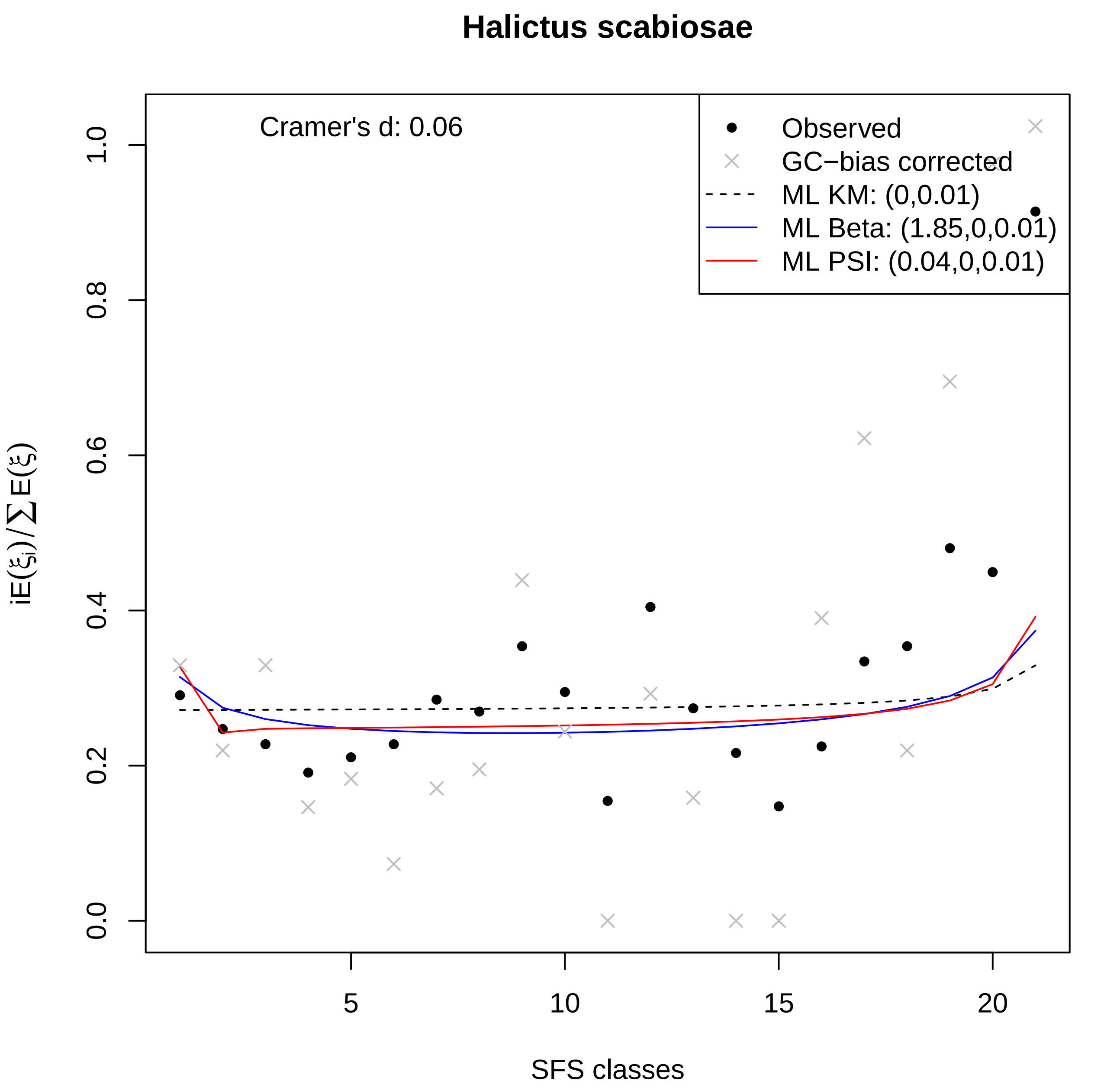

Halictus scabiosae
Cramer's d: 0.06
Observed
GC−bias corrected
ML KM: (0,0.01)
ML Beta: (1.85,0,0.01)
ML PSI: (0.04,0,0.01)
0.0
0.2
0.4
0.6
0.8
1.0
5
10
15
20
SFS classes
iE(ξi)/∑E(ξ)

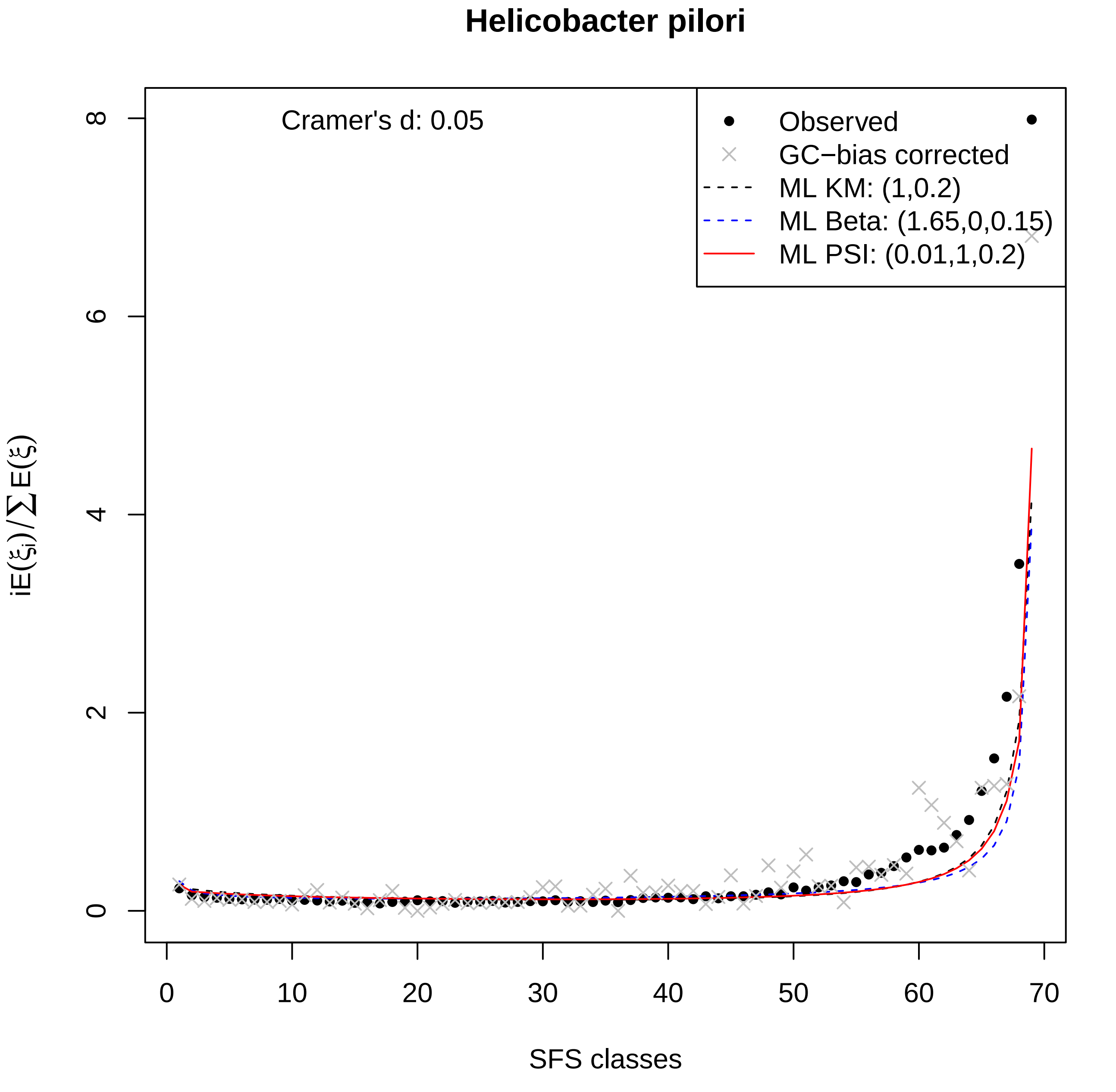

Helicobacter pilori
Cramer's d: 0.05
Observed
GC−bias corrected
ML KM: (1,0.2)
ML Beta: (1.65,0,0.15)
ML PSI: (0.01,1,0.2)
$iE(\xi_i)/\sum E(\xi)$
SFS classes
0
2
4
6
8
0
10
20
30
40
50
60
70

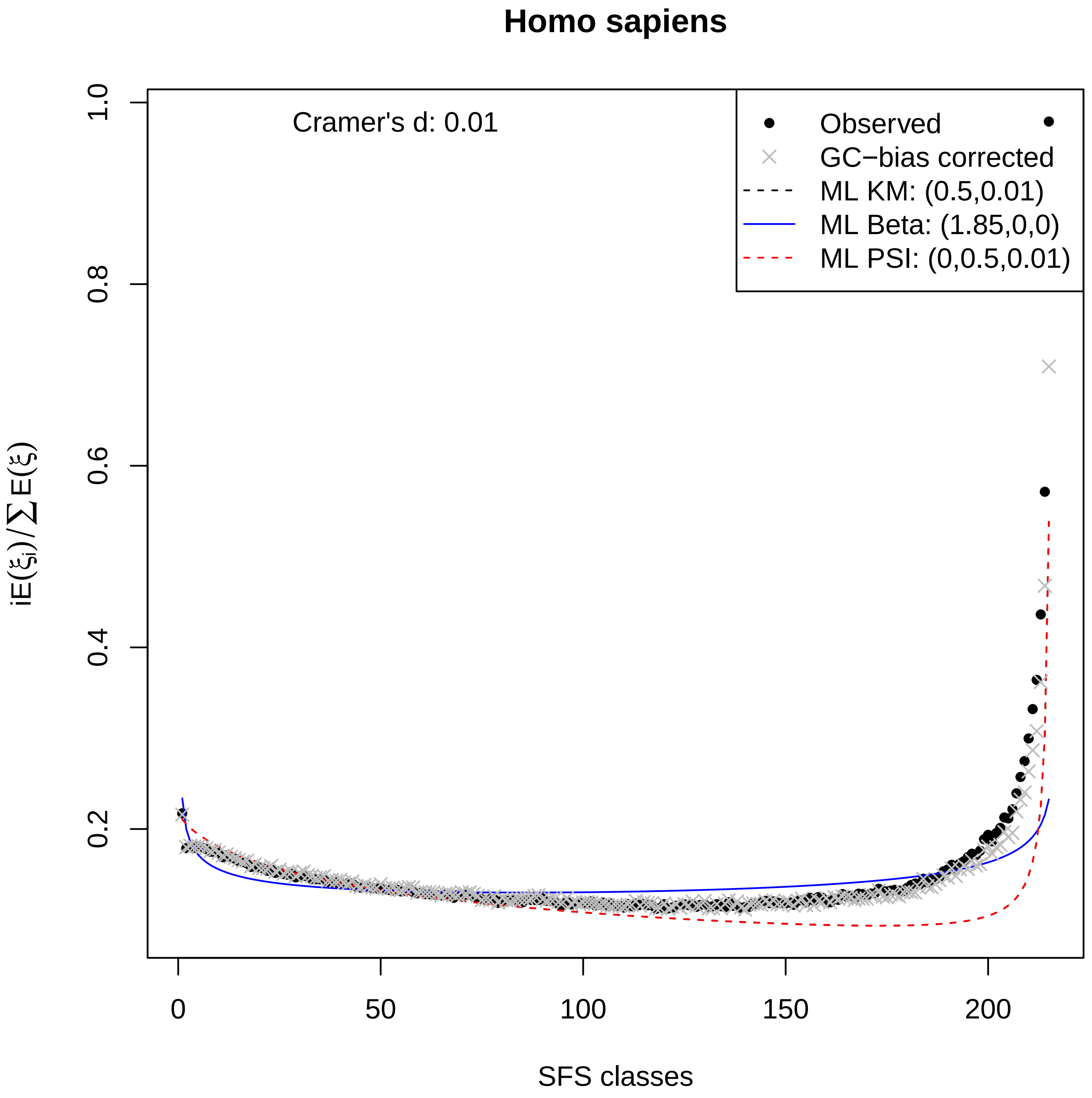

Homo sapiens
Cramer's d: 0.01
Observed
GC−bias corrected
ML KM: (0.5,0.01)
ML Beta: (1.85,0,0)
ML PSI: (0,0.5,0.01)
$iE(\xi_i)/\sum E(\xi)$
SFS classes
0
50
100
150
200
0.2
0.4
0.6
0.8
1.0

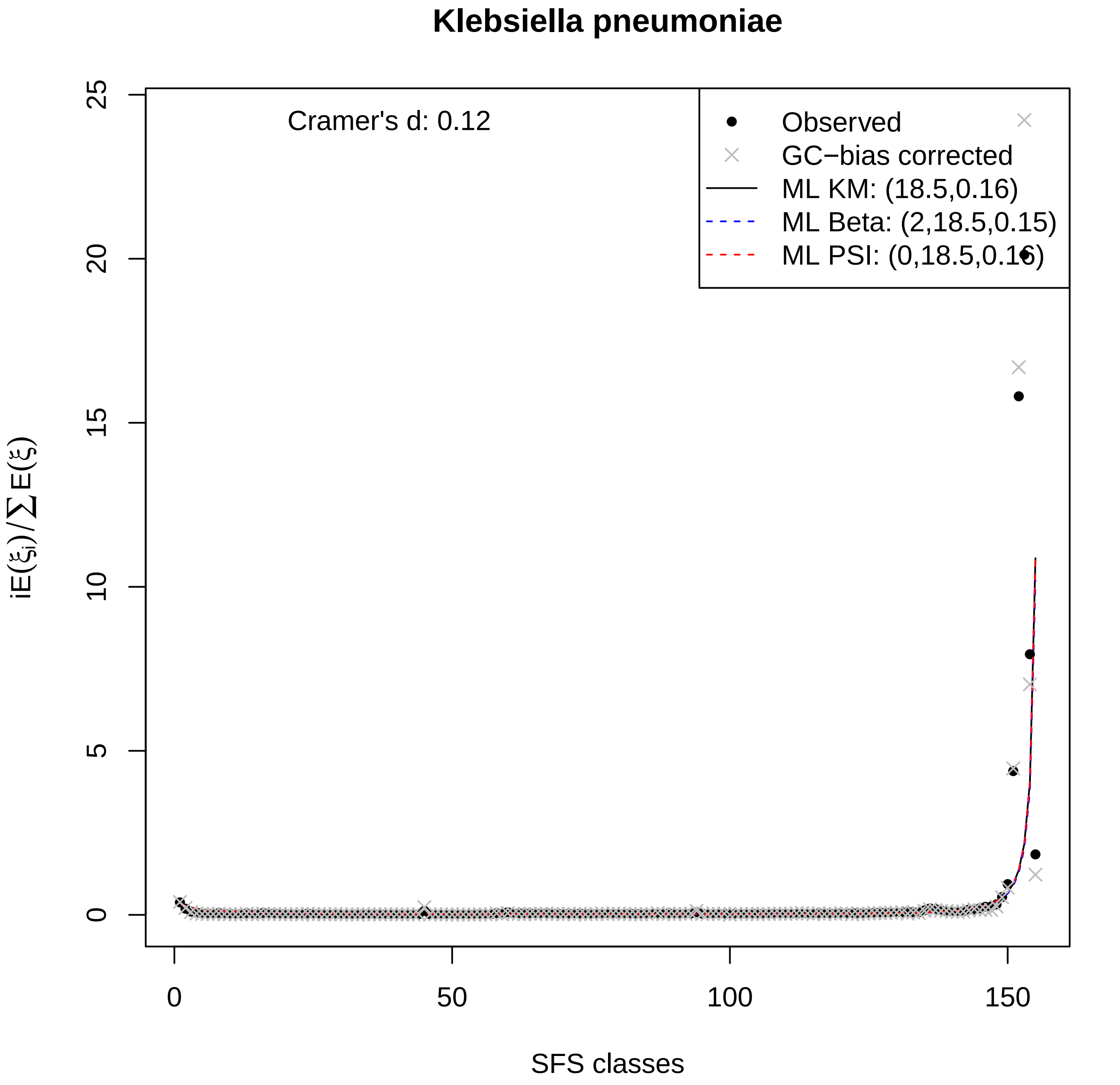

Klebsiella pneumoniae
Cramer's d: 0.12
Observed
GC−bias corrected
ML KM: (18.5,0.16)
ML Beta: (2,18.5,0.15)
ML PSI: (0,18.5,0.16)
0
5
10
15
20
25
0
50
100
150
SFS classes
$iE(\xi_i)/\sum E(\xi)$

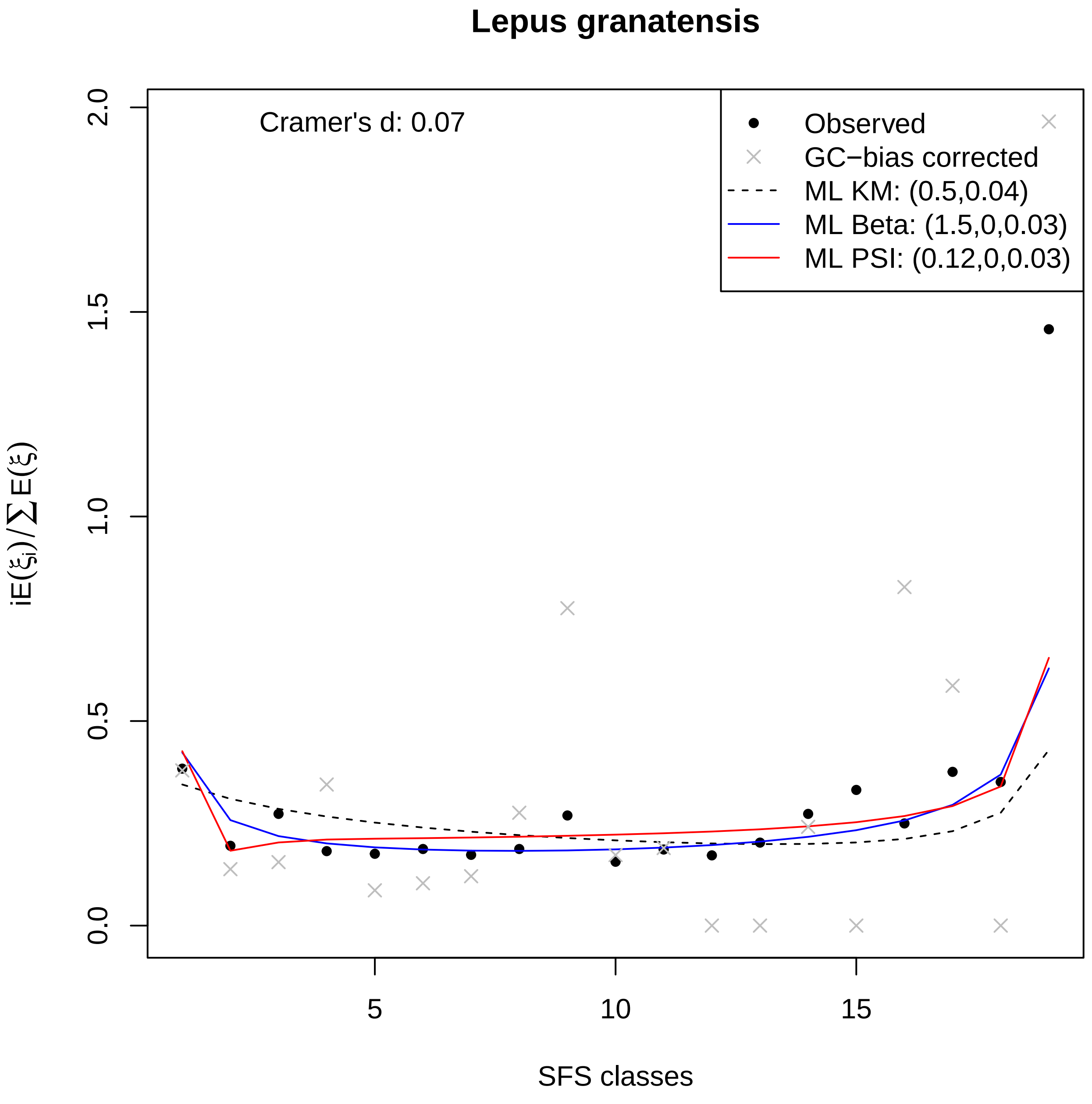

Lepus granatensis
Cramer's d: 0.07
Observed
GC−bias corrected
ML KM: (0.5,0.04)
ML Beta: (1.5,0,0.03)
ML PSI: (0.12,0,0.03)
$iE(\xi_i)/\sum E(\xi)$
SFS classes
0.0
0.5
1.0
1.5
2.0
5
10
15

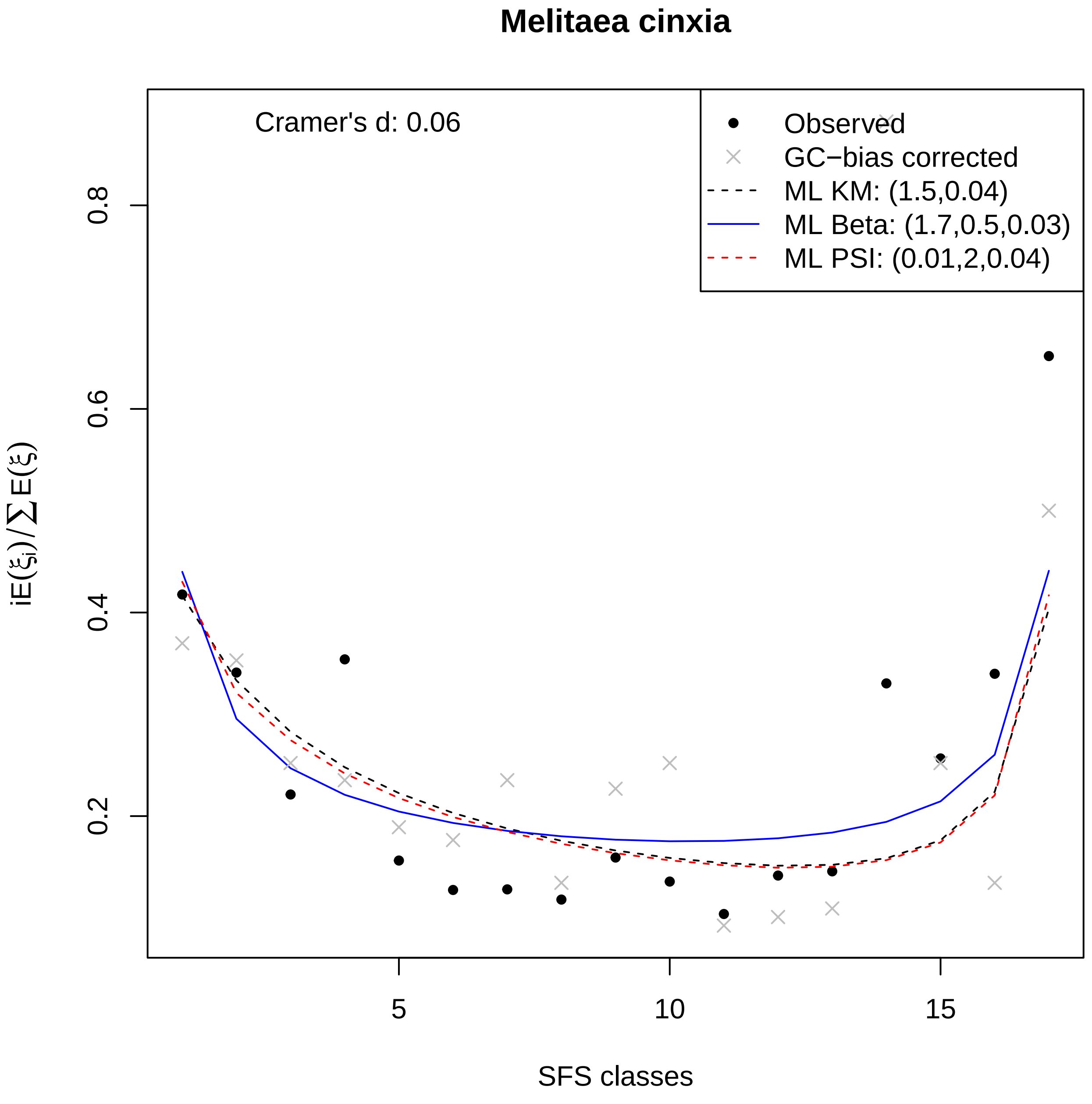
Melitaea cinxia
Cramer's d: 0.06
Observed
GC−bias corrected
ML KM: (1.5,0.04)
ML Beta: (1.7,0.5,0.03)
ML PSI: (0.01,2,0.04)
0.2
0.4
0.6
0.8
5
10
15
SFS classes
$iE(\xi_i)/\sum E(\xi)$

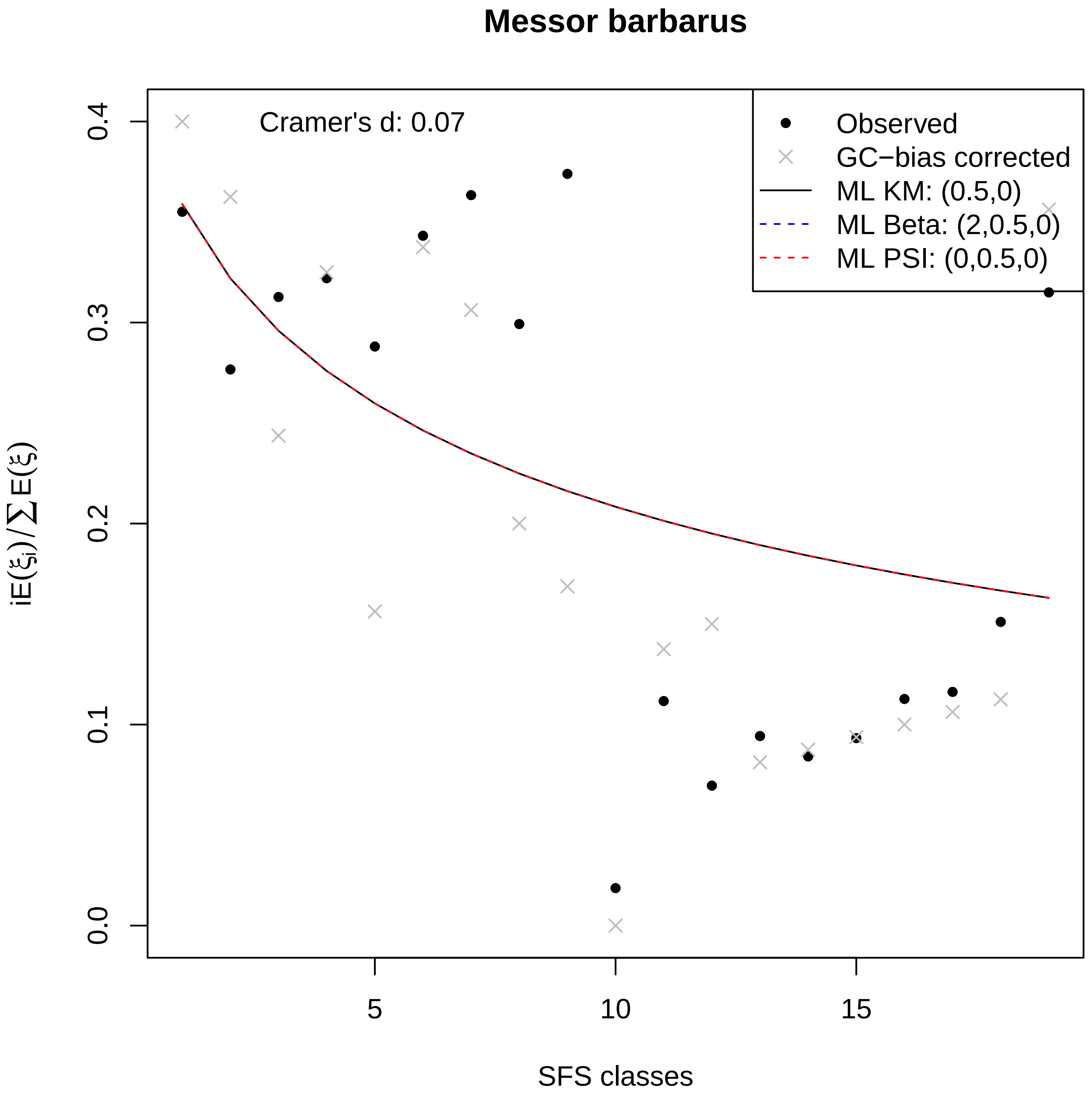

Messor barbarus
Cramer's d: 0.07
Observed
GC−bias corrected
ML KM: (0.5,0)
ML Beta: (2,0.5,0)
ML PSI: (0,0.5,0)
0.0
0.1
0.2
0.3
0.4
5
10
15
SFS classes
$iE(\xi_i)/\sum E(\xi)$

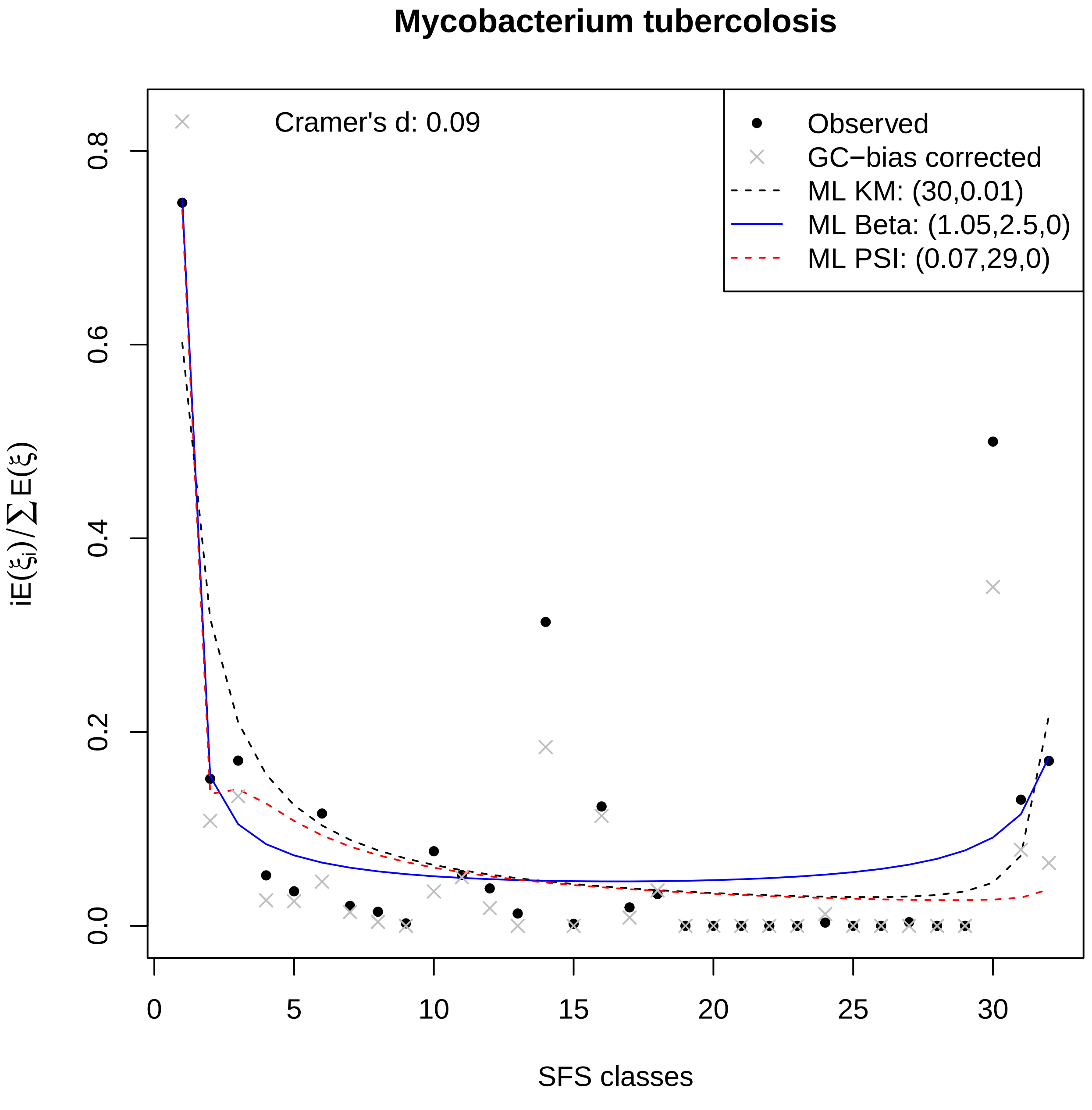

Mycobacterium tubercolosis
Cramer's d: 0.09
Observed
GC−bias corrected
ML KM: (30,0.01)
ML Beta: (1.05,2.5,0)
ML PSI: (0.07,29,0)
$iE(\xi_i)/\sum E(\xi)$
0.0
0.2
0.4
0.6
0.8
0
5
10
15
20
25
30
SFS classes

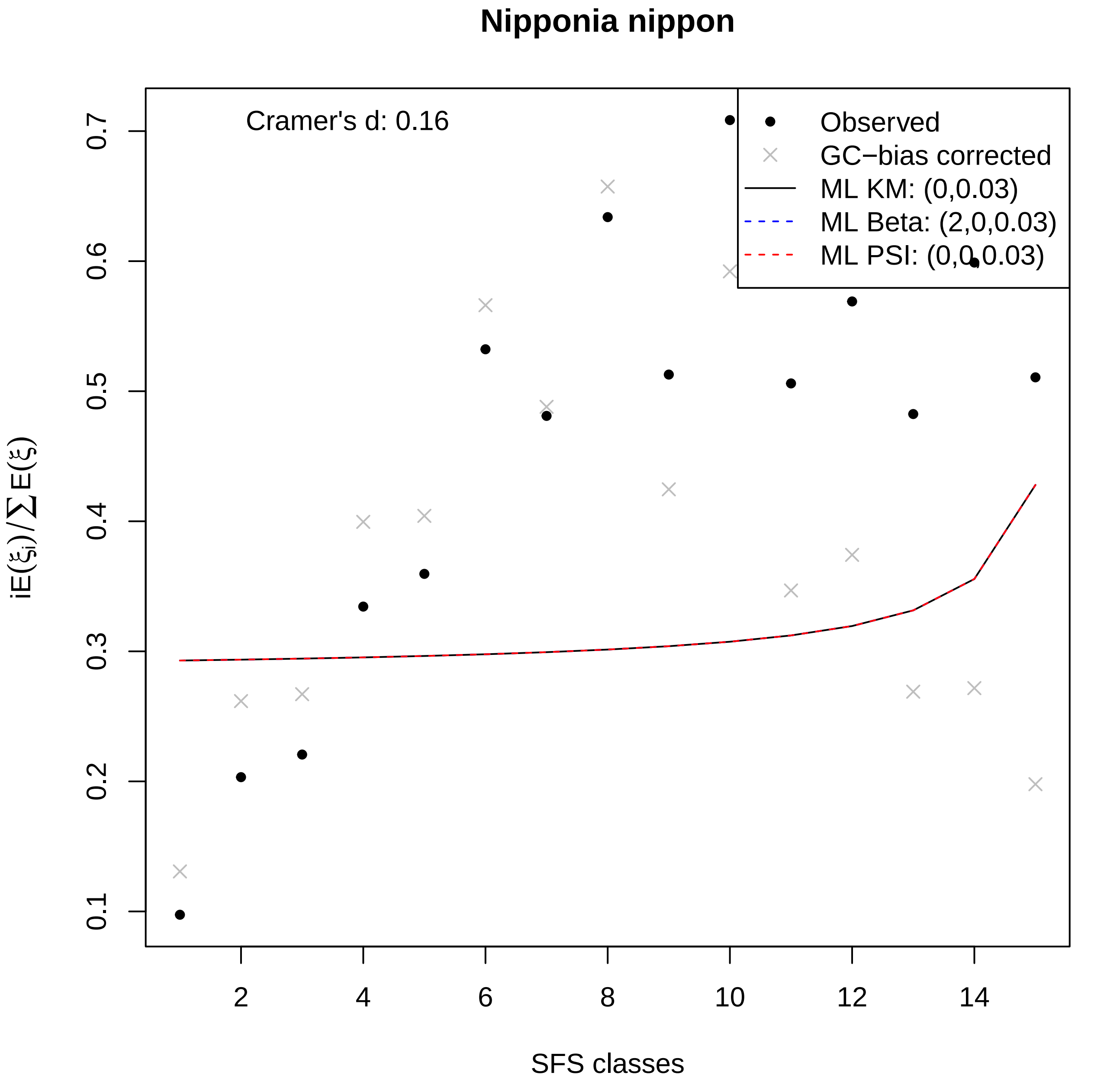

Nipponia nippon
Cramer's d: 0.16
Observed
GC−bias corrected
ML KM: (0,0.03)
ML Beta: (2,0,0.03)
ML PSI: (0,0,0.03)
0.1
0.2
0.3
0.4
0.5
0.6
0.7
2
4
6
8
10
12
14
SFS classes
$iE(\xi_i)/\sum E(\xi)$

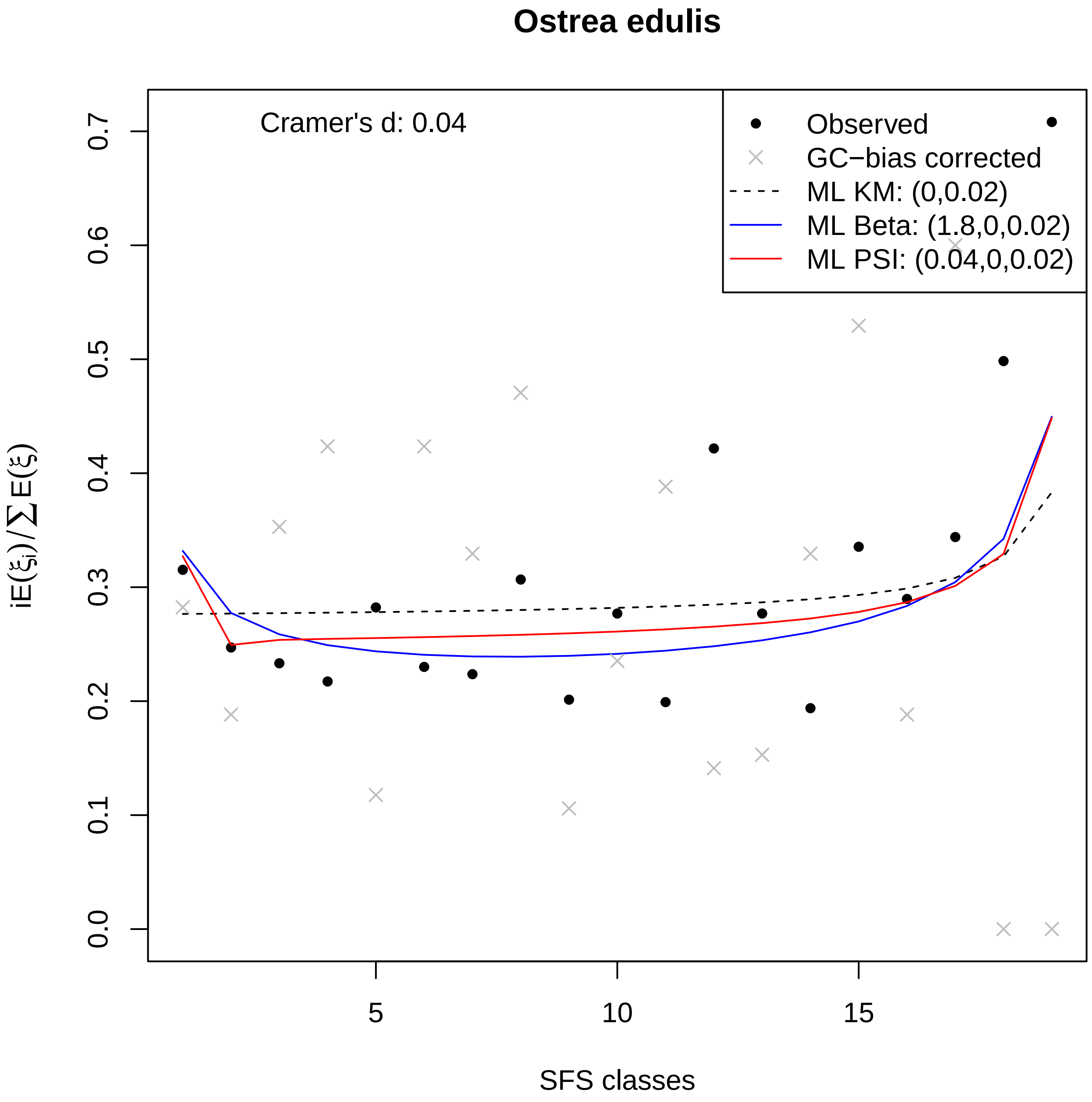
Ostrea edulis
Cramer's d: 0.04
Observed
GC−bias corrected
ML KM: (0,0.02)
ML Beta: (1.8,0,0.02)
ML PSI: (0.04,0,0.02)
0.0
0.1
0.2
0.3
0.4
0.5
0.6
0.7
$iE(\xi_i)/\sum E(\xi)$
5
10
15
SFS classes

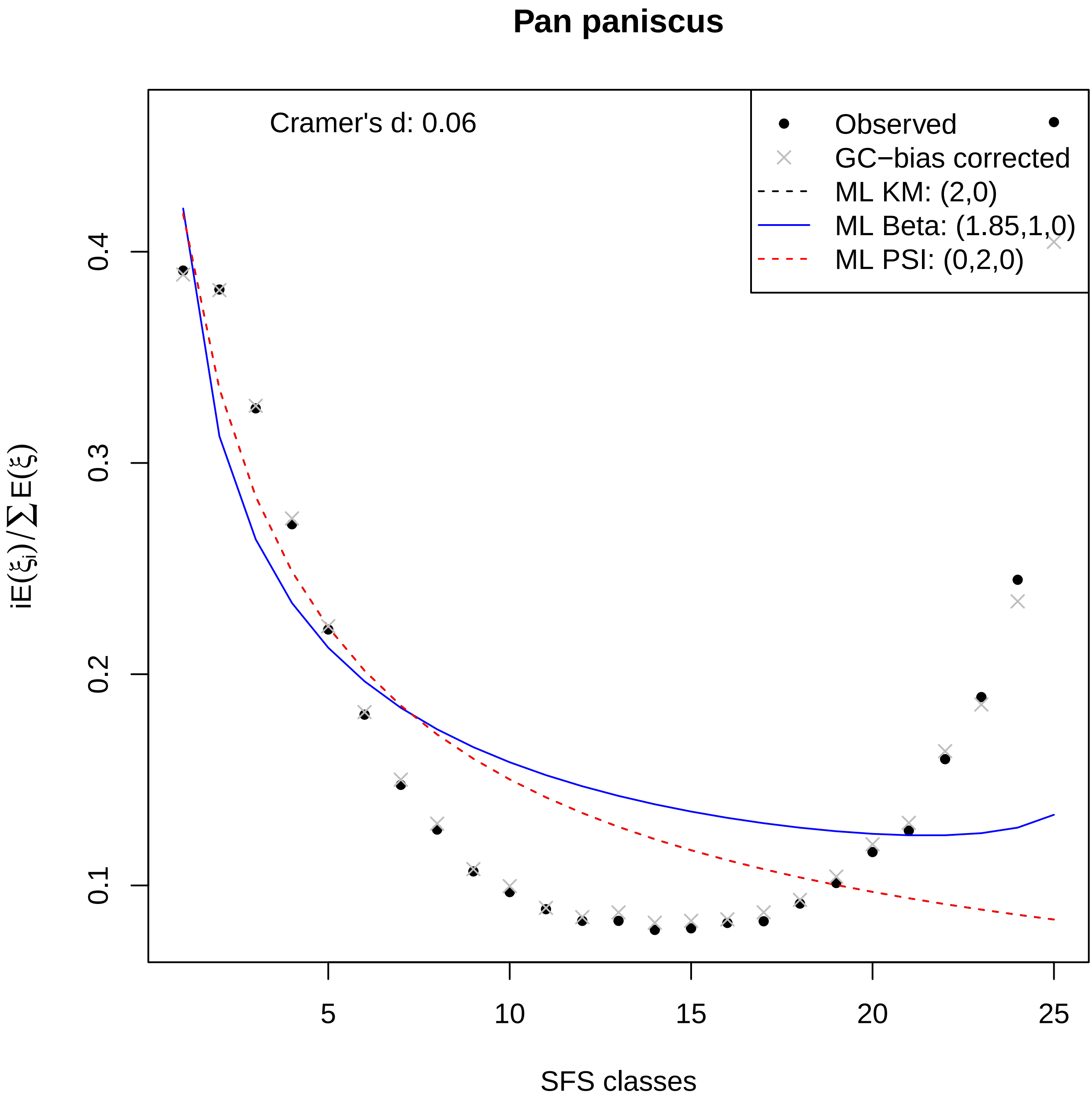

Pan paniscus
Cramer's d: 0.06
Observed
GC−bias corrected
ML KM: (2,0)
ML Beta: (1.85,1,0)
ML PSI: (0,2,0)
0.1
0.2
0.3
0.4
5
10
15
20
25
SFS classes
$iE(\xi_i)/\sum E(\xi)$

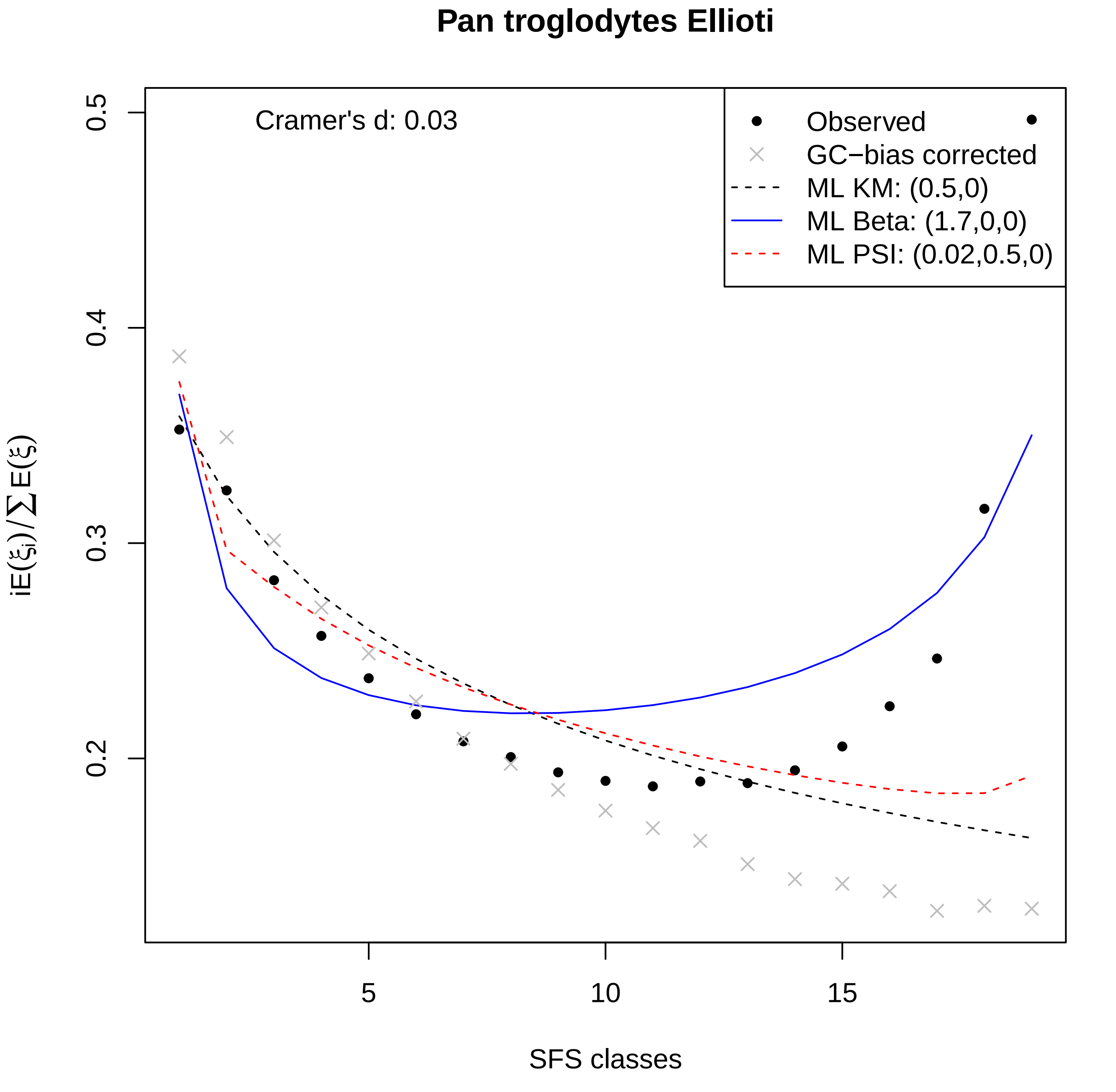

Pan troglodytes Ellioti
Cramer's d: 0.03
Observed
GC−bias corrected
ML KM: (0.5,0)
ML Beta: (1.7,0,0)
ML PSI: (0.02,0.5,0)
0.2
0.3
0.4
0.5
5
10
15
SFS classes
$iE(\xi_i)/\sum E(\xi)$

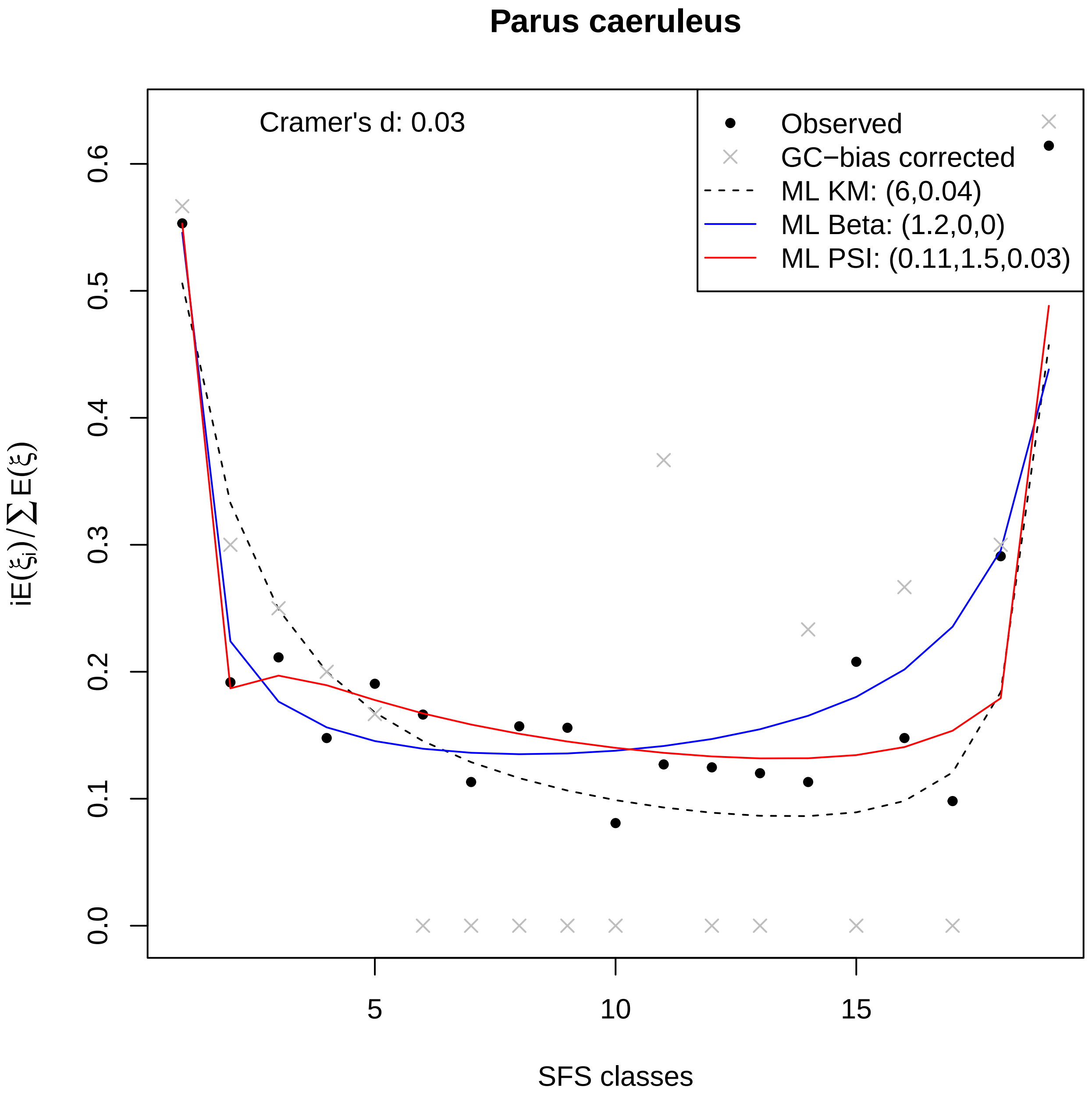

Parus caeruleus
Cramer's d: 0.03
Observed
GC−bias corrected
ML KM: (6,0.04)
ML Beta: (1.2,0,0)
ML PSI: (0.11,1.5,0.03)
0.0
0.1
0.2
0.3
0.4
0.5
0.6
5
10
15
SFS classes
iE(ξi)/∑E(ξ)

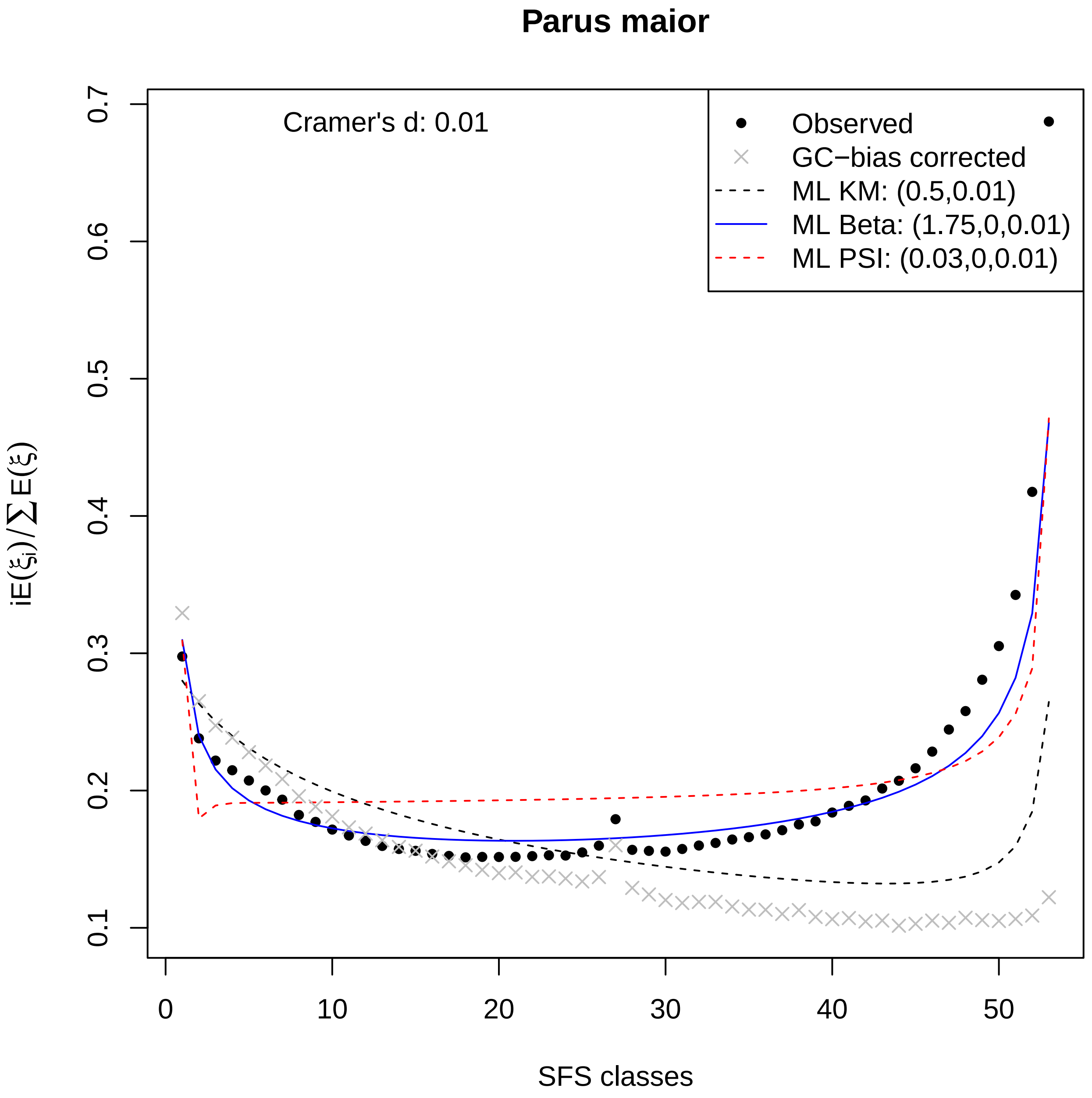

Parus maior
Cramer's d: 0.01
Observed
GC−bias corrected
ML KM: (0.5,0.01)
ML Beta: (1.75,0,0.01)
ML PSI: (0.03,0,0.01)
$iE(\xi_i)/\sum E(\xi)$
SFS classes
0.1
0.2
0.3
0.4
0.5
0.6
0.7
0
10
20
30
40
50

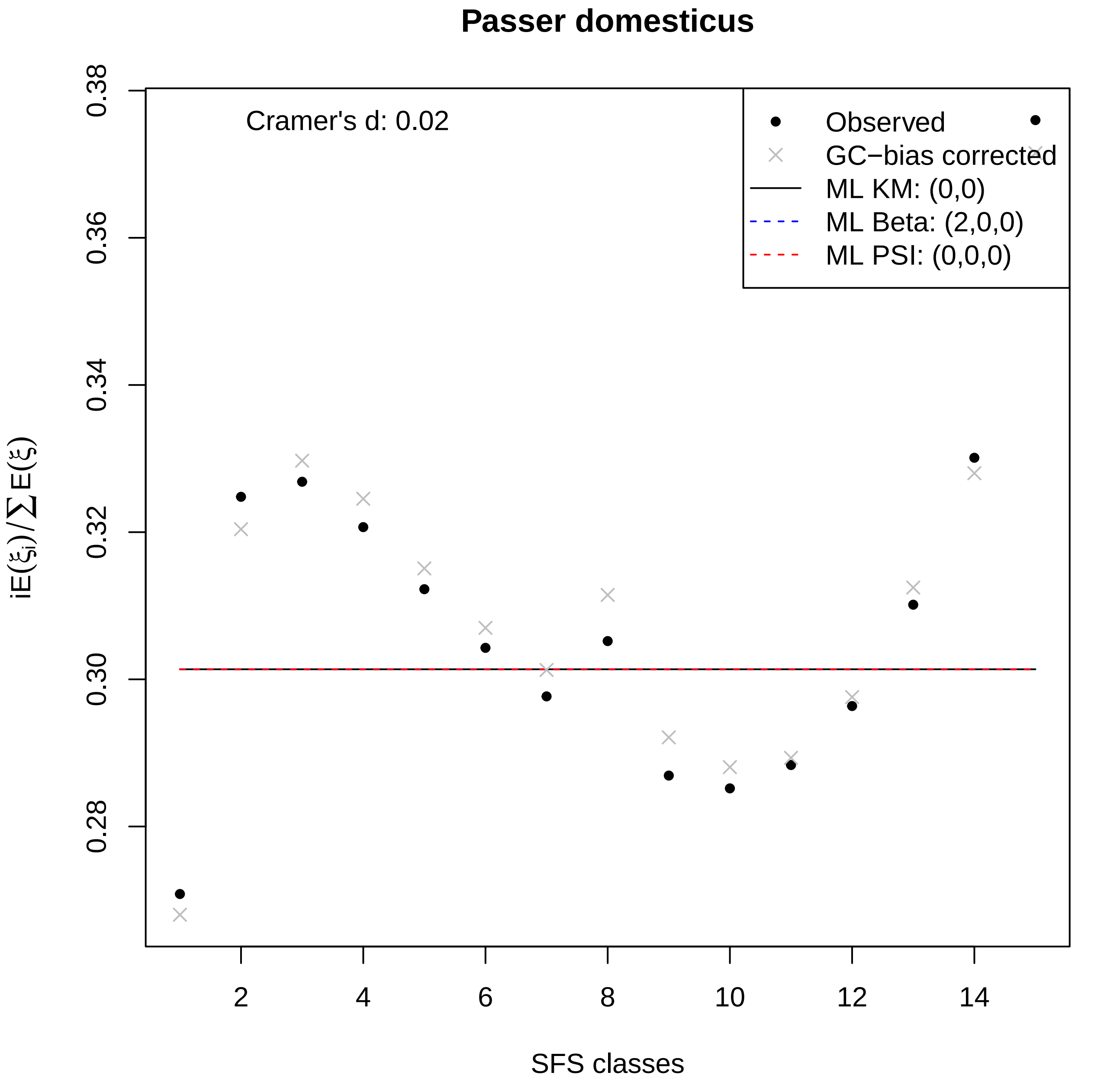

Passer domesticus
Cramer's d: 0.02
Observed
GC−bias corrected
ML KM: (0,0)
ML Beta: (2,0,0)
ML PSI: (0,0,0)
0.38
0.36
0.34
0.32
0.30
0.28
2
4
6
8
10
12
14
SFS classes

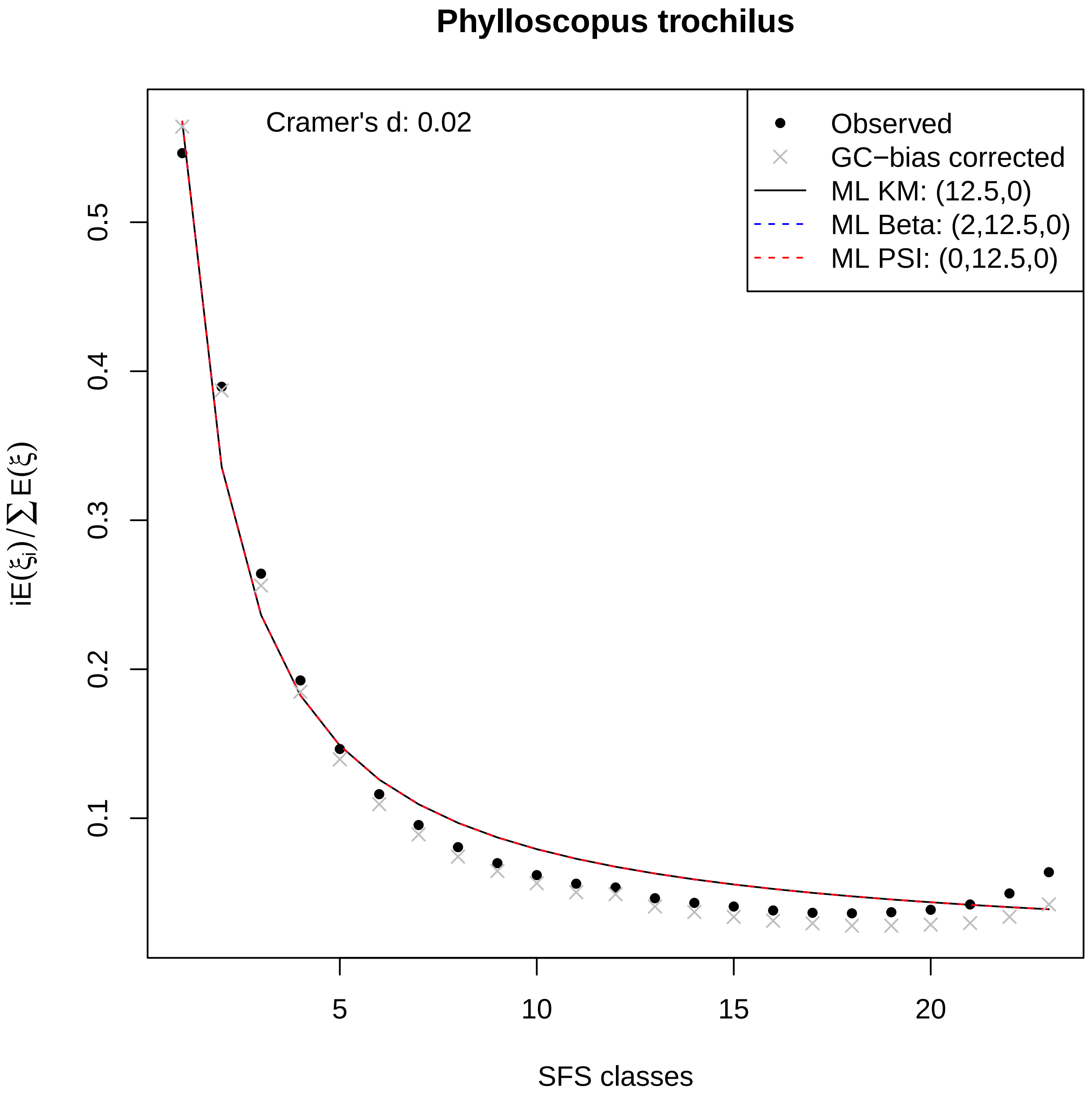

Phylloscopus trochilus
Cramer's d: 0.02
Observed
GC−bias corrected
ML KM: (12.5,0)
ML Beta: (2,12.5,0)
ML PSI: (0,12.5,0)
0.1
0.2
0.3
0.4
0.5
5
10
15
20
SFS classes
$iE(\xi_i)/\sum E(\xi)$

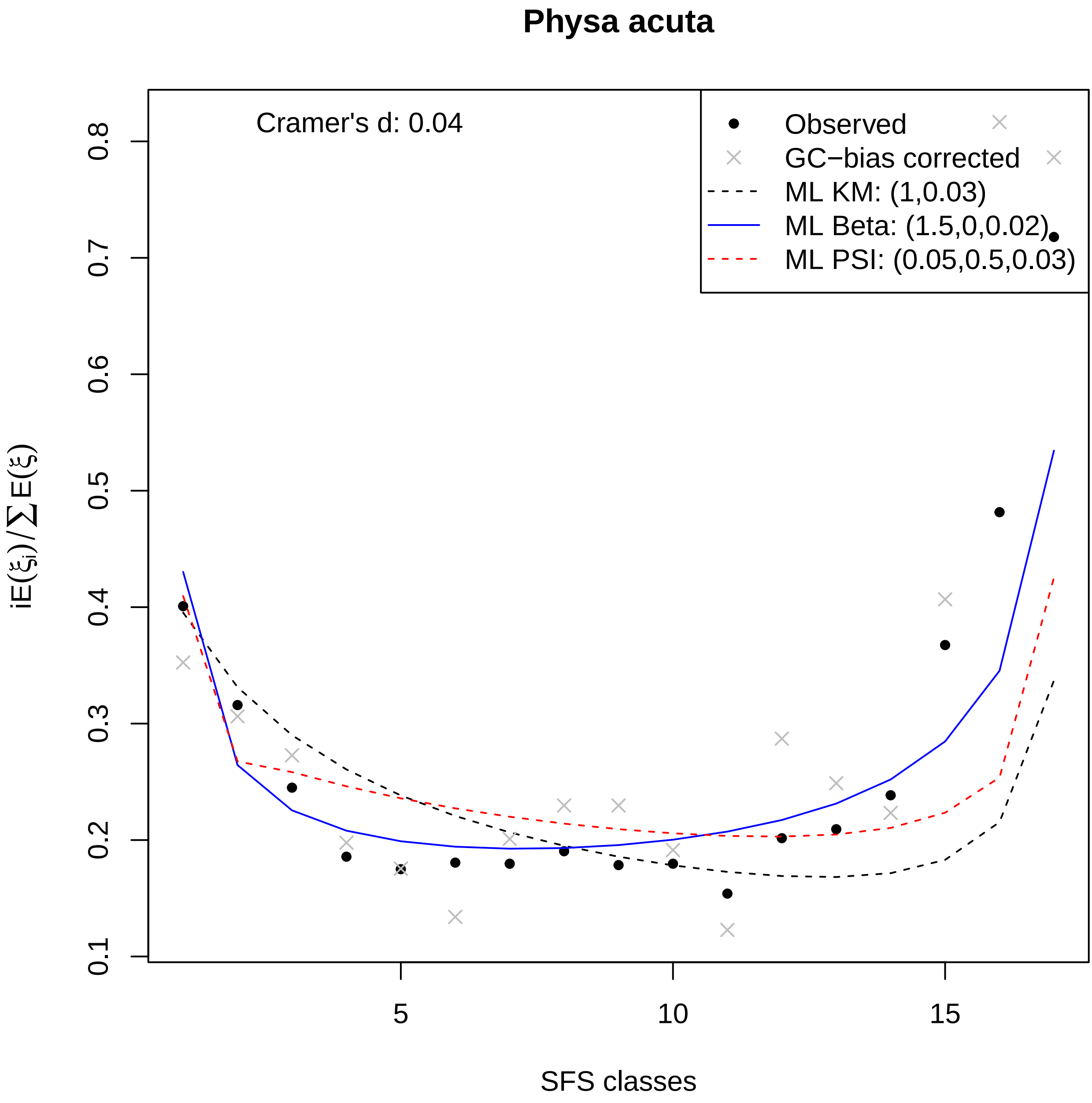
Physa acuta
Cramer's d: 0.04
Observed
GC−bias corrected
ML KM: (1,0.03)
ML Beta: (1.5,0,0.02)
ML PSI: (0.05,0.5,0.03)
0.1
0.2
0.3
0.4
0.5
0.6
0.7
0.8
5
10
15
SFS classes
iE(ξi)/∑E(ξ)

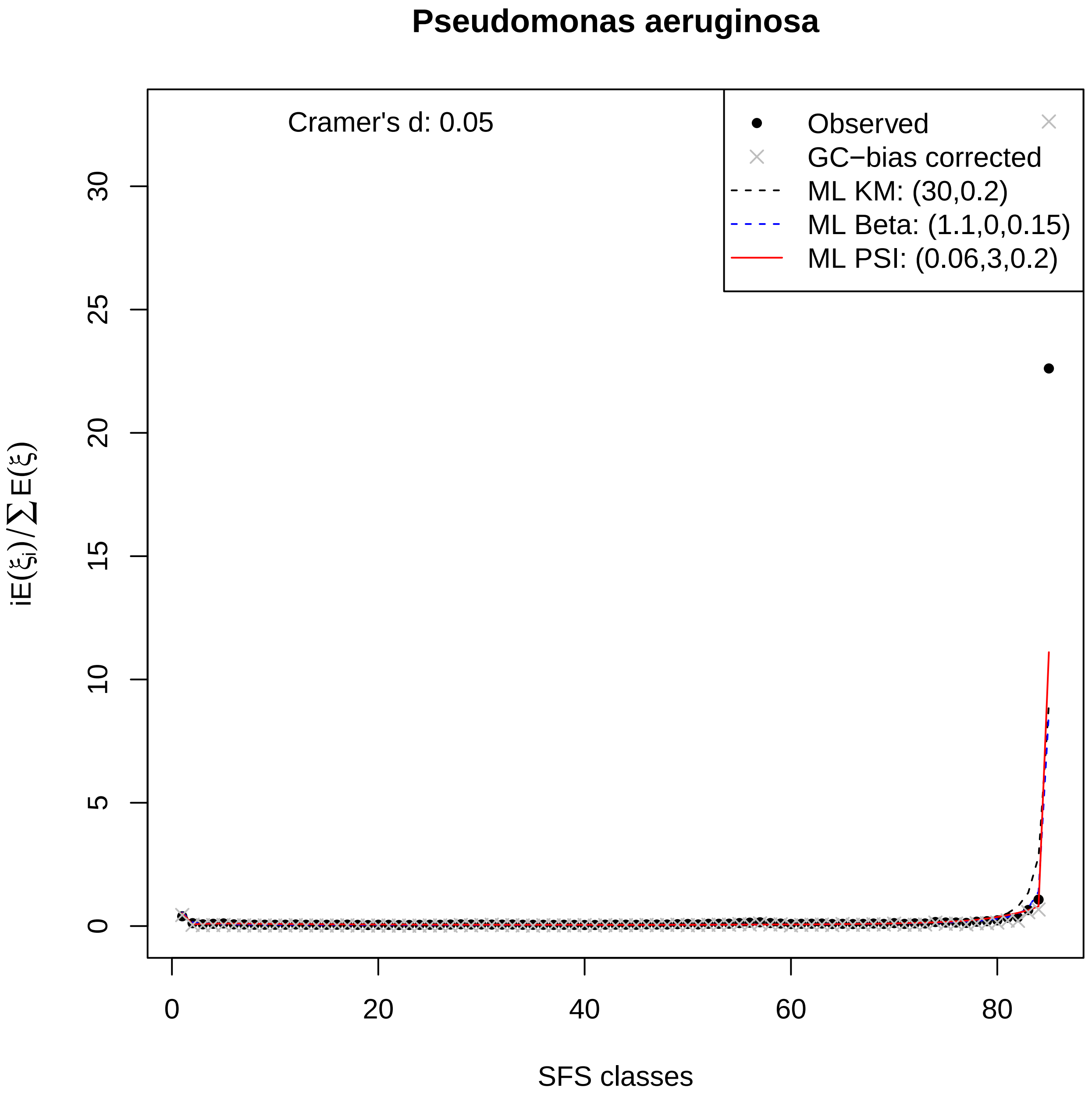

Pseudomonas aeruginosa
Cramer's d: 0.05
Observed
GC−bias corrected
ML KM: (30,0.2)
ML Beta: (1.1,0,0.15)
ML PSI: (0.06,3,0.2)
$iE(\xi_i)/\sum E(\xi)$
0
5
10
15
20
25
30
0
20
40
60
80
SFS classes

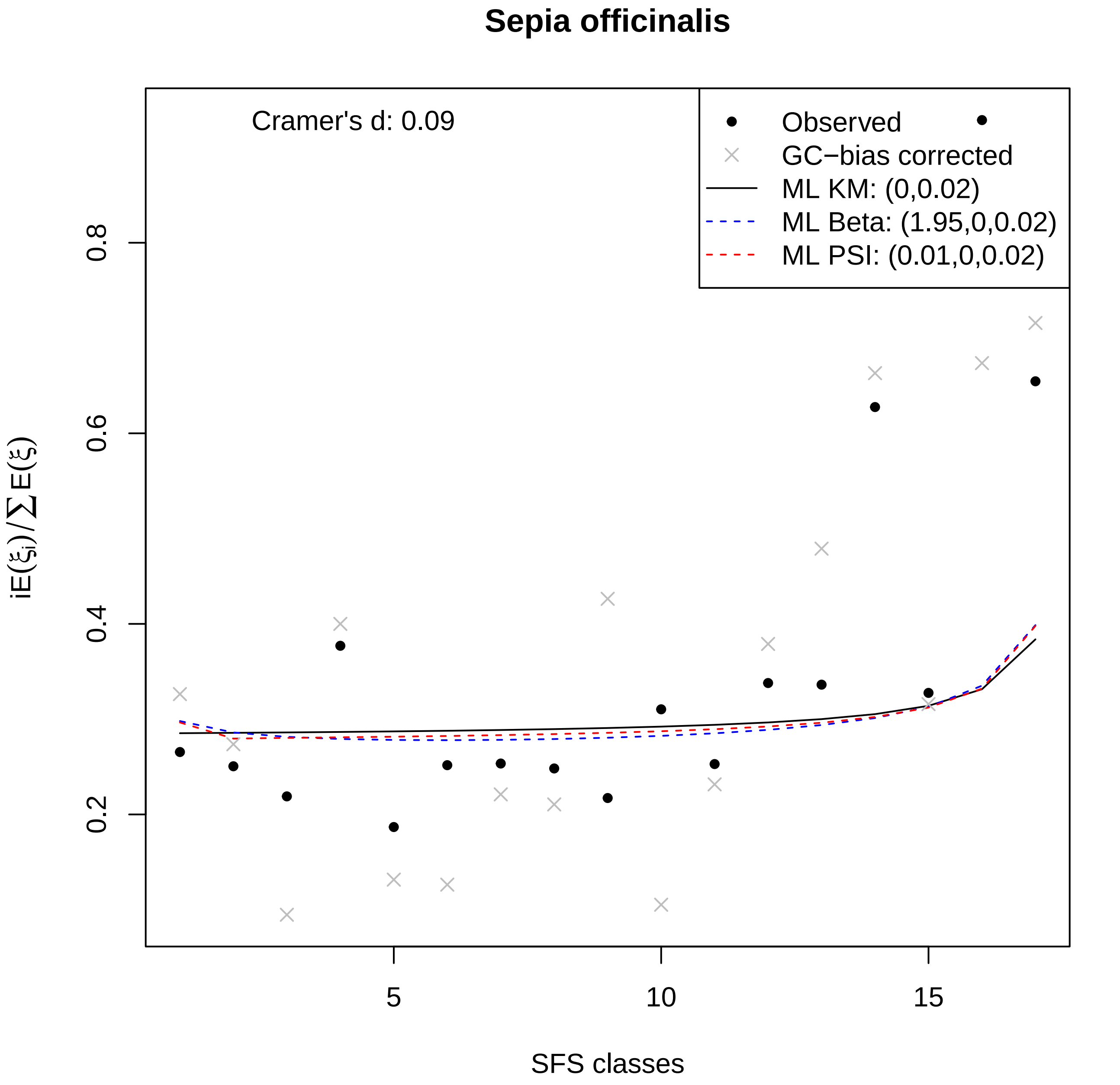
Sepia officinalis
Cramer's d: 0.09
Observed
GC−bias corrected
ML KM: (0,0.02)
ML Beta: (1.95,0,0.02)
ML PSI: (0.01,0,0.02)
$iE(\xi_i)/\sum E(\xi)$
0.2
0.4
0.6
0.8
5
10
15
SFS classes

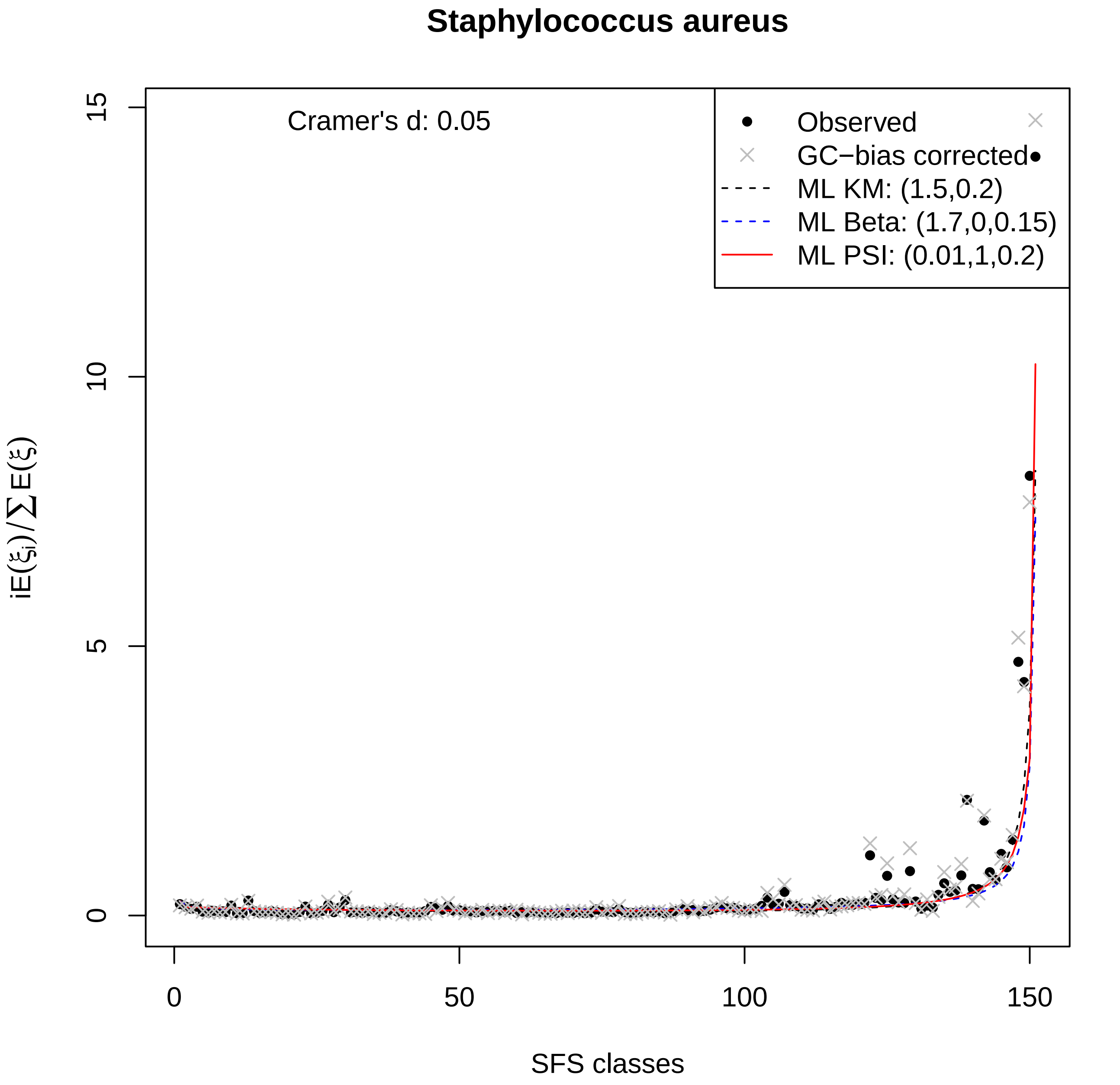
Staphylococcus aureus
Cramer's d: 0.05
Observed
GC−bias corrected
ML KM: (1.5,0.2)
ML Beta: (1.7,0,0.15)
ML PSI: (0.01,1,0.2)
iE(ξi)/∑E(ξ)
SFS classes
0
5
10
15
0
50
100
150

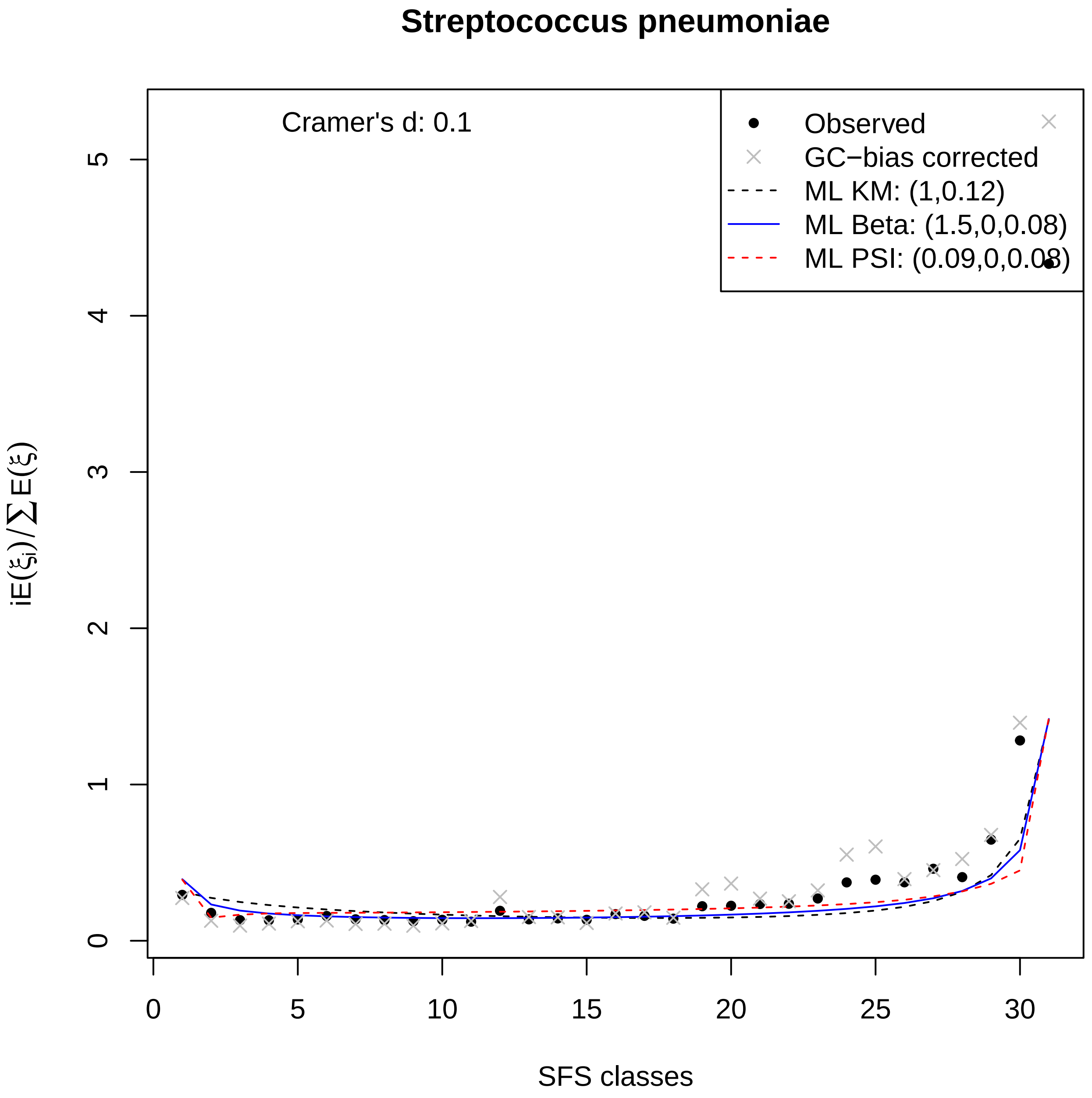

Streptococcus pneumoniae
Cramer's d: 0.1
Observed
GC−bias corrected
ML KM: (1,0.12)
ML Beta: (1.5,0,0.08)
ML PSI: (0.09,0,0.08)
$iE(\xi_i)/\sum E(\xi)$
SFS classes
0
1
2
3
4
5
0
5
10
15
20
25
30

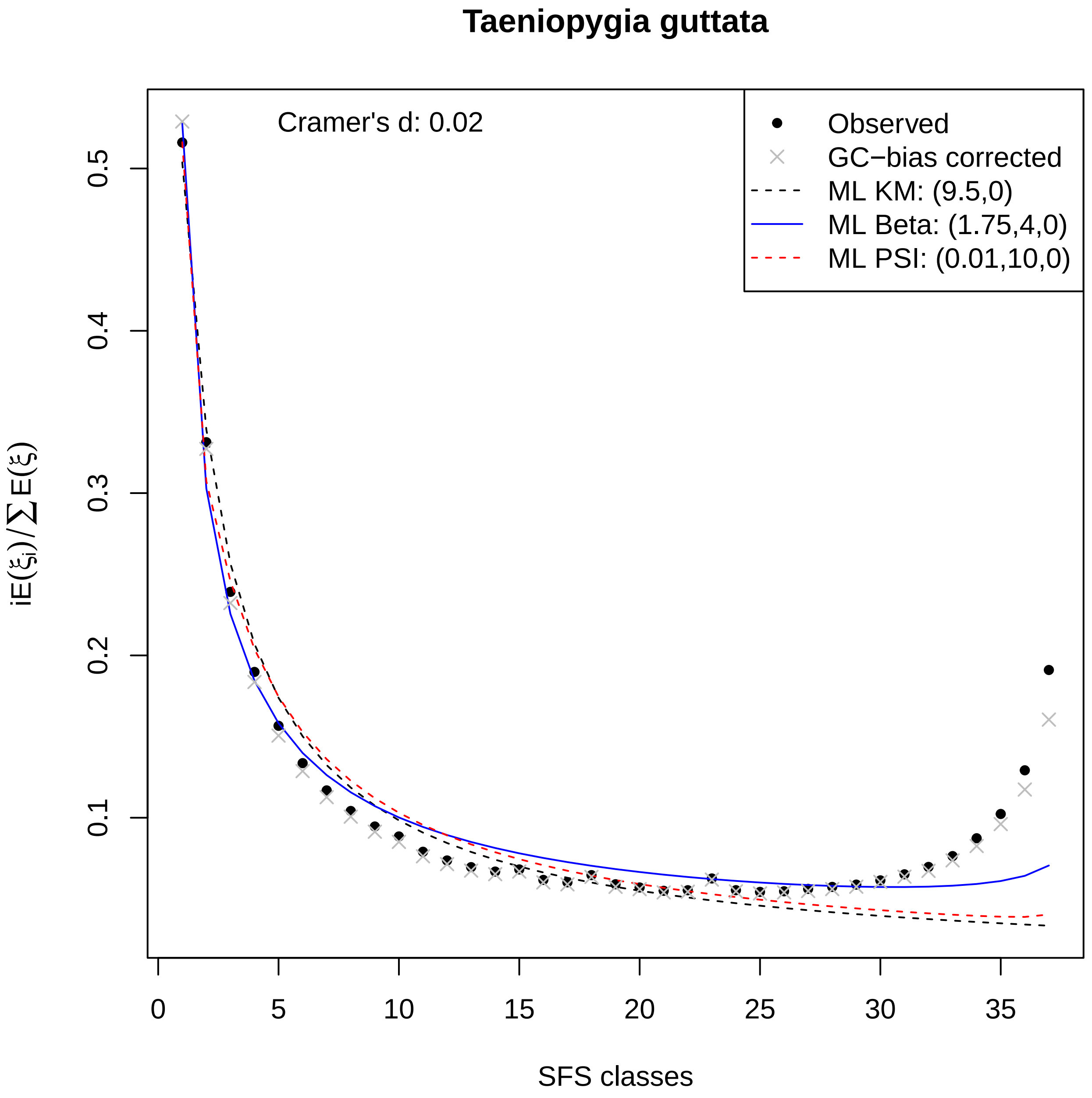

Taeniopygia guttata
Cramer's d: 0.02
Observed
GC−bias corrected
ML KM: (9.5,0)
ML Beta: (1.75,4,0)
ML PSI: (0.01,10,0)
$iE(\xi_i)/\sum E(\xi)$
SFS classes
0.1
0.2
0.3
0.4
0.5
0
5
10
15
20
25
30
35

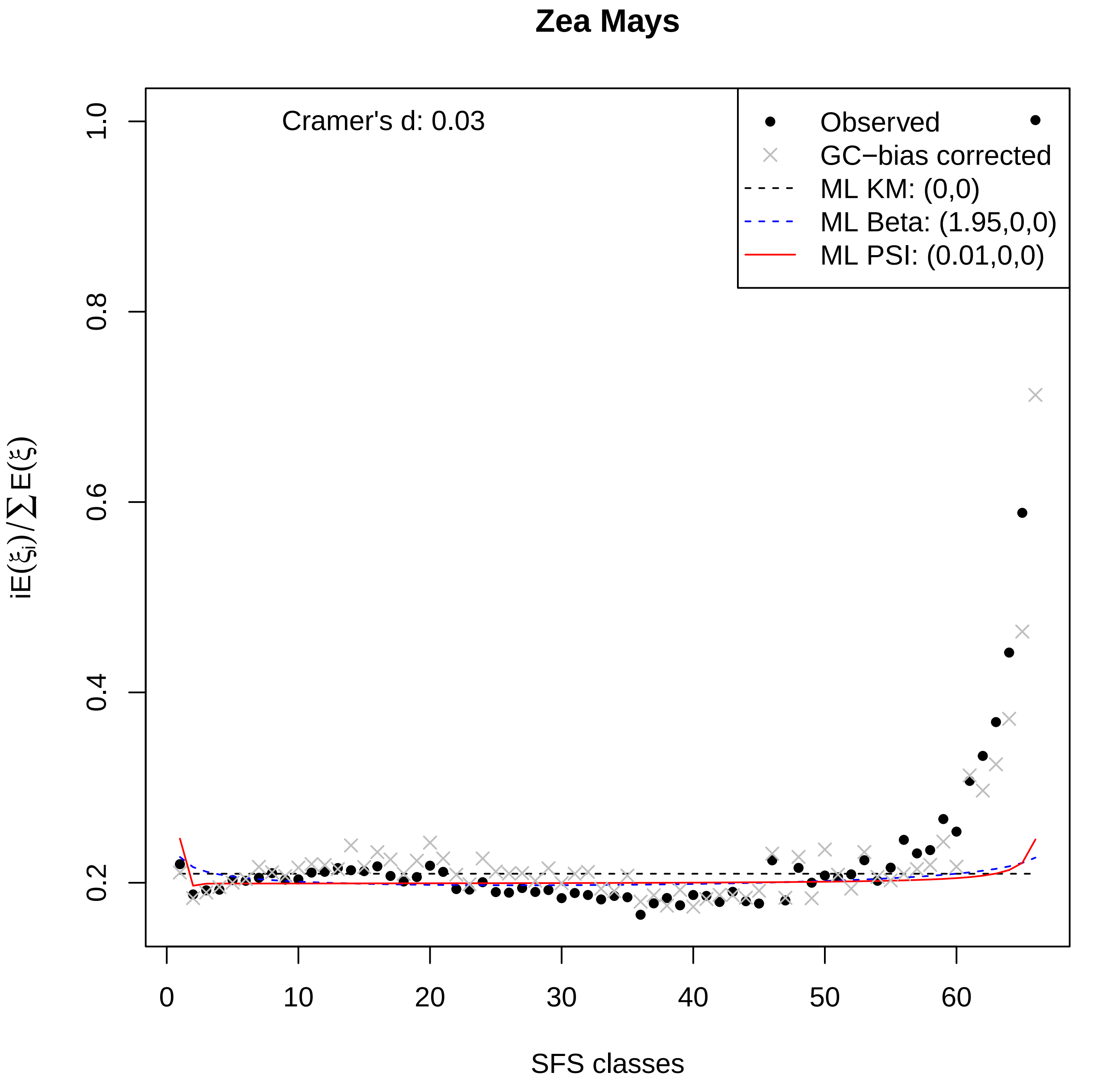
Zea Mays
Cramer's d: 0.03
Observed
GC−bias corrected
ML KM: (0,0)
ML Beta: (1.95,0,0)
ML PSI: (0.01,0,0)
0.2
0.4
0.6
0.8
1.0
0
10
20
30
40
50
60
SFS classes
$iE(\xi_i)/\sum E(\xi)$
